# Intravital dynamic and correlative imaging reveals diffusion-dominated canalicular and flow-augmented ductular bile flux

**DOI:** 10.1101/778803

**Authors:** Nachiket Vartak, Georgia Guenther, Florian Joly, Amruta Damle-Vartak, Gudrun Wibbelt, Jörns Fickel, Simone Jörs, Brigitte Begher-Tibbe, Adrian Friebel, Kasimir Wansing, Ahmed Ghallab, Marie Rosselin, Noemie Boissier, Irene Vignon-Clementel, Christian Hedberg, Fabian Geisler, Heribert Hofer, Peter Jansen, Stefan Hoehme, Dirk Drasdo, Jan G. Hengstler

## Abstract

Small-molecule flux in tissue-microdomains is essential for organ function, but knowledge of this process is scant due to the lack of suitable methods. We developed two independent techniques that allow the quantification of advection (flow) and diffusion in individual bile canaliculi and in interlobular bile ducts of intact livers in living mice, namely Fluorescence Loss After Photoactivation (FLAP) and Intravital Arbitrary Region Image Correlation Spectroscopy (IVARICS). The results challenge the prevailing ‘mechano-osmotic’ theory of canalicular bile flow. After active transport across hepatocyte membranes bile acids are transported in the canaliculi primarily by diffusion. Only in the interlobular ducts, diffusion is augmented by regulatable advection. Photoactivation of fluorescein bis-(5-carboxymethoxy-2-nitrobenzyl)-ether (CMNB-caged fluorescein) in entire lobules demonstrated the establishment of diffusive gradients in the bile canalicular network and the sink function of interlobular ducts. In contrast to the bile canalicular network, vectorial transport was detected and quantified in the mesh of interlobular bile ducts. In conclusion, the liver consists of a diffusion dominated canalicular domain, where hepatocytes secrete small molecules and generate a concentration gradient and a flow-augmented ductular domain, where regulated water influx creates unidirectional advection that augments the diffusive flux.

**One Sentence Summary/Keywords:** Bile flux proceeds by diffusion in canaliculi, augmented by advection in ducts.

## Main Text

The flux of small molecules through tissue compartments and vessel networks is a fundamental process supporting organ function. Analysis of flux in microscopic vessels, cells and tissue compartments in living organisms remains intractable due to their inaccessibility to conventional rheological and ultrasonic methods. In this work, we used the liver as an exemplary organ and quantified transport in its microscopic biliary conduits. Liver tissue architecture consists of lobules — functional units comprised of blood capillaries called sinusoids, hepatocytes, and a canalicular network formed by hepatocyte apical membranes (Fig 1A). The canalicular networks are linked to interlobular bile ducts (IBDs), which progressively converge into larger ducts and finally the extrahepatic bile duct (EHBD)(1).

**Fig. 1.**
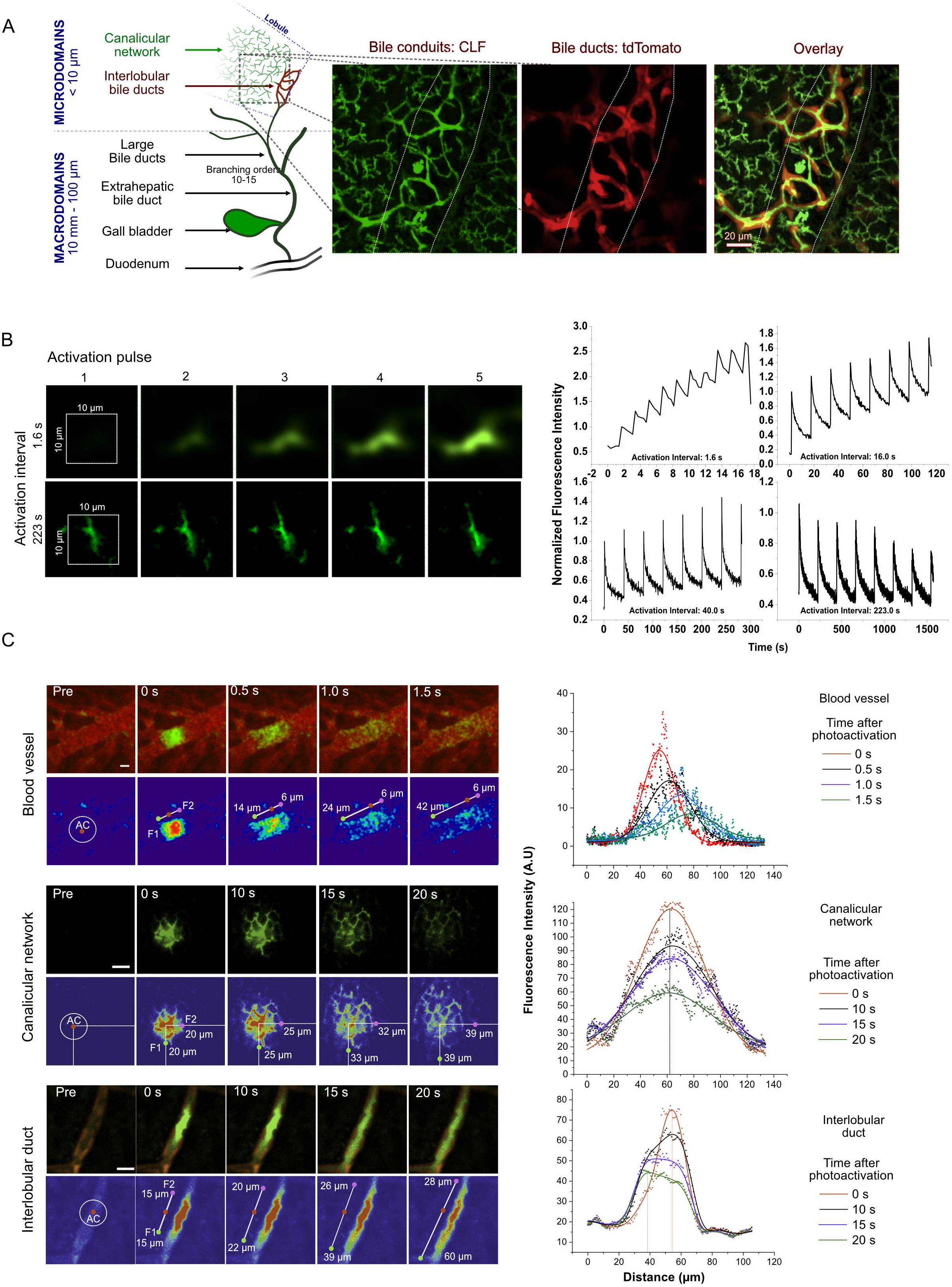
Definition of diffusion-dominated and advection-augmented compartments of the biliary system and identification of the upper-bound of flux kinetics. **A**. The diagram depicts the microdomains analyzed in the present study connected to macroscopic structures of the downstream biliary tract. Confocal maximum intensity projection (20 μm) of HNF1beta/CreER-tdTomato labeled bile ducts (red) and CLF-enriched canalicular network and duct lumen (green) 10 min after tail-vein injection of CLF in live mice. White dotted line indicates the portal vein. **B.** Representative sequence of repeating CMNB-Fluo (CMNB-Fluo) photoactivation, with pulses spaced 1.6 s or 223 s apart, showing accumulating fluorescence due to insufficient depletion of a 10 μm×10 μm activation region by canalicular bile flux. The graph shows quantification of intensities, indicating that CMNB-Fluo requires >40 s to be cleared from a 10 μm×10 μm region by canalicular flux. **C.** Photoactivation of CMNB-Fluo-Dextran (10Kd) in a central vein blood vessel and CMNB-Fluo in the canalicular network and duct showing the activated region (white circle) with its centroid (red, AC), and the distance travelled by the fluorescence fronts (F1, F2) from the centroid due to bile flux. Ducts show vectorial flux (F1>F2) in contrast to symmetric dispersion in the canalicular network (F1~F2). The upper panel shows raw images (green: CMNB-Fluo, red: tdTomato marking ducts) while the lower panel represents the same with a polychromatic color table and annotations. Scale bars: 20 μm. The graphs depict the temporal evolution of spatial intensity profiles after photoactivation, with the intensity centre-of-mass drifting in IBDs but a remaining static in the canalicular network. Photoactivated region: White dotted circle. Scale bar: 20 μm.

The prevailing concept ascribes the movement of bile (solvent) and its constituent small molecule solutes (bile acids, xenobiotics etc.) to solvent fluid advection due to osmotic water influx and canalicular membrane contractility (2). Hepatocyte-mediated active transport of bile acids and other organic solutes into the canalicular lumen is expected to generate an osmotic gradient that drives the movement of water and causes flow of canalicular bile. Bile would then be pushed through the canalicular network with an increasing velocity towards IBDs, supported by peristalsis-like contractions of canalicular membranes (3). In the present study, we established <u>I</u>ntra<u>v</u>ital <u>A</u>rbitrary <u>R</u>egion <u>I</u>mage <u>C</u>orrelation <u>S</u>pectroscopy (*IVA*RICS), <u>f</u>luorescence <u>l</u>oss <u>a</u>fter <u>p</u>hotoactivation (FLAP) and time-lapse microscopy techniques to quantify flux in microdomains in live organisms. Applying these techniques to bile canaliculi and interlobular bile ducts revealed a diffusion-dominated canalicular and flow-augmented ductular domain through which bile flux occurs. We present a mechanistic model of bile flux that corroborates existing knowledge of bile clearance, but is not compatible with the prevailing mechano-osmotic concept of bile flow.

## Materials and Methods

Materials and Methods for animal handling and preparation for microscopy, experimental details of intravital imaging procedures, acquisition parameters and analysis routines for photoactivation and IVARICS, in silico modeling, and synthesis of fluorescent bile acid analogs are provided in the Supporting Information.

## Results

Intravital analysis of bile flux was enabled by HNF1beta/CreER-reporter mice which express tdTomato specifically in cholangiocytes (4), and through the use of fluorescent analogs of bile salts. Fluorescent analogs such as cholyl-lysyl fluorescein (CLF) are transported by hepatocytes from sinusoids to canaliculi, allowing direct visualization of the canalicular network. Confocal or 2-photon imaging of live mouse livers show the canalicular networks and connecting interlobular bile ducts, depicting the basic lobular architecture in which biliary flux occurs (Fig. 1A).

### Determination of local flux mechanisms and the bounds of flux kinetics in the liver canalicular network

Bile flux may occur either by advection or by inherent diffusion of the bile salts through the canalicular space, or a combination of both (convection). To investigate the local flux mechanisms, we utilized fluorescence loss after photo-activation (FLAP) of a photoactivatable analog of fluorescein (CMNB-Fluorescein) in intravital microscopy of mouse livers. Following tail vein injection, CMNB-Fluorescein (CMNB-Fluo) was photo-activated at time points when it was almost entirely localized in the canalicular network. Repeated photoactivation of CMNB-Fluo was performed in a pulsed manner on predefined regions of known spatial dimensions (Fig. 1B). By varying the time-interval between the photoactivation pulses, a race-condition is set up in the defined region between photoactivation pulses which increase fluorescence, and the underlying flux mechanism which depletes fluorescence. Accumulation of fluorescence with subsequent pulses indicates insufficiency of the underlying flux to remove the fluorescent material in the time interval between pulses. An upper-bound was established by decreasing the activation pulse rate until the underlying flux rate could finally remove the fluorescent material in the time-interval between activation pulses. Assuming pure advection in the canaliculi, the upper limit of the flux rate was determined to be **~0.05 μm/s** as the ratio of the maximum length of the activated region and the minimum interval between pulses in which it is depleted. Correspondingly, assuming pure diffusion, a lower bound of **0.5 μm^2^/s** is obtained for the diffusion coefficient from the ratio of the area of the activated region and the time-interval required to deplete fluorescence in the region. These limits of the flux rates of advection or diffusion suggest that the inherent diffusion of small molecule solutes such as bile acids dominates their transport, rather than advection of the solvent.

### FLAP differentiates the diffusion-dominated and advection-augmented compartments of the biliary system

To determine the mechanism of bile flux, we examined the nature of the fluorescence loss in the photoactivated (FLAP) region after a single photoactivation pulse in a blood vessel, canalicular network and interlobular bile duct. The blood vessel represents a flow-dominated system, and thus the photoactivated mass disperses by molecular diffusion in the solvent, while moving along the with the vectorial blood flow (Fig. 1C, Mov. S1). We next measured the loss of fluorescence in a circular photoactivated region in the bile canalicular network, since the diameter of individual canaliculi approaches the optical resolution limit and is too small to precisely define a region within the lumen. The dispersion of photoactivated material is symmetric with a corresponding symmetric spread outside this region, irrespective of the lobular location of the activated zone (Fig 1C, Mov. S2). The location of the fluorescence maximum within the activated zone remains invariant over time. Alternative activation geometries (stripe, 3D) led to identical conclusions (Fig. S1A, Mov. S3, Mov. S10). This indicates that the local biliary flux in canaliculi is **not vectorial** but rather the dispersion of bile salts is symmetric and omnidirectional in the canalicular network. This implies molecular diffusion rather than vectorial flow as the primary flux mechanism in the canalicular network, consistent with the extremely low upper bound of pure advection in the canalicular network. The same intravital photoactivation technique was applied to intralobular bile ducts (Fig. 1C) that were specifically visualized through the cholangiocyte-specific tdTomato-fluorescence-labeling in the HNF1beta/CreER-tdTomato mice. Contrary to observations in canaliculi but similar to blood, dispersion of fluorescence was asymmetrical with vectorial bias along one direction in the IBD. This indicates the presence of advection, in addition to diffusion, that conducts bile flux in the interlobular ducts. However, advection in the interlobular bile duct is substantially slower compared to the blood vessel.

To demonstrate diffusion-dominance in the canalicular network, we performed FLAP experiments at the Canal of Hering (CoH) - connections of the canalicular network to IBDs. These junctional regions would represent the transition zone from the diffusion-dominated canalicular lumen to the advection-augmented ductular lumen. Photoactivated CMNB-Fluo in the canalicular network adjacent to the CoH (i.e. PV zone) transited to the IBD and diffused symmetrically into the network as expected (Fig. 2A). However, photoactivated CMNB-Fluo in the CoH also transited retrograde into the canalicular network (Fig. 2B). This retrograde flux of fluorescence from the CoH into the canalicular network confirmed that in the canalicular network, the inherent diffusion of small molecules overrules the meagre (if any) advection towards the bile duct.

**Fig. 2.**
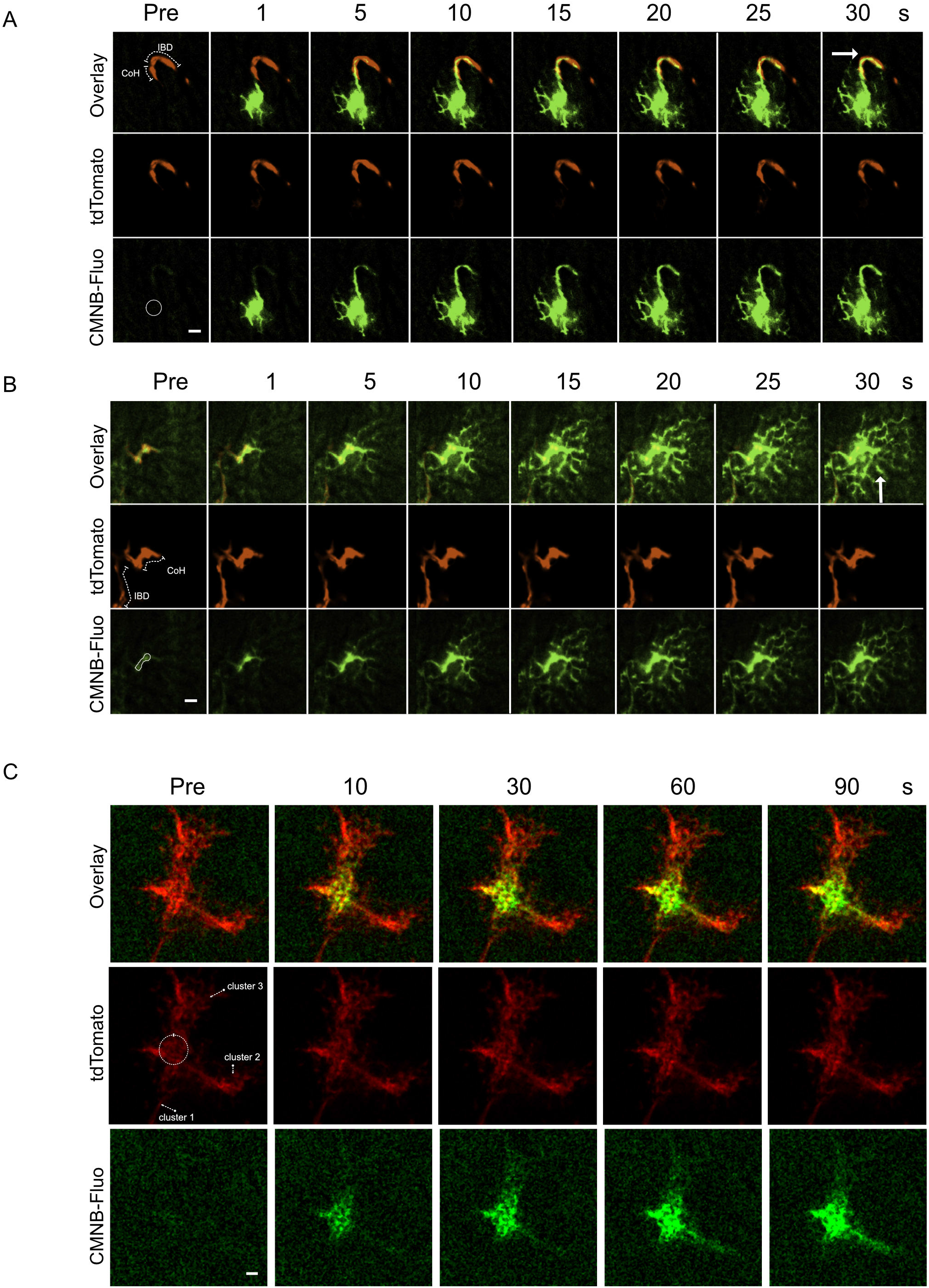
Demonstration of bile acid flux within and between diffusion-dominated canaliculi network and advection-augmented ductular mesh. **A.** Representative images of photoactivation in the PV zone adjacent to a CoH junction. **B.** Photoactivation in the CoH junction (right) showing ductular transit and retrograde flux into the canalicular network, respectively. Duct and CoH are annotated as dotted lines. **C.** Photoactivation in a ductular mesh surrounding a portal vein. Clusters of branches of the ductular network are indicated. Photoactivated region: White dotted circle. Scale bar: 20 μm.

We then confirmed that unidirectional flow in individual ductules (Fig. 1C) leads to a similar net vectorial flow in the larger ductular mesh surrounding portal veins. We photoactivated a region of the mesh encompassing several ductules, themselves further connected to other branch clusters. CMNB-Fluo preferentially migrated to only some of all available connected clusters of branches. This demonstrates vectorial flow in the larger interlobular biliary network (Fig. 2C).

The insignificant canalicular advection under basal conditions led us to test if conventional interventions that are known to increase extrahepatic bile flow could induce canalicular advection. Secretin is known to stimulate bile flow from the bile duct epithelium (5) and is expected to increase extrahepatic bile flow without increasing canalicular flux. In contrast, taurocholate (TCA) is excreted into canaliculi and expected to cause hyperosmotic water influx (6) to increase canalicular bile flow. Secretin as well as TCA infusion both induced an increase in extrahepatic bile flow from basal levels of ~1 μl/min/g liver weight, upto ~3 μl/min/g liver weight (Fig 3A), recapitulating previous reports (7,8).

**Fig. 3.**
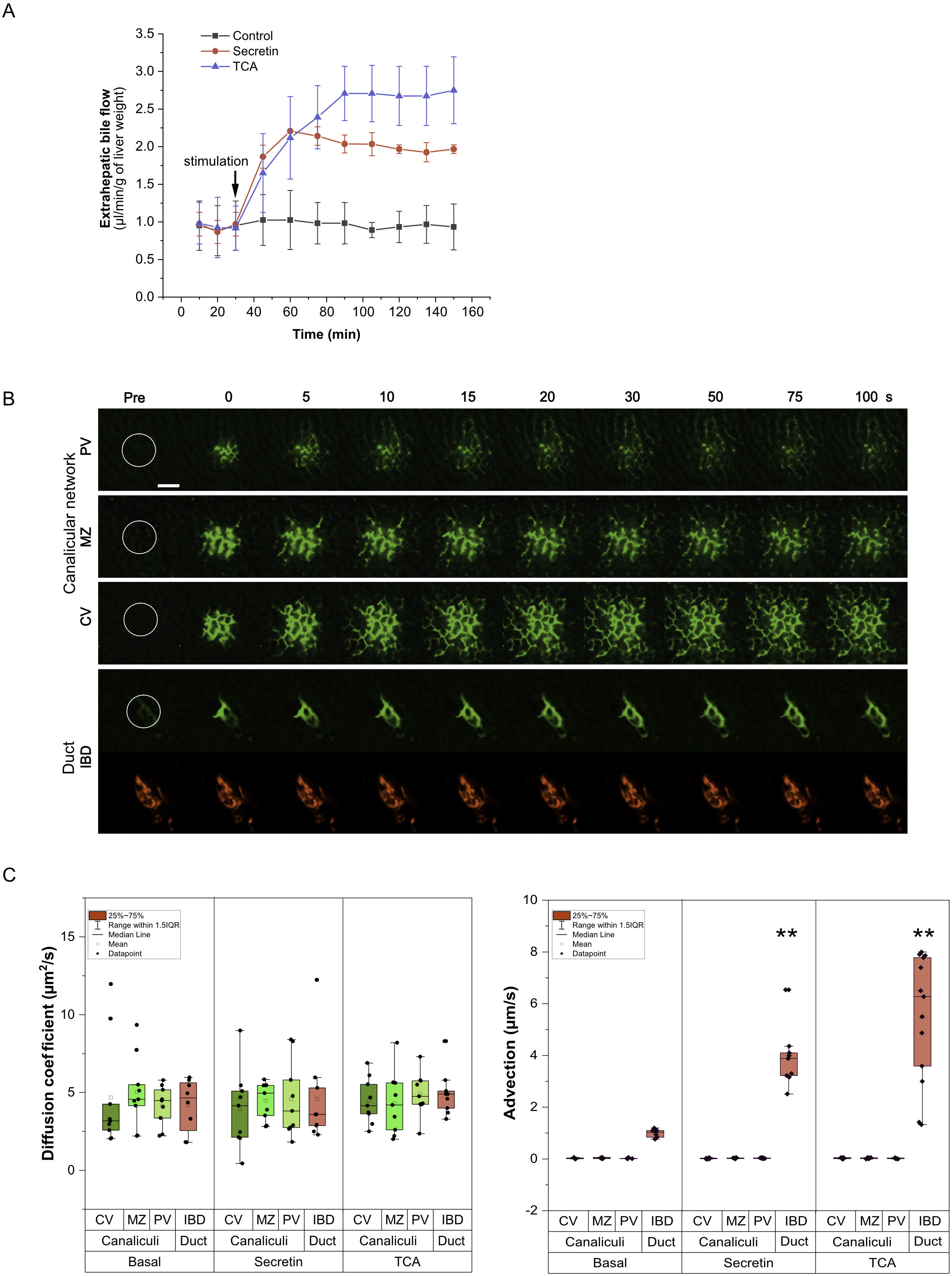
Quantification of canalicular flux in basal and stimulated conditions. **A**. Stimulation of extrahepatic bile flow after administration of a secretin bolus or TCA infusion. Stimulation was provided 30 min after extrahepatic bile collection started (arrow). Data indicated mean ± S.D of 3 mice per condition. **B.** Representative time-lapse images following photo-activation in a defined region (white circle) of CMNB-Fluo (green) in canalicular networks adjacent to the central vein (CV), portal vein (PV) or in the mid-zone (MZ), as well as in interlobular bile ducts marked by tdTomato (red). Scale bars: 50 μm. **C.** Quantification of diffusion coefficients from the half-life of intensity decay and advection velocity from shift of intensity centre-of-mass for various lobular zones (CV: pericentral; MZ: midzonal; PV: periportal) under basal conditions or stimulation with secretin and TCA. Values indicate measurements in n>8 mice for each condition. Dots in the box plots represent the data of individual mice. **p<0.01 compared to controls.

Photoactivation in various lobular zones and interlobular bile ducts was performed (Fig. 3B, Mov. S4) and the half-life of clearance was determined (Fig. S1B). Relative contributions of diffusion and advection were determined by quantification of the dispersion and displacement of Gaussian photo-activation profiles for the diffusion coefficient and velocity, respectively (9) (Supporting information: Photoactivation).

Under basal conditions, diffusion was similar in the interlobular ducts and all lobular zones of the canalicular network, ranging between median diffusion coefficients of **2-5 μm^2^/s** (Fig. 3C, Table S1). However, the advective velocity in canalicular networks was **less than 0.02 μm/s**. In contrast, bile ducts showed an advective velocity ranging from **1-1.2 μm/s**. These findings reinforce the deduction that biliary flux is diffusion-dominated in the canalicular network but show that it is augmented by advection in the bile ducts. Following secretin administration or TCA infusion, advection in the canalicular network did not significantly increase and the diffusion coefficient remained similar to that under basal conditions. In the interlobular bile ducts, the diffusion coefficient also remained unchanged from that of basal conditions, but the median advection velocity increased to up to **3.8 μm/s after secretin administration** and up to **6.2 μm/s during TCA-infusion** (Fig. 3C, Table S1). These findings are consistent with the known property of cholangiocytes to secrete water upon secretin stimulation (10) or when excess bile salts are detected in the bloodstream (11). Yet, even under these conditions, we find that the canalicular bile flux remains diffusion-dominated.

### Quantification of flux parameters for diffusion and advection - <u>Intravital Arbitrary Region Image Correlation Spectroscopy (IVARICS)</u>

The quantification methods used for FLAP (9) assume a homogeneous fluid medium without diffusion barriers. Non-ideal conditions in a living mouse such as inherent inhomogeneity of the canalicular network, its finite size, boundary at the central vein and the sink at the CoH/IBD may affect the photo-activation profiles. To remove the effect of these non-ideal morphological constraints on determination of flux parameters, we established an orthogonal approach to determine diffusion and advection rates in the biliary compartments – namely, <u>Intravital Arbitrary Region Image Correlation Spectroscopy</u> (IVARICS). Image correlation spectroscopies (ICS) (12,13) are based on rapid acquisition of fluorescence fluctuations caused by molecular movements (flux) in a defined region and the generation of corresponding correlation maps. The method is intrinsically local and yields apparent diffusion coefficients and advection velocities for the acquisition region. Image correlation techniques have distinct limitations from those of FLAP. They require a priori knowledge of the range in which flux parameters are expected, on which basis image acquisition and fitting parameters have to be set. In the case of in vivo imaging, light scattering by tissue and movements of the animal during image acquisition must be compensated for during analysis (see Supporting Information: IVARICS). Nonetheless, due to its fundamentally different imaging acquisition, analysis and limitations, IVARICS represents a complementary and orthogonal approach to FLAP analysis.

We first determined the feasibility of applying image correlation methods in vivo, by acquiring conventional spatial and temporal ICS sequences in the liver sinusoids, hepatocytes and canaliculi of mice infused with CLF. Since sinusoidal blood flow comprises both diffusion and advection, while the hepatocyte cytoplasm comprises only diffusion, these compartments served as *in situ* controls. IVARICS sequences could be used to extract the flux mechanism – sinusoidal blood flow was measured to be ~60 μm/s and diffusion coefficients were ~2 μm^2^/s in all compartments (Fig. 4A). No advection could be detected in the hepatocyte cytoplasm as well as the canalicular lumen. The effect of animal movements due to heartbeat, respiration and intestinal peristalsis (Fig. S2, Mov. S5) could be mitigated during post-processing using pruning of slow autocorrelations and the use of the recently reported arbitrary region raster ICS (RICS) algorithm (13) (Fig. 4B, see Supporting Information). Spatial and temporal image autocorrelations (RICS/TICS) were independently fitted to appropriate functions (see Supporting Information: IVARICS) to determine diffusion coefficients and velocities (Fig. 3C, D, S2, Table S2, S3).

**Fig. 4.**
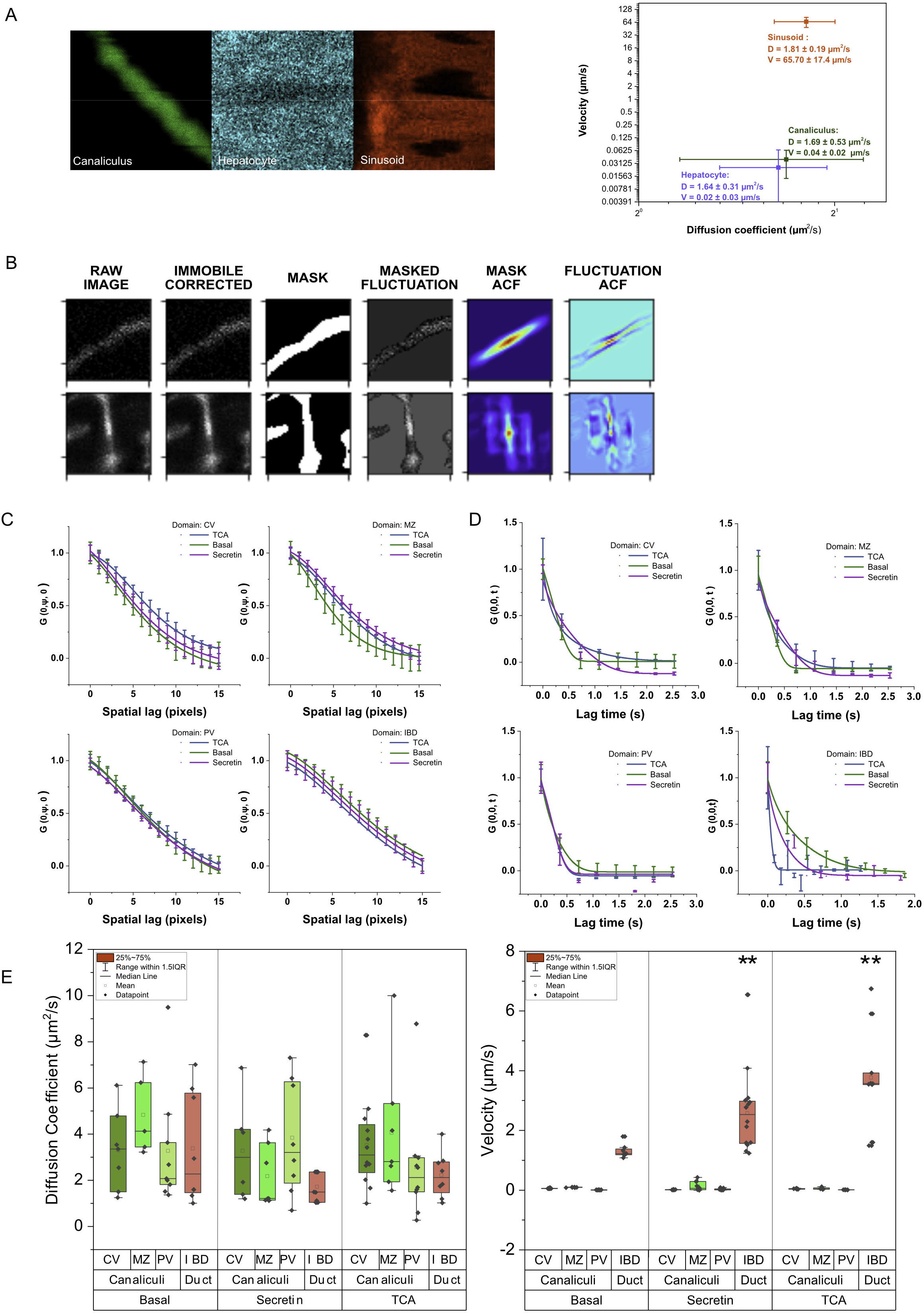
Establishment of IVARICS. **A.** Representative images acquired for ICS analysis in canaliculi, hepatocyte cytoplasm and sinusoids. The graph shows quantification of diffusion coefficient and velocity in each compartment. **B.** Workflow of IVARICS analysis. Representative fluctuation images of fluorescein loaded canaliculi and ducts are acquired. A binary mask representing the structures of interest is created. Autocorrelation (ACF) maps are generated for both the intensity images and masks. A ratio of the intensity ACF and mask ACF yields a fluctuation ACF devoid of movement and shape effects. **C.** Average normalized spatial (y)-autocorrelation for various liver domains with fits for 1-population 3D-diffusion. **D.** Average normalized temporal autocorrelation for various liver domains with fits for 1-population 3D-diffusion and advection. **E.** Diffusion coefficients and velocities derived from spatial (RICS) and temporal (TICS) analysis for various canalicular zones (green) and interlobular bile ducts (red) under basal conditions or stimulation with secretin and TCA. Values indicate measurements of n>4 mice for each condition. The dots in the box plots represent the data of individual mice. **p<0.01 compared to controls.

Diffusion coefficients and velocities derived from IVARICS analysis of CLF or fluorescein-infused mice confirmed the findings of the FLAP experiments (Fig. 3C, D, E). Median ductular advection velocity was measured to be ~1 μm/s under basal conditions, which increased up to a value of 2.4 μm/s by the action of secretin on the duct, and up to 3.7 μm/s through TCA infusion (Fig. 4E). Flow in the canalicular network remained negligible and was neither affected by secretin administration, nor by TCA infusion. Diffusion coefficients in the canalicular network and interlobular ducts ranged between 2.4-6 μm^2^/s. Thus, the general conclusion of the FLAP experiments was validated by *IVARICS*.

### Simulation of Bile Flux in the liver

To validate if a system with diffusion-dominated canalicular flux, augmented by advection in the duct, is capable of accurately describing the behavior of bile acids in liver canalicular networks and ducts, we developed a deterministic Navier-Stokes *in silico model* in realistic canalicular geometries of the liver tissue. Confocal scans of immunofluorescence stained liver tissue showing canalicular network and interlobular bile ducts were generated as described previously (14) and digitized into triangulated meshes that served as the simulation space (Fig. 5A). Concentrations of fluorescein or CLF generated within various liver compartments during our experiments were estimated by establishing instrument calibration curves in murine bile in vitro (Fig. S3A). These concentrations, and the diffusion coefficients as well as velocities for the canalicular network and IBDs determined thus far were used as input parameters for the simulations.

**Fig. 5.**
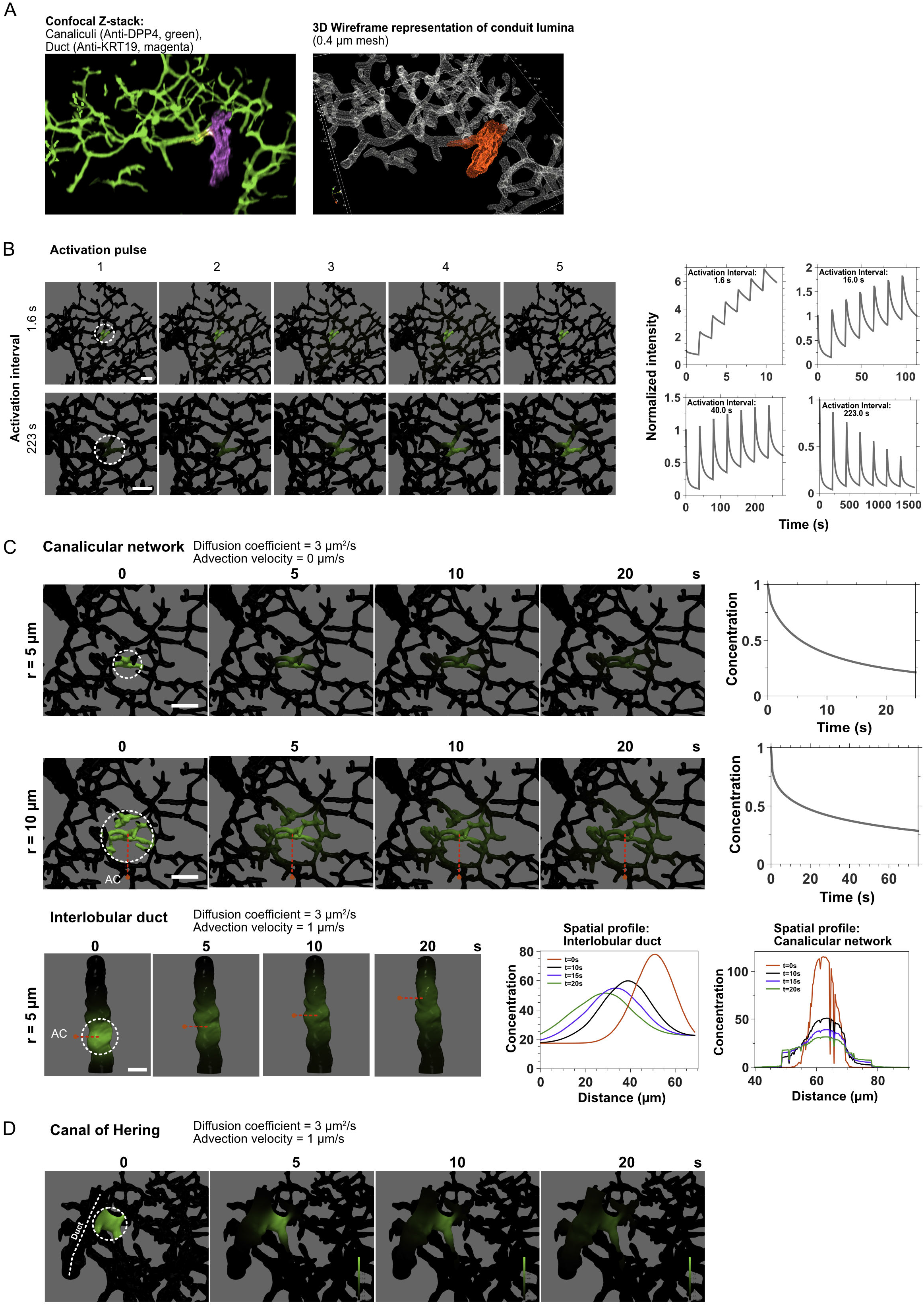
Simulation of bile acid flux mechanisms. **A.** Representative 3D immunostained confocal stack (left) identifying the canalicular network and interlobular bile duct and its digitization to a polygonal mesh (right). **B.** Simulation of repeated photoactivation with various time intervals in a 10 μm radius spherical region encompassing a canaliculus. Graphs show quantification of mean intensity in the activated spherical region with various activation intervals (compare: Fig. 1B). **C.** Simulation of single photoactivation in the canalicular network of varying radii (top, middle) and interlobular ducts (bottom), showing directional shift in the center of mass of the spatial intensity. Graphs show temporal profiles in the canalicular network (compare: Fig. S1B), spatial profiles in the IBD and canalicular network (compare: Fig. 1C). **D.** Simulation of photoactivation in the Canal of Hering, showing retrograde diffusion to the canalicular network (compare: Fig. 1D). All simulations were performed with experimentally derived flux parameters (D = 3 μm^2^/s, V = 0 μm/s in canaliculi, 1 μm/s in ducts). Photoactivated region: White dotted circle. Scale bar: 10 μm.

Simulations of repeated activation as described earlier (Fig. 1B) quantitatively reproduced the upper-bound of flux rates (Fig. 5B). Simulations of single photoactivations in the canalicular network reproduced symmetric canalicular dispersion and unidirectional displacement of intensity in the IBD (Fig. 5C) as was observed experimentally (Fig. 1C). The retrograde flux of fluorescence into the canalicular network from the CoH (Fig. 1D) was also accurately predicted by the simulations (Fig. 5D). The half-times of depletion in silico for a given area were similar to those observed in vivo (Fig. 5C, Fig. S1B), and varied proportionately to the area of photoactivation. In summary, the ‘first-principles’ simulation accurately recapitulated all experimental observations for using experimentally derived input parameters for diffusion and advection in the various zones. Thus, the simulation reinforced our findings of a diffusion-dominated canalicular network with negligible (if any) advection and an advection-augmented ductular system.

### Factors (not) influencing global canalicular clearance rates

Water influx due to the osmotic gradient created by bile salts, wherein the cholic acid moiety represents the osmotically-active moiety, is the presumed *raison d’etre* for canalicular advection. Since our results so far indicated no such advection in the canalicular network, we addressed the effect of osmosis on global clearance of the bile salt analog cholyl-lysyl-fluorescein (CLF) from the liver lobule under basal or stimulated extrahepatic bile flow (TCA infusion, secretin administration) conditions. In particular, the effect of TCA to increase extrahepatic bile flow is ascribed to increased canalicular flow due to the aforementioned osmotic gradient (2).

Following intravenous injection of CLF into the tail-vein, intravital time-lapse microscopy showed that the rate of CLF clearance from the canalicular network was unchanged between basal conditions or secretin- and TCA-stimulated conditions (Fig. 6A, B, D, Mov. S6). To rule out the possibility that the CLF molecule itself created a strong, perhaps maximal, osmotic gradient, we performed calibrated quantitative comparison of canalicular excretion of its fluorophore (fluorescein) with and without co-injection of its osmotically active component, cholic acid (15). Co-injection of cholic acid along with fluorescein did not significantly alter the kinetics of fluorescein excretion from the canalicular network (Fig. 6C, D, Mov. S7). These observations are incompatible with a role of osmosis in canalicular excretion. Since the ‘difference in osmotic potential’ assumption could not explain the differences in the half-life of clearance of CLF and fluorescein, we hypothesized that intrinsic transporter activity of the hepatocyte generates their different clearance kinetics. CLF and fluorescein differ in molecular size, structure and transporter specificity (16,17) and hence direct comparisons between the two compounds as substrates for transporter activity are complex. We therefore synthesized cholic acid derivatives CL-ATTO405 and CL-ATTO565, conjugated with 2 spectrally distinct, but chemically similar fluorophores. Co-injection of both compounds into the tail vein showed that hepatocyte export of CL-ATTO405 occurred at a much higher rate than that of CL-ATTO565. This shows that the transporter activity differs between the two compounds. However, in both cases, canalicular depletion closely followed the hepatocyte export kinetics. Thus, the higher export of CL-ATTO405 did not saturate the diffusive canalicular flux. Rather, hepatocyte export was the rate limiting step for both compounds, and canalicular intensity remained correlated to the intensity in the hepatocytes (Fig. S3C, Table S4). Taken together, these results show that for bile acid concentrations of up to 400 μM (calibration curve, Fig. S3A) the hepatocyte export activity, rather than osmotic potential, determines canalicular clearance rates of bile acid analogs.

**Fig. 6.**
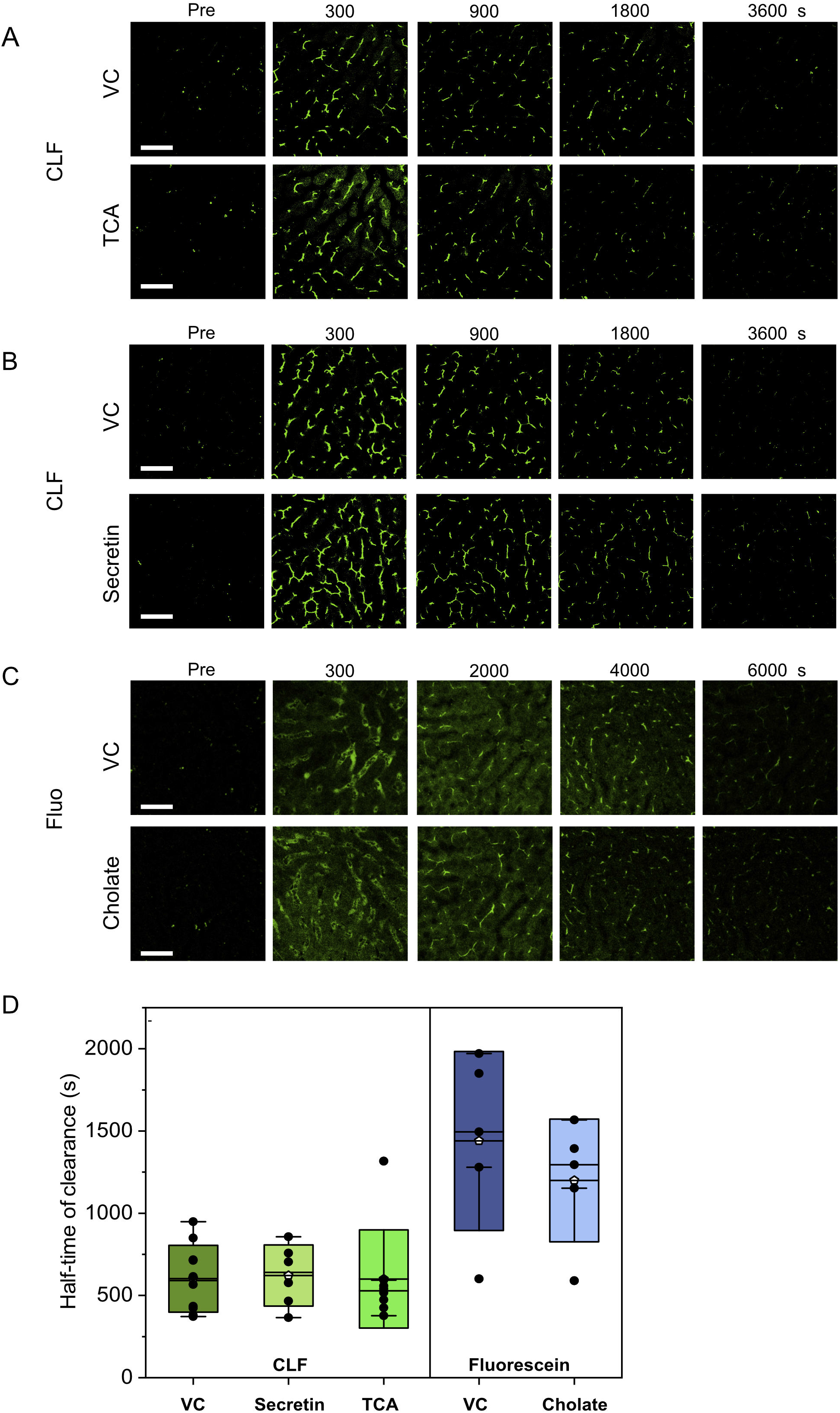
TCA, secretin and cholate do not influence canalicular clearance. **A.** Representative time-lapse images of the canalicular clearance of CLF with and without TCA infusion. **B.** Representative time-lapse images of the canalicular clearance of CLF with and without secretin administration. **C.** Representative time-lapse images of the canalicular clearance of fluorescein with and without cholate injection. **D.** Quantitative comparison of clearance kinetics under conditions described in A, B, C, for n > 6 mice each, with no significant difference due to secretin, TCA or cholate treatment for the respective fluorophore. Scale bar: 50 μM.

### Visualization of the lobule-wide diffusion gradient

A diffusive system such as the canalicular network should establish higher concentrations at the closed (pericentral) end than at the sink (periportal) end. Such a gradient is a necessary condition to drive diffusive flux out of the network. However, local concentrations in canaliculi depend not only on canalicular flux, but also on hepatocyte export. To investigate the influence of these factors, we measured the lobule-wide spatiotemporal distribution and clearance of carboxymethyl-fluorescein diacetate (CMFDA). Intracellular esterases cleave acetate groups from the non-fluorescent CMFDA which enters hepatocytes to generate the 400-fold brighter fluorescein molecule, which is excreted then into the canaliculi. Following tail-vein injection, fluorescein appears earlier and with higher intensity in periportal than in pericentral canaliculi (Fig. 7A, Movie S8). Within approximately an hour, the intensities in the periportal and pericentral canaliculi equilibrate to similar levels. These experiments show that the canalicular fluorescein excretion by hepatocytes occurs in a PV-CV gradient. This transport-imposed gradient would antagonize and mask a diffusion-derived CV-PV canalicular gradient. To compare the relative clearance rates in periportal versus pericentral canaliculi, temporal intensity profiles were normalized to their maximal values. Normalized profiles showed a higher depletion rate in periportal than pericentral canaliculi, hinting at a diffusion-mediated gradient masked by an antidromic transport gradient.

**Fig. 7.**
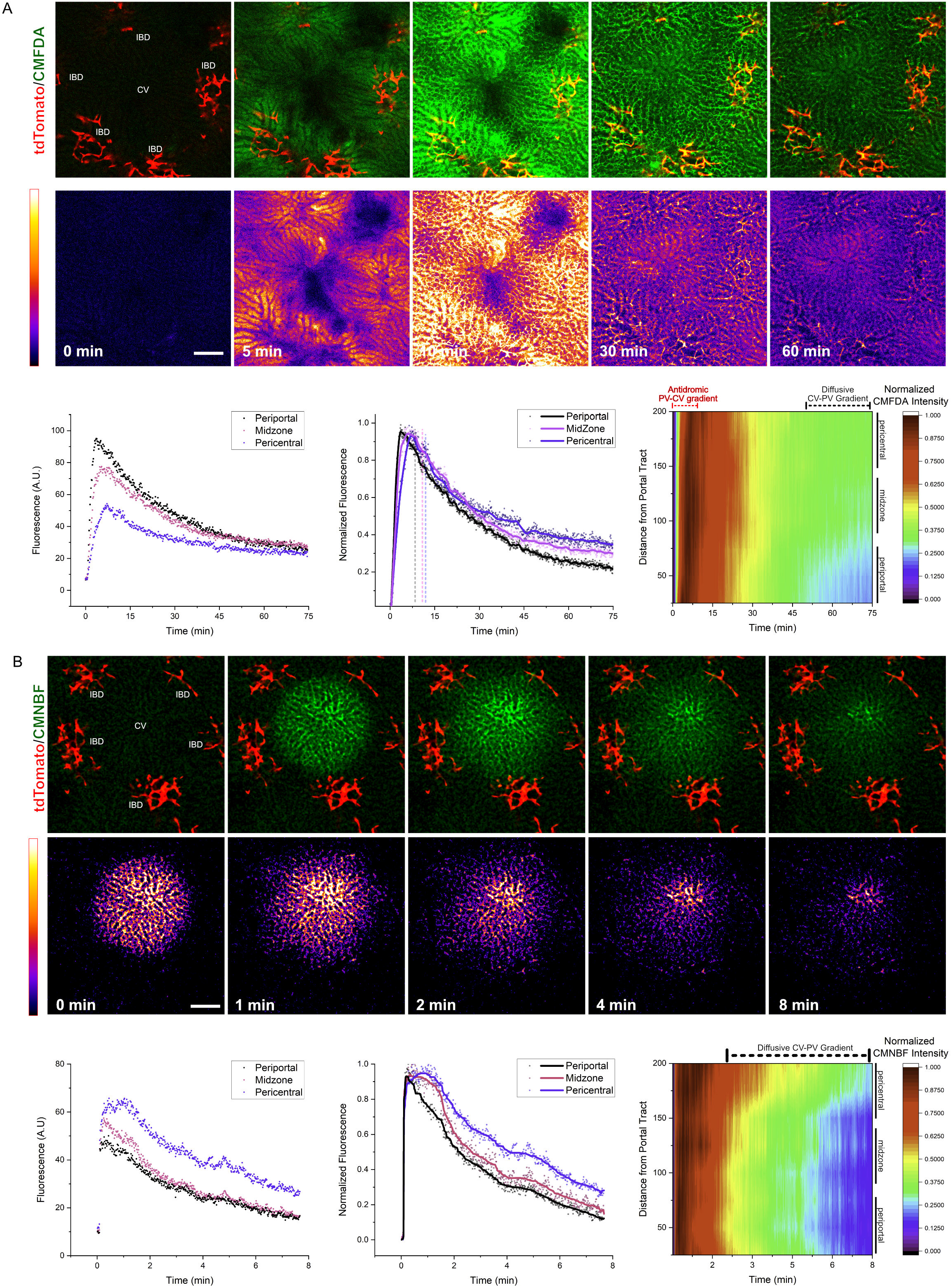
Visualization of lobule-wide bile flux gradients. **A.** Representative time-lapse imaging of a liver lobule with tdTomato-positive interlobular ducts (IBD, red), infused with CMFDA (green). Bottom row shows false color images for visualization of CMFDA intensity. Graphs show raw and normalized fluorescence intensity indicating higher depletion rate in the periportal versus pericentral zones. Contour plot shows the hepatocyte-transport induced antidromic PV-CV gradient and diffusion-derived CV-PV gradient appear over time. **B.** Representative time-lapse imaging of a liver lobule with tdTomato-positive interlobular ducts (red), infused with CMNB-Fluo after photoactivation in the entire lobule. Bottom row shows false colour images for visualization of CMNBF intensity. Graphs show raw and normalized fluorescence intensity indicating higher depletion rate in the periportal versus pericentral zones. Contour plot shows the appearance of the diffusion-derived CV-PV gradient appear over time. Scale bar: 100 μm. CV indicates central vein.

To demonstrate the existence of this diffusion-gradient, we used a method that allows the sole analysis of the influence of diffusion on the canalicular intensity, without the interference of the hepatocellular export. For this purpose, we photoactivated CMNB-Fluo lobule-wide approximately 1 hour after tail-vein injection, when the material is predominantly in the canaliculi and not in the hepatocyte cytoplasm. Immediately after photoactivation, a CV-PV gradient in canalicular intensity was visible which steepened over time (Fig.7B, Movie S9). After normalization to maximal intensities, the faster clearance rate of the periportal compared to pericentral canaliculi became even more apparent. Taken together, we conclude that diffusion indeed establishes a CV-PV canalicular gradient, but which is antagonized by higher hepatocellular export in periportal zone.

## Discussion and Conclusion

In this work, we combined advanced intravital imaging with simulation techniques in real geometries to investigate the flux characteristics of tissue microdomains. IVARICS is an adaptation of image correlation spectroscopy, wherein the decay in correlation between pixels due to fluctuation of the signal created by molecular flux is used to compute the rate of flux, with specialized corrections for periodic animal movements caused by its heartbeat, respiration and peristalsis. This allowed us to quantitatively decipher the mode of transport and determine its rate for small molecules in a sub-micron scale vessel network of arbitrary geometry. Utilizing FLAP studies, correlative imaging, rheological measurements and custom chemical probes, we have addressed a specific long-standing problem in liver physiology.

Diffusion-dominated canalicular bile flux contradicts the concept of advective bile flow in the canaliculi that has been reported in medical text books since decades. The osmotic theory of bile flux states that bile flows through the canaliculi countercurrent to the direction of sinusoidal blood flow, due to osmolytes including bile acids being actively exported into the canalicular lumen, simultaneously drawing water along the osmotic gradient (2). Since the canalicular network is closed at the pericentral end, water influx results in a unidirectional flow towards the bile duct. While the theoretical concept of the osmotic theory was proposed already in 1959 (18), corroboratory experiments were presented in the following decades, comprehensively reviewed by Boyer (2). A general feature of these experiments is the intravenous administration of compounds such as bile salts (8,19–22) with simultaneous measurement of increased extrahepatic bile volume and biliary compound excretion. Since bile acids and other organic solutes are secreted by hepatocytes into the bile canaliculi, and extrahepatic bile flow varied approximately linearly with amount of infused compound, it was concluded that the canaliculi are the anatomical structure where flow is initiated. Further, because the apparently linear relationship seemed to contain an inherent offset, the resultant extrahepatic bile flow was proposed to consist of a bile-salt dependent fraction which varies with the amount of administered bile acid, and a bile-salt independent fraction attributed to other osmolytes than bile acids, and/or to water secretion by cholangiocytes (2). As long as the focus is on macro-scale correlations between the infusion and clearance of compounds from blood and the excreted volume of extrahepatic bile as well as the excreted amounts of compounds, the osmotic flow concept creates no contradiction with experiment.

Yet, it has also been recognized early on that the postulated linear relationship between infused compounds and extrahepatic bile flow is only approximate (18,19,22–24). This unsatisfactory linearity has been dealt with by introducing a ‘correction for delay in transit in the biliary tree’ when comparing two compounds (22). Nahrwold and Grossman (24) detected threshold effects in the induction of increased extrahepatic bile flow – an observation incompatible with the osmotic theory without further assumptions. Wheeler (19) specifically recognizes through theoretical considerations that diffusion could also explain the experimentally observed bile salt excretion, and laments that direct canalicular flow measurements are not available. In later work (25), Wheeler and Ramos cautiously state that the osmotic theory “would provide an adequate explanation” but agree that for such observations, it does “not seem necessary to invoke a specific mechanism for active water transport.” Indeed, as long as the focus is on macro-scale extrahepatic bile flow, the precise mechanism of canalicular transport (diffusion/advection/convection) is not relevant in order to derive these correlations. However, this mechanism becomes relevant for other questions such as the influence of pathological biliary pressure in canaliculi or the site of action of choleretic drugs.

For example, the concept of ‘osmotic canalicular bile flow’ has been used to mathematically model the velocity of canalicular bile flow and the resulting pressure, claiming a bile velocity in periportal canaliculi of up to ~12μm/s in mice (3). Based on the osmotic concept of bile flow, another study simulated pericentral canalicular pressure in individual patients with NASH and calculated increases above 3000 Pa due to altered canalicular morphology. This was interpreted ‘as a new component of the NAFLD pathophysiology’ (26). The applicability of these models crucially depends on what the *actual* molecular mechanisms of canalicular flux are. Leaving such conclusions (26) unchallenged may encourage the development of unfounded therapies that focus on modifying canalicular flow and pressure.

To our knowledge, no previous study in these intervening 40 years has measured flow and diffusion in bile canaliculi and interlobular bile ducts. Therefore, the part of the osmotic theory that bile flows countercurrent to sinusoidal blood in the canaliculi remained untested. In the present study, we directly investigated the flux mechanism in these microdomains and exclude the canaliculi as the anatomical site where flow originates. The osmotic concept did not consider that the transport of solutes may also occur through a standing water compartment by molecular diffusion. The techniques developed in our study allow to experimentally quantify flow and diffusion in bile canaliculi and ducts of intact livers *in situ*, and thus to validate or falsify previous model predictions.

Our results show that bile acids are transported primarily through diffusion in the canaliculi, while being augmented by directional advection in the interlobular ducts. Through these observations, we deduced the compartmentalization of the liver into a diffusion-dominated canalicular domain where hepatocytes actively secrete small molecules and generate a concentration gradient that drives diffusive flux towards the IBDs (Fig. 8A). In IBDs, regulated and inducible water influx creates unidirectional advection which augments the diffusive flux. Water influx into ducts can be caused either directly by hormones such as secretin (27), or indirectly by bile salts themselves as demonstrated for TCA (10), explaining the corresponding increase in extra-hepatic bile flow. This inducible increase in advection is restricted to the bile ducts, while the canalicular network remains nothing more than a standing-water zone.

**Fig. 8.**
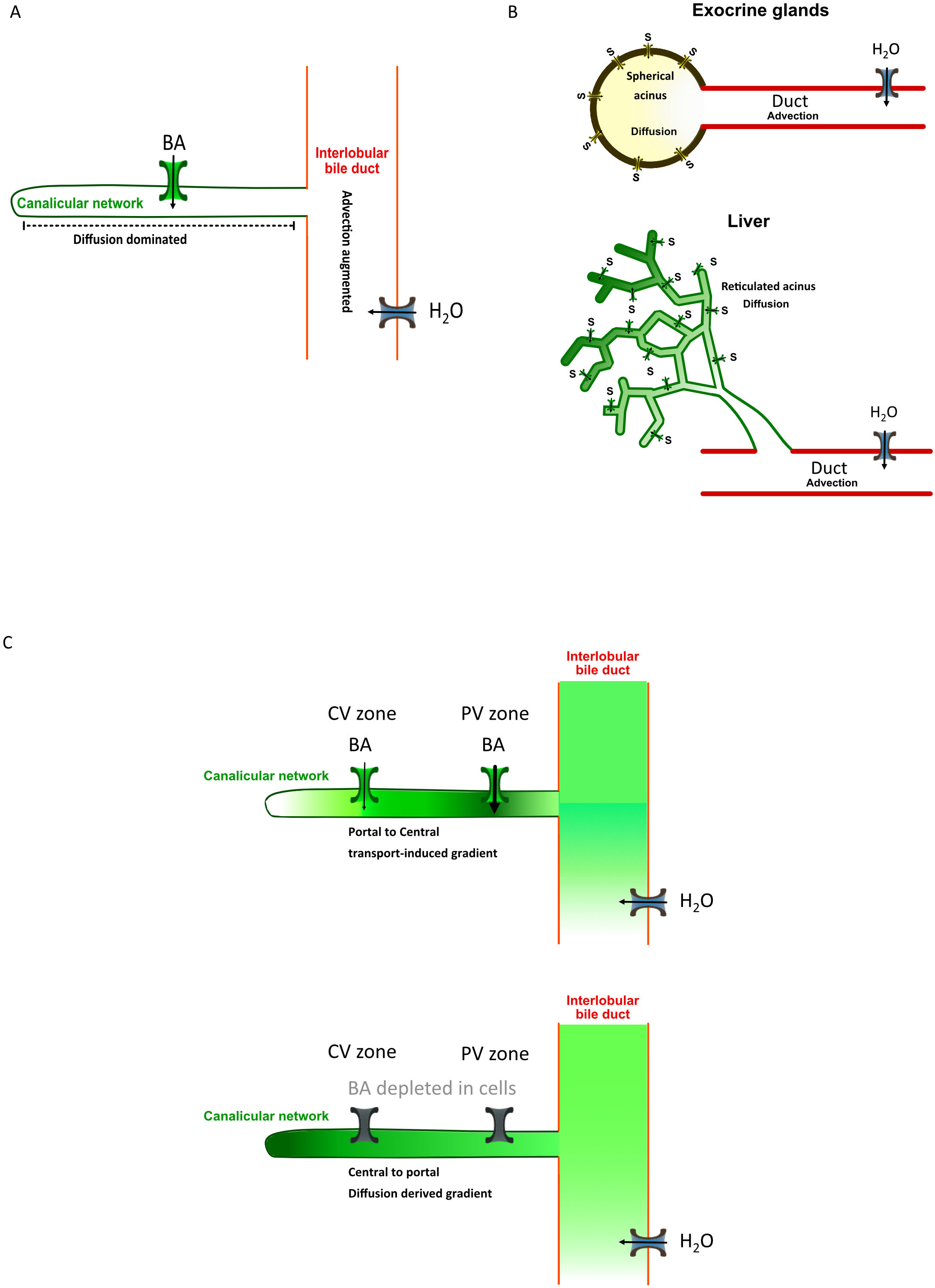
Flux organization in the liver. **A.** Schematic representation of the diffusion-dominated domain, the canalicular network and the advection-augmented domain, the interlobular bile duct in the liver. Bile acids (BA) are secreted in the diffusion-dominated zone, while water is secreted in the advection-augmented zone. **B.** Comparison of liver tissue topography to that of a generic exocrine gland in terms of solute secretion zone and water secretion zone, implying that the liver canalicular network is analogous to an exocrine acinus, only modified geometrically from a sphere to a reticulated network. **C.** Zonation of canalicular bile acid concentrations. Upper panel: a central to portal gradient is established, if CMNB Fluorescein is photoactivated only in the canaliculi. Lower panel: Establishment of a portal to central gradient due to the higher excretion rate of periportal compared to pericentral hepatocytes.

We note that the principle of liver physiology presented in this work, with a diffusion-dominated standing-water domain connected to an inducible flow-augmented excretion duct, seems a general principle applicable to exocrine glands (Fig. 8A, B). All exocrine glands contain an acinus, surrounded by epithelial cells that secrete their products into the acinar lumen. The lumen represents a reservoir where products accumulate to high concentrations, which is connected to an excretion duct in which flow can be induced on demand. A similar design enables the function of all ~42 types of mammalian exocrine glands (28), of which the liver is the largest. For example, lactating mammary glands synthetize milk proteins consistently in the acini, but secretion of milk is induced through the action of hormones such as prolactin on the mammary ducts. From this perspective, the canalicular network may be envisaged as a highly reticulated form of the conventional spheroidal acinus (Fig. 8B). The reticulated geometry is probably imposed due to the necessity of hepatocytes to maximize their contact area with the also reticulated sinusoidal network, which dictates their positioning. This modified acinus is connected to flow-augmented bile ducts of the liver, following the general principle of exocrine glands.

The reticulated acinus (bile canaliculi) of the liver implies reduced degrees of freedom for molecular flux, compared to a spherical acinus and contributes to complex zonation phenomena. Due to diffusion, a gradient with higher bile acid concentrations at the pericentral closed end of bile canaliculi is established, while concentrations at the periportal open end are lower (Fig. 7D, upper panel). However, the diffusive CV-PV gradient is antagonized by an antidromic gradient, caused by the higher canalicular excretion rate of PV than PC hepatocytes (Fig. 7E, lower panel). The preferential PV excretion of bile acids and other compounds is in agreement with previous studies, injecting [^3^H]taurocholate into the portal vein of isolated perfused rat livers, which led to a steep PV-CV gradient of radioactive tracer in liver tissue (29). Vice versa, the same authors also show that retrograde perfusion with [^3^H]taurocholate injection into the central vein led to a CV-PV gradient (30). These experiments suggest that uptake from the sinusoid into hepatocytes depends on the sinusoidal blood concentration that decreases during passage through the lobule. As the excretion and the diffusion driven gradients are antidromic (Fig. 8C), it is difficult to visualize the diffusive gradient by the use of conventional fluorescent ‘always-on’ tracers such as CLF or CMFDA. However, the diffusive CV-PV gradient can be demonstrated by photoactivatable fluorophores, e.g. CMNB-Fluo, if photoactivation is performed when most of the compound has been excreted to the canaliculi. Preferential excretion of bile acids in the PV zone leads to diffusive equilibration into the duct, but also rest of the canalicular network towards the CV zone. This retrograde loading means that bile acids are kinetically-trapped in the network at a relatively high concentration. In case of a demand of bile acids in the intestine after food intake, low pH in the stomach (and duodenum) triggers S-cells in the mucosa of the duodenum to release secretin into the blood, which causes water influx into bile ducts via cholangiocytes and thereby flow of bile to the gall bladder, which releases its content into the duodenum under the control of other hormones, such as cholecystokinin(31). Therefore, hepatocytes represent a domain of continuous bile acid synthesis, canaliculi serve as an intermediate store, while the duct acts as a logistic center that serves the food intake dependent needs of the intestine for bile acids. Photoactivation in the present study was performed for small regions, (e.g. radius = 10 μm), to analyze local flux properties, as well as for large areas comprising entire lobular sections (radius=200 μm). These experiments clearly demonstrated that the diffusion-dominated flux in canaliculi, and advection-augmented flux in ducts holds at the scale of the lobule. Diffusion gradients could be demonstrated over the lobular canalicular network, while vectorial transport was observed in the mesh of interlobular bile ducts around portal veins. Photoactivation of entire lobules also demonstrated the sink function of interlobular duct meshes since canalicular intensities decreased faster in the neighborhood of these structures, while fluorescent material was cleared more slowly in the pericentral zone. It should be considered that a flow-dominated mechanism would create a gradient of the opposite orientation with lower concentrations in the lobule center (CV zone).

Why does the excretion of bile acids into the canaliculi, does not cause any measurable advection? Advection is not the inevitable result of the transport of osmotically active compounds into a biological network with a closed end. Bile acids, as hydrophobic molecules, are incorporated in phospholipid micelles, preventing them from exerting their full osmotic potential (15). Rather, the effect of inorganic ionic balance is clearly known to play a complex role in bile acid excretion and extrahepatic bile flow (25,32). Yet, the secretion of bile acids by hepatocytes will necessitate simultaneous uptake of anions such as Cl^-^ to maintain plasma membrane potential (33). Homeostatic ion exchange may counteract the development of any net osmotic pressure in the canalicular lumen. The present analyses show that advection does not exceed 0.02 μm/s in the canalicular network, suggesting that as far as a flow is concerned, the osmotic effect of osmotic ion exchange is exerted primarily in the biliary ducts - not the canalicular network.

Canaliculi are known to contract autonomously and *asynchronously* at a low frequency of 6 events/hour through a calcium-dependent mechanism (34). This has been proposed to cause propulsion of bile in the canaliculi. Again, the absence of any significant advection in the canalicular network indicates that in the absence of *synchronized* contractions along the CV-PV axis, merely mixing (rather than pumping) of canalicular contents may occur.

One of the surprising aspects of flux in the canalicular network is the low diffusion coefficient of 2.4 – 4.6 μm^2^/s (CI: 95%), compared to ~200 μm^2^/s in water in vitro (Fig. S3B). Biliary constituents such as albumin and phospholipid micelles may interact with bile acids to reduce their diffusivity. The apparent value of a diffusion coefficient may represent anomalous diffusion processes such as (i) polydispersive diffusion (35) - the convolved diffusivity and binding interactions of multiple populations ranging from micellar to freely diffusing bile acid molecules, or (ii) Knudsen diffusion - due to the relatively small size of the diffusible volume compared to the mean path length. Nonetheless, even the relatively low apparent diffusion coefficient is sufficient to account for observed lobular clearance rates, as verified by the diffusion simulation in realistic canalicular geometries. Moreover, we show that hepatocyte cytoplasmic concentration is the strongest factor of influence of canalicular concentrations for bile salt analogs over the entire clearance period (Fig. S3C). Thus, we corroborate previous reports (36,37), that energy-dependent export from hepatocytes to the canalicular lumen is a stringent bottleneck for bile acid clearance. Diffusive flux through the canalicular network in not limiting. Due to high periportal transport rates, high concentrations of bile acids are created near the interlobular bile ducts, reducing the distance bile acids have to cross by diffusion. Effectively, the functional diffusive zone may be smaller than the actual anatomical size of the lobule.

In summary, this work demonstrates the flux mechanisms that operate in the various liver microdomains and provides a generalized methodology to perform quantitative intravital evaluation of flux-associated organ function. The liver follows the principle of parsimony – the energy invested in active transport across membranes is entropically dissipated for canalicular diffusion, to be augmented by energy-driven regulated flow in the ducts as required for physiology.

## Author contributions

Conceptualization, Methodology, NV, JGH, DD, FJ, CH; Investigation, NV, GG, FJ, MR; Writing – Original draft, NV, JGH, DD; Writing – Review and Editing, NV, JH, JF, DD; Resources, FG, SJ, GW, HH; Funding acquisition, JH, DD, NV; Data curation, Validation, JF, BBT, AG, KW, AF, SH, IVC, NB; Supervision, Project administration, JH, DD, NV, HH, SH, CH.

## Supporting Information

### Materials and Methods

#### Animal source, handling and preparation for microscopy

Male C57BL/6N mice, 8-10 weeks old were obtained from Charles River (Sulzfeld, Germany), HNF1b/CreER-reporter mice were obtained from Dr. Fabian Geisler. These ***R26Tom Hnf1b-CreER*** mice allow the labelling of bile duct cholangiocytes through HNF1b-dependent tamoxifen-induced Cre recombination of o the R26 promoter with the tdTomato locus. The specificity of the HNF1b dependent tdTomato expression in cholangiocytes has been extensively validated through CK19 immunostaining and other markers, with a false positive rate of less than 1/8000 (4,38), and this mouse model is commonly used for cholangiocyte lineage tracing.

Animal preparation for intravital imaging was performed according to a published method (39). Briefly, mice were anaesthetized with a combination of ketamine (64 mg/kg), xylazine (7.2 mg/kg) and acepromazine (1.7 mg/kg) given intraperitoneally. The abdomen of the animal was shaved, and a ~1.5 cm midline incision made to expose the xiphoid process which was retracted to allow dissection of the falciform ligament. The left lobe of the liver was gently exteriorized, and the animal inverted onto a 49 mm x 74 mm glass coverslip of 0.17-0.16 mm thickness (Ocon 159, Logitech, Glasgow, UK) mounted within a custom-made imaging platform. The liver was covered with sterile saline-soaked gauze to prevent dehydration. Additionally, a subcutaneous injection of 0.5 ml saline was given. During imaging mice were kept under ongoing anesthesia of 1-2% isoflurane in a climate-controlled chamber.

All experiments were approved by the local animal protection agency.

##### Intravenous administration

###### Bolus injection

1. CLF (Corning, Product no. 451041), fluorescein (Thermo-Fischer Scientific, Cat no. F1300) and CMNB-Fluo (Thermo-Fischer Scientific, Cat. no. C20050) were dissolved in phosphate-buffered saline (PBS) to a concentration of 1 mg/ml and administered intravenously through a tail-vein catheter at a dose of 1 mg×kg^−1^.
2. Lyophilized human Secretin (TOCRIS, Cat. No. 1918) was dissolved in PBS to a concentration of 75 U/ml and administered intravenously through a tail-vein catheter into mice at a dose of 3.0 U×kg^−1^.

###### Infusion

For experiments involving infusion, intravenous tail-vein catheters in mice were infused via a syringe pump (Perkin-Elmer).

1. CLF and fluorescein were dissolved to a concentration of 1 mg/ml in PBS and infused at a rate of 50 μL×kg^−1^×min^−1^.
2. TCA (Sigma-Aldrich, Cat. no. 86339) was infused at a rate of 660 nmol×kg^−1^×min^−1^ (40). The total infused volume was not allowed to exceed 200 μL.
3. CMNB-Fluo-Dextran was utilized for photoactivation in blood vessels, and synthesised as follows: <u>Reagents and Materials:</u> CMNB-caged fluorescein SE (Invitrogen C20050) Aminodextran, 10 kDa (Invitrogen D1860) Zeba Desalt Spin Column (Pierce 89889) 0.1 M Na2B4O7, pH 8.5 15-mL polypropylene conical tubes (such as Falcon 352097) 1.5-mL microcentrifuge tubes Vortexing mixer Centrifuge capable of spinning15-mL tubes at 1000 x g <u>Procedure</u>
  1. Measure out 3.5 - 4 mg of aminodextran and add into the Invitrogen-supplied tinted tube containing 1 mg of CMNB-caged fluorescein SE. In our hands this ratio gives an average loading of ~ 2.5 dye molecules per dextran.
  2. Add 500 μL of 0.1 M Na2B4O7 (sodium borate) buffer to the tube.
  3. Cap and vortex for 30 seconds to dissolve the aminodextran and caged fluorescein.
  4. Let react overnight on a vortexing mixer.
  5. Twist off the bottom closure on a Zeba spin column. Place column in a 15-mL conical tube.
  6. Centrifuge at 1000 x g for 2 min with the dot facing the center of the rotor. Use column immediately after compacting.
  7. Place compacted column in a new 15-mL conical tube.
  8. Pool reaction mixture and transfer to the center of the compacted column resin bed. Replace cap.
  9. Centrifuge at 1000 x g for 2 min with the dot facing the center of the rotor. After the spin, the eluent and the top of the column will be roughly the same shade of yellow in colour.
  10. Transfer the yellow eluent (~350 μL) to a 1.5-mL microcentrifuge tube.
  11. Lyophilize to dryness
  12. Re-suspend the resulting ~ 2-3 mg of pale-yellow foam in water to a final concentration of 1% w/v.
  13. Store this solution at −20 °C in dark tubes.

#### Extrahepatic bile collection

Collection of extrahepatic bile was achieved by ligation of the common bile duct, followed by catheterization of the extrahepatic bile duct just before it enters the gall bladder fundus (41). Bile was collected from anaesthetized mice, with or without secretin/TCA administration, in preweighed Eppendorf tubes and the quantity collected was measured every 10 minutes. Results are reported as amount of bile (μL) collected in unit time (min) for unit weight of the liver (g).

#### General imaging setup

Imaging was performed on a Zeiss LSM 880 confocal microscope equipped with a 40× N.A 1.4 PLAN-APOCHROMAT oil-immersion objective, a 20× N.A 0.8 PLAN air objective, 2 GaAsP and 1 PMT detector, an automated stage and climate-control chamber. Excitation was achieved with the Argon 488 laser line for CLF, fluorescein and CMNB-Fluo, the HeNe 561 line for tdTomato. Emission bands for CLF, fluorescein and CMNB-Fluo were set at 498-551 nm, while 571-630 nm were used for tdTomato. Imaging resolution was set at 512 × 512 pixels, with 8-bit acquisition depth and a frame rate of 1s. According to the required field-of-view, i.e, (250 μm)^2^ for local activation and imaging or (1.1 mm) ^2^ for lobule wide activation and imaging, either the 40× or 20× objective was chosen as appropriate.

#### Photo-activation

##### Imaging

Photoactivation of CMNB-Fluo was performed with a 405 nm diode laser in a manually defined circular ROI of 5-40 μm radius. Typical in vivo photoactivation sequences consisted of images acquired without activation for 5 frames at 1s/frame, followed by 2s of photoactivation in the ROI and subsequent imaging for upto 10 min min at 1s/frame. For photoactivation in the blood-vessel, frame time was reduced to 100 ms/frame and acquisition resolution was dropped to 128×128 pixels to compensate for fast blood flow. For in vitro photoactivation, 10 μm CMNB-Fluo was dissolved in distilled water or mouse bile (collected as described above) and frame times were reduced to 100 ms/frame with a acquisition resolution of 128×128 pixels, to compensate for the faster diffusion of small molecules in water.

##### Analysis

###### 1. Diffusion

The centroid of the photoactivated area was identified using 1D-line profiles of the first photoactivation image in x and y. Gaussian fitting of these line profiles was used to identify the full width half-maximum (FWHM) of the profiles in X and Y.

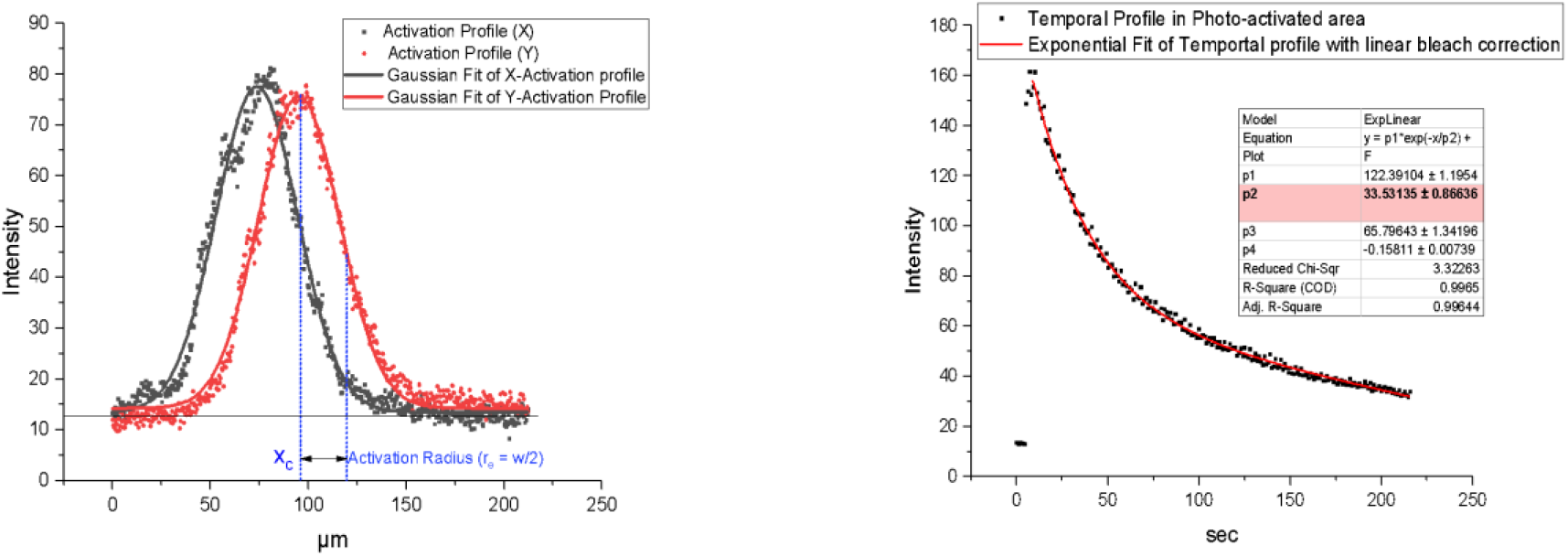

The effective radius r_e_ was taken as FWHM/2. Nominal radius r_n_ corresponds to the user-defined photoactivation area and was derived directly from instrument metadata.

For determination of half-life, the temporal decay of mean intensity in the activated area was measured and fitted to a mono-exponential decay and linear function. The linear function accounted for bleaching that may occur over the monitoring interval.

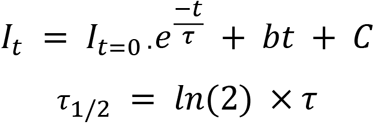

where *I_t_* is the intensity at time t, and b is the bleaching constant, C is the offset in intensity, *τ* is the exponential time constant and *τ*_1/2_ is the half-life of decay.

The diffusion coefficient, D, was then determined using the Soumpasis method (9) as:

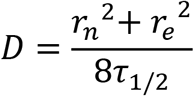

###### 2. Advection

In order to determine advection velocity, the location of the centroid of intensity in the spatial activation profiles (see above), were obtained over all frames in a time-lapse sequence, following photoactivation. The mean-square displacement of the centroid per unit time gave the velocity v per frame.

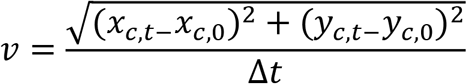

Where x_c_ and y_c_ are the location of the centroids at time t and t=0 and Δ*t* is the time between frames. The mean velocity over all the frames in the time-lapse sequence was derived and reported.

Analysis for performed using FIJI/ImageJ (http://www.fiji.sc) through manual tracking of the fluorescence centroid over time for advection velocity, and determination of intensity decay over time to determine a half-life and area of activation for diffusion coefficients.

#### IVARICS

##### Theory

Spatial and Temporal Image Correlation Spectroscopy (RICS/TICS) is part of a family of methods where the autocorrelation of fluorescence signals is analysed to extract information regarding molecular events. Fundamentally, the magnitude of autocorrelation decays in time as molecular events progressively change the distribution of fluorophores, i.e, decorrelates them.

ICS exploits the raster imaging design of a confocal scanning microscope, wherein pixels are acquired sequentially to form a line, lines are acquired sequentially to form an image and images are acquired sequentially to produce a stack. Thus, the temporal autocorrelations in time produced by underlying molecular events are encoded in the spatial dimensions of the image (or stack). Autocorrelation (G (x,y,t)) curves obtained over space can then be used to derive information about the molecular processes involved.

The generalized discrete normalized spatiotemporal autocorrelation function for a fluctuating signal I with dimensions X, Y, T is defined explicitly as:

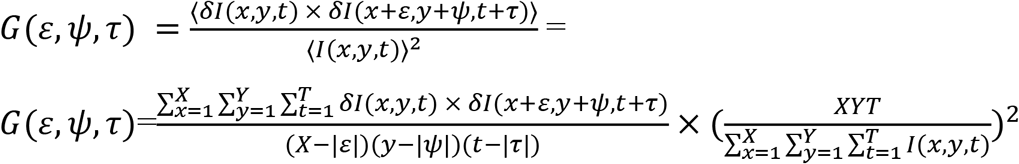

where the fluctuation *δI*(*x*, *y*, *t*) is the deviation from the mean *I*(*x*, *y*, *t*)of the signal *I*:

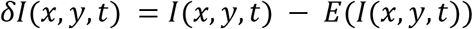

ε,ψ,τ are the correlation lags and E(I(x,y,t) represents the mean intensity of the pixels in dimensions X, Y and T, respectively.

##### Considerations for in vivo ICS imaging

The precise magnitude and shape of the autocorrelation function during in vivo imaging is affected by several experimental and mechanistic parameters. These parameters, and procedural aspects for each of the methods is described below.

- <u>Concentration of fluorophores</u>: The magnitude of the autocorrelation function G varies inversely with the number of molecules (N). The measured autocorrelation is scaled by (1/N). The in vivo concentration of fluorophores in liver canalicular compartments is largely determined by the amount injected into the animal and the degree by which hepatocytes take up the fluorophore from the blood and export it into canaliculi.
- <u>Fluorophore intermittency</u>: Most molecular fluorescent emitters switch between bright and dark states intermittently at timescales of a few seconds. Since the pixel acquisition times utilized in RICS typically range from 2-32 μs, RICS sequences are not significantly affected by fluorophore blinking.
- <u>Binding events</u>: As most fluorophores are hydrophobic to some degree, molecular binding events with hydrophobic structures such as membranes occur in biological samples. These events typically reduce the mobility of the fluorophore, and in the present work are assumed to affect molecular flux, which may influence the apparent diffusion coefficient.
- <u>Instrumental correlations from detector noise</u>: Dark current in detectors used on confocal microscopes is usually correlated and may contribute to the autocorrelation. The absence of correlated dark current-related noise in the GaAsP detector used in our setup was confirmed by acquiring image sequences without laser illumination and confirming that the autocorrelation function has a value of 0 across all pixel lags. Shot noise from the detector is easily eliminated by excluding the (0,0) point from the RICS fitting.
- <u>Instrumental correlations due to scanner movement</u>: The regular movement of the scanner creates autocorrelation that is convolved with the autocorrelation due to molecular fluctuations. Ths autocorrelation due to the scanner movement is given as:

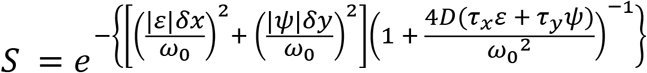

where ω_0_ is the beam-waist in x and y,

ε and ψ are the spatial lag in pixel in x and y respectively,
δx and δy are the pixel dimensions in x and y respectively,
D is the diffusion coefficient,
*τ*_x_ and *τ*_y_ are the pixel-dwell times and line times in x and y respectively.
- <u>Instrumental PSF size and shape</u>: Confocal point-spread functions (PSFs) are theoretically expected to be an elliptical 3D Gaussian, which is symmetric in the x and y directions (beam-waist) and elongated in the z-direction (axial distance). The dimensions by which the PSF affect the autocorrelation function are accounted for by the ω_0_ and ω_z_ parameters representing the beam-waist and axial-distance, respectively. For measuring autocorrelation, it is necessary to set the pixel dimensions δx and δy < ω_0_. In practice, the PSF may also be asymmetric due to inconsistencies in component manufacturing or beam alignment of the microscope. These asymmetries are represented as a shape-factor γ. For a theoretical confocal PSF, γ = 2^(−3/2)^ = 0.3535, and ω_0_ and ω_z_ are in the range of 0.4μm and 0.8 μm at the focal point. However, these parameters are substantially affected during in vivo imaging due to tissue scattering, with beam-widths expanding upto 4-fold and 5-fold in the (x,y) and (z) directions respectively (42). Since it is not possible to precisely determine the changes in these parameters in a live animal during intravital imaging, ω_0_ and ω_z_ are left as free parameters with constraints rather than as fixed parameters during fitting (see below).
- <u>Presence of immobile or slowly moving populations</u>: Slow moving structures or immobile populations which display little to no fluorescence fluctuations can significantly affect the autocorrelation function by artificially stretching the autocorrelation curve. The influence of these structures is corrected by subtracting the moving average of the stack from each frame, and rescaling the autocorrelation function. This operation represents a high-pass filter that removes slow or zero fluctuations from the image sequences. The moving average for a raw intensity image at time t is defined as:

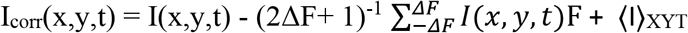

where I(x, y, t) is the frame F at time t, ΔF is the moving average window size, and 〈I〉_XYT_ is the average intensity of the entire stack. The autocorrelation function G obtained from I_corr_(x,y,t) is re-scaled to compensate for the net reduced intensity due to the subtraction of the mean as follows:

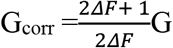 For all experiments in the present work, we used the moving average window size of ΔF = 1.
- <u>Animal movements</u>: In vivo imaging sequences are affected by movements in the organ of interest in the animal due to breathing, heart rate and intestinal peristalsis. These movements typically occur in the range of 14 ms −1s in a mouse. To avoid these artifacts in the spatial autocorrelation, we restricted the maximum pixel acquisition time to 8 μs, and the maximum image acquisition time to 250 μs. Since these movements occur on a slower timescale compared to molecular fluctuations, the high-pass filter applied to remove slowly moving structures effectively filters of the effect of animal movements. The temporal autocorrelation of the images is more significantly affected by animal movements despite correction by moving average filtering. Due to their periodic nature animal movements manifest as recurring spikes in the temporal autocorrelation of a stack, despite a general decay in the correlation amplitude. A band-pass FFT filter of frequency equivalent to 1/4Δt was applied to compensate for animal movements, where Δt is the time lag between frames (Fig. S2). Utilization of such a filter, however, can have the undesirable outcome of removing the autocorrelation decays of the flux processes we are interested in. As a result, the filter must be applied judiciously, manually and only ICS sequences where mouse movement occurs at frequencies different from the fluctuation frequency were further analyzed.
- <u>Irregular regions of interest</u>: Generally, ICS is performed on square regions where the number of times a particular spatial lag is sampled is a linear function. Since the liver canalicular structures and bile ducts measured are not conducive to assigning square regions due to their irregular shape, we adapted the Arbitrary region RICS algorithm (ARICS) (13), wherein the autocorrelation function G in a particular irregular structure is normalized to the number of times each spatial lag is sampled by calculating the autocorrelation of a binary mask of the irregular structure. In our case, time-lapse sequences were then thresholded for fluorescent structures representing the canalicular network or interlobular ducts to obtain the respective binary masks. Spatial autocorrelation maps were calculated from the binary masks containing exclusively the ‘shape-autocorrelation’ without information on the flux. According to the ARICS algorithm, these sets of autocorrelation maps were then used to derive the autocorrelation representing only the fluorescence fluctuation due to the flux mechanism, as follows:

∘ High-pass filter for removal of slow/immobile structures with ΔF = 1 was applied to obtain the corrected RICS sequence.
∘ For each frame image I, a binary mask M was generated using Otsu thresholding, where areas representing the biological structure of interest were valued as 1, while the background was valued as 0.
∘ The masked image (I × M) was computed through pixel-wise multiplication of I and M.
∘ The fluctuation image F can now be defined as the element difference of all nonzero pixels in I × M and the mean of all non-zero pixel in I × M:

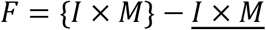
∘ To avoid boundary and aliasing effects, M and F were 0-padded with (2 × dimension + 1) zeros in the x,y, and z dimensions and the autocorrelation A(M) and A(F) was calculated using the Wiener–Khinchin theorem. Briefly, the autocorrelation of a function is the **real part** of the **product** of the **Fourier transform of the function** with the **complex conjugate of the Fourier transform of that function**.

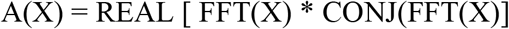
∘ The unnormalized autocorrelation due to fluctuation G is now defined as:

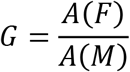
∘ To obtain the spatial autocorrelation function: **G_RICS_**, the spatial autocorrelation at each time point is normalized with the average squared intensity of the pixels in (I × M) at that time point. The spatial autocorrelations for all frames are averaged to obtain the normalized autocorrelation function G_norm_.
∘ G_norm_ is rescaled to correct for moving average subtraction (as stated above) to obtain the final autocorrelation function G_RICS_.
∘ To obtain the temporal autocorrelation function **G_TICS_**, the value of G at spatial lag (0,0) was calculated as above and normalized by the average squared intensity of every pixel in (I × M) over the entire stack.

The spatial autocorrelation in the y-direction was utilized due to few pixels in the x-direction for canaliculi, and was fitting to a 1-population 3D-diffusion function. The temporal autocorrelation was fitted to a 1-population 2component diffusion and velocity function.

###### Limitations of ICS for in vivo imaging

While ICS represents a powerful method for accessing molecular flux mechanisms in live animals, we encountered several challenges in its application. These are summarized below:

1. A priori knowledge: ICS sequences require some knowledge of the range in which flux is expected, as image acquisition parameters are set accordingly. For a given set of image acquisition parameters, it is possible that faster and slower autocorrelations arising from different populations are not represented well in the experimental autocorrelation decay. In the case of in vivo imaging, where biological fluids typically contain multiple diffusing populations, ICS yields only an apparent diffusion coefficient. Further, as in our case, a balance has to be struck between imaging speeds that allow measurement of the autocorrelation decay of molecular flux, while avoiding the effect of animal movements that occur over longer time periods. This is not always possible.
2. Dependence on non-linear fitting: ICS data provides diffusion coefficients and advection velocities from non-linear fitting of the autocorrelation decay. Diffusion results in hyperbolic autocorrelation decays, while advection results in Gaussian autocorrelation decay (43). An experimental in vivo autocorrelation curve with diffusion and velocity is therefore a convolution of hyperbolic and Gaussian decays Fitting of these curves is nontrivial and requires the utilization of global fitting procedures for a robust fit.
3. Structure effects: The geometry of the structure in which flux occurs also affects the autocorrelation function. In relevance this work, a single canaliculus is a tubular structure of only upto 0.8 μm in diameter, but extending upto 15 μm in length. Thus, the number of pixels along the diameter is much fewer and usually not sufficient to obtain a good autocorrelation function, restricting us to using the line correlation instead for RICS analysis (44). Further, the geometry of the biological structure may also affect
4. Requirement of a representative model: The autocorrelation decays are affected by several parameters such as binding and unbinding, number of populations with flux and the flux speeds, geometry of the

##### Imaging and Preprocessing

In addition to standard imaging parameters described previously, the following specific imaging parameters were: pixel dwell-times ranging from 2-32 μs, and pixel sizes adjusted for 2-4× oversampling in XY. Microscope pinhole size was adjusted to provide a confocal Z-depth that encompasses the canaliculi (~0.5 μm). Rapid scanning was performed for 20 frames.

Typical representation of metadata relevant for IVARICS is provided below:

File format: CZI
Dimensions - X: 128.0 Y: 128.0 T: 20.0
Pixel size: 1.66053164193e-07 m
Pixel dwell-time: 8.24242424242e-06 s
Scan line time: 0.00108503030303 s
Frame time: 0.316509090909 s
PSF - beam waist: 324.676083907 nm
PSF - axial distance: 865.238181818 nm

Raw image stacks were processed for removal of immobile populations and assigning arbitrary regions for analysis as described above.

##### Analysis of Molecular Flux by Diffusion and Advection

Fluctuations of the fluorophore intensity at each pixel occur due to molecular flux of fluorophores. Autocorrelation in these fluctuations over time and space can be used to compute diffusion coefficients and advection velocities through ICS analysis (45).

For a single population of fluorophores with diffusion and advection, the spatial autocorrelation due to diffusion is given as:

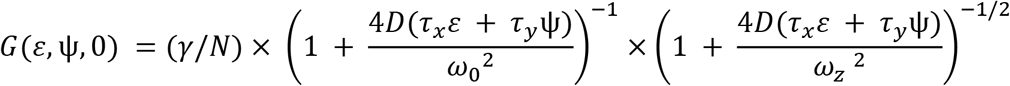

The temporal autocorrelation of the image sequence is given as:

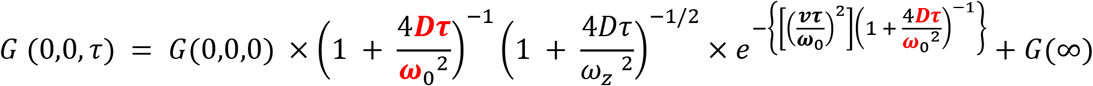

where ω_0_ is the beam-waist in x and y, and ω_z_ is the axial distance

ε and ψ are the spatial lag in pixel in x and y respectively,
δx and δy are the pixel dimensions in x and y respectively,
*τ*_x_ and *τ*_y_ are the pixel-dwell times and line times in x and y respectively,
τ is the lag time between frames,
γ is the PSF shape factor,
N is the number of fluorescent entities per PSF,
D is the effective diffusion coefficient,
and v is the advection velocity.

Flux parameters can be independently derived from the spatial and temporal ICS sequences through fitting of the spatial autocorrelation maps for each frame as:

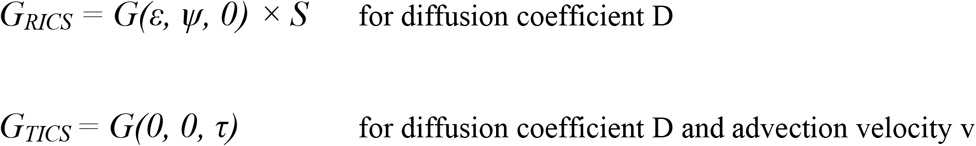

To prevent bias from the strong correlation at G (0,0) due to shot noise, the *ε*, *φ* = 0 points were excluded from fitting.

Python code for the functions computing the above parameters and extracting relevant metadata from the raw image files is provided as a Python scripts **RICSAnalysis.py and TICSAnalysis.py** (see Appendix).

#### Generation of an instrument calibration curve

In order to generate instrument calibration curves, we collected bile from mice and dissolved varying concentrations of CLF (0-700 μM) in mouse bile. These solutions were then imaged under our intravital imaging conditions in 8-well glass-bottom dishes (ibidi) at multiple locations per well. Mean intensities of the images acquired per concentration of CLF were then utilized to plot the calibration curve.

#### In silico modeling

##### 1. Digitization of liver tissue geometry

Mouse liver tissue slices of approximately 70 μm thickness were fixed and immunostained with Alexa 488-labeled anti-DPP4 and Alexa 594-labeled anti-KRT19 antibodies as described previously (14). DPP4 is a marker for apical membranes and allows visualization of canaliculi. KRT19 is a cholangiocyte specific marker that allows visualization of interlobular bile ducts. Confocal Z-stacks were obtained using imaging parameters as described earlier and Z-stacks were segmented to obtain specific canalicular and duct signals (ImageJ). The volume of the reconstructed isosurface of the reconstructed canalicular network and ducts was then discretized in a polyhedral mesh with 1.75 million volume elements (“cfMesh”, Creative-Fields, UK). The outer surface was divided into three sub-domains: the inlet and outlet of the bile duct and the wall of the bile duct and bile canaliculi.

##### 2. Molecular transport simulation in bile

###### Theoretical considerations

Molecular transport in bile was modelled by an advection-diffusion equation. For a species of concentration 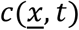 (*t*=time, 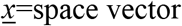) this equation reads:

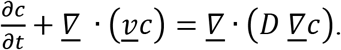

*D* is the diffusion coefficient and assumed to be constant. The local velocity vector field 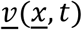 has been determined from the Navier-Stokes equation of an incompressible Newtonian fluid:

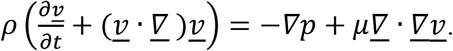

where 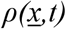 is the local fluid density, 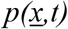 is the hydrostatic pressure, and *μ* is the dynamic viscosity. Incompressibility implies 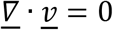 from the mass conservation, steady state implies 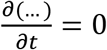.

###### Numerical implementation

The time-independent flow was computed by solving the stationary incompressible Navier-Stokes equations using the finite-volume methods implemented in the OpenFOAM toolbox (46). A large time-step transient solver was used to solve the pressure–velocity coupling, the PIMPLE (merged PISO-SIMPLE) algorithm. Spatial convergence was checked computing the Grid Convergence Index (GCI) (47) and the numerical convergence was ensured. The flow velocity was fed into a transient advection-diffusion equation for the concentration of labeled molecules. In order to minimize numerical diffusion and precisely capture the diffusion process, a *SuperBee* flux-limiter was used.

The diffusion simulations (for 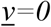) in continuum space were verified against particle-based simulations in one-, two-, and three dimensions in a stochastic compartmental approach on a regular lattice (solving a multivariate master equation) with various initial activation zones and analytical results in those settings that were amenable to analytical solution.

###### Simulation

Bile was considered as a Newtonian fluid of a *10^6^ μm^2^/s* viscosity and the molecular diffusion coefficient in bile was set to 3 μm^2^/s, the in vivo diffusion coefficient of bile salts determined experimentally. A fixed bile velocity was imposed normal to the inlet of the bile-duct, set as *1 μm/s*, the in-plane components being set to zero, and a fixed pressure was constrained at its exit. A non-slip condition 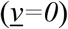 was chosen for the flow at the walls. For the concentration, a no-gradient boundary condition was set on all the boundaries.

The following situations of particular interest were then simulated:

- Repeated activation (Fig. 4B): At activation times *t*=*k t_a_*, (*n* see Fig. 4B), *c* was elevated by a constant value *c*_0_ in the activated zone.
- Activation experiments in Fig. 4C, D (BC=bile canaliculi): At *t*=*t_0_*=*0, c* was set to a constant value *c_0_* in the activated zone. The activated bile salt concentration value was without loss of generality fixed as initial condition at *c_0_=10^6^ M* in the chosen activation zone of the bile canaliculi and *0 M* elsewhere.

1. *R=5 μm* radius spheres in the BC: Decay of a sphere of radius *= 5 μm* at approximately *30 μm* Euclidian distance from the BD, *T_1/2_ = 5.5* s;
2. *R=10 μm* radius spheres in the BC: Decay of a sphere of radius *= 15 μm* in a midzonal canalicular region, *T_1/2_ =15.75* s;
3. *R=5 μm* spheres at the interface between canaliculi and duct (CoH) with *1 μm/s* inflow in the BD: Decay of a sphere of radius *= 5 μm* at the interface between canaliculi and duct (CoH), *T_1/2_ = 2.28 sec*. The speed of advection in the duct is *1 μm/sec* and virtually 0 in the BC, hence in-between in the CoH. The half-life times in cases (1.) and (2.) are between the value obtained for pure diffusion for a 1d domain and a 2d domain in a homogeneous isotropic medium: In a 1-dimensional homogeneous, practically infinite domain with initial condition *c(x,t=0)=c_0_ >0* in *-R ≤ x ≤ R (R:* radius), *c_0_ =0 otherwise*, the half-life time has been numerically determined as: 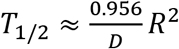, i.e. *T*_1/2_ ≈ *7.96 sec* for R=5*μm* and *T_1/2_ ≈ 31.87 sec* for R=10*μm*. In a 2-dimensional homogeneous, isotropic, practically infinite domain with initial condition *c(r, t=0)=c_0_ >0* in *-R ≤ r ≤ R (r* being the radius around the origin at *(x,y)=(0,0)* in polar coordinates), *c_0_ =0 otherwise*, the half-life time has been numerically determined as 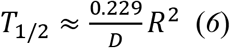, i.e. *T*_1/2_ ≈ *1.91 sec* for R=5*μm* and *T_1/2_ ≈ 7.63 sec* for R=10*μm*. Simulations in which the diffusing species reaches the impermeable domain border before the concentration in the activated domain has dropped to half of its initial value (i.e., *c_0_/2)*, were not utilized. Moreover, we found that the mean-square displacement on a 2-dimensional square network is between the mean-square displacement of a 1 dimensional homogeneous domain, and a 2 dimensional homogeneous, isotropic domain, consistent with the findings for cases (1.) and (2.) above.
- Activation in the bile duct (Fig. 4C): The BD in the domain was elongated towards the same length as in the experiment. The initial value 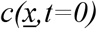 was extrapolated from the experimental results: 1) a Gaussian function was fitted on the experimental curve *c(x,t=0)* (Fig 1C, bottom) and 2) this curve was imposed in the bile along the main axis of an equivalent cylinder (*x* being the coordinate along this axis; the concentration is constant within each cross-section to this axis). Data was sampled along the centerline of arclength s of the duct for post-processing. For the activation in the bile canaliculi (Fig. 4C) the initial concentration profile was determined as for the BD and the simulation executed as explained for Fig. 4D.

#### General synthesis route for fluorescent cholic acid derivatives

NMR data for compound struture verification are provided (Table S5).

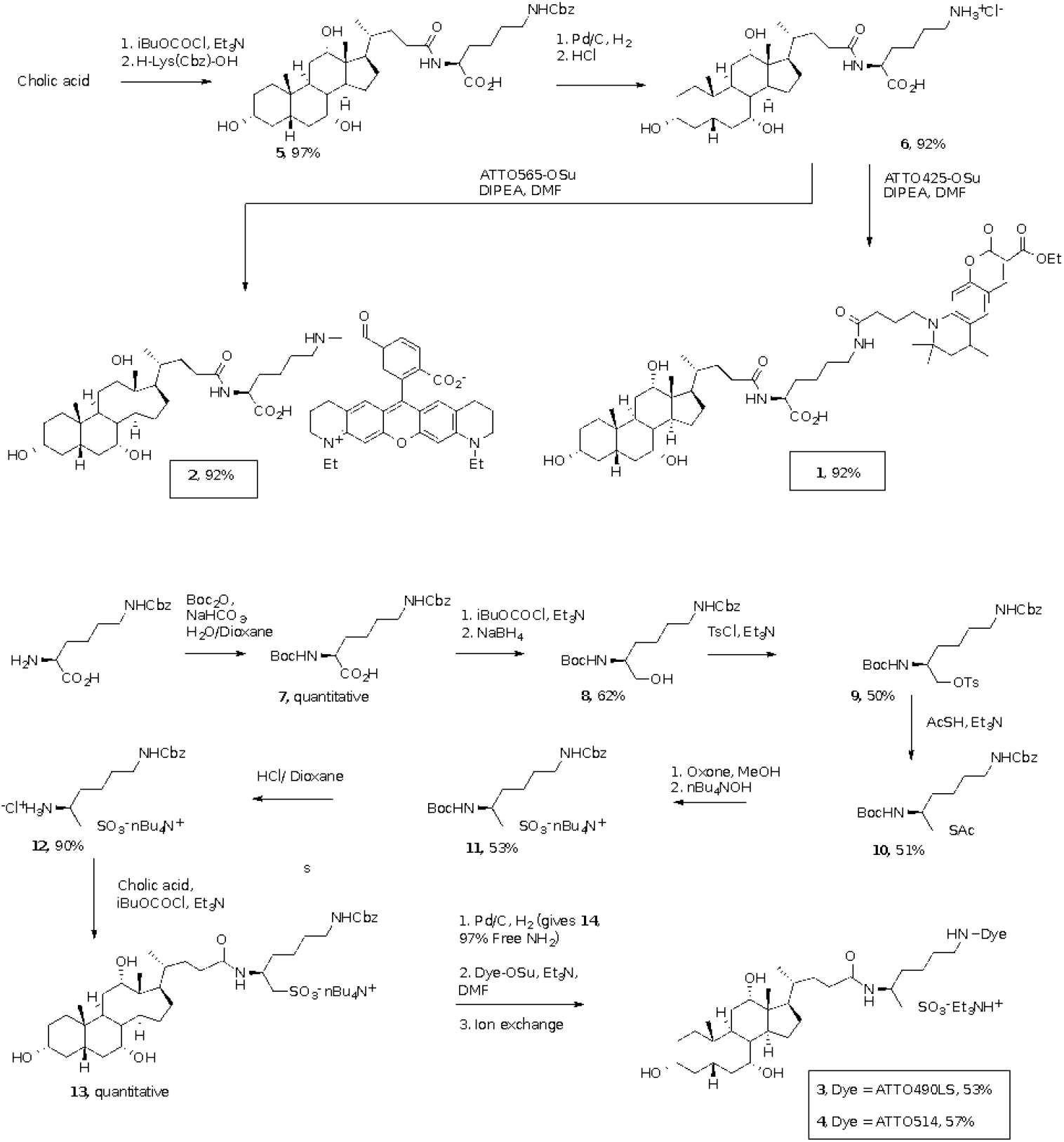

##### Cholic-Lys(Z)-OH (5)

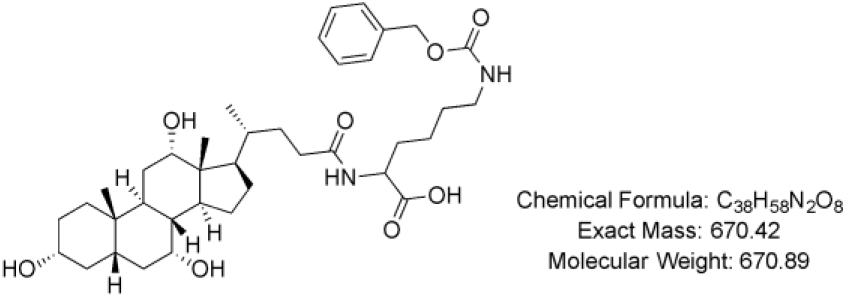

In a round bottom flask, cholic acid (2.0 g, 4.89 mmol) was dissolved in acetone (20 mL), triethylamine (1.4 equiv., 1.0 mL, 6.85 mmol) was added and the mixture was stirred for 20 min. After cooling to 0°C with an ice bath, ethyl chloroformate (0.46 mL, 4.89 mmol) was added dropwise and the mixture was allowed to stir for 90 min. In parallel, another round bottom flask was charged with a fresh solution of aqueous NaOH (0.5 M), H-Lys(Z)-OH (1.37 g, 4.89 mmol) was added and the reaction stirred for 60 min. The resulting mixture was transferred to the first flask and stirred overnight at room temperature. The pH of the mixture was adjusted to 2-2.5 with a 1M HCl solution and the compound was extracted with EtOAc. The organic layer was washed with water, dried over Na_2_SO_4_, filtered and concentrated to give compound **5** (3.20 g, 4.77 mmol) as a white solid, in 97% yield.

Rf: 0.28 CH_2_Cl_2_/MeOH (8:2)

^1^H NMR (600 MHz, MeOD) δ 7.34 (d, *J* = 4.4 Hz, 4H), 7.29 (q, *J* = 4.3 Hz, 1H), 5.06 (s, 2H), 4.35 (dd, *J* = 9.2, 4.8 Hz, 1H), 3.95 (s, 1H), 3.79 (d, *J* = 3.2 Hz, 1H), 3.37 (t, *J* = 4.5 Hz, 1H), 3.12 (t, *J* = 6.9 Hz, 2H), 2.35 – 2.15 (m, 4H), 2.03 – 1.92 (m, 2H), 1.92 – 1.78 (m, 5H), 1.77 – 1.63 (m, 3H), 1.63 – 1.50 (m, 7H), 1.47 – 1.26 (m, 7H), 1.10 (qd, *J* = 12.0, 5.6 Hz, 1H), 1.03 (d, *J* = 6.6 Hz, 3H), 0.98 (td, *J* = 15.6, 14.2, 4.6 Hz, 1H), 0.91 (s, 3H), 0.71 (s, 3H).

^13^C NMR (151 MHz, MeOD) δ 176.9, 175.6, 158.9, 138.4, 129.4, 128.9, 128.7, 79.5, 74.0, 72.9, 69.0, 67.3, 53.5, 48.1, 47.5, 43.2, 43.0, 41.5, 41.0, 40.5, 36.8, 36.5, 35.9, 33.8, 33.2, 32.2, 31.2, 30.4, 29.6, 28.7, 27.9, 24.2, 24.2, 23.2, 17.8, 13.0.

HR-MS (ESI+, m/z) calcd for C_38_H_58_N_2_O_8_ [(M + Na)^+^] 693.4085, found 693.4069.

##### Cholic-Lys(NH_3_^+^Cl^-^)-OH (6)

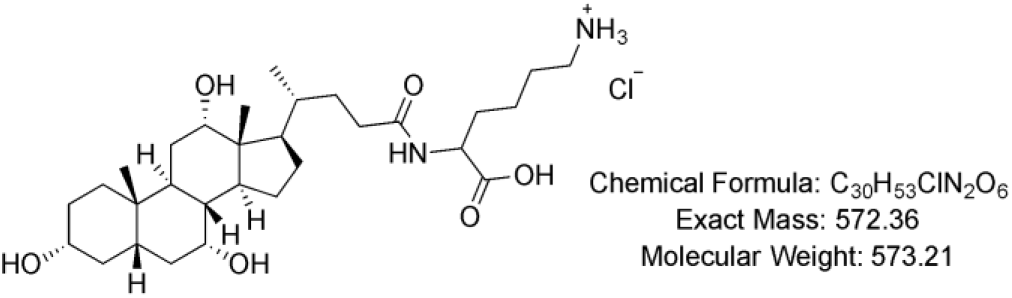

Cholic-Lys(Z) OH (**5**, 1.5 g, 2.24 mmol) was dissolved in EtOH (50 mL) and 10% of Pd/C was added. The solution was submitted to a hydrogen atmosphere overnight. After filtration through celite, the filtrate was evaporated to dryness. The product was then dissolved in water and the pH was adjusted to 2-2.5 with 1M HCl solution. The mixture was extracted with EtOAc and the aqueous layer was freeze-dried to obtain compound **6** as a white powder in quantitative yield.

^1^H NMR (400 MHz, MeOD) δ 4.40 (dd, *J* = 9.2, 5.0 Hz, 1H), 3.98 (t, *J* = 4.0 Hz, 1H), 3.82 (q, *J* = 3.0 Hz, 1H), 3.46 – 3.38 (m, 1H), 2.96 (t, *J* = 7.7 Hz, 2H), 2.41 – 2.15 (m, 4H), 2.08 – 1.94 (m, 2H), 1.93 – 1.67 (m, 9H), 1.65 – 1.26 (m, 14H), 1.06 (d, *J* = 6.4 Hz, 3H), 1.03 – 0.97 (m, 1H), 0.94 (s, 3H), 0.74 (s, 3H).

^13^C NMR (100 MHz, MeOD) δ 177.1, 174.1, 74.0, 72.8, 69.0, 53.5, 48.0, 47.5, 43.2, 43.0, 41.0, 40.5, 40.4, 36.9, 36.5, 35.9, 33.6, 33.1, 31.9, 31.2, 29.6, 28.7, 28.0, 27.9, 24.2, 23.9, 23.1, 17.8, 13.0.

HR-MS (ESI+, m/z) calcd for C_30_H_52_N_2_O_3_ [(M + H)^+^] 537.3898, found 537.3881.

##### Cholic-Lys(ATTO425)-OH (1)

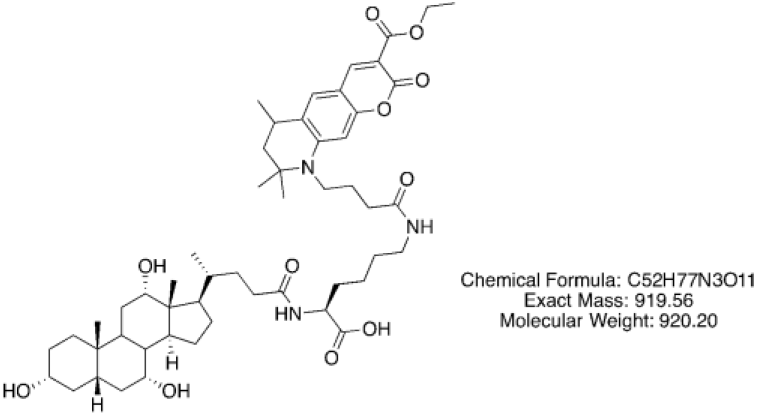

The amino compound MR15 (5.90 mg, 0.01 mmol) was dissolved in anhydrous DMF (0.4 mL), DIPEA (2.0 equiv., 3.70 x10^−3^ mL, 0.02 mmol) was added to adjust the pH around 8-9 followed by the addition of the succinimide ester of the fluorescent dye ATTO425 (1.0 equiv., 5.00 mg, 0.01 mmol). The mixture was stirred overnight in the dark and under argon atmosphere. The reaction was followed by LCMS, 0.5 equivalent of compound MR15 and DIPEA were added until complete consumption of the fluorescent dye. The reaction was complete after addition of 1.7 equivalent. The mixture was concentrated and purified by silica column chromatography eluted with 30% of EtOAc/MeOH/AcOH/H_2_O (3:3:3:2) in EtOAc. Compound **1** (8.50 mg, 9.24 x10^−3^ mmol) was obtained as a yellow powder in 92% yield.

^1^H NMR (600 MHz, MeOD) δ 8.57 (s, 1H), 7.42 (s, 1H), 6.49 (s, 1H), 4.32 (q, *J* = 7.1 Hz, 2H), 4.29 (d, *J* = 6.8 Hz, 1H), 3.93 (t, *J* = 6.0 Hz, 1H), 3.76 (t, *J* = 2.9 Hz, 1H), 3.59 – 3.51 (m, 1H), 3.39 – 3.33 (m, 1H), 3.22 (t, *J* = 6.8 Hz, 2H), 2.89 (td, *J* = 9.3, 6.5, 3.6 Hz, 1H), 2.30 (m, 3H), 2.27 – 2.23 (m, 1H), 2.15 (td, *J* = 13.7, 11.3, 6.6 Hz, 1H), 2.00 – 1.82 (m, 9H), 1.82 – 1.76 (m, 2H), 1.75 – 1.62 (m, 3H), 1.61 – 1.47 (m, 9H), 1.43 (s, 3H), 1.38 (q, *J* = 6.8 Hz, 10H), 1.31 (s, 7H), 1.06 (dq, *J* = 12.2, 6.6, 6.2 Hz, 1H), 1.01 (d, *J* = 6.4 Hz, 3H), 0.89 (s, 3H), 0.66 (d, *J* = 1.8 Hz, 3H).

^13^C NMR (151 MHz, MeOD) δ 176.1, 175.0, 165.7, 158.5, 153.2, 151.0, 128.7, 128.0, 109.1, 108.1, 97.8, 74.0, 72.9, 69.0, 62.1, 57.7, 55.5, 48.1, 47.5, 46.8, 46.0, 43.2, 43.0, 41.0, 40.5, 40.4, 36.9, 36.5, 35.9, 35.9, 34.4, 33.9, 33.5, 33.3, 31.2, 30.1, 29.7, 29.6, 28.7, 28.0, 27.9, 26.1, 25.3, 24.3, 24.2, 23.2, 19.9, 17.8, 14.6, 13.1.

HR-MS (ESI+, m/z) calcd for C_52_H_77_N_3_O_11_ [(M + Na)^+^] 942.5450, found 942.5435.

##### Cholic-Lys(ATTO565)-OH (2)

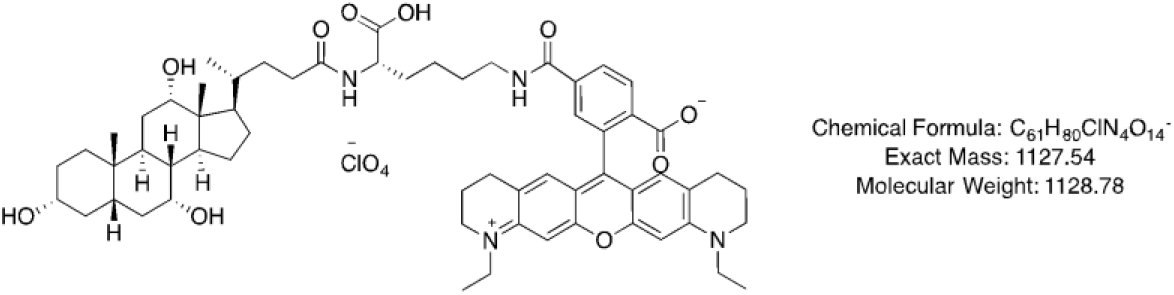

The amino compound **6** (1.8 equiv., 6.80 mg, 0.01 mmol) was dissolved in anhydrous DMF (0.4 mL), DIPEA (4 equiv., 3.00 x10^−3^ mL, 0.03 mmol) was added to adjust the pH around 8-9 followed by the addition of the succinimide ester of the fluorescent dye ATTO565 (1.0 equiv., 5.00 mg, 7.00 x10^−3^ mmol). The mixture was stirred overnight in the dark and under argon atmosphere. The reaction was followed by LCMS, 1 equivalent of compound **6** and DIPEA were added until complete consumption of the fluorescent dye. The reaction was complete after addition of 3 equivalents. The mixture was concentrated and purified by silica column chromatography eluted with 30% of EtOAc/MeOH/AcOH/H_2_O (3:3:3:2) in EtOAc. Compound **2** (7.30 mg, 6.47 x10^−3^ mmol) was obtained as a pink powder in 92% yield.

^1^H NMR (600 MHz, MeOD) δ 8.14 (d, *J* = 8.1 Hz, 1H), 8.10 – 8.05 (d, *J* = 8.0 Hz 1H), 7.68 (d, *J* = 1.7 Hz, 1H), 6.89 (d, *J* = 6.3 Hz, 4H), 4.26 (bs, 1H), 3.92 (s, 1H), 3.77 (d, *J* = 3.3 Hz, 1H), 3.66 (dt, *J* = 17.0, 7.1 Hz, 4H), 3.57 (t, *J* = 5.9 Hz, 4H), 3.42 – 3.36 (m, 3H), 2.71 (d, *J* = 6.4 Hz, 4H), 2.34 – 2.21 (m, 3H), 1.94 (m, 7H), 1.88 – 1.83 (m, 2H), 1.79 (m, 2H), 1.71 (m, 2H), 1.68 – 1.62 (m, 2H), 1.60 (m, 2H), 1.54 – 1.48 (m, 2H), 1.43 (m, 4H), 1.33 (m, 12H), 1.12 – 1.03 (m, 1H), 1.00 (d, *J* = 6.3 Hz, 3H), 0.96 (dd, *J* = 14.4, 3.4 Hz, 1H), 0.90 (s, 3H), 0.68 (s, 3H).

^13^C NMR (151 MHz, MeOD) δ 176.1, 168.7, 160.1, 158.5, 154.7, 144.3, 136.5, 133.9, 131.1, 129.6, 129.2, 126.4, 126.4, 115.0, 95.7, 79.5, 74.0, 72.9, 69.0, 58.3, 55.8, 50.4, 49.8, 48.1, 47.8, 47.5, 43.2, 43.0, 41.1, 41.0, 40.5, 36.9, 36.5, 35.9, 35.9, 34.3, 33.5, 33.3, 31.2, 30.1, 29.6, 28.8, 28.5, 27.9, 24.4, 24.2, 23.2, 22.8, 22.1, 18.4, 17.8, 13.1, 11.3.

HR-MS (ESI+, m/z) calcd for C_61_H_81_N_4_O_10_ [(M + H)^+^] 1030.6020, found 1030.6017.

##### Taurin Lysine

##### Boc-Lys(Z)-OH (7)

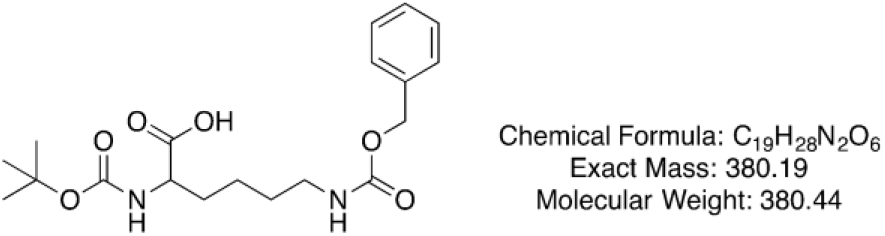

H-Lys(Z)-H (2.00 g, 7.13 mmol) was dissolved in 30 mL of H_2_O and NaHCO_3_ (3 equiv., 1.80 g, 21.40 mmol) was added. The mixture was cooled to 0°C before to add Boc_2_O (1.5 equiv., 2.30 g, 10.70 mmol) and cold 1,4-dioxane (90 mL). The reaction was stirred overnight and evaporated to dryness to give compound **7** in 99% yield (2.70 g, 7.10 mmol).

^1^H NMR (400 MHz, DMSO) δ 7.42 – 7.29 (m, 5H), 7.24 (t, *J* = 5.8 Hz, 1H), 7.02 (d, *J* = 8.0 Hz, 1H), 5.01 (s, 2H), 3.83 (td, *J* = 8.5, 7.9, 4.6 Hz, 1H), 2.98 (q, *J* = 6.5 Hz, 2H), 1.70 – 1.49 (m, 2H), 1.38 (s, 9H), 1.37 – 1.23 (m, 4H).

^13^C NMR (100 MHz, DMSO) δ 174.3, 156.1, 155.6, 137.3, 128.4, 127.7, 79.2, 78.0, 65.1, 53.5, 30.4, 29.0, 28.2, 22.9.

##### Boc-Lys(Z)-CH_2_OH (8)

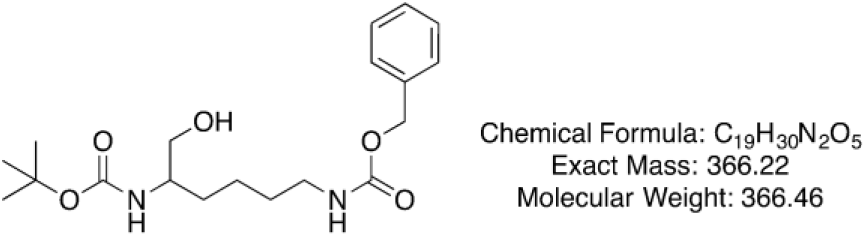

Boc-Lys(Z)-OH (**7**, 0.70 g, 1.80 mmol) and Et_3_N (1.1 equiv., 0.28 mL, 2.00 mmol) were dissolved in anhydrous DMF (10 mL), the mixture was cooled to −10°C and ethylchloroformate (1.1 equiv., 0.19 mL, 2.00 mmol) was added dropwise. The reaction stirred for 30 min under argon atmosphere, filtered and rinsed with THF (3x). The filtrate was cooled to 0°C, NaBH_4_ (3 equiv., 0.23 g, 6.00 mmol) was added as well as 70 mL of MeOH and the reaction was stirred for 1 h. The mixture was carefully quenched with HCl 2N and concentrated. After addition of H_2_O, the product was extracted with EtOAc and the organic layer washed with HCl 1N, H_2_O, sat. NaHCO_3_ and brine (2x), dried over Na_2_SO_4_, filtered and concentrated. Purification on silica column chromatography eluted with Petrol/EtOAc (2:8) gave compound **8** (0.88 g, 2.39 mmol) in 62% yield.

Rf: 0.37 EtOAc/PET (8:2)

^1^H NMR (400 MHz, MeOD) δ 7.34 (d, *J* = 4.3 Hz, 4H), 7.29 (dt, *J* = 9.1, 4.2 Hz, 1H), 5.06 (s, 2H), 3.55 – 3.38 (m, 3H), 3.12 (t, *J* = 6.8 Hz, 2H), 1.63 – 1.43 (m, 4H), 1.43 (s, 9H), 1.36 (m, 2H).

^13^C NMR (100 MHz, MeOD) δ 158.8, 158.2, 138.3, 129.4, 128.9, 128.7, 79.9, 79.4, 67.2, 65.3, 53.6, 41.6, 31.9, 30.7, 28.8, 24.2.

HR-MS (ESI+, m/z) calcd for C_19_H_30_N_2_O_5_ [(M + Na)^+^] 389.2047, found 349.2051.

##### Boc-Lys(Z)-CH2OTs (9)

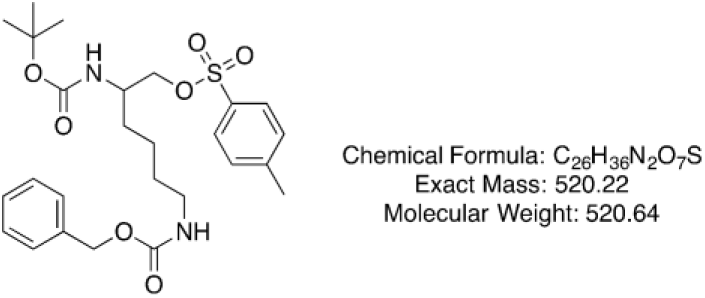

Boc-Lys(Z)-CH_2_OH (**8**, 1.80 g, 4.93 mmol) was dissolved in 10 mL of anhydrous CH_2_Cl_2_, cooled to 0°C and TsCl (1.5 equiv., 1.40 g, 7.40 mmol) was added followed by Et_3_N (2 equiv., 1.40 mL, 9.80 mmol). The reaction was stirred overnight, poured into H_2_O and extracted with CH_2_Cl_2_. The product was dried over Na_2_SO_4_, filtered, concentrated and purified by silica column chromatography eluted with CHCl_3_. Compound **9** (1.28 g, 2.40 mmol) was obtained in 50% yield.

^1^H NMR (400 MHz, CDCl_3_) δ 7.78 (d, *J* = 8.2 Hz, 2H), 7.43 – 7.27 (m, 7H), 5.09 (s, 2H), 4.76 (s, 1H), 4.62 (s, 1H), 4.10 – 3.92 (m, 2H), 3.72 (m, 0H), 3.15 (s, 2H), 2.45 (s, 3H), 1.39 (s, 15H).

^13^C NMR (100 MHz, MeOD) δ 158.9, 157.9, 146.5, 138.4, 134.2, 131.1, 129.4, 129.1, 128.9, 128.8, 80.2, 79.5, 72.7, 67.3, 52.5, 41.5, 31.3, 30.5, 28.7, 24.0, 21.6.

HR-MS (ESI+, m/z) calcd for C_26_H_36_N_2_O_7_S [(M + Na)^+^] 543.2135, found 543.2112.

##### Boc-Lys(Z)-CH_2_SCOCH_3_ (10)

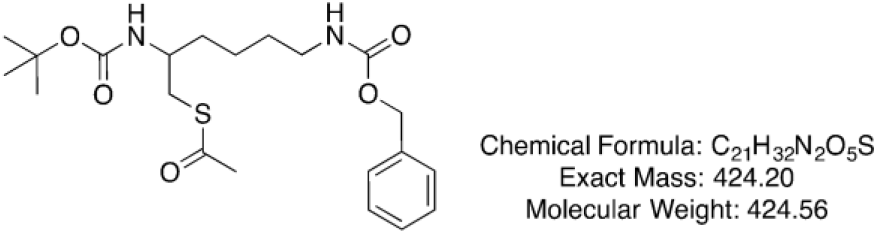

Compound **9** (0.23 g, 0.44 mmol) was dissolved in CHCl_3_ (5 mL) and DIPEA (5 equiv., 0.38 mL, 2.21 mmol) was added followed by thioacetic acid (5 equiv., 0.16 mL, 2.21 mmol) dissolved in 1 mL of CHCl_3_. The mixture was stirred overnight under argon atmosphere and followed by LCMS. 3 equivalents of thioacetic acid and DIPEA were added every day and the reaction was complete the third day. The mixture was washed with brine, extracted with CHCl_3_, dried over Na_2_SO_4_, filtered and concentrated. The residual solid was dissolved in CH_2_Cl_2_ and treated with active charcoal overnight. After filtration and purification on silica column chromatography eluted with CHCl_3_, compound **10** (0.10 g, 0.23 mmol) was obtained in 51% yield.

Rf: 0.60 CHCl_3_/MeOH (95:5)

^1^H NMR (400 MHz, Chloroform-*d*) δ 7.26 – 7.15 (m, 5H), 4.96 (s, 2H), 4.69 (s, 1H), 4.38 (s, 1H), 3.59 (s, 1H), 3.05 (q, *J* = 6.8 Hz, 2H), 2.98 – 2.78 (m, 2H), 2.21 (s, 3H), 1.47 – 1.31 (m, 4H), 1.28 (s, 9H), 1.26 – 1.17 (m, 2H).

^13^C NMR (100 MHz, CDCl_3_) δ 195.7, 156.5, 155.6, 136.6, 128.5, 128.1, 128.1, 79.4, 66.6, 50.3, 40.7, 33.9, 30.6, 29.5, 28.4, 23.0.

HR-MS (ESI+, m/z) calcd for C_21_H_32_N_2_O_5_S [(M + Na)^+^] 447.1924, found 447.1919.

##### Boc-Lys(Z)-CH_2_SO_3_^-^nBu_4_^+^ (11)

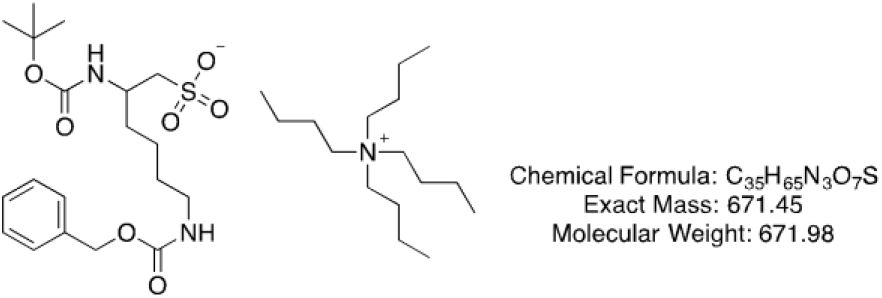

Compound **10** (0.10 g, 0.23 mmol) was dissolved in MeOH (5 mL) and a solution of oxone (3.5 equiv., 0.25 g, 0.80 mmol) in 5 mL of H_2_O was added. The mixture was stirred for 1.5 h, tetrabutylammonium hydroxide solution 40 wt. % in H_2_O (0.34 mL) in 2mL of H_2_O was added and the stirring was continued for 16 h. The methanol was concentrated under vacuum, the compound diluted with H_2_O and extracted with CH_2_Cl_2_, dried over Na_2_SO_4_, filtered and concentrated. Purification by silica column chromatography and elution with CHCl_3_ gave compound **11** (0.08 g, 0.12 mmol) in 53% yield.

Rf: 0.24 CHCl_3_/MeOH (9:1)

^1^H NMR (600 MHz, Chloroform-*d*) δ 7.36 – 7.32 (m, 4H), 7.32 – 7.27 (m, 1H), 5.14 – 4.99 (m, 2H), 3.91 – 3.83 (m, 1H), 3.28 – 3.25 (m, 8H), 3.24 – 3.11 (m, 2H), 3.04 (ddd, *J* = 58.4, 14.2, 4.5 Hz, 2H), 1.65 – 1.61 (m, 8H), 1.58 – 1.47 (m, 2H), 1.43 (q, *J* = 7.3 Hz, 10H), 1.39 (s, 11H), 1.00 (t, *J* = 7.3 Hz, 6H).

^13^C NMR (151 MHz, CDCl_3_) δ 156.7, 156.0, 137.2, 128.5, 128.1, 128.0, 78.5, 66.4, 59.0, 53.6, 48.1, 40.4, 32.8, 29.1, 28.6, 24.2, 23.0, 19.9, 13.8.

HR-MS (ESI+, m/z) calcd for C_19_H_30_N_2_O_7_S [(M + H)^+^] 431.1846, found 431.1839.

##### Cl^-^NH_3_^+^-Lys(Z)-CH_2_SO_3_^-^nBu_4_+ (12)

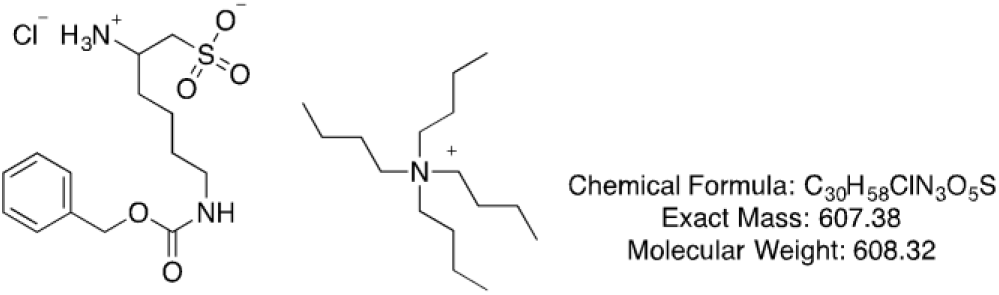

At 0°C, compound **11** was dissolved in 4M HCl/dioxane solution and the mixture was stirred for 2 h at rt. Evaporation to dryness gave the chloride salt **12** (0.06 mg, 0.11 mmol) in 90% yield.

^1^H NMR (400 MHz, Deuterium Oxide) δ 6.99 (m, 5H), 4.67 (s, 2H), 3.25 – 3.19 (m, 1H), 2.79 – 2.72 (m, 8H), 2.70 (s, 4H), 1.32 (q, *J* = 7.6 Hz, 2H), 1.20 (p, *J* = 7.8 Hz, 8H), 1.08 (q, *J* = 7.0 Hz, 2H), 0.91 (q, *J* = 7.4 Hz, 10H), 0.50 (t, *J* = 7.4 Hz, 12H).

^13^C NMR (151 MHz, MeOD) δ 159.0, 138.4, 129.5, 129.0, 128.8, 67.4, 59.5, 59.5, 59.5, 52.8, 50.3, 41.2, 33.3, 30.4, 24.8, 23.2, 20.7, 20.7, 20.7, 13.9.

HR-MS (ESI+, m/z) calcd for C_14_H_22_N_2_O_5_S [(M + H)^+^] 331.1322, found 331.1303.

##### Chol-Lys(Z)-CH_2_SO_3_^-^nBu_4_^+^ (13)

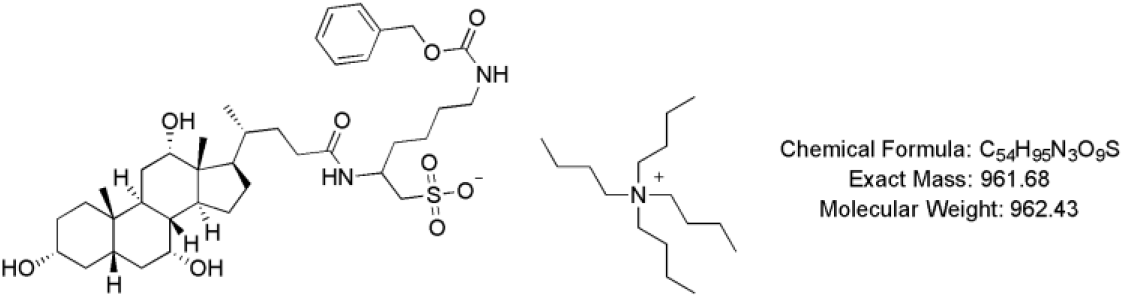

In a round bottom flask, cholic acid (21 mg, 0.05 mmol) was dissolved in acetone (0.6 mL), Et_3_N (1.4 equiv., 0.01 mL, 0.07 mmol) was added and the mixture was stirred for 10 min. After cooling to 0°C with an ice bath, ethyl chloroformate (5.0×10^−3^ mL, 0.05 mmol) was added dropwise and the mixture was allowed to stir for 90 min. In parallel, another round bottom flask was charged with a fresh solution of 0.5 M NaOH (1 mL), compound **12** (0.03 g, 0.05 mmol) was added and the reaction stirred for 60 min. The resulting mixture was transferred to flask (1) and stirred overnight at rt. After concentration to dryness, compound **13** (0.05 g, 0.05 mmol) occurred as a white solid in quantitative yield.

^1^H NMR (600 MHz, Methanol-*d*_4_) δ 7.41 – 7.25 (m, 5H), 5.06 (s, 2H), 4.26 (dq, *J* = 11.0, 5.6 Hz, 1H), 3.95 (dt, *J* = 6.3, 3.0 Hz, 1H), 3.79 (p, *J* = 3.0 Hz, 1H), 3.38 (m, 1H), 3.26 – 3.23 (m, 8H), 3.11 (t, *J* = 6.9 Hz, 2H), 3.03 – 2.90 (m, 3H), 2.33 – 2.21 (m, 4H), 2.10 (m, 1H), 2.01 – 1.93 (m, 2H), 1.90 – 1.85 (m, 2H), 1.83 – 1.79 (m, 2H), 1.74 (dd, *J* = 6.4, 3.7 Hz, 1H), 1.66 (m, 9H), 1.59 – 1.52 (m, 8H), 1.42 (m, 10H), 1.35 – 1.31 (m, 2H), 1.24 – 1.22 (m, 2H), 1.10 (m, 1H), 1.04 – 1.01 (m, 15H), 0.98 (m, 1H), 0.91 (s, 3H), 0.70 (s, 3H).

^13^C NMR (151 MHz, MeOD) δ 183.5, 176.0, 158.9, 138.5, 129.4, 128.9, 128.7, 74.0, 72.9, 69.0, 67.2, 59.5, 56.0, 48.1, 47.7, 47.5, 43.2, 43.0, 41.6, 41.0, 40.5, 36.9, 36.5, 35.9, 34.4, 33.3, 31.2, 30.5, 29.6, 28.7, 27.9, 24.8, 24.2, 24.1, 23.2, 20.7, 17.8, 14.0, 13.0.

HR-MS (ESI+, m/z) calcd for C_38_H_60_N_2_O_9_S [(M + Na)^+^] 743.3912, found 743.3913.

##### Chol-Lys(NH_2_)-CH_2_SO_3_^-^nBu_4_^+^ (14)

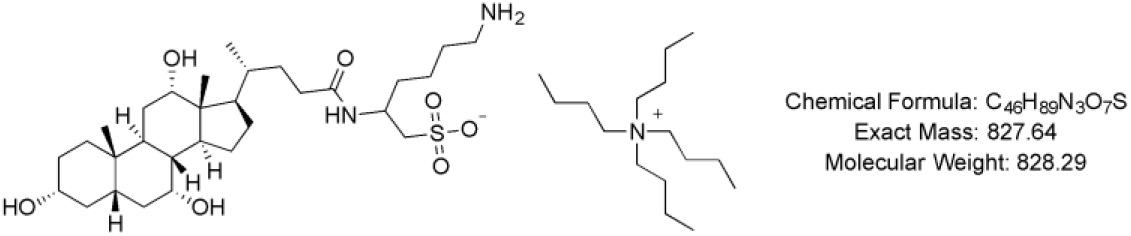

Compound **13** (0.05 g, 0.05 mmol) was dissolved in EtOH/THF (1:1, 5 ml), 10% of Pd/C (0.05 g) was added and the mixture submitted to a hydrogen atmosphere overnight. After filtration through celite and rinsing with EtOH, the compound was freeze-dried and **14** was obtained as a yellowish solid in 97% yield.

^1^H NMR (600 MHz, Methanol-*d*_4_) δ 4.29 (s, 1H), 3.95 (s, 1H), 3.79 (s, 1H), 3.41 – 3.34 (m, 1H), 3.26 – 3.21 (m, 5H), 3.08 – 2.99 (m, 2H), 2.93 (dd, *J* = 14.0, 7.1 Hz, 2H), 2.26 (m, 3H), 2.02 – 1.94 (m, 3H), 1.88 (d, *J* = 13.3 Hz, 3H), 1.83 – 1.78 (m, 2H), 1.69 – 1.64 (m, 7H), 1.57 (m, 5H), 1.44 – 1.39 (m, 9H), 1.33 – 1.23 (m, 4H), 1.14 – 1.08 (m, 1H), 1.04 – 1.00 (m, 12H), 0.99 – 0.96 (m, 1H), 0.92 (s, 3H), 0.71 (s, 3H).

^13^C NMR (151 MHz, MeOD) δ 176.0, 74.0, 72.9, 69.0, 59.5, 55.6, 52.0, 44.0, 43.2, 43.0, 41.0, 40.5, 36.9, 36.7, 36.5, 35.9, 34.4, 32.2, 31.9, 31.2, 29.6, 28.7, 27.9, 24.8, 24.2, 23.5, 23.2, 20.7, 17.8, 13.9, 13.0.

##### Chol-Lys(ATTO490LS)-CH_2_SO_3_^-^nBu_4_^+^ (3)

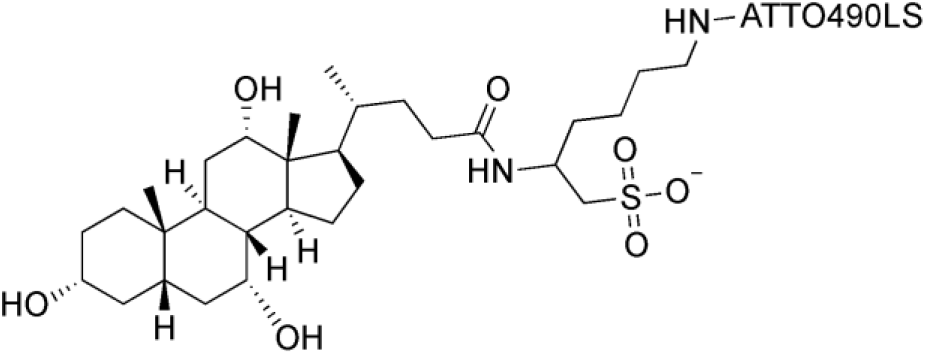

MW(nBu_4_ salt) = 1484 g.mol^−1^

MW(Et_3_N salt) = 1344 g.mol^−1^

M= 1242

Compound **14** (10.40 mg, 12.60 x10^−3^ mmol, 2 equiv.) was dissolved in 0.5 mL of dry DMF and DIPEA (2.20 μL, 12.60 x10^−3^ mmol, 2 equiv.) was added to the solution, the pH reached

## Supporting Figure Legends

**Figure S1.**
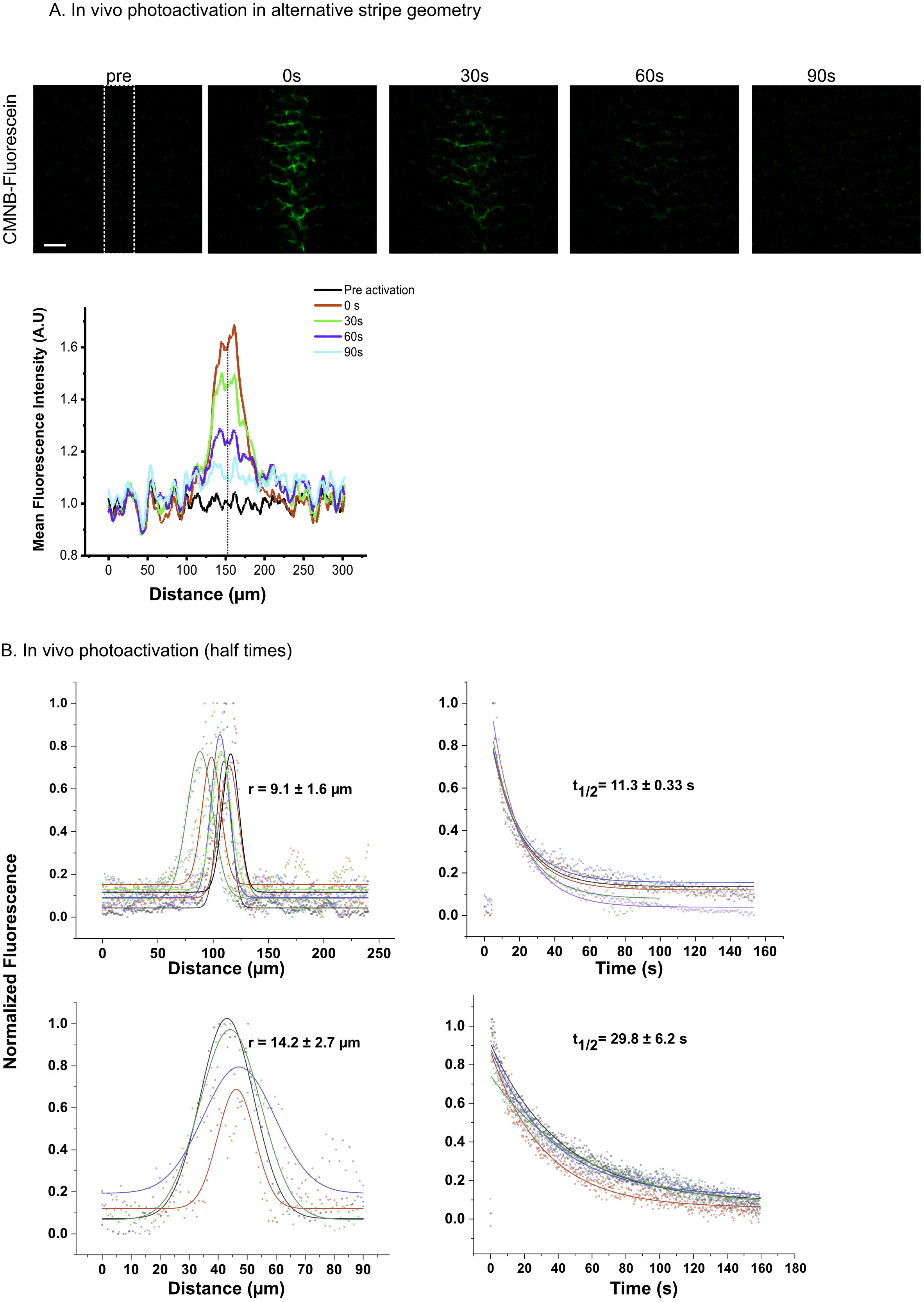
A. Alternative geometry of activation does not influence spatial decay symmetry. In vivo time-lapse imaging of the decay of CMNB-Fluo from the canalicular network, following activation (white dotted line) in a stripe geometry. Graph shows the symmetric decay with no shift of the center of mass of the intensity (black line). Scale bar: 30 μm. **B. Area of photoactivation influences half-life of decay.** Graphs show the circular spatial activation profiles CMNB-Fluo of varying radius, with correspondingly varying half-life of temporal decay as predicted by the diffusion equation t_1/2_ = r^2^ / 4D. The individual colors indicate independent photoactivations using the same conditions.

**Figure S2.**
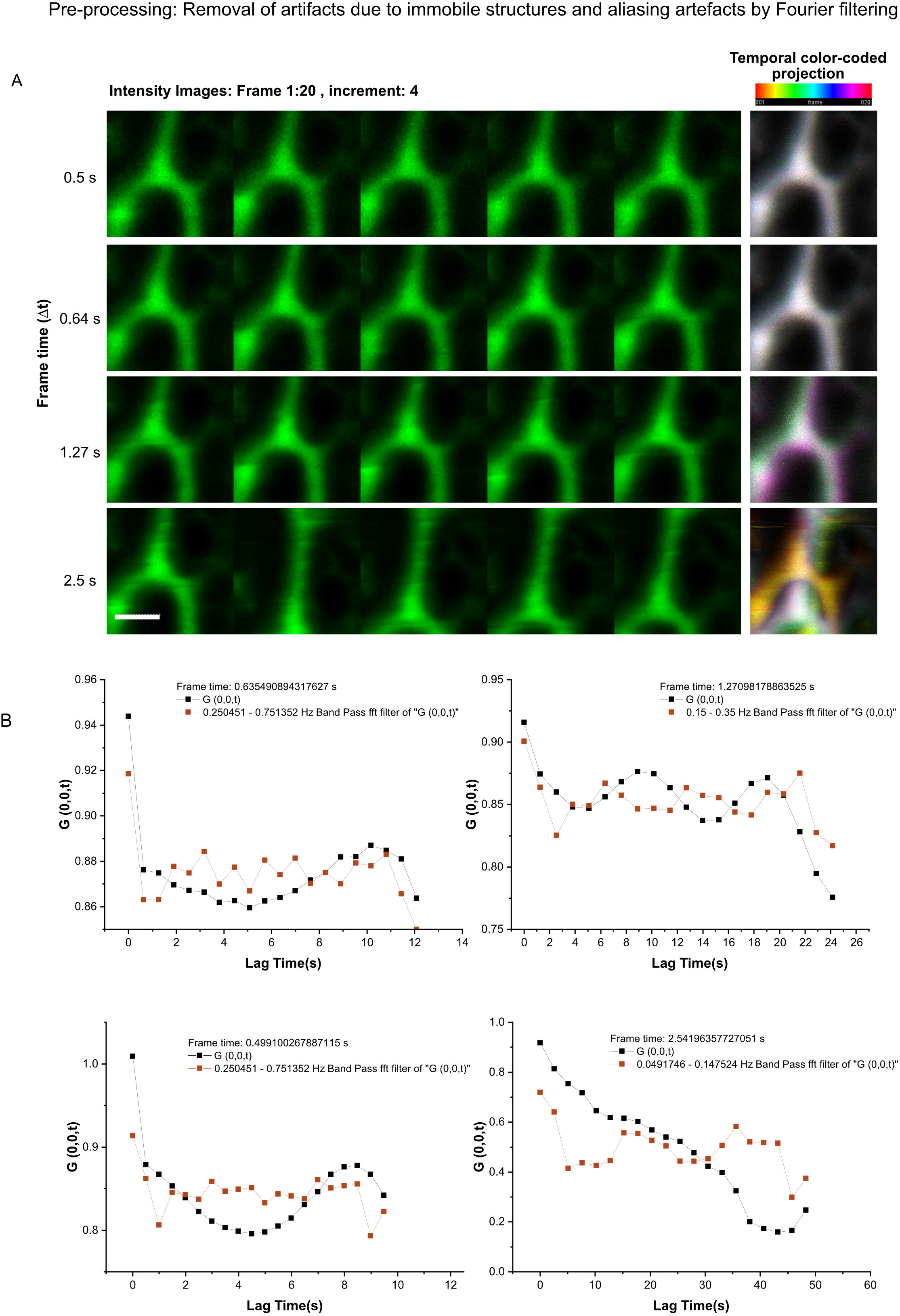
**A.** Representative ICS imaging sequences taken of an IBD taken at various frame times showing animal movements in raw intensity. The temporal color-coded project illustrates the deviations introduced. Scale bar: 10 μm. **B.** The graphs show the periodic variation of the temporal autocorrelation due to animal movements (black) and corresponding band-pass Fourier-filtered autocorrelation curves to remove these variations (red).

**Figure S3.**
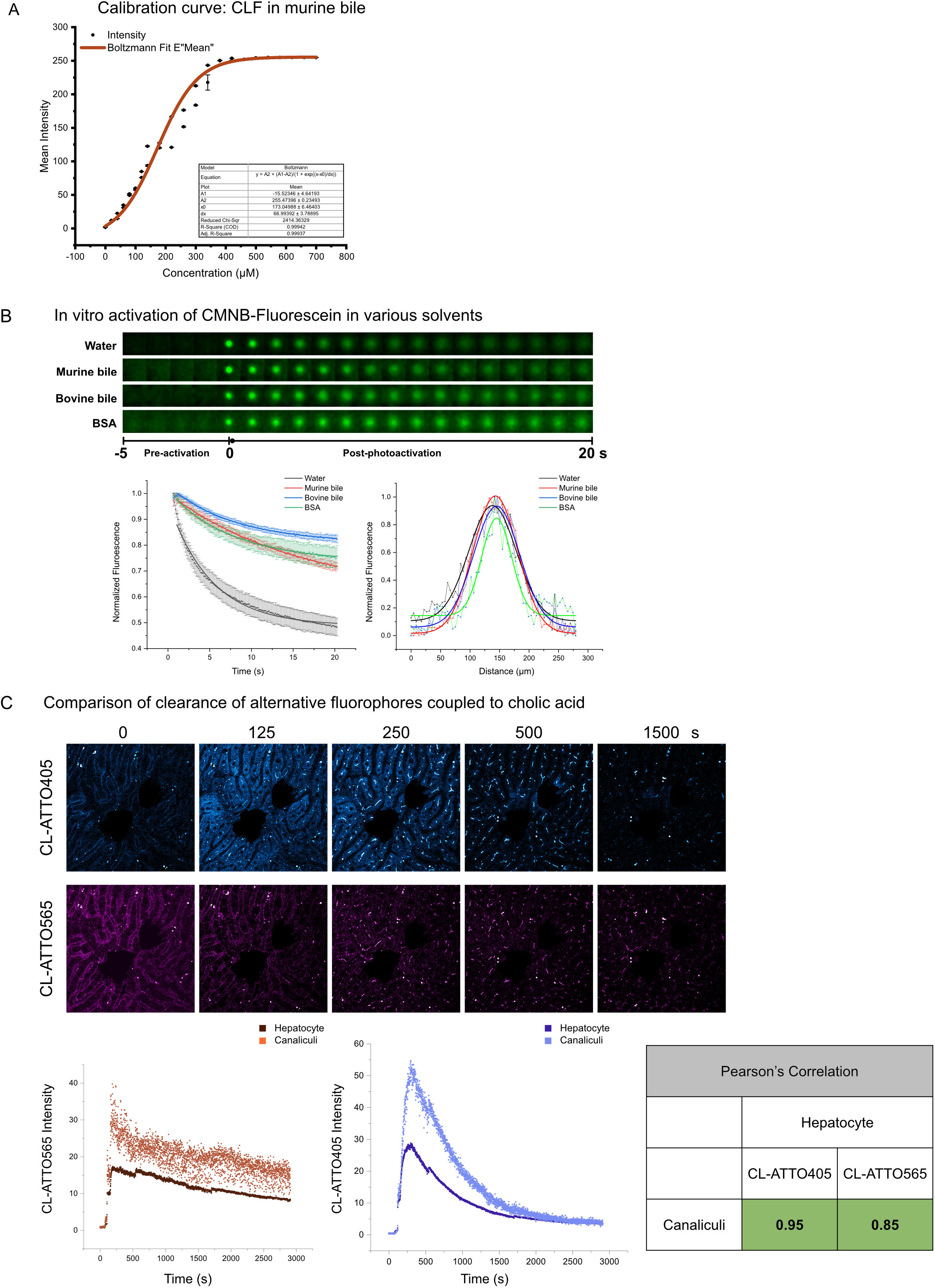
**A.** Calibration curve of CLF in mouse bile, indicating the non-linearity in lower concentration ranges, near-linearity in most of the concentration range up to 400 μM of CLF and detector saturation beyond 400 μM. The Boltzmann fit provides a convenient function for interconversion of concentrations and intensity values for CLF in bile. **B.** Kymographs and intensity quantification following Photoactivation of CMNB-Fluo in vitro in water, mouse bile, bovine bile, or 1% albumin solution showing that diffusion of fluorescein can be significantly reduced by binding to hydrophobic components (proteins, micelles, etc.) known to be present in bile. **C.** Time-lapse imaging (top) and temporal clearance of a poorly-exported bile salt analog CL-ATTO565(red) and a well-exported analog CL-ATTO405 (blue), showing a strong correlation between hepatocyte intensity and canalicular intensity over the time required for clearance. This indicates that hepatocyte export kinetics to the canaliculi is a major determinant of canalicular clearance, and that the canalicular excretion rate is not rate-limiting for overall clearance of the bile salt analogues. The table provides the Pearson correlation coefficients between hepatocyte intensity and canalicular intensity over time.

**Figure S4:**
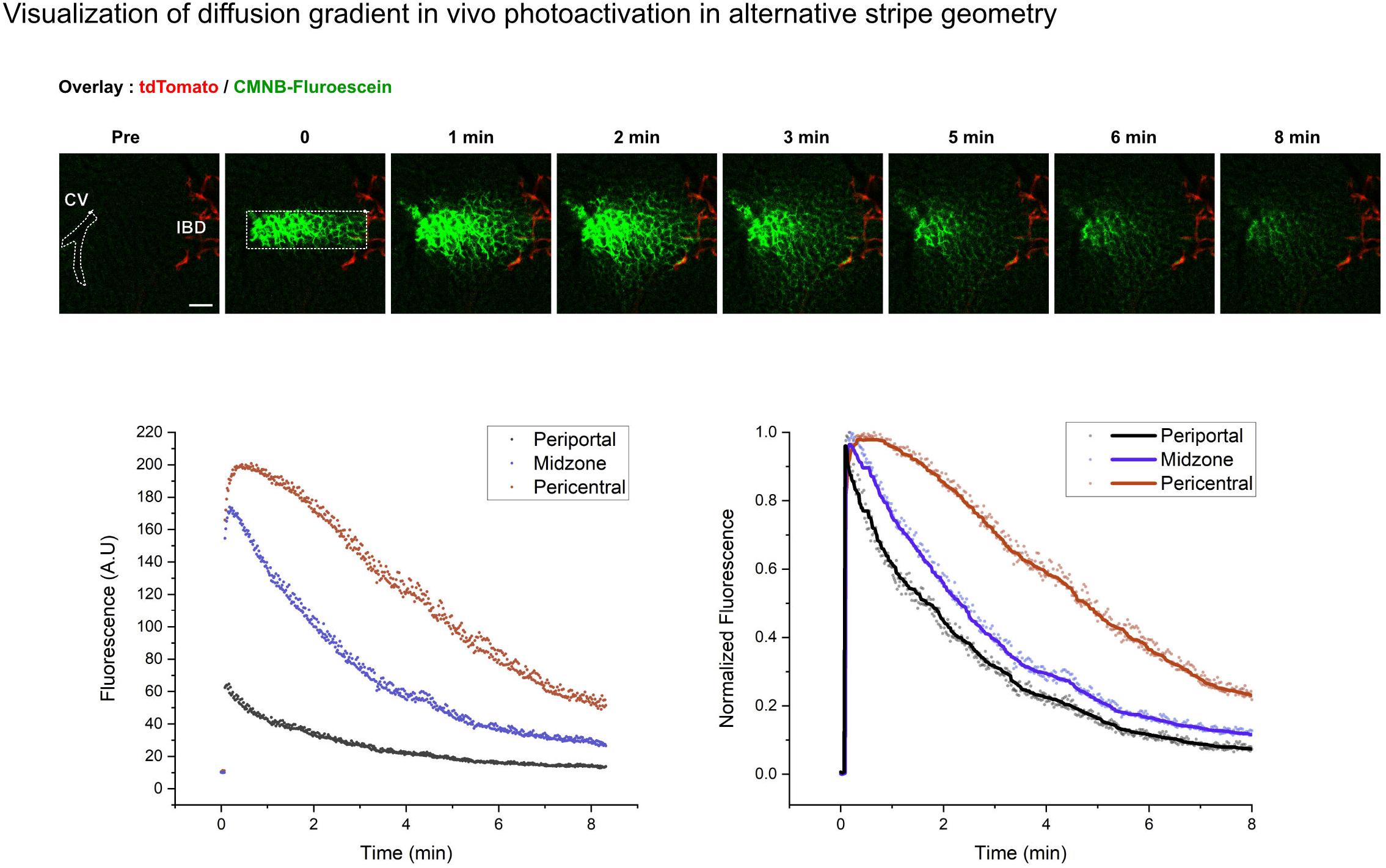
Representative time-lapse imaging of a liver lobule with tdTomato-positive interlobular ducts (red), infused with CMNB-Fluo after photoactivation a rectangular stripe bridging the pericentral (proximal to CV) and periportal zones (proximal to IBD), showing the diffusive CV-PV gradient. Graphs show raw and normalized fluorescence intensity indicating higher depletion rate in the periportal versus pericentral zones.

**Table S1: Results and data for velocity and diffusion in liver microcompartments obtained from fluorescence loss after photoactivation**

Diffusion coefficients and velocities calculated using the Soumpasis method (see Supporting Information - Photo-activation) for single photoactivations performed in CV, PV, MZ and IBD zones of the liver, under treatment conditions corresponding to basal (control), secretin bolus or TCA-infusion. Each row represents an independent photoactivation sequence. Zero values in velocity columns indicate that no shift could be detected in the centroid of the photoactivated region over time. Cases where a half-life could not be determined as marked ‘n.d’ and were not used. These data are graphically represented in Fig. 2C.

**Table S2: Results and data for diffusion in liver microcompartments obtained from spatial ICS.**

Diffusion coefficients (D) derived from fitting of spatial autocorrelation decays in the y-direction (RICS-y) in CV, PV, MZ and IBD zones of the liver, under treatment conditions corresponding to basal (control), secretin bolus or TCA-infusion. The ‘Region’ column classifies the liver domains into canalicular or ductular zones. The ‘w0’ and ‘wz’ columns correspond to the beam waist and axial beam height used as fit parameters (see Supporting Information - IVARICS). The ‘COD: R^2’ column provides the coefficient of determination of the non-linear global fit to a 1-population 3D-diffusion function, and the ‘Bound encountered’ column records if the bound was encountered during fitting iterations. Raw data files are referenced in the filename column and corresponding autocorrelation curves and fits are available (see Appendix). ICS data with fitting COD < 0.8 or in which the fitting bound was encountered (marked red) were not used for further analysis. These data are graphically represented in Fig. 3E.

**Table S3**: **Results and data for diffusion and velocity in liver microcompartments obtained from temporal ICS.**

Diffusion coefficients (D) and advection velocities (V) derived from fitting of temporal autocorrelation decays in CV, PV, MZ and IBD zones of the liver, under treatment conditions corresponding to basal (control), secretin bolus or TCA-infusion. The ‘Region’ column classifies the liver domains into canalicular or ductular zones. The ‘w0’ and ‘wz’ columns correspond to the beam waist and axial beam height used as fit parameters (see Supporting Information - IVARICS). The ‘COD : R^2’ column provides the coefficient of determination of the non-linear global fit for a 1-population diffusion and velocity function, and the ‘Bound encountered’ column records if the bound was encountered for either D or V during fitting iterations. Raw data files are referenced in the ‘File’ column and corresponding autocorrelation curves and fits are available (see Appendix). ICS data with fitting COD < 0.8 or in which the fitting bound was encountered (marked red) were not used for further analysis. These data are graphically represented in Fig. 3E.

**Table S4: NMR spectra of custom synthesized cholic acid derivatives**

NMR spectra of custom synthesized cholic acid derivatives (see Supporting Information - General synthesis route for fluorescent cholic acid derivatives) are provided, showing peaks relevant to confirm the chemical structure of the synthesized compound.

## Supporting Movies

**Movie S1:** Photoactivation of CMNB-Fluo-Dextran in blood vessel

**Movie S2:** Photoactivation of CMNB-Fluo in various domains of the liver biliary network.

**Movie S3:** Photoactivation of CMNB-Fluo in using stripe geometry in the liver canalicular network.

**Movie S4:** Photoactivation of CMNB-Fluo in various domains of the liver biliary network under basal, secretin- or TCA-stimulated conditions.

**Movie S5:** Aliasing effects in RICS sequences due to animal movements and their representation in the spatial autocorrelation.

**Movie S6:** Clearance of CLF from the liver canalicular network under basal, secretin- or TCA-stimulated conditions.

**Movie S7:** Clearance of fluorescein in its native form with and without the presence of cholic acid, or as CLF (fluorescein conjugated with cholic acid).

**Movie S8:** Visualization of lobule-wide antidromic transport-induced PV-CV and diffusive CV-PV gradients through CMFDA infusion

**Movie S9:** Visualization of lobule-wide antidromic diffusive CV-PV gradients through photoactivation of CMNB-Fluo

**Movie S10:** Photoactivation of CMNB-Fluo in 3D showing symmetric dispersion in the liver canalicular network.

## Appendix I

Compilation of raw spatial and temporal autocorrelation curves, and individual fits are provided.

Raw data files and analysis code are available at: https://lab.vartak.org/repository/bile-flux/

