## Supplementary material for "Intravital dynamic and correlative imaging reveals diffusion-dominated canalicular and flow-augmented ductular bile flux": Table S1

### Table S1: Results and data for velocity and diffusion in liver microcompartments obtained from fluorescence loss after photoactivation

| Sequences | Velocity µm/s Canaliculi CV Basal | Diffusion $\mu m^2/s$ Canaliculi CV Basal | Velocity µm/s Canaliculi MZ Basal | Diffusion $\mu m^2/s$ Canaliculi MZ Basal | Velocity µm/s Canaliculi PV Basal | Diffusion $\mu m^2/s$ Canaliculi PV Basal | Velocity µm/s Duct IBD Basal | Diffusion $\mu m^2/s$ Duct IBD Basal | Velocity µm/s Canaliculi CV Secretin | Diffusion $\mu m^2/s$ Canaliculi CV Secretin | Velocity µm/s Canaliculi MZ Secretin | Diffusion $\mu m^2/s$ Canaliculi MZ Secretin | Velocity µm/s Canaliculi PV Secretin | Diffusion $\mu m^2/s$ Canaliculi PV Secretin | Velocity µm/s Duct IBD Secretin | Diffusion $\mu m^2/s$ Duct IBD Secretin | Velocity µm/s Canaliculi CV TCA | Diffusion $\mu m^2/s$ Canaliculi CV TCA | Velocity µm/s Canaliculi MZ TCA | Diffusion $\mu m^2/s$ Canaliculi MZ TCA | Velocity µm/s Canaliculi PV TCA | Diffusion $\mu m^2/s$ Canaliculi PV TCA | Velocity µm/s Duct IBD TCA | Diffusion $\mu m^2/s$ Duct IBD TCA |
|---|---|---|---|---|---|---|---|---|---|---|---|---|---|---|---|---|---|---|---|---|---|---|---|---|
| 1 | 0.01 | 11.96171 | 0.05638 | 4.55946 | 0.01667 | 3.36065 | 1.02852 | 1.80329 | 0.00889 | 0.44163 | 0.025 | 5.45371 | 0.03611 | 8.28734 | 2.51056 | 3.60312 | 0.00314 | 2.5 | 0.04089 | 8.2 | 0.04433 | 4.3 | 7.9 | 3.7 |
| 2 | 0 | 2.07459 | 0.00767 | 7.73468 | 0.01367 | 4.55946 | 1.12745 | 3.30941 | 0.01267 | 2.07459 | 0.007 | 2.82112 | 0.01367 | 8.40462 | 6.53843 | 2.49563 | 0.05263 | 6.1 | 0.00725 | 2.6 | 0.0274 | 7.3 | 8 | 3.3 |
| 3 | 0.038 | 4.26106 | 0.06 | 2.23747 | 0 | 4.07288 | 0.76726 | 1.80754 | 0.01 | 5.45371 | 0.03 | 5.45371 | 0.04 | 2.90233 | 4.35895 | 5.96291 | 0.00778 | 6.9 | 0.01544 | 4.2 | 0.00395 | 5.5 | 6.5 | 4 |
| 4 | 0.026 | 9.75088 | 0.034 | 9.33593 | 0.01 | 5.46837 | 1.1 | n.d | 0.03 | 8.97666 | 0.05 | 2.86016 | 0.05 | 1.82177 | 3.16 | 12.24007 | 0 | 4.16 | 0 | 4.83 | 0.01 | 4.23 | 3 | 4.6 |
| 5 | 0.055 | 3.2955 | 0.007 | 5.4991 | 0.005 | 4.4852 | 1.07 | 5.4583 | 0.05 | 2.1286 | 0.02 | 5.8392 | 0.01 | 4.775 | 3.22 | 2.28362 | 0.02 | 3.73 | 0.01 | 5.65 | 0.03 | 4.25 | 4.863 | 5.1 |
| 6 | 0 | 2.0421 | 0 | 4.1519 | 0.054 | 2.3116 | 0.84 | 5.9656 | 0 | 2.4565 | 0.02 | 3.5252 | 0.02 | 2.7555 | 4 | 2.93458 | 0.03 | 2.98 | 0.05 | 2 | 0.03 | 2.36 | 1.418 | 8.3 |
| 7 | 0 | 2.9635 | 0 | 4.4574 | 0 | 5.1716 | 0.94 | 5.8044 | 0 | 4.1603 | 0.04 | 4.9633 | 0.05 | 2.6647 | 4.1 | 2.88 | 0.05 | 5.53 | 0.04 | 3.44 | 0.02 | 5.75 | 7.78 | 4.9 |
| 8 | 0 | 3.1838 | 0 | 5.1144 | 0 | 5.8015 | 0.8 | 4.3501 | 0.03 | 5.0912 | 0.01 | 5.4724 | 0.03 | 5.8072 | 3.3 | 3.59 | 0.05 | 3.63 | 0.05 | 2.21 | 0.04 | 4.76 | 3.59 | 4.9 |
| 9 | 0 | 2.5832 | 0 | 2.2114 | 0 | 2.2161 | 1.19 | 4.9526 | 0.01 | 4.7039 | 0.05 | 3.8971 | 0.02 | 3.8127 | 3.88 | 5.3 | 0.05 | 4.68 | 0.04 | 5.61 | 0 | 5.78 | 1.33 | 5.8 |
| 10 | | | | | | | | | | | | | | | | | | | | | | | 7.86 | |
| 11 | | | | | | | | | | | | | | | | | | | | | | | 6.27 | |
| 12 | | | | | | | | | | | | | | | | | | | | | | | 5.5 | |
| 13 | | | | | | | | | | | | | | | | | | | | | | | 7.39 | |
