## Supplementary material for "Intravital dynamic and correlative imaging reveals diffusion-dominated canalicular and flow-augmented ductular bile flux": Table S2

### Table S2: Results and data for diffusion in liver microcompartments obtained from spatial ICS.

| File | Mouse | Treatment | Zone | D (RICS Y) $m^2.s^{-1}$ | w 0 $m^2$ | w z $m^2$ | COD : R^2 | BOUND ENCOUNTERED | REGION |
|---|---|---|---|---|---|---|---|---|---|
| [CV]"Image 16.czi_RICS"! | 1 | Basal | CV | 1.49818E-12 | 3.24E-07 | 8.65E-7 | 0.99768 | -- | Canaliculi |
| [CV]"Image 33.czi_RICS"! | 3 | Basal | CV | 2.54589E-12 | 3.24E-07 | 8.65E-7 | 0.84014 | -- | Canaliculi |
| [CV]"Image 19.czi_RICS"! | 2 | Basal | CV | 1E-12 | 3.24E-07 | 8.65E-7 | 0.99207 | YES | Canaliculi |
| [CV]"Image 17.czi_RICS"! | 1 | Basal | CV | 1.25462E-12 | 3.24E-07 | 8.65E-7 | 0.97896 | -- | Canaliculi |
| [CV]"Image 20.czi_RICS"! | 2 | Basal | CV | 4.79231E-12 | 3.24E-07 | 8.65E-7 | 0.95964 | -- | Canaliculi |
| [CV]"Image 30.czi_RICS"! | 3 | Basal | CV | 3.3581E-12 | 3.24E-07 | 8.65E-7 | 0.72333 | -- | Canaliculi |
| [CV]"Image 28.czi_RICS"! | 2 | Basal | CV | 3.54218E-12 | 3.24E-07 | 8.65E-7 | 0.83348 | -- | Canaliculi |
| [CV]"Image 18.czi_RICS"! | 1 | Basal | CV | 1E-12 | 3.24E-07 | 8.65E-7 | -0.15385 | YES | Canaliculi |
| [CV]"Image 41.czi_RICS"! | 4 | Basal | CV | 6.11408E-12 | 3.24E-07 | 8.65E-7 | 0.98638 | -- | Canaliculi |
| [IBD]"Image 9.czi_RICS"! | 2 | Basal | IBD | 1.33824E-12 | 3.24E-07 | 8.65E-7 | 0.99993 | -- | Duct |
| [IBD]"Image 11.czi_RICS"! | 2 | Basal | IBD | 5.5818E-12 | 3.24E-07 | 8.65E-7 | 0.99984 | -- | Duct |
| [IBD]"Image 37.czi_RICS"! | 3 | Basal | IBD | 2.94268E-12 | 3.24E-07 | 8.65E-7 | 0.99894 | -- | Duct |
| [IBD]"Image 38.czi_RICS"! | 3 | Basal | IBD | 1.59971E-12 | 3.24E-07 | 8.65E-7 | 0.99923 | -- | Duct |
| [IBD]"Image 40.czi_RICS"! | 3 | Basal | IBD | 1E-12 | 3.24E-07 | 8.65E-7 | 0.99575 | YES | Duct |
| [IBD]"Image 8.czi_RICS"! | 2 | Basal | IBD | 5.96825E-12 | 3.24E-07 | 8.65E-7 | 0.99976 | -- | Duct |
| [IBD]"Image 7.czi_RICS"! | 2 | Basal | IBD | 7.01209E-12 | 3.24E-07 | 8.65E-7 | 0.99988 | -- | Duct |
| [IBD]"Image 4.czi_RICS"! | 1 | Basal | IBD | 1.00847E-12 | 3.24E-07 | 8.65E-7 | 0.99638 | -- | Duct |
| [IBD]"Image 3.czi_RICS"! | 1 | Basal | IBD | 1.59564E-12 | 3.24E-07 | 8.65E-7 | 0.99804 | -- | Duct |
| [MZ]"Image 18.czi_RICS"! | 1 | Basal | MZ | 3.44514E-12 | 3.24E-07 | 8.65E-7 | 0.97732 | -- | Canaliculi |
| [MZ]"Image 23.czi_RICS"! | 2 | Basal | MZ | 1E-12 | 3.24E-07 | 8.65E-7 | 0.9952 | YES | Canaliculi |
| [MZ]"Image 24.czi_RICS"! | 3 | Basal | MZ | 1E-12 | 3.24E-07 | 8.65E-7 | 0.99341 | YES | Canaliculi |
| [MZ]"Image 25.czi_RICS"! | 3 | Basal | MZ | 7.12829E-12 | 3.24E-07 | 8.65E-7 | 0.99575 | -- | Canaliculi |
| [MZ]"Image 26.czi_RICS"! | 3 | Basal | MZ | 4.12935E-12 | 3.24E-07 | 8.65E-7 | 0.96603 | -- | Canaliculi |
| [MZ]"Image 42.czi_RICS"! | 4 | Basal | MZ | 1E-12 | 3.24E-07 | 8.65E-7 | 0.81336 | YES | Canaliculi |
| [MZ]"Image 22.czi_RICS"! | 2 | Basal | MZ | 6.23353E-12 | 3.24E-07 | 8.65E-7 | 0.95823 | -- | Canaliculi |
| [MZ]"Image 20.czi_RICS"! | 1 | Basal | MZ | 1.09383E-12 | 3.24E-07 | 8.65E-7 | 0.57057 | -- | Canaliculi |
| [MZ]"Image 19.czi_RICS"! | 1 | Basal | MZ | 3.22588E-12 | 3.24E-07 | 8.65E-7 | 0.98562 | -- | Canaliculi |
| [MZ]"Image 21.czi_RICS"! | 2 | Basal | MZ | 1E-12 | 3.24E-07 | 8.65E-7 | 0.99924 | YES | Canaliculi |
| [PV]"Image 9.czi_RICS"! | 2 | Basal | PV | 1E-12 | 2.93E-07 | 8.65E-7 | 0.92741 | YES | Canaliculi |
| [PV]"Image 10.czi_RICS"! | 2 | Basal | PV | 1.36758E-12 | 2.93E-07 | 8.65E-7 | 0.91011 | -- | Canaliculi |
| [PV]"Image 11.czi_RICS"! | 3 | Basal | PV | 3.63753E-12 | 2.93E-07 | 8.65E-7 | 0.91804 | -- | Canaliculi |
| [PV]"Image 12.czi_RICS"! | 3 | Basal | PV | 2.00446E-12 | 2.93E-07 | 8.65E-7 | 0.93913 | -- | Canaliculi |
| [PV]"Image 13.czi_RICS"! | 3 | Basal | PV | 9.49267E-12 | 2.93E-07 | 8.65E-7 | 0.91069 | -- | Canaliculi |
| [PV]"Image 4.czi_RICS"! | 1 | Basal | PV | 4.86048E-12 | 2.93E-07 | 8.65E-7 | 0.86755 | -- | Canaliculi |
| [PV]"Image 43.czi_RICS"! | 4 | Basal | PV | 2.68047E-12 | 2.93E-07 | 8.65E-7 | 0.99594 | -- | Canaliculi |
| [PV]"Image 5.czi_RICS"! | 1 | Basal | PV | 1.5133E-12 | 2.93E-07 | 8.65E-7 | 0.97466 | -- | Canaliculi |
| [PV]"Image 6.czi_RICS"! | 1 | Basal | PV | 2.07567E-12 | 2.93E-07 | 8.65E-7 | 0.9453 | -- | Canaliculi |
| [PV]"Image 7.czi_RICS"! | 2 | Basal | PV | 1.81998E-12 | 2.93E-07 | 8.65E-7 | 0.87697 | -- | Canaliculi |
| [PV]"Image 8.czi_RICS"! | 2 | Basal | PV | 1E-12 | 2.93E-07 | 8.65E-7 | 0.8544 | YES | Canaliculi |
| [SecretinCV]"Image 125.czi_RICS"! | 5 | Secretin | CV | 1.92024E-12 | 3.24E-07 | 8.65E-7 | 0.98964 | -- | Canaliculi |
| [SecretinCV]"Image 130.czi_RICS"! | 5 | Secretin | CV | 6.87061E-12 | 3.24E-07 | 8.65E-7 | 0.97349 | -- | Canaliculi |
| [SecretinCV]"Image 44.czi_RICS"! | 2 | Secretin | CV | 1.38789E-12 | 3.24E-07 | 8.65E-7 | 0.98632 | -- | Canaliculi |
| [SecretinCV]"Image 25.czi_RICS"! | 1 | Secretin | CV | 4.21237E-12 | 3.24E-07 | 8.65E-7 | 0.99719 | -- | Canaliculi |
| [SecretinCV]"Image 28.czi_RICS"! | 1 | Secretin | CV | 1.20258E-12 | 3.24E-07 | 8.65E-7 | 0.98255 | -- | Canaliculi |
| [SecretinCV]"Image 128.czi_RICS"! | 5 | Secretin | CV | 4.07264E-12 | 3.24E-07 | 8.65E-7 | 0.98393 | -- | Canaliculi |
| [SecretinIBD]"Image 88.czi_RICS"! | 3 | Secretin | IBD | 1.03927E-12 | 3.24E-07 | 8.65E-7 | 0.99218 | -- | Duct |
| [SecretinIBD]"Image 73.czi_RICS"! | 3 | Secretin | IBD | 1.4928E-12 | 3.24E-07 | 8.65E-7 | 0.98192 | -- | Duct |
| [SecretinIBD]"Image 83.czi_RICS"! | 3 | Secretin | IBD | 2.36198E-12 | 3.24E-07 | 8.65E-7 | 0.98613 | -- | Duct |
| [SecretinIBD]"Image 50.czi_RICS"! | 2 | Secretin | IBD | 2.36198E-12 | 3.24E-07 | 8.65E-7 | 0.95283 | -- | Duct |
| [SecretinIBD]"Image 49.czi_RICS"! | 2 | Secretin | IBD | 1.03927E-12 | 3.24E-07 | 8.65E-7 | 0.96287 | -- | Duct |
| [SecretinIBD]"Image 48.czi_RICS"! | 2 | Secretin | IBD | 1.12066E-12 | 3.24E-07 | 8.65E-7 | 0.97781 | -- | Duct |
| [SecretinIBD]"Image 23.czi_RICS"! | 1 | Secretin | IBD | 2.36198E-12 | 3.24E-07 | 8.65E-7 | 0.94994 | -- | Duct |
| [SecretinIBD]"Image 22.czi_RICS"! | 1 | Secretin | IBD | 1.48217E-12 | 3.24E-07 | 8.65E-7 | 0.98723 | -- | Duct |
| [SecretinIBD]"Image 21.czi_RICS"! | 1 | Secretin | IBD | 2.36198E-12 | 3.24E-07 | 8.65E-7 | 0.99306 | -- | Duct |
| [SecretinIBD]"Image 20.czi_RICS"! | 1 | Secretin | IBD | 2.35172E-12 | 3.24E-07 | 8.65E-7 | 0.98424 | -- | Duct |
| [SecretinIBD]"Image 19.czi_RICS"! | 1 | Secretin | IBD | 2.36198E-12 | 3.24E-07 | 8.65E-7 | 0.9955 | -- | Duct |
| [SecretinIBD]"Image 77.czi_RICS"! | 3 | Secretin | IBD | 1.03927E-12 | 3.24E-07 | 8.65E-7 | 0.9661 | -- | Duct |
| [SecretinIBD]"Image 79.czi_RICS"! | 3 | Secretin | IBD | 1.04168E-12 | 3.24E-07 | 8.65E-7 | 0.98205 | -- | Duct |
| [SecretinMZ]"Image 120.czi_RICS"! | 3 | Secretin | MZ | 1E-12 | 2.93E-07 | 8.65E-7 | 0.93201 | YES | Canaliculi |
| [SecretinMZ]"Image 119.czi_RICS"! | 2 | Secretin | MZ | 1.20737E-12 | 2.93E-07 | 8.65E-7 | 0.90432 | -- | Canaliculi |
| [SecretinMZ]"Image 118.czi_RICS"! | 2 | Secretin | MZ | 1.1435E-12 | 2.93E-07 | 8.65E-7 | 0.92354 | -- | Canaliculi |
| [SecretinMZ]"Image 115.czi_RICS"! | 2 | Secretin | MZ | 1.21463E-12 | 2.93E-07 | 8.65E-7 | 0.87047 | -- | Canaliculi |
| [SecretinMZ]"Image 114.czi_RICS"! | 2 | Secretin | MZ | 1E-12 | 2.93E-07 | 8.65E-7 | 0.87458 | YES | Canaliculi |
| [SecretinMZ]"Image 113.czi_RICS"! | 2 | Secretin | MZ | 1E-12 | 2.93E-07 | 8.65E-7 | 0.85312 | YES | Canaliculi |
| [SecretinMZ]"Image 122.czi_RICS"! | 3 | Secretin | MZ | 2.75571E-12 | 2.93E-07 | 8.65E-7 | 0.9924 | -- | Canaliculi |
| [SecretinMZ]"Image 26.czi_RICS"! | 1 | Secretin | MZ | 3.62623E-12 | 2.93E-07 | 8.65E-7 | 0.98986 | -- | Canaliculi |
| [SecretinMZ]"Image 29.czi_RICS"! | 1 | Secretin | MZ | 4.18184E-12 | 2.93E-07 | 8.65E-7 | 0.99883 | -- | Canaliculi |
| [SecretinMZ]"Image 121.czi_RICS"! | 3 | Secretin | MZ | 1.1212E-12 | 2.93E-07 | 8.65E-7 | 0.94022 | -- | Canaliculi |
| [SecretinPV]"Image 47.czi_RICS"! | 2 | Secretin | PV | 6.4247E-12 | 3.24E-07 | 8.65E-7 | 0.99697 | -- | Canaliculi |
| [SecretinPV]"Image 104.czi_RICS"! | 3 | Secretin | PV | 2.86065E-12 | 3.24E-07 | 8.65E-7 | 0.98808 | -- | Canaliculi |
| [SecretinPV]"Image 103.czi_RICS"! | 3 | Secretin | PV | 6.10632E-12 | 3.24E-07 | 8.65E-7 | 0.95755 | -- | Canaliculi |
| [SecretinPV]"Image 100.czi_RICS"! | 3 | Secretin | PV | 2.20081E-12 | 3.24E-07 | 8.65E-7 | 0.9974 | -- | Canaliculi |
| [SecretinPV]"Image 108.czi_RICS"! | 4 | Secretin | PV | 1.55538E-12 | 3.24E-07 | 8.65E-7 | 0.99155 | -- | Canaliculi |
| [SecretinPV]"Image 109.czi_RICS"! | 3 | Secretin | PV | 7.30286E-12 | 3.24E-07 | 8.65E-7 | 0.99476 | -- | Canaliculi |
| [SecretinPV]"Image 107.czi_RICS"! | 4 | Secretin | PV | 7.0212E-13 | 3.24E-07 | 8.65E-7 | 0.99589 | -- | Canaliculi |
| [SecretinPV]"Image 105.czi_RICS"! | 4 | Secretin | PV | 3.55175E-12 | 3.24E-07 | 8.65E-7 | 0.99517 | -- | Canaliculi |
| [TCACV]"Image 294.czi_RICS"! | 4 | TCA | CV | 1E-12 | 3.24E-07 | 8.65E-7 | 0.98316 | -- | Canaliculi |
| [TCACV]"Image 237.czi_RICS"! | 1 | TCA | CV | 1.66102E-12 | 3.24E-07 | 8.65E-7 | 0.98457 | -- | Canaliculi |
| [TCACV]"Image 290.czi_RICS"! | 3 | TCA | CV | 2.64286E-12 | 3.24E-07 | 8.65E-7 | 0.99699 | -- | Canaliculi |
| [TCACV]"Image 289.czi_RICS"! | 3 | TCA | CV | 2.77094E-12 | 3.24E-07 | 8.65E-7 | 0.96047 | -- | Canaliculi |
| [TCACV]"Image 288.czi_RICS"! | 2 | TCA | CV | 5.10148E-12 | 3.24E-07 | 8.65E-7 | 0.74136 | -- | Canaliculi |
| [TCACV]"Image 287.czi_RICS"! | 2 | TCA | CV | 2.69749E-12 | 3.24E-07 | 8.65E-7 | 0.97011 | -- | Canaliculi |
| [TCACV]"Image 286.czi_RICS"! | 2 | TCA | CV | 4.15997E-12 | 3.24E-07 | 8.65E-7 | 0.98225 | -- | Canaliculi |
| [TCACV]"Image 295.czi_RICS"! | 4 | TCA | CV | 4.66769E-12 | 3.24E-07 | 8.65E-7 | 0.80377 | -- | Canaliculi |
| [TCACV]"Image 291.czi_RICS"! | 3 | TCA | CV | 3.78054E-12 | 3.24E-07 | 8.65E-7 | 0.91438 | -- | Canaliculi |
| [TCACV]"Image 243.czi_RICS"! | 1 | TCA | CV | 2.02501E-12 | 3.24E-07 | 8.65E-7 | 0.96537 | -- | Canaliculi |
| [TCACV]"Image 240.czi_RICS"! | 1 | TCA | CV | 8.28384E-12 | 3.24E-07 | 8.65E-7 | 0.97838 | -- | Canaliculi |
| [TCACV]"Image 241.czi_RICS"! | 1 | TCA | CV | 3.42236E-12 | 3.24E-07 | 8.65E-7 | 0.98753 | -- | Canaliculi |
| [TCAIBD]"Image 210.czi_RICS"! | 1 | TCA | IBD | 2.75571E-12 | 3.55E-07 | 8.65E-7 | 0.99647 | -- | Duct |
| [TCAIBD]"Image 241.czi_RICS"! | 4 | TCA | IBD | 1.8296E-12 | 3.55E-07 | 8.65E-7 | 0.938 | -- | Duct |
| [TCAIBD]"Image 254.czi_RICS"! | 3 | TCA | IBD | 1.74212E-12 | 3.55E-07 | 8.65E-7 | 0.93364 | -- | Duct |
| [TCAIBD]"Image 253.czi_RICS"! | 3 | TCA | IBD | 1.19333E-12 | 3.55E-07 | 8.65E-7 | 0.94008 | -- | Duct |
| [TCAIBD]"Image 250.czi_RICS"! | 3 | TCA | IBD | 2.40771E-12 | 3.55E-07 | 8.65E-7 | 0.9982 | -- | Duct |
| [TCAIBD]"Image 249.czi_RICS"! | 3 | TCA | IBD | 1E-12 | 3.55E-07 | 8.65E-7 | 0.95582 | YES | Duct |
| [TCAIBD]"Image 245.czi_RICS"! | 2 | TCA | IBD | 1E-12 | 3.55E-07 | 8.65E-7 | 0.98845 | YES | Duct |
| [TCAIBD]"Image 244.czi_RICS"! | 2 | TCA | IBD | 1E-12 | 3.55E-07 | 8.65E-7 | 0.97347 | YES | Duct |
| [TCAIBD]"Image 243.czi_RICS"! | 2 | TCA | IBD | 2.83722E-12 | 3.55E-07 | 8.65E-7 | 0.99163 | -- | Duct |
| [TCAIBD]"Image 220.czi_RICS"! | 1 | TCA | IBD | 1.02124E-12 | 3.55E-07 | 8.65E-7 | 0.94137 | -- | Duct |
| [TCAIBD]"Image 215.czi_RICS"! | 1 | TCA | IBD | 4.0044E-12 | 3.55E-07 | 8.65E-7 | 0.9903 | -- | Duct |
| [TCAMZ]"Image 249.czi_RICS"! | 1 | TCA | MZ | 1E-11 | 3.55E-07 | 8.65E-7 | 0.96142 | YES | Canaliculi |
| [TCAMZ]"Image 276.czi_RICS"! | 2 | TCA | MZ | 5.32468E-12 | 3.55E-07 | 8.65E-7 | 0.98978 | -- | Canaliculi |
| [TCAMZ]"Image 277.czi_RICS"! | 2 | TCA | MZ | 1E-11 | 3.55E-07 | 8.65E-7 | 0.98993 | YES | Canaliculi |
| [TCAMZ]"Image 278.czi_RICS"! | 2 | TCA | MZ | 2.81375E-12 | 3.55E-07 | 8.65E-7 | 0.98971 | -- | Canaliculi |
| [TCAMZ]"Image 279.czi_RICS"! | 3 | TCA | MZ | 1.93655E-12 | 3.55E-07 | 8.65E-7 | 0.99819 | -- | Canaliculi |
| [TCAMZ]"Image 280.czi_RICS"! | 3 | TCA | MZ | 1E-11 | 3.55E-07 | 8.65E-7 | 0.98157 | YES | Canaliculi |
| [TCAMZ]"Image 281.czi_RICS"! | 3 | TCA | MZ | 1.56079E-12 | 3.55E-07 | 8.65E-7 | 0.99302 | -- | Canaliculi |
| [TCAMZ]"Image 282.czi_RICS"! | 4 | TCA | MZ | 2.61983E-12 | 3.55E-07 | 8.65E-7 | 0.99942 | -- | Canaliculi |
| [TCAMZ]"Image 283.czi_RICS"! | 4 | TCA | MZ | 4.15682E-12 | 3.55E-07 | 8.65E-7 | 0.99012 | -- | Canaliculi |
| [TCAMZ]"Image 255.czi_RICS"! | 1 | TCA | MZ | 1E-12 | 3.55E-07 | 8.65E-7 | 0.96552 | YES | Canaliculi |
| [TCAPV]"Image 266.czi_RICS"! | 2 | TCA | PV | 2.62999E-12 | 2.97E-07 | 8.65E-7 | 0.89279 | -- | Canaliculi |
| [TCAPV]"Image 265.czi_RICS"! | 2 | TCA | PV | 2.97681E-12 | 2.97E-07 | 8.65E-7 | 0.99139 | -- | Canaliculi |
| [TCAPV]"Image 264.czi_RICS"! | 2 | TCA | PV | 1.60585E-12 | 2.97E-07 | 8.65E-7 | 0.99967 | -- | Canaliculi |
| [TCAPV]"Image 230.czi_RICS"! | 1 | TCA | PV | 2.5499E-12 | 2.97E-07 | 8.65E-7 | 0.83953 | -- | Canaliculi |
| [TCAPV]"Image 228.czi_RICS"! | 1 | TCA | PV | 1.693E-12 | 2.97E-07 | 8.65E-7 | 0.99709 | -- | Canaliculi |
| [TCAPV]"Image 267.czi_RICS"! | 2 | TCA | PV | 2.73224E-13 | 2.97E-07 | 8.65E-7 | 0.99634 | -- | Canaliculi |
| [TCAPV]"Image 263.czi_RICS"! | 3 | TCA | PV | 3.05566E-12 | 2.97E-07 | 8.65E-7 | 0.97546 | -- | Canaliculi |
| [TCAPV]"Image 271.czi_RICS"! | 3 | TCA | PV | 8.77452E-12 | 2.97E-07 | 8.65E-7 | 0.99589 | -- | Canaliculi |
| [TCAPV]"Image 270.czi_RICS"! | 3 | TCA | PV | 5.97421E-13 | 2.97E-07 | 8.65E-7 | 0.99717 | -- | Canaliculi |
| [TCAPV]"Image 269.czi_RICS"! | 2 | TCA | PV | 1.50485E-12 | 2.97E-07 | 8.65E-7 | 0.96273 | -- | Canaliculi |

--
