## Supplementary material for "Intravital dynamic and correlative imaging reveals diffusion-dominated canalicular and flow-augmented ductular bile flux": Table S3

### Table S3: Results and data for diffusion and velocity in liver microcompartments obtained from temporal ICS.

| File | MOUSE | TREATMENT | ZONE | REGION | D $m^2.s^{-1}$ | V $m.s^{-1}$ | BOUND ENCOUNTERED | wz $m^2$ | w0 $m^2$ | COD : R^2 |
|---|---|---|---|---|---|---|---|---|---|---|
| [BasalCV]"Image 16.czi_TICS"! | 1 | Basal | CV | Canaliculi | 1.18176E-12 | 8.42508E-8 | -- | 3.27E-7 | 8.67E-7 | 0.994 |
| [BasalCV]"Image 17.czi_TICS"! | 1 | Basal | CV | Canaliculi | 3.739E-12 | 5.55958E-8 | -- | 3.27E-7 | 8.67E-7 | 0.982 |
| [BasalCV]"Image 18.czi_TICS"! | 1 | Basal | CV | Canaliculi | 3.80206E-12 | 6.04792E-8 | -- | 3.27E-7 | 8.67E-7 | 0.970 |
| [BasalCV]"Image 19.czi_TICS"! | 1 | Basal | CV | Canaliculi | 3.53596E-12 | 5.91102E-8 | -- | 3.27E-7 | 8.67E-7 | 0.986 |
| [BasalCV]"Image 20.czi_TICS"! | 2 | Basal | CV | Canaliculi | 3.66822E-12 | 5.83783E-8 | -- | 3.27E-7 | 8.67E-7 | 0.973 |
| [BasalCV]"Image 27.czi_TICS"! | 2 | Basal | CV | Canaliculi | 4.03967E-12 | 5.255E-8 | -- | 3.27E-7 | 8.67E-7 | 0.931 |
| [BasalCV]"Image 28.czi_TICS"! | 2 | Basal | CV | Canaliculi | 4.11419E-12 | 5.22795E-8 | -- | 3.27E-7 | 8.67E-7 | 0.979 |
| [BasalCV]"Image 30.czi_TICS"! | 2 | Basal | CV | Canaliculi | 3.70122E-12 | 5.25868E-8 | -- | 3.27E-7 | 8.67E-7 | 0.985 |
| [BasalCV]"Image 31.czi_TICS"! | 3 | Basal | CV | Canaliculi | 4.20724E-12 | 5.27659E-8 | -- | 3.27E-7 | 8.67E-7 | 0.990 |
| [BasalCV]"Image 32.czi_TICS"! | 3 | Basal | CV | Canaliculi | 3.81011E-12 | 5.33105E-8 | -- | 3.27E-7 | 8.67E-7 | 0.989 |
| [BasalCV]"Image 33.czi_TICS"! | 3 | Basal | CV | Canaliculi | 2.9889E-12 | 5.36648E-8 | -- | 3.27E-7 | 8.67E-7 | 0.958 |
| [BasalCV]"Image 34.czi_TICS"! | 3 | Basal | CV | Canaliculi | 3.04783E-12 | 6.11854E-8 | -- | 3.27E-7 | 8.67E-7 | 0.985 |
| [BasalIBD]"Image 3.czi_TICS"! | 3 | Basal | IBD | Duct | 2.21957E-12 | 1.19284E-6 | -- | 3.27E-7 | 8.67E-7 | 0.997 |
| [BasalIBD]"Image 11.czi_TICS"! | 2 | Basal | IBD | Duct | 1.63381E-12 | 1.22692E-6 | -- | 3.27E-7 | 8.67E-7 | 0.982 |
| [BasalIBD]"Image 4.czi_TICS"! | 2 | Basal | IBD | Duct | 1.70066E-12 | 1.32989E-6 | -- | 3.27E-7 | 8.67E-7 | 0.988 |
| [BasalIBD]"Image 40.czi_TICS"! | 2 | Basal | IBD | Duct | 4.6465E-12 | 1.08381E-6 | -- | 3.27E-7 | 8.67E-7 | 0.997 |
| [BasalIBD]"Image 9.czi_TICS"! | 2 | Basal | IBD | Duct | 3.38727E-12 | 1.1777E-6 | -- | 3.27E-7 | 8.67E-7 | 0.987 |
| [BasalIBD]"Image 7.czi_TICS"! | 1 | Basal | IBD | Duct | 6.12926E-12 | 1.21795E-6 | -- | 3.27E-7 | 8.67E-7 | 0.988 |
| [BasalIBD]"Image 38.czi_TICS"! | 3 | Basal | IBD | Duct | 1.33852E-12 | 1.42736E-6 | -- | 3.27E-7 | 8.67E-7 | 0.993 |
| [BasalIBD]"Image 8.czi_TICS"! | 1 | Basal | IBD | Duct | 2.56958E-12 | 1.79467E-6 | -- | 3.27E-7 | 8.67E-7 | 0.973 |
| [BasalMZ]"Image 42.czi_TICS"! | 2 | Basal | MZ | Canaliculi | 1.96536E-12 | 8.148E-8 | -- | 3.27E-7 | 8.67E-7 | 0.968 |
| [BasalMZ]"Image 19.czi_TICS"! | 3 | Basal | MZ | Canaliculi | 1.30366E-12 | 9.71264E-8 | -- | 3.27E-7 | 8.67E-7 | 0.974 |
| [BasalMZ]"Image 20.czi_TICS"! | 1 | Basal | MZ | Canaliculi | 6.66721E-13 | 2.01757E-7 | Bound encountered for V | 3.27E-7 | 8.67E-7 | 0.980 |
| [BasalMZ]"Image 21.czi_TICS"! | 1 | Basal | MZ | Canaliculi | 1.05576E-12 | 1E-8 | Bound encountered for V | 3.27E-7 | 8.67E-7 | 0.775 |
| [BasalMZ]"Image 22.czi_TICS"! | 1 | Basal | MZ | Canaliculi | 1.1292E-12 | 9.74731E-8 | -- | 3.27E-7 | 8.67E-7 | 0.966 |
| [BasalMZ]"Image 23.czi_TICS"! | 2 | Basal | MZ | Canaliculi | 1.45343E-12 | 9.65728E-8 | -- | 3.27E-7 | 8.67E-7 | 0.965 |
| [BasalMZ]"Image 24.czi_TICS"! | 2 | Basal | MZ | Canaliculi | 1.30339E-12 | 9.8641E-8 | -- | 3.27E-7 | 8.67E-7 | 0.956 |
| [BasalMZ]"Image 25.czi_TICS"! | 3 | Basal | MZ | Canaliculi | 1.26068E-12 | 9.55133E-8 | -- | 3.27E-7 | 8.67E-7 | 0.969 |
| [BasalMZ]"Image 26.czi_TICS"! | 3 | Basal | MZ | Canaliculi | 1.25106E-12 | 1E-8 | Bound encountered for V | 3.27E-7 | 8.67E-7 | 0.973 |
| [BasalPV]"Image 9.czi_TICS"! | 3 | Basal | PV | Canaliculi | 1.57285E-12 | 9.09692E-9 | -- | 3.27E-7 | 8.67E-7 | 0.988 |
| [BasalPV]"Image 8.czi_TICS"! | 3 | Basal | PV | Canaliculi | 1.62824E-12 | 6.08665E-9 | -- | 3.27E-7 | 8.67E-7 | 0.994 |
| [BasalPV]"Image 6.czi_TICS"! | 2 | Basal | PV | Canaliculi | 1.72562E-12 | 5.29324E-9 | -- | 3.27E-7 | 8.67E-7 | 0.981 |
| [BasalPV]"Image 5.czi_TICS"! | 2 | Basal | PV | Canaliculi | 1.60713E-12 | 7.19867E-9 | -- | 3.27E-7 | 8.67E-7 | 0.994 |
| [BasalPV]"Image 43.czi_TICS"! | 2 | Basal | PV | Canaliculi | 2.00987E-12 | 6.00927E-9 | -- | 3.27E-7 | 8.67E-7 | 0.981 |
| [BasalPV]"Image 13.czi_TICS"! | 2 | Basal | PV | Canaliculi | 1.615E-12 | 6.50355E-9 | -- | 3.27E-7 | 8.67E-7 | 0.971 |
| [BasalPV]"Image 12.czi_TICS"! | 4 | Basal | PV | Canaliculi | 1.8448E-12 | 1.62118E-8 | -- | 3.27E-7 | 8.67E-7 | 0.971 |
| [BasalPV]"Image 11.czi_TICS"! | 1 | Basal | PV | Canaliculi | 1.74818E-12 | 9.58409E-9 | -- | 3.27E-7 | 8.67E-7 | 0.970 |
| [BasalPV]"Image 7.czi_TICS"! | 3 | Basal | PV | Canaliculi | 1.24783E-12 | 1.21617E-8 | -- | 3.27E-7 | 8.67E-7 | 0.967 |
| [SecretinCV]"Image 44.czi_TICS"! | 2 | Secretin | CV | Canaliculi | 1E-12 | 1.13035E-8 | -- | 3.27E-7 | 8.67E-7 | 0.994 |
| [SecretinCV]"Image 25.czi_TICS"! | 1 | Secretin | CV | Canaliculi | 8E-12 | 1.44363E-8 | -- | 3.27E-7 | 8.67E-7 | 0.992 |
| [SecretinCV]"Image 28.czi_TICS"! | 1 | Secretin | CV | Canaliculi | 1.72221E-12 | 1E-8 | Bound encountered for V | 3.27E-7 | 8.67E-7 | 0.893 |
| [SecretinCV]"Image 127.czi_TICS"! | 2 | Secretin | CV | Canaliculi | 1.4041E-12 | 2.65302E-8 | -- | 3.27E-7 | 8.67E-7 | 0.924 |
| [SecretinCV]"Image 129.czi_TICS"! | 2 | Secretin | CV | Canaliculi | 1E-12 | 1.1367E-8 | -- | 3.27E-7 | 8.67E-7 | 0.900 |
| [SecretinCV]"Image 130.czi_TICS"! | 2 | Secretin | CV | Canaliculi | 5.15317E-12 | 1.96569E-8 | -- | 3.27E-7 | 8.67E-7 | 0.834 |
| [SecretinCV]"Image 132.czi_TICS"! | 3 | Secretin | CV | Canaliculi | 2.97974E-12 | 2.09782E-8 | -- | 3.27E-7 | 8.67E-7 | 0.913 |
| [SecretinCV]"Image 126.czi_TICS"! | 2 | Secretin | CV | Canaliculi | 1.0039E-12 | 1.03163E-8 | -- | 3.27E-7 | 8.67E-7 | 0.938 |
| [SecretinIBD]"Image 78.czi_TICS"! | 4 | Secretin | IBD | Duct | 1.14514E-12 | 2.29418E-6 | -- | 3.27E-7 | 8.67E-7 | 0.954 |
| [SecretinIBD]"Image 75.czi_TICS"! | 4 | Secretin | IBD | Duct | 1.14514E-12 | 4.08394E-6 | -- | 3.27E-7 | 8.67E-7 | 0.986 |
| [SecretinIBD]"Image 73.czi_TICS"! | 4 | Secretin | IBD | Duct | 1.14514E-12 | 2.77659E-6 | -- | 3.27E-7 | 8.67E-7 | 0.993 |
| [SecretinIBD]"Image 71.czi_TICS"! | 4 | Secretin | IBD | Duct | 1.14514E-12 | 2.76806E-6 | -- | 3.27E-7 | 8.67E-7 | 0.984 |
| [SecretinIBD]"Image 70.czi_TICS"! | 4 | Secretin | IBD | Duct | 1.14514E-12 | 2.87676E-6 | -- | 3.27E-7 | 8.67E-7 | 0.972 |
| [SecretinIBD]"Image 66.czi_TICS"! | 4 | Secretin | IBD | Duct | 1.14514E-12 | 3.00857E-6 | -- | 3.27E-7 | 8.67E-7 | 0.994 |
| [SecretinIBD]"Image 65.czi_TICS"! | 4 | Secretin | IBD | Duct | 1.14514E-12 | 3.08493E-6 | -- | 3.27E-7 | 8.67E-7 | 0.951 |
| [SecretinIBD]"Image 64.czi_TICS"! | 4 | Secretin | IBD | Duct | 1.14514E-12 | 2.94745E-6 | -- | 3.27E-7 | 8.67E-7 | 0.977 |
| [SecretinIBD]"Image 19.czi_TICS"! | 3 | Secretin | IBD | Duct | 4.53061E-12 | 1.23704E-6 | -- | 3.27E-7 | 8.67E-7 | 0.935 |
| [SecretinIBD]"Image 20.czi_TICS"! | 3 | Secretin | IBD | Duct | 4.53061E-12 | 1.51869E-6 | -- | 3.27E-7 | 8.67E-7 | 0.921 |
| [SecretinIBD]"Image 21.czi_TICS"! | 2 | Secretin | IBD | Duct | 4.53061E-12 | 2.12734E-6 | -- | 3.27E-7 | 8.67E-7 | 0.912 |
| [SecretinIBD]"Image 22.czi_TICS"! | 1 | Secretin | IBD | Duct | 4.53061E-12 | 1.59171E-6 | -- | 3.27E-7 | 8.67E-7 | 0.920 |
| [SecretinIBD]"Image 23.czi_TICS"! | 1 | Secretin | IBD | Duct | 4.53061E-12 | 1.53872E-6 | -- | 3.27E-7 | 8.67E-7 | 0.959 |
| [SecretinIBD]"Image 49.czi_TICS"! | 1 | Secretin | IBD | Duct | 4.53061E-12 | 1.58995E-6 | -- | 3.27E-7 | 8.67E-7 | 0.995 |
| [SecretinIBD]"Image 50.czi_TICS"! | 2 | Secretin | IBD | Duct | 4.53061E-12 | 1.30694E-6 | -- | 3.27E-7 | 8.67E-7 | 0.995 |
| [SecretinIBD]"Image 87.czi_TICS"! | 5 | Secretin | IBD | Duct | 1.14514E-12 | 6.54445E-6 | -- | 3.27E-7 | 8.67E-7 | 0.991 |
| [SecretinMZ]"Image 106.czi_TICS"! | 2 | Secretin | MZ | Canaliculi | 1.71435E-12 | 9.50191E-8 | -- | 3.27E-7 | 8.67E-7 | 0.988 |
| [SecretinMZ]"Image 118.czi_TICS"! | 3 | Secretin | MZ | Canaliculi | -- | -- | -- | 3.27E-7 | 8.67E-7 | 0.945 |
| [SecretinMZ]"Image 117.czi_TICS"! | 3 | Secretin | MZ | Canaliculi | 2.68369E-13 | 2.89778E-7 | -- | 3.27E-7 | 8.67E-7 | 0.975 |
| [SecretinMZ]"Image 116.czi_TICS"! | 3 | Secretin | MZ | Canaliculi | 2.31509E-12 | 4.26097E-8 | -- | 3.27E-7 | 8.67E-7 | 0.929 |
| [SecretinMZ]"Image 112.czi_TICS"! | 2 | Secretin | MZ | Canaliculi | 3.23216E-12 | 2.89778E-7 | -- | 3.27E-7 | 8.67E-7 | 0.942 |
| [SecretinMZ]"Image 110.czi_TICS"! | 2 | Secretin | MZ | Canaliculi | 2.31509E-12 | 4.26097E-7 | -- | 3.27E-7 | 8.67E-7 | 0.938 |
| [SecretinMZ]"Image 109.czi_TICS"! | 2 | Secretin | MZ | Canaliculi | 3.23216E-12 | 1.37737E-7 | -- | 3.27E-7 | 8.67E-7 | 0.940 |
| [SecretinMZ]"Image 108.czi_TICS"! | 2 | Secretin | MZ | Canaliculi | 6.21605E-12 | 1.80729E-8 | -- | 3.27E-7 | 8.67E-7 | 0.946 |
| [SecretinMZ]"Image 113.czi_TICS"! | 1 | Secretin | MZ | Canaliculi | 5.15556E-12 | 4.51043E-12 | -- | 3.27E-7 | 8.67E-7 | 0.948 |
| [SecretinMZ]"Image 120.czi_TICS"! | 2 | Secretin | MZ | Canaliculi | 4.47879E-12 | 1.47086E-8 | -- | 3.27E-7 | 8.67E-7 | 0.937 |
| [SecretinMZ]"Image 119.czi_TICS"! | 1 | Secretin | MZ | Canaliculi | 4.36967E-12 | 1.21681E-8 | -- | 3.27E-7 | 8.67E-7 | 0.949 |
| [SecretinMZ]"Image 121.czi_TICS"! | 1 | Secretin | MZ | Canaliculi | 3.71992E-12 | 6.00744E-8 | -- | 3.27E-7 | 8.67E-7 | 0.990 |
| [SecretinPV]"Image 103.czi_TICS"! | 4 | Secretin | PV | Canaliculi | 4.10077E-12 | 1.00937E-8 | -- | 3.27E-7 | 8.67E-7 | 0.990 |
| [SecretinPV]"Image 124.czi_TICS"! | 5 | Secretin | PV | Canaliculi | 5.26229E-8 | 6.98677E-8 | -- | 3.27E-7 | 8.67E-7 | 0.994 |
| [SecretinPV]"Image 120.czi_TICS"! | 5 | Secretin | PV | Canaliculi | 9.31348E-8 | 4.56768E-8 | -- | 3.27E-7 | 8.67E-7 | 0.978 |
| [SecretinPV]"Image 127.czi_TICS"! | 5 | Secretin | PV | Canaliculi | 8.44591E-8 | 1.4226E-9 | -- | 3.27E-7 | 8.67E-7 | 0.989 |
| [SecretinPV]"Image 126.czi_TICS"! | 5 | Secretin | PV | Canaliculi | 6.18967E-8 | 9.053E-8 | -- | 3.27E-7 | 8.67E-7 | 0.995 |
| [SecretinPV]"Image 107.czi_TICS"! | 4 | Secretin | PV | Canaliculi | 2.68923E-12 | 1.74327E-8 | -- | 3.27E-7 | 8.67E-7 | 0.995 |
| [SecretinPV]"Image 105.czi_TICS"! | 4 | Secretin | PV | Canaliculi | 3.04836E-12 | 1.50228E-8 | -- | 3.27E-7 | 8.67E-7 | 0.991 |
| [SecretinPV]"Image 100.czi_TICS"! | 3 | Secretin | PV | Canaliculi | 1.10218E-12 | 1.03134E-8 | -- | 3.27E-7 | 8.67E-7 | 0.900 |
| [SecretinPV]"Image 125.czi_TICS"! | 5 | Secretin | PV | Canaliculi | 1E-7 | 2.29031E-8 | -- | 3.27E-7 | 8.67E-7 | 0.996 |
| [SecretinPV]"Image 87.czi_TICS"! | 2 | Secretin | PV | Canaliculi | 2.58577E-12 | 1.85649E-8 | -- | 3.27E-7 | 8.67E-7 | 0.973 |
| [SecretinPV]"Image 86.czi_TICS"! | 2 | Secretin | PV | Canaliculi | 8.87311E-12 | 1.11513E-8 | -- | 3.27E-7 | 8.67E-7 | 0.994 |
| [SecretinPV]"Image 122.czi_TICS"! | 5 | Secretin | PV | Canaliculi | 2.43186E-12 | 2.58088E-8 | -- | 3.27E-7 | 8.67E-7 | 0.993 |
| [SecretinPV]"Image 123.czi_TICS"! | 5 | Secretin | PV | Canaliculi | 1.60763E-12 | 4.74883E-8 | -- | 3.27E-7 | 8.67E-7 | 0.995 |
| [SecretinPV]"Image 101.czi_TICS"! | 3 | Secretin | PV | Canaliculi | 1.72515E-12 | 3.56632E-8 | -- | 3.27E-7 | 8.67E-7 | 0.981 |
| [SecretinPV]"Image 121.czi_TICS"! | 3 | Secretin | PV | Canaliculi | 7.18143E-12 | 0 | No convergence | 3.27E-7 | 8.67E-7 | 0.982 |
| [SecretinPV]"Image 102.czi_TICS"! | 1 | Secretin | PV | Canaliculi | 1E-12 | 8.60124E-8 | Bound encountered | 3.27E-7 | 8.67E-7 | 0.863 |
| [TCACV]"Image 50.czi_TICS"! | 2 | TCA | CV | Canaliculi | 1.38879E-12 | 3.2569E-8 | -- | 3.27E-7 | 8.67E-7 | 0.956 |
| [TCACV]"Image 49.czi_TICS"! | 2 | TCA | CV | Canaliculi | 1.40267E-12 | 5.07107E-8 | -- | 3.27E-7 | 8.67E-7 | 0.914 |
| [TCACV]"Image 48.czi_TICS"! | 2 | TCA | CV | Canaliculi | 1.07089E-12 | 6.44886E-8 | -- | 3.27E-7 | 8.67E-7 | 0.952 |
| [TCACV]"Image 47.czi_TICS"! | 2 | TCA | CV | Canaliculi | 2.13527E-12 | 3.76973E-8 | No convergence | 3.27E-7 | 8.67E-7 | 0.005 |
| [TCACV]"Image 37.czi_TICS"! | 3 | TCA | CV | Canaliculi | 1.25937E-12 | 4.09761E-8 | -- | 3.27E-7 | 8.67E-7 | 0.983 |
| [TCACV]"Image 35.czi_TICS"! | 2 | TCA | CV | Canaliculi | 1.21799E-12 | 3.29049E-8 | -- | 3.27E-7 | 8.67E-7 | 0.959 |
| [TCACV]"Image 36.czi_TICS"! | 2 | TCA | CV | Canaliculi | 1.3328E-12 | 5.60994E-8 | -- | 3.27E-7 | 8.67E-7 | 0.951 |
| [TCAIBD]"Image 52.czi_TICS"! | 4 | TCA | IBD | Duct | 1.43918E-12 | 5.90791E-6 | -- | 3.27E-7 | 8.67E-7 | 0.994 |
| [TCAIBD]"Image 53.czi_TICS"! | 1 | TCA | IBD | Duct | 5.57762E-12 | 6.74516E-6 | -- | 3.27E-7 | 8.67E-7 | 0.954 |
| [TCAIBD]"Image 2.czi_TICS"! | 1 | TCA | IBD | Duct | 5E-13 | 3.17979E-6 | Bound encountered for D | 3.27E-7 | 8.67E-7 | 0.994 |
| [TCAIBD]"Image 3.czi_TICS"! | 2 | TCA | IBD | Duct | 5E-13 | 3.3326E-6 | Bound encountered for D | 3.27E-7 | 8.67E-7 | 0.765 |
| [TCAIBD]"Image 4.czi_TICS"! | 3 | TCA | IBD | Duct | 3.67423E-12 | 1.48967E-6 | -- | 3.27E-7 | 8.67E-7 | 0.995 |
| [TCAIBD]"Image 51.czi_TICS"! | 4 | TCA | IBD | Duct | 1.48913E-12 | 3.92781E-6 | -- | 3.27E-7 | 8.67E-7 | 0.971 |
| [TCAIBD]"Image 49.czi_TICS"! | 2 | TCA | IBD | Duct | 1.0017E-12 | 1.59947E-6 | -- | 3.27E-7 | 8.67E-7 | 0.972 |
| [TCAIBD]"Image 48.czi_TICS"! | 2 | TCA | IBD | Duct | 1.53059E-12 | 3.58227E-6 | -- | 3.27E-7 | 8.67E-7 | 0.902 |
| [TCAIBD]"Image 47.czi_TICS"! | 2 | TCA | IBD | Duct | 1.21648E-12 | 3.54761E-6 | -- | 3.27E-7 | 8.67E-7 | 0.992 |
| [TCAIBD]"Image 33.czi_TICS"! | 2 | TCA | IBD | Duct | 1.06459E-12 | 3.53527E-6 | -- | 3.27E-7 | 8.67E-7 | 0.994 |
| [TCAIBD]"Image 32.czi_TICS"! | 1 | TCA | IBD | Duct | 5.75384E-13 | 3.57115E-6 | -- | 3.27E-7 | 8.67E-7 | 0.995 |
| [TCAIBD]"Image 28.czi_TICS"! | 3 | TCA | IBD | Duct | 5E-13 | 3.62365E-6 | Bound encountered for D | 3.27E-7 | 8.67E-7 | 0.768 |
| [TCAMZ]"Image 254.czi_TICS"! | 2 | TCA | MZ | Canaliculi | 2.18474E-12 | 1.17031E-7 | -- | 3.27E-7 | 8.67E-7 | 0.948 |
| [TCAMZ]"Image 39.czi_TICS"! | 1 | TCA | MZ | Canaliculi | 4.50469E-12 | 4.54672E-8 | -- | 3.27E-7 | 8.67E-7 | 0.955 |
| [TCAMZ]"Image 38.czi_TICS"! | 1 | TCA | MZ | Canaliculi | 7.98937E-12 | 2.52759E-8 | -- | 3.27E-7 | 8.67E-7 | 0.955 |
| [TCAMZ]"Image 40.czi_TICS"! | 1 | TCA | MZ | Canaliculi | 7.73268E-12 | 1.63498E-8 | Bound encountered for D | 3.27E-7 | 8.67E-7 | 0.949 |
| [TCAMZ]"Image 251.czi_TICS"! | 1 | TCA | MZ | Canaliculi | 3.84478E-12 | 3.9621E-8 | -- | 3.27E-7 | 8.67E-7 | 0.948 |
| [TCAMZ]"Image 252.czi_TICS"! | 2 | TCA | MZ | Canaliculi | 4.55105E-12 | 7.96576E-8 | -- | 3.27E-7 | 8.67E-7 | 0.954 |
| [TCAMZ]"Image 249.czi_TICS"! | 2 | TCA | MZ | Canaliculi | 3.28118E-12 | 3.33721E-8 | -- | 3.27E-7 | 8.67E-7 | 0.949 |
| [TCAPV]"Image 31.czi_TICS"! | 4 | TCA | PV | Canaliculi | 1.09517E-12 | 9.58437E-9 | -- | 3.27E-7 | 8.67E-7 | 0.973 |
| [TCAPV]"Image 42.czi_TICS"! | 3 | TCA | PV | Canaliculi | 1.28869E-12 | 1.23412E-8 | -- | 3.27E-7 | 8.67E-7 | 0.905 |
| [TCAPV]"Image 44.czi_TICS"! | 3 | TCA | PV | Canaliculi | 1.31665E-12 | 1.27885E-8 | Bound encountered for V | 3.27E-7 | 8.67E-7 | 0.958 |
| [TCAPV]"Image 45.czi_TICS"! | 3 | TCA | PV | Canaliculi | 1.1481E-12 | 1.16795E-8 | -- | 3.27E-7 | 8.67E-7 | 0.961 |
| [TCAPV]"Image 29.czi_TICS"! | 3 | TCA | PV | Canaliculi | 2.1322E-12 | 1.46048E-8 | -- | 3.27E-7 | 8.67E-7 | 0.971 |
| [TCAPV]"Image 43.czi_TICS"! | 3 | TCA | PV | Canaliculi | 4.49304E-12 | 6.64152E-9 | -- | 3.27E-7 | 8.67E-7 | 0.975 |
| [TCAPV]"Image 30.czi_TICS"! | 4 | TCA | PV | Canaliculi | 1.92093E-12 | 1.44997E-8 | -- | 3.27E-7 | 8.67E-7 | 0.973 |
