## Supplementary material for "Intravital dynamic and correlative imaging reveals diffusion-dominated canalicular and flow-augmented ductular bile flux": Table S4

### Table S4: NMR spectra of custom synthesized cholic acid derivatives

#### NMR Spectra and HRMS data

#### Compound 5

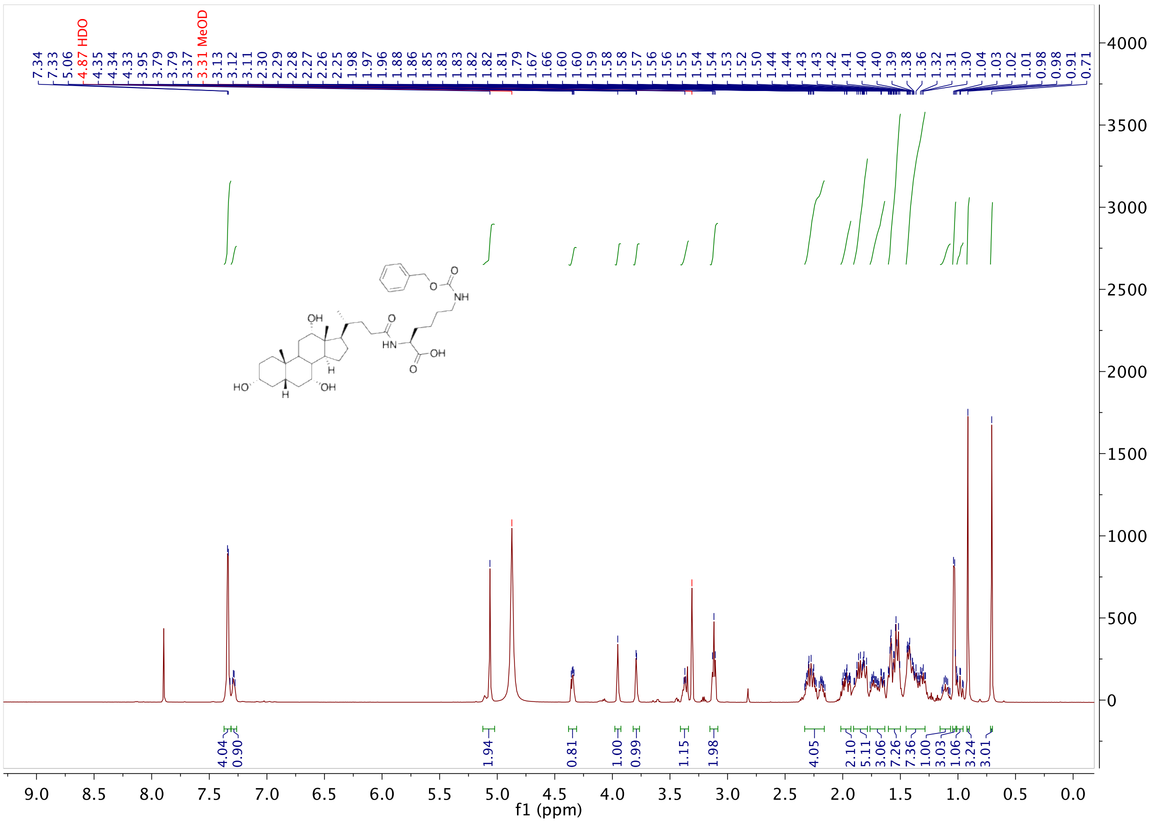


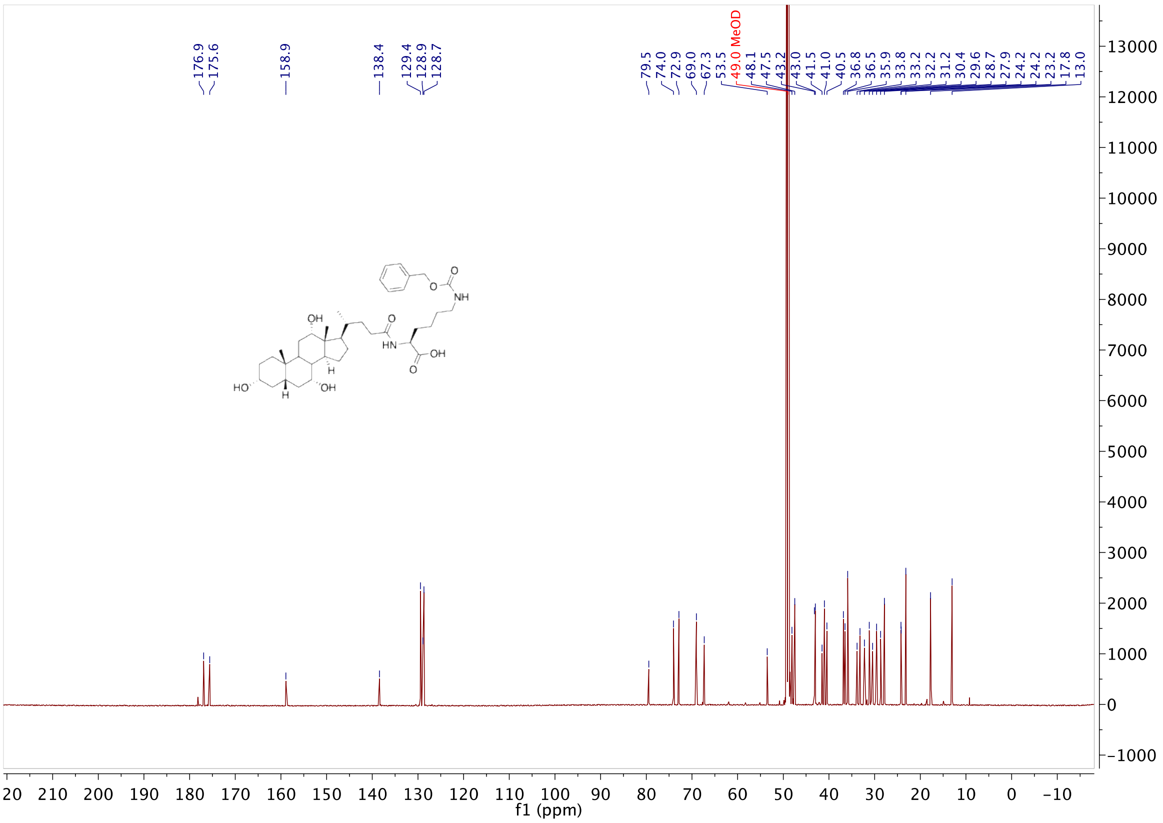

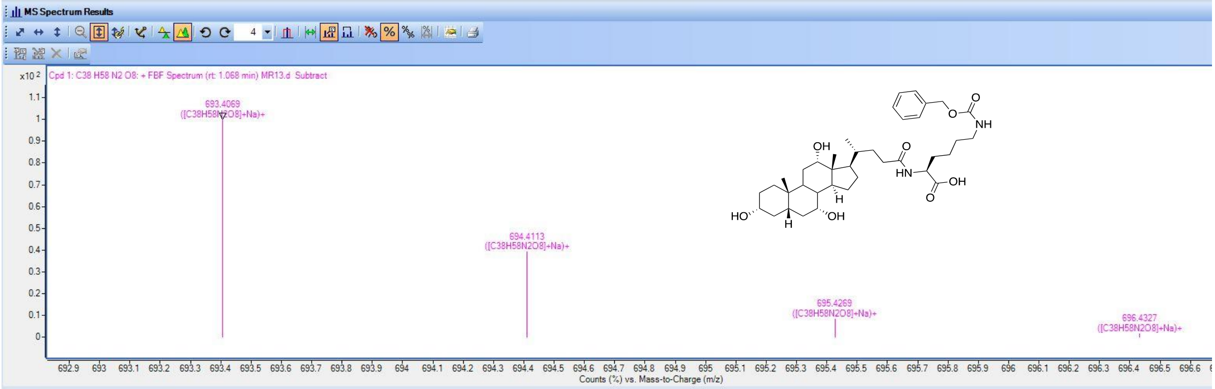


Compound List

| Label | Show/Hide | Cpd | File | ID Source | Name | Formula | Flags (Tgt) | Saturated | RT | RT (DB) |
|---|---|---|---|---|---|---|---|---|---|---|
| Cpd 1: C38 H58 N2 O8 | ☑ | 1 | MR13.d | FBF | | C38 H58 N2 O8 | | | 0.988 | |

| Best | ID Source | Name | Formula | Score | Score (RT) | RT Diff | Diff (ppm) | Score (Lib) | Score (DB) | Score (MFG) |
|---|---|---|---|---|---|---|---|---|---|---|
| ⊙ | FBF | | C38 H58 N2 O8 | 88.32 | | | -1.39 | | | |

| m/z | Species | Ion Formula | Score (MFG) | Score (MFG, MS/MS) | Score (MS) | Score (mass) | Score (iso. abund) | Score (iso. spacing) | Height |
|---|---|---|---|---|---|---|---|---|---|
| 693.4069 | (M+Na)+ | | | | 88.32 | 99.53 | 94.58 | 58.37 | 129706 |

| m/z | m/z (Calc) | Diff (ppm) | Height | Height (Calc) | Height % | Height % (Calc) | Height Sum% (Calc) | Height Sum % |
|---|---|---|---|---|---|---|---|---|
| 693.4069 | 693.4085 | 2.39 | 129706 | 124360.8 | 100 | 100 | 64.3 | 67 |
| 694.4113 | 694.4118 | 0.76 | 50779.2 | 53229.2 | 39.1 | 42.8 | 27.5 | 26.2 |
| 695.4269 | 695.4148 | -17.51 | 10786.3 | 13157.9 | 8.3 | 10.6 | 6.8 | 5.6 |
| 696.4327 | 696.4176 | -21.72 | 1988.2 | 2382.5 | 1.5 | 1.9 | 1.2 | 1 |
| 697.4313 | 697.4203 | -15.76 | 217.3 | 346.7 | 0.2 | 0.3 | 0.2 | 0.1 |

| m/z | Species | Ion Formula | Score (MFG) | Score (MFG, MS/MS) | Score (MS) | Score (mass) | Score (iso. abund) | Score (iso. spacing) | Height |
|---|---|---|---|---|---|---|---|---|---|
| 671.4263 | (M+H)+ | | | | 66.69 | 75.23 | 90.48 | 21.03 | 82100.5 |

#### Compound 6

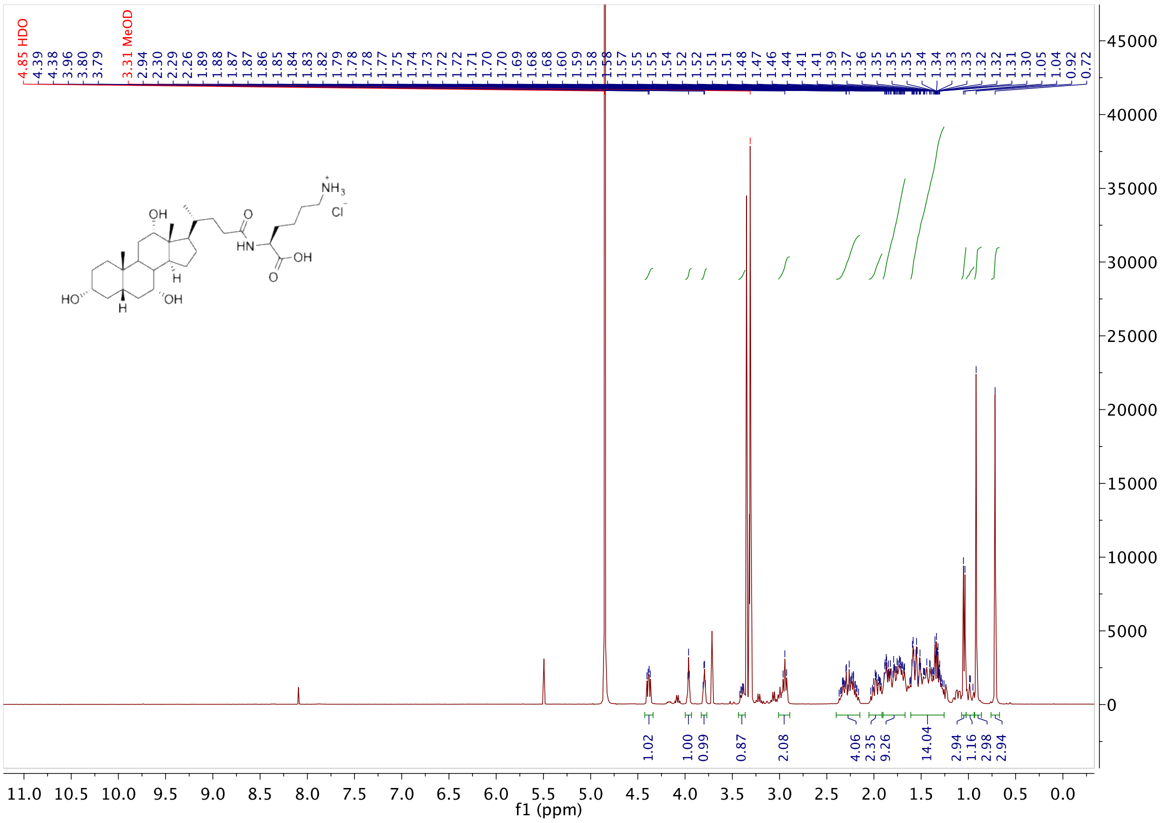

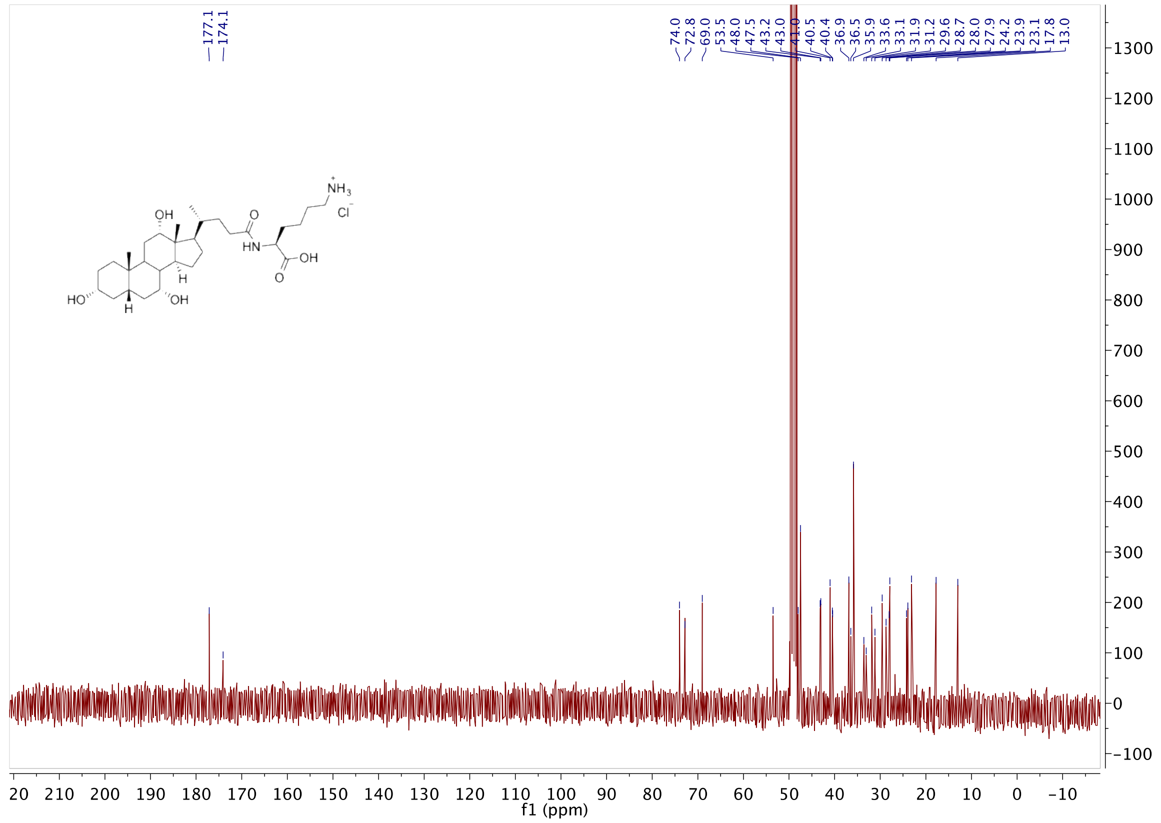


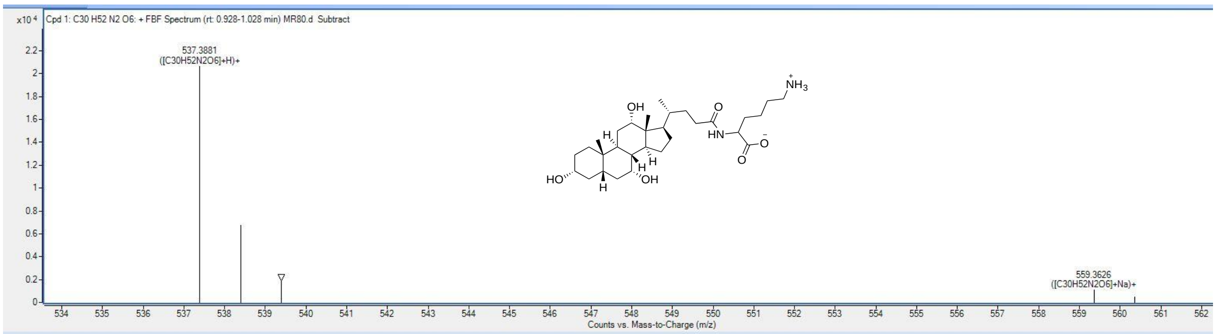


**Compound List**

Automatically Show Columns

| Label | Show/Hide | Cpd | File | ID Source | Name | Formula | Flags (Tgt) | Saturated | RT | RT (DB) | m/z |
|---|---|---|---|---|---|---|---|---|---|---|---|
| Cpd 1: C30 H52 N2 O6 | ☑ | 1 | MR80.d | FBF | | C30 H52 N2 O6 | | | 0.939 | | |

| Best | ID Source | Name | Formula | Score | Score (RT) | RT Diff | Diff (ppm) | Score (Lib) | Score (DB) | Score (MFG) | Num Sp |
|---|---|---|---|---|---|---|---|---|---|---|---|
| ⦿ | FBF | | C30 H52 N2 O6 | 92.04 | | | 4.01 | | | | |

| m/z | Species | Ion Formula | Score (MFG) | Score (MFG, MS/MS) | Score (MS) | Score (mass) | Score (iso. abund) | Score (iso. spacing) | Height |
|---|---|---|---|---|---|---|---|---|---|
| 537.3881 | (M+H)+ | | | | 92.04 | 88.88 | 92.24 | 98.11 | 20822.8 |

| m/z | m/z (Calc) | Diff (ppm) | Height | Height (Calc) | Height % | Height % (Calc) | Height Sum% (Calc) | Height Sum % |
|---|---|---|---|---|---|---|---|---|
| 537.3881 | 537.3898 | 3.12 | 20661.9 | 20822.8 | 100 | 100 | 71 | 70.4 |
| 538.3916 | 538.3931 | 2.81 | 6741.8 | 7083.1 | 32.6 | 34 | 24.1 | 23 |
| 539.3931 | 539.396 | 5.36 | 1926.8 | 1424.6 | 9.3 | 6.8 | 4.9 | 6.6 |

| m/z | Species | Ion Formula | Score (MFG) | Score (MFG, MS/MS) | Score (MS) | Score (mass) | Score (iso. abund) | Score (iso. spacing) | Height |
|---|---|---|---|---|---|---|---|---|---|
| 559.3626 | (M+Na)+ | | | | 17.55 | 1.78 | 40.47 | 21.57 | 1155.8 |

### Compound 1

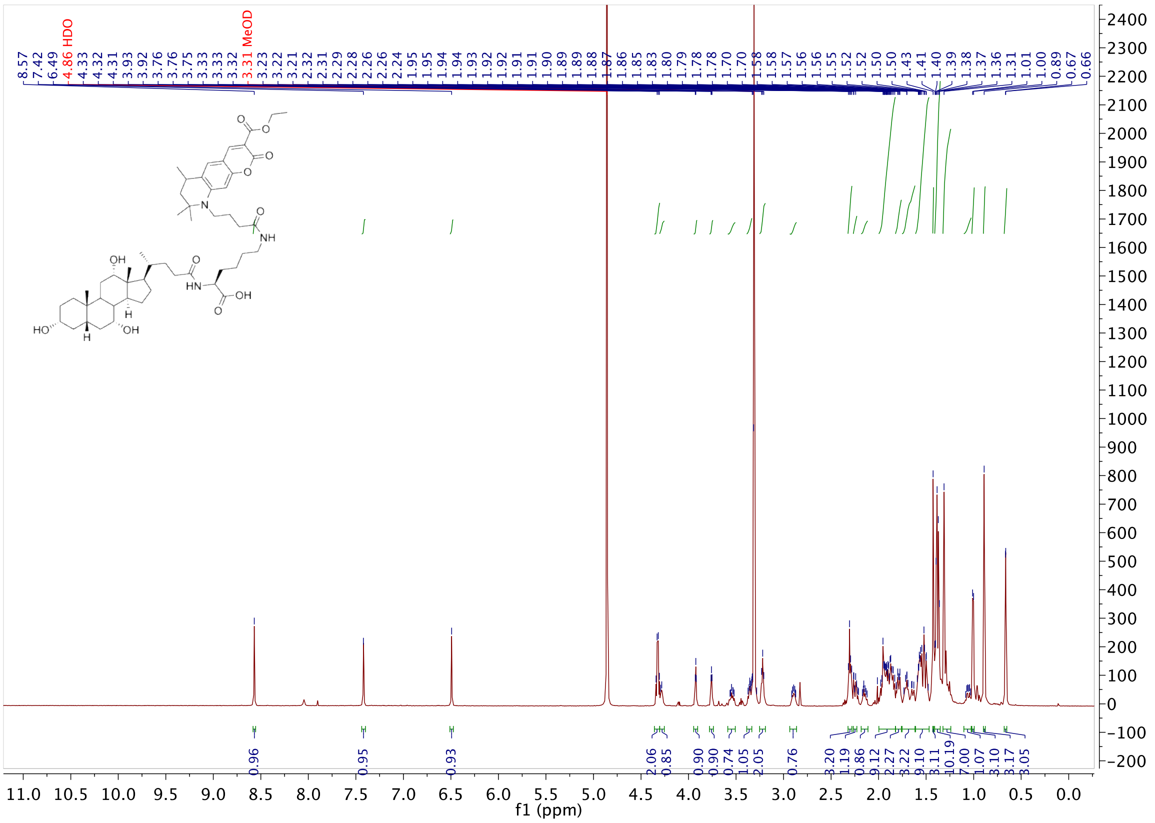
8.57
7.42
6.49
4.86 HDO
4.33
4.32
4.31
3.93
3.92
3.76
3.76
3.75
3.33
3.33
3.32
3.31 MeOD
3.23
3.22
3.21
2.32
2.31
2.29
2.28
2.26
2.26
2.24
1.95
1.95
1.94
1.94
1.93
1.92
1.92
1.91
1.91
1.90
1.89
1.89
1.88
1.87
1.86
1.85
1.83
1.80
1.79
1.78
1.78
1.70
1.70
1.58
1.58
1.57
1.56
1.56
1.55
1.52
1.52
1.50
1.50
1.43
1.41
1.40
1.39
1.38
1.37
1.36
1.31
1.01
1.00
0.89
0.67
0.66
OH
HN
OH
HO
H
H
OH
NH
N
O
0.96
0.95
0.93
2.06
0.85
0.90
0.90
0.74
1.05
2.05
0.76
3.20
1.19
0.86
9.12
2.27
3.22
9.10
3.11
10.19
7.00
1.07
3.10
3.17
3.05
f1 (ppm)

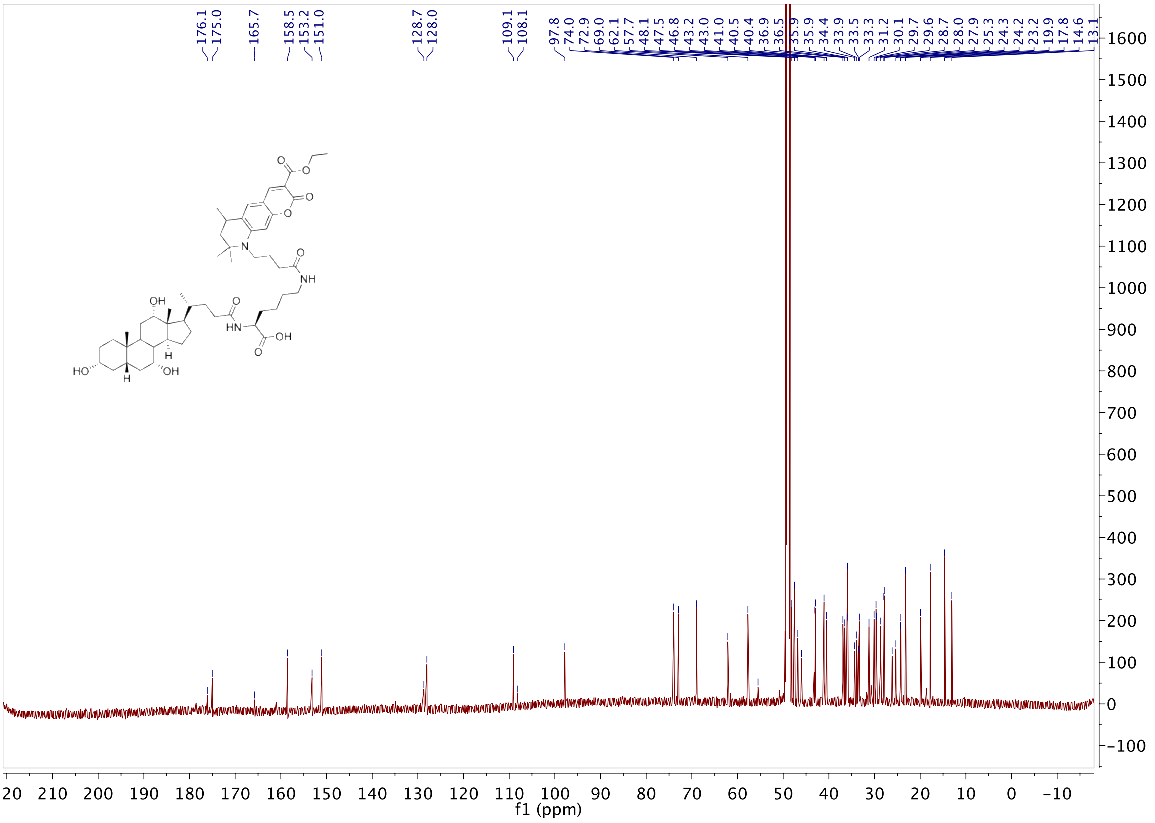
176.1
175.0
165.7
158.5
153.2
151.0
128.7
128.0
109.1
108.1
97.8
74.0
72.9
69.0
62.1
57.7
48.1
47.5
46.8
43.2
43.0
41.0
40.5
40.4
36.9
36.5
35.9
35.9
34.4
33.9
33.5
33.3
31.2
30.1
29.7
29.6
28.7
28.0
27.9
25.3
24.3
24.2
23.2
19.9
17.8
14.6
13.1
f1 (ppm)

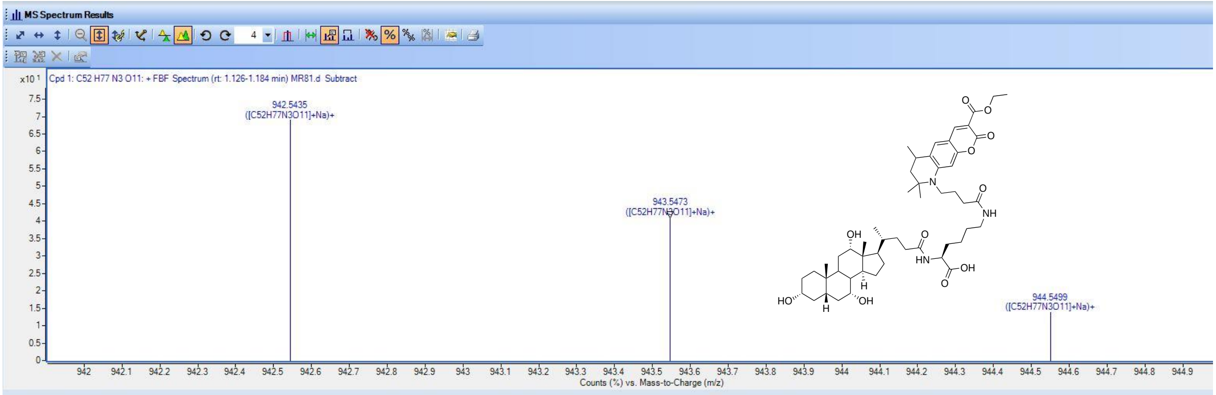


Compound List

| Label | Show/Hide | Cpd | File | ID Source | Name | Formula | Flags (Tgt) | Saturated | RT | RT (DB) | m/z |
|---|---|---|---|---|---|---|---|---|---|---|---|
| Cpd 1: C52 H77 N3 O11 | ☑ | 1 | MR81.d | FBF | | C52 H77 N3 O11 | | | 1.076 | | |

| Best | ID Source | Name | Formula | Score | Score (RT) | RT Diff | Diff (ppm) | Score (Lib) | Score (DB) | Score (MFG) | Num Spe |
|---|---|---|---|---|---|---|---|---|---|---|---|
| ⦿ | FBF | | C52 H77 N3 O11 | 98.02 | | | 1.51 | | | | |

| m/z | Species | Ion Formula | Score (MFG) | Score (MFG, MS/MS) | Score (MS) | Score (mass) | Score (iso. abund) | Score (iso. spacing) | Height |
|---|---|---|---|---|---|---|---|---|---|
| 942.5435 | (M+Na)+ | | | | 98.02 | 96.62 | 98.99 | 99.65 | 13349.7 |

| m/z | m/z (Calc) | Diff (ppm) | Height | Height (Calc) | Height % | Height % (Calc) | Height Sum% (Calc) | Height Sum % |
|---|---|---|---|---|---|---|---|---|
| 942.5435 | 942.545 | 1.6 | 13158.7 | 13349.7 | 100 | 100 | 54.9 | 54.1 |
| 943.5473 | 943.5483 | 1.1 | 7839.9 | 7828.6 | 59.6 | 58.6 | 32.2 | 32.2 |
| 944.5499 | 944.5513 | 1.49 | 2647.2 | 2556.3 | 20.1 | 19.1 | 10.5 | 10.9 |
| 945.5527 | 945.5542 | 1.58 | 690.7 | 601.9 | 5.2 | 4.5 | 2.5 | 2.8 |

| m/z | Species | Ion Formula | Score (MFG) | Score (MFG, MS/MS) | Score (MS) | Score (mass) | Score (iso. abund) | Score (iso. spacing) | Height |
|---|---|---|---|---|---|---|---|---|---|
| 920.5617 | (M+H)+ | | | | 97.01 | 96.17 | 95.96 | 99.98 | 19094.7 |

#### Compound 2

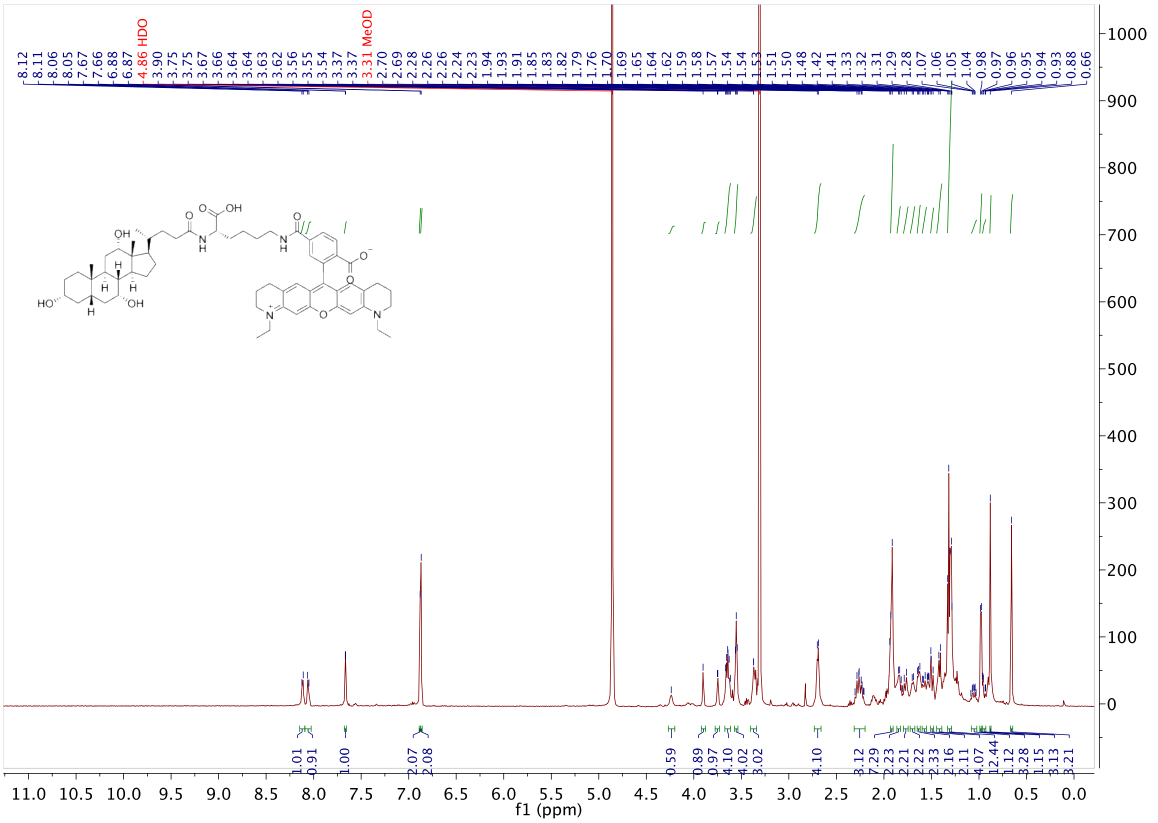

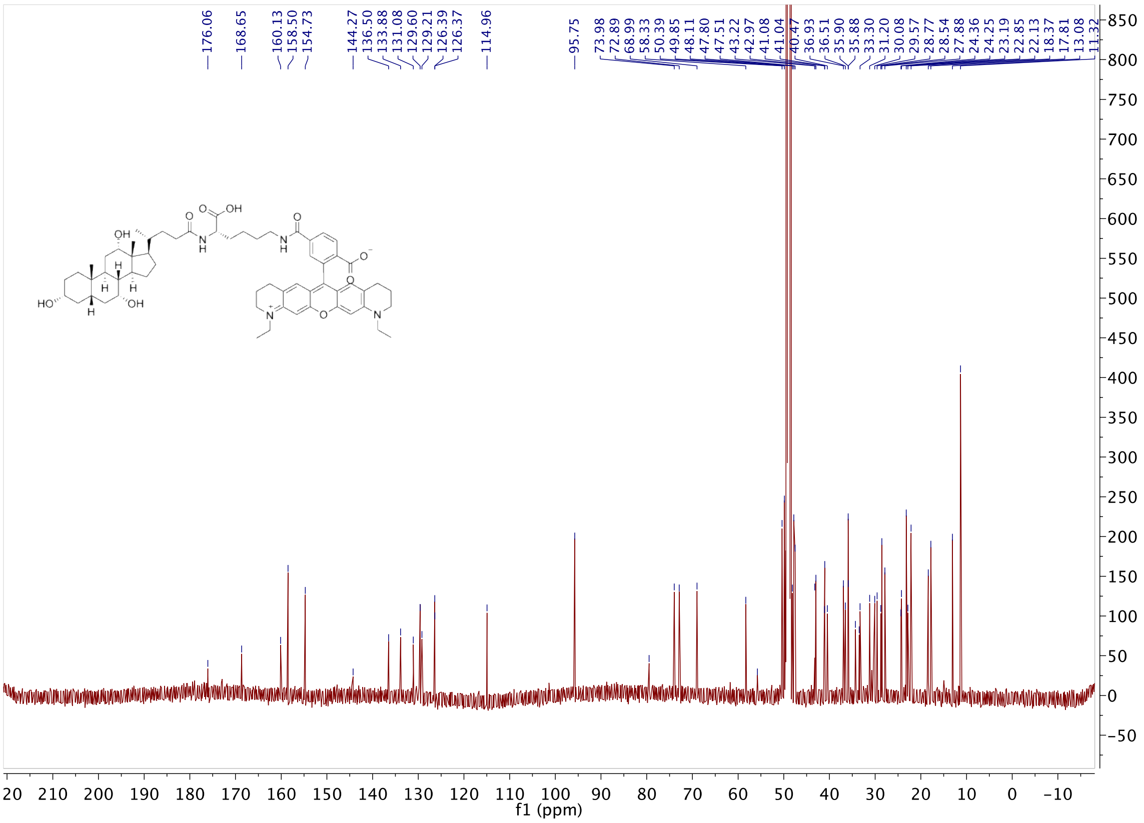


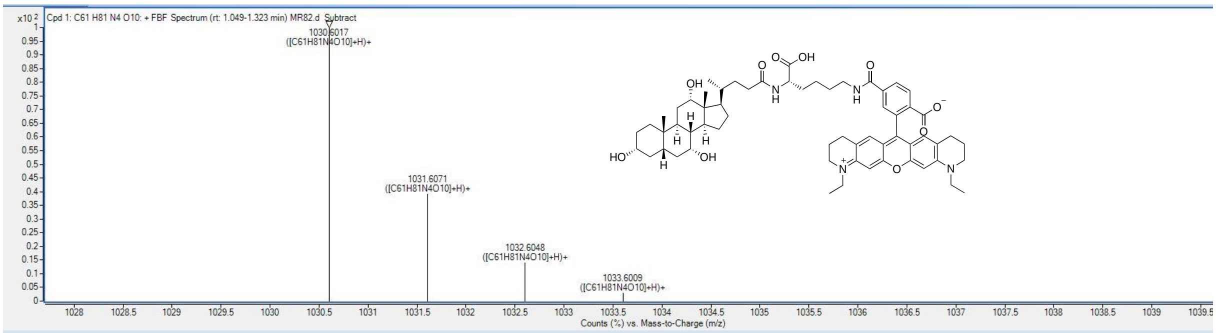


**Compound List**

Automatically Show Columns

| Label | Show/Hide | Cpd | File | ID Source | Name | Formula | Flags (Tgt) | Saturated | RT | RT (DB) | m/z |
|---|---|---|---|---|---|---|---|---|---|---|---|
| Cpd 1: C61 H81 N4 O10 | ☑ | 1 | MR82.d | FBF | | C61 H81 N4 O10 | low score | | 1.066 | | |

| Best | ID Source | Name | Formula | Score | Score (RT) | RT Diff | Diff (ppm) | Score (Lib) | Score (DB) | Score (MFG) | Num Sp |
|---|---|---|---|---|---|---|---|---|---|---|---|
| ⊙ | FBF | | C61 H81 N4 O10 | 73.36 | | | 2.01 | | | | |

| m/z | Species | Ion Formula | Score (MFG) | Score (MFG, MS/MS) | Score (MS) | Score (mass) | Score (iso. abund) | Score (iso. spacing) | Height |
|---|---|---|---|---|---|---|---|---|---|
| 1030.6017 | (M+H)+ | | | | 73.36 | 98.97 | 18.04 | 88.54 | 2588.7 |
| 1052.5802 | (M+Na)+ | | | | 5.82 | 6.78 | 0.06 | 10.82 | 245 |

| m/z | m/z (Calc) | Diff (ppm) | Height | Height (Calc) | Height % | Height % (Calc) | Height Sum% (Calc) | Height Sum % |
|---|---|---|---|---|---|---|---|---|
| 1030.6017 | 1030.6025 | 0.84 | 2588.7 | 2010.3 | 100 | 100 | 49.8 | 64.2 |
| 1031.6071 | 1031.6058 | -1.26 | 1010.2 | 1382.3 | 39 | 68.8 | 34.3 | 25 |
| 1032.6048 | 1032.6089 | 3.91 | 360.5 | 509.3 | 13.9 | 25.3 | 12.6 | 8.9 |
| 1033.6009 | 1033.6118 | 10.59 | 75 | 132.4 | 2.9 | 6.6 | 3.3 | 1.9 |

#### Compound 7

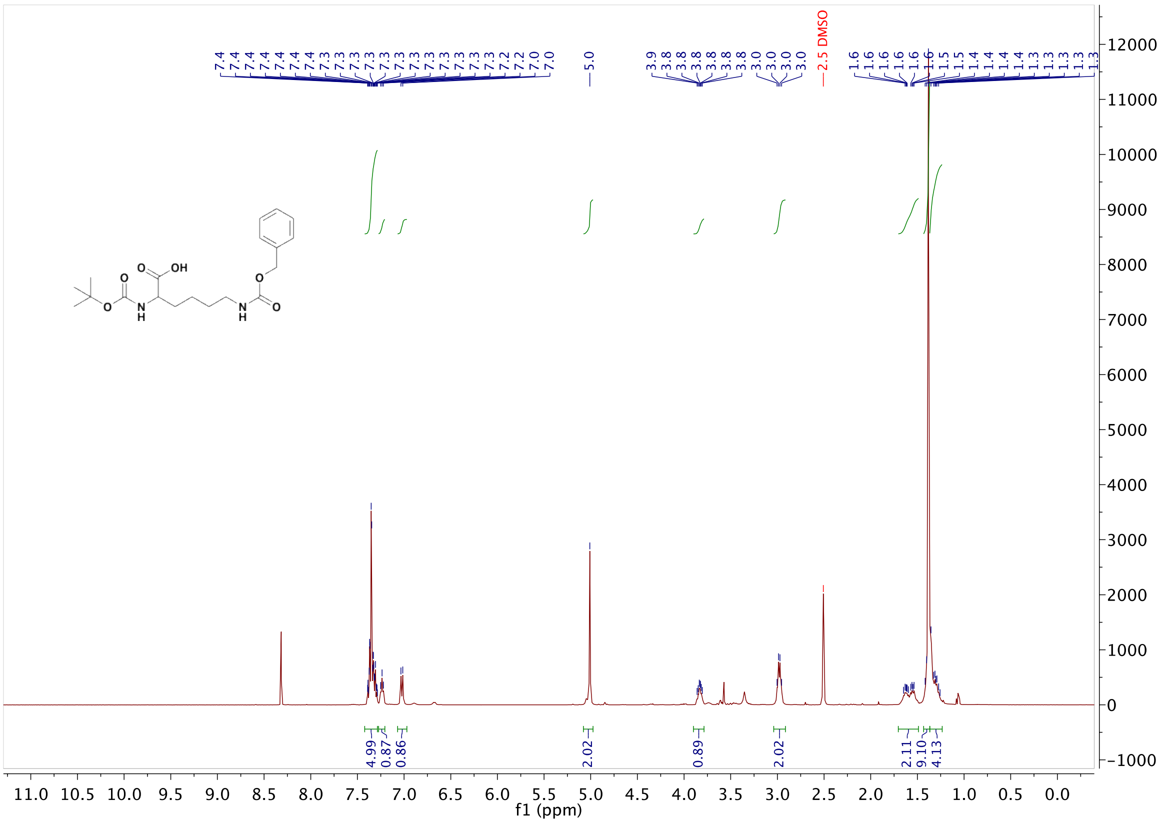

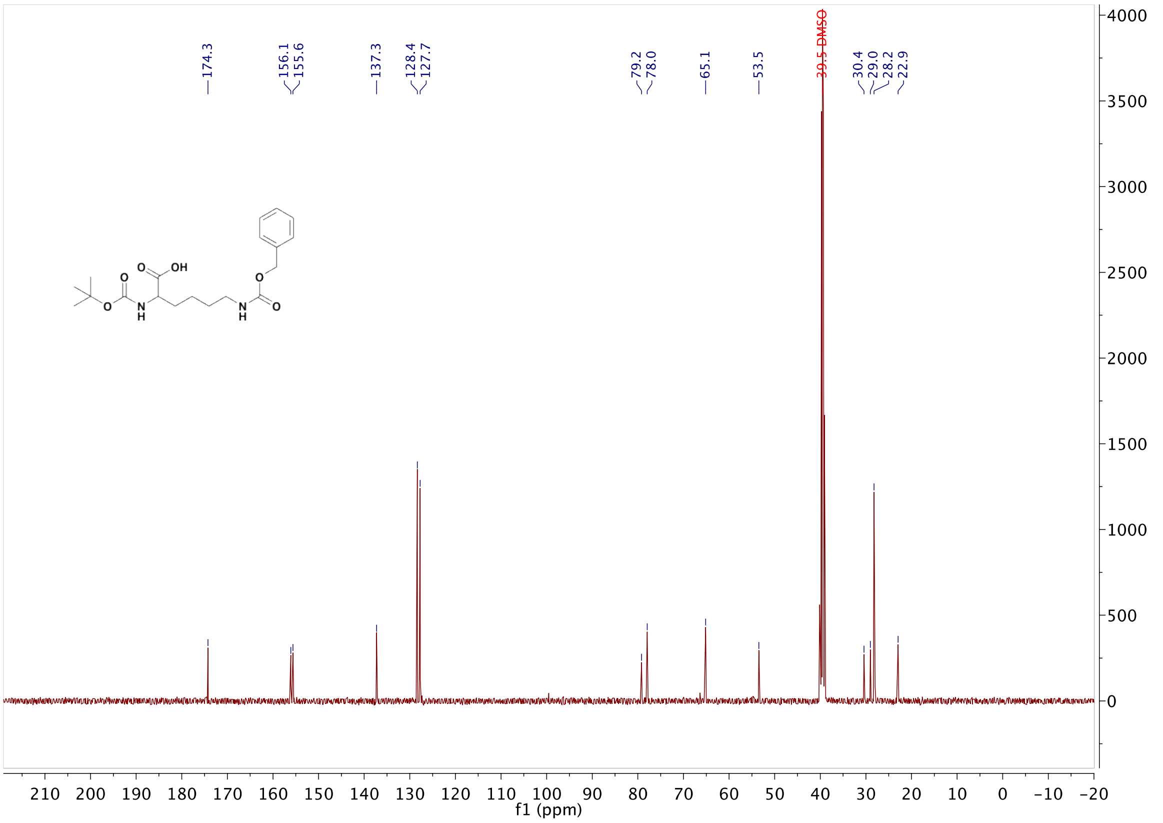

#### Compound 8

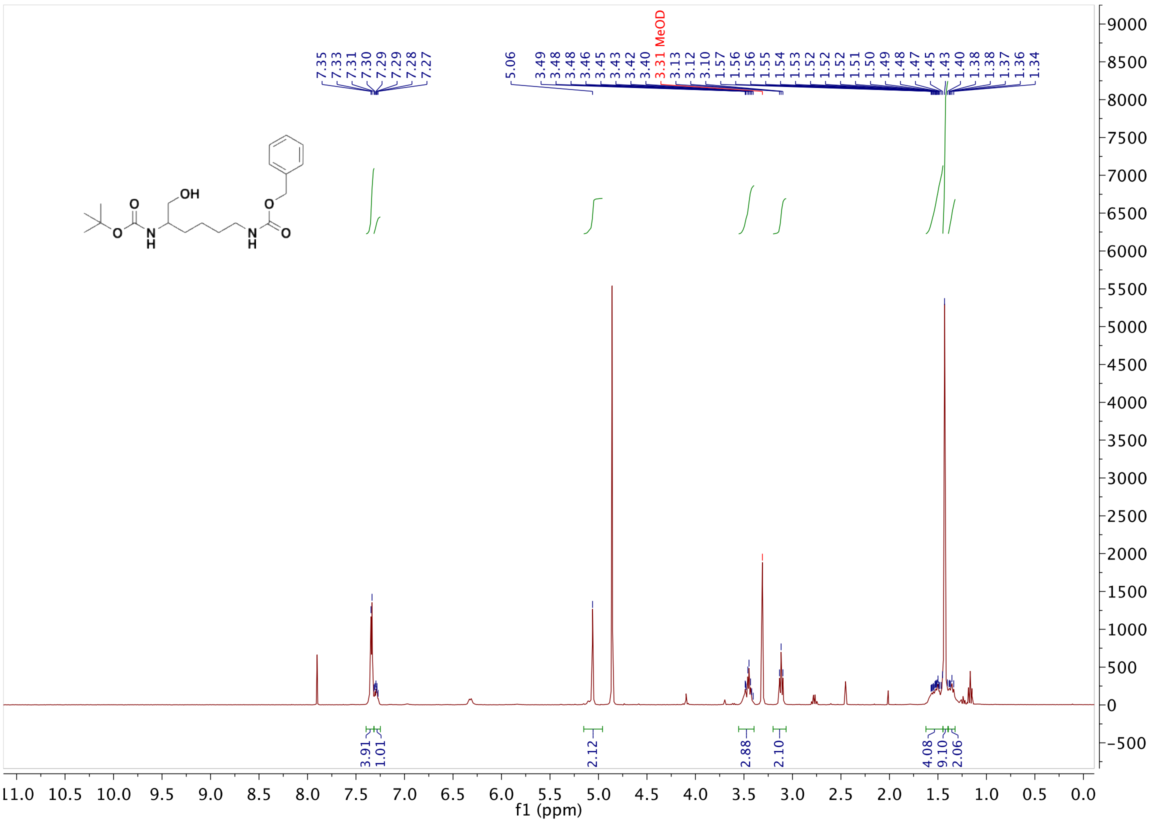

7.35 7.33 7.31 7.30 7.29 7.29 7.28 7.27
5.06
3.49 3.48 3.48 3.46 3.45 3.43 3.42 3.40
3.31 MeOD
3.13 3.12 3.10 1.57 1.56 1.56 1.55 1.54 1.53 1.52 1.52 1.52 1.51 1.50 1.49 1.48 1.47 1.45 1.43 1.40 1.38 1.38 1.37 1.36 1.34
OH
O
N
H
O
N
H
O
3.91 1.01 2.12 2.88 2.10 4.08 9.10 2.06
f1 (ppm)


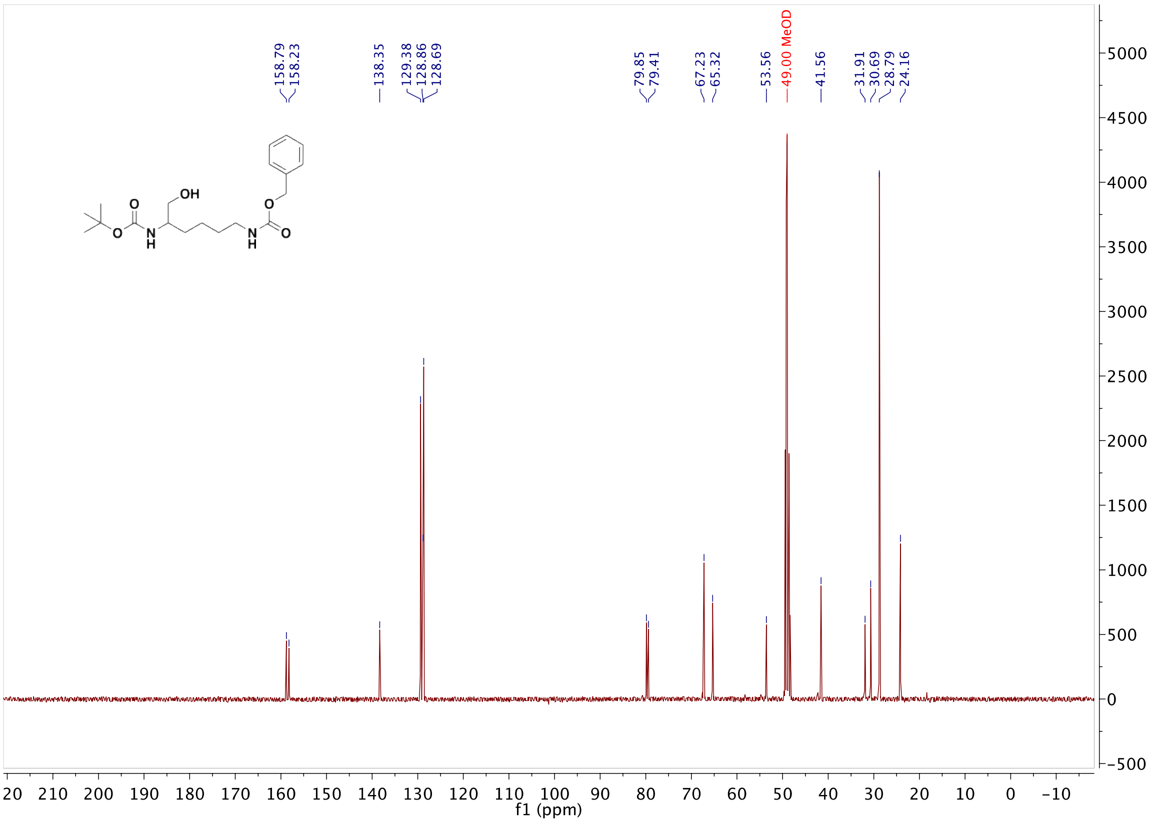

158.79 158.23
138.35
129.38 128.86 128.69
79.85 79.41
67.23 65.32
53.56
49.00 MeOD
41.56
31.91 30.69 28.79 24.16
OH
O
N
H
O
N
H
O
f1 (ppm)

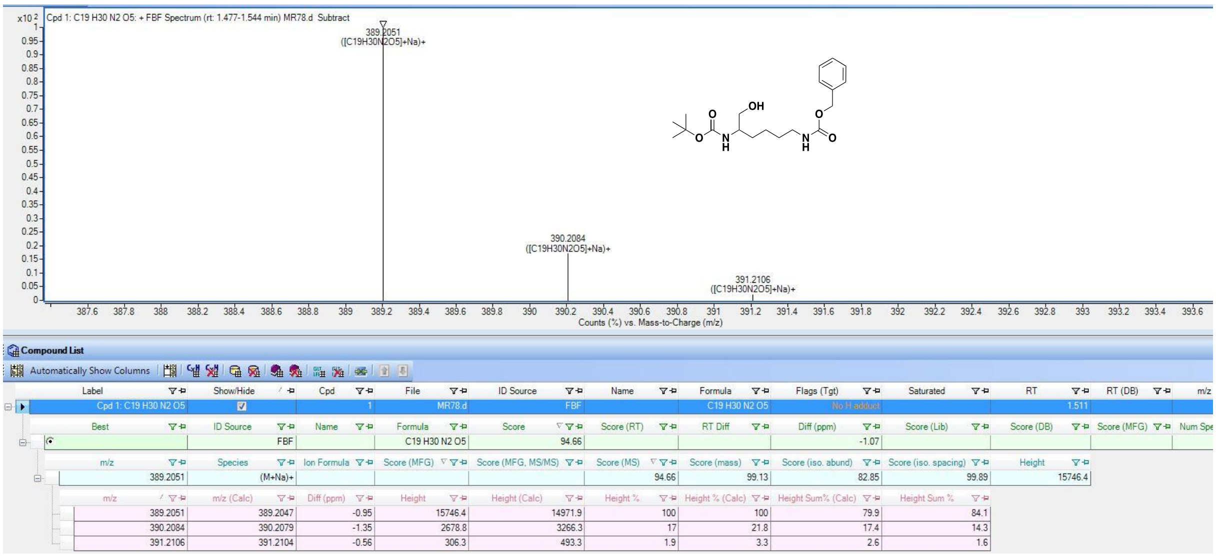
Cpd 1: C19 H30 N2 O5: + FBF Spectrum (rt: 1.477-1.544 min) MR78.d Subtract
389.2051
([C19H30N2O5]+Na)+
390.2084
([C19H30N2O5]+Na)+
391.2106
([C19H30N2O5]+Na)+
Counts (%) vs. Mass-to-Charge (m/z)
OH
O
N
H
O
Compound List
Automatically Show Columns
Cpd 1: C19 H30 N2 O5
MR78.d
FBF
C19 H30 N2 O5
No H adduct
1.511
C19 H30 N2 O5
94.66
-1.07
389.2051
(M+Na)+
94.66
99.13
82.85
99.89
15746.4
389.2051 389.2047 -0.95 15746.4 14971.9 100 100 79.9 84.1
390.2084 390.2079 -1.35 2678.8 3266.3 17 21.8 17.4 14.3
391.2106 391.2104 -0.56 306.3 493.3 1.9 3.3 2.6 1.6

#### Compound 9

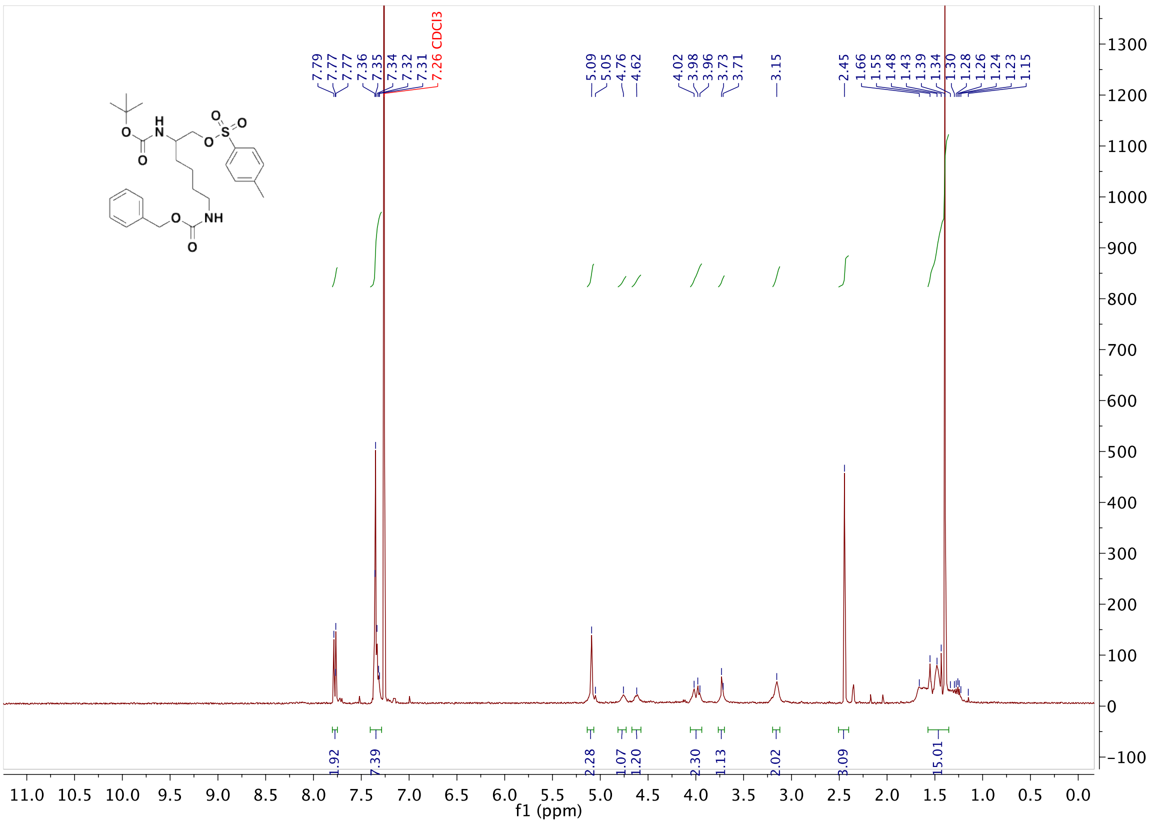
7.79
7.77
7.36
7.35
7.34
7.32
7.31
7.26 CDCl3
5.09
5.05
4.76
4.62
4.02
3.98
3.96
3.73
3.71
3.15
2.45
1.66
1.55
1.48
1.43
1.39
1.34
1.30
1.28
1.26
1.24
1.23
1.15
1.92
7.39
2.28
1.07
1.20
2.30
1.13
2.02
3.09
15.01
f1 (ppm)
H
N
O
O
S
O
O
NH
O
O

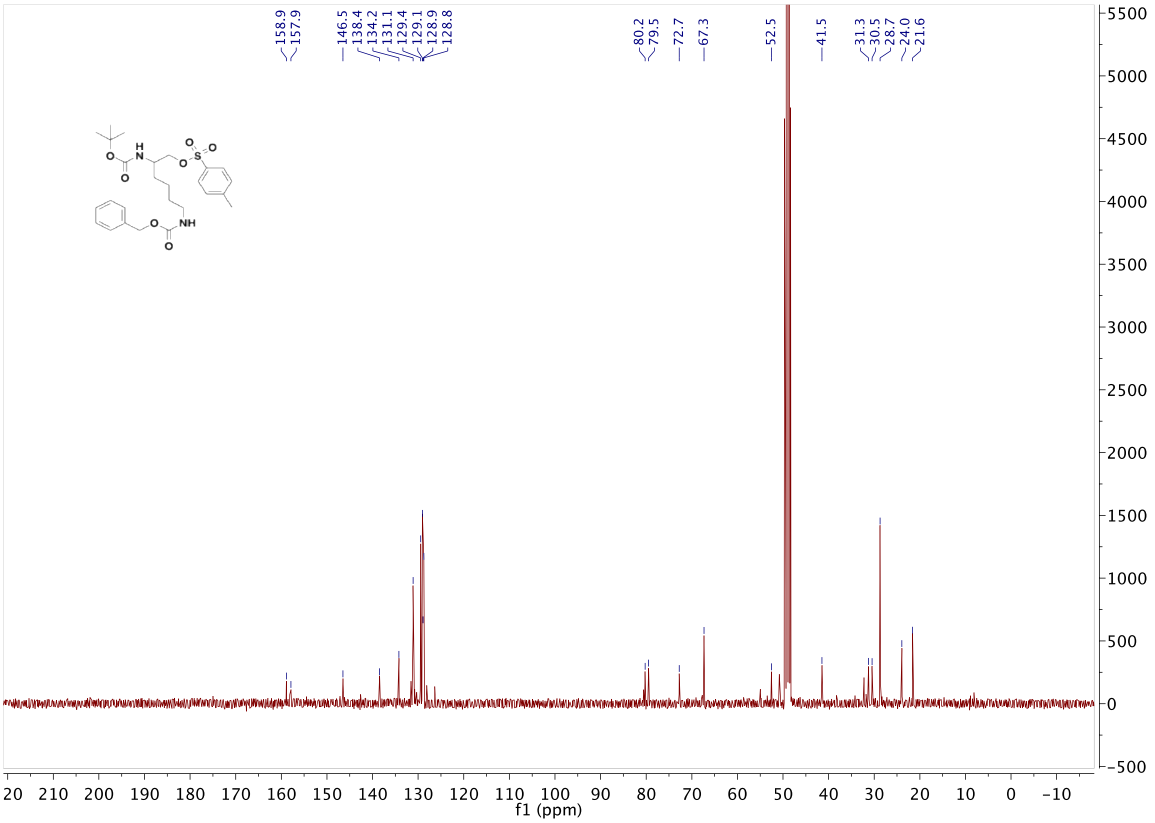


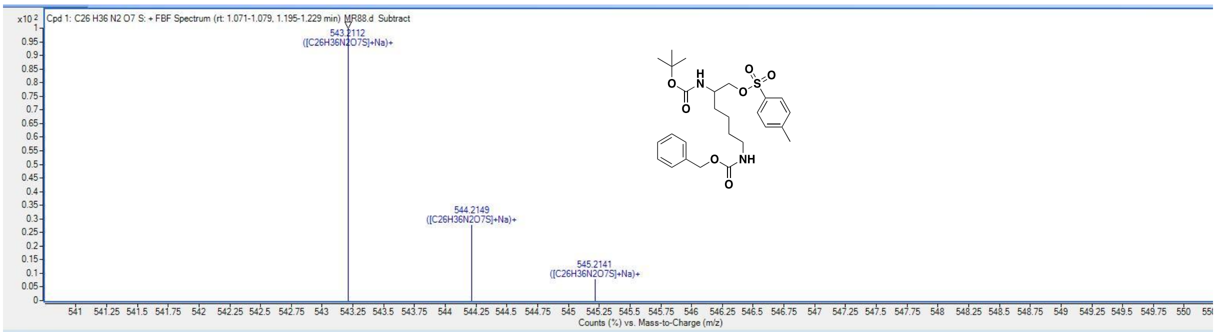


**Compound List**

Automatically Show Columns

| Label | Show/Hide | Cpd | File | ID Source | Name | Formula | Flags (Tgt) | Saturated | RT | RT (DB) | m/z |
|---|---|---|---|---|---|---|---|---|---|---|---|
| Cpd 1: C26 H36 N2 O7 S | ☑ | 1 | MR88.d | FBF | | C26 H36 N2 O7 S | No H adduct | | 1.096 | | |

| Best | ID Source | Name | Formula | Score | Score (RT) | RT Diff | Diff (ppm) | Score (Lib) | Score (DB) | Score (MFG) | Num Spe |
|---|---|---|---|---|---|---|---|---|---|---|---|
| ⊙ | FBF | | C26 H36 N2 O7 S | 88.92 | | | 4.12 | | | | |

| m/z | Species | Ion Formula | Score (MFG) | Score (MFG, MS/MS) | Score (MS) | Score (mass) | Score (iso. abund) | Score (iso. spacing) | Height |
|---|---|---|---|---|---|---|---|---|---|
| 543.2112 | (M+Na)+ | | | | 88.92 | 83.45 | 91.02 | 97.37 | 42427.1 |

| m/z | m/z (Calc) | Diff (ppm) | Height | Height (Calc) | Height % | Height % (Calc) | Height Sum% (Calc) | Height Sum % |
|---|---|---|---|---|---|---|---|---|
| 543.2112 | 543.2135 | 4.31 | 42427.1 | 40842.7 | 100 | 100 | 71.1 | 73.8 |
| 544.2149 | 544.2167 | 3.26 | 11749.4 | 12384.2 | 27.7 | 30.3 | 21.6 | 20.4 |
| 545.2141 | 545.215 | 1.74 | 3279 | 4228.5 | 7.7 | 10.4 | 7.4 | 5.7 |

**Compound 10**

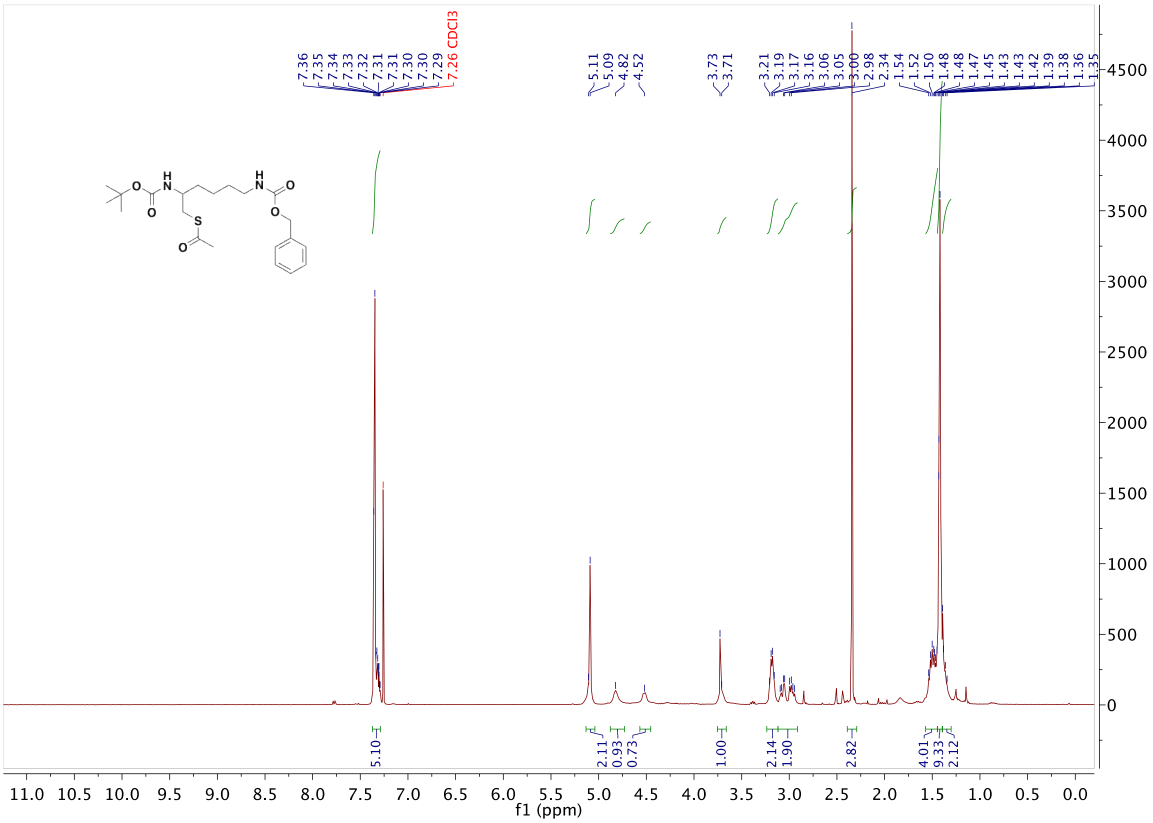
7.36
7.35
7.34
7.33
7.32
7.31
7.31
7.30
7.30
7.29
7.26 CDCl3
5.11
5.09
4.82
4.52
3.73
3.71
3.21
3.19
3.17
3.16
3.06
3.05
3.00
2.98
2.34
1.54
1.52
1.50
1.48
1.48
1.47
1.45
1.43
1.43
1.42
1.39
1.38
1.36
1.35
5.10
2.11
0.93
0.73
1.00
2.14
1.90
2.82
4.01
9.33
2.12
f1 (ppm)

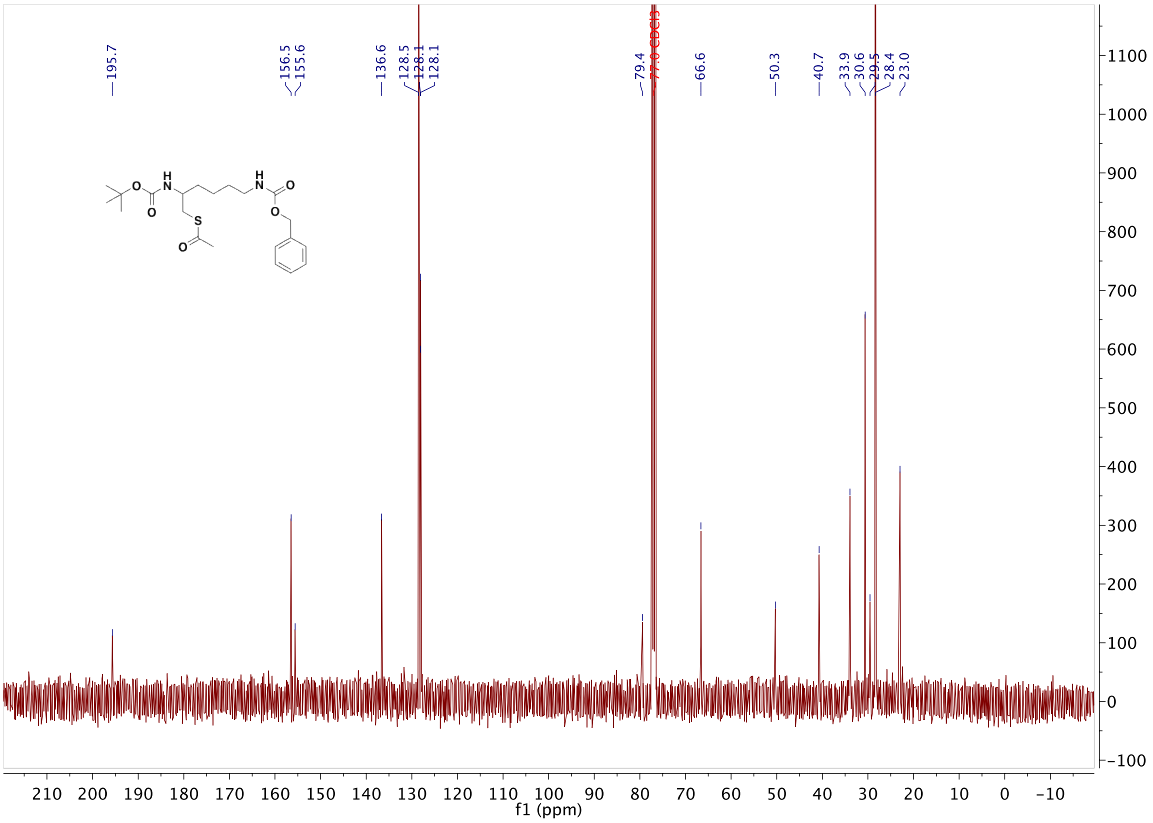
195.7
156.5
155.6
136.6
128.5
128.1
128.1
79.4
77.0 CDCl3
66.6
50.3
40.7
33.9
30.6
29.5
28.4
23.0
f1 (ppm)

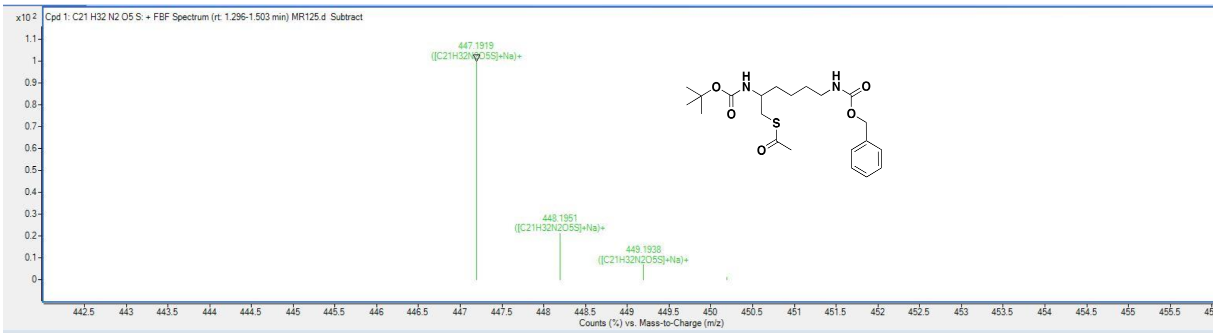


Compound List

| Label | Show/Hide | Cpd | File | ID Source | Name | Formula | Flags (Tgt) | Saturated | RT | RT (DB) |
|---|---|---|---|---|---|---|---|---|---|---|
| Cpd 1: C21 H32 N2 O5 S | ☑ | 1 | MR125.d | FBF | | C21 H32 N2 O5 S | | | 1.304 | |

| Best | ID Source | Name | Formula | Score | Score (RT) | RT Diff | Diff (ppm) | Score (Lib) | Score (DB) | Score (MFG) |
|---|---|---|---|---|---|---|---|---|---|---|
| ⦿ | FBF | | C21 H32 N2 O5 S | 96.6 | | | 0.85 | | | |

| m/z | Species | Ion Formula | Score (MFG) | Score (MFG, MS/MS) | Score (MS) | Score (mass) | Score (iso. abund) | Score (iso. spacing) | Height |
|---|---|---|---|---|---|---|---|---|---|
| 447.1919 | (M+Na)+ | | | | 96.6 | 99.3 | 90.63 | 98.37 | 40098 |

| m/z | m/z (Calc) | Diff (ppm) | Height | Height (Calc) | Height % | Height % (Calc) | Height Sum% (Calc) | Height Sum % |
|---|---|---|---|---|---|---|---|---|
| 447.1919 | 447.1924 | 1.08 | 40098 | 38305.4 | 100 | 100 | 74.2 | 77.7 |
| 448.1951 | 448.1955 | 0.98 | 8427.1 | 9496.6 | 21 | 24.8 | 18.4 | 16.3 |
| 449.1938 | 449.1929 | -2.16 | 2648 | 3235.8 | 6.6 | 8.4 | 6.3 | 5.1 |
| 450.1965 | 450.1943 | -5.06 | 459 | 594.3 | 1.1 | 1.6 | 1.2 | 0.9 |

| m/z | Species | Ion Formula | Score (MFG) | Score (MFG, MS/MS) | Score (MS) | Score (mass) | Score (iso. abund) | Score (iso. spacing) | Height |
|---|---|---|---|---|---|---|---|---|---|
| 425.2174 | (M+H)+ | | | | 4.01 | 8.42 | 0 | 0 | 139.5 |

#### Compound 11

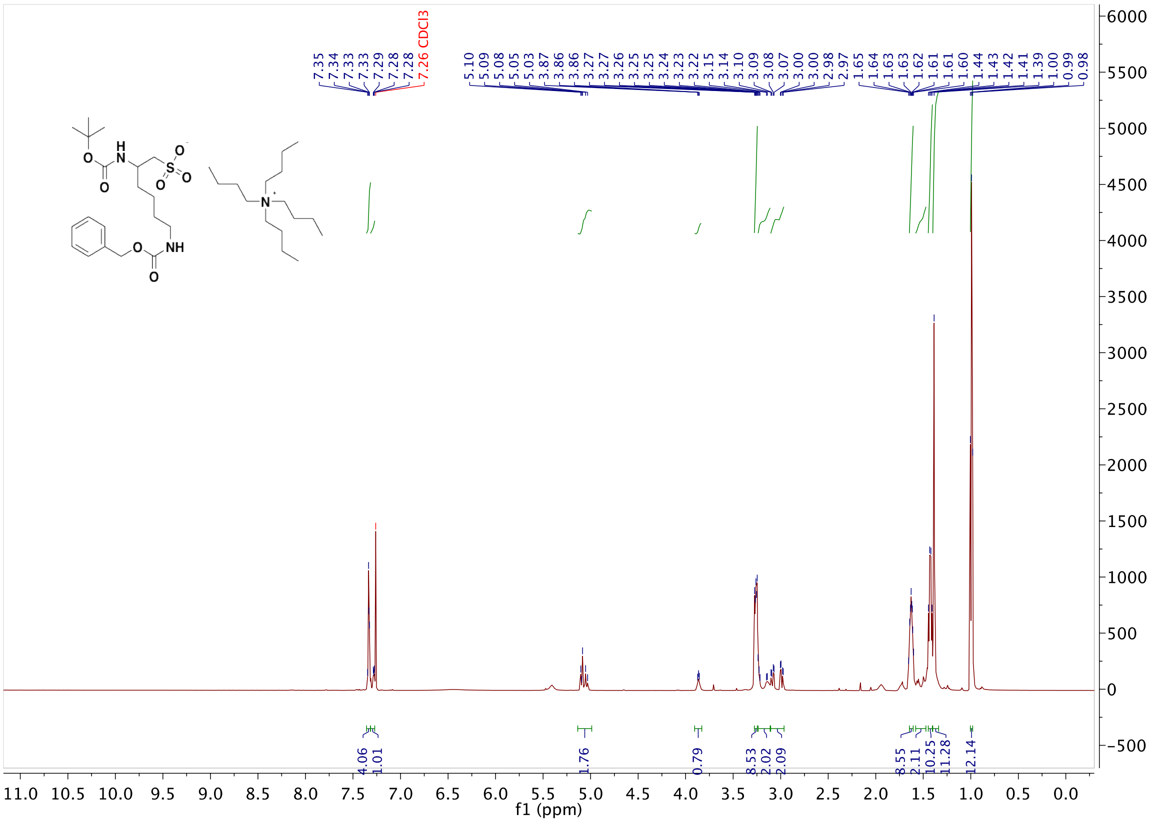

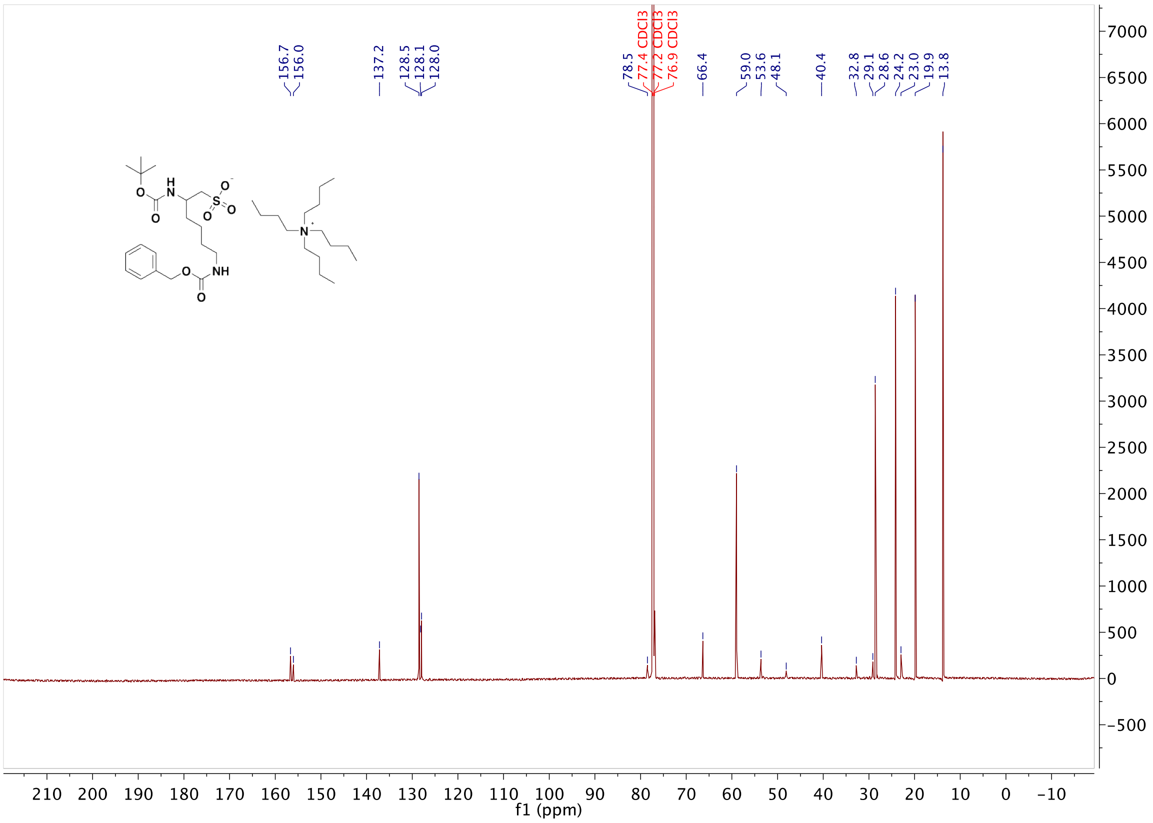


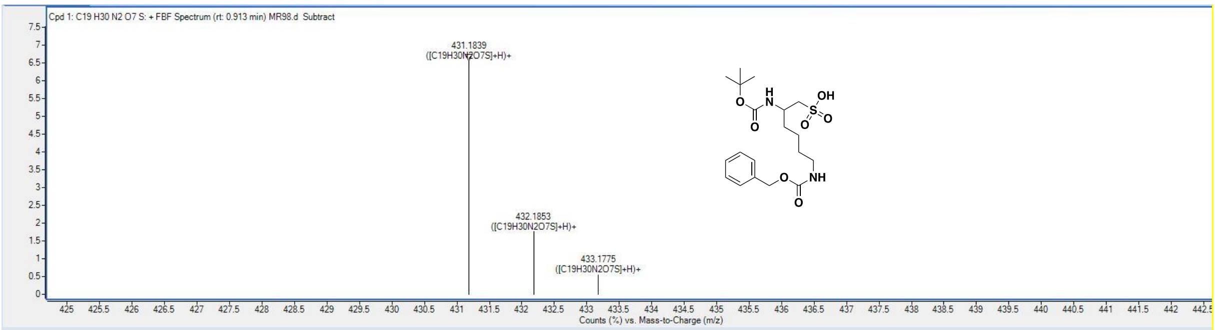


**Compound List**

Automatically Show Columns

| Label | Show/Hide | Cpd | File | ID Source | Name | Formula | Flags (Tgt) | Saturated | RT | RT (DB) | m/z |
|---|---|---|---|---|---|---|---|---|---|---|---|
| Cpd 1: C19 H30 N2 O7 S | ☑ | 1 | MR98.d | FBF | | C19 H30 N2 O7 S | | | 0.847 | | |

| Best | ID Source | Name | Formula | Score | Score (RT) | RT Diff | Diff (ppm) | Score (Lib) | Score (DB) | Score (MFG) | Num Spe |
|---|---|---|---|---|---|---|---|---|---|---|---|
| ⊙ | FBF | | C19 H30 N2 O7 S | 80.92 | | | 3.78 | | | | |

| m/z | Species | Ion Formula | Score (MFG) | Score (MFG, MS/MS) | Score (MS) | Score (mass) | Score (iso. abund) | Score (iso. spacing) | Height |
|---|---|---|---|---|---|---|---|---|---|
| 431.1839 | (M+H)+ | | | | 80.92 | 88.84 | 85.97 | 58.99 | 11166.9 |

| m/z | m/z (Calc) | Diff (ppm) | Height | Height (Calc) | Height % | Height % (Calc) | Height Sum% (Calc) | Height Sum % |
|---|---|---|---|---|---|---|---|---|
| 431.1839 | 431.1846 | 1.79 | 10812.9 | 11166.9 | 100 | 100 | 76.3 | 73.9 |
| 432.1853 | 432.1877 | 5.59 | 2900.4 | 2534.1 | 26.8 | 22.7 | 17.3 | 19.8 |
| 433.1775 | 433.1849 | 17.19 | 922.6 | 934.9 | 8.5 | 8.4 | 6.4 | 6.3 |

| m/z | Species | Ion Formula | Score (MFG) | Score (MFG, MS/MS) | Score (MS) | Score (mass) | Score (iso. abund) | Score (iso. spacing) | Height |
|---|---|---|---|---|---|---|---|---|---|
| 453.1643 | (M+Na)+ | | | | 79.36 | 87.82 | 76.22 | 66.22 | 164708 |

**Compound 12**

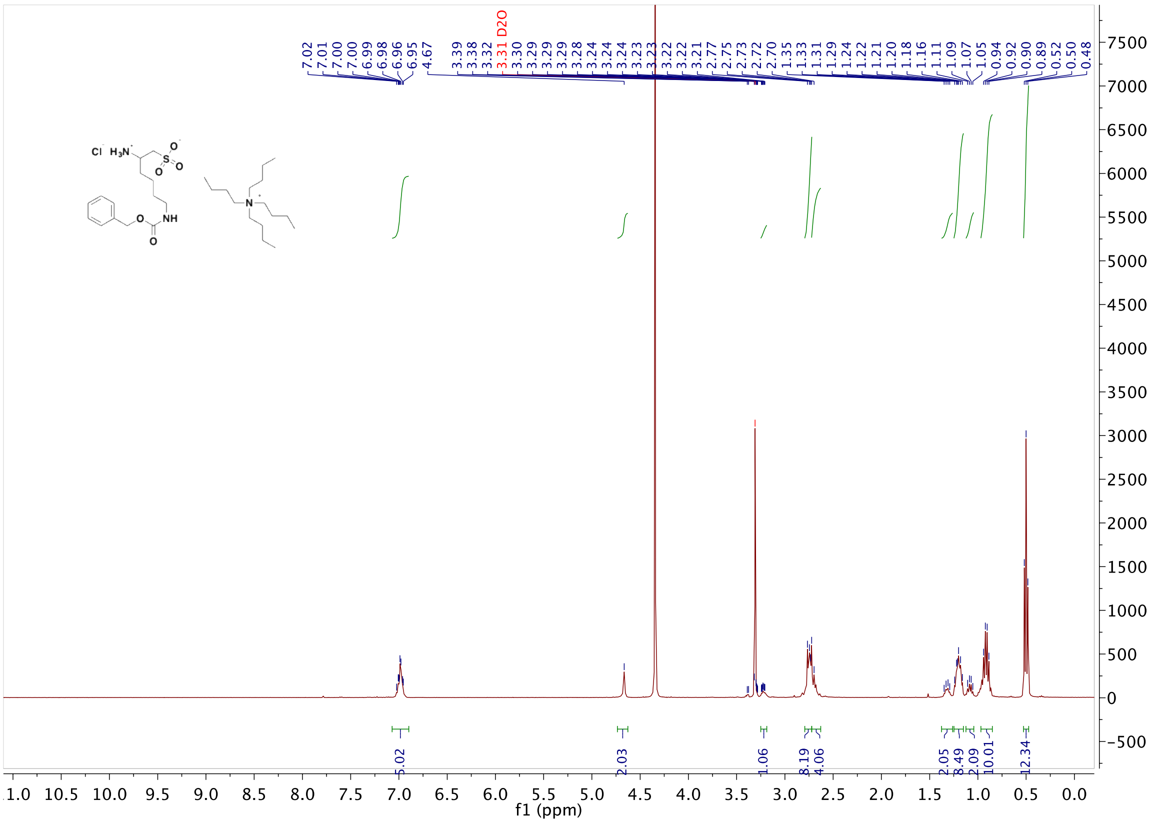

7.02
7.01
7.00
7.00
6.99
6.98
6.96
6.95
4.67
3.39
3.38
3.32
3.31 D2O
3.30
3.29
3.29
3.29
3.28
3.24
3.24
3.24
3.23
3.23
3.22
3.22
3.21
2.77
2.75
2.73
2.72
2.70
1.35
1.33
1.31
1.29
1.24
1.22
1.21
1.20
1.18
1.16
1.11
1.09
1.07
1.05
0.94
0.92
0.90
0.89
0.52
0.50
0.48
5.02
2.03
1.06
8.19
4.06
2.05
8.49
2.09
10.01
12.34
f1 (ppm)


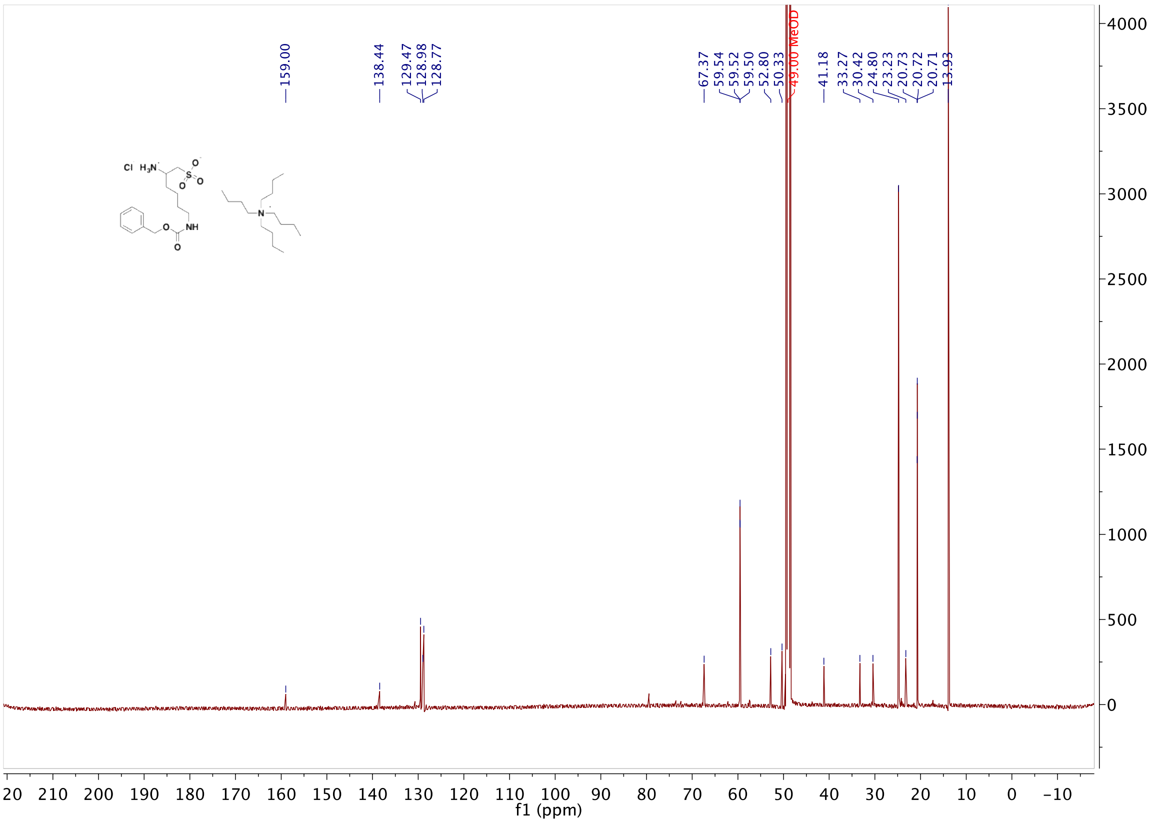

159.00
138.44
129.47
128.98
128.77
67.37
59.54
59.52
59.50
52.80
50.33
49.00 MeOD
41.18
33.27
30.42
24.80
23.23
20.73
20.72
20.71
13.93
f1 (ppm)

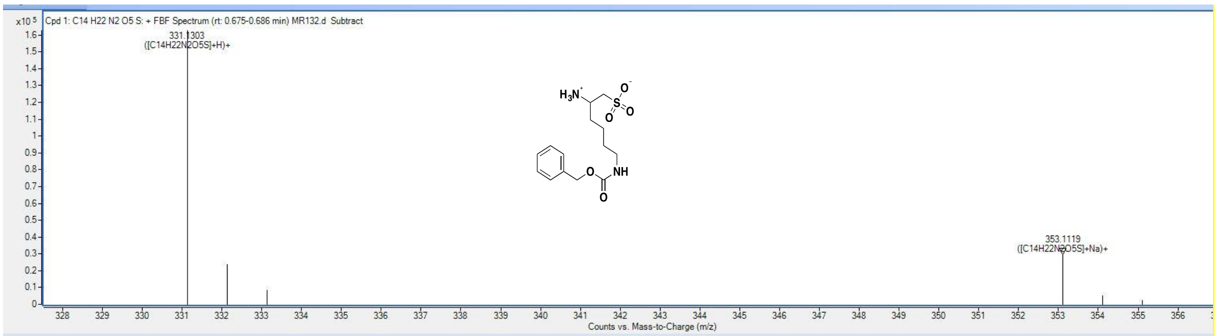


Compound List

Automatically Show Columns

| Label | Show/Hide | Cpd | File | ID Source | Name | Formula | Flags (Tgt) | Saturated | RT | RT (DB) | m/z |
|---|---|---|---|---|---|---|---|---|---|---|---|
| Cpd 1: C14 H22 N2 O5 S | ☑ | 1 | MR132.d | FBF | | C14 H22 N2 O5 S | | | 0.631 | | |

| Best | ID Source | Name | Formula | Score | Score (RT) | RT Diff | Diff (ppm) | Score (Lib) | Score (DB) | Score (MFG) | Num Sp |
|---|---|---|---|---|---|---|---|---|---|---|---|
| ⊙ | FBF | | C14 H22 N2 O5 S | 87.4 | | | 5.88 | | | | |

| m/z | Species | Ion Formula | Score (MFG) | Score (MFG, MS/MS) | Score (MS) | Score (mass) | Score (iso. abund) | Score (iso. spacing) | Height |
|---|---|---|---|---|---|---|---|---|---|
| 331.1303 | (M+H)+ | | | | 87.4 | 78.54 | 92.48 | 99.01 | 162638.7 |

| m/z | m/z (Calc) | Diff (ppm) | Height | Height (Calc) | Height % | Height % (Calc) | Height Sum% (Calc) | Height Sum % |
|---|---|---|---|---|---|---|---|---|
| 331.1303 | 331.1322 | 5.82 | 162638.7 | 157011.7 | 100 | 100 | 80.6 | 83.5 |
| 332.1331 | 332.1352 | 6.21 | 23745.4 | 26876 | 14.6 | 17.1 | 13.8 | 12.2 |
| 333.1306 | 333.1313 | 2.23 | 8306.5 | 10802.8 | 5.1 | 6.9 | 5.5 | 4.3 |

| m/z | Species | Ion Formula | Score (MFG) | Score (MFG, MS/MS) | Score (MS) | Score (mass) | Score (iso. abund) | Score (iso. spacing) | Height |
|---|---|---|---|---|---|---|---|---|---|
| 353.1119 | (M+Na)+ | | | | 86.85 | 73.93 | 98.66 | 98.51 | 29777.2 |

### Compound 13

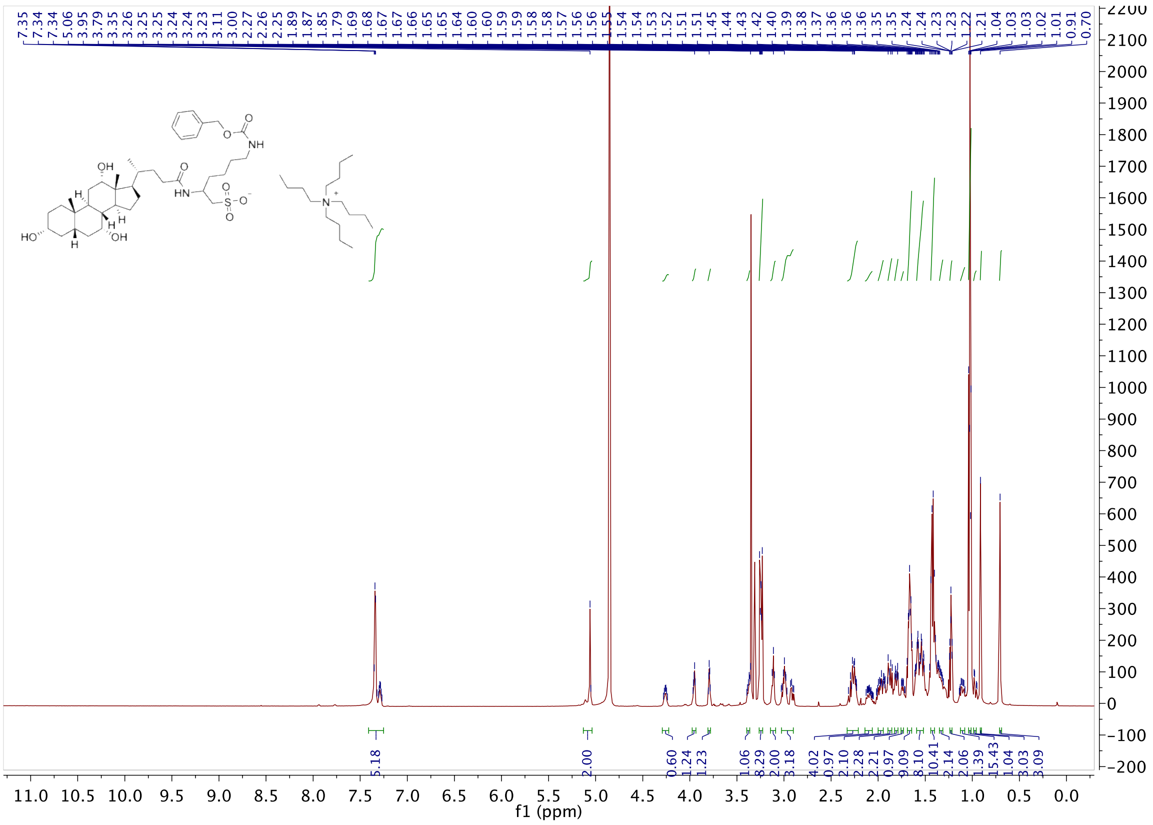

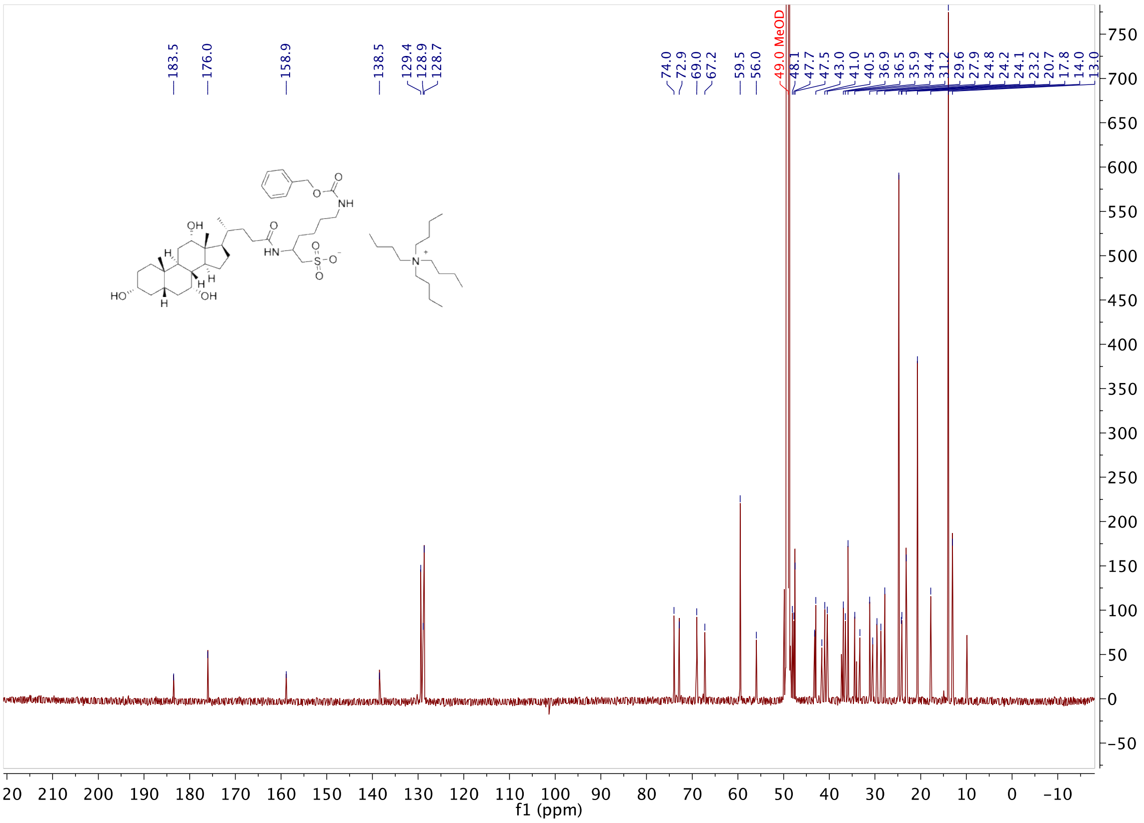


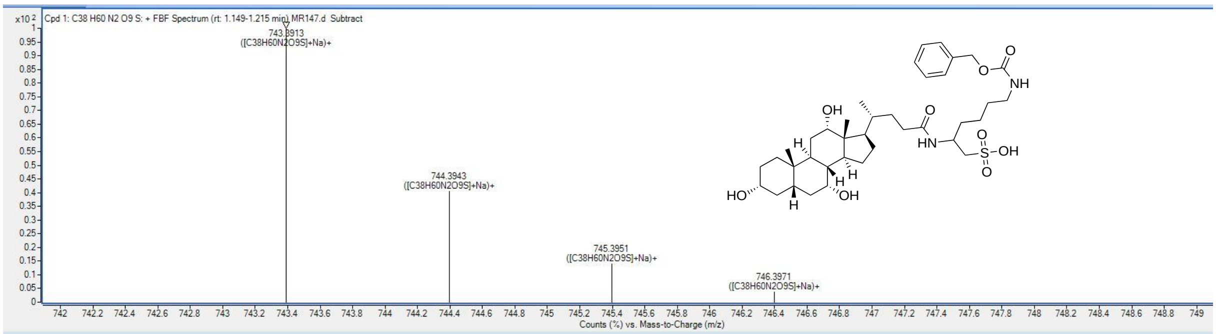


**Compound List**

Automatically Show Columns

| Label | Show/Hide | Cpd | File | ID Source | Name | Formula | Flags (Tgt) | Saturated | RT | RT (DB) |
|---|---|---|---|---|---|---|---|---|---|---|
| Cpd 1: C38 H60 N2 O9 S | ☑ | 1 | MR147.d | FBF | | C38 H60 N2 O9 S | | | 1.083 | |

| Best | ID Source | Name | Formula | Score | Score (RT) | RT Diff | Diff (ppm) | Score (Lib) | Score (DB) | Score (MFG) | Num Sp |
|---|---|---|---|---|---|---|---|---|---|---|---|
| ⦿ | FBF | | C38 H60 N2 O9 S | 98.97 | | | -0.2 | | | | |

| m/z | Species | Ion Formula | Score (MFG) | Score (MFG, MS/MS) | Score (MS) | Score (mass) | Score (iso. abund) | Score (iso. spacing) | Height |
|---|---|---|---|---|---|---|---|---|---|
| 743.3913 | (M+Na)+ | | | | 98.97 | 99.94 | 96.97 | 99.45 | 63070.5 |

| m/z | m/z (Calc) | Diff (ppm) | Height | Height (Calc) | Height % | Height % (Calc) | Height Sum% (Calc) | Height Sum % |
|---|---|---|---|---|---|---|---|---|
| 743.3913 | 743.3912 | -0.14 | 63070.5 | 61083.5 | 100 | 100 | 61.2 | 63.2 |
| 744.3943 | 744.3944 | 0.13 | 25539.6 | 26664.7 | 40.5 | 43.7 | 26.7 | 25.6 |
| 745.3951 | 745.3943 | -1.08 | 8804.3 | 9544.1 | 14 | 15.6 | 9.6 | 8.8 |
| 746.3971 | 746.3954 | -2.38 | 2329.5 | 2451.5 | 3.7 | 4 | 2.5 | 2.3 |

| m/z | Species | Ion Formula | Score (MFG) | Score (MFG, MS/MS) | Score (MS) | Score (mass) | Score (iso. abund) | Score (iso. spacing) | Height |
|---|---|---|---|---|---|---|---|---|---|
| 721.4085 | (M+H)+ | | | | 59.28 | 99.75 | 0 | 49.47 | 1259.3 |

**Compound 14**

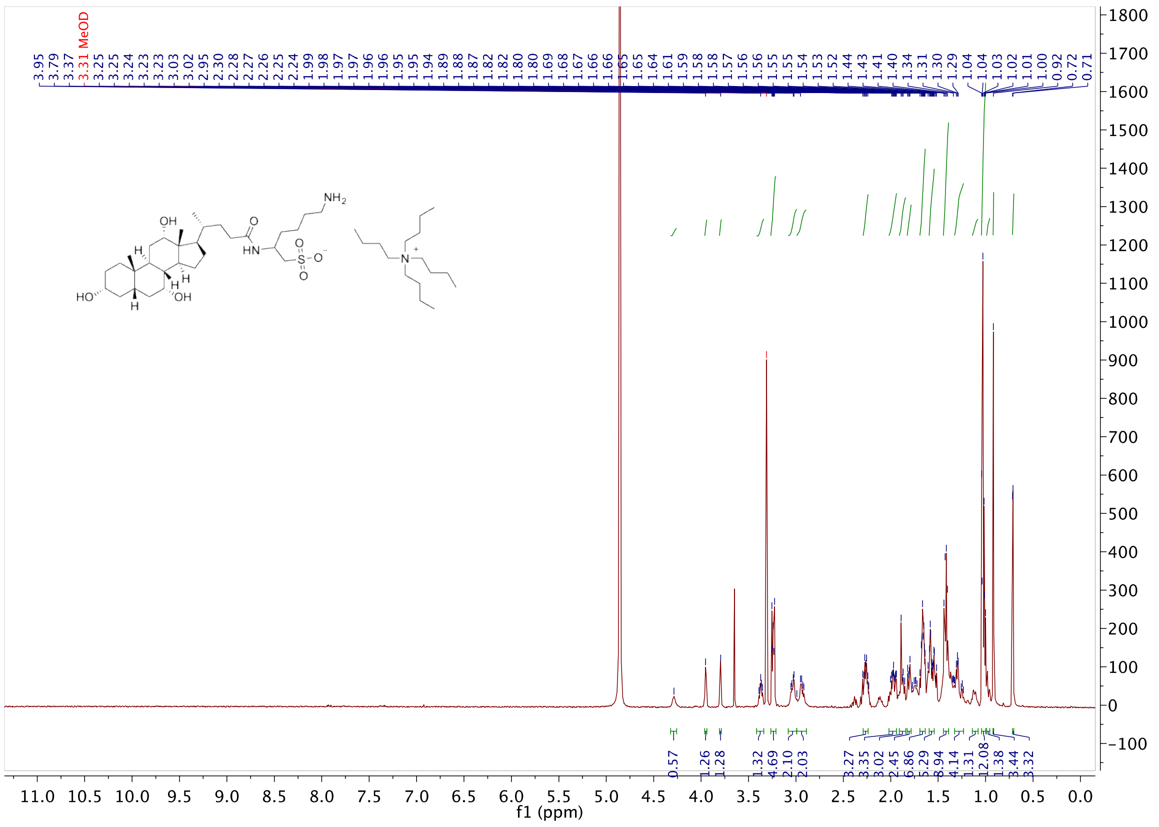

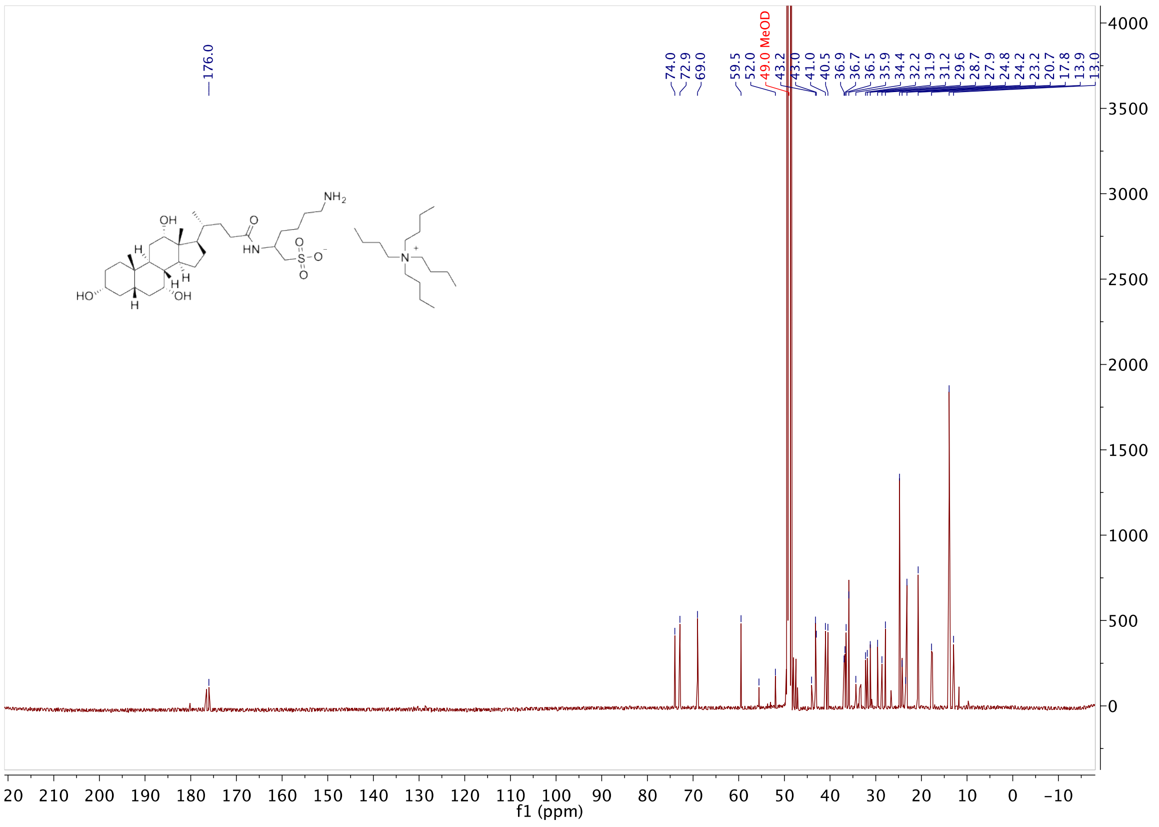

#### Compound 3

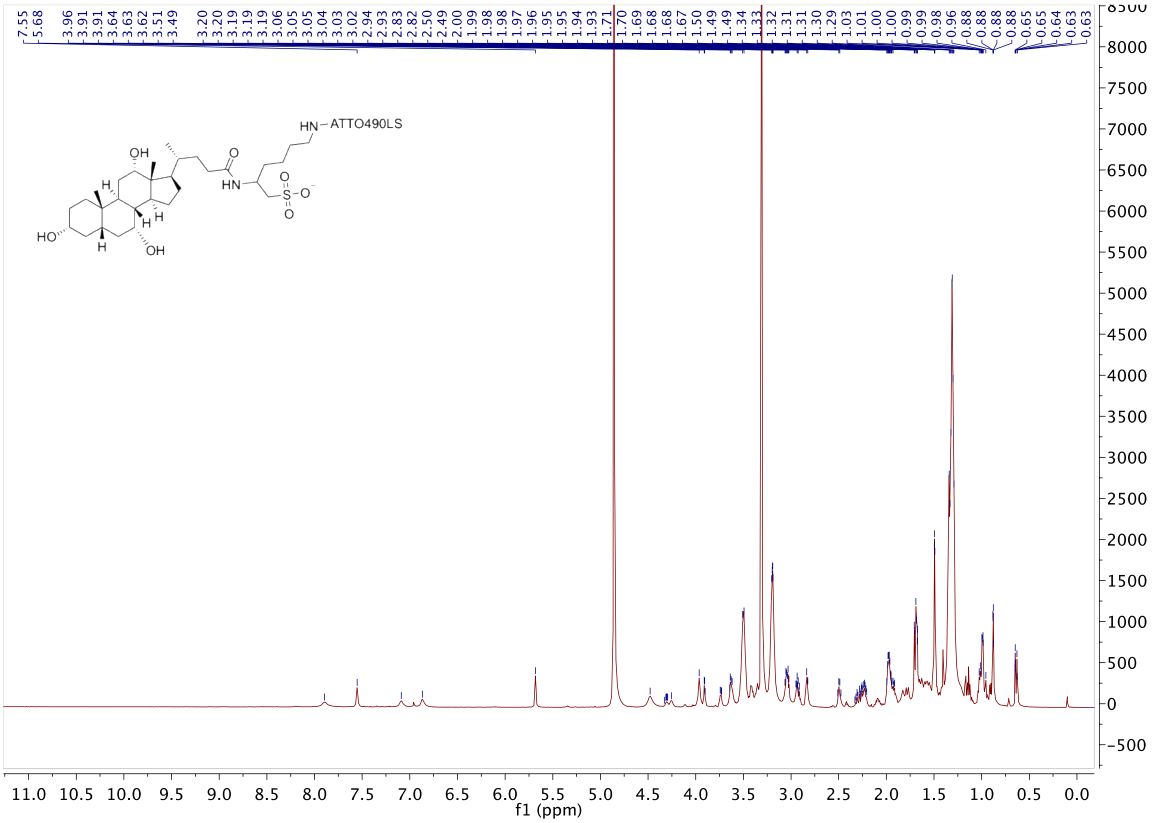

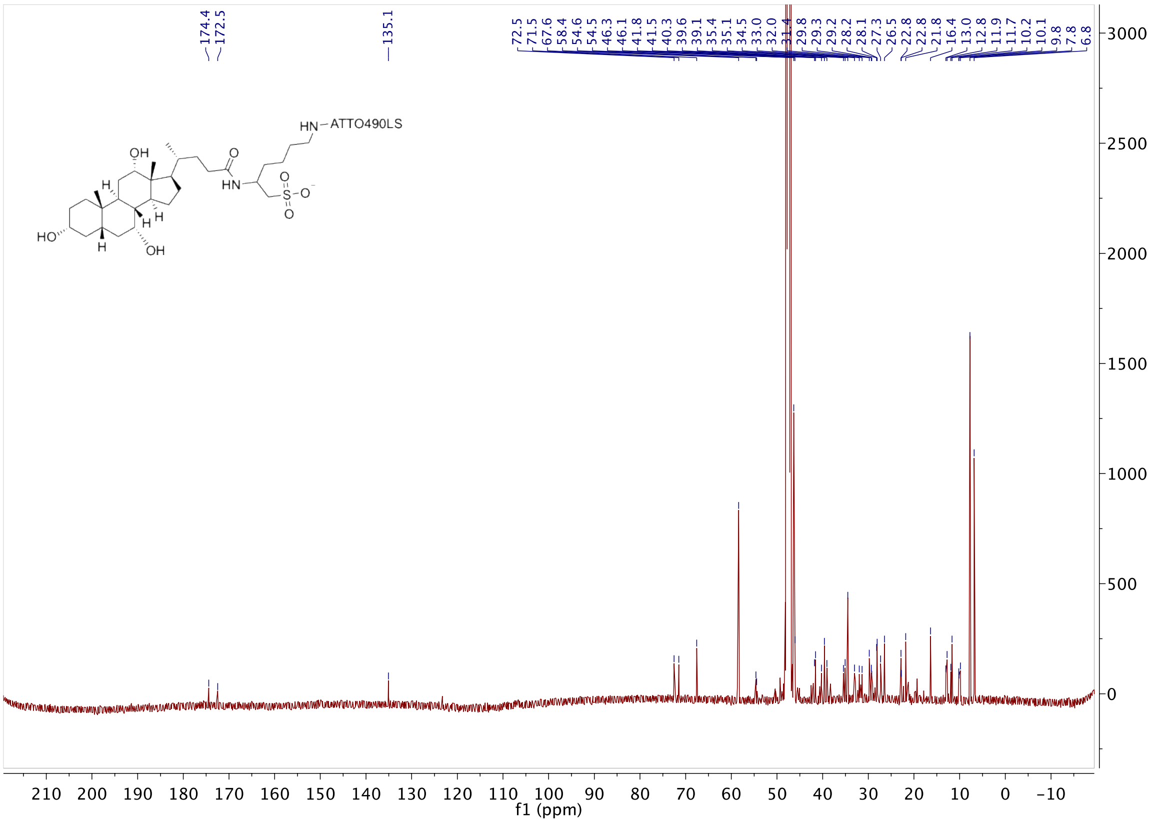

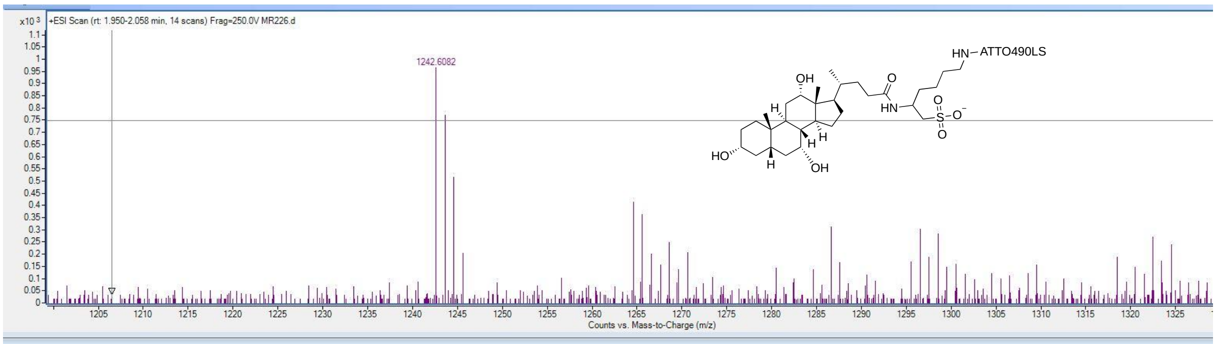

#### Compound 4

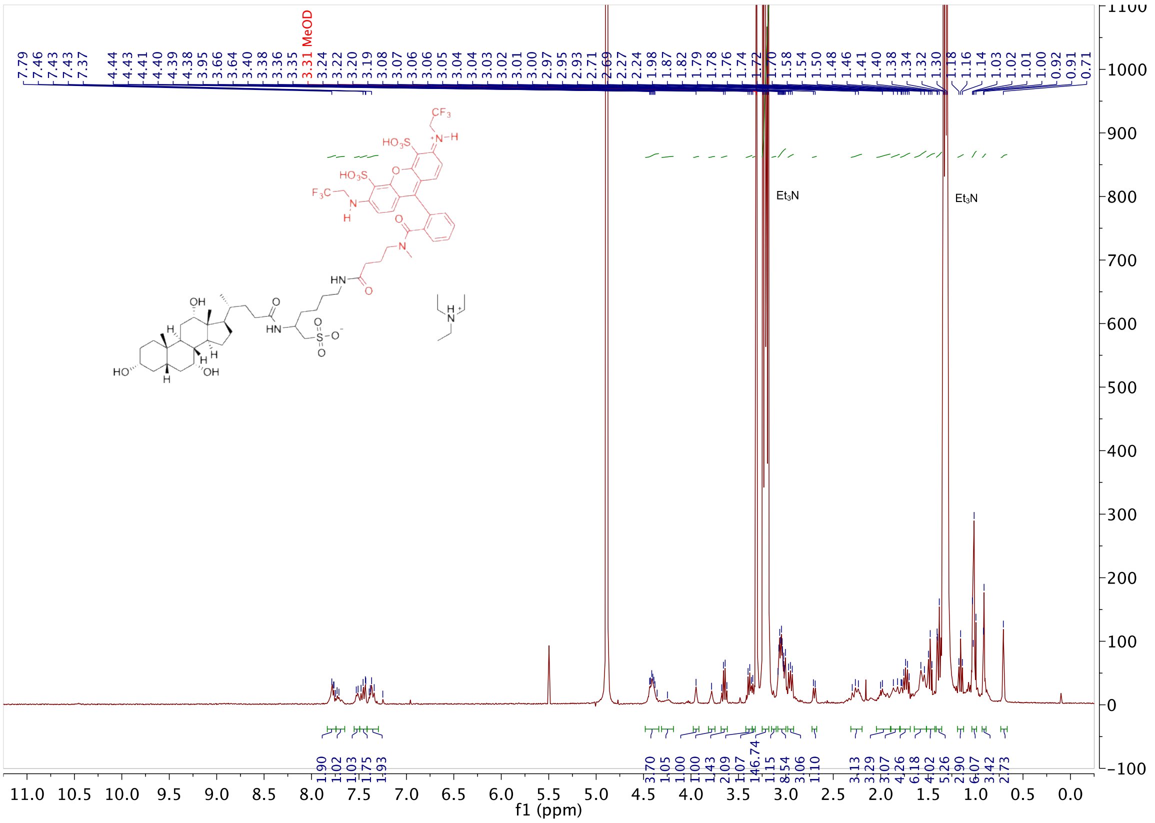

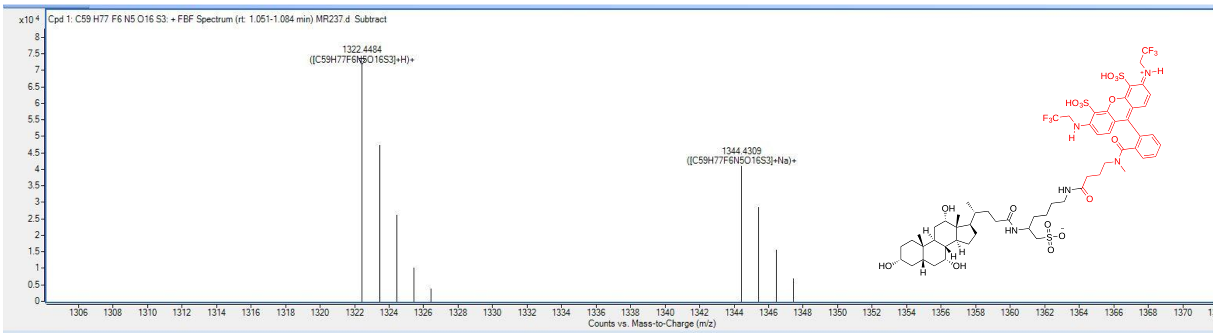


**Compound List**

Automatically Show Columns

| Label | Show/Hide | Cpd | File | ID Source | Name | Formula | Flags (Tgt) | Saturated | RT | RT (DB) | m/z |
|---|---|---|---|---|---|---|---|---|---|---|---|
| Cpd 1: C59 H77 F6 N5 O16 S3 | ☑ | 1 | MR237.d | FBF | | C59 H77 F6 N5 O16... | | | 1.051 | | |

| Best | ID Source | Name | Formula | Score | Score (RT) | RT Diff | Diff (ppm) | Score (Lib) | Score (DB) | Score (MFG) | Num Spe |
|---|---|---|---|---|---|---|---|---|---|---|---|
| ⊙ | FBF | | C59 H77 F6 N5 O16... | 97.49 | | | 1.08 | | | | |

| m/z | Species | Ion Formula | Score (MFG) | Score (MFG, MS/MS) | Score (MS) | Score (mass) | Score (iso. abund) | Score (iso. spacing) | Height |
|---|---|---|---|---|---|---|---|---|---|
| 1322.4484 | (M+H)+ | | | | 97.49 | 97.17 | 96.53 | 99.28 | 71685.2 |

| m/z | m/z (Calc) | Diff (ppm) | Height | Height (Calc) | Height % | Height % (Calc) | Height Sum% (Calc) | Height Sum % |
|---|---|---|---|---|---|---|---|---|
| 1322.4484 | 1322.4504 | 1.54 | 71685.2 | 68249.9 | 100 | 100 | 43 | 45.1 |
| 1323.452 | 1323.4535 | 1.18 | 47180.9 | 47443.7 | 65.8 | 69.5 | 29.9 | 29.7 |
| 1324.4521 | 1324.453 | 0.7 | 26060.1 | 27650.8 | 36.4 | 40.5 | 17.4 | 16.4 |
| 1325.4541 | 1325.4538 | -0.2 | 10135.7 | 11507.8 | 14.1 | 16.9 | 7.2 | 6.4 |
| 1326.4508 | 1326.454 | 2.4 | 3828.7 | 4038.3 | 5.3 | 5.9 | 2.5 | 2.4 |

| m/z | Species | Ion Formula | Score (MFG) | Score (MFG, MS/MS) | Score (MS) | Score (mass) | Score (iso. abund) | Score (iso. spacing) | Height |
|---|---|---|---|---|---|---|---|---|---|
| 1344.4309 | (M+Na)+ | | | | 95.53 | 98.61 | 87.02 | 99.57 | 40969.2 |
