## Appendix for "Intravital dynamic and correlative imaging reveals diffusion-dominated canalicular and flow-augmented ductular bile flux"

**IDENTIFIERS**

Data and graphs are identified as [Treatment_Domain] Filename.!

Treatments: Basal–Control, Secretin– Secretin bolus, TCA–Taurocholate infusion

Domains: CV – Central vein, PV – Portal vein, MZ– Midzone, IBD– Interlobular bile duct

Eg: [TCACV] Image 40.czi! refers to data from central vein canalicular network under TCA treatment.

**RAW DATA** is available at: https://lab.vartak.org/repository/bile-flux/

Raw data is arranged as in a folder hierarchy as: Treatment >> Domain >> Files

**CROSS-REFERENCE**

Data in this appendix is related to: **Fig. 3C, D, E**

Data and results from this appendix are summarized in **Supplementary Tables S3 and S4**

Data is generated using the Python scripts: **RICSAnalysis.py** and **TICSAnalysis.py**

**CONTENTS**


| Treatment | Data | Domain | Page | |
|---|---|---|---|---|
| | | | **RICS** | **TICS** |
| **Basal** | Experimental Autocorrelation Curves | CV | 1-9 | 133-144 |
| | | IBD | 10-18 | 145-152 |
| | | MZ | 18-28 | 153-161 |
| | | PV | 29-39 | 162-170 |
| | Fitting | CV, IBD, MZ, PV | 121-124 | 257-260 |
| **Secretin** | Experimental Autocorrelation Curves | CV | 40-45 | 171-178 |
| | | IBD | 46-59 | 179-195 |
| | | MZ | 60-69 | 196-207 |
| | | PV | 70-77 | 208-223 |
| | Fitting | | 125-128 | 261-264 |
| **TCA** | Experimental Autocorrelation Curves | CV | 78-89 | 224-230 |
| | | IBD | 90-100 | 231-242 |
| | | MZ | 101-110 | 243-249 |
| | | PV | 111-120 | 250-256 |
| | Fitting | CV, IBD, MZ, PV | 129-132 | 265-268 |

Treatment: Basal , Domain: CV , File: Image 16.czi

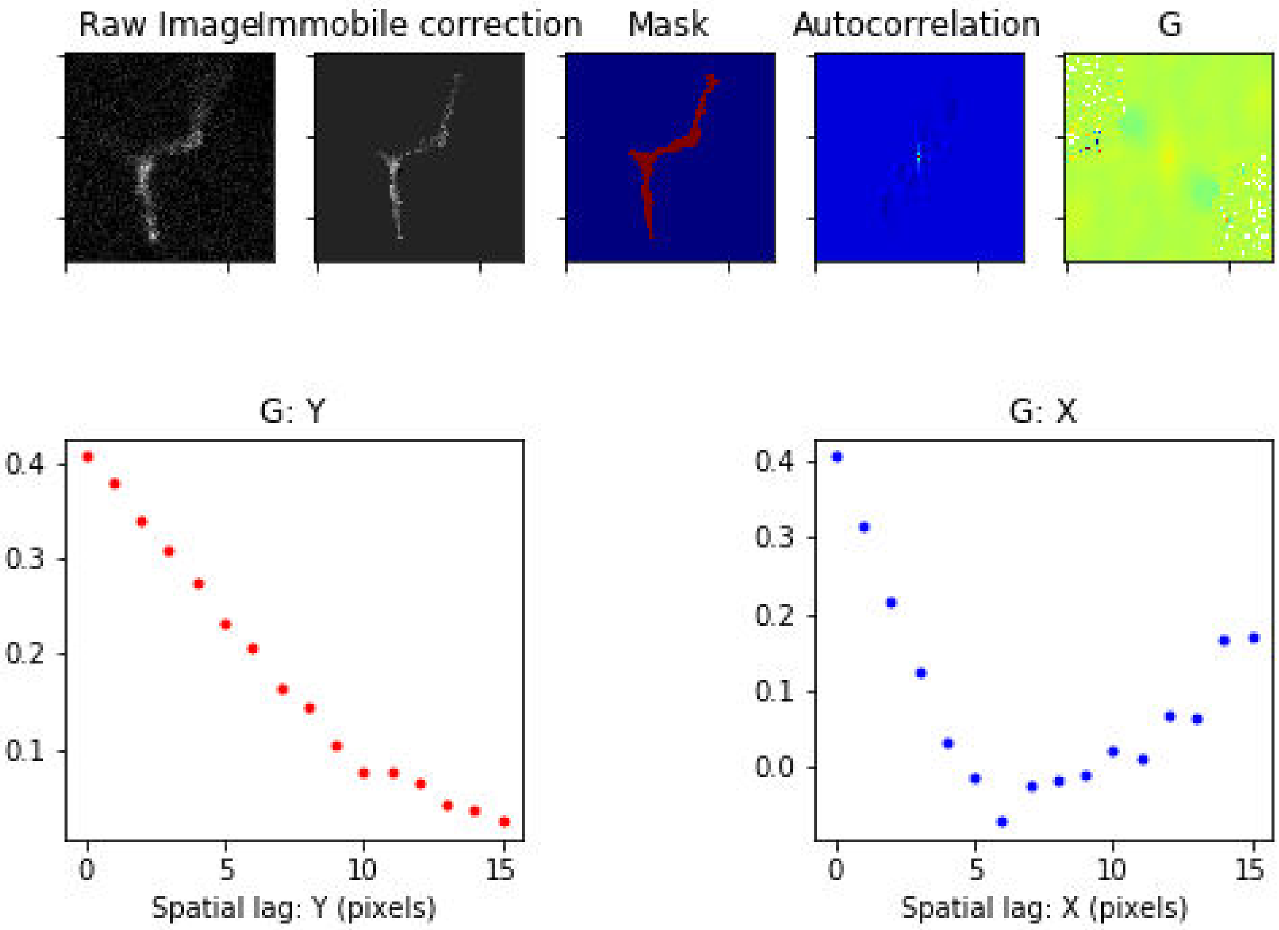

Treatment: Basal , Domain: CV , File: Image 17.czi

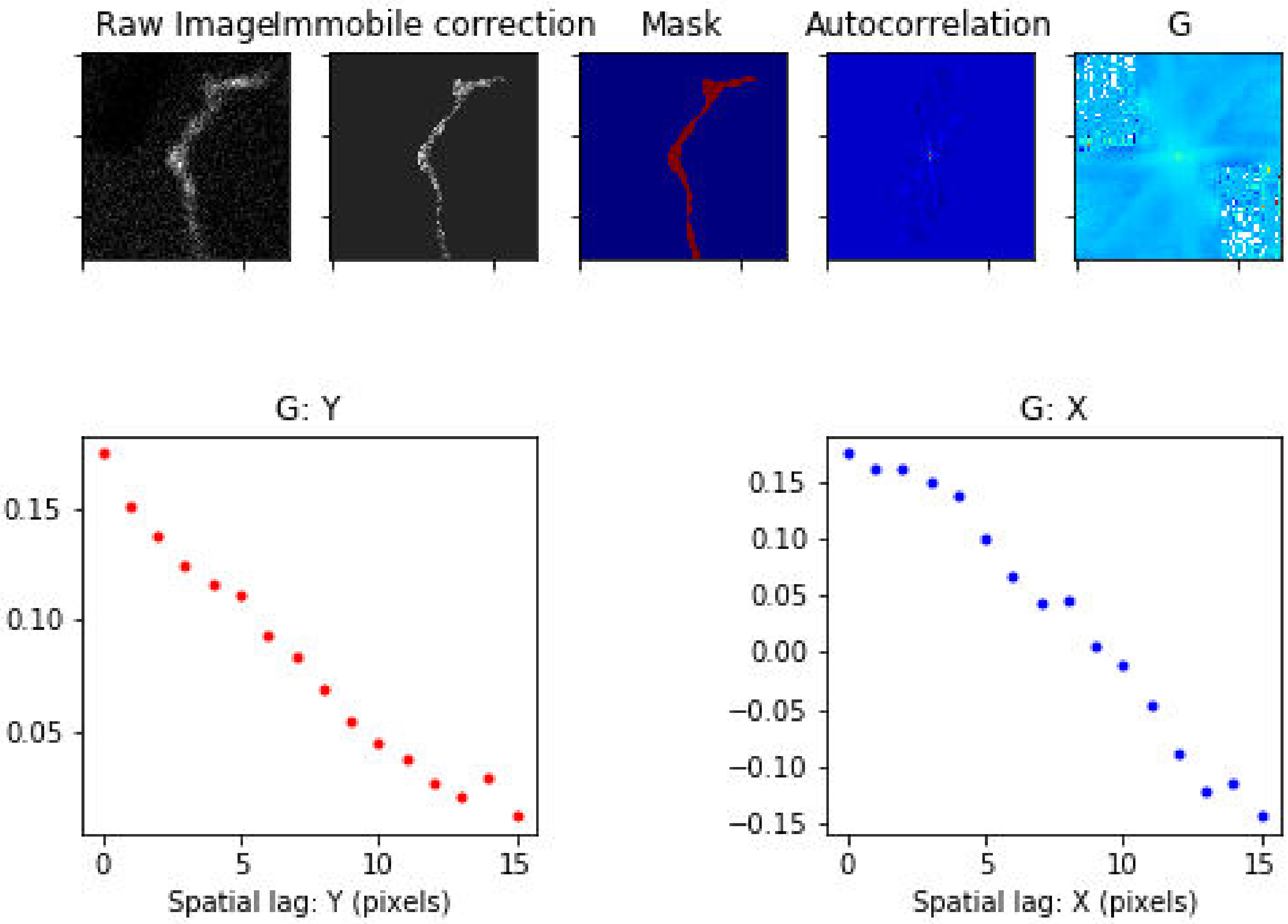

Treatment: Basal , Domain: CV , File: Image 18.czi

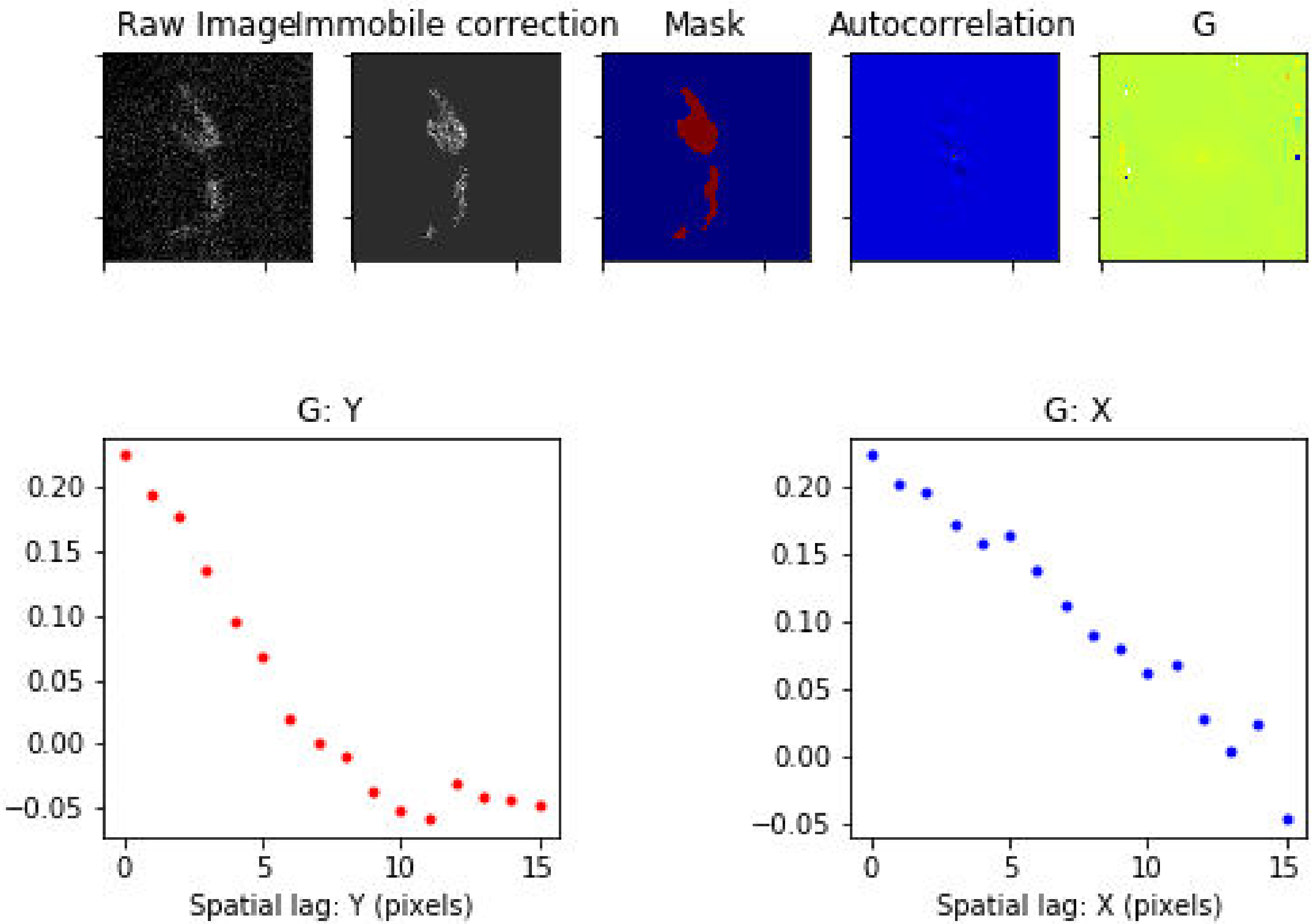

### Treatment: Basal , Domain: CV , File: Image 19.czi

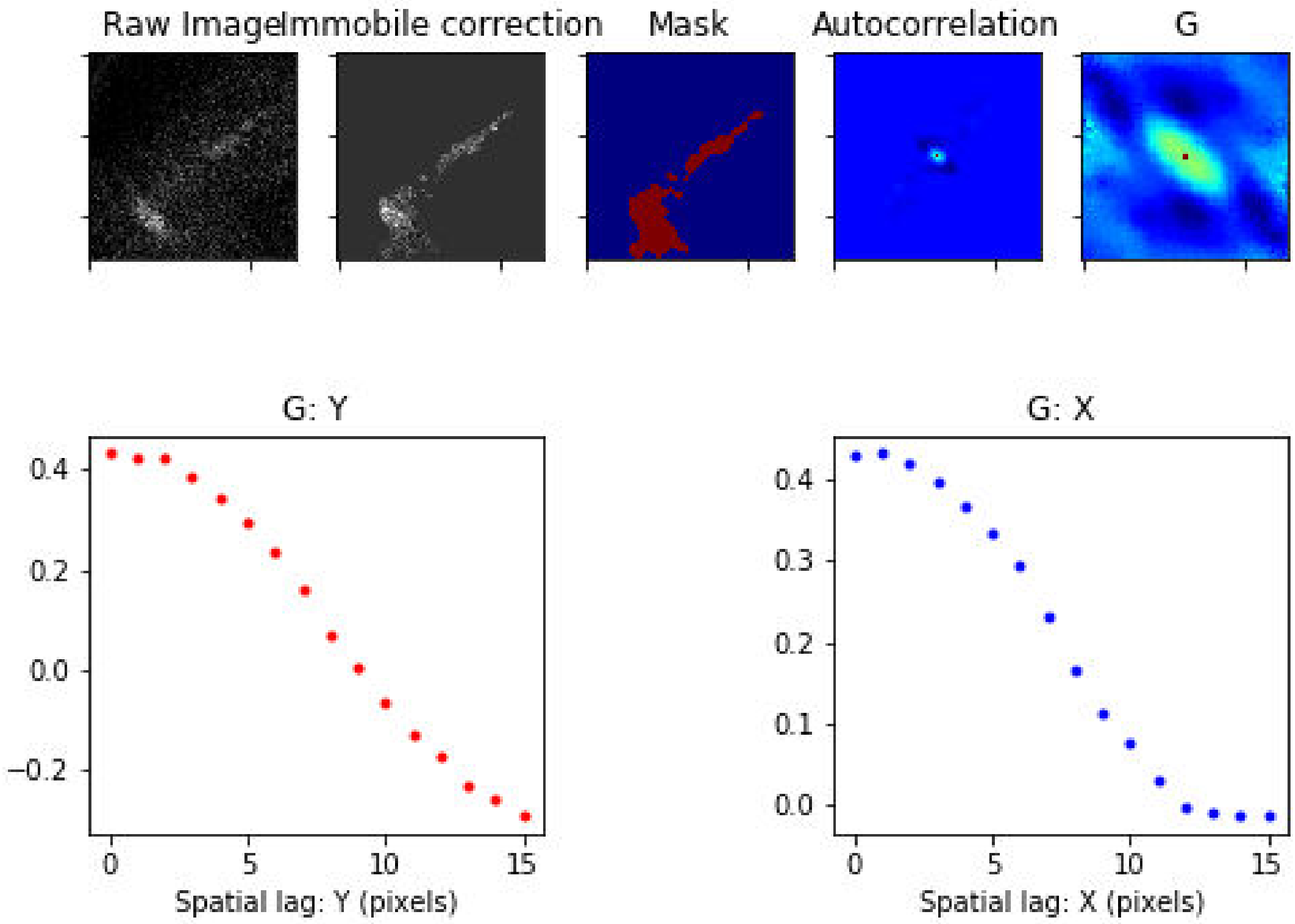

### Treatment: Basal , Domain: CV , File: Image 20.czi

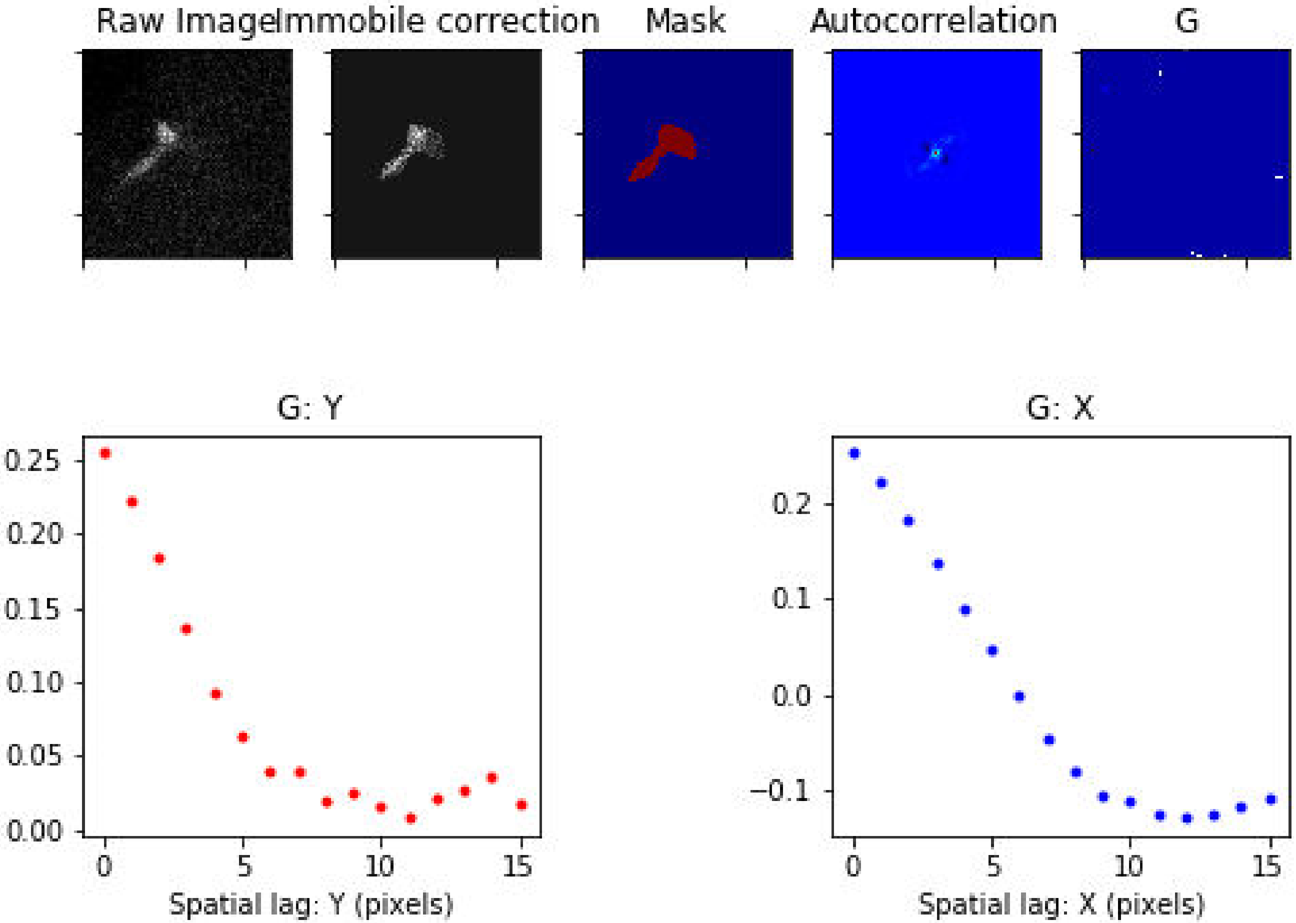

Treatment: Basal , Domain: CV , File: Image 28.czi

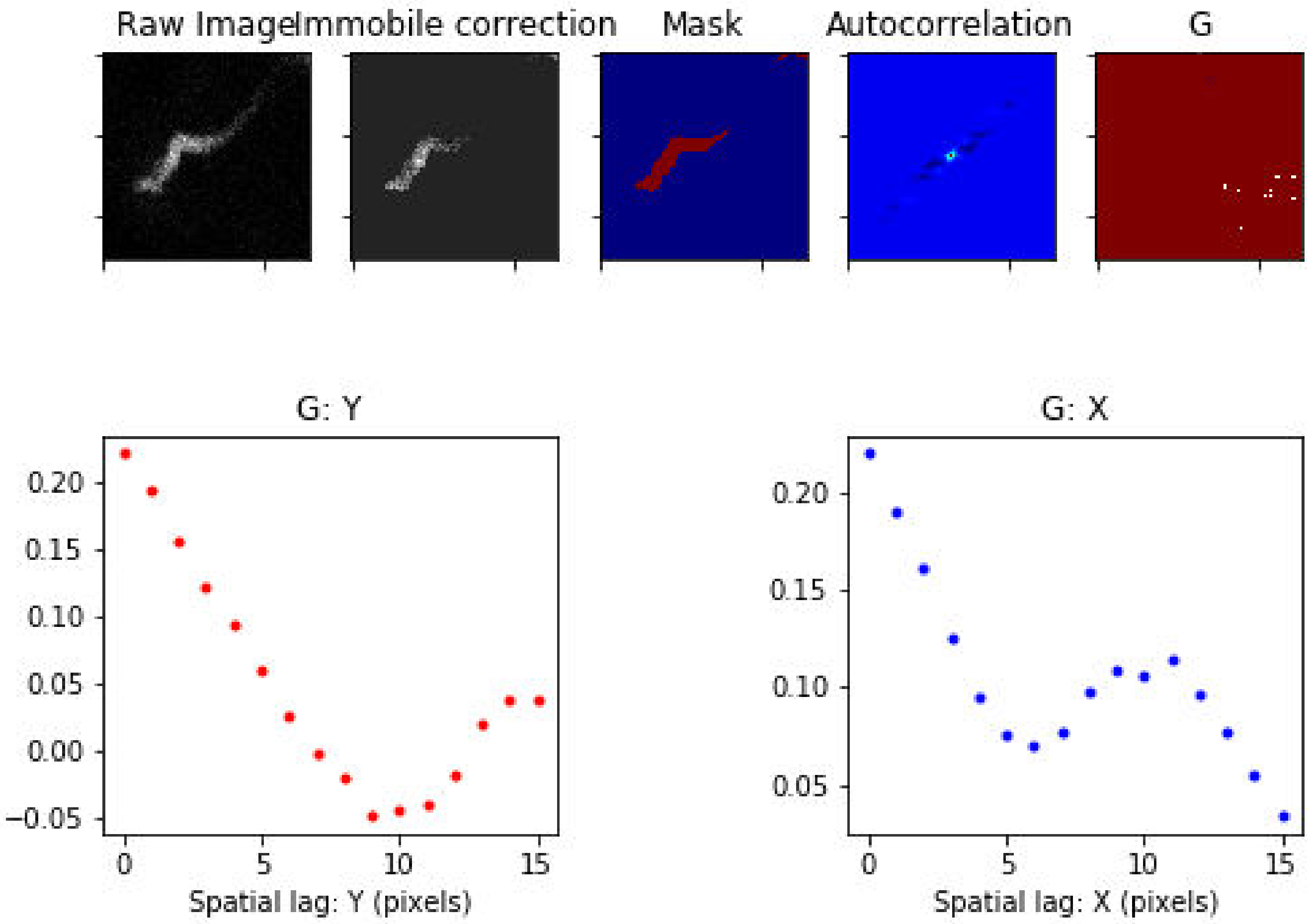

Treatment: Basal , Domain: CV , File: Image 30.czi

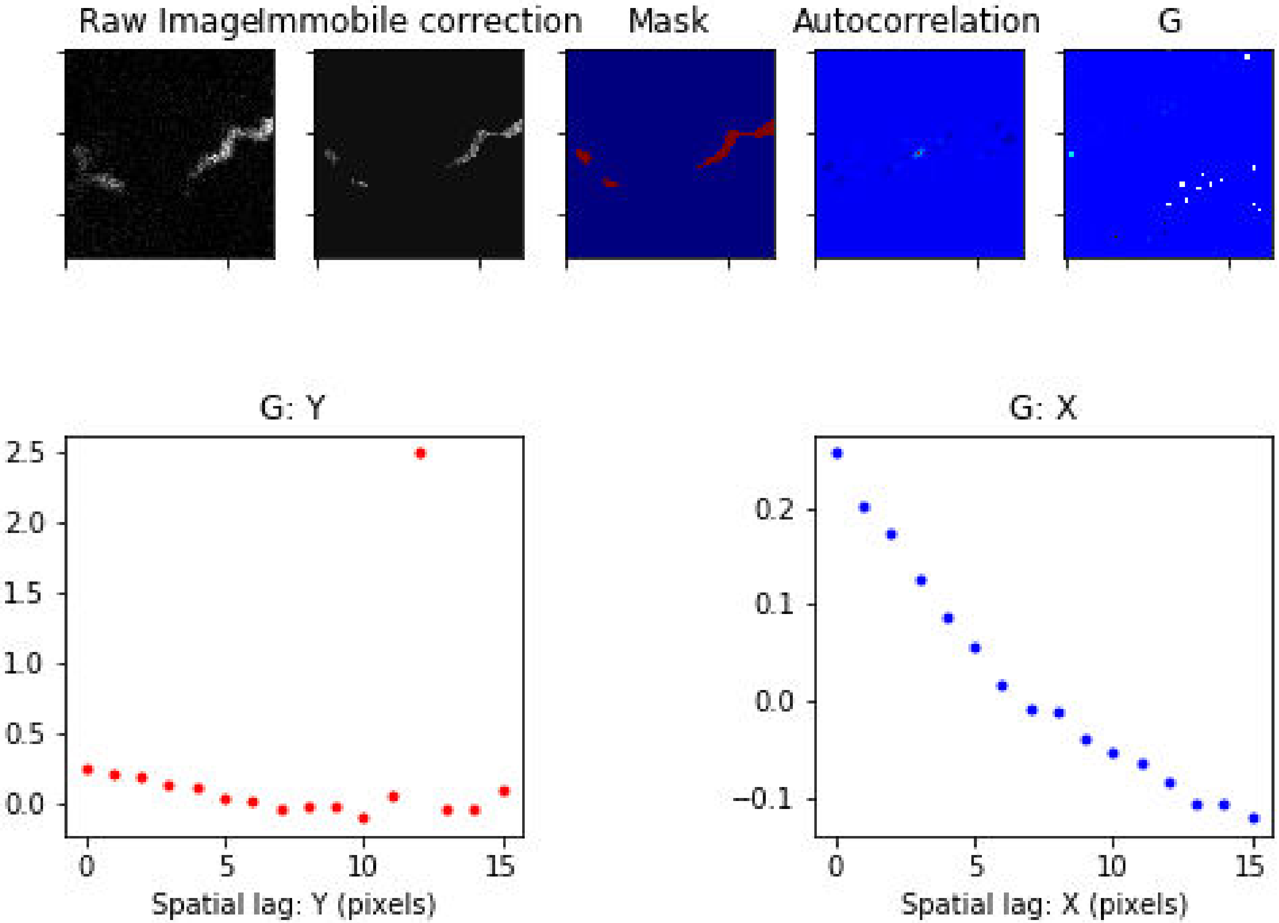

### Treatment: Basal , Domain: CV , File: Image 33.czi

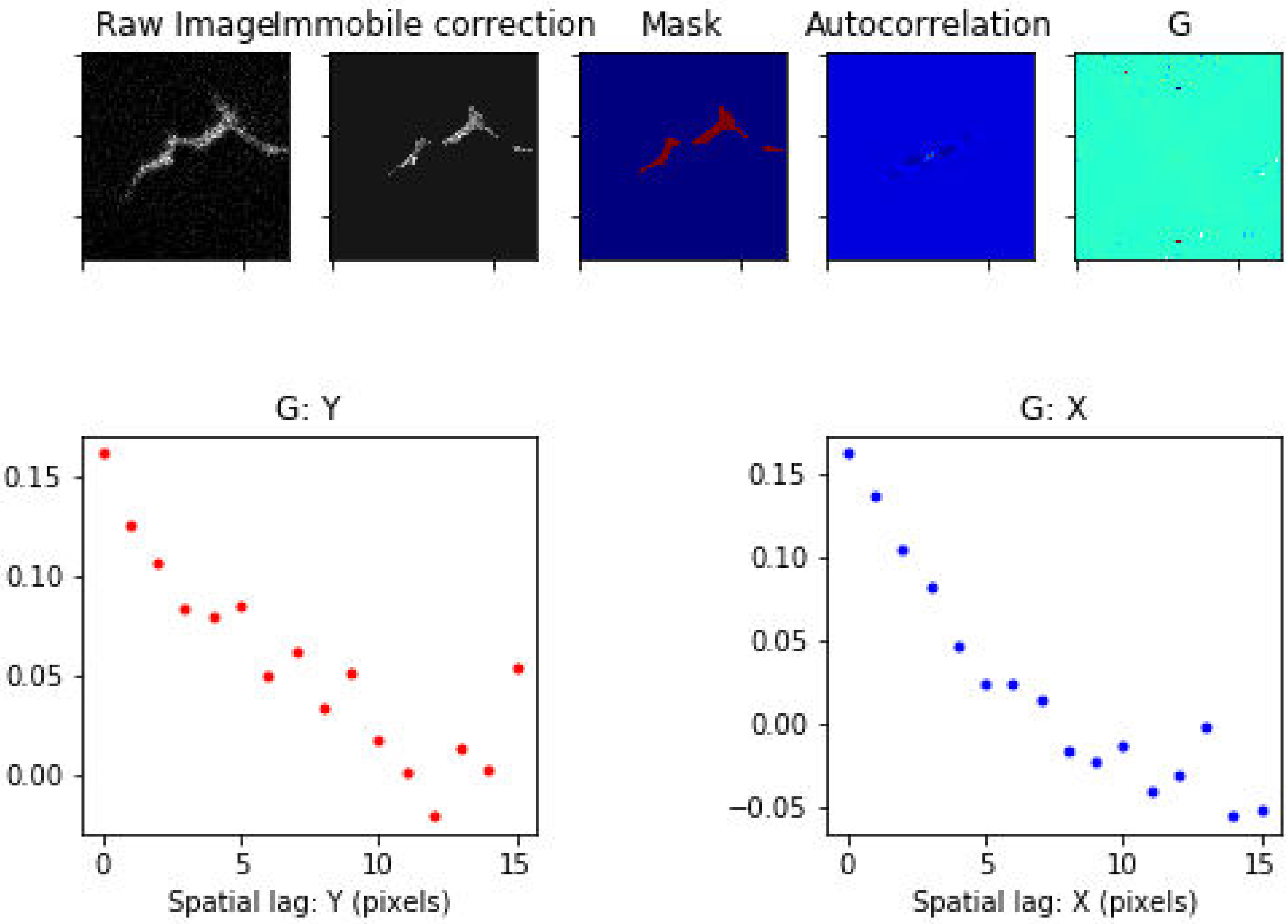

Treatment: Basal , Domain: CV , File: Image 41.czi

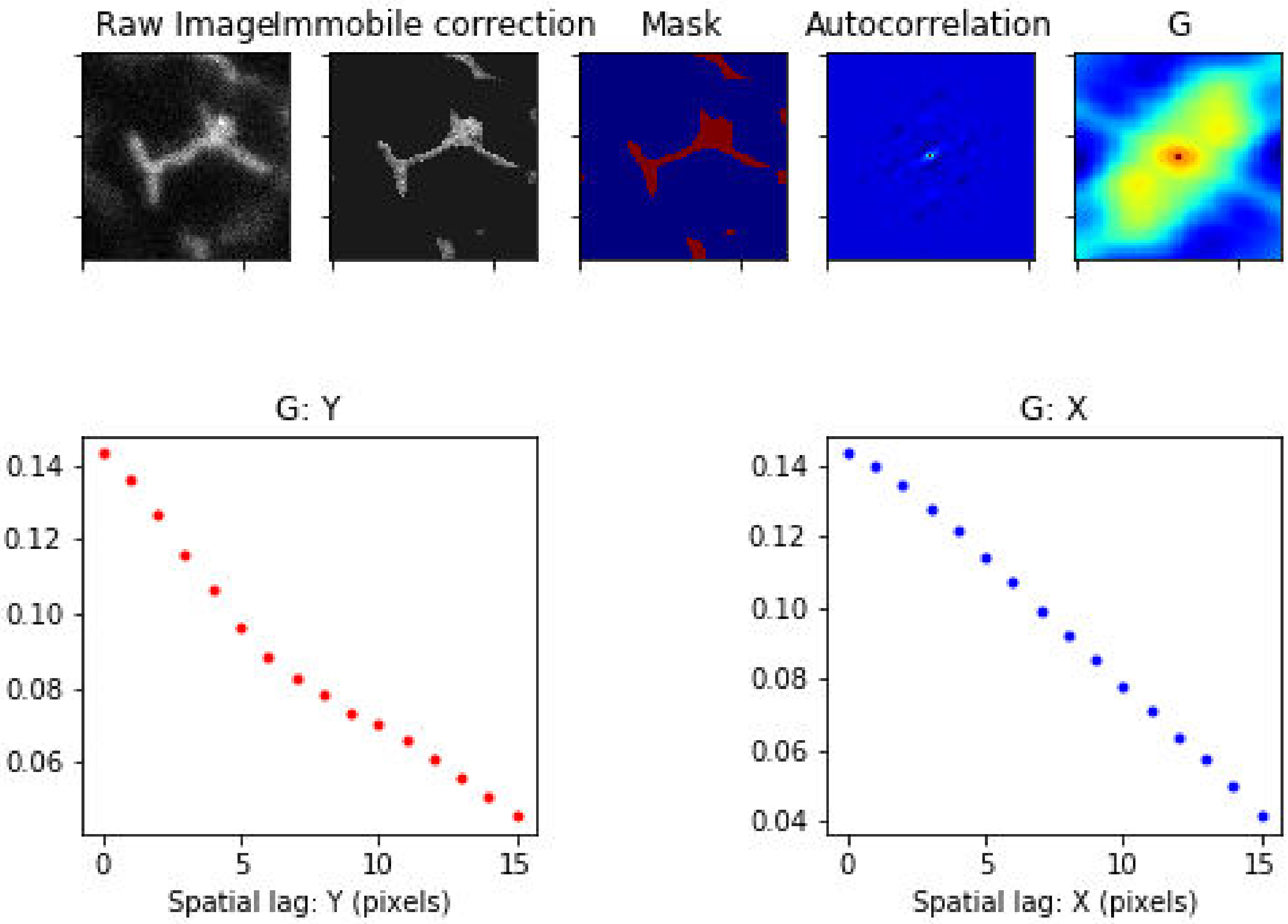

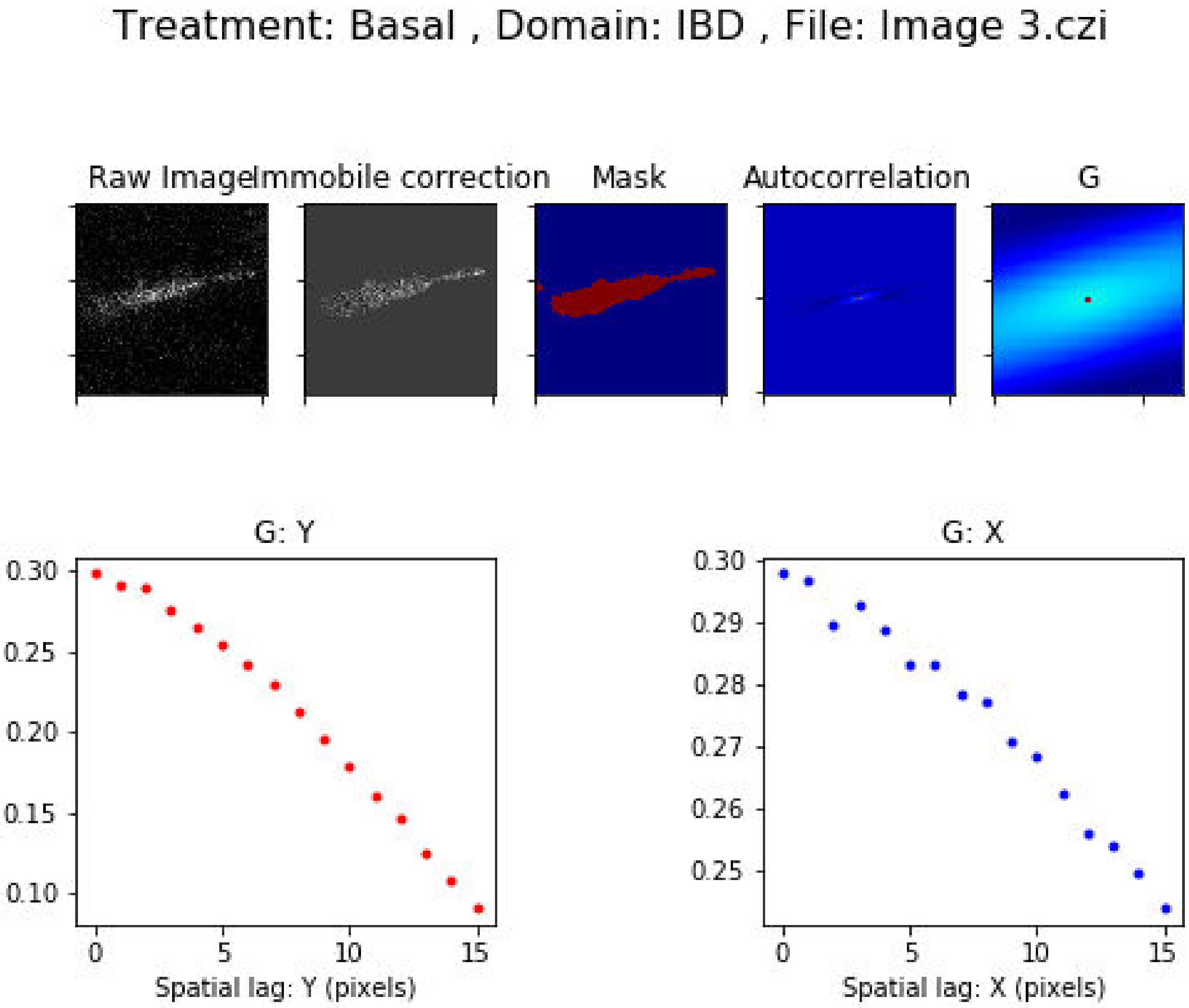
Treatment: Basal , Domain: IBD , File: Image 3.czi
Raw Image
Immobile correction
Mask
Autocorrelation
G
G: Y
0.30
0.25
0.20
0.15
0.10
0
5
10
15
Spatial lag: Y (pixels)
G: X
0.30
0.29
0.28
0.27
0.26
0.25
0
5
10
15
Spatial lag: X (pixels)

Treatment: Basal , Domain: IBD , File: Image 4.czi

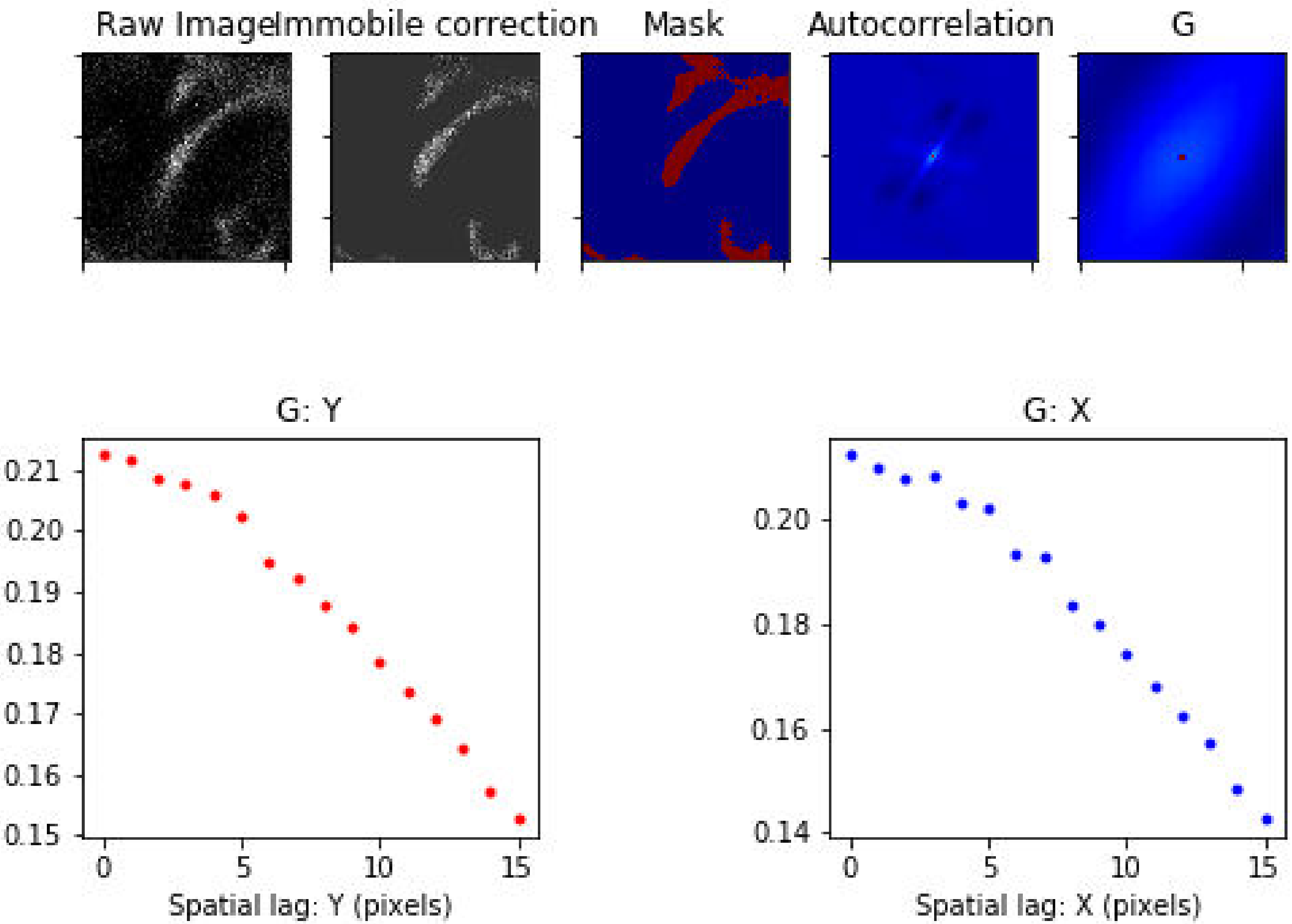

#### Treatment: Basal , Domain: IBD , File: Image 7.czi

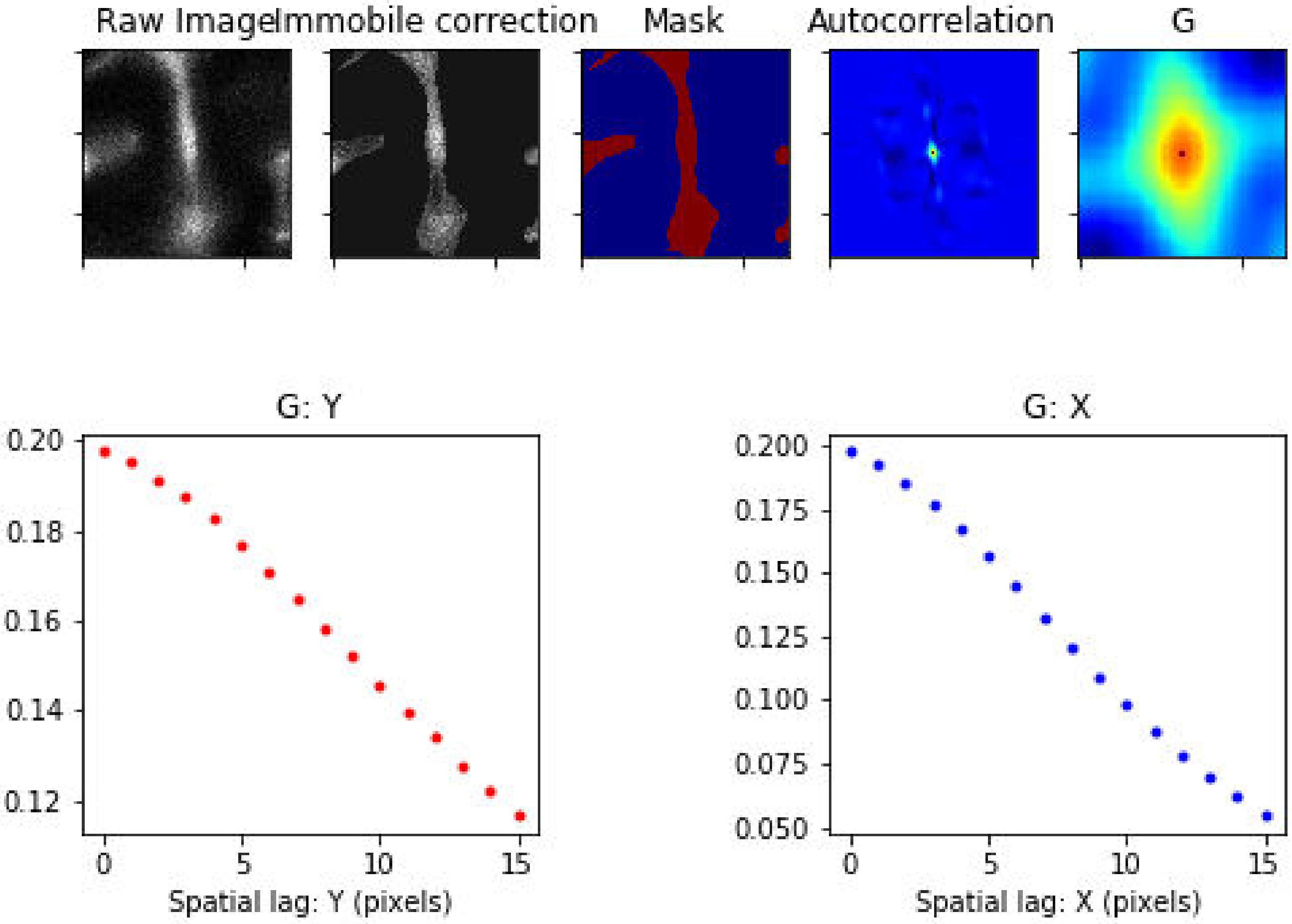

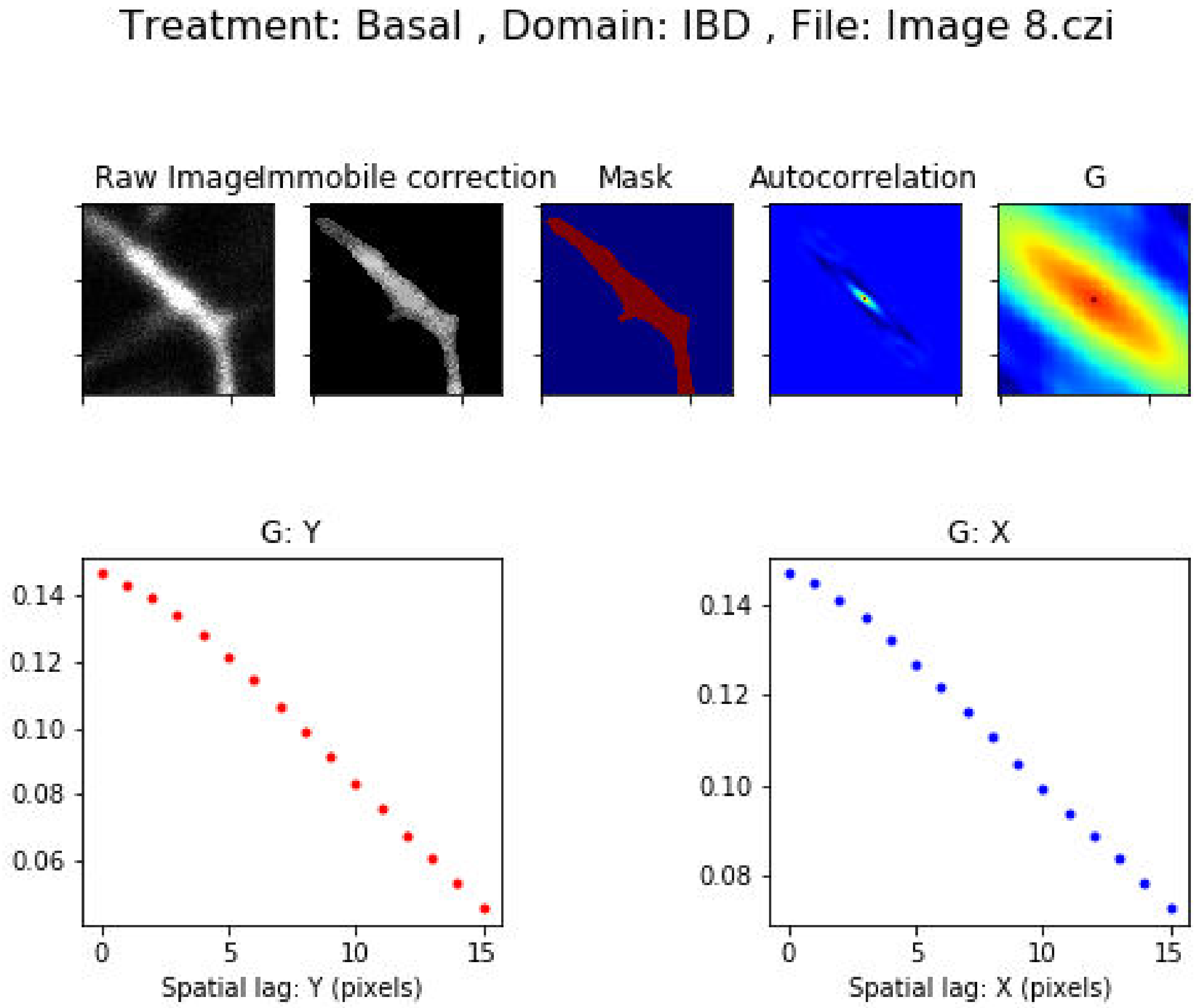
Treatment: Basal , Domain: IBD , File: Image 8.czi
Raw Image
Immobile correction
Mask
Autocorrelation
G
G: Y
0.14
0.12
0.10
0.08
0.06
0
5
10
15
Spatial lag: Y (pixels)
G: X
0.14
0.12
0.10
0.08
0
5
10
15
Spatial lag: X (pixels)

Treatment: Basal , Domain: IBD , File: Image 9.czi

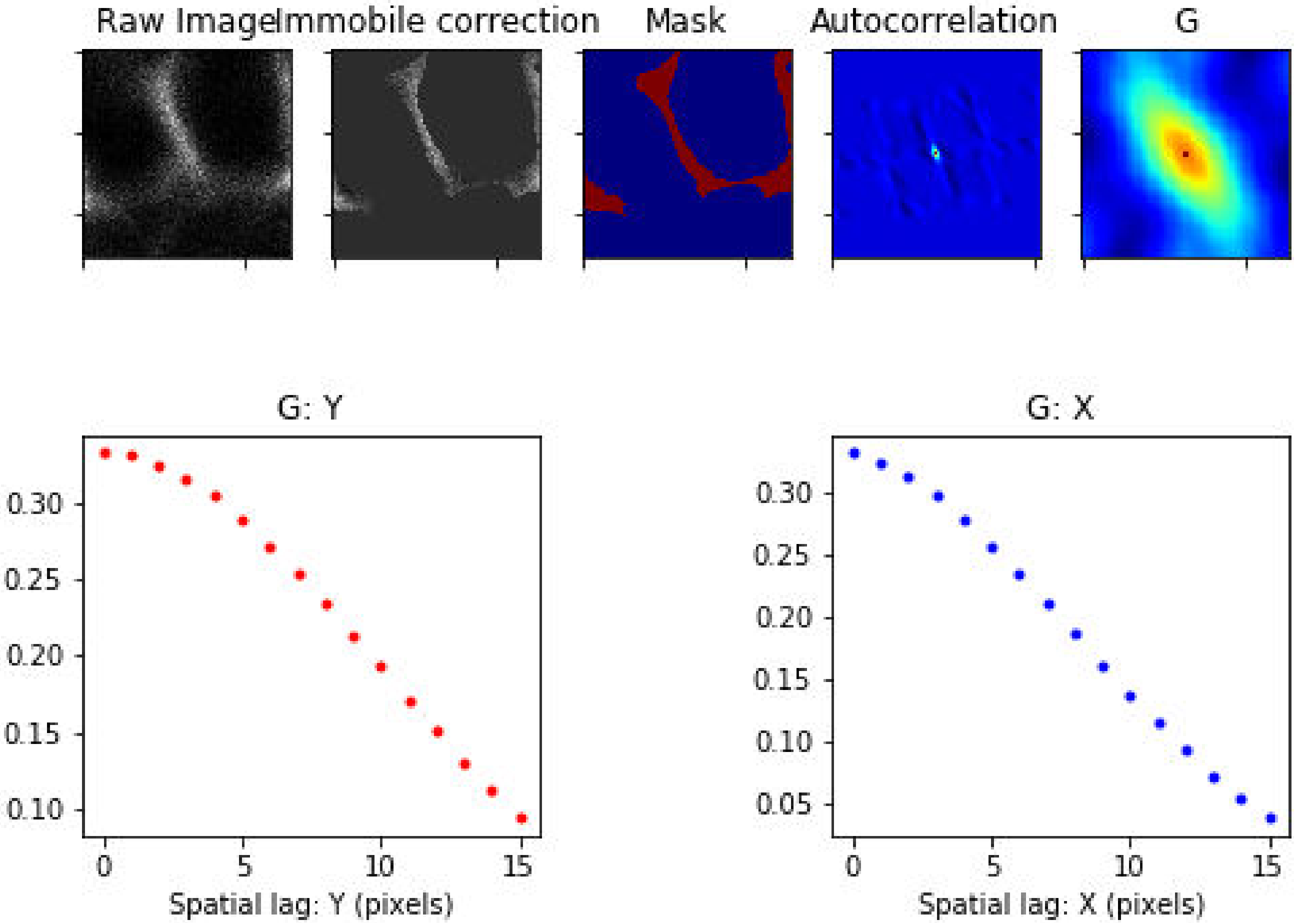

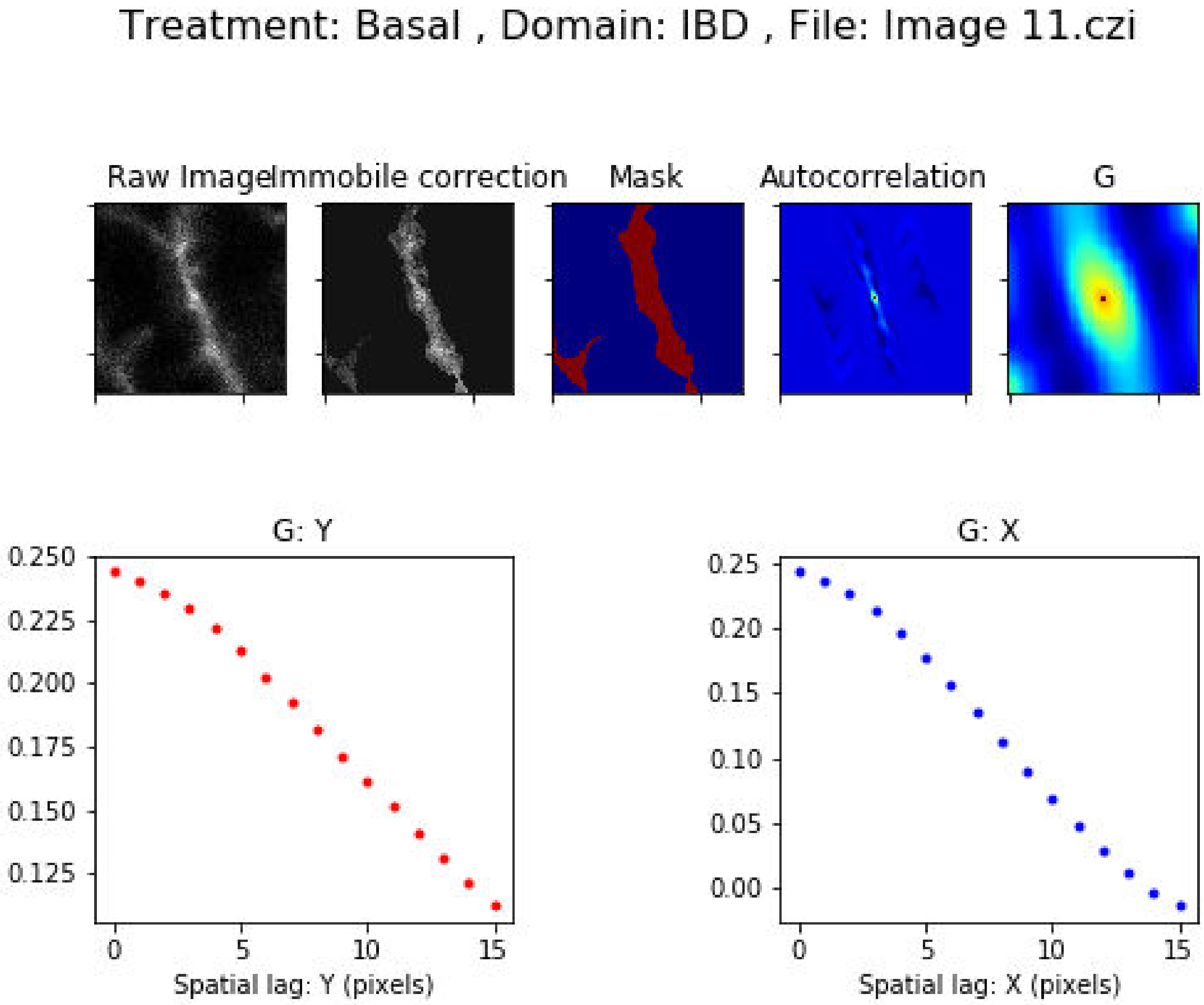
Treatment: Basal , Domain: IBD , File: Image 11.czi
Raw Image
Immobile correction
Mask
Autocorrelation
G
G: Y
Spatial lag: Y (pixels)
G: X
Spatial lag: X (pixels)

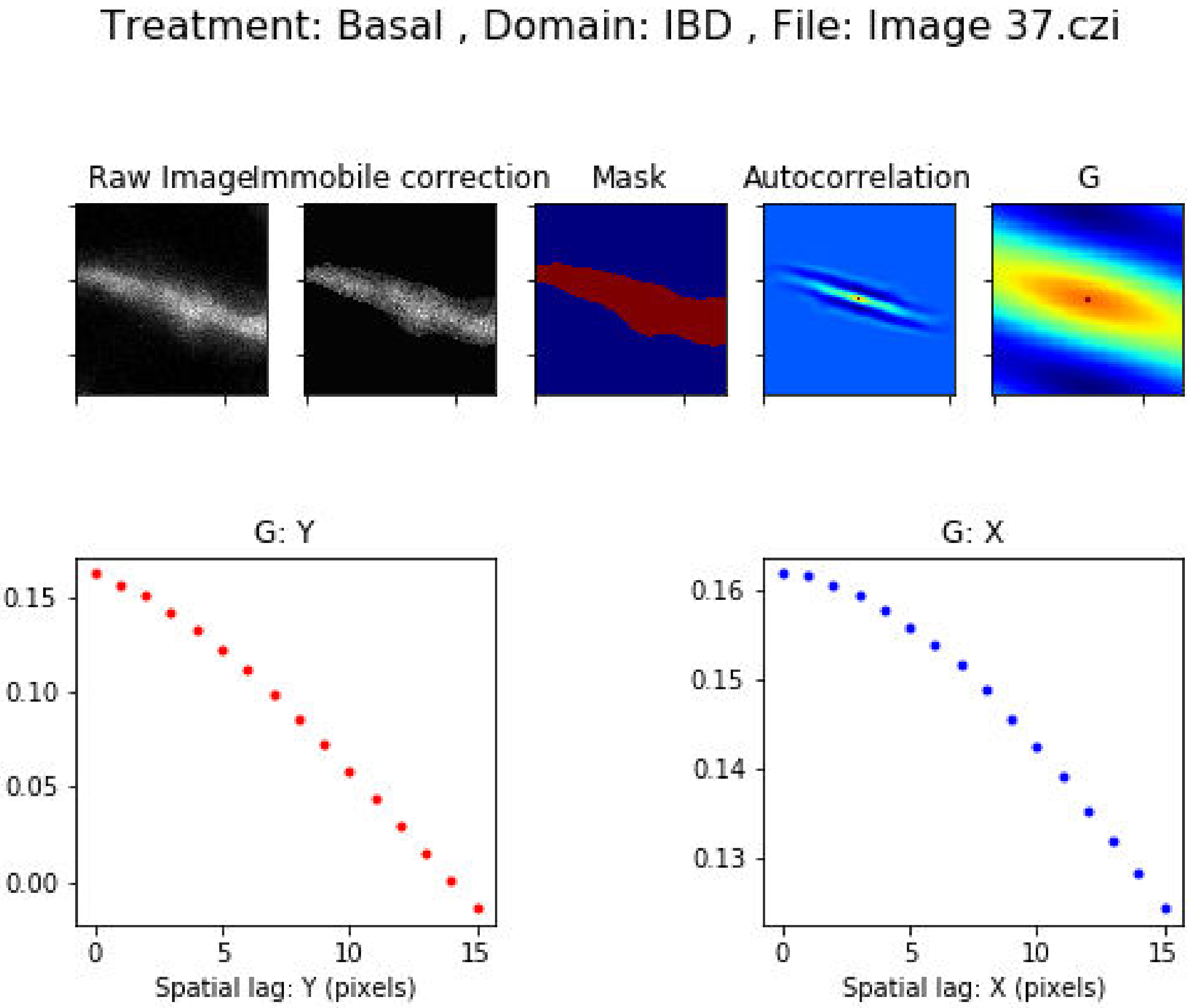
Treatment: Basal , Domain: IBD , File: Image 37.czi
Raw Image
Immobile correction
Mask
Autocorrelation
G
G: Y
0.15
0.10
0.05
0.00
0
5
10
15
Spatial lag: Y (pixels)
G: X
0.16
0.15
0.14
0.13
0
5
10
15
Spatial lag: X (pixels)

Treatment: Basal , Domain: IBD , File: Image 38.czi

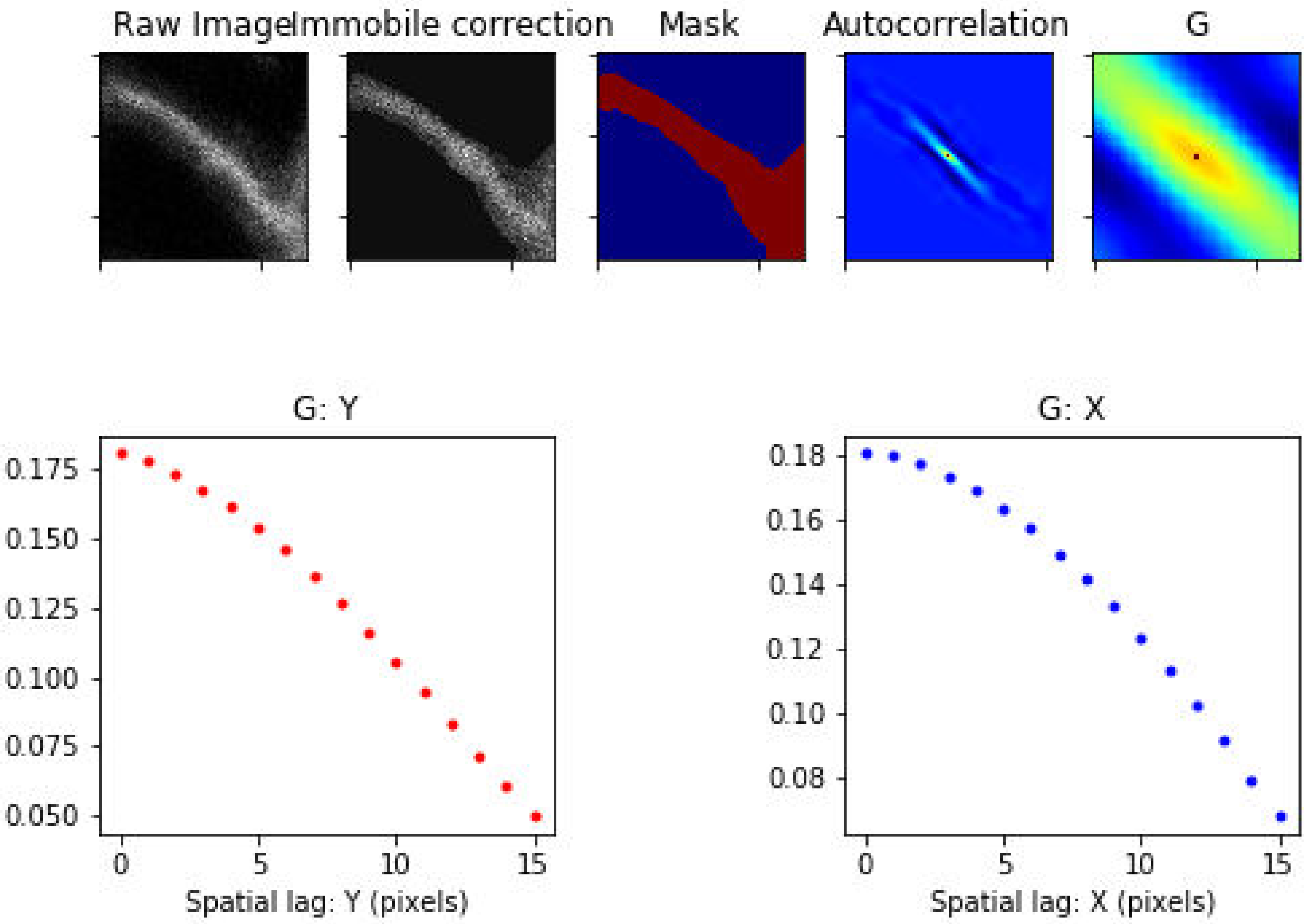

Treatment: Basal , Domain: IBD , File: Image 40.czi

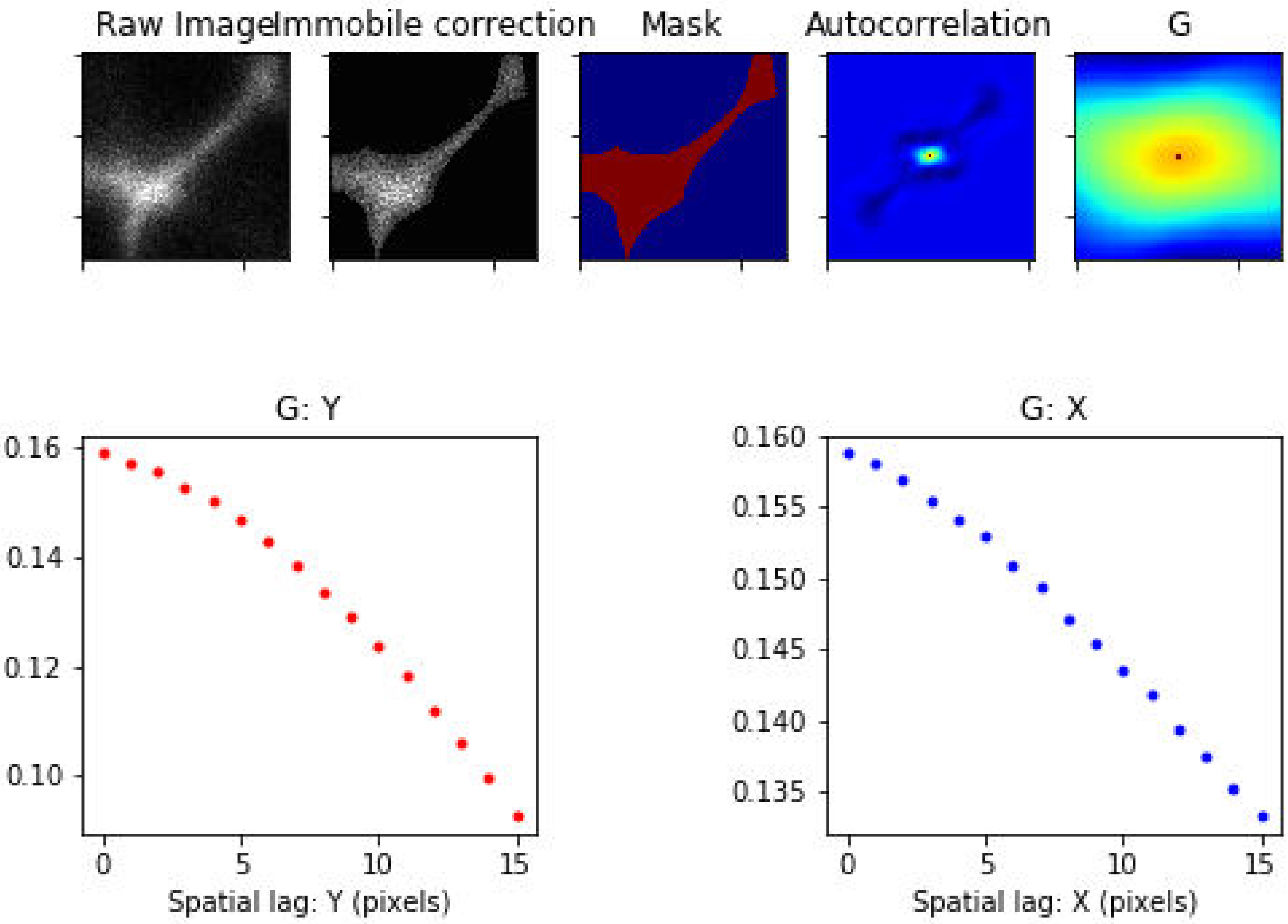

### Treatment: Basal , Domain: MZ , File: Image 18.czi

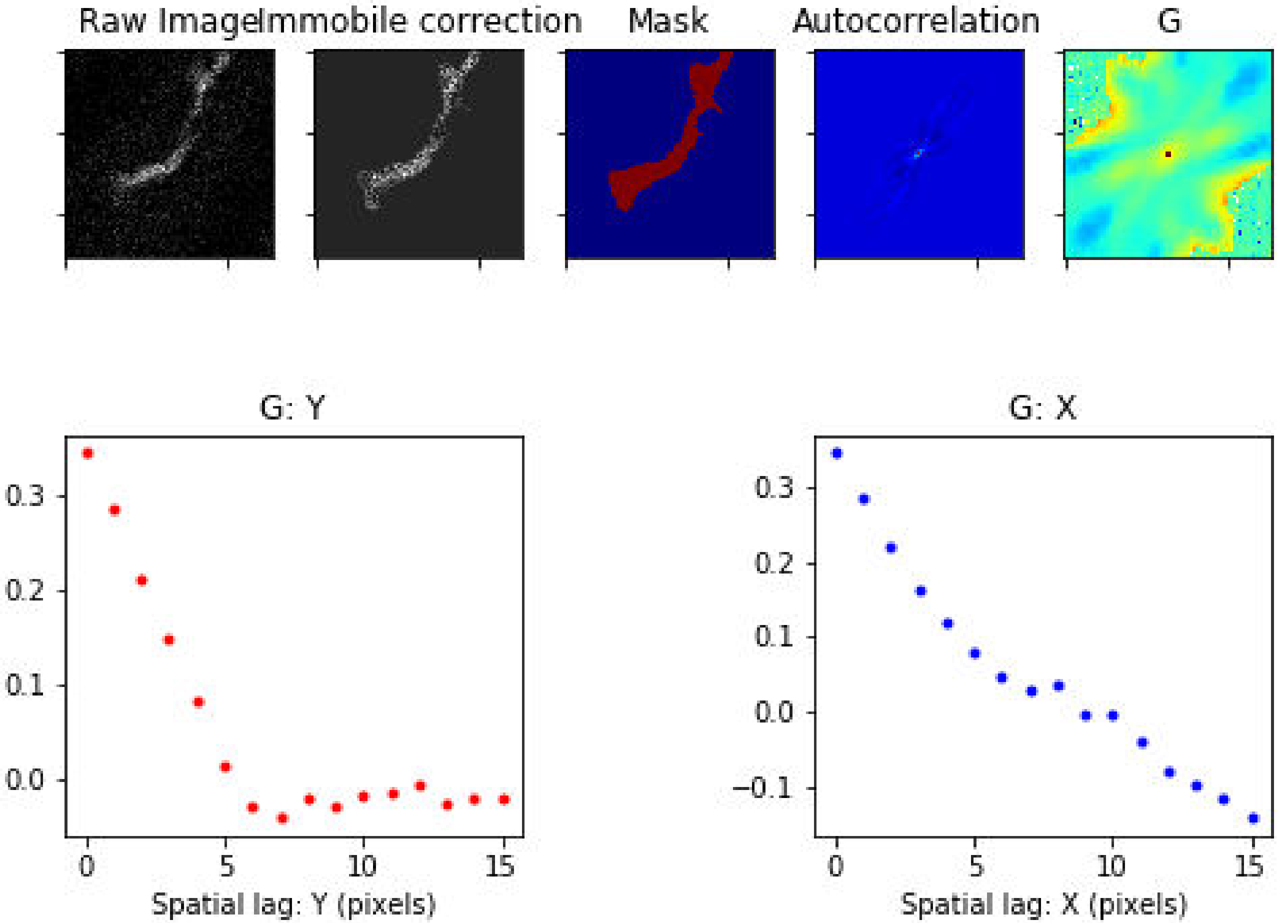

Treatment: Basal , Domain: MZ , File: Image 19.czi

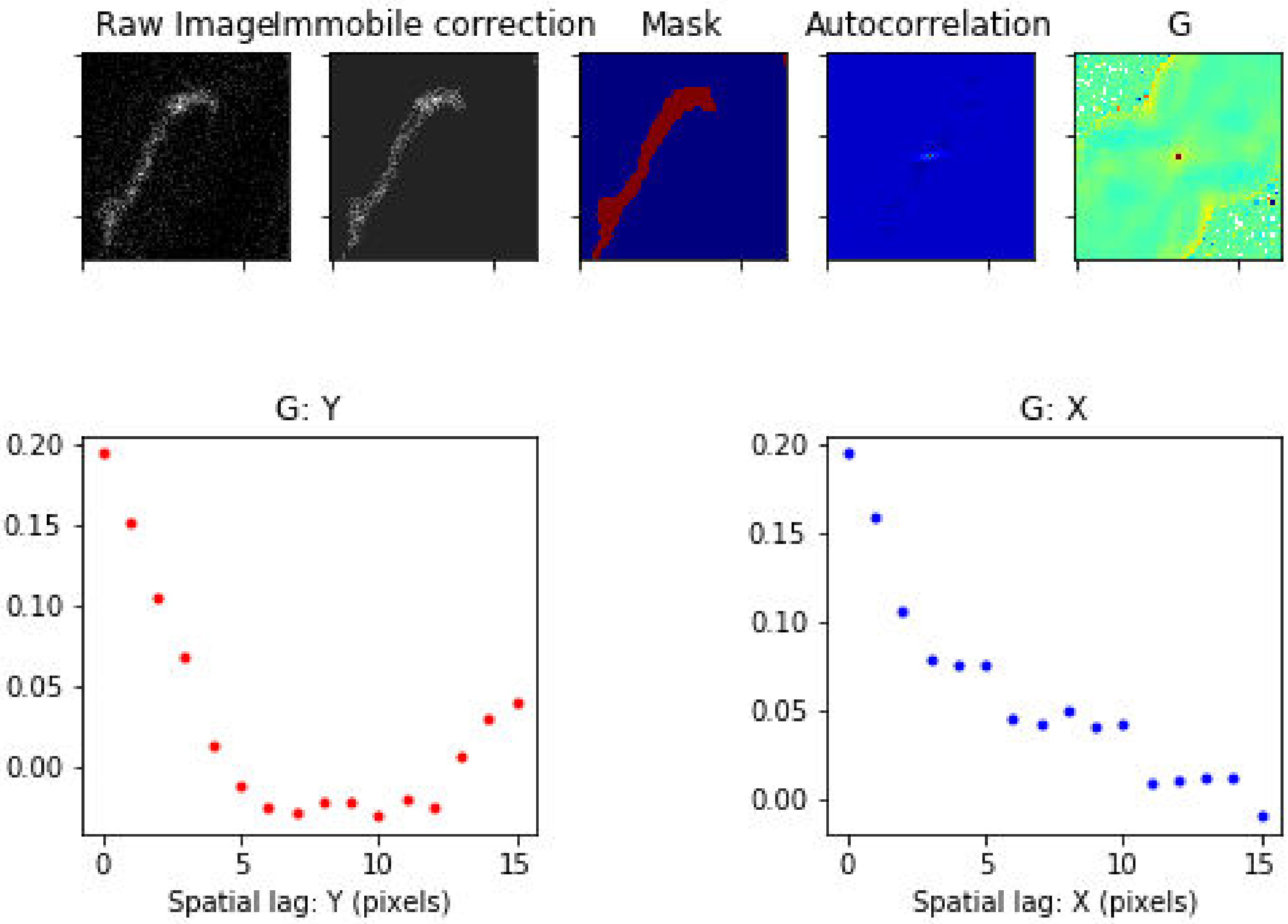

Treatment: Basal , Domain: MZ , File: Image 20.czi

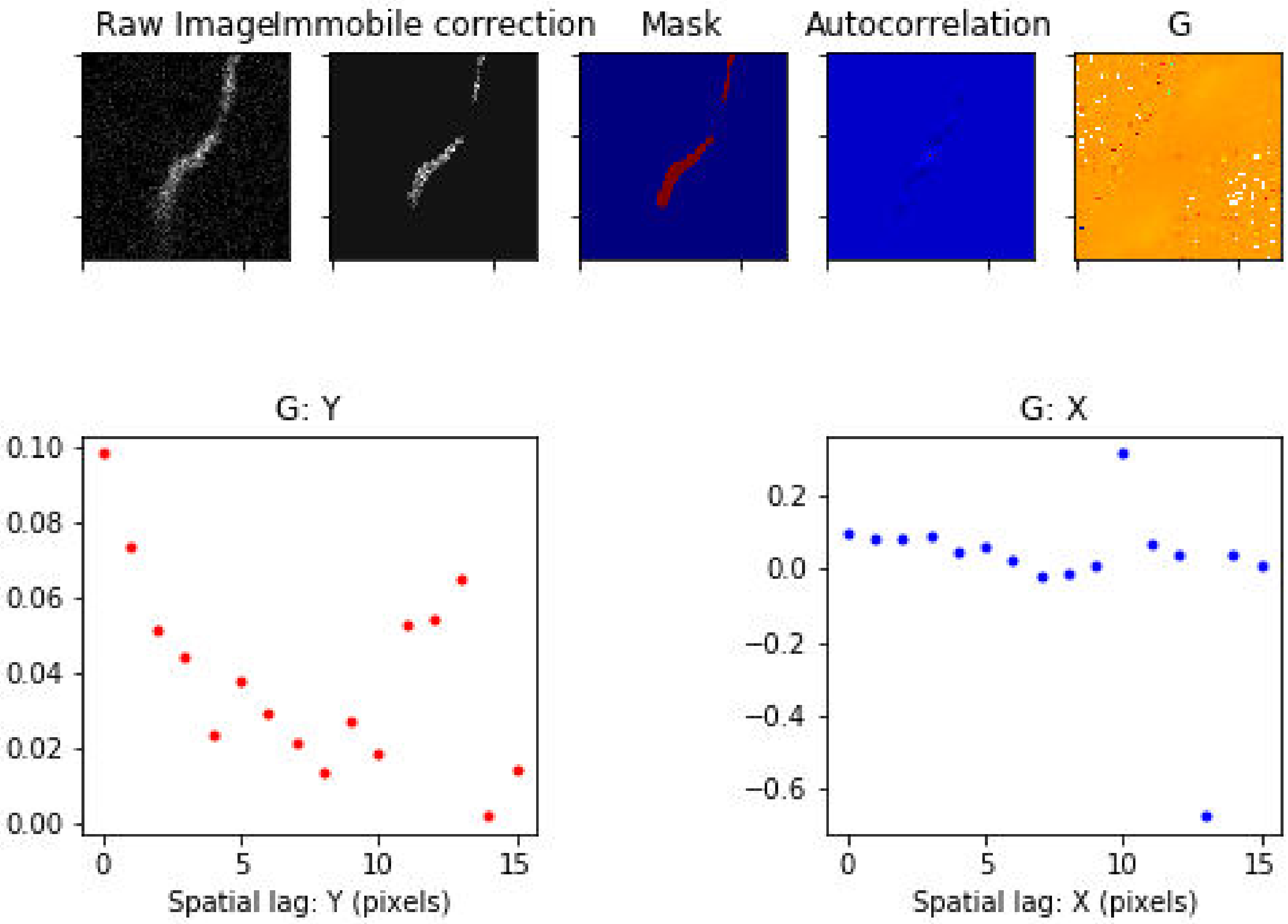

Treatment: Basal , Domain: MZ , File: Image 21.czi

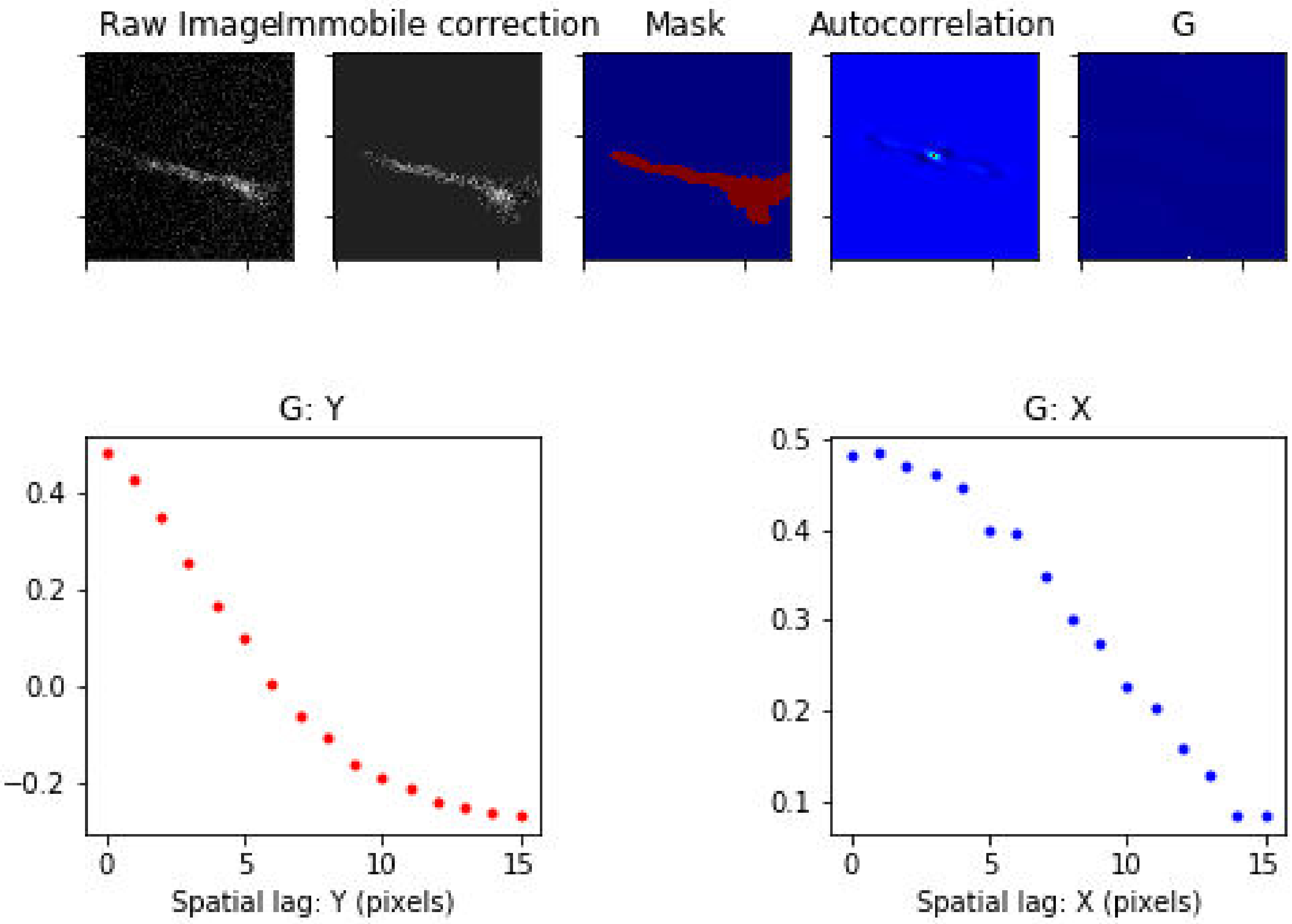

Treatment: Basal , Domain: MZ , File: Image 22.czi

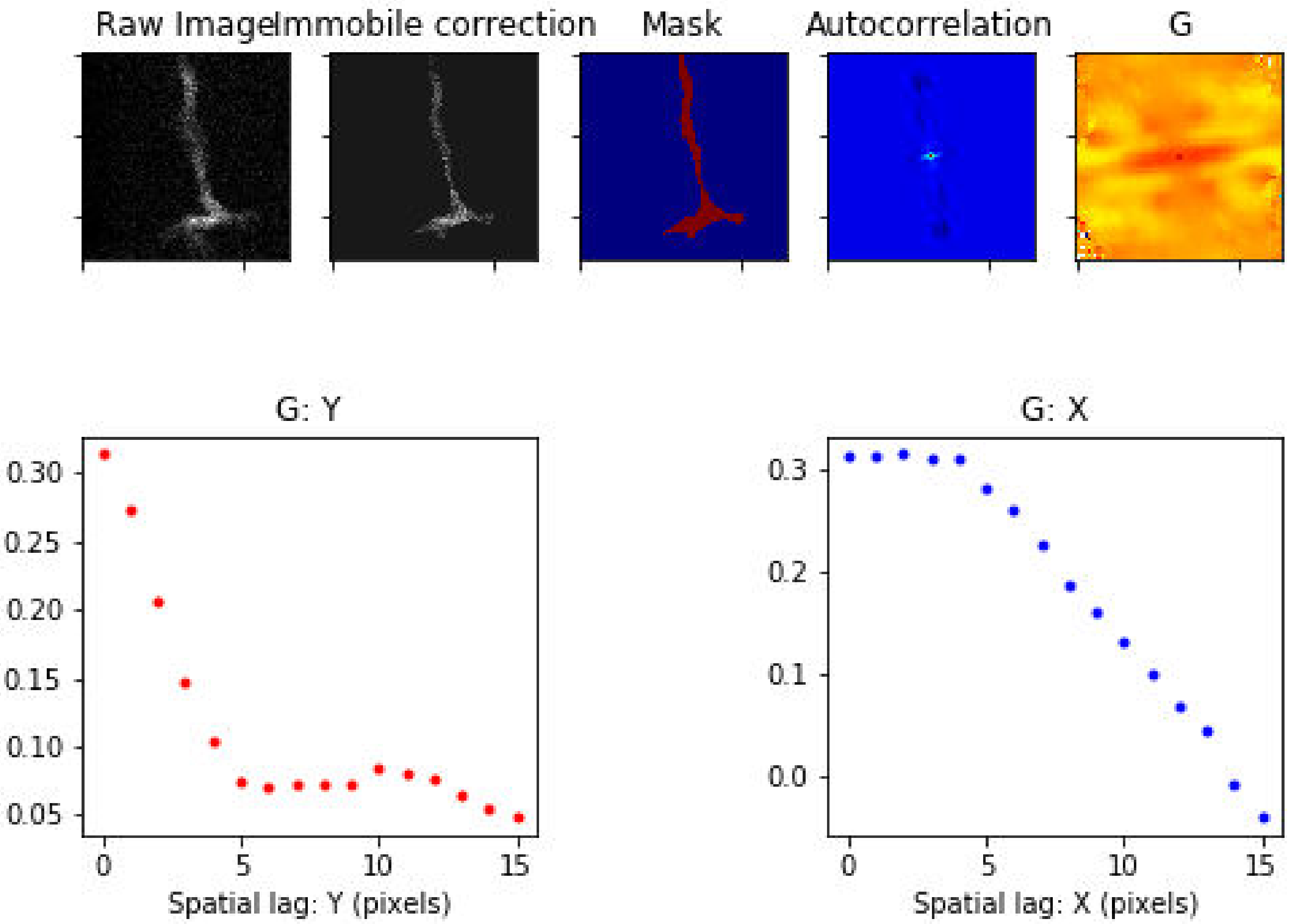

### Treatment: Basal , Domain: MZ , File: Image 23.czi

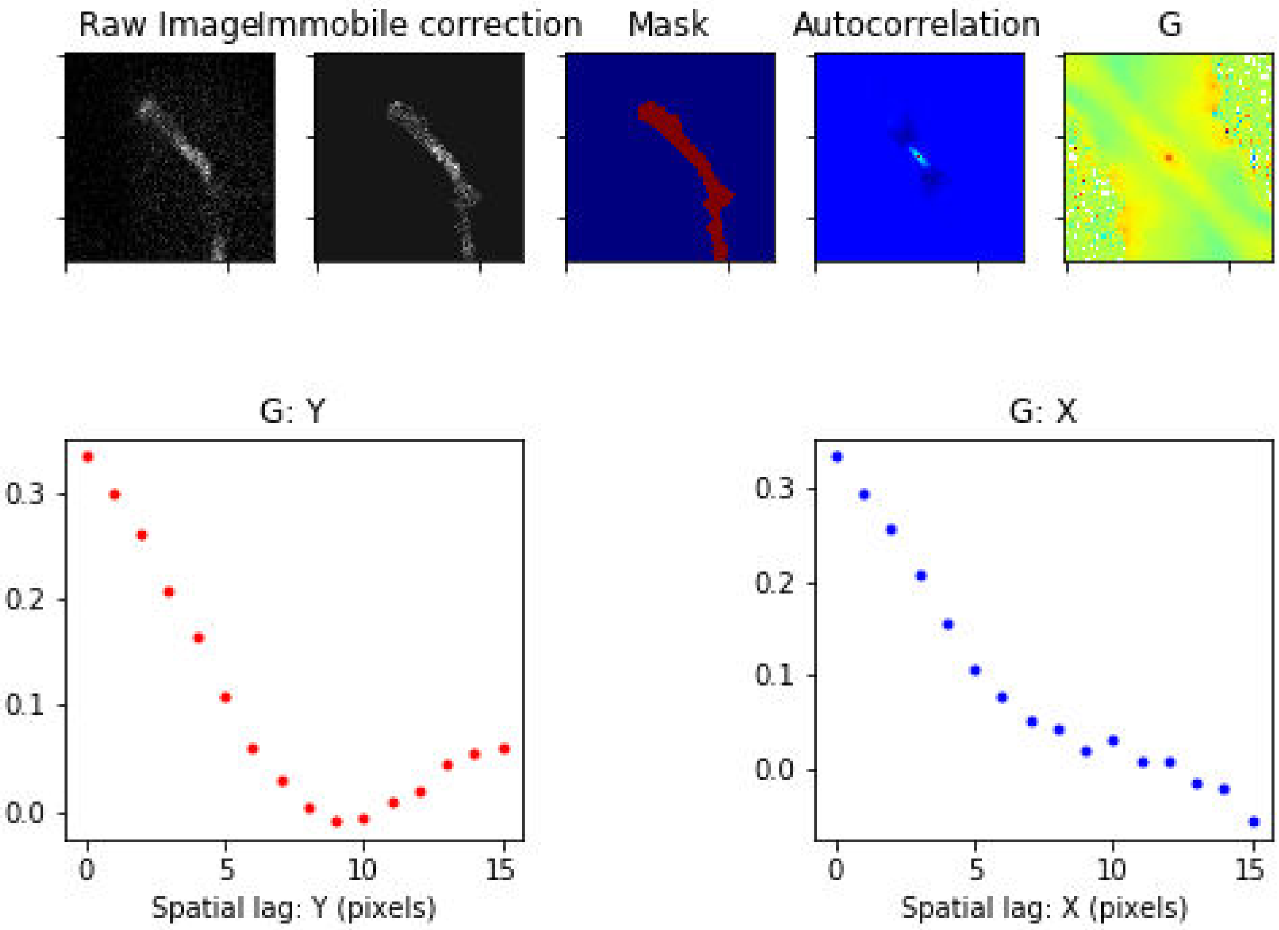

### Treatment: Basal , Domain: MZ , File: Image 24.czi

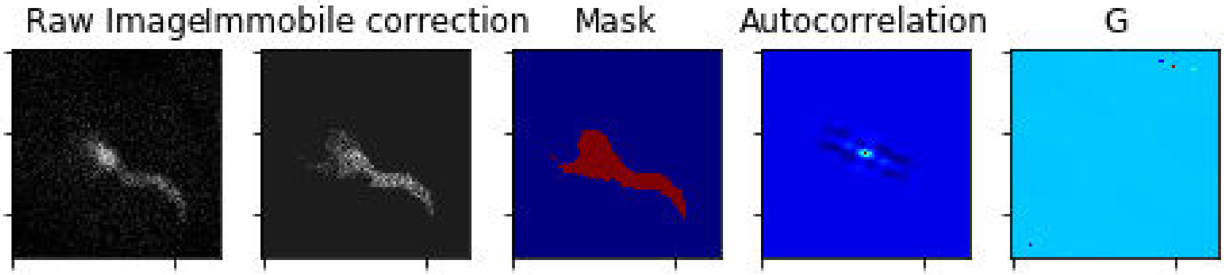


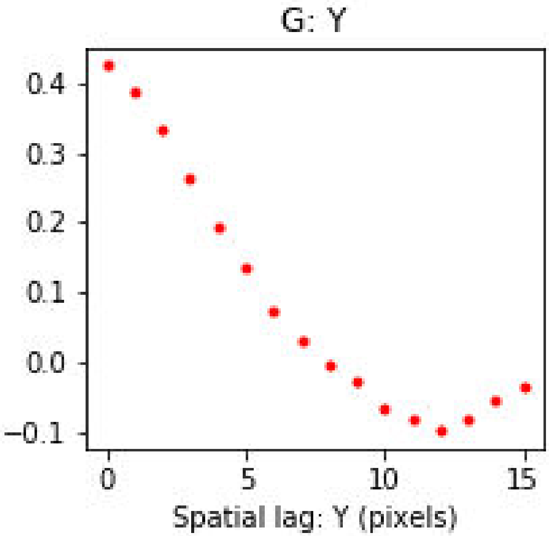


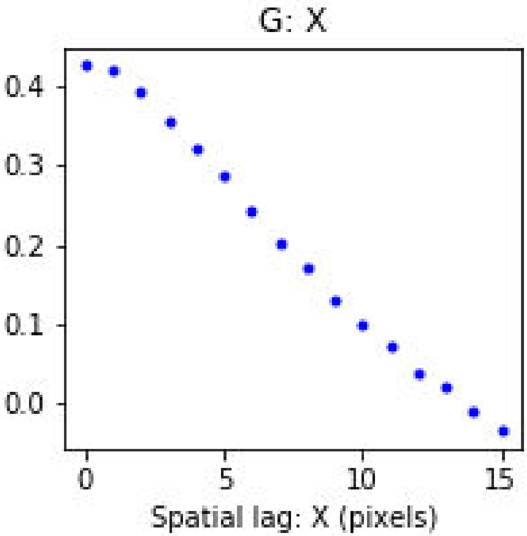

Treatment: Basal , Domain: MZ , File: Image 25.czi

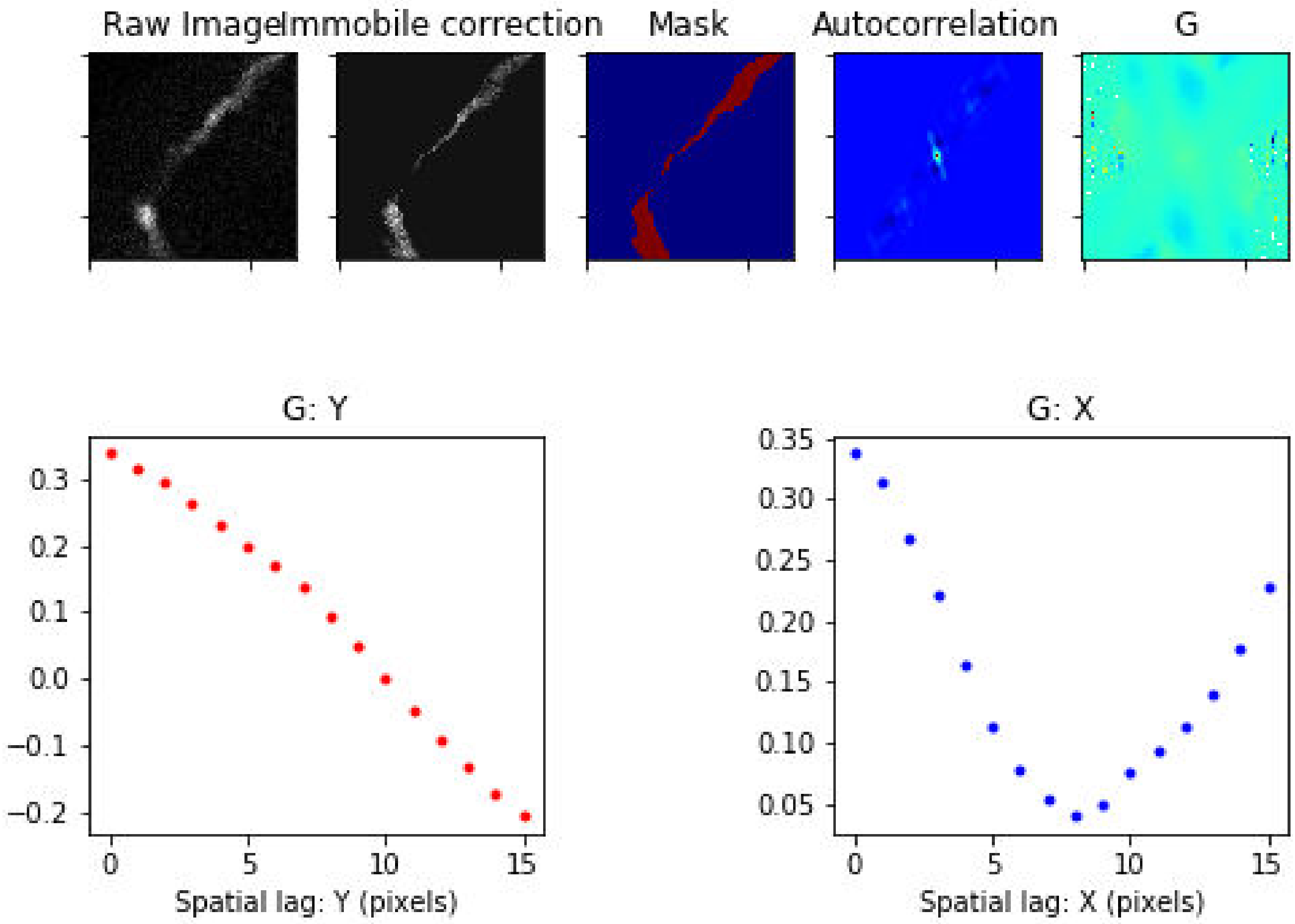

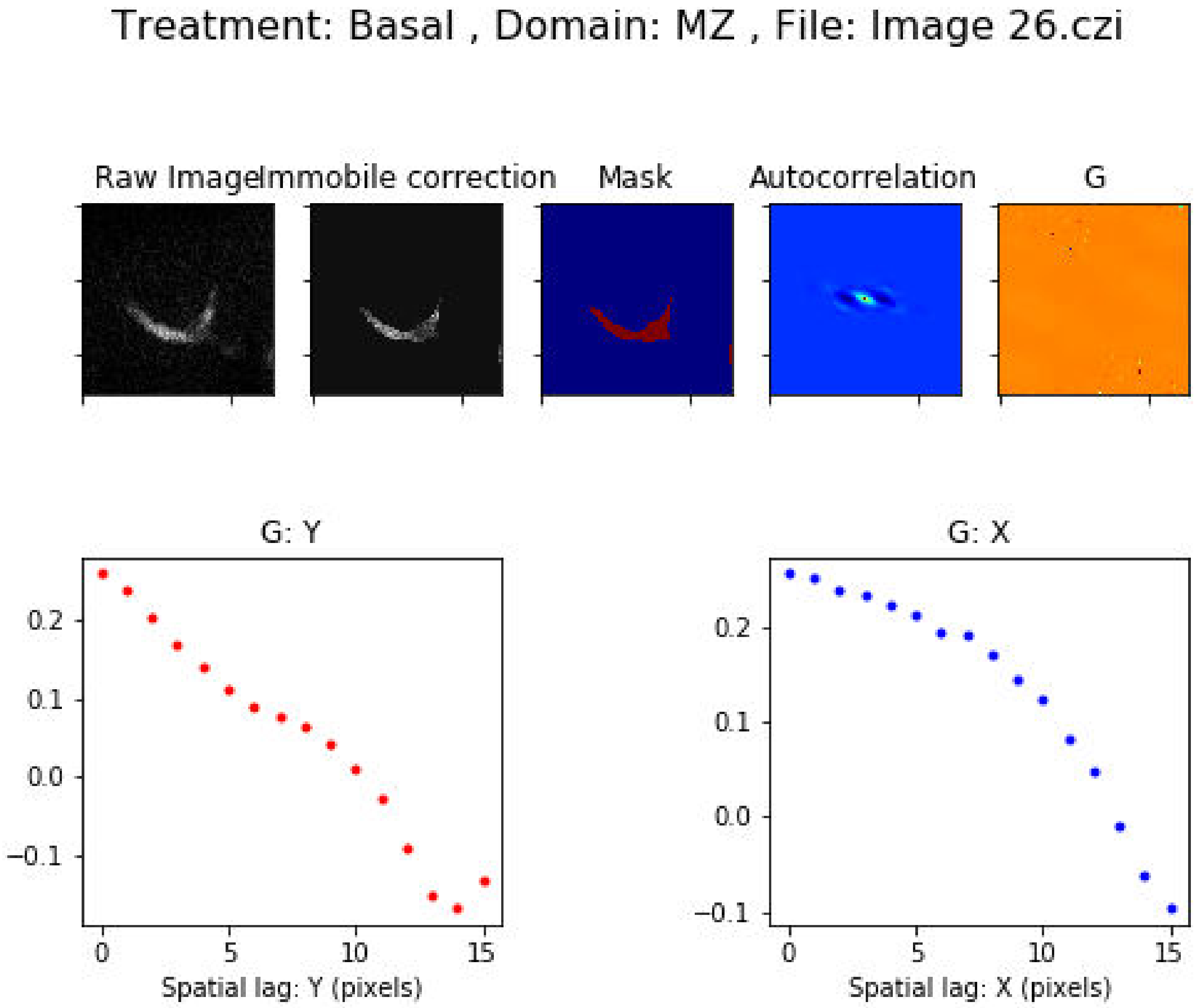
Treatment: Basal , Domain: MZ , File: Image 26.czi
Raw Image
Immobile correction
Mask
Autocorrelation
G
G: Y
0.2
0.1
0.0
−0.1
0
5
10
15
Spatial lag: Y (pixels)
G: X
0.2
0.1
0.0
−0.1
0
5
10
15
Spatial lag: X (pixels)

### Treatment: Basal , Domain: MZ , File: Image 42.czi

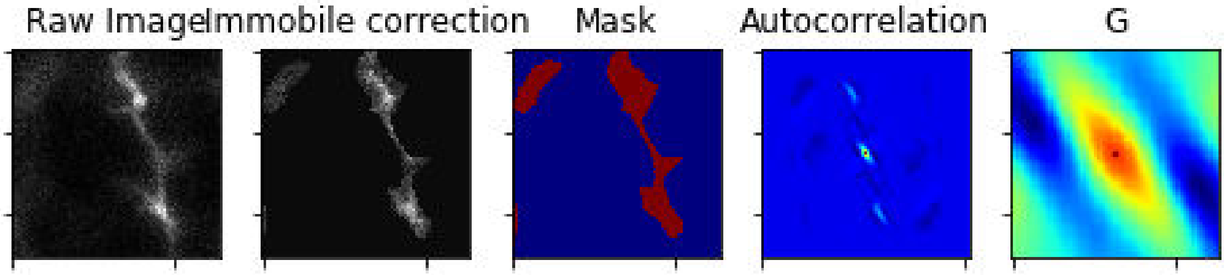


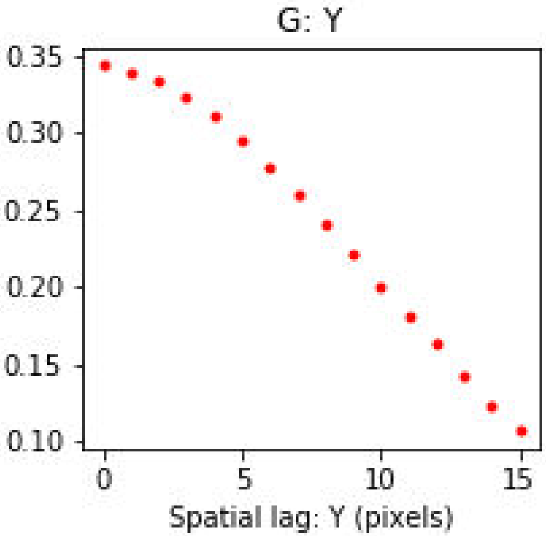


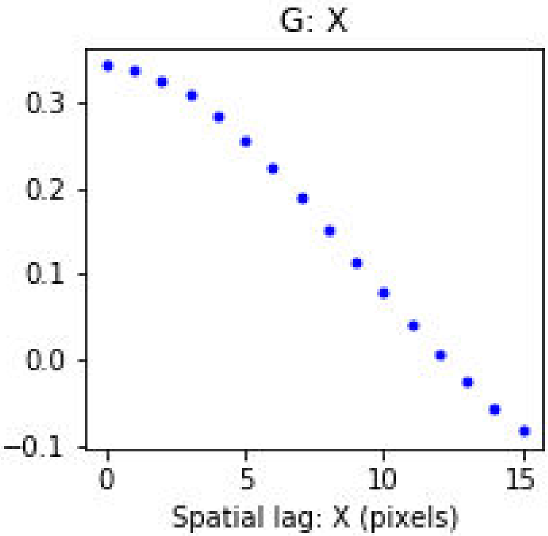

Treatment: Basal , Domain: PV , File: Image 4.czi

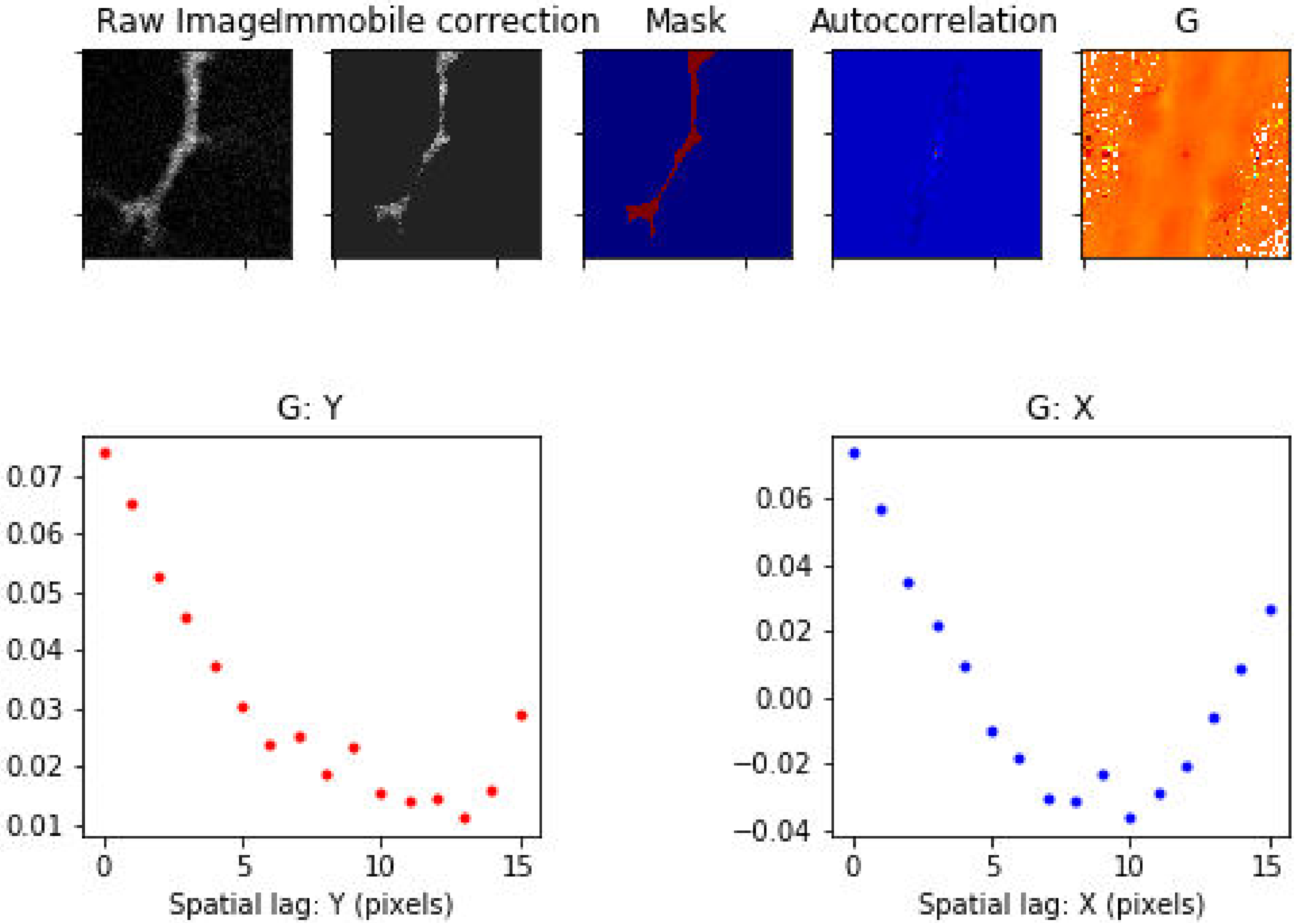

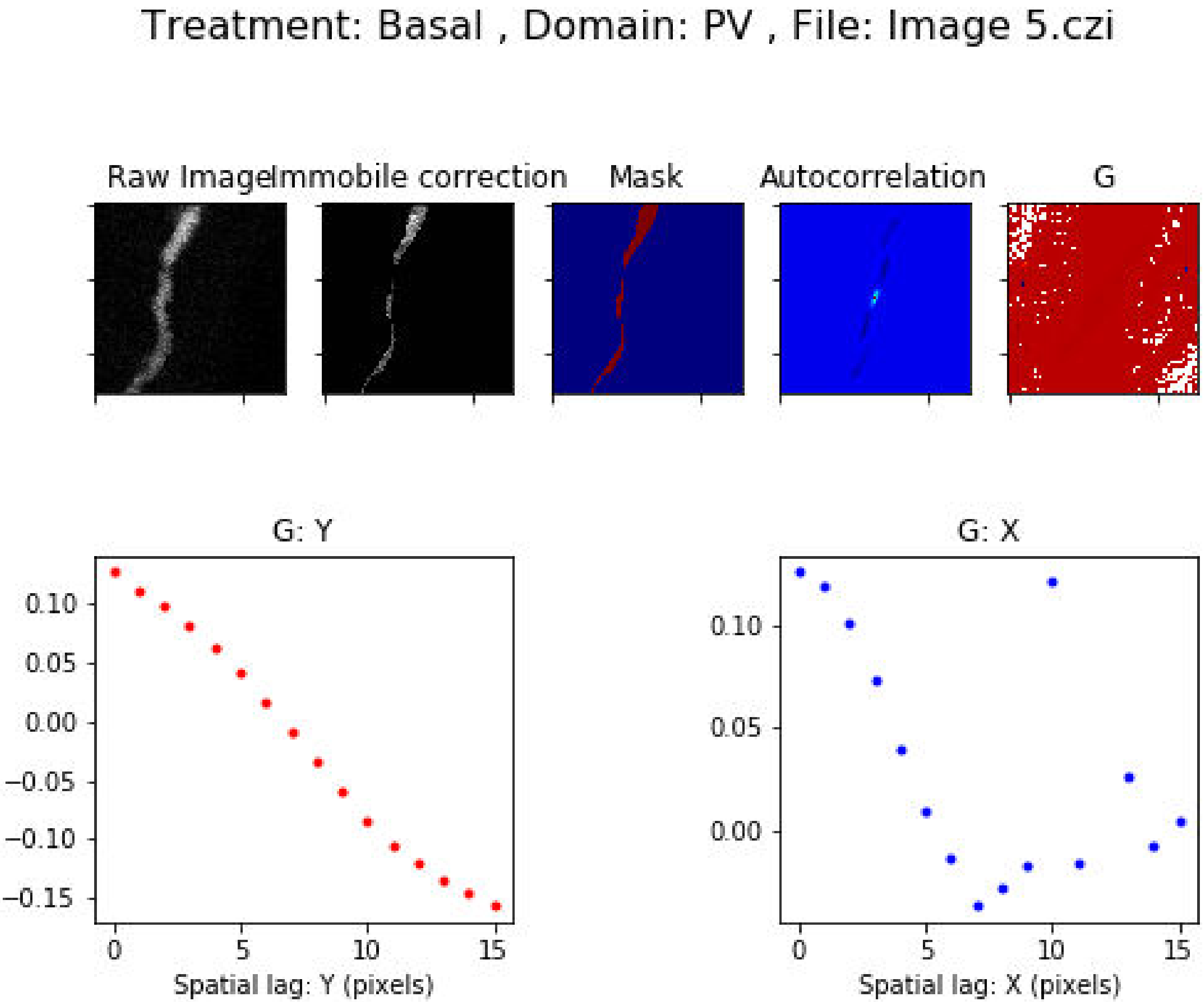

Treatment: Basal , Domain: PV , File: Image 5.czi
Raw Image
Immobile correction
Mask
Autocorrelation
G
G: Y
0.10
0.05
0.00
−0.05
−0.10
−0.15
0
5
10
15
Spatial lag: Y (pixels)
G: X
0.10
0.05
0.00
0
5
10
15
Spatial lag: X (pixels)

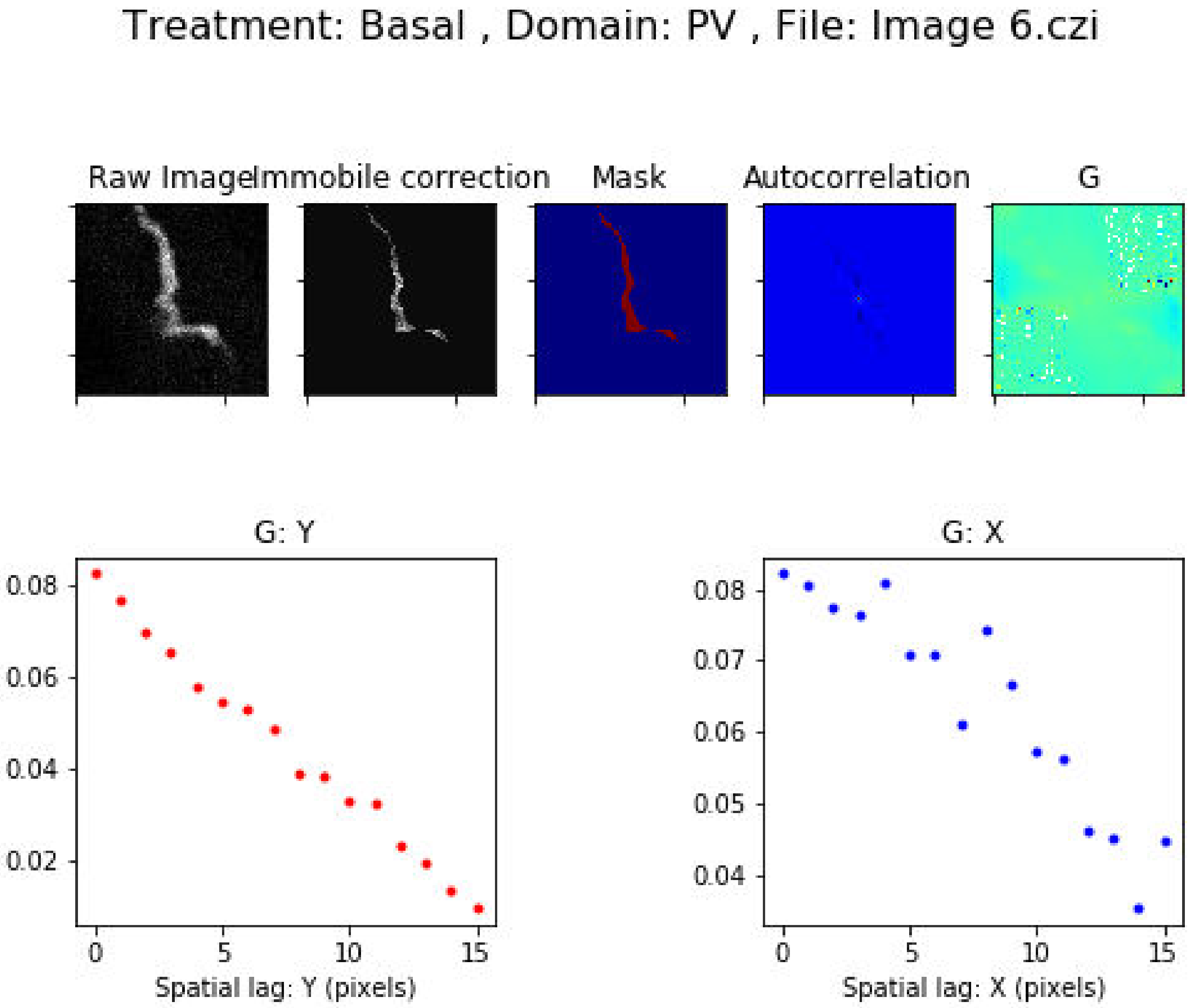
Treatment: Basal , Domain: PV , File: Image 6.czi
Raw Image
Immobile correction
Mask
Autocorrelation
G
G: Y
0.08
0.06
0.04
0.02
0
5
10
15
Spatial lag: Y (pixels)
G: X
0.08
0.07
0.06
0.05
0.04
0
5
10
15
Spatial lag: X (pixels)

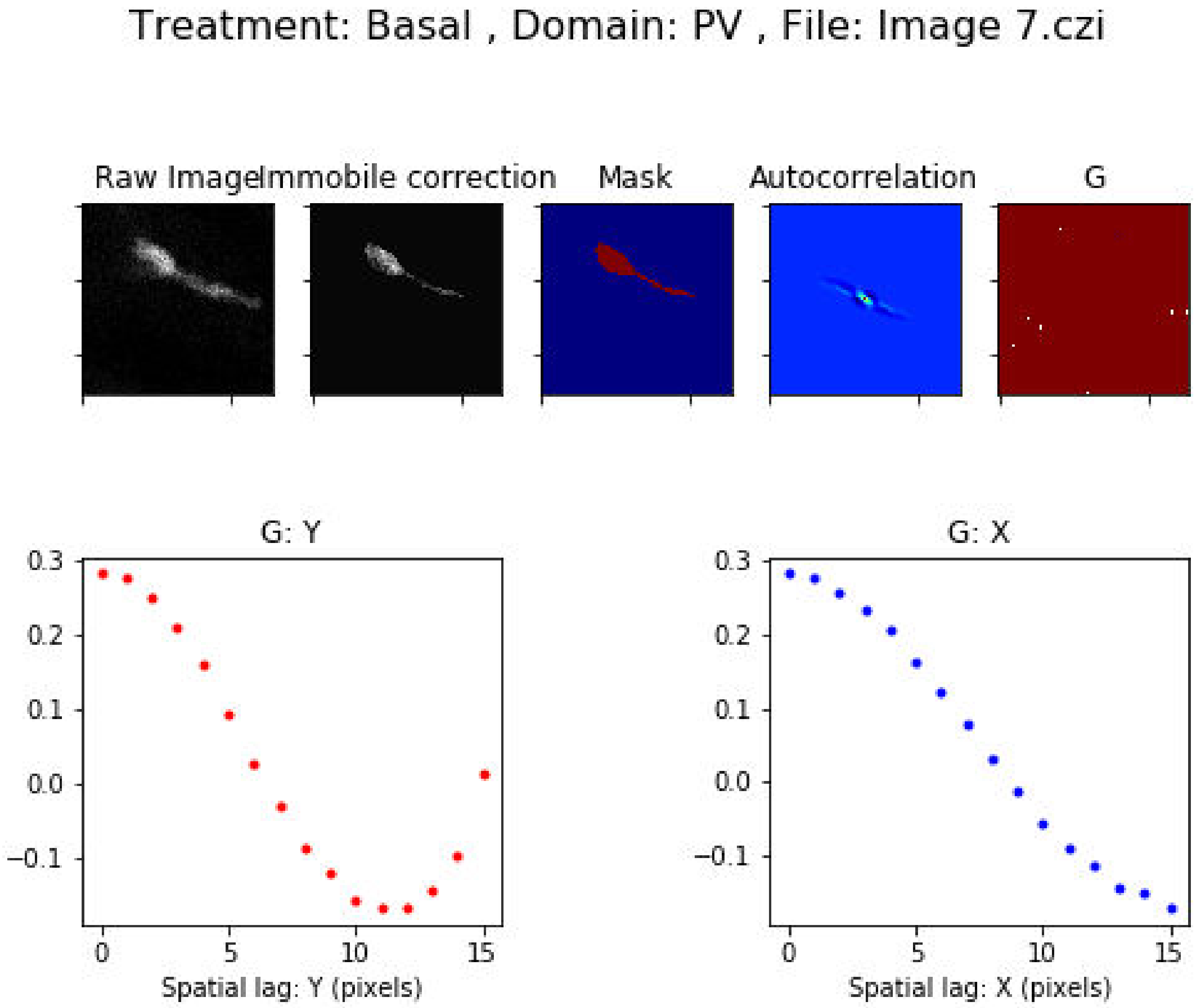

Treatment: Basal , Domain: PV , File: Image 7.czi
Raw Image
Immobile correction
Mask
Autocorrelation
G
G: Y
0.3
0.2
0.1
0.0
−0.1
0
5
10
15
Spatial lag: Y (pixels)
G: X
0.3
0.2
0.1
0.0
−0.1
0
5
10
15
Spatial lag: X (pixels)

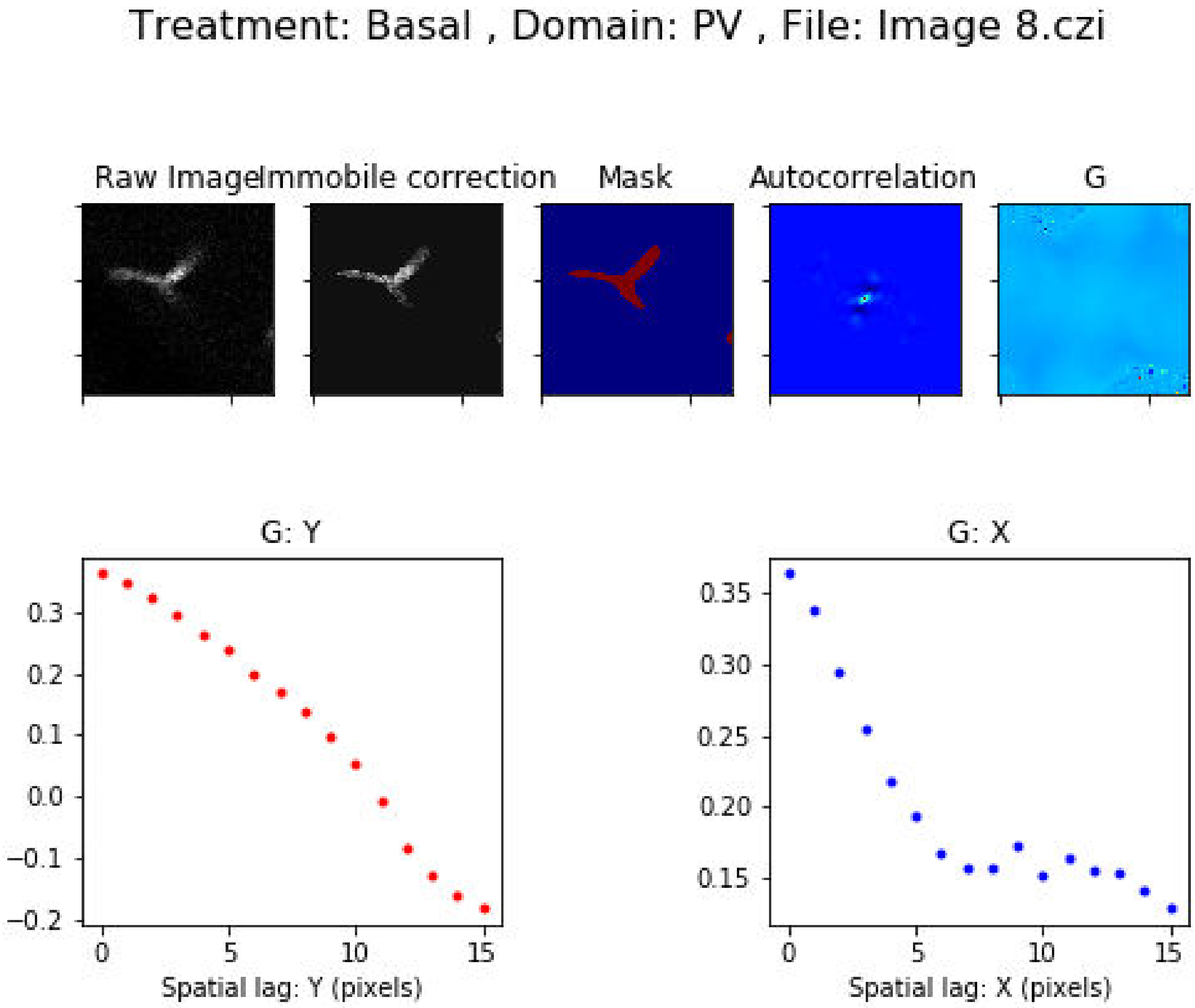
Treatment: Basal , Domain: PV , File: Image 8.czi
Raw Image
Immobile correction
Mask
Autocorrelation
G
G: Y
G: X
Spatial lag: Y (pixels)
Spatial lag: X (pixels)
0.3
0.2
0.1
0.0
−0.1
−0.2
0.35
0.30
0.25
0.20
0.15
0
5
10
15

Treatment: Basal , Domain: PV , File: Image 9.czi

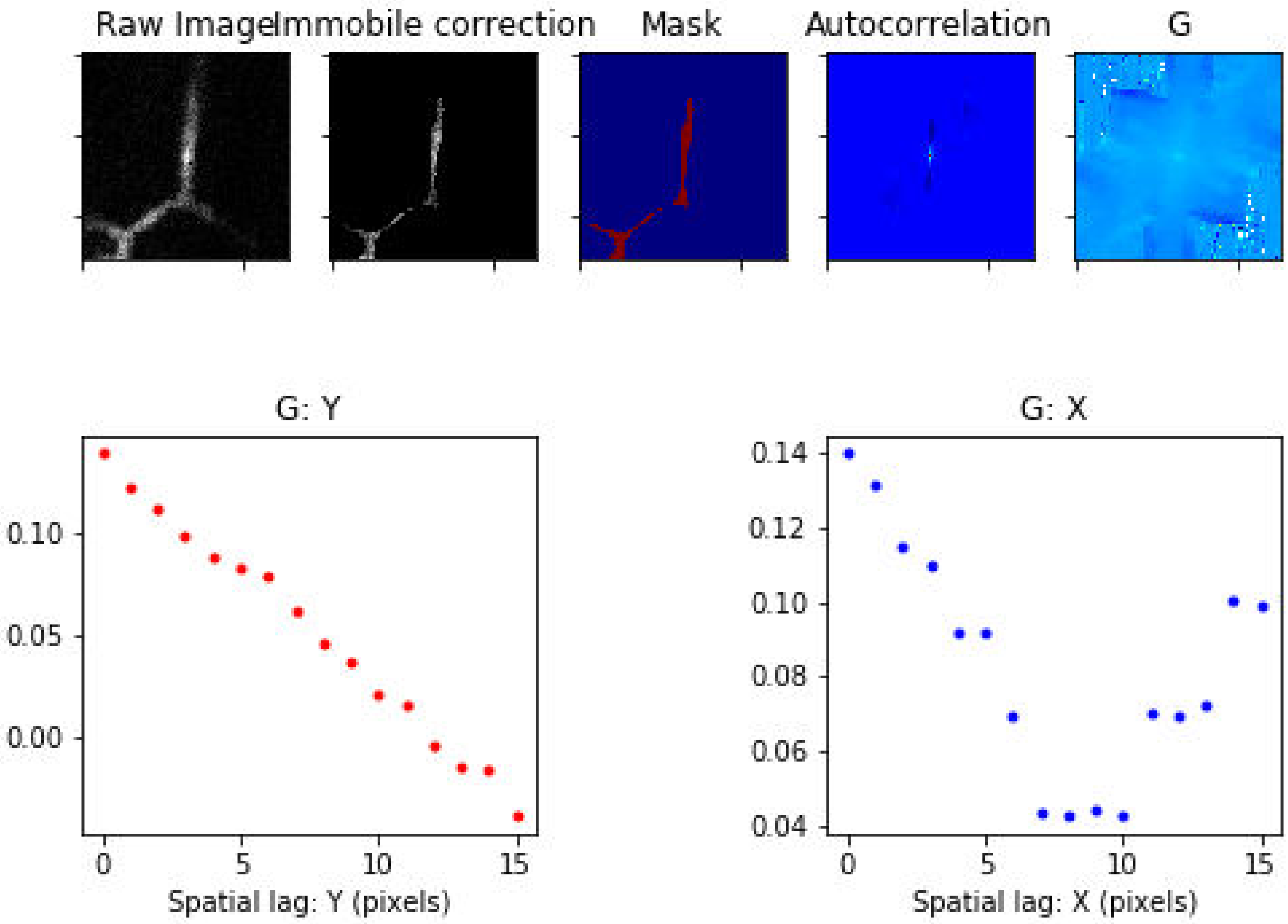

Treatment: Basal , Domain: PV , File: Image 10.czi

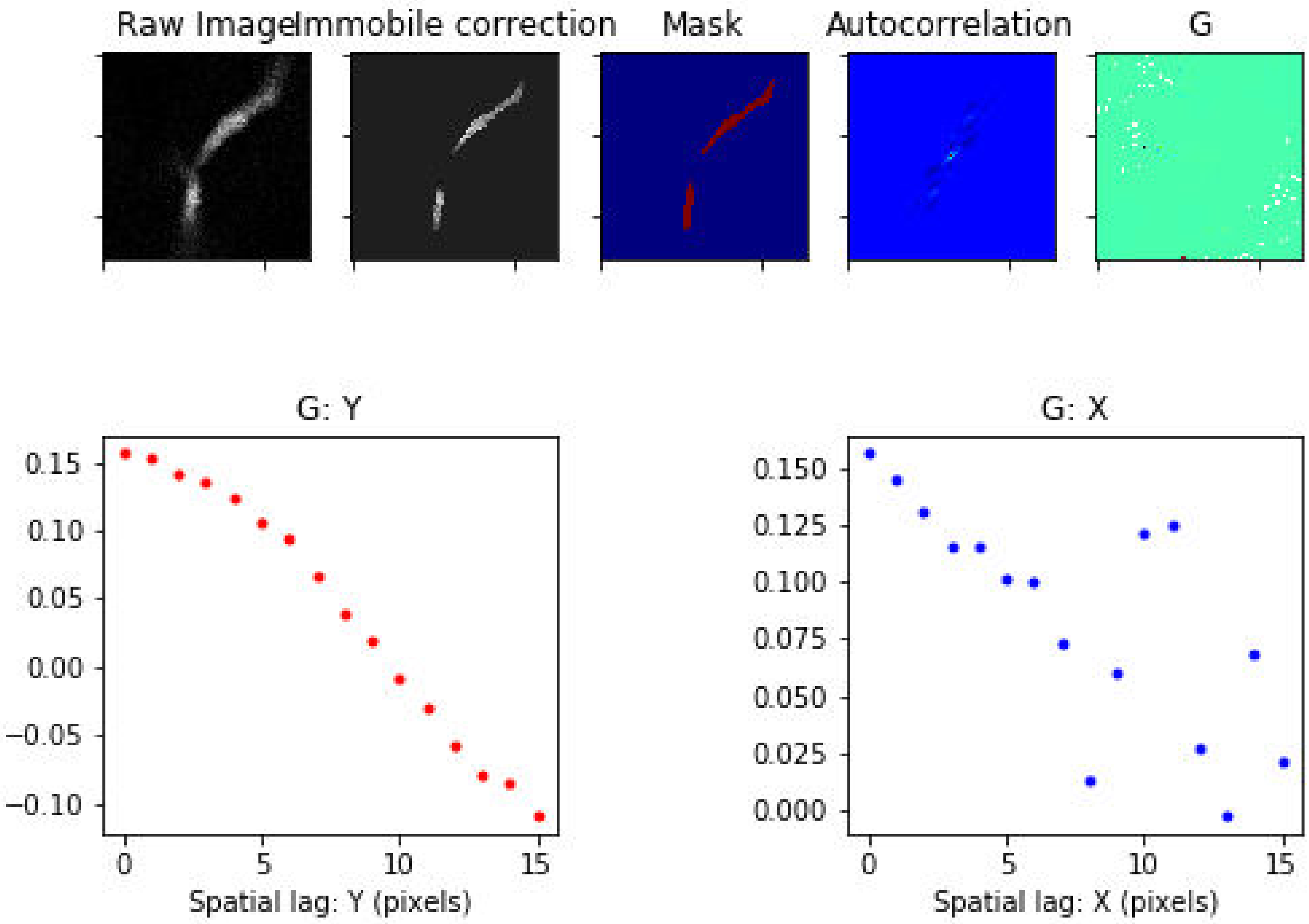

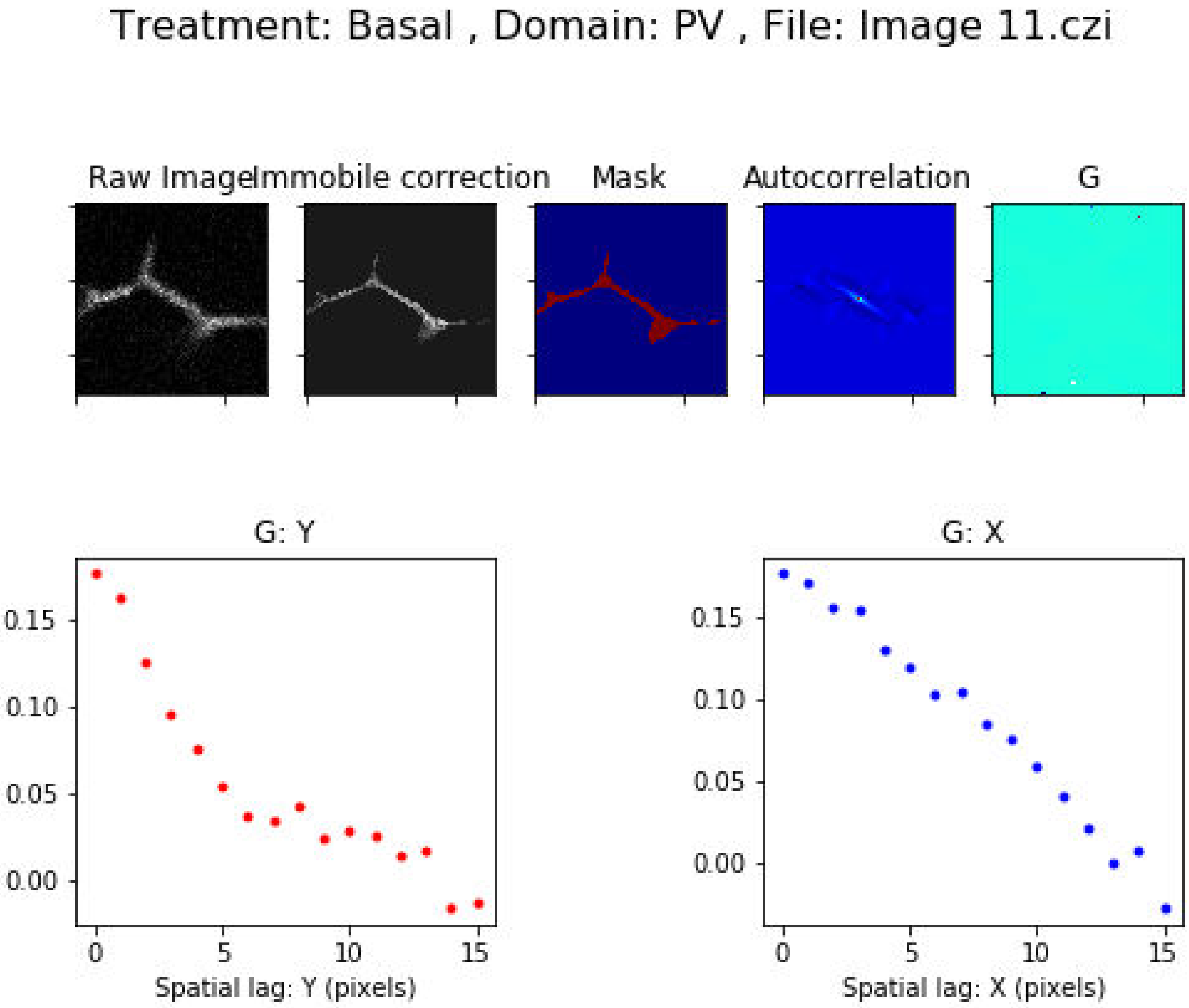
Treatment: Basal , Domain: PV , File: Image 11.czi
Raw Image
Immobile correction
Mask
Autocorrelation
G
G: Y
G: X
0.15
0.10
0.05
0.00
0
5
10
15
Spatial lag: Y (pixels)
Spatial lag: X (pixels)

Treatment: Basal , Domain: PV , File: Image 12.czi

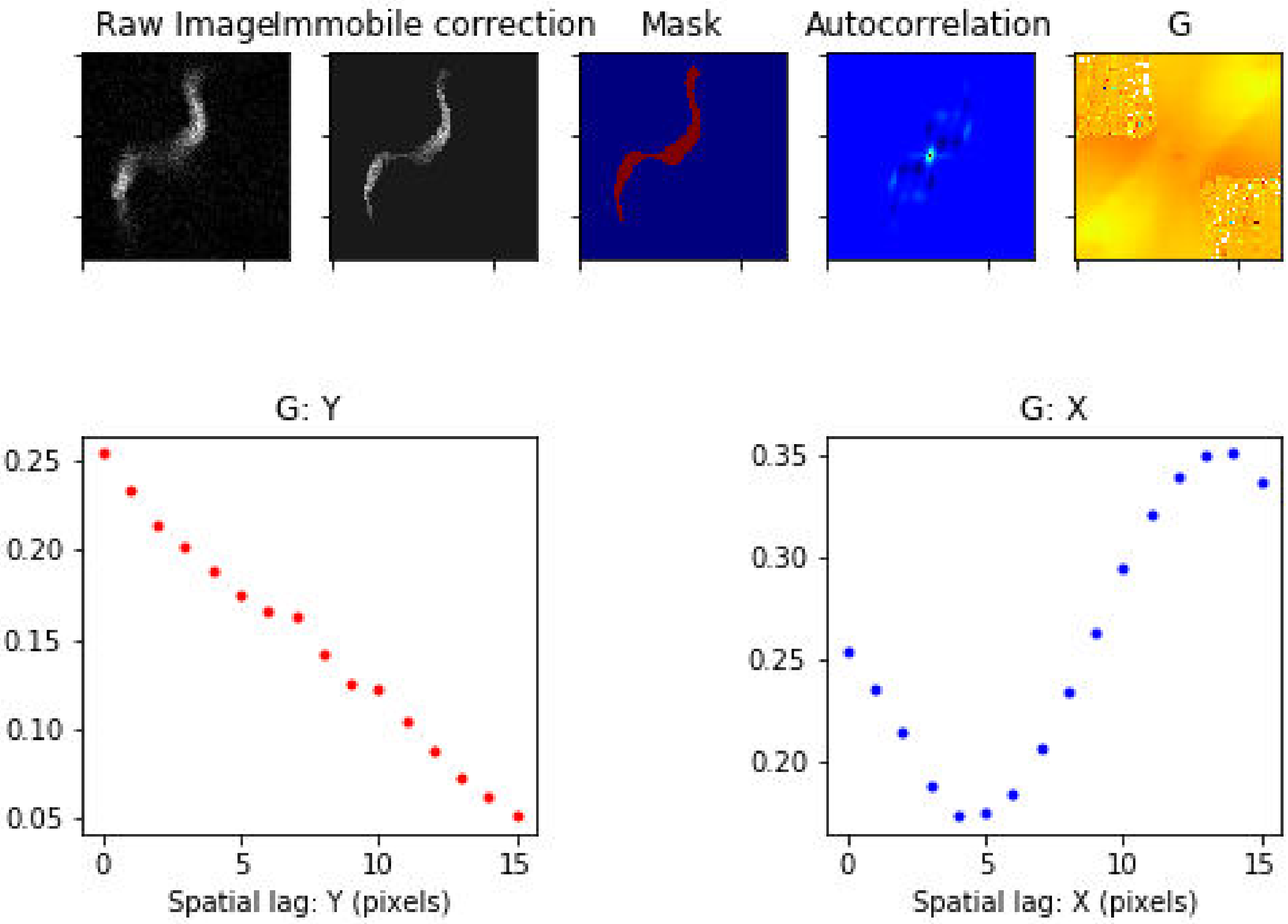

Treatment: Basal , Domain: PV , File: Image 13.czi

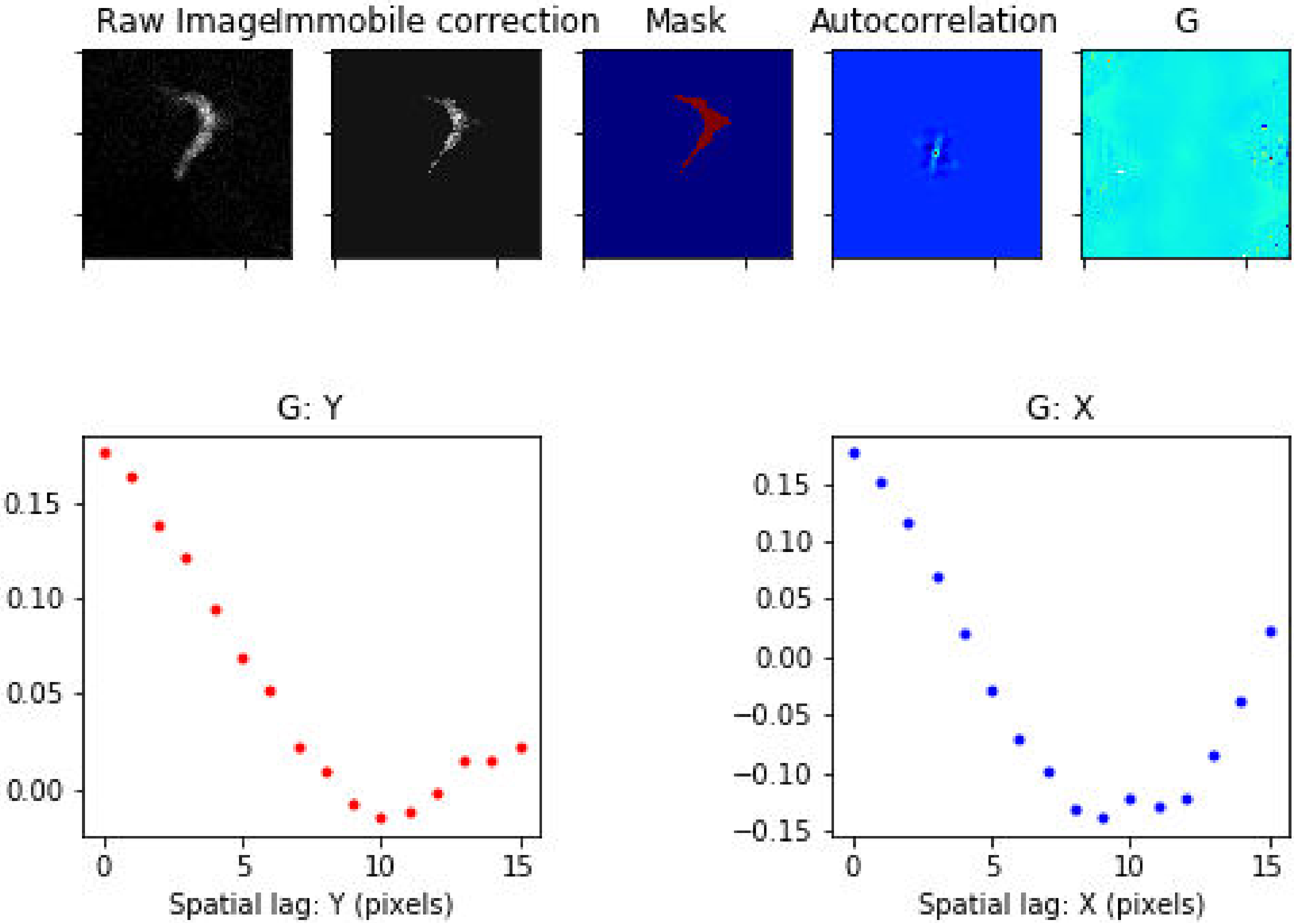

### Treatment: Basal , Domain: PV , File: Image 43.czi

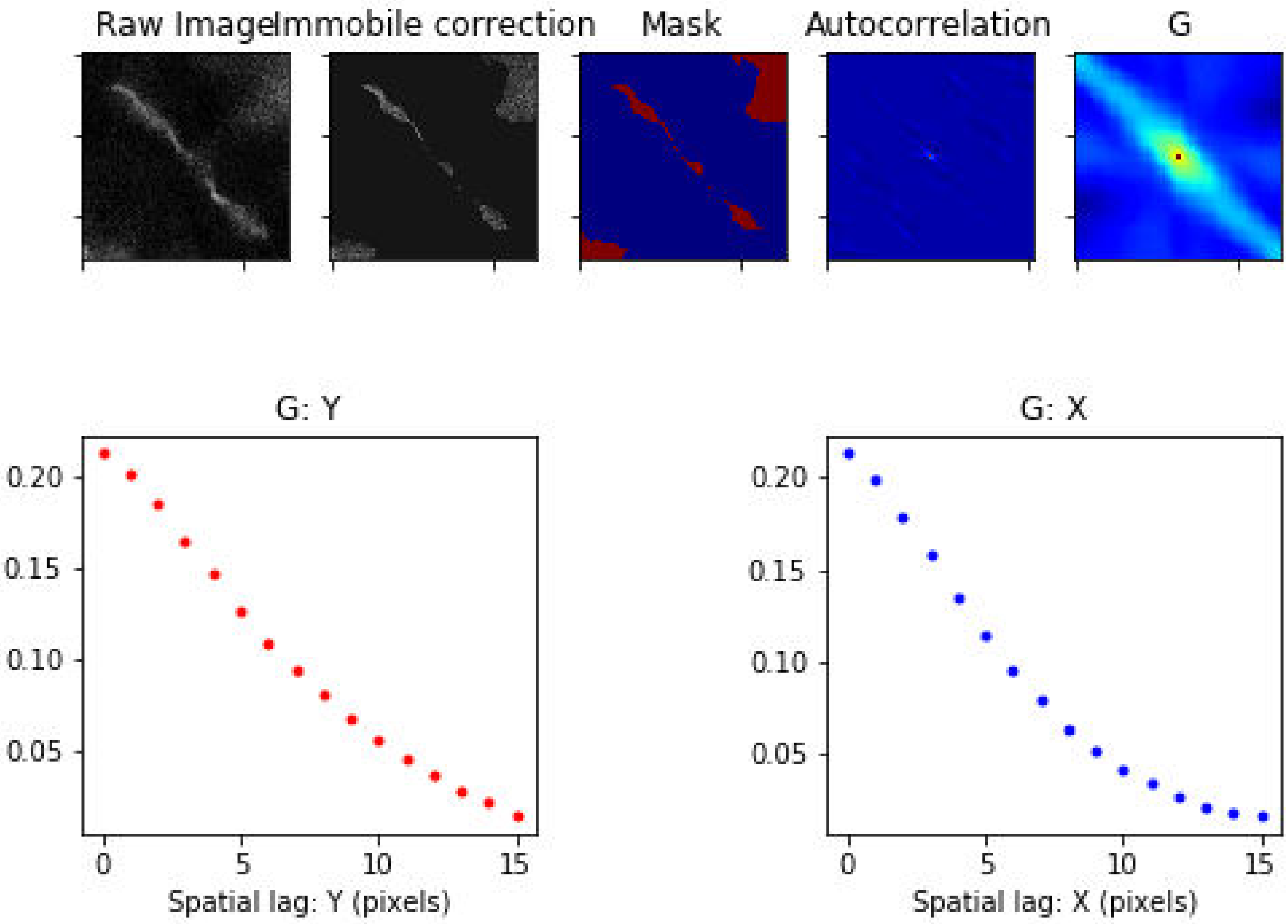

Treatment: Secretin , Domain: CV , File: Image 25.czi

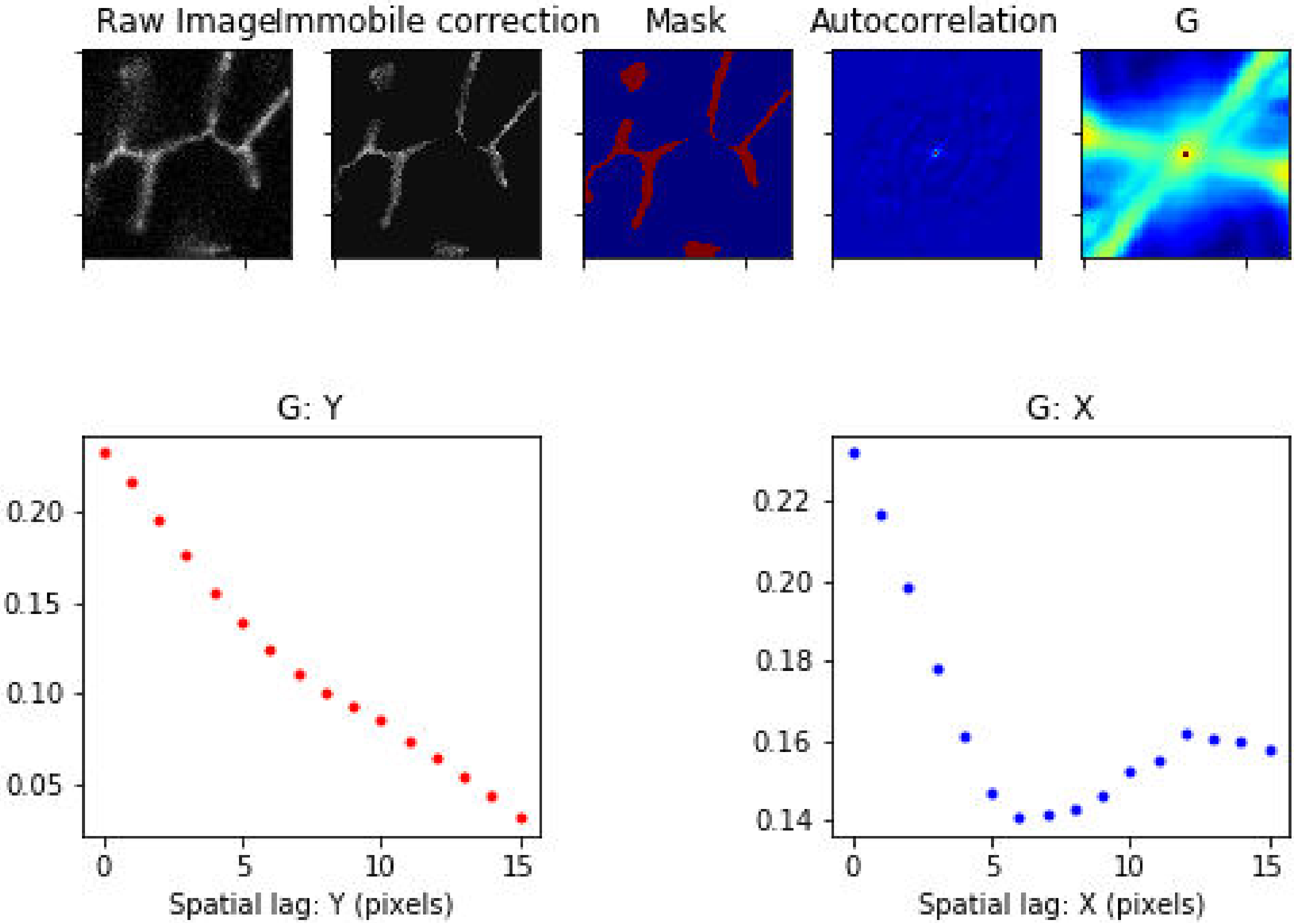

Treatment: Secretin , Domain: CV , File: Image 28.czi

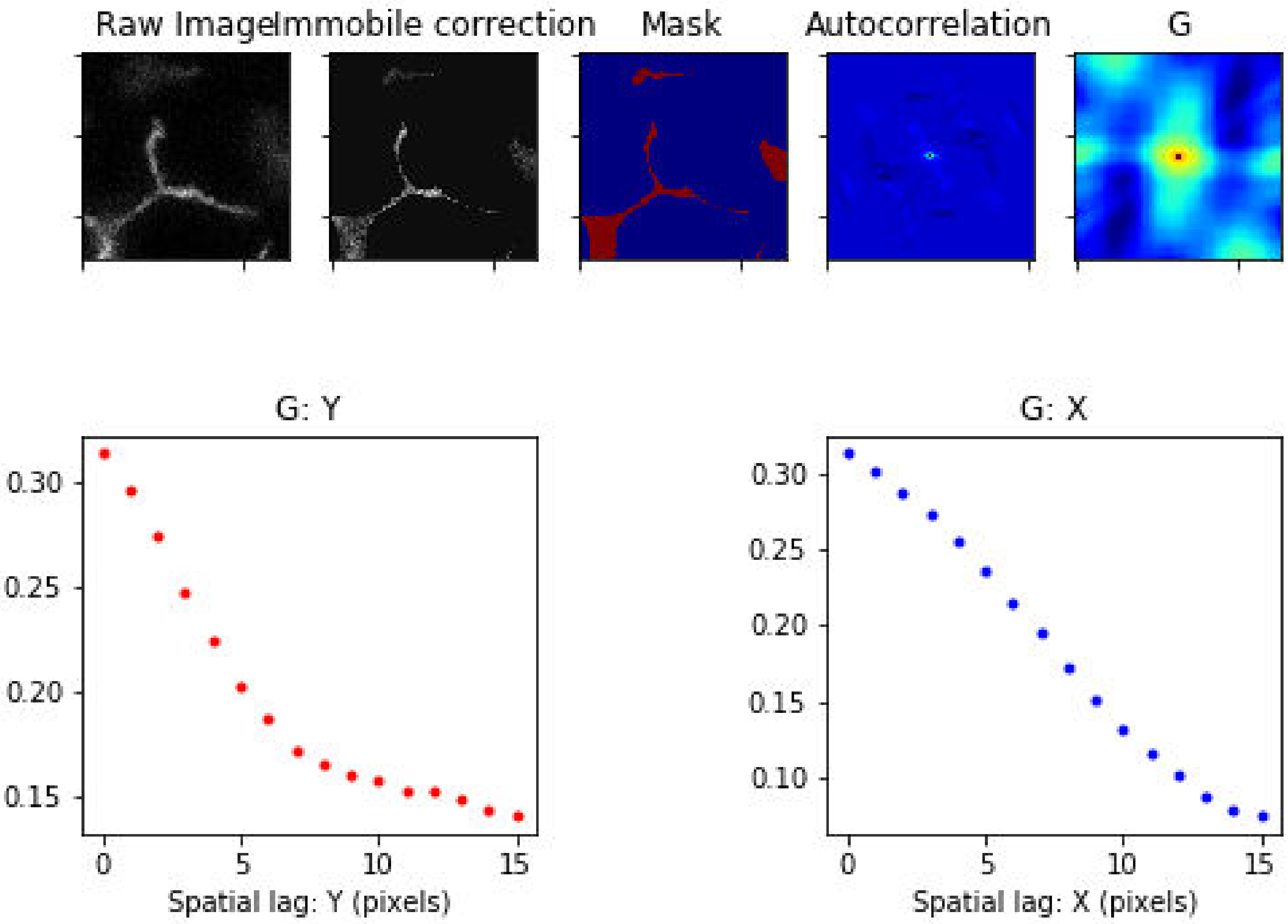

Treatment: Secretin , Domain: CV , File: Image 44.czi

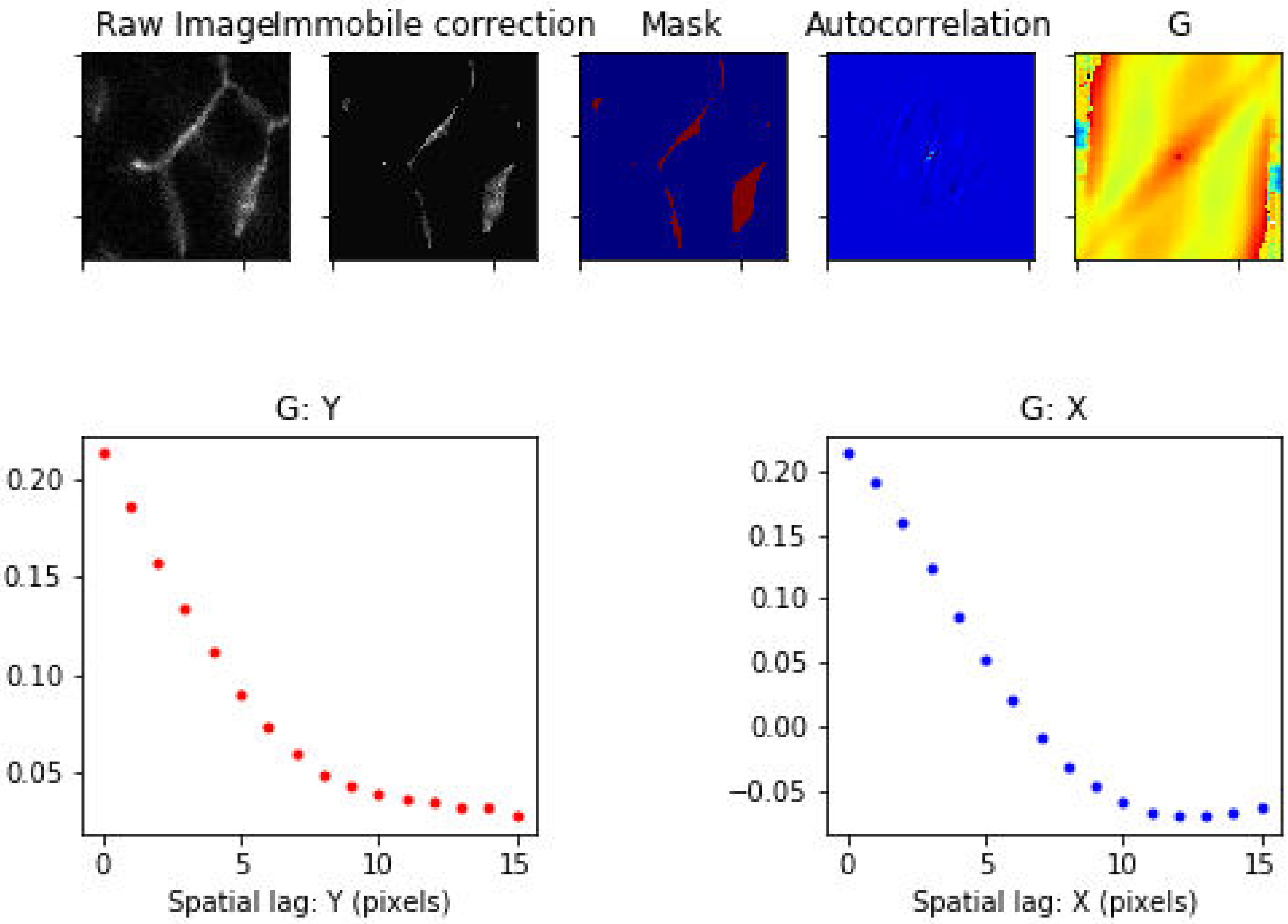

Treatment: Secretin , Domain: CV , File: Image 125.czi

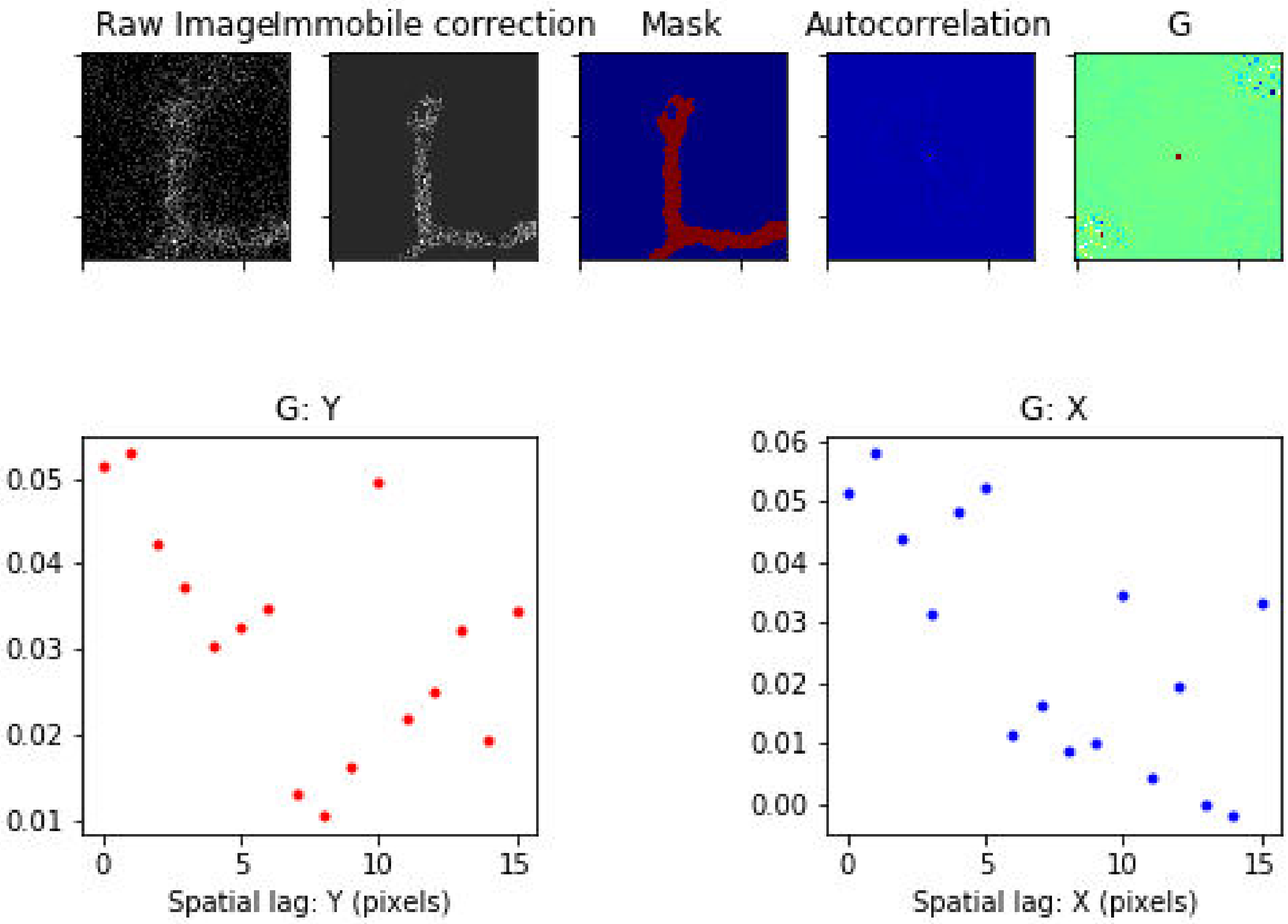

### Treatment: Secretin , Domain: CV , File: Image 128.czi

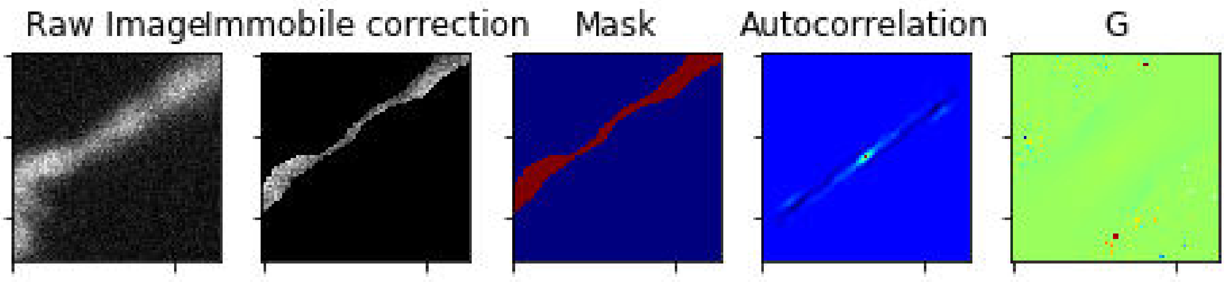


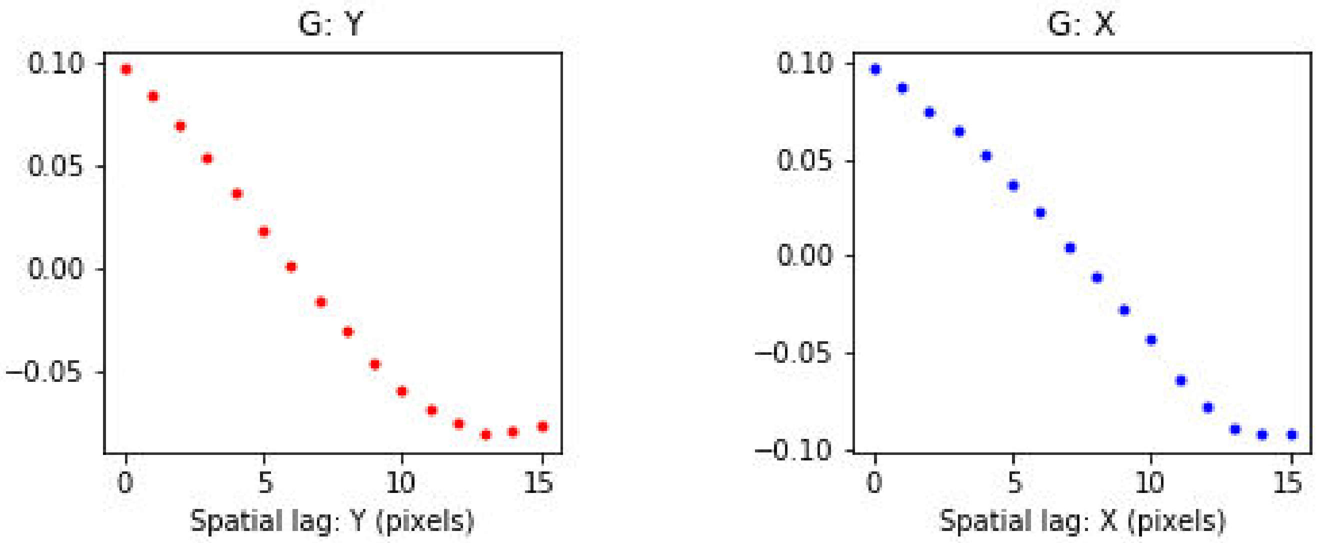

Treatment: Secretin , Domain: CV , File: Image 130.czi

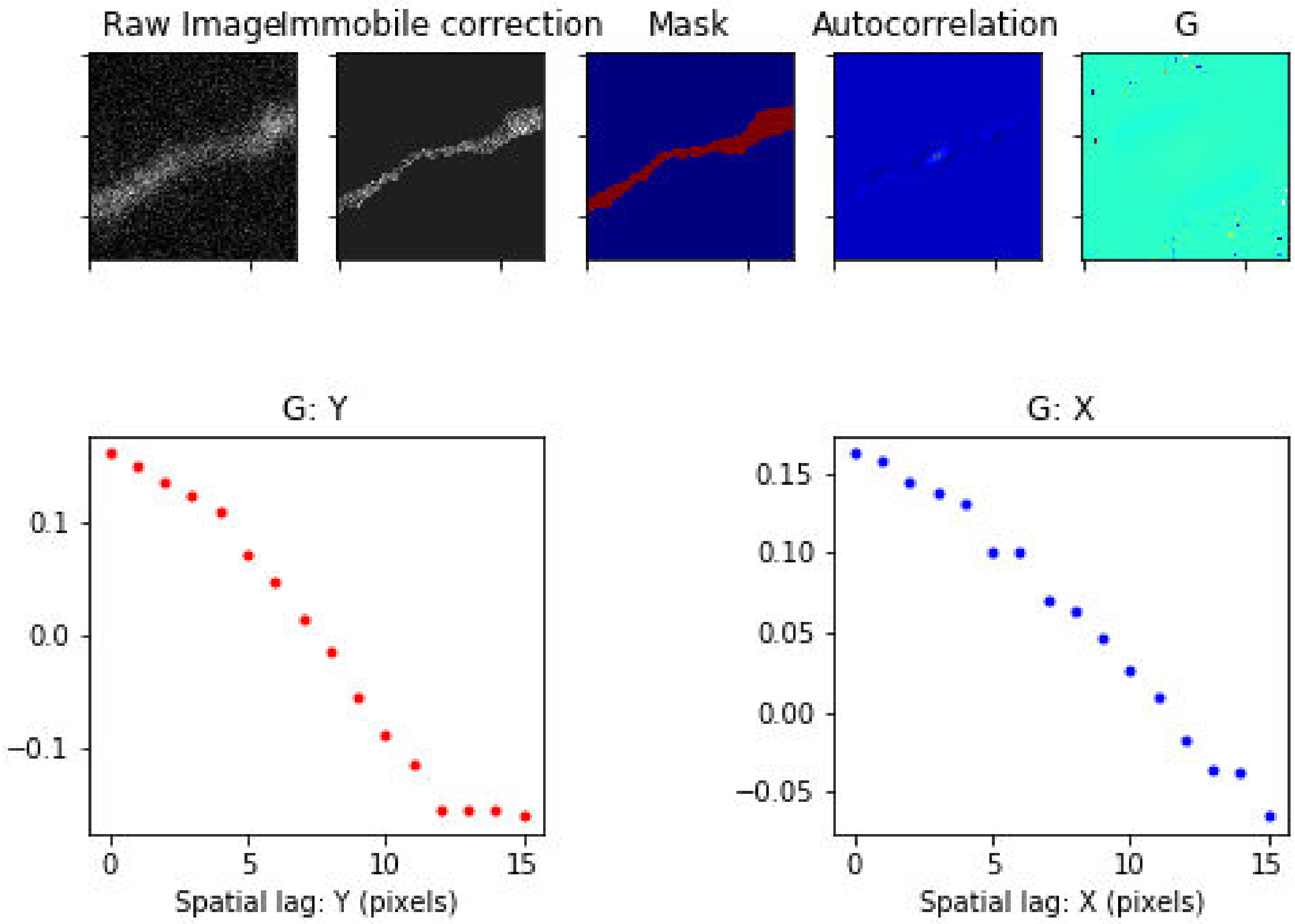

Treatment: Secretin , Domain: IBD , File: Image 19.czi

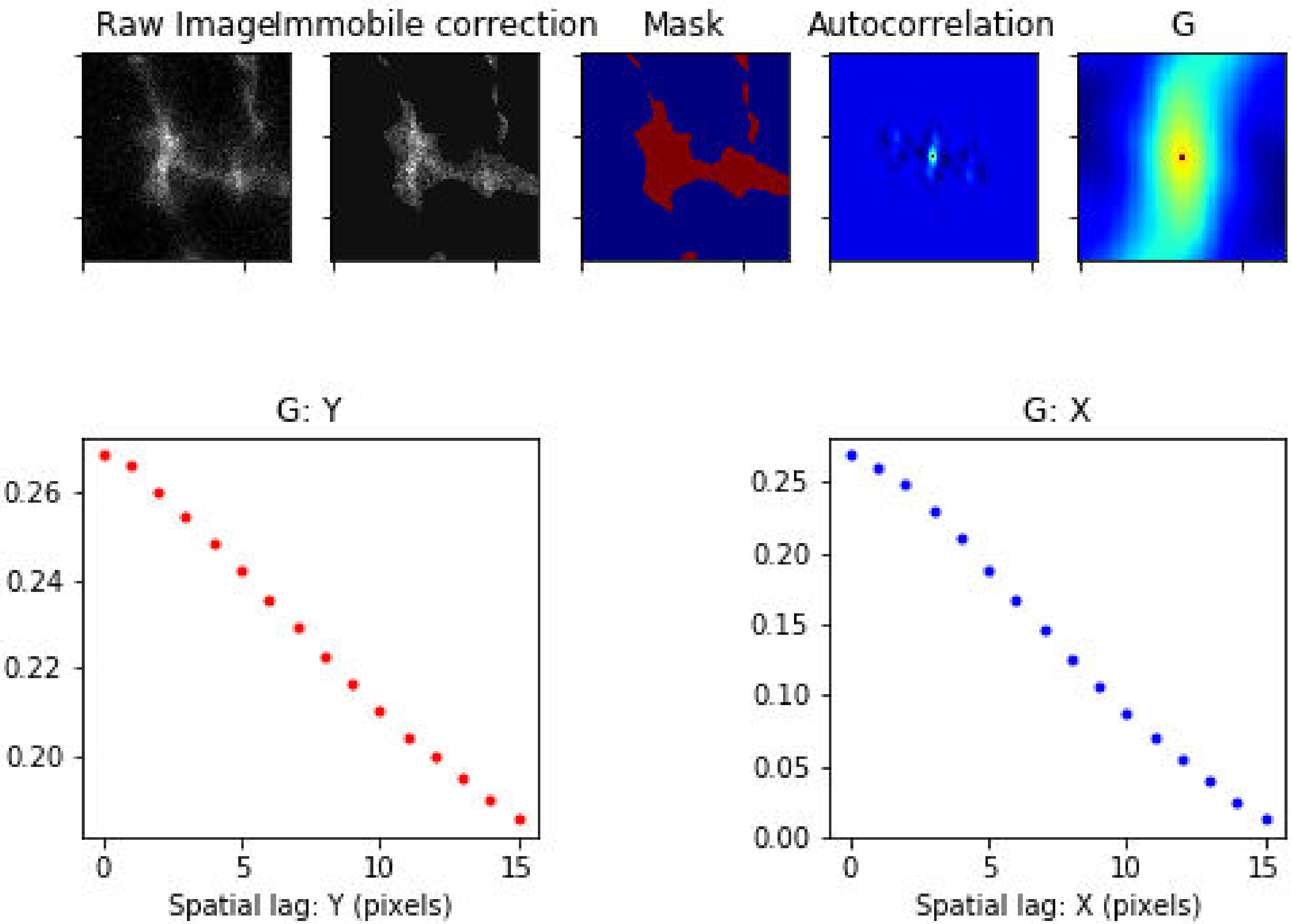

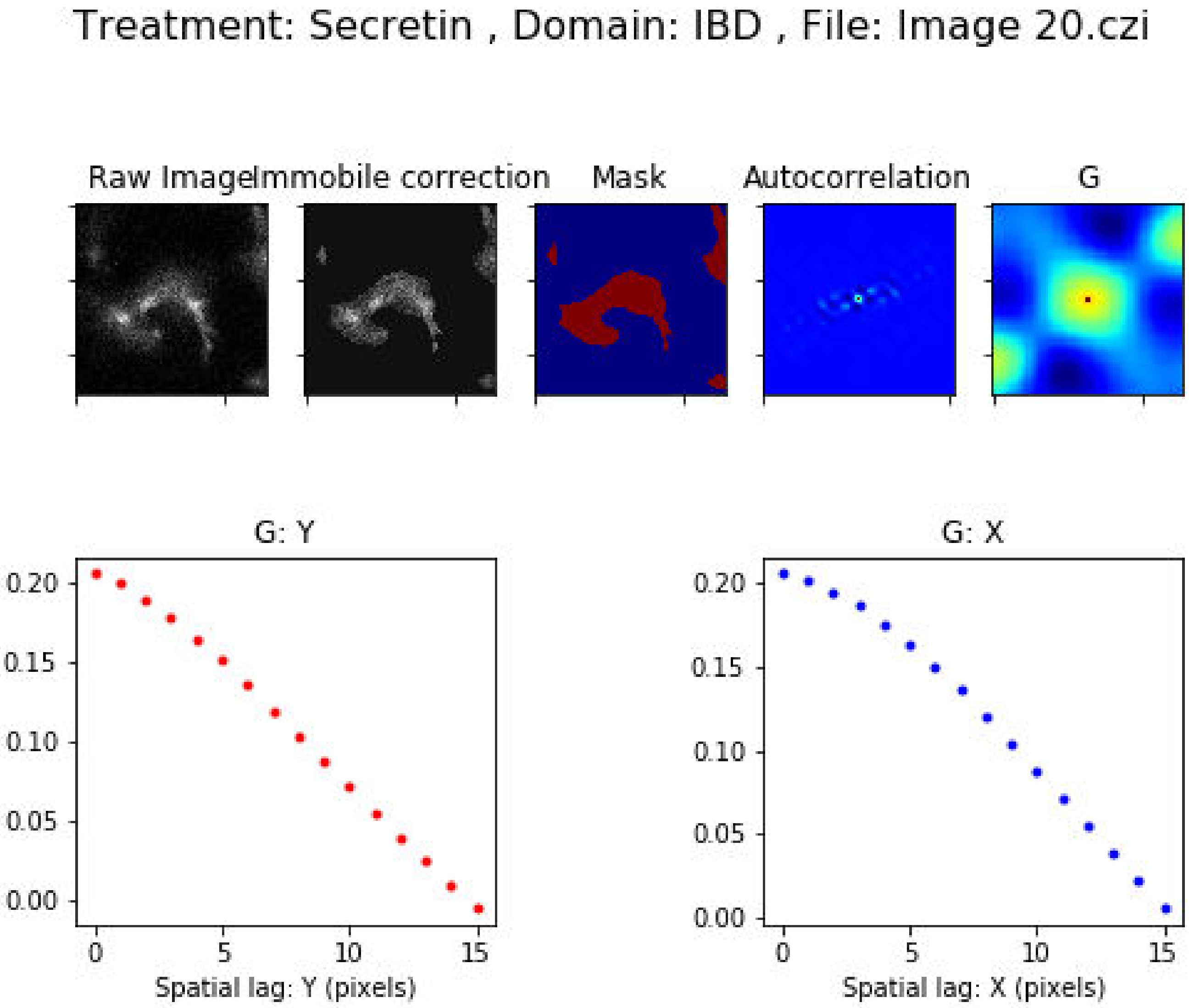

Treatment: Secretin , Domain: IBD , File: Image 20.czi
Raw Image
Immobile correction
Mask
Autocorrelation
G
G: Y
0.20
0.15
0.10
0.05
0.00
0
5
10
15
Spatial lag: Y (pixels)
G: X
0.20
0.15
0.10
0.05
0.00
0
5
10
15
Spatial lag: X (pixels)

Treatment: Secretin , Domain: IBD , File: Image 21.czi

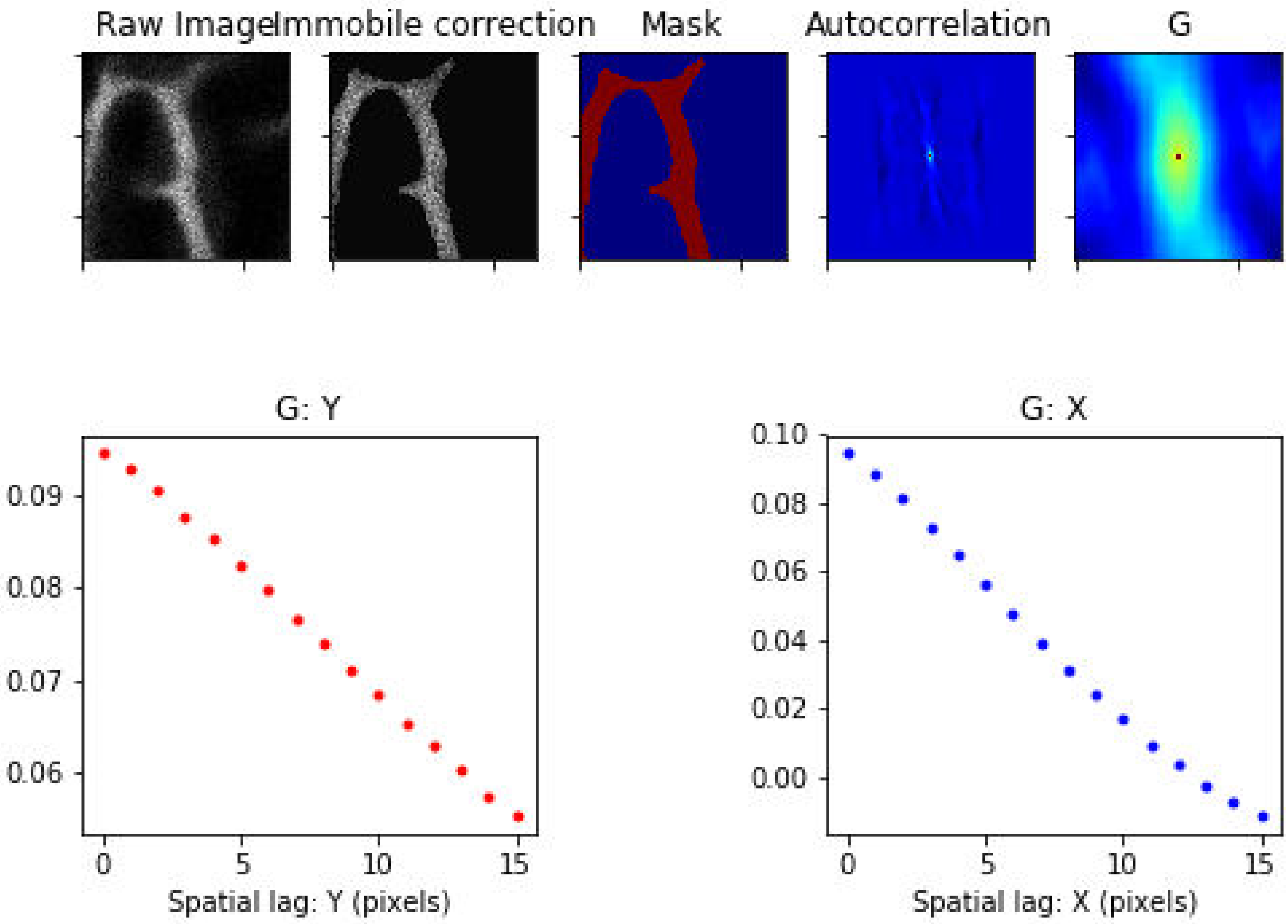

Treatment: Secretin , Domain: IBD , File: Image 22.czi

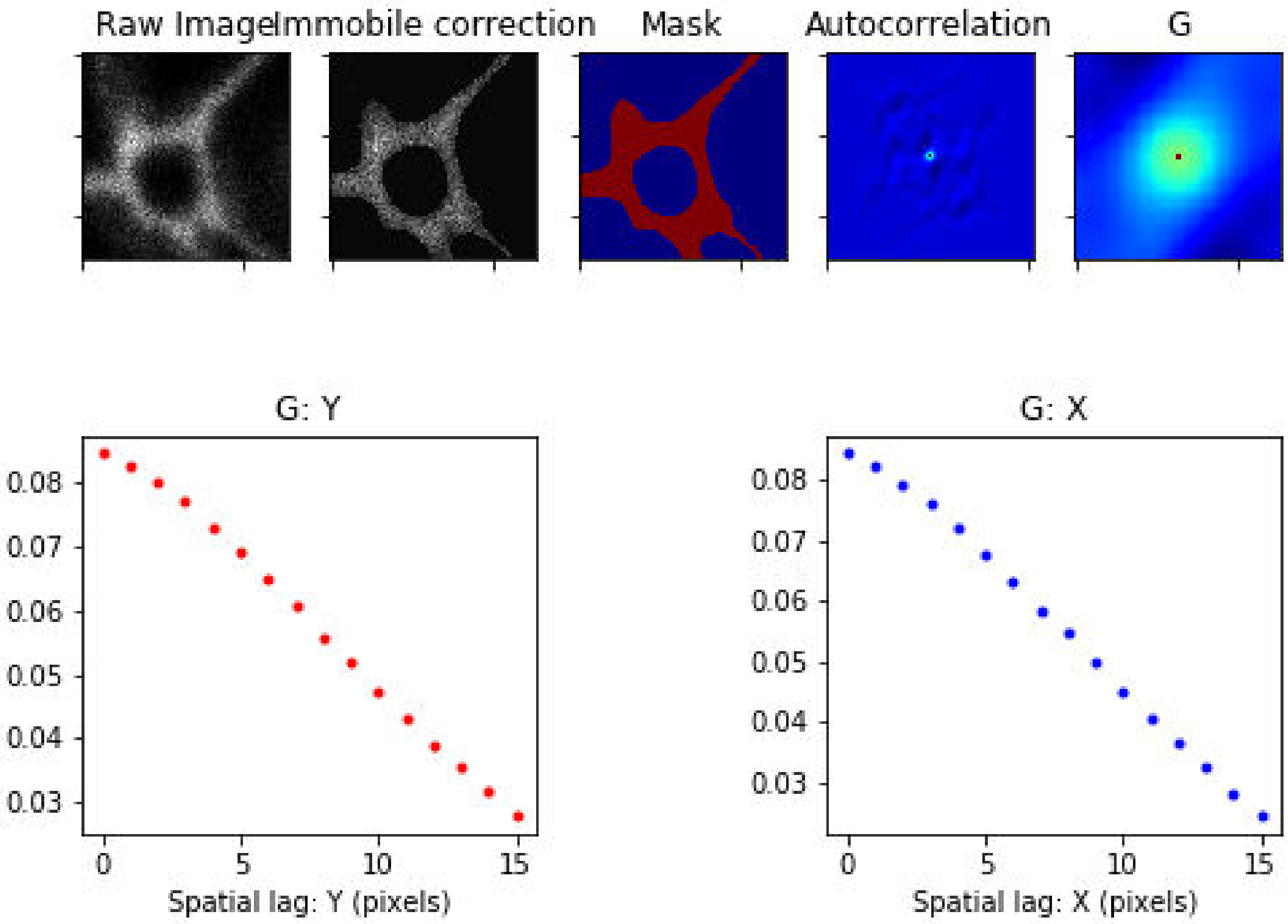

Treatment: Secretin , Domain: IBD , File: Image 23.czi

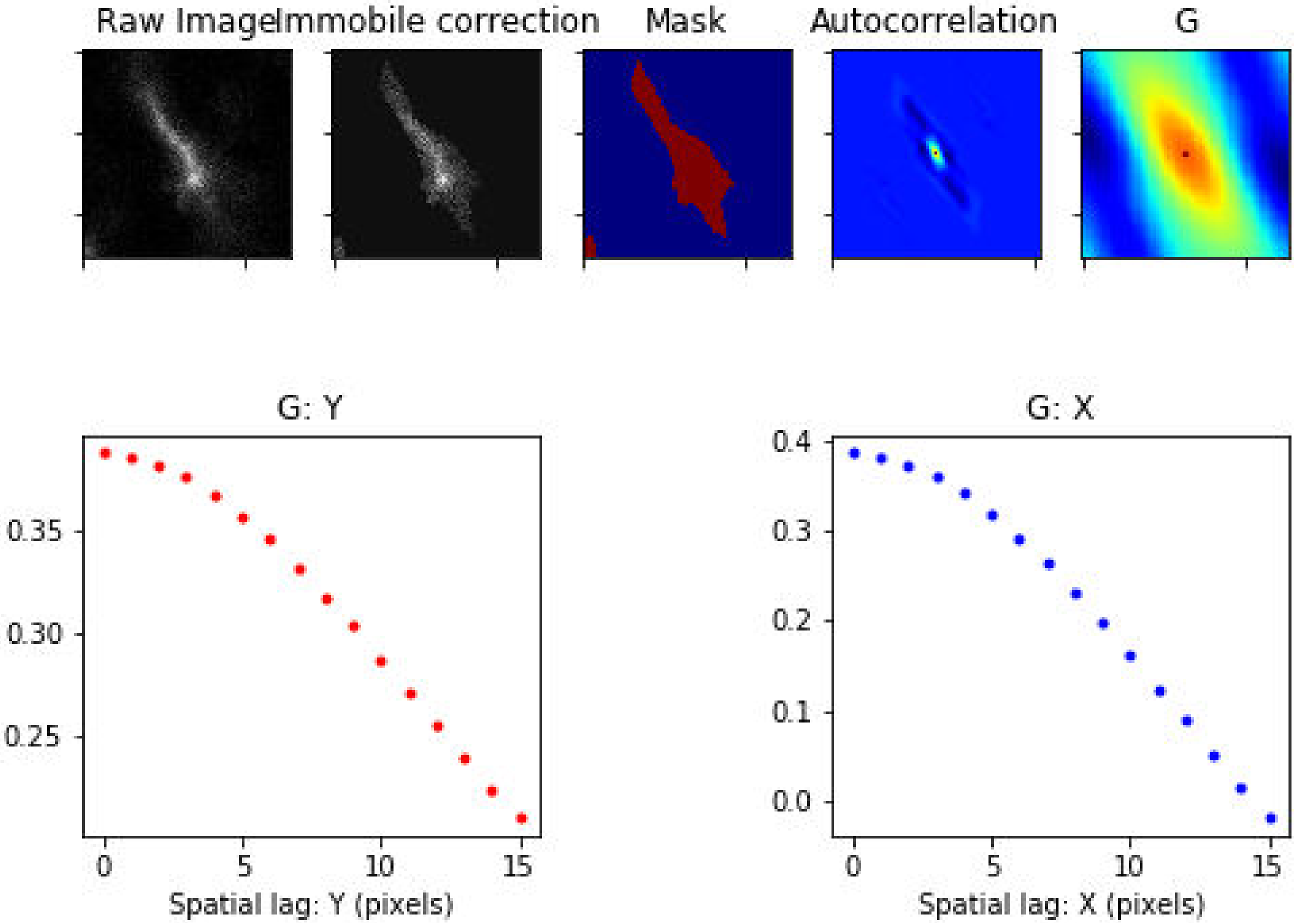

Treatment: Secretin , Domain: IBD , File: Image 48.czi

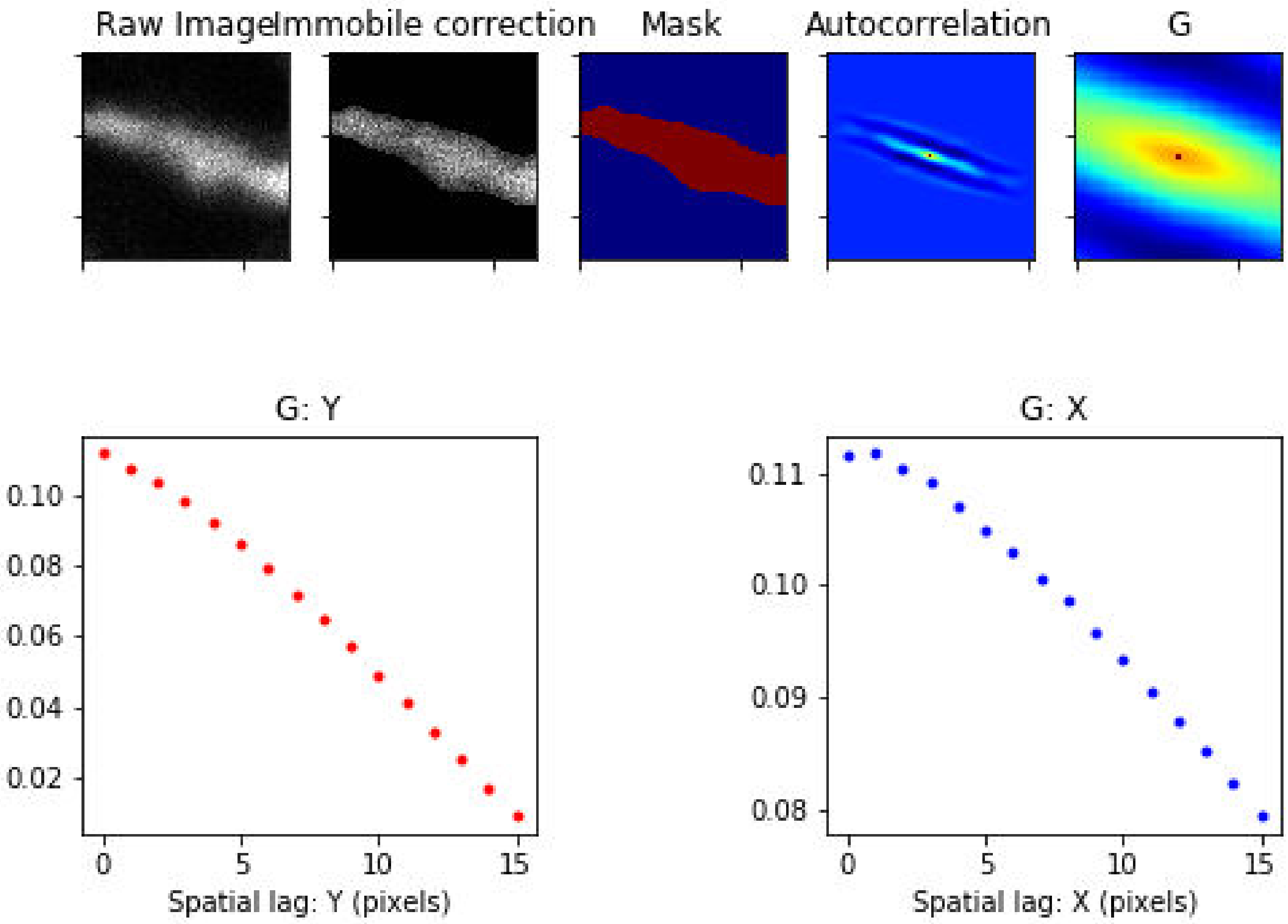

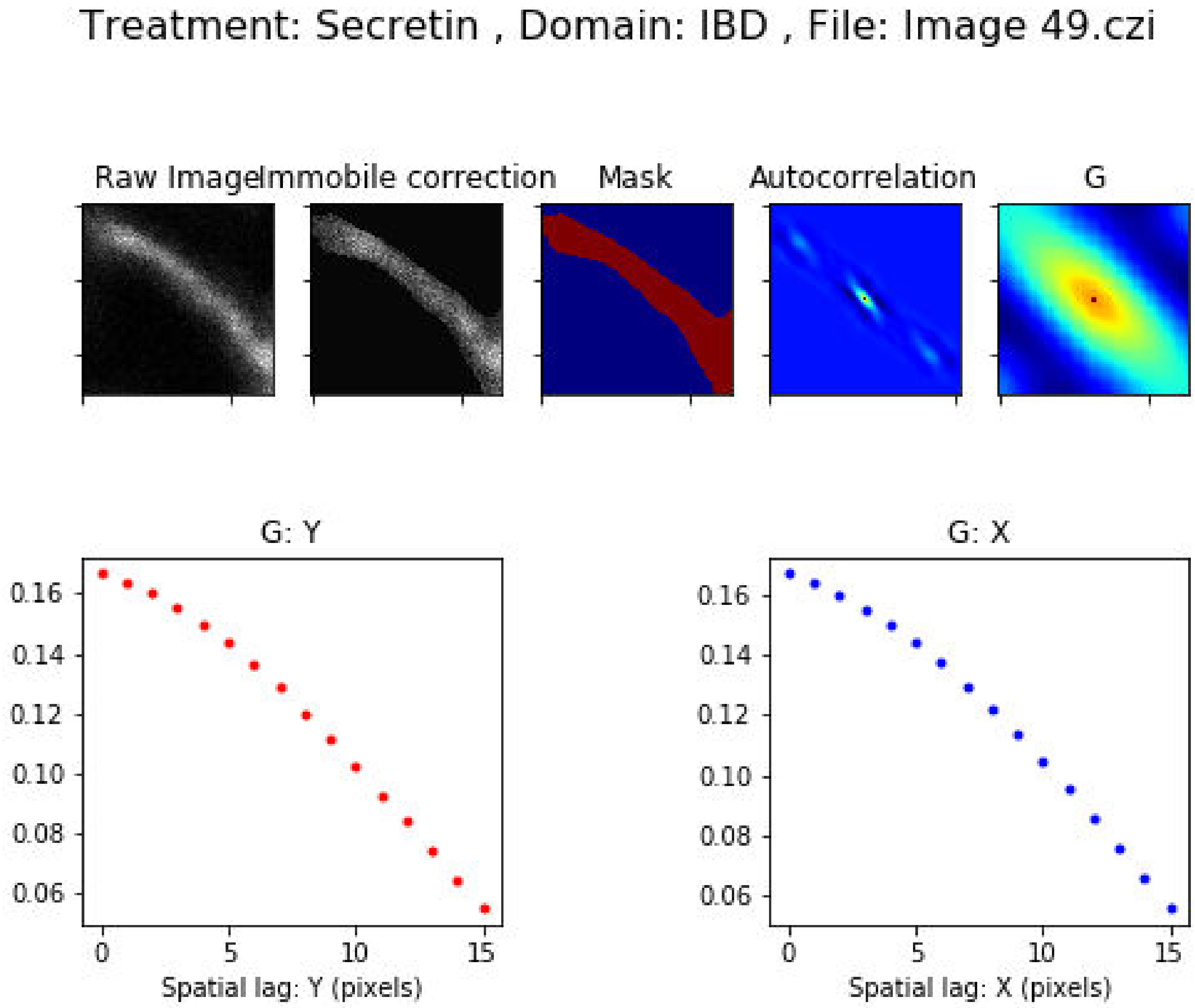
Treatment: Secretin , Domain: IBD , File: Image 49.czi
Raw Image
Immobile correction
Mask
Autocorrelation
G
G: Y
0.16
0.14
0.12
0.10
0.08
0.06
0
5
10
15
Spatial lag: Y (pixels)
G: X
0.16
0.14
0.12
0.10
0.08
0.06
0
5
10
15
Spatial lag: X (pixels)

Treatment: Secretin , Domain: IBD , File: Image 50.czi

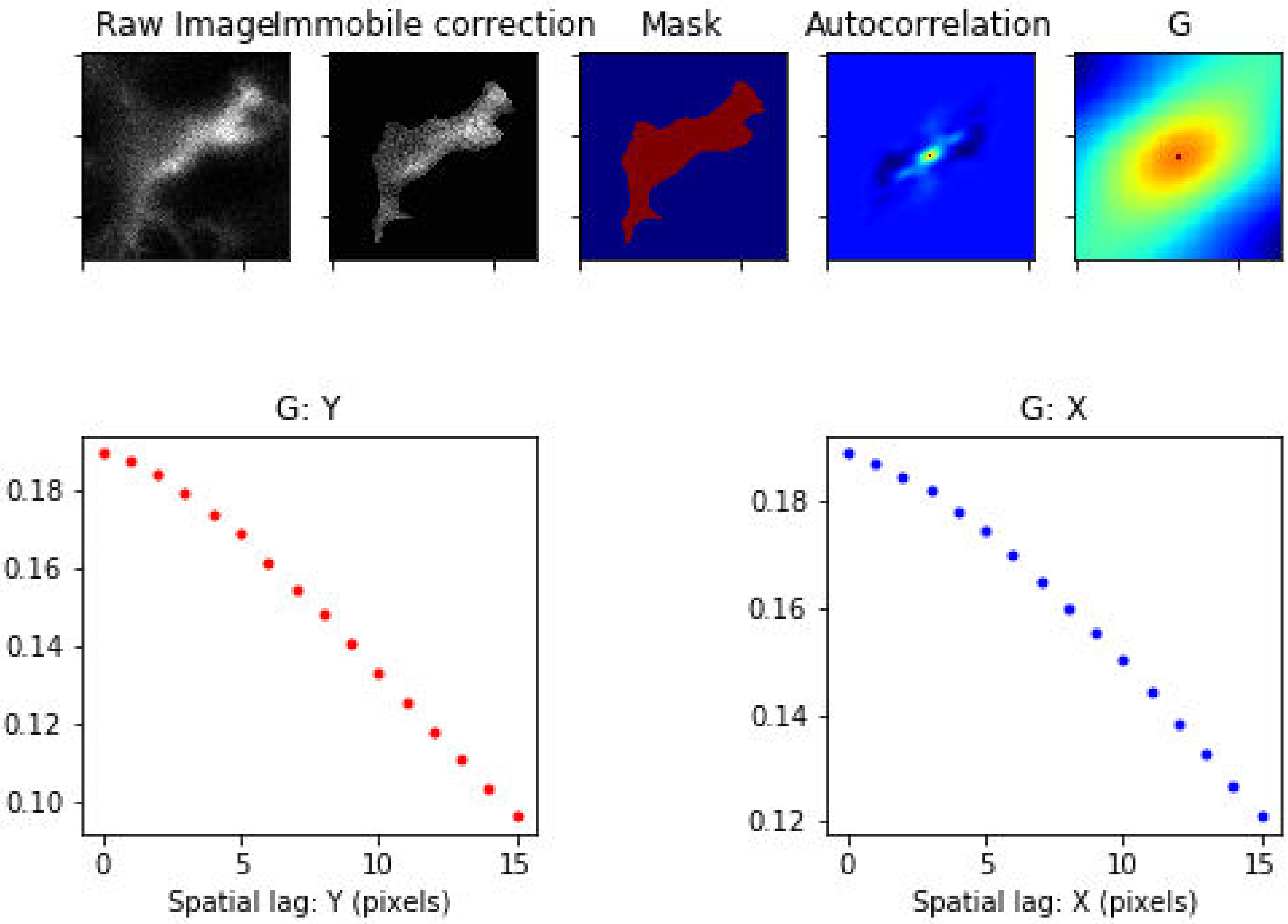

### Treatment: Secretin , Domain: IBD , File: Image 51.czi

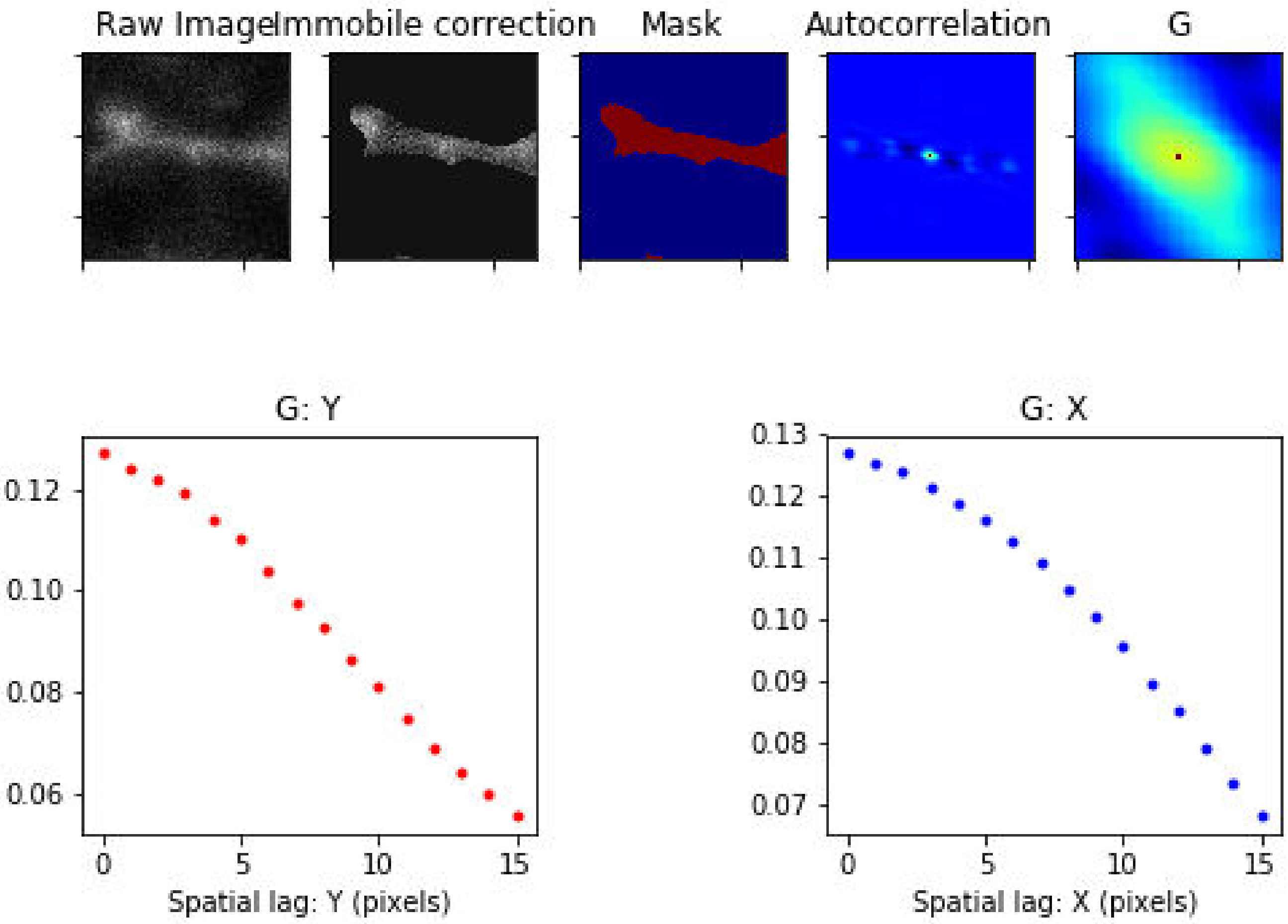

Treatment: Secretin , Domain: IBD , File: Image 73.czi

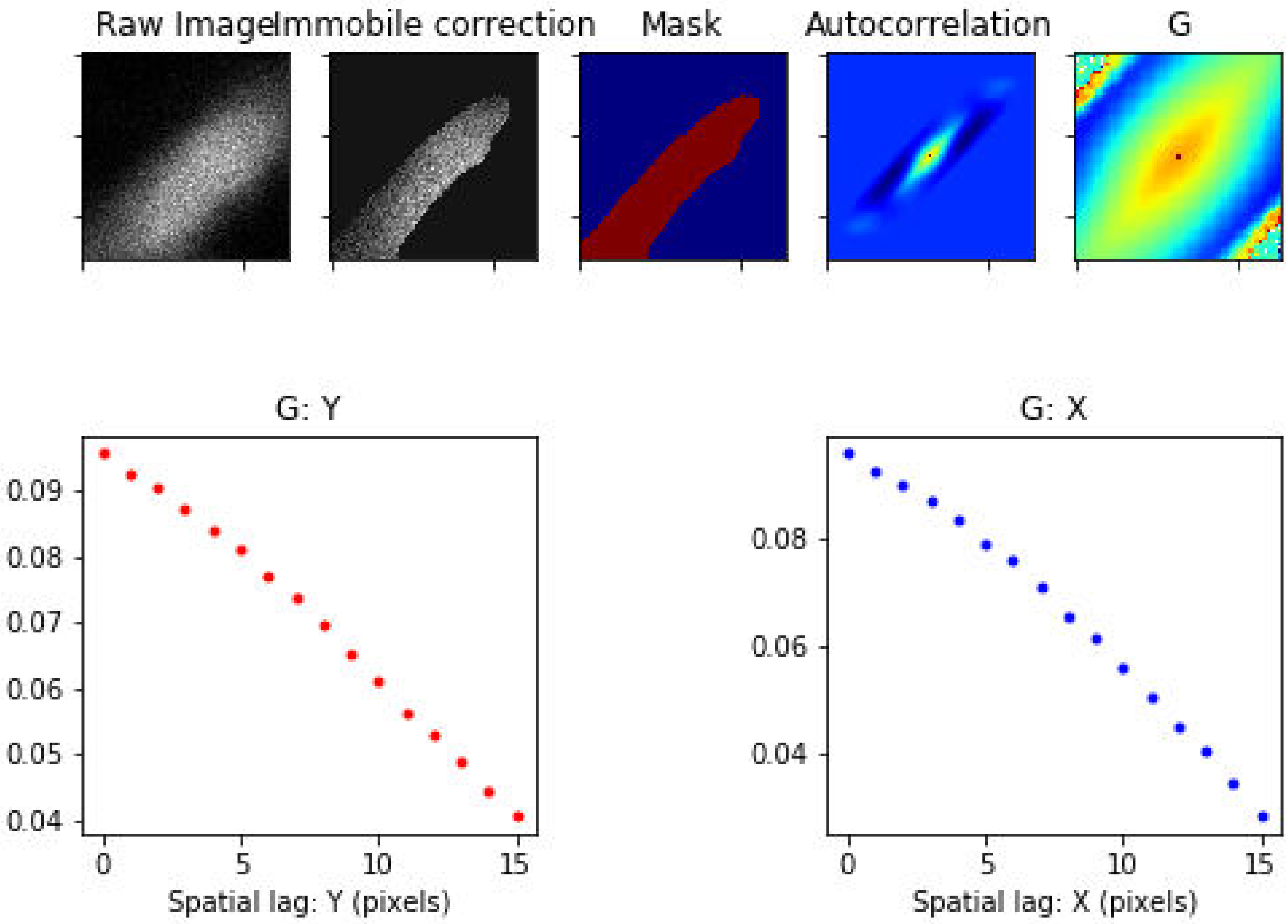

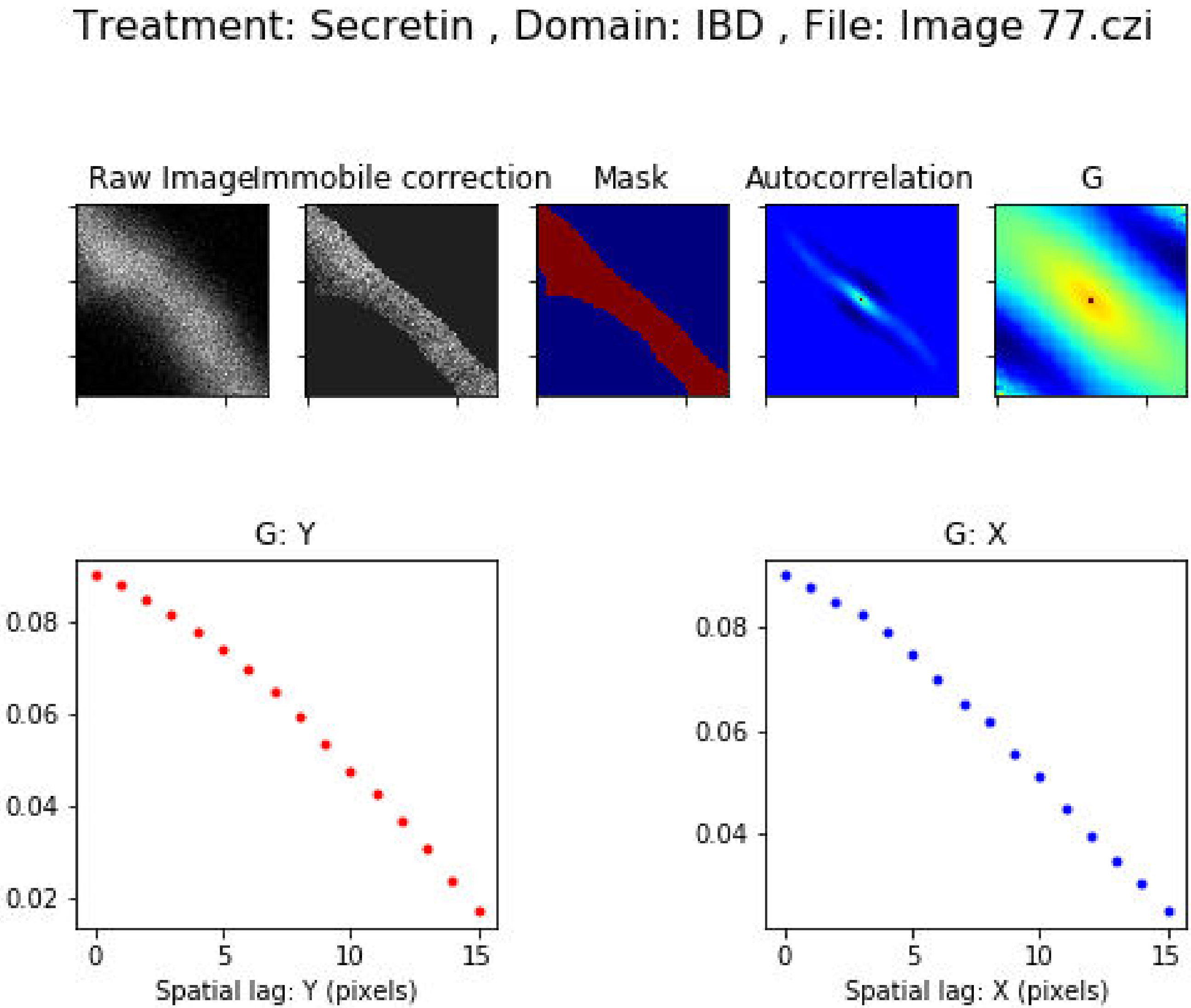
Treatment: Secretin , Domain: IBD , File: Image 77.czi
Raw Image
Immobile correction
Mask
Autocorrelation
G
G: Y
0.08
0.06
0.04
0.02
0
5
10
15
Spatial lag: Y (pixels)
G: X
0.08
0.06
0.04
0
5
10
15
Spatial lag: X (pixels)

### Treatment: Secretin , Domain: IBD , File: Image 79.czi

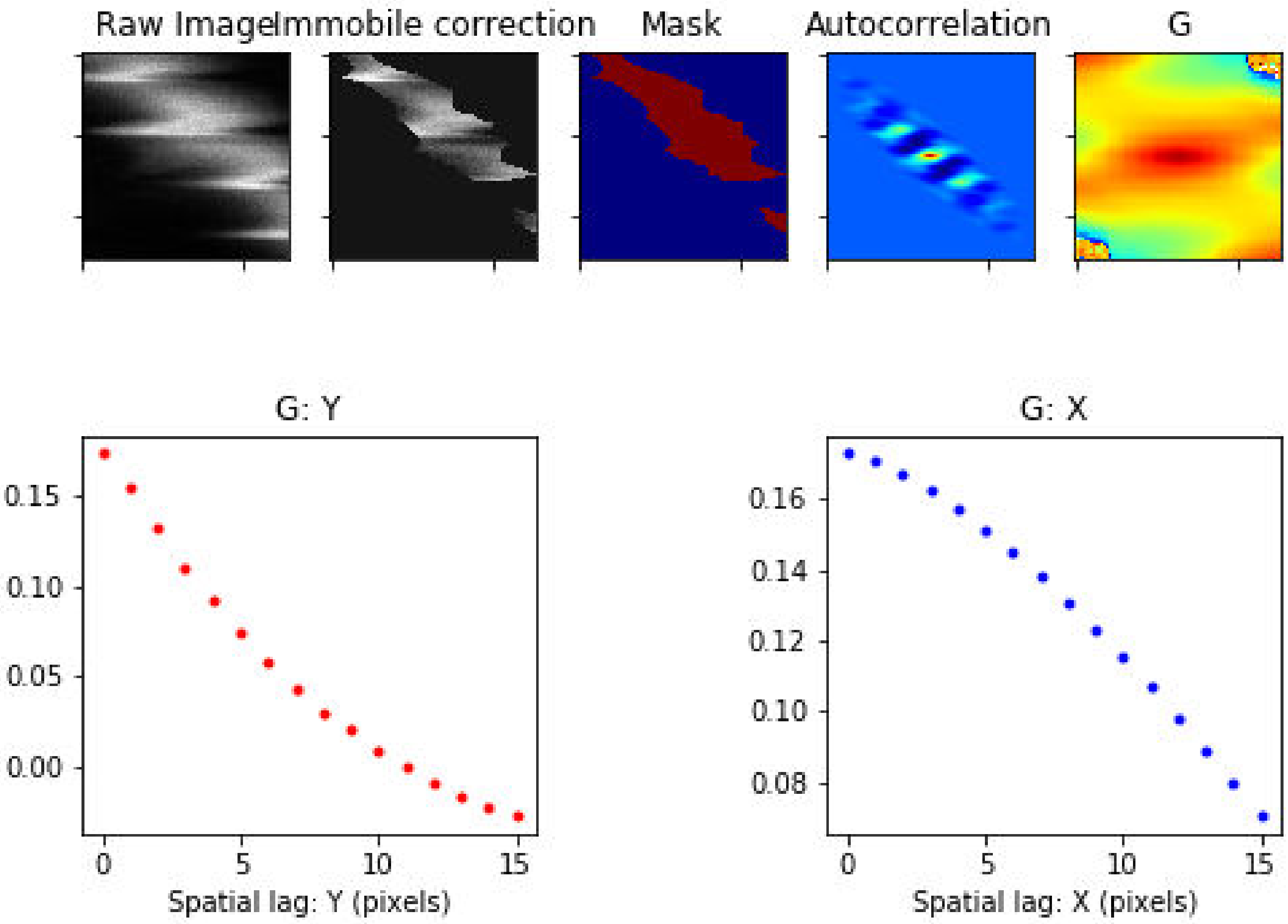

Treatment: Secretin , Domain: IBD , File: Image 83.czi

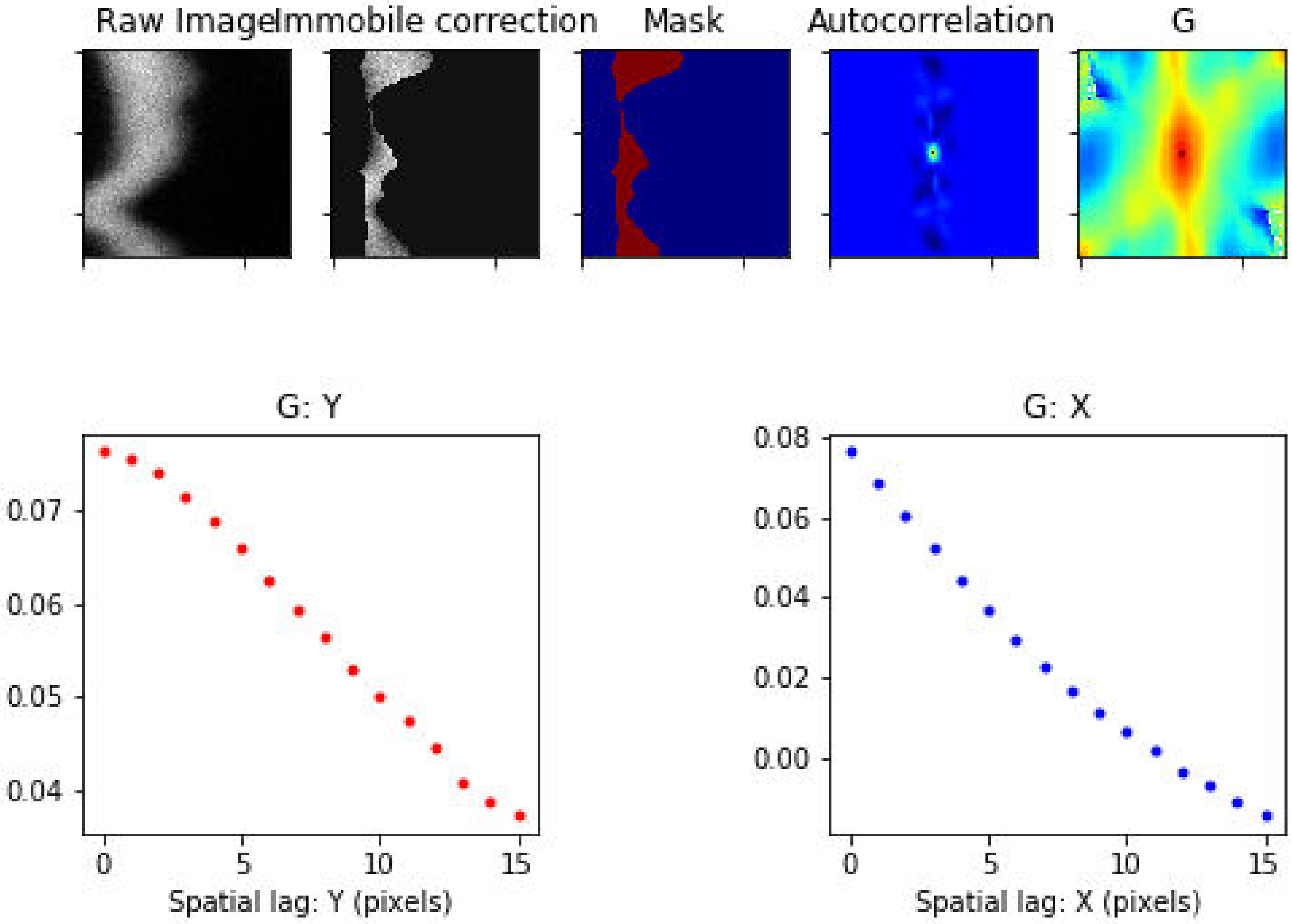

### Treatment: Secretin , Domain: IBD , File: Image 88.czi

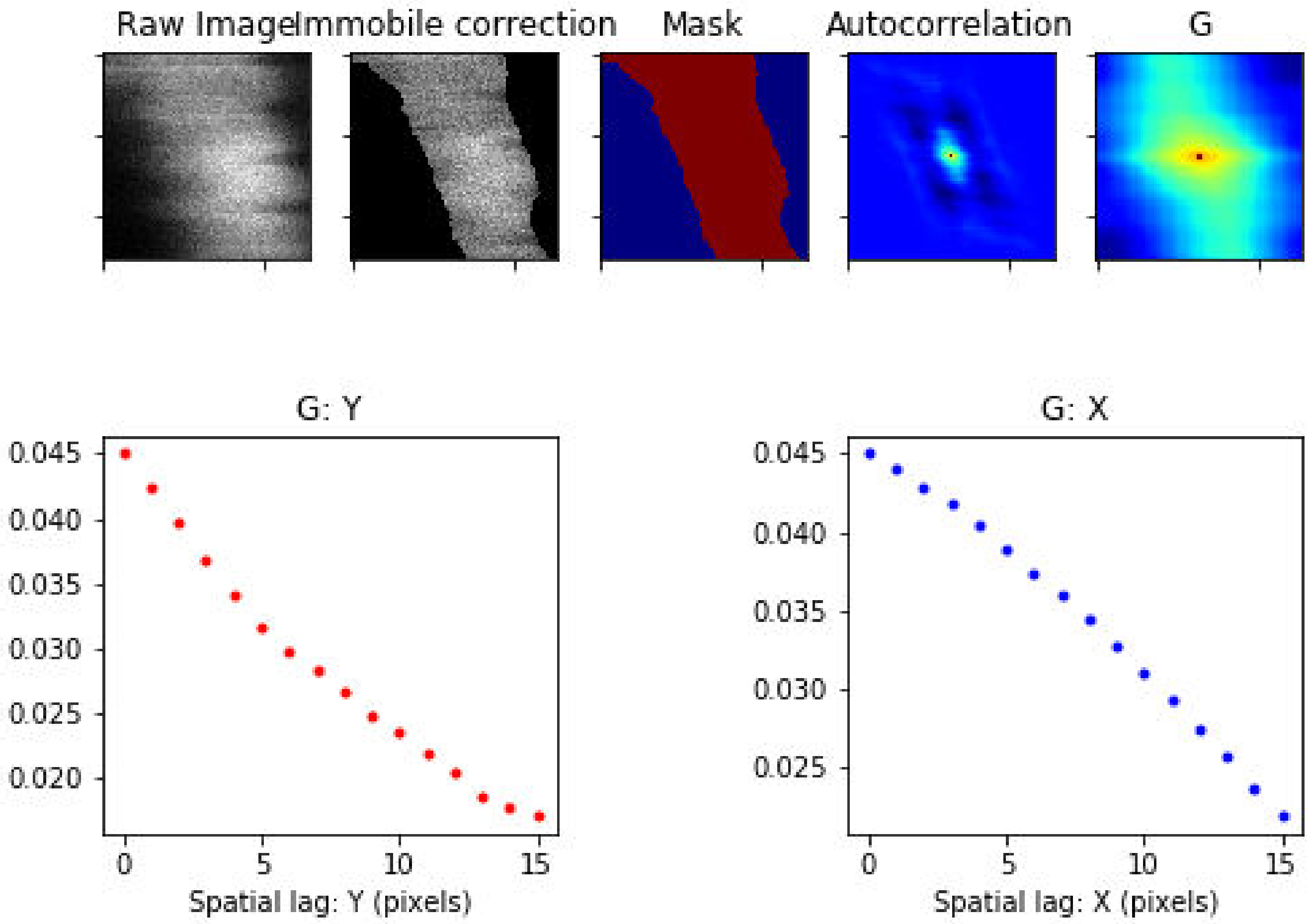

Treatment: Secretin , Domain: MZ , File: Image 26.czi

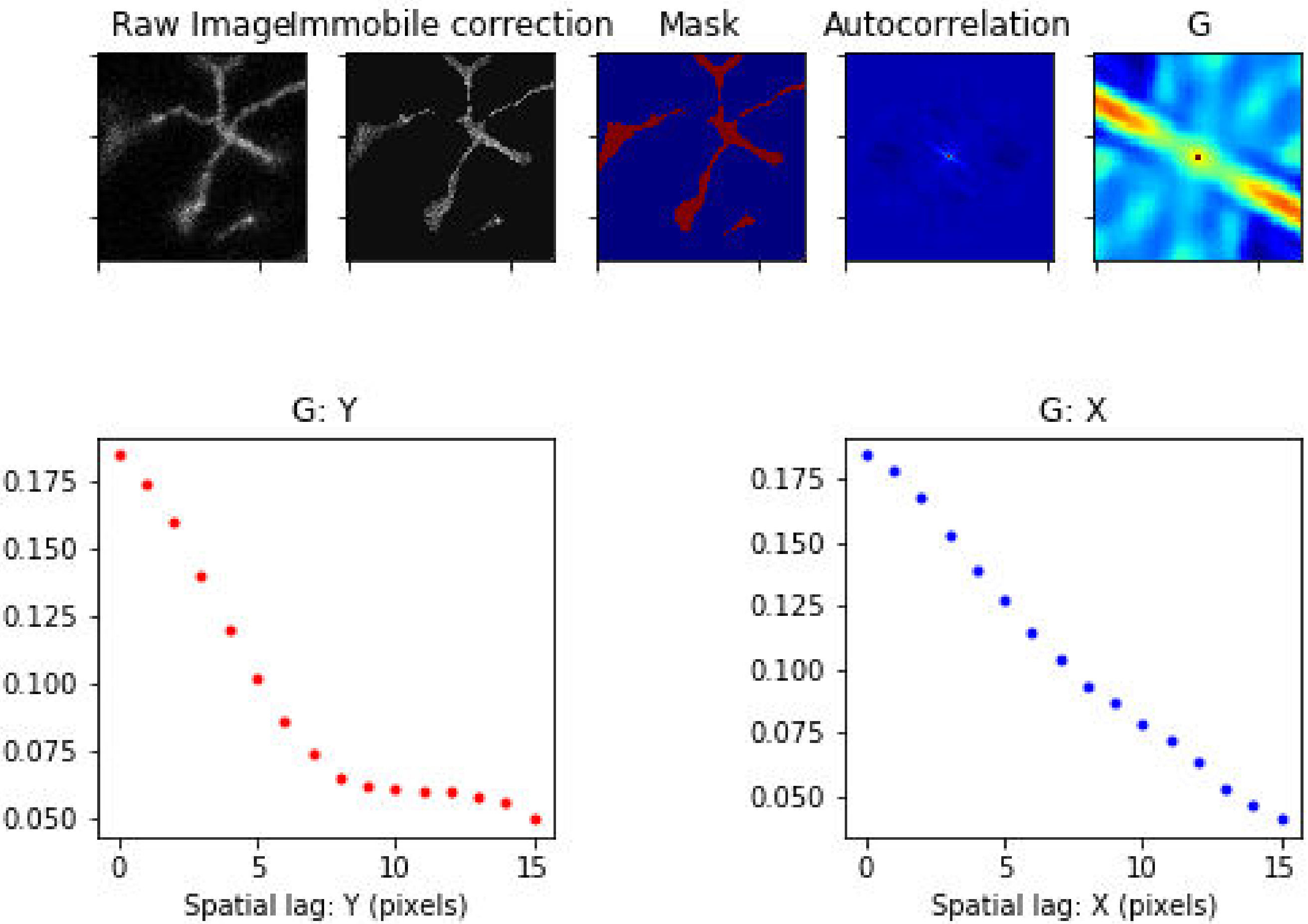

Treatment: Secretin , Domain: MZ , File: Image 29.czi

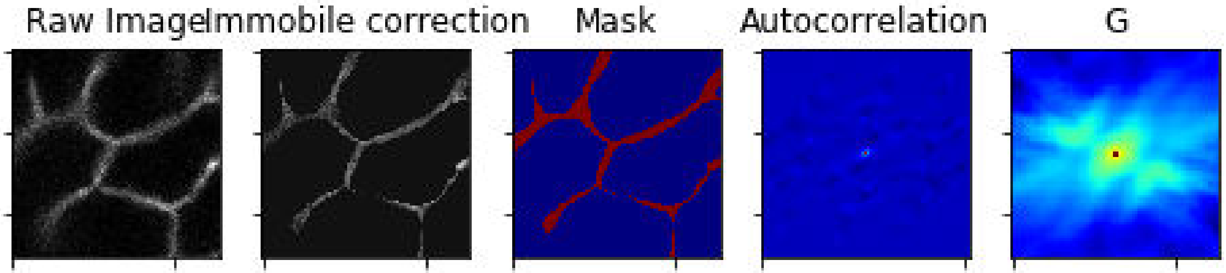


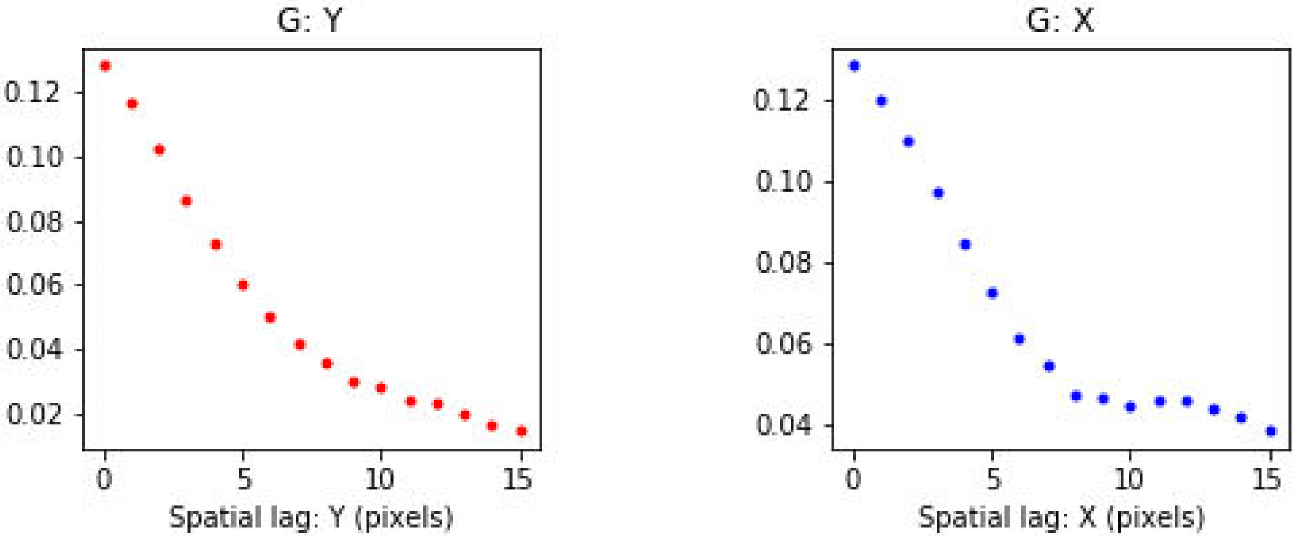

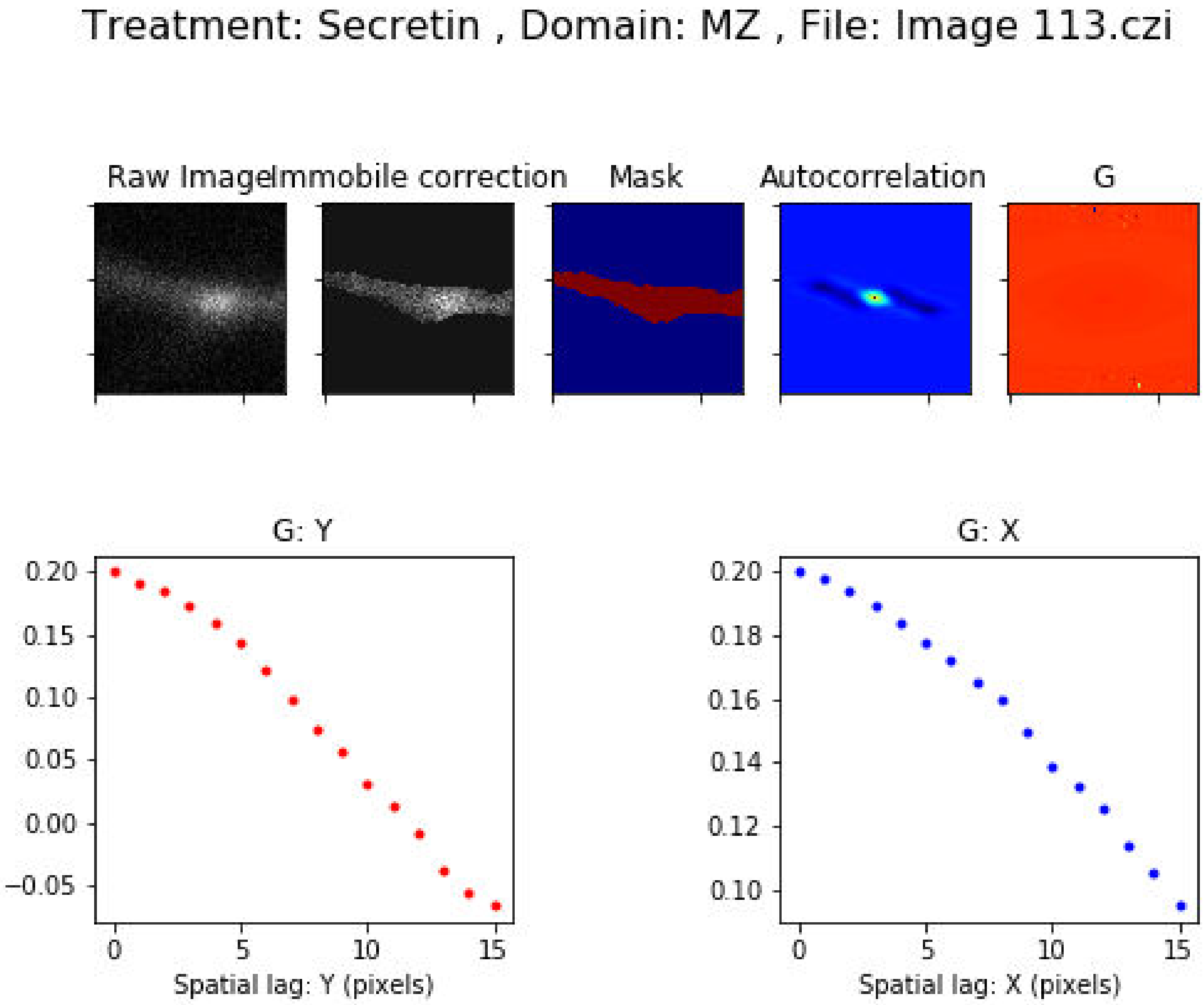
Treatment: Secretin , Domain: MZ , File: Image 113.czi
Raw Image
Immobile correction
Mask
Autocorrelation
G
G: Y
0.20
0.15
0.10
0.05
0.00
−0.05
0
5
10
15
Spatial lag: Y (pixels)
G: X
0.20
0.18
0.16
0.14
0.12
0.10
0
5
10
15
Spatial lag: X (pixels)

Treatment: Secretin , Domain: MZ , File: Image 114.czi

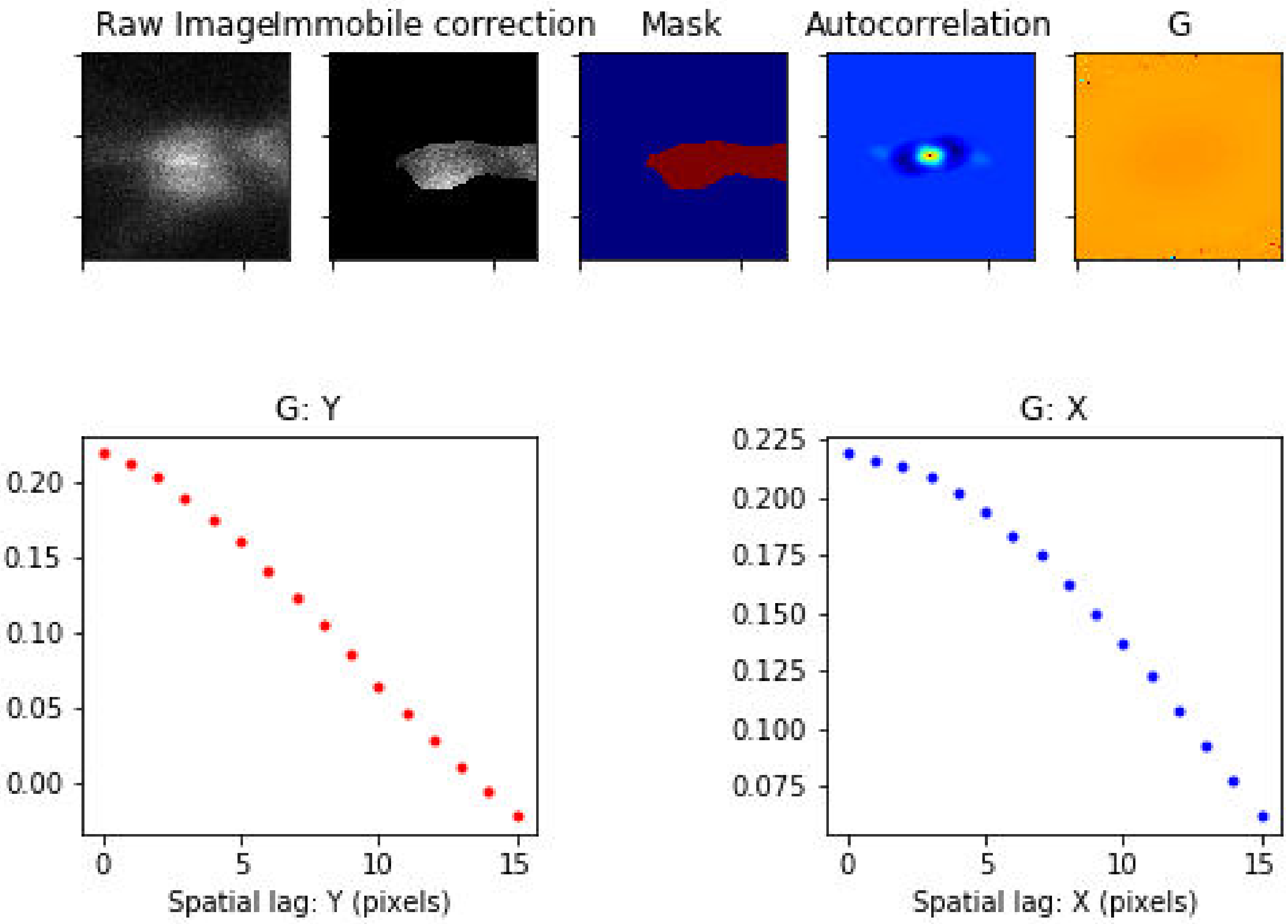

Treatment: Secretin , Domain: MZ , File: Image 115.czi

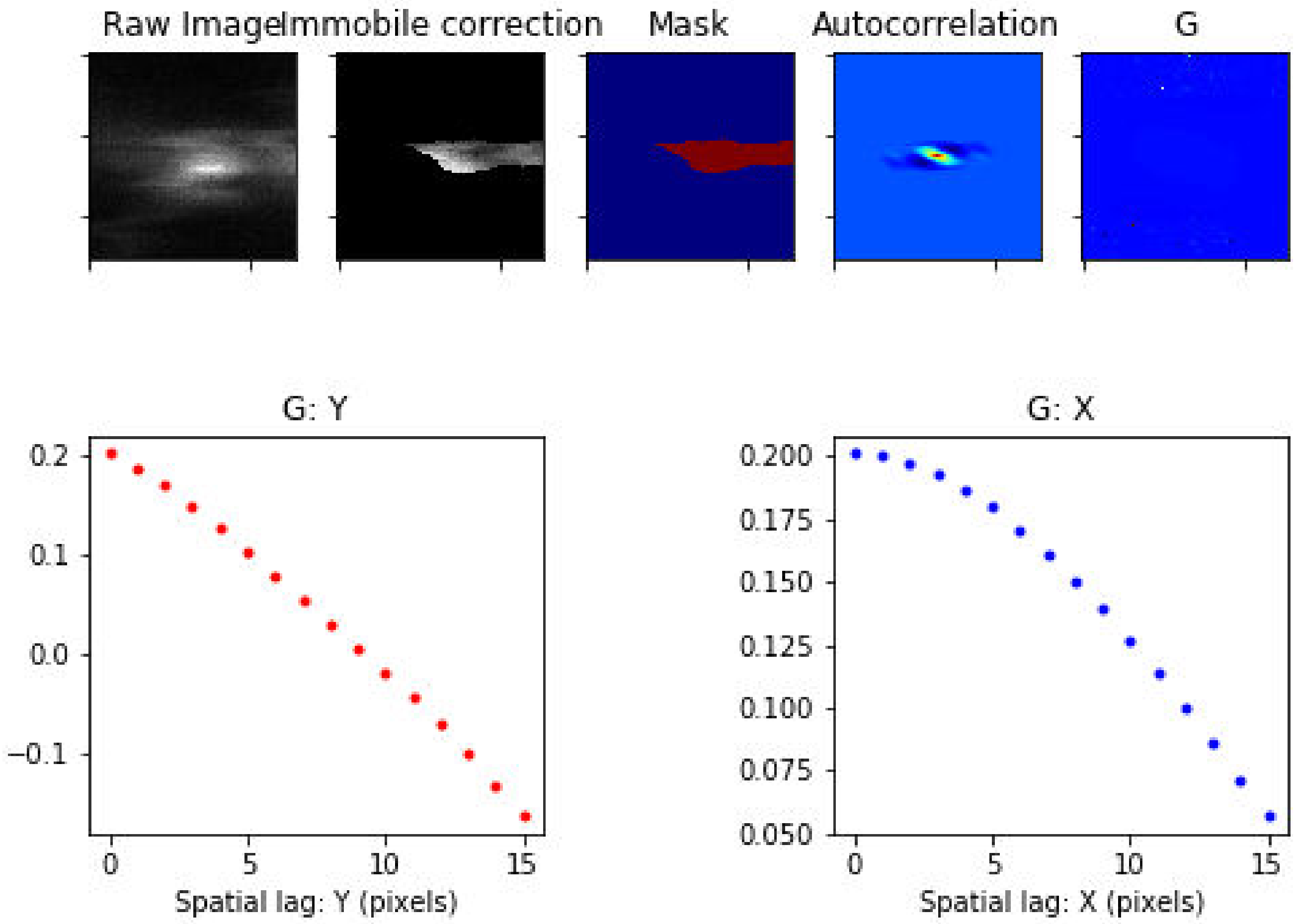

### Treatment: Secretin , Domain: MZ , File: Image 118.czi

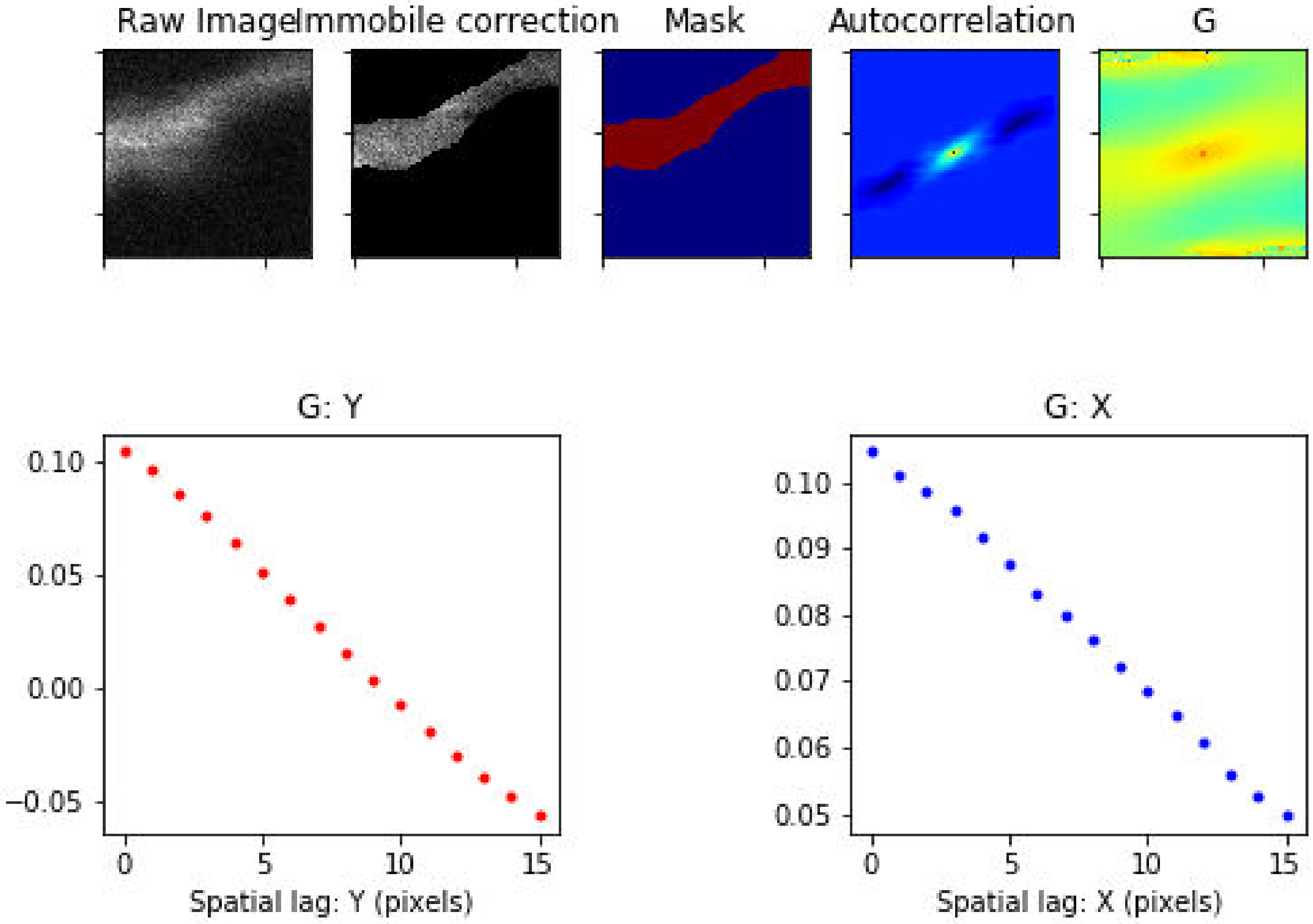

Treatment: Secretin , Domain: MZ , File: Image 119.czi

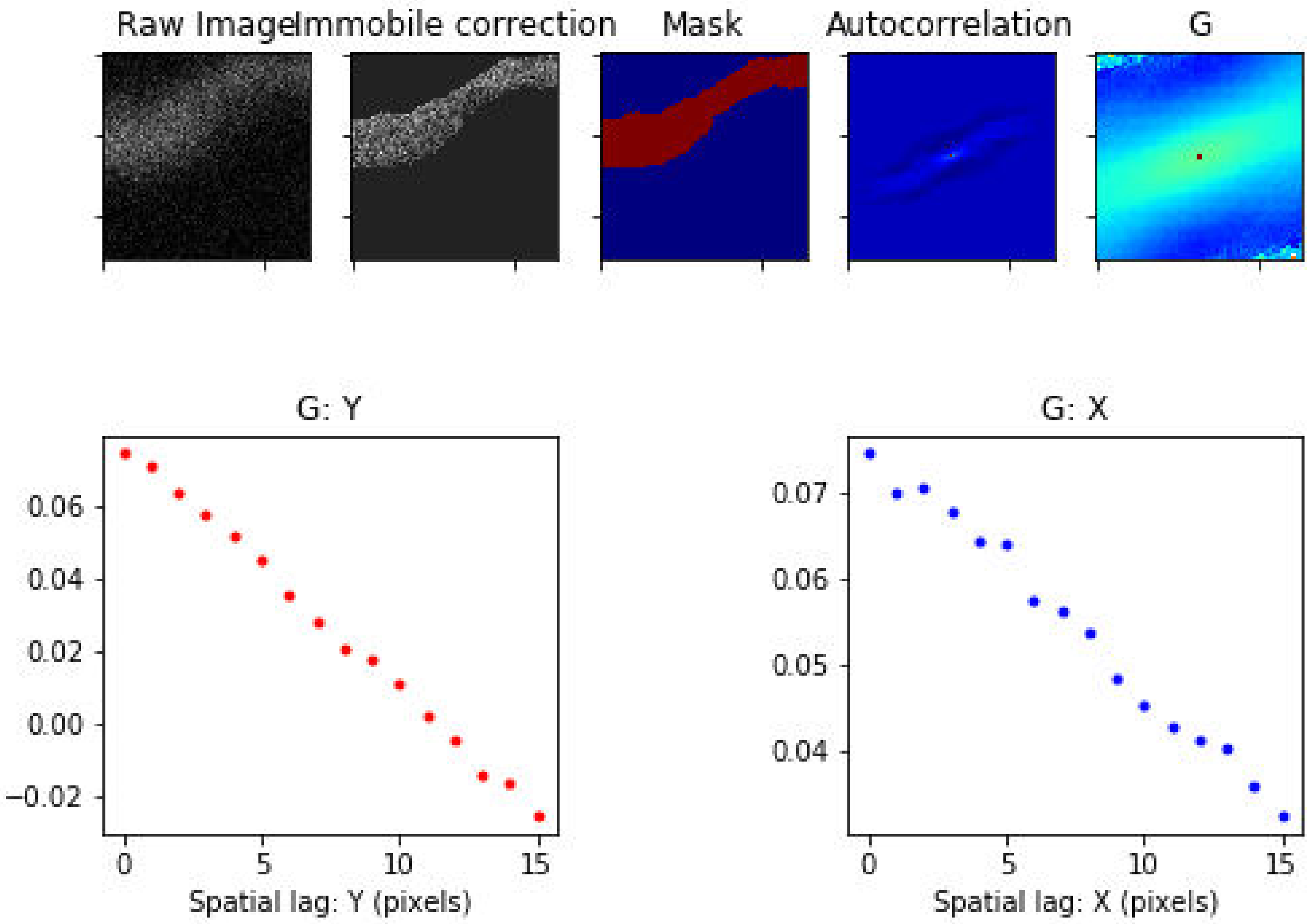

### Treatment: Secretin , Domain: MZ , File: Image 120.czi

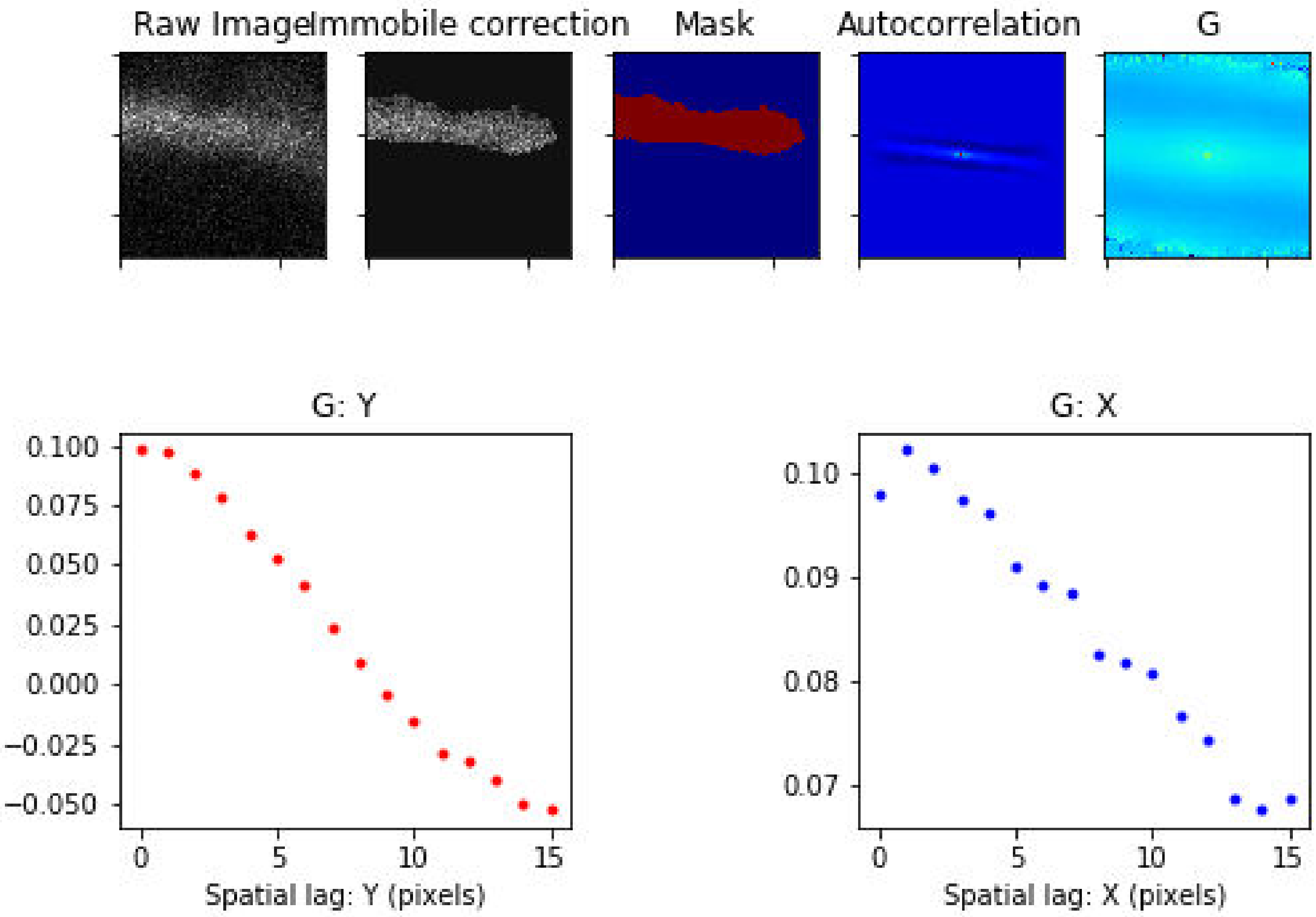

### Treatment: Secretin , Domain: MZ , File: Image 121.czi

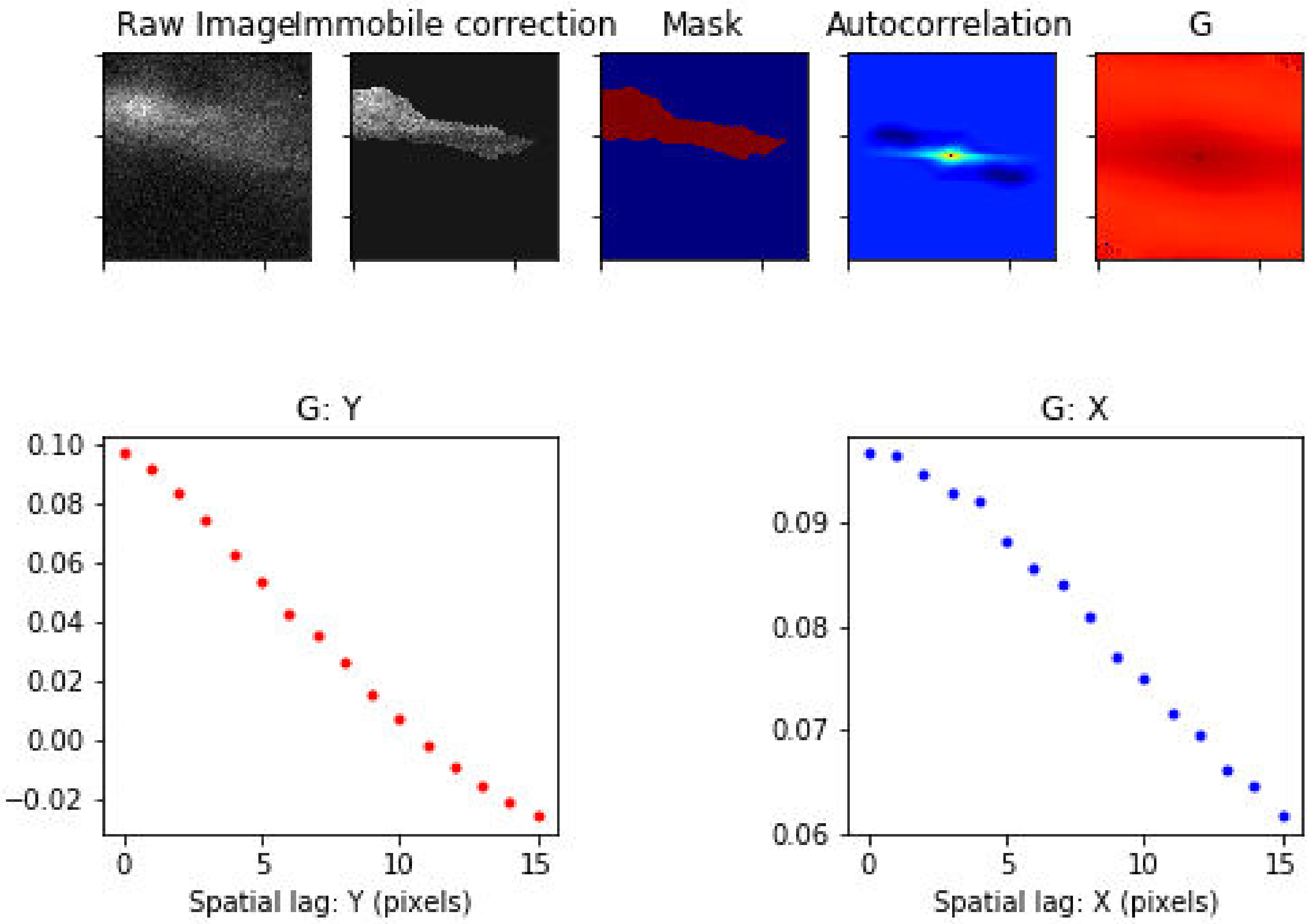

Treatment: Secretin , Domain: MZ , File: Image 122.czi

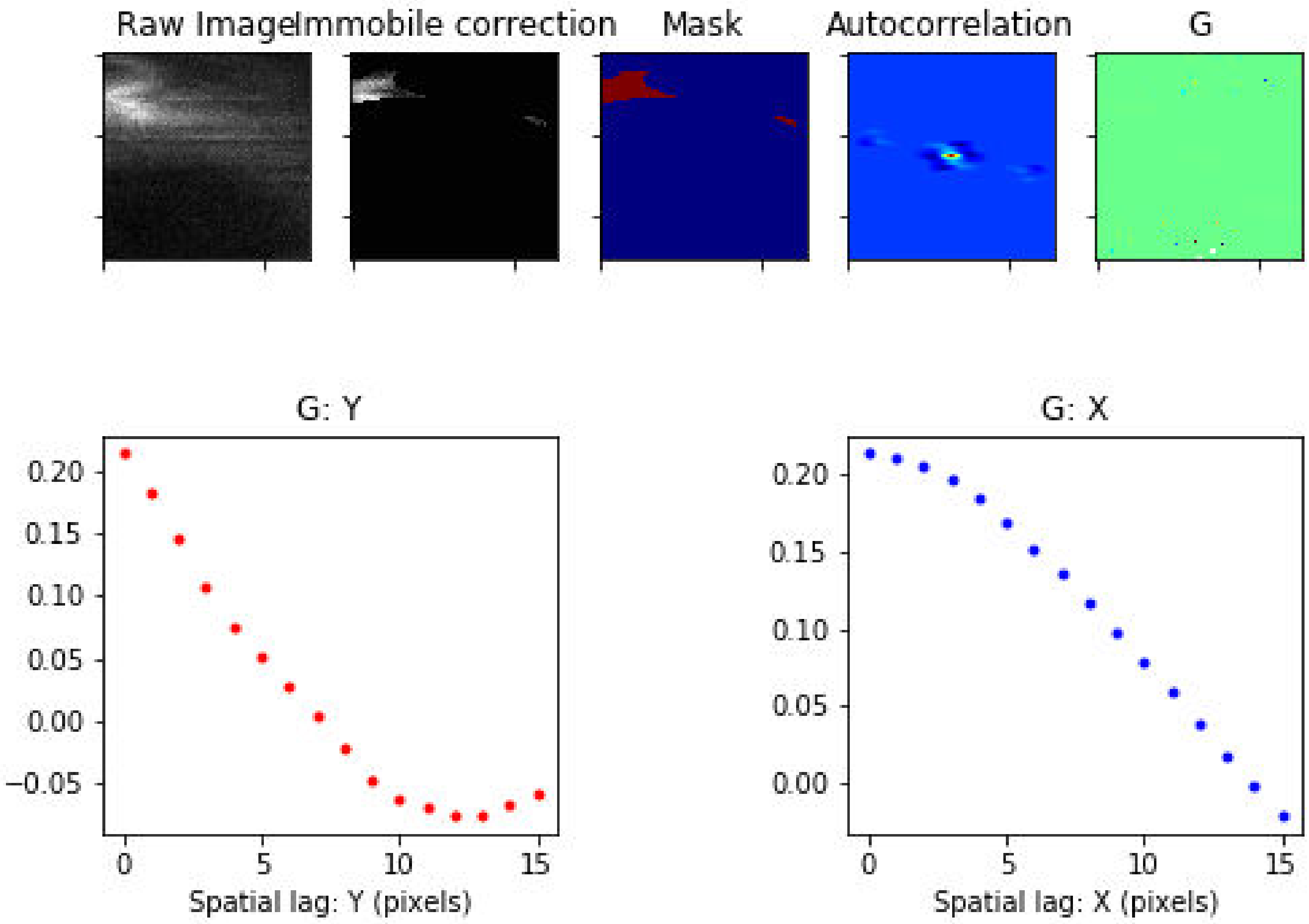

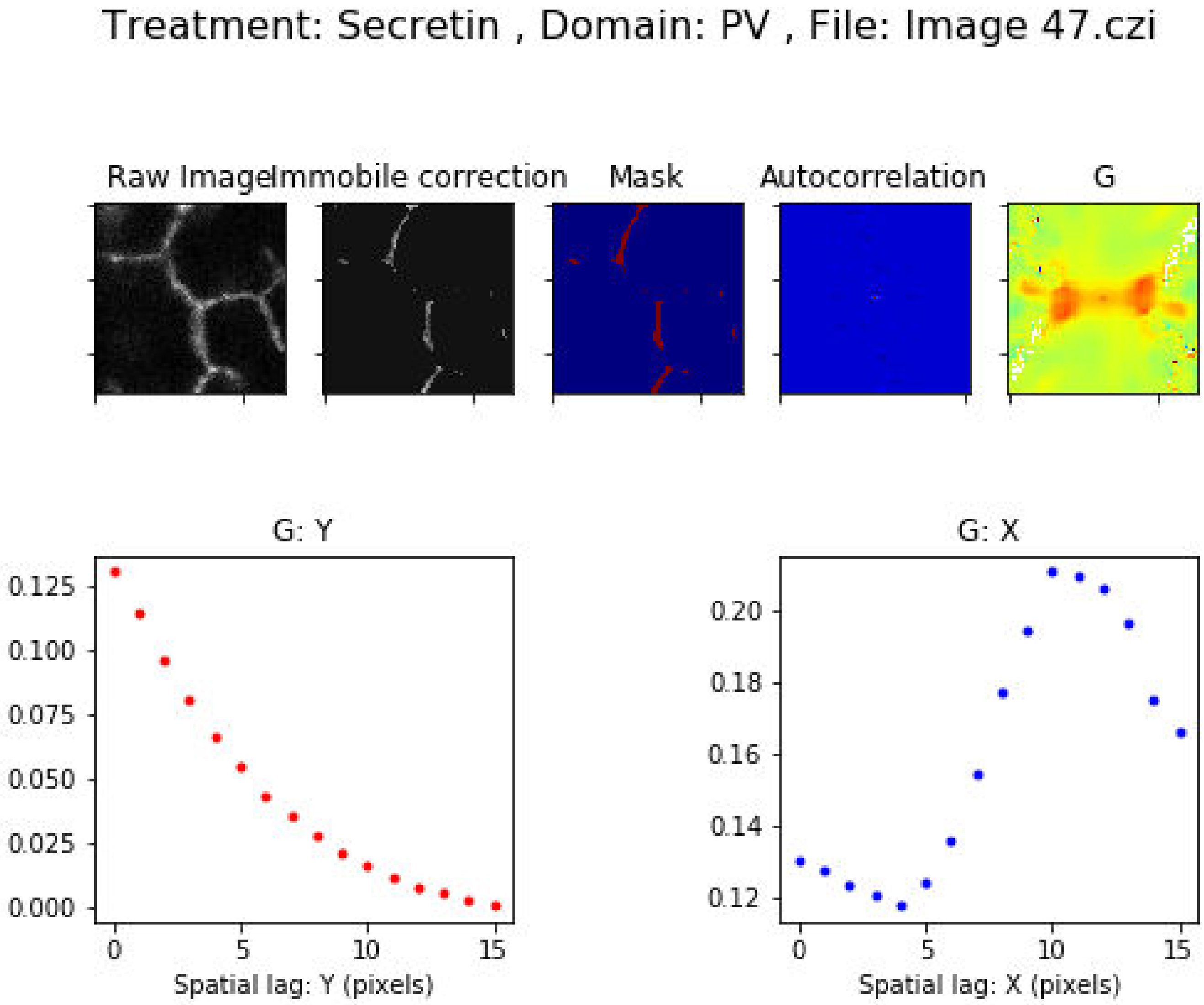
Treatment: Secretin , Domain: PV , File: Image 47.czi
Raw Image
Immobile correction
Mask
Autocorrelation
G
G: Y
0.125
0.100
0.075
0.050
0.025
0.000
0
5
10
15
Spatial lag: Y (pixels)
G: X
0.20
0.18
0.16
0.14
0.12
0
5
10
15
Spatial lag: X (pixels)

Treatment: Secretin , Domain: PV , File: Image 100.czi

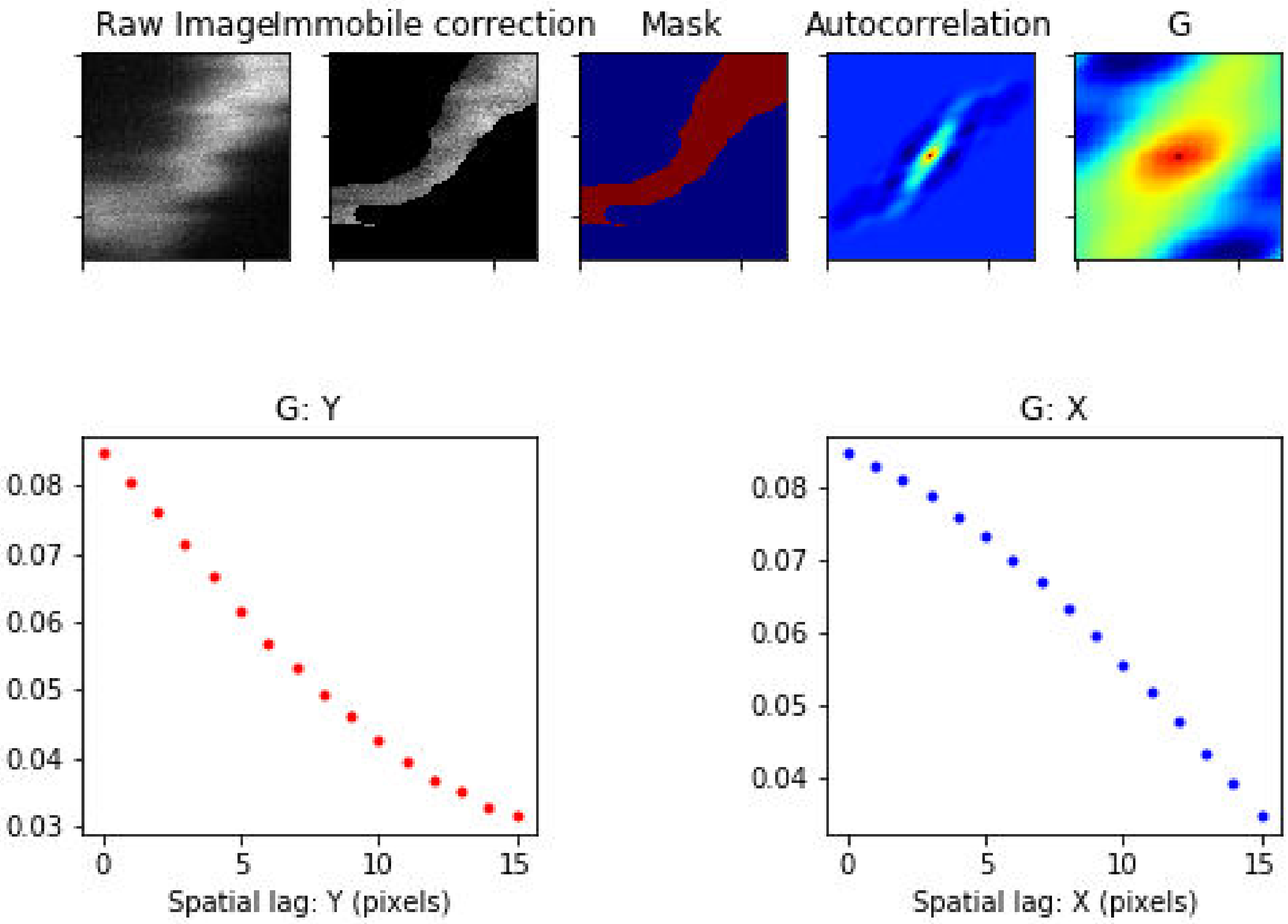

Treatment: Secretin , Domain: PV , File: Image 101.czi

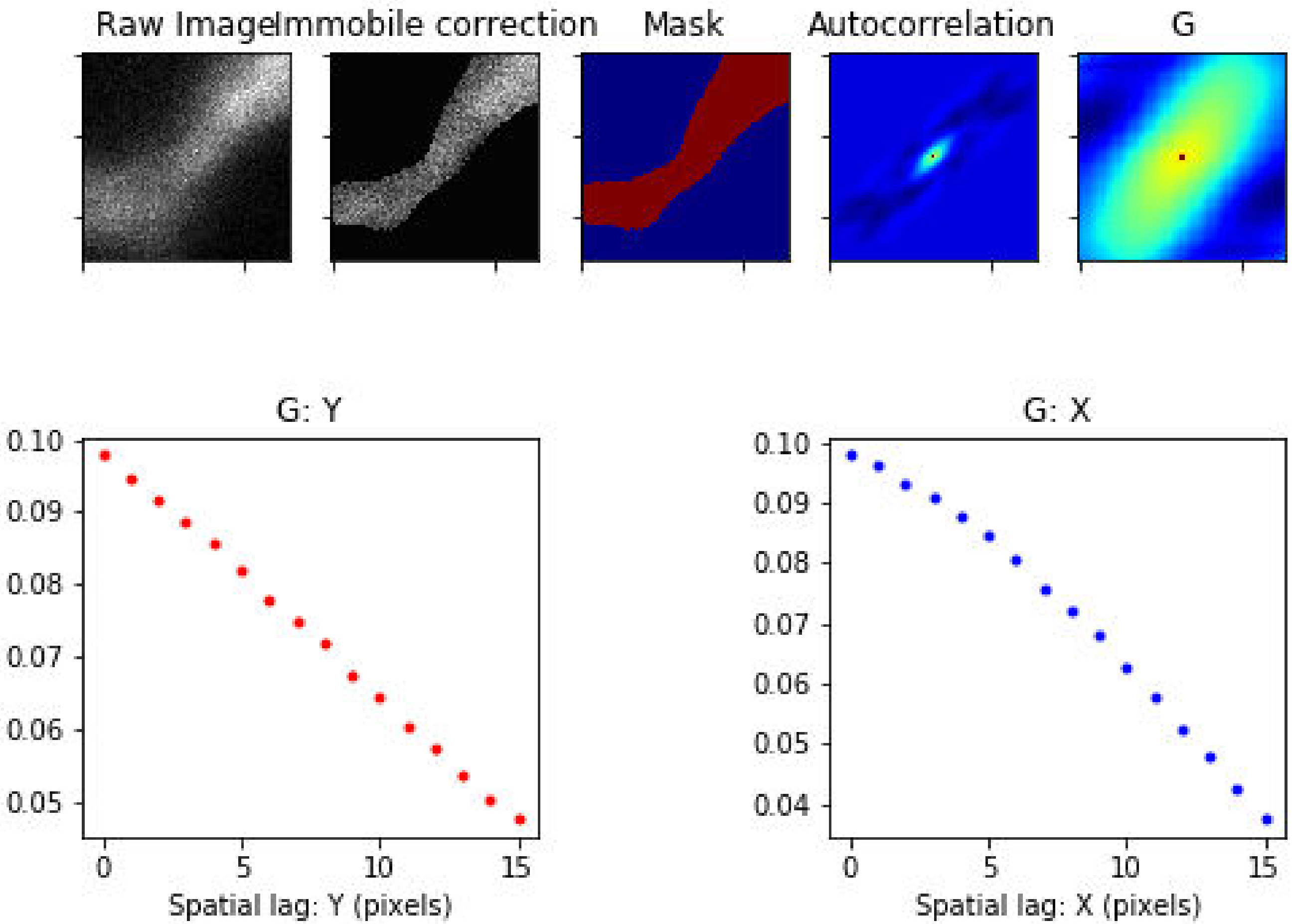

Treatment: Secretin , Domain: PV , File: Image 104.czi

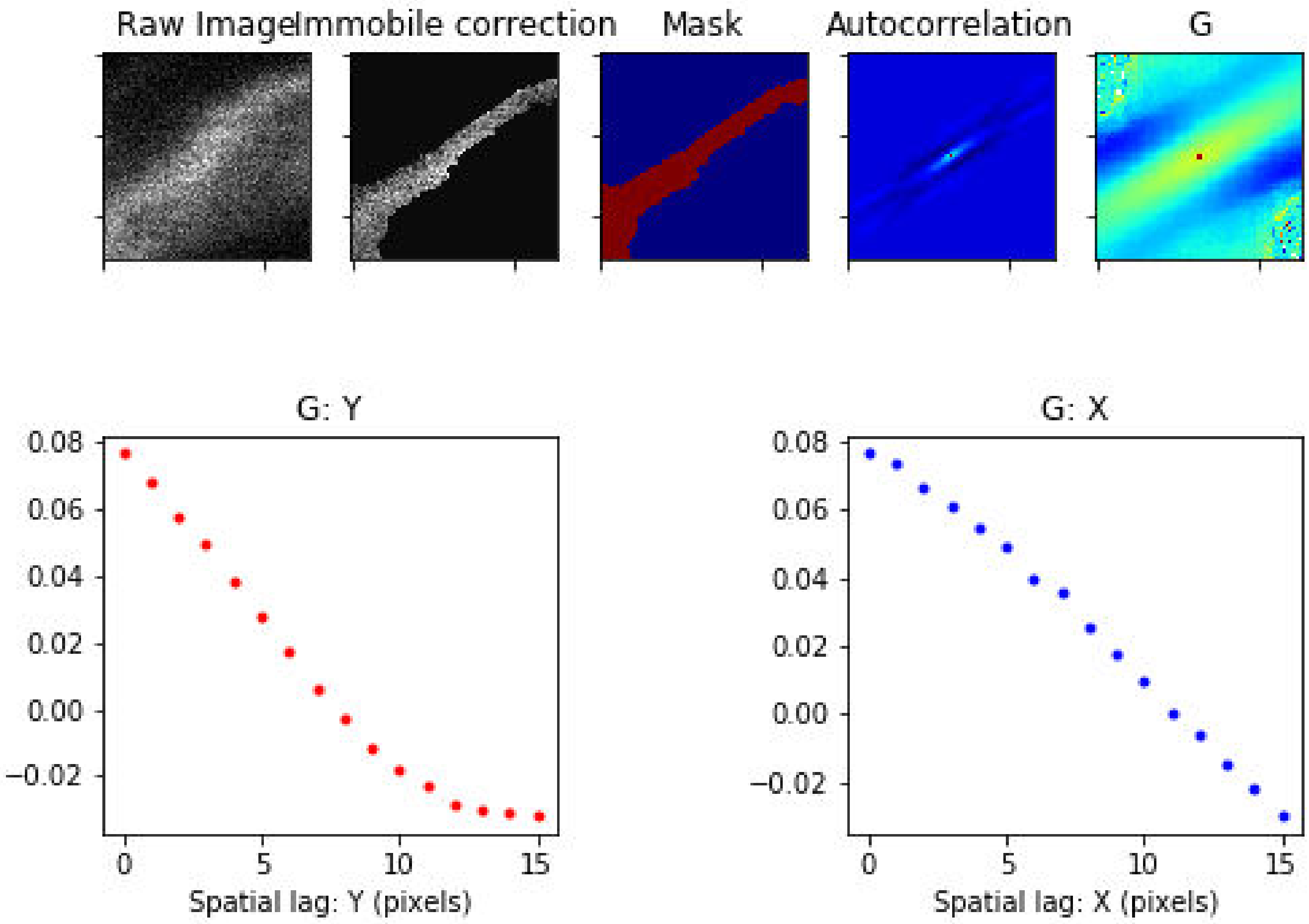

Treatment: Secretin , Domain: PV , File: Image 105.czi

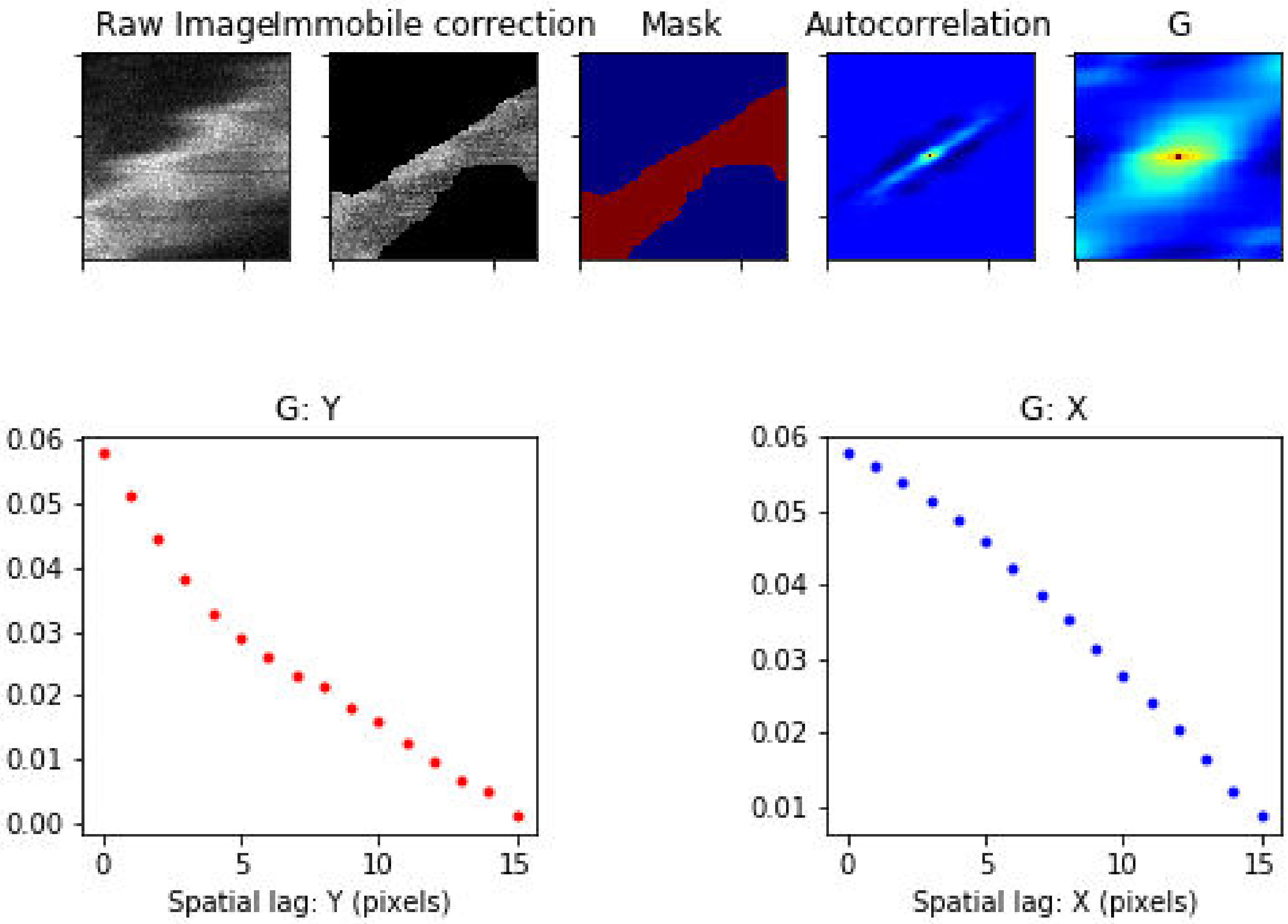

### Treatment: Secretin , Domain: PV , File: Image 107.czi

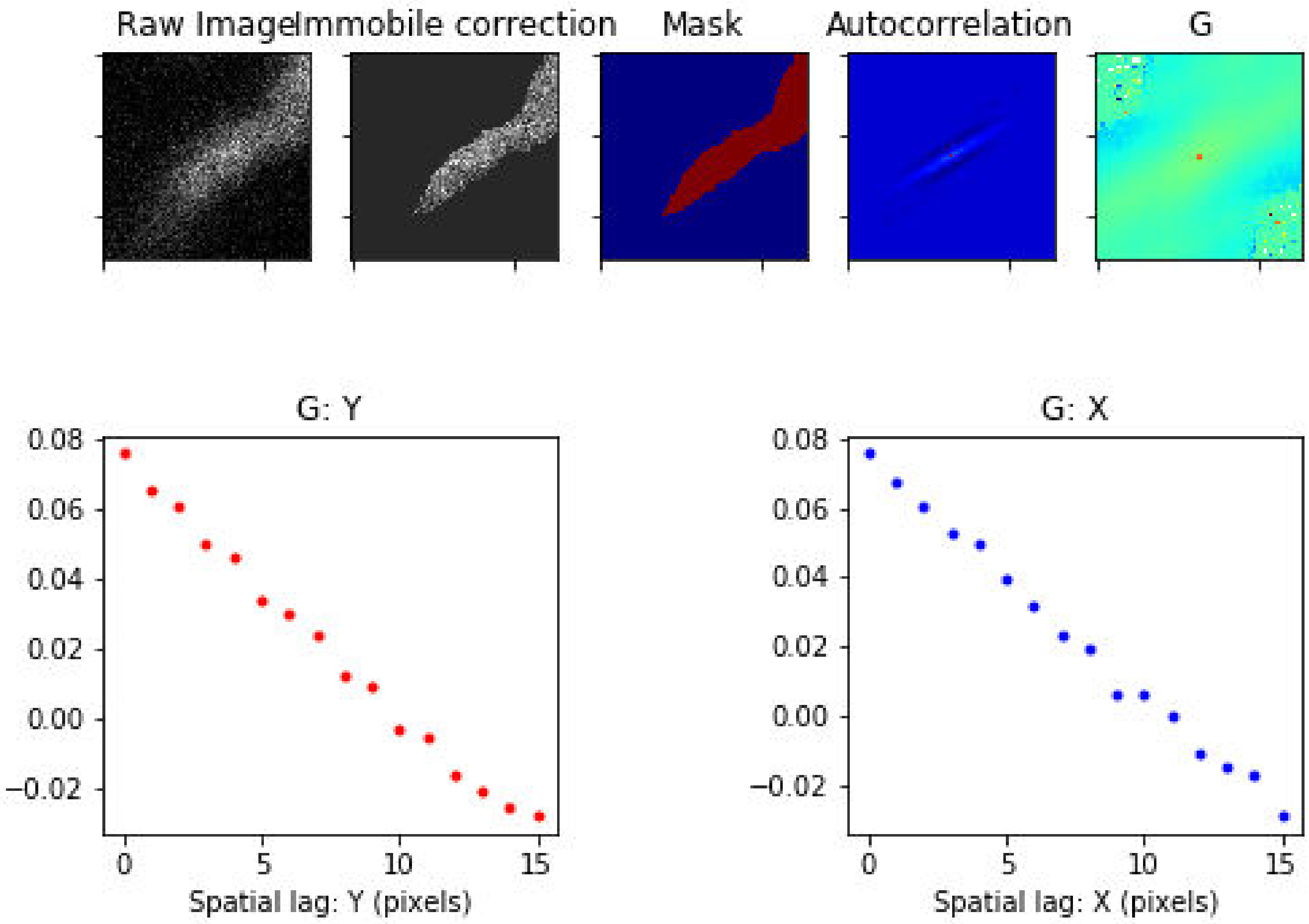

Treatment: Secretin , Domain: PV , File: Image 108.czi

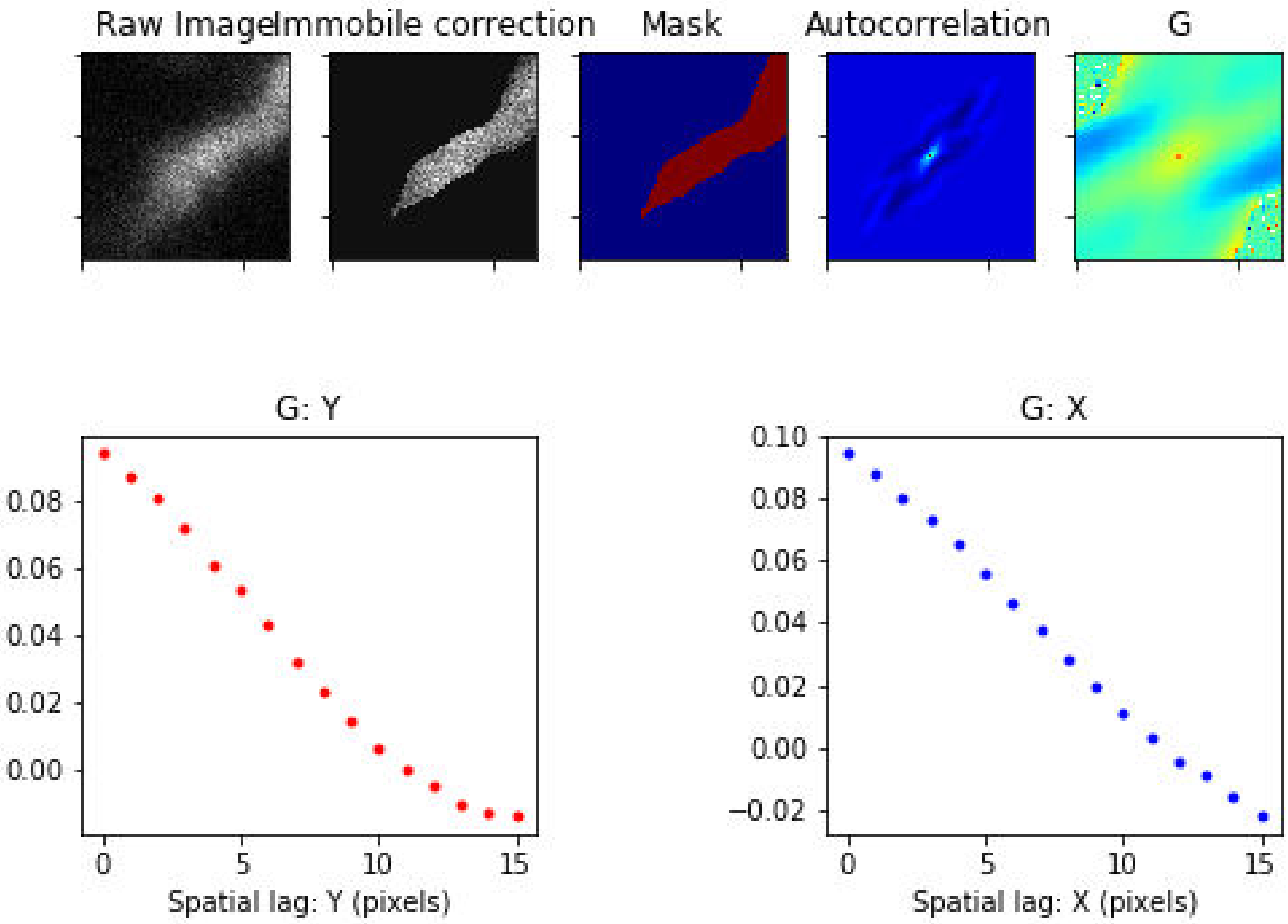

Treatment: Secretin , Domain: PV , File: Image 109.czi

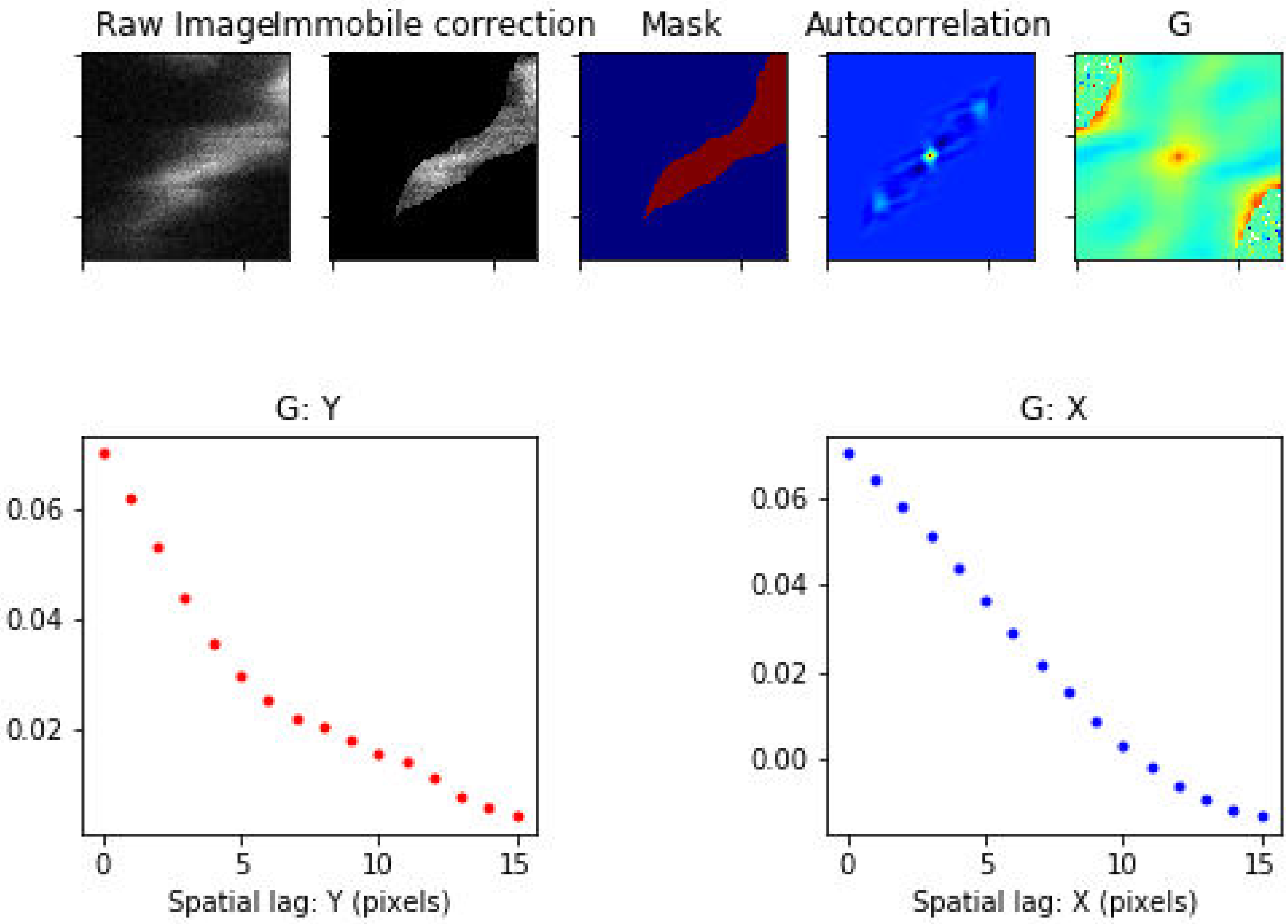

Treatment: TCA , Domain: CV , File: Image 237.czi

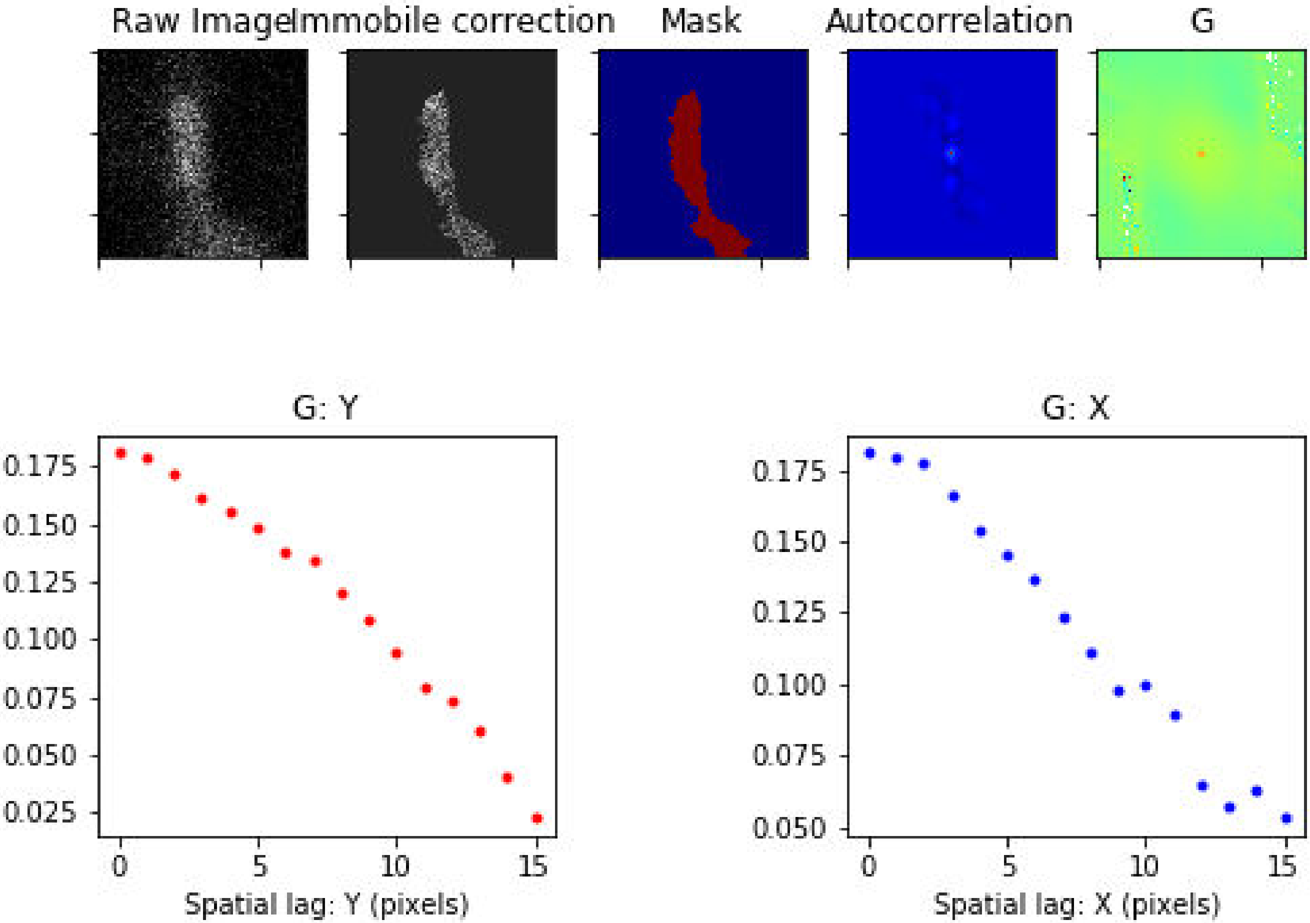

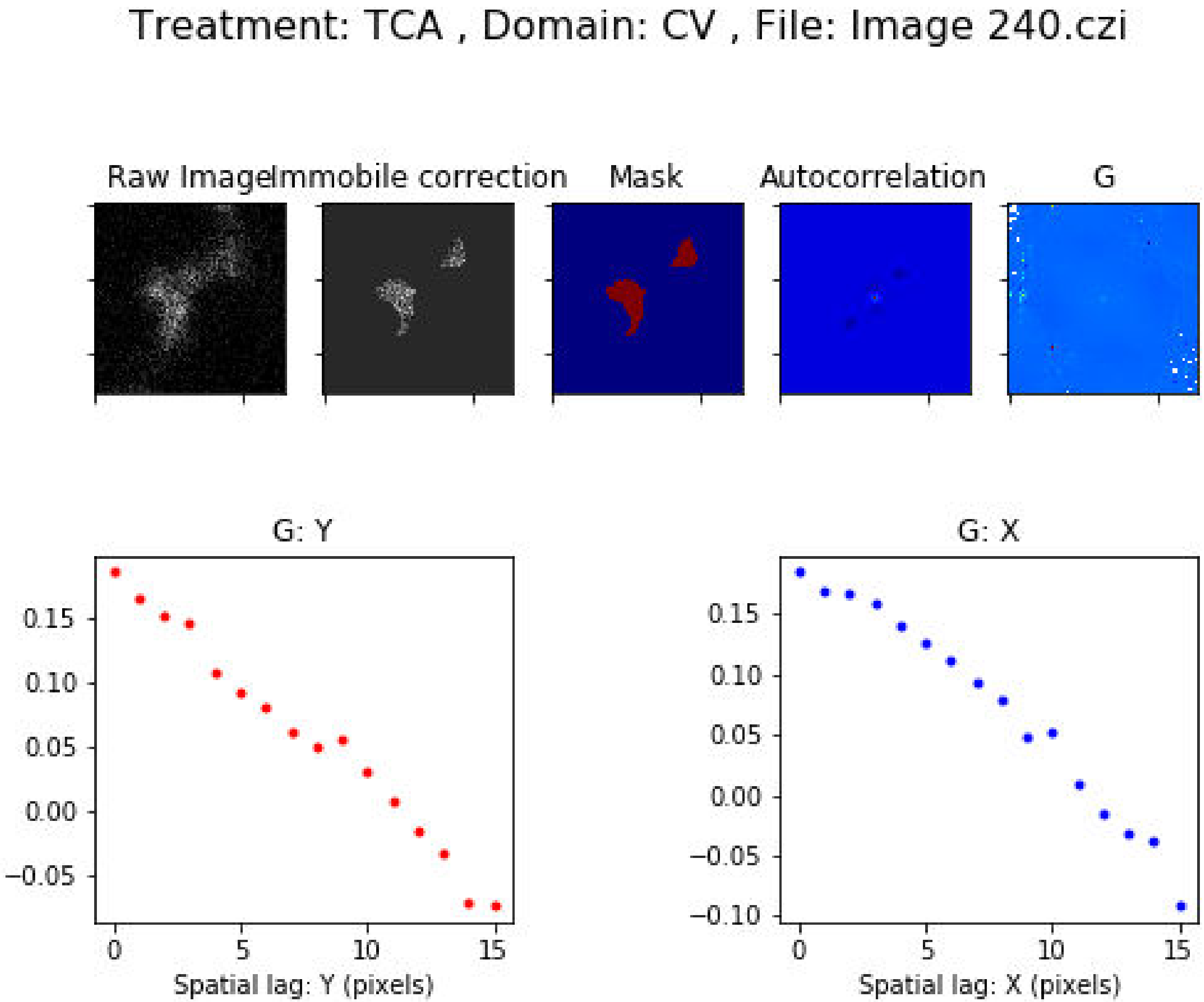
Treatment: TCA , Domain: CV , File: Image 240.czi
Raw Image
Immobile correction
Mask
Autocorrelation
G
G: Y
0.15
0.10
0.05
0.00
−0.05
0
5
10
15
Spatial lag: Y (pixels)
G: X
0.15
0.10
0.05
0.00
−0.05
−0.10
0
5
10
15
Spatial lag: X (pixels)

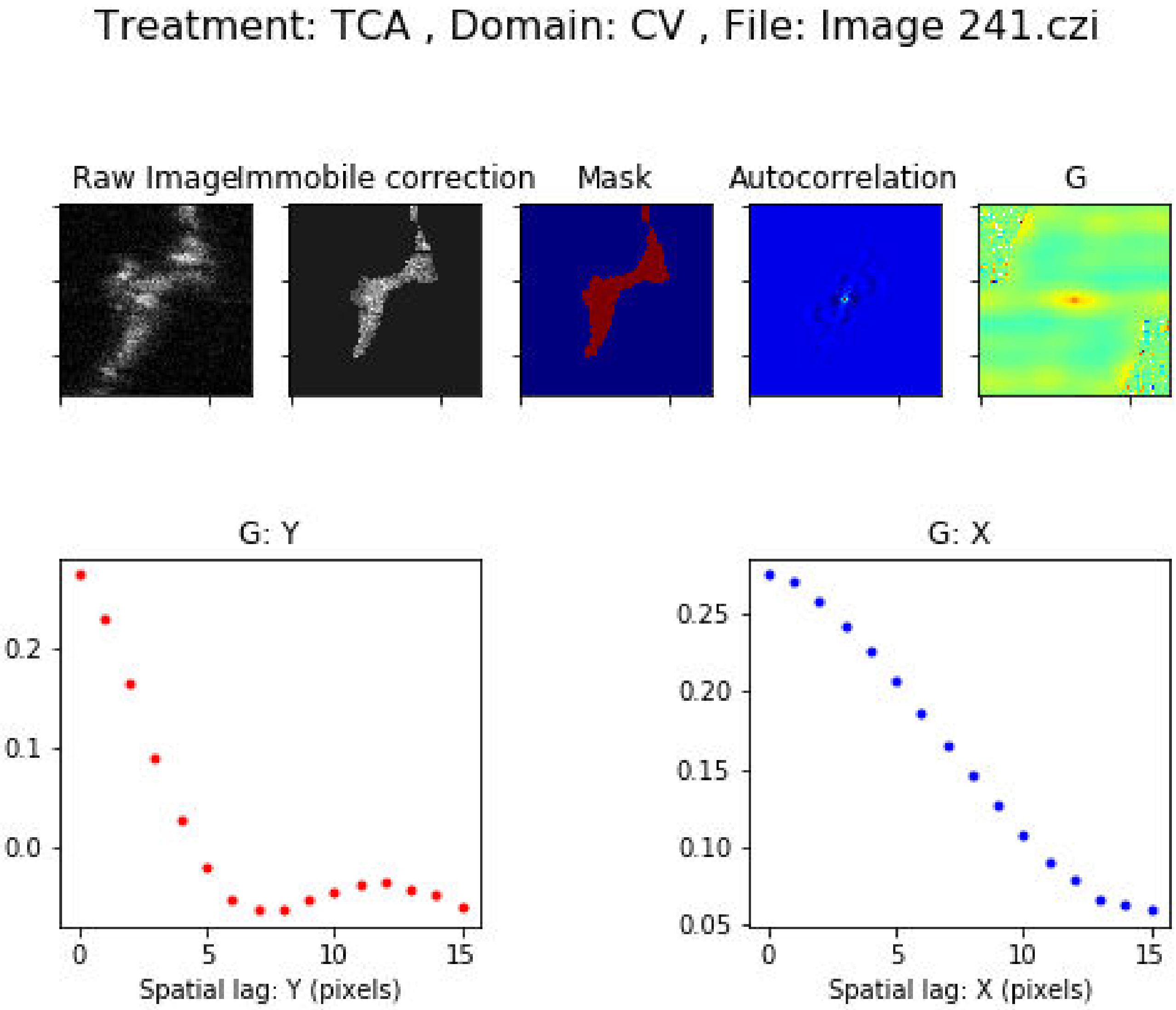
Treatment: TCA , Domain: CV , File: Image 241.czi
Raw Image
Immobile correction
Mask
Autocorrelation
G
G: Y
0.2
0.1
0.0
0
5
10
15
Spatial lag: Y (pixels)
G: X
0.25
0.20
0.15
0.10
0.05
0
5
10
15
Spatial lag: X (pixels)

### Treatment: TCA , Domain: CV , File: Image 243.czi

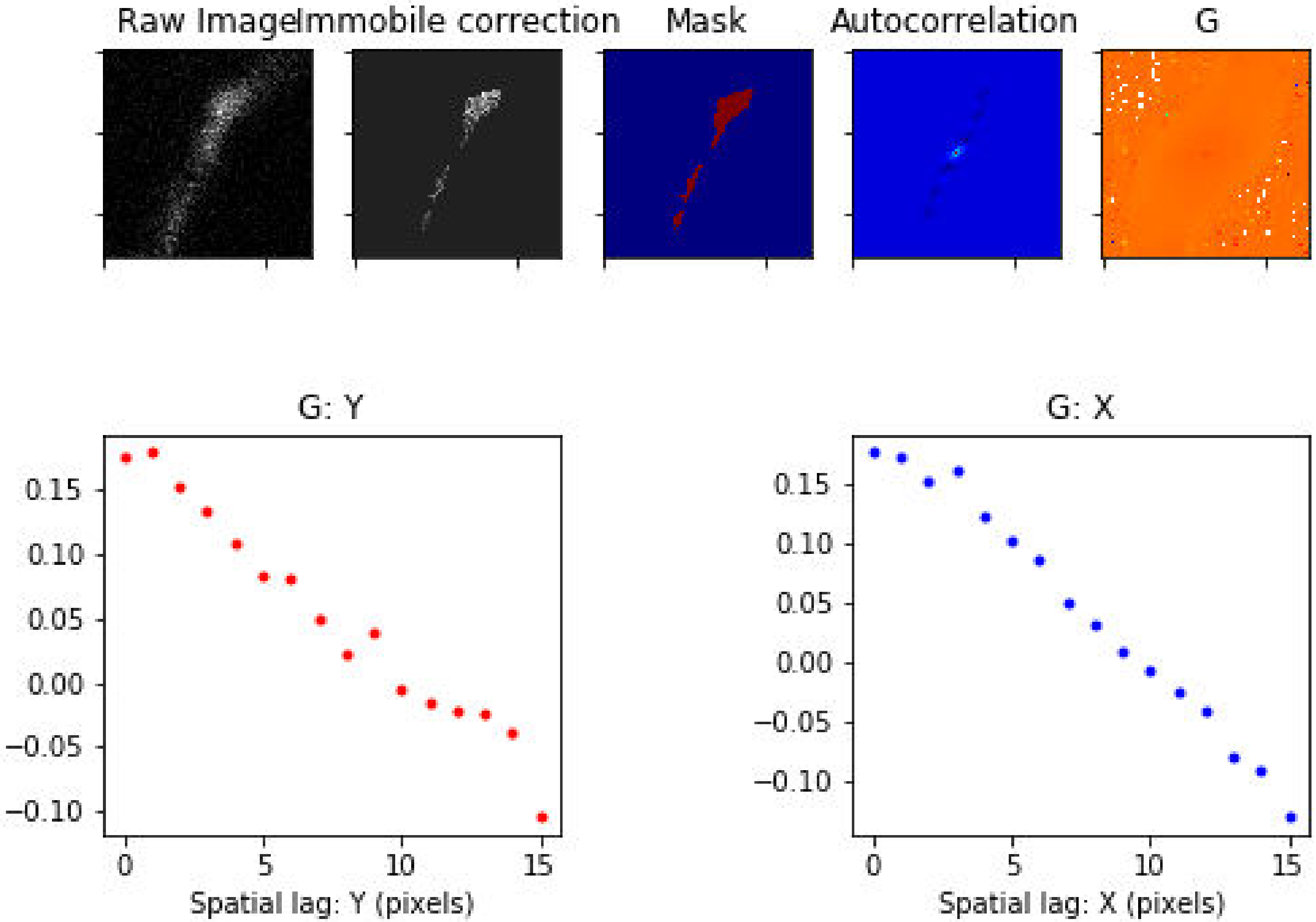

Treatment: TCA , Domain: CV , File: Image 286.czi

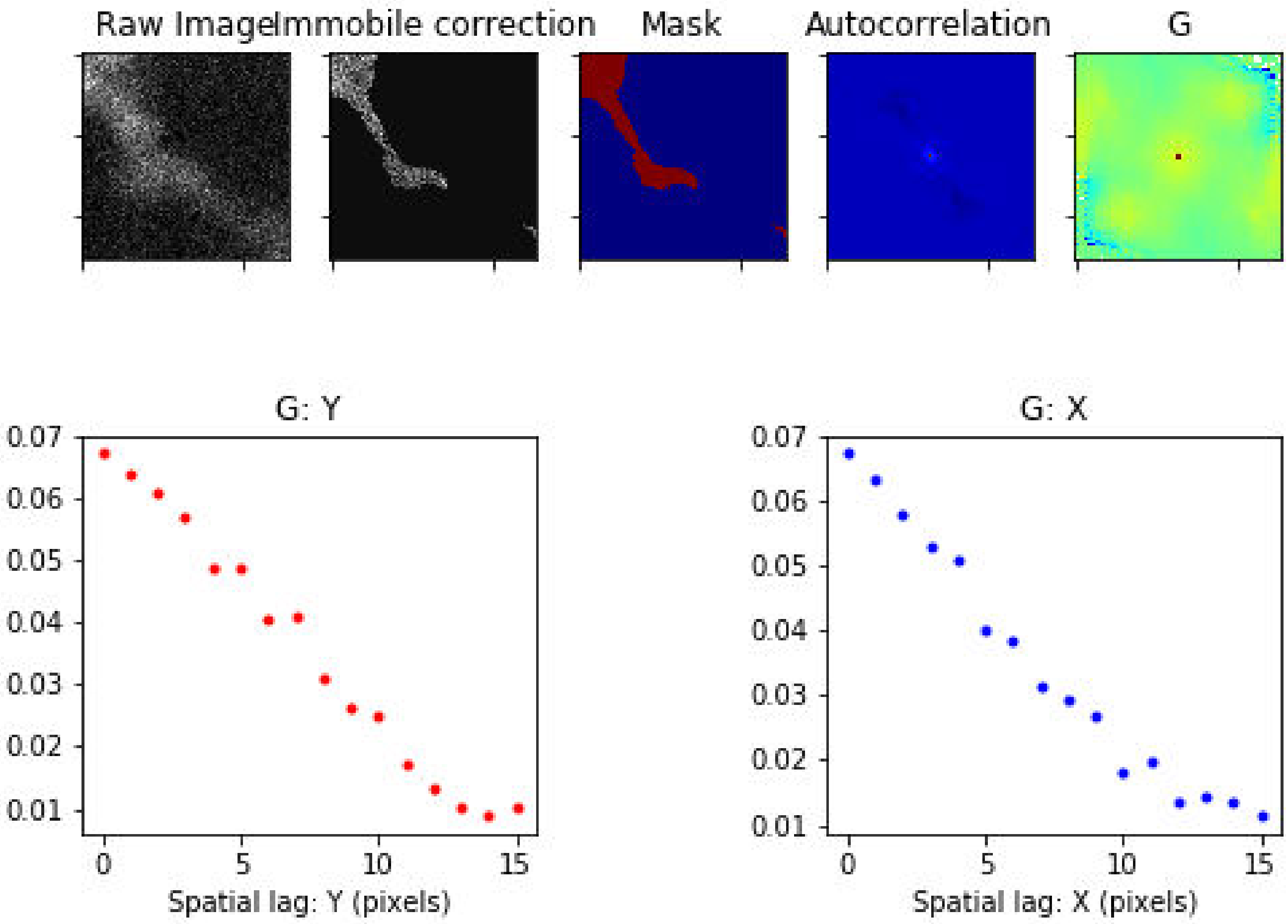

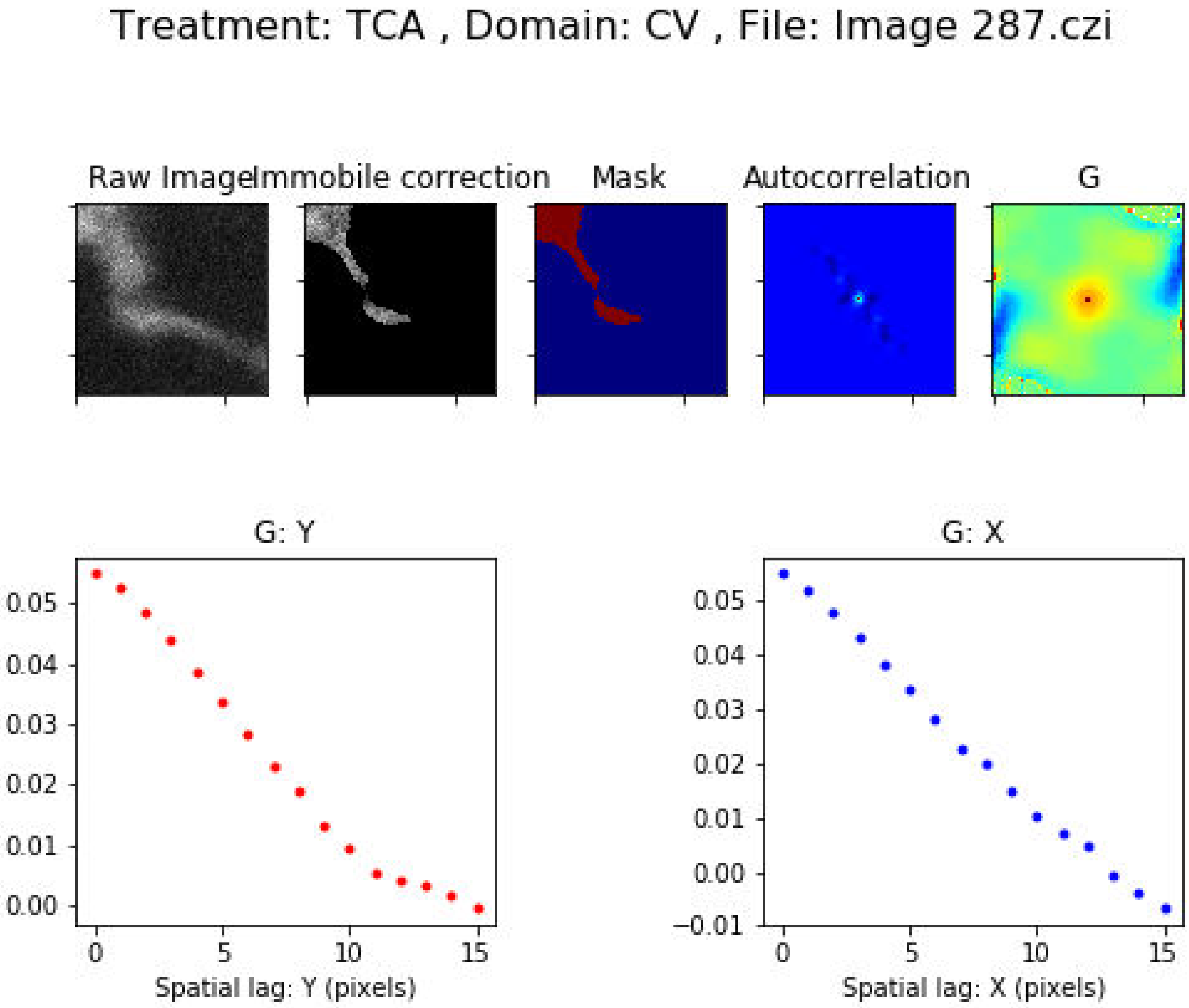
Treatment: TCA , Domain: CV , File: Image 287.czi
Raw Image
Immobile correction
Mask
Autocorrelation
G
G: Y
0.05
0.04
0.03
0.02
0.01
0.00
0
5
10
15
Spatial lag: Y (pixels)
G: X
0.05
0.04
0.03
0.02
0.01
0.00
−0.01
0
5
10
15
Spatial lag: X (pixels)

Treatment: TCA , Domain: CV , File: Image 288.czi

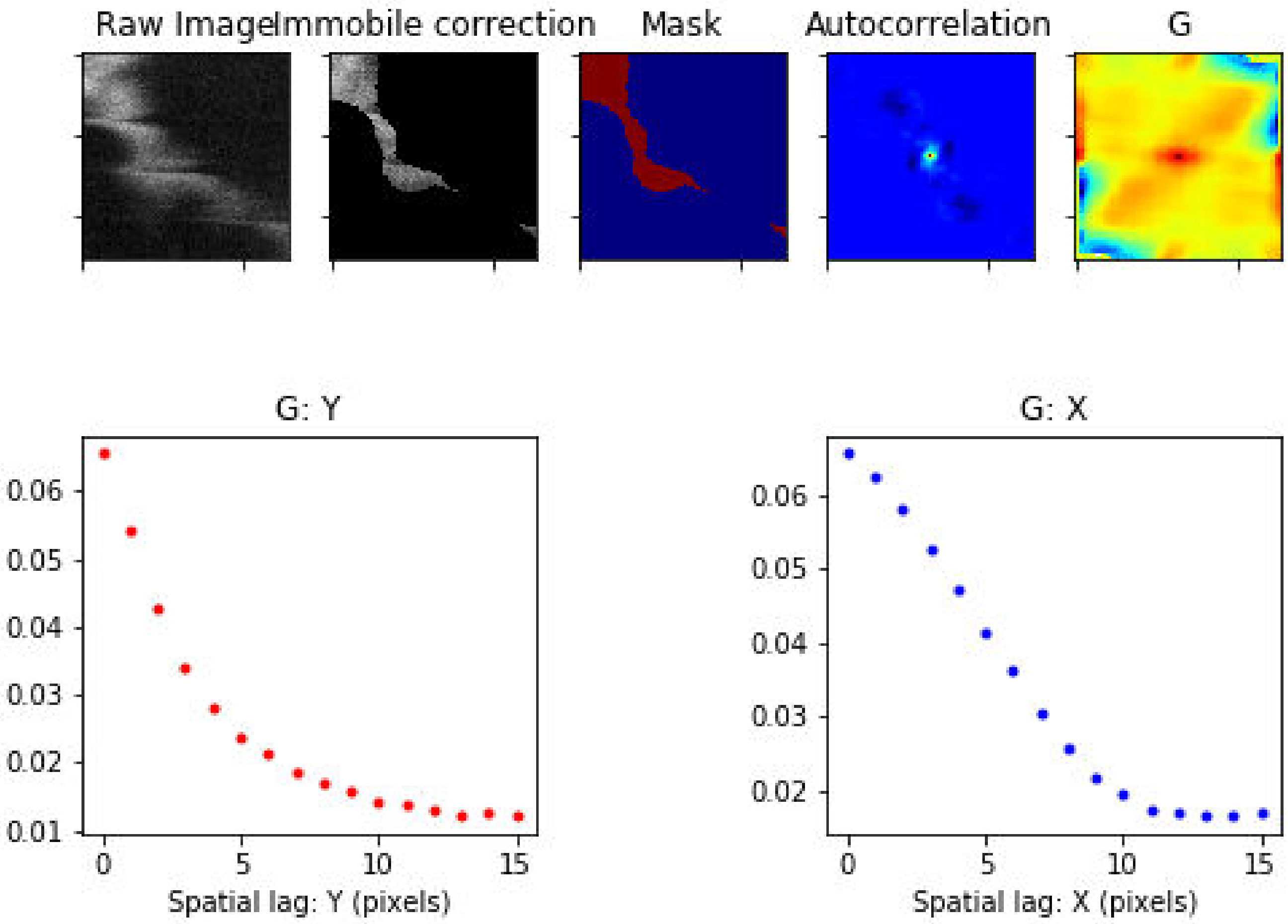

Treatment: TCA , Domain: CV , File: Image 289.czi

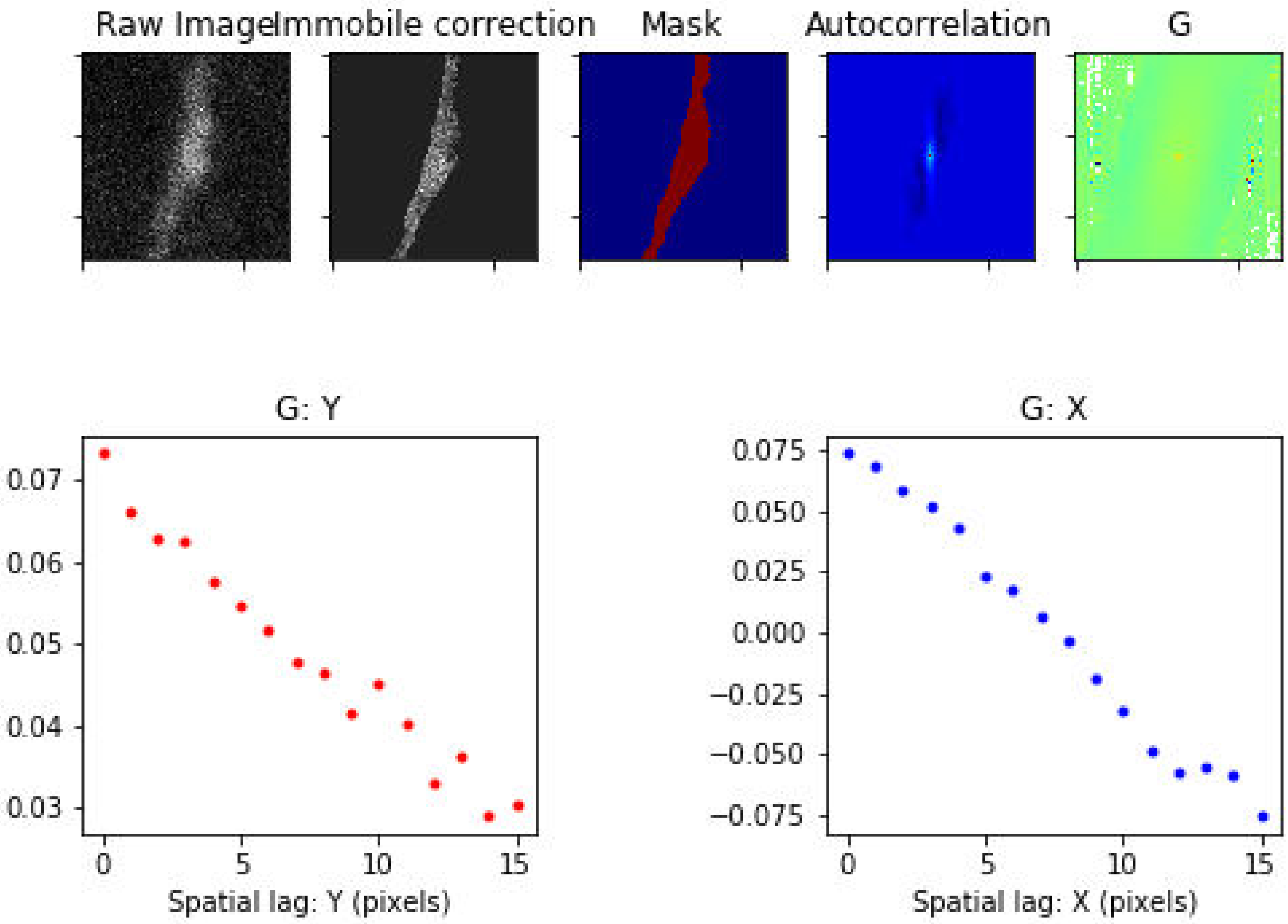

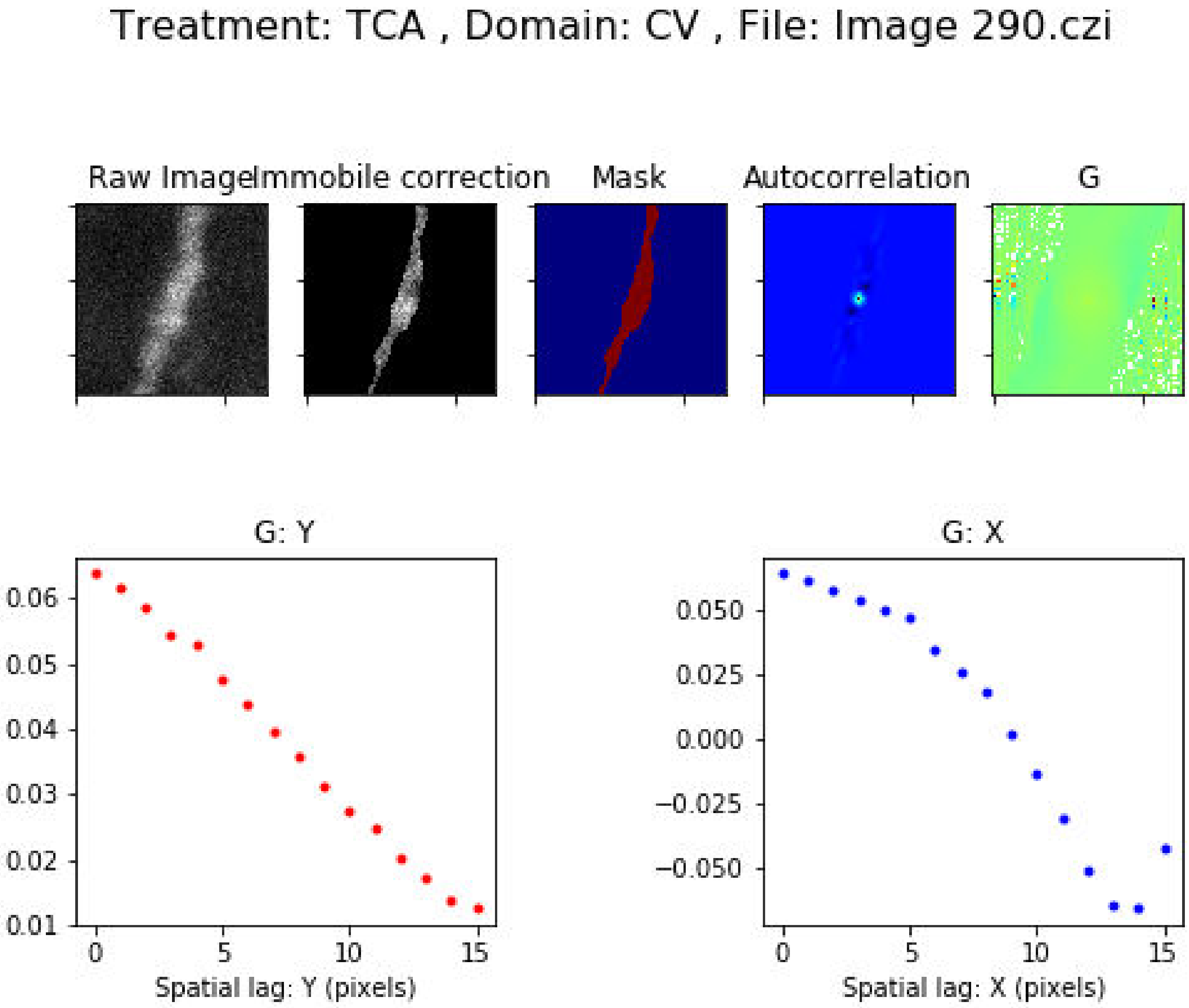
Treatment: TCA , Domain: CV , File: Image 290.czi
Raw Image
Immobile correction
Mask
Autocorrelation
G
G: Y
0.06
0.05
0.04
0.03
0.02
0.01
0
5
10
15
Spatial lag: Y (pixels)
G: X
0.050
0.025
0.000
−0.025
−0.050
0
5
10
15
Spatial lag: X (pixels)

Treatment: TCA , Domain: CV , File: Image 291.czi

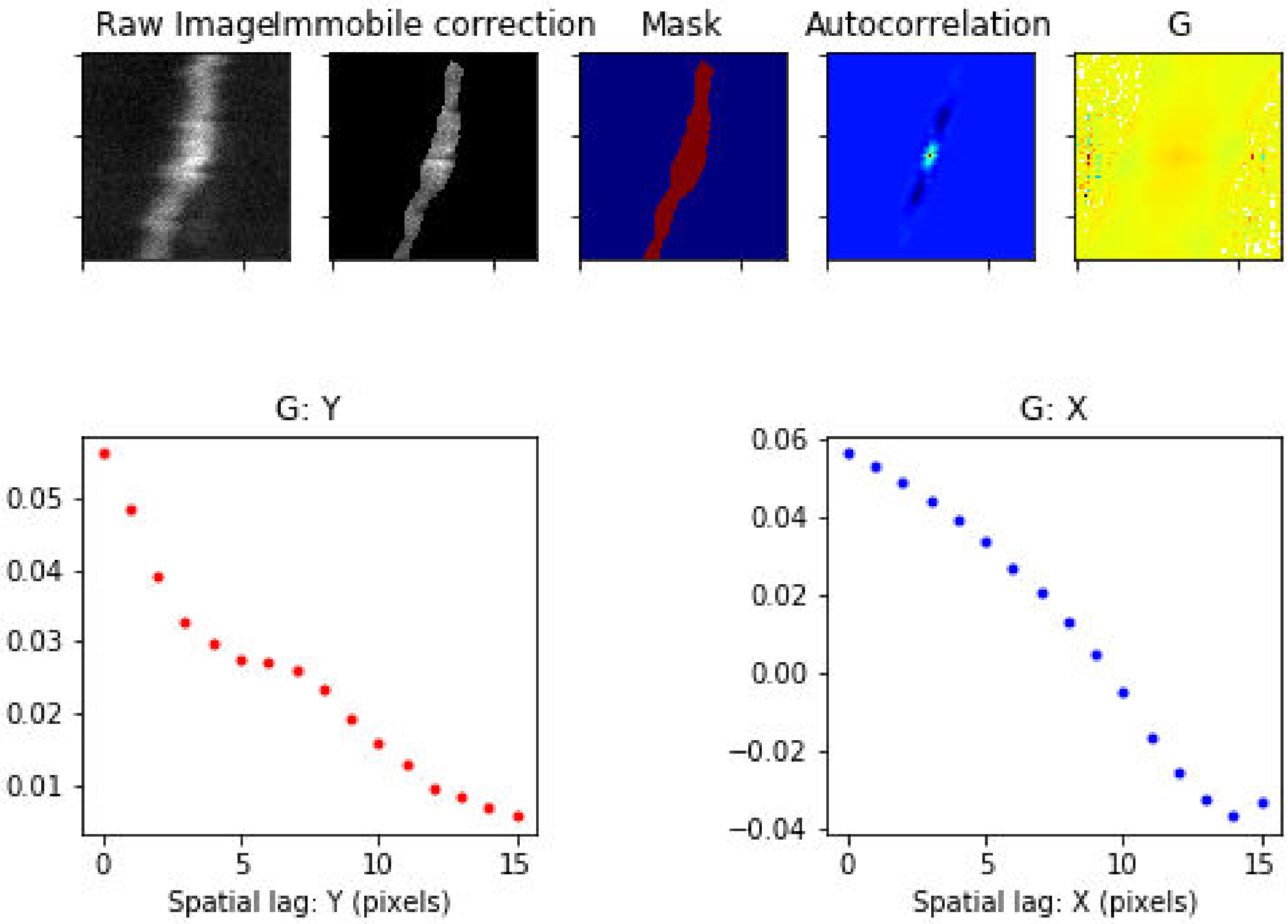

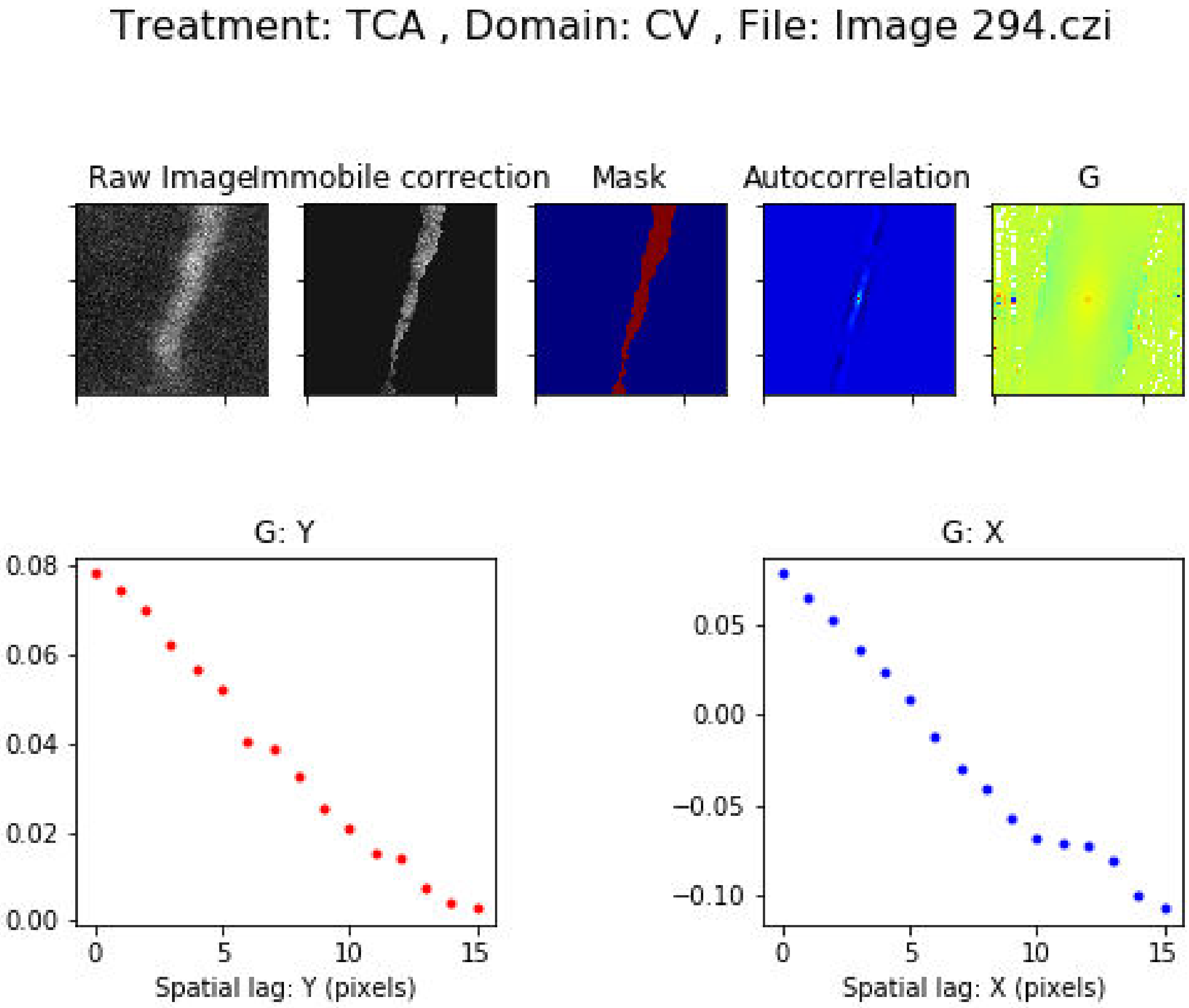

Treatment: TCA , Domain: CV , File: Image 294.czi
Raw Image
Immobile correction
Mask
Autocorrelation
G
G: Y
0.08
0.06
0.04
0.02
0.00
0
5
10
15
Spatial lag: Y (pixels)
G: X
0.05
0.00
−0.05
−0.10
0
5
10
15
Spatial lag: X (pixels)

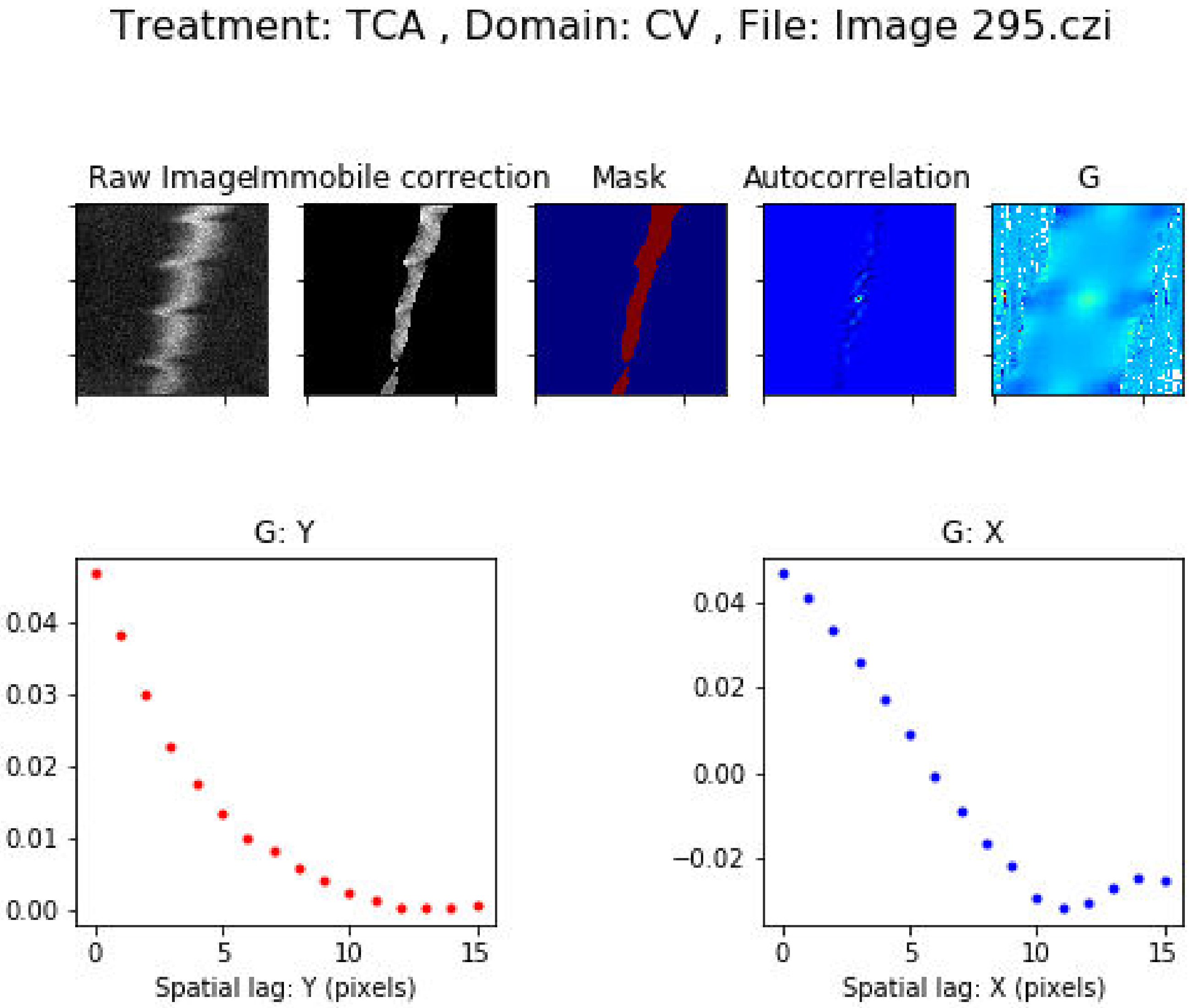

Treatment: TCA , Domain: CV , File: Image 295.czi
Raw Image
Immobile correction
Mask
Autocorrelation
G
G: Y
0.04
0.03
0.02
0.01
0.00
0
5
10
15
Spatial lag: Y (pixels)
G: X
0.04
0.02
0.00
−0.02
0
5
10
15
Spatial lag: X (pixels)

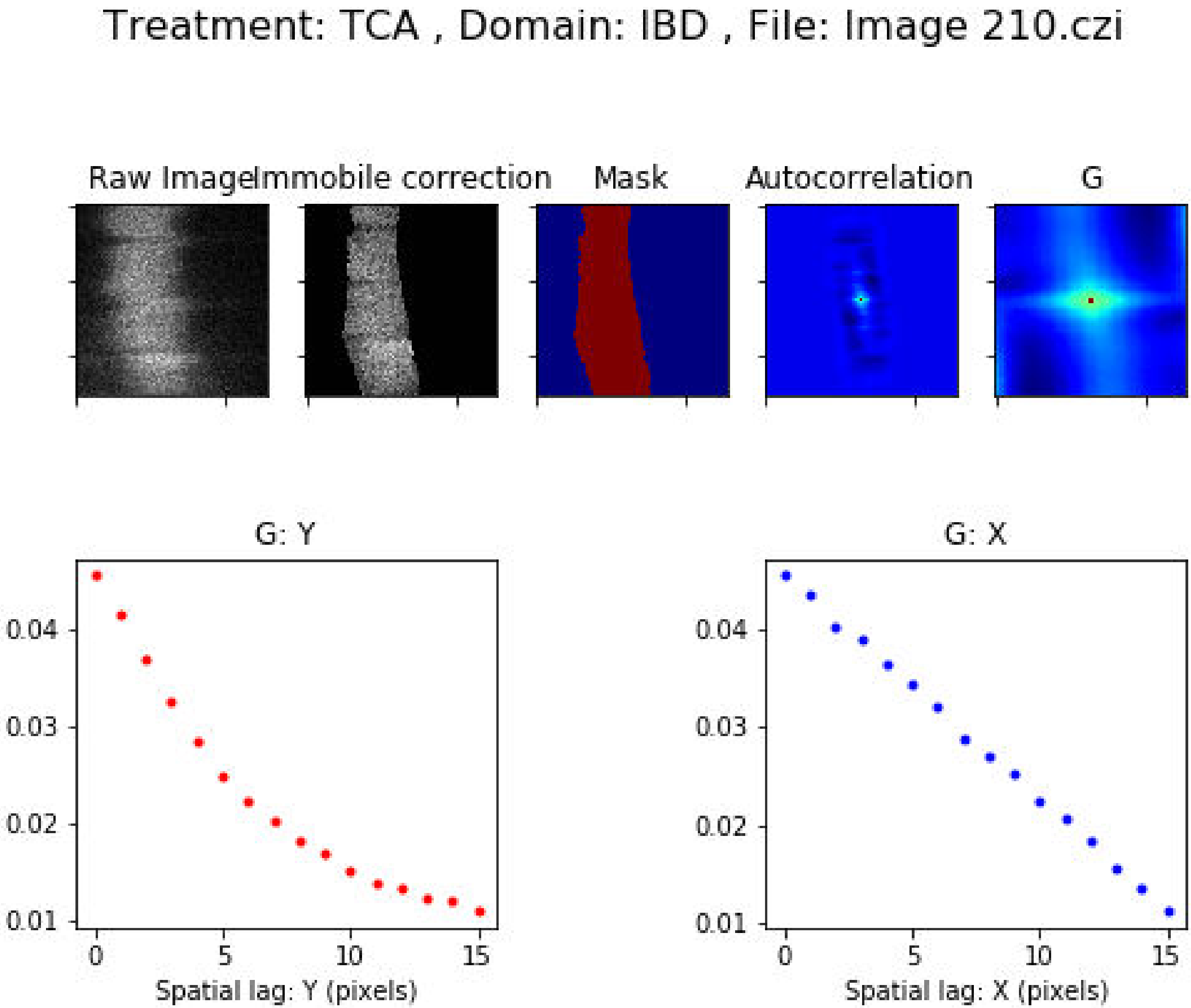

Treatment: TCA , Domain: IBD , File: Image 210.czi
Raw Image
Immobile correction
Mask
Autocorrelation
G
G: Y
0.04
0.03
0.02
0.01
0
5
10
15
Spatial lag: Y (pixels)
G: X
0.04
0.03
0.02
0.01
0
5
10
15
Spatial lag: X (pixels)

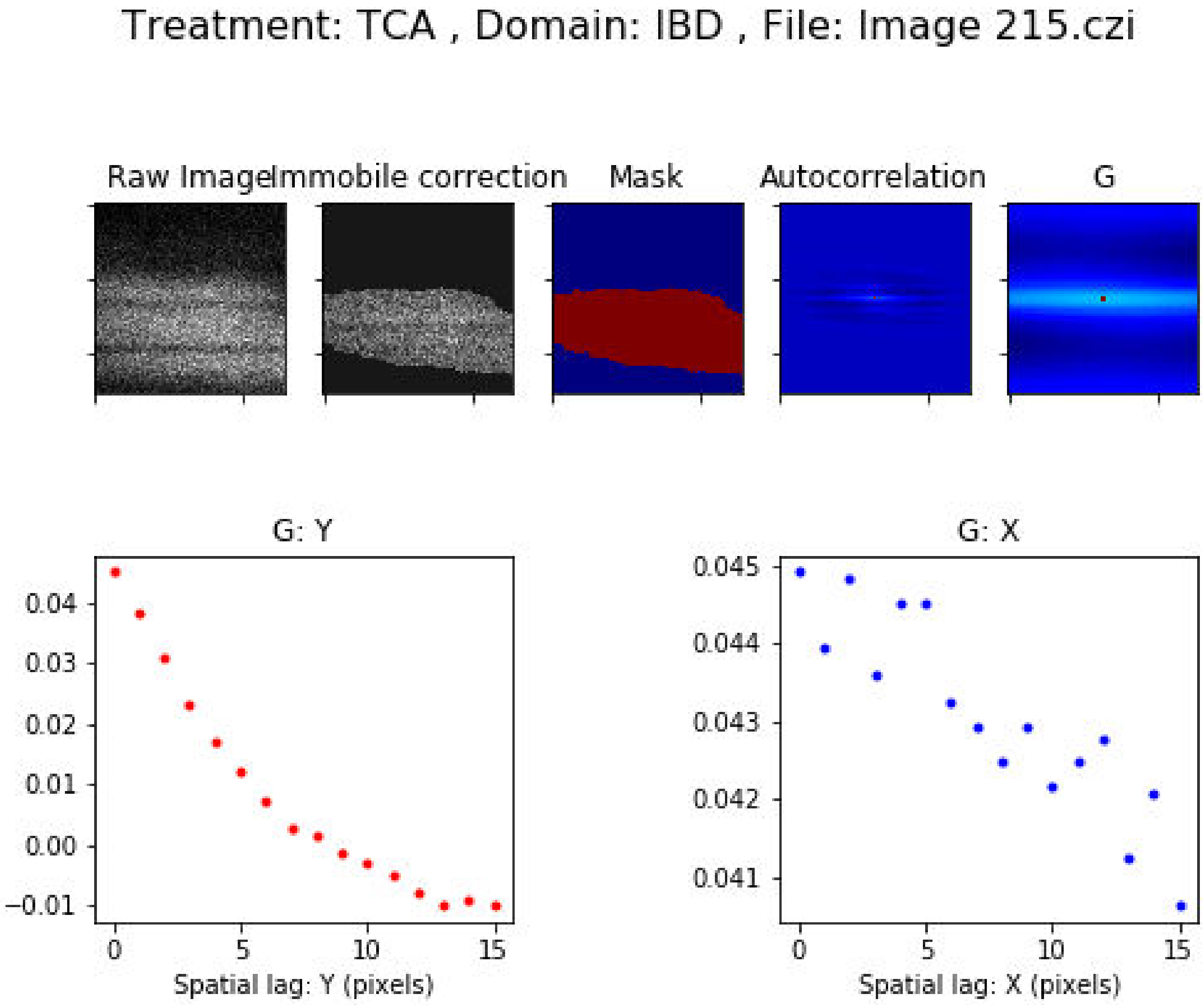
Treatment: TCA , Domain: IBD , File: Image 215.czi
Raw Image
Immobile correction
Mask
Autocorrelation
G
G: Y
0.04
0.03
0.02
0.01
0.00
−0.01
0
5
10
15
Spatial lag: Y (pixels)
G: X
0.045
0.044
0.043
0.042
0.041
0
5
10
15
Spatial lag: X (pixels)

Treatment: TCA , Domain: IBD , File: Image 220.czi

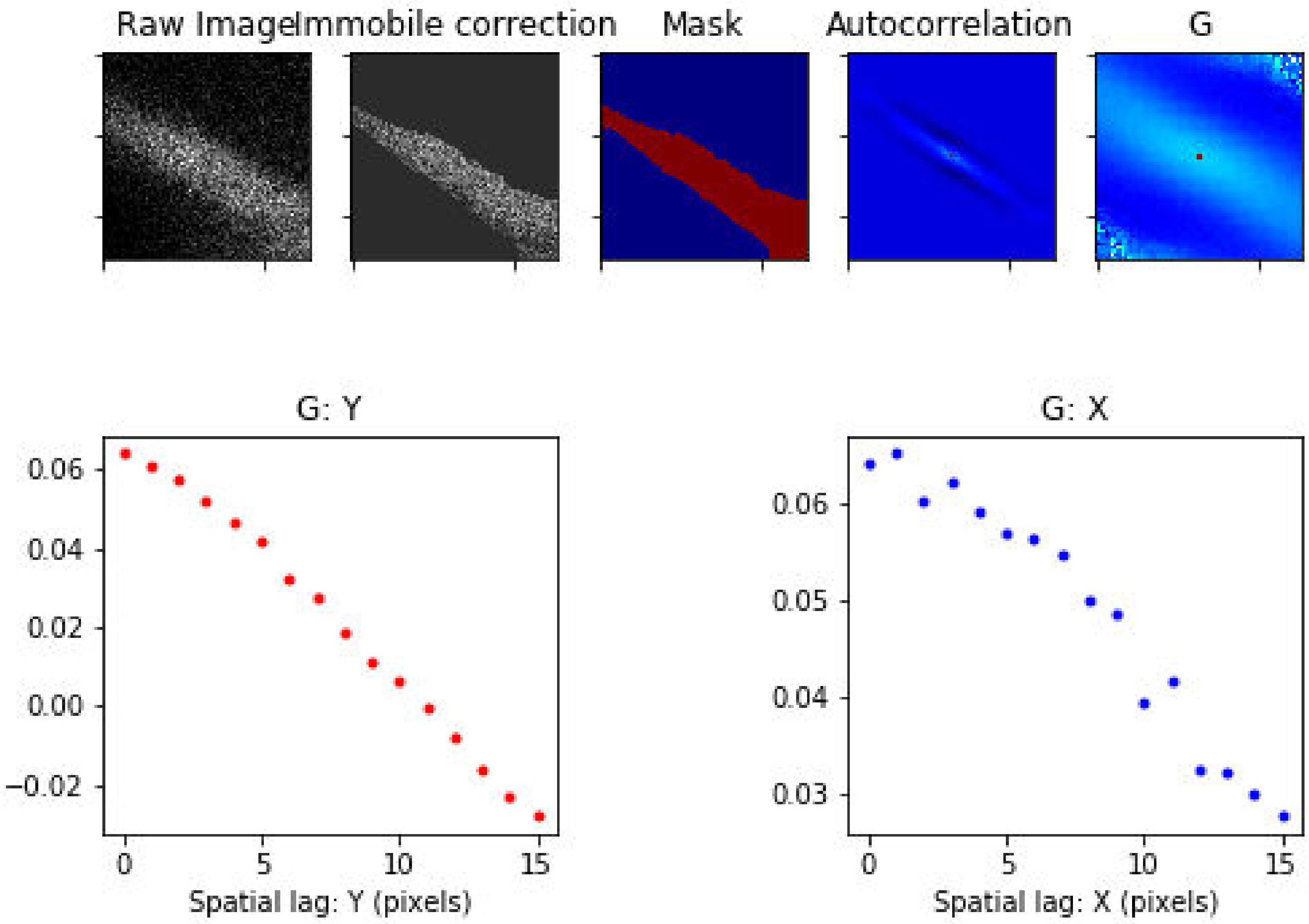

Treatment: TCA , Domain: IBD , File: Image 241.czi

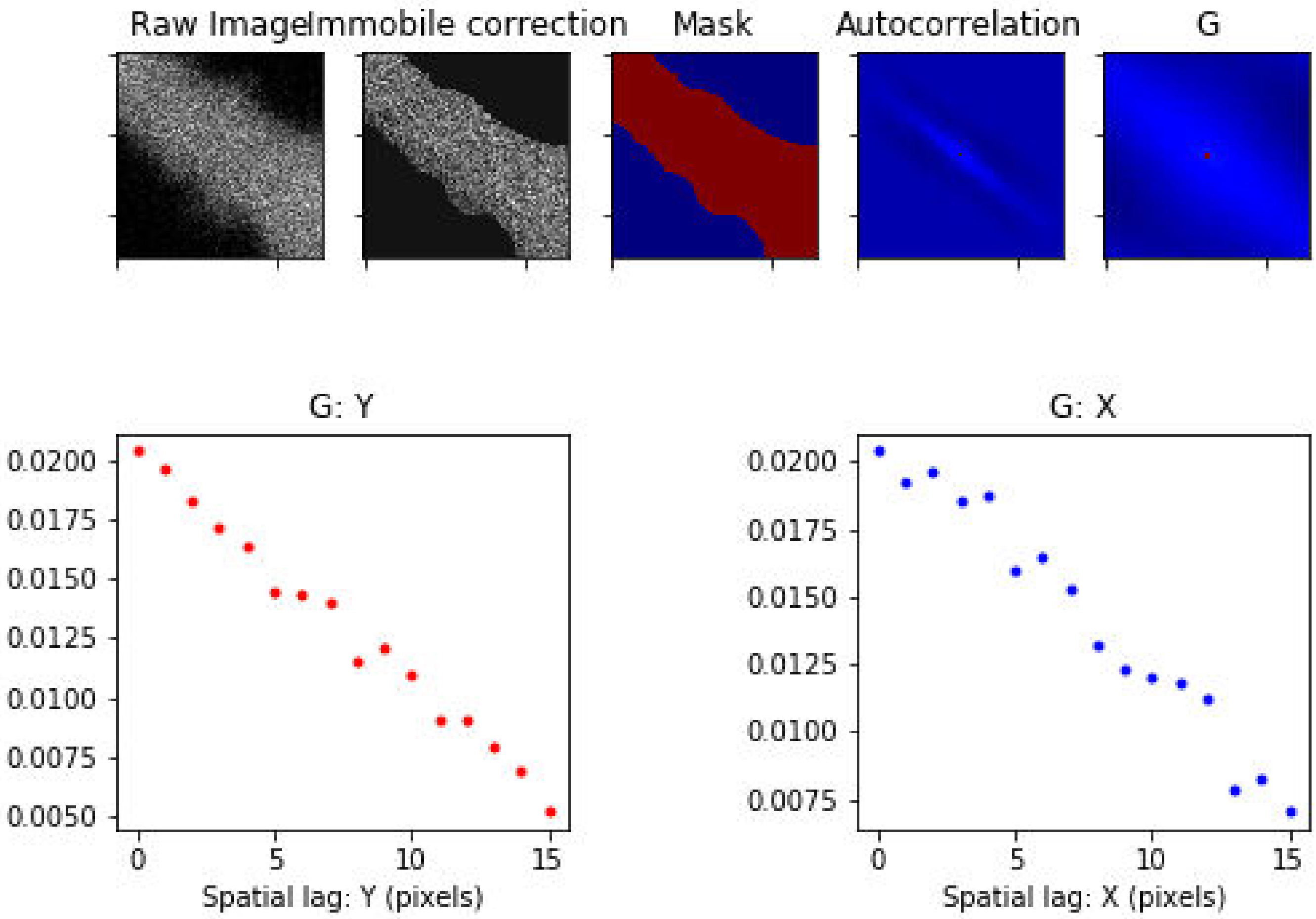

Treatment: TCA , Domain: IBD , File: Image 243.czi

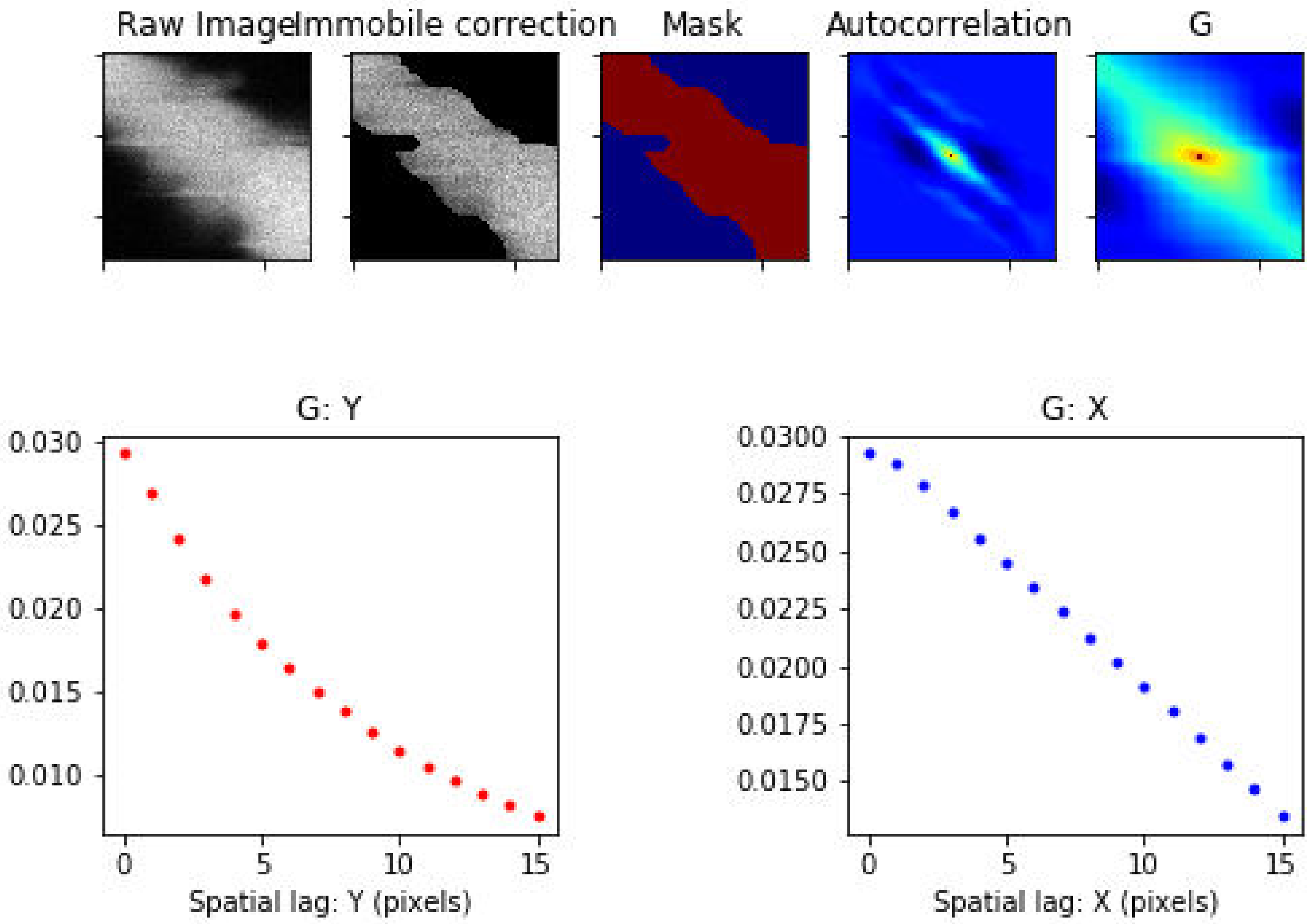

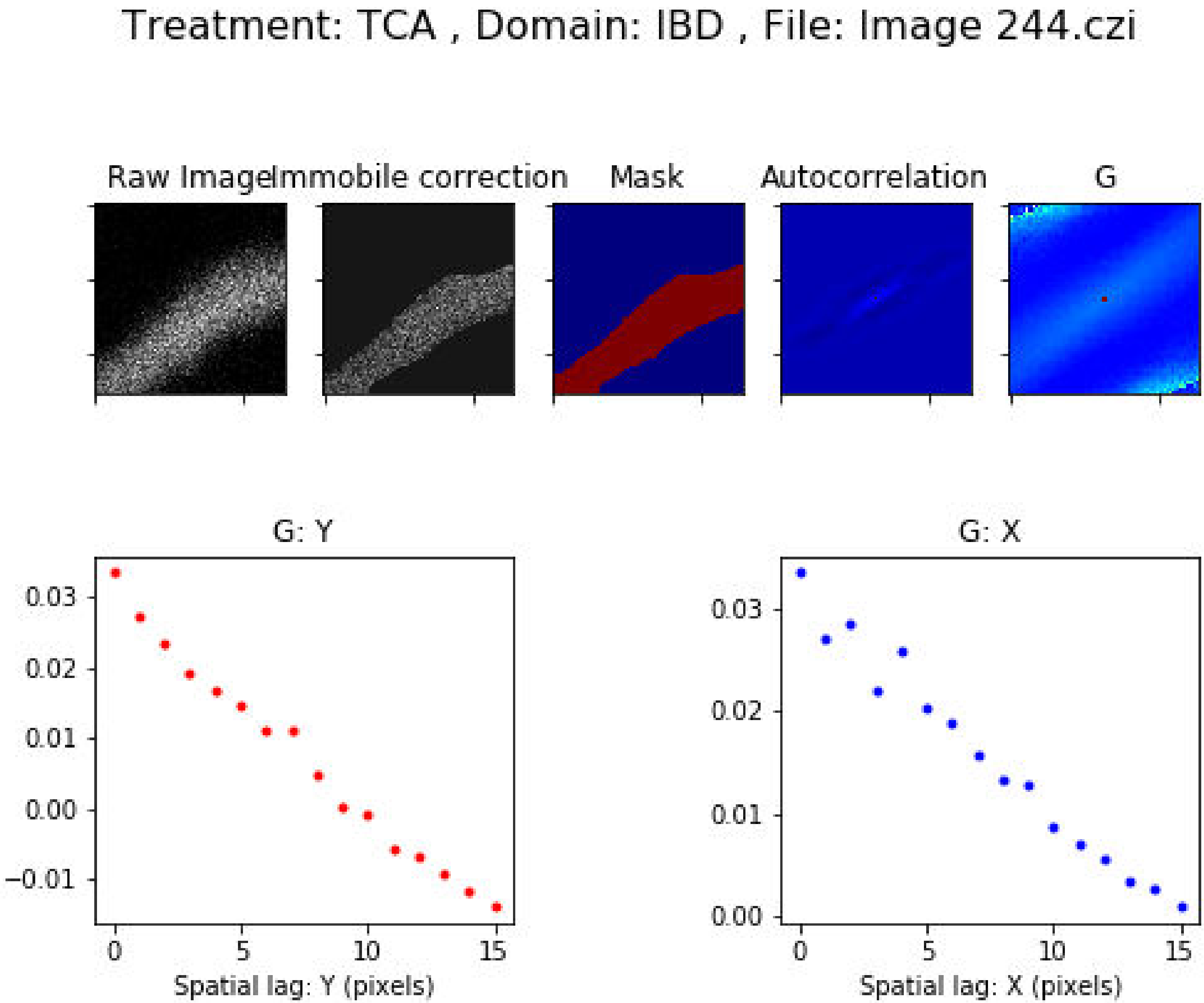

Treatment: TCA , Domain: IBD , File: Image 244.czi
Raw Image
Immobile correction
Mask
Autocorrelation
G
G: Y
0.03
0.02
0.01
0.00
−0.01
0
5
10
15
Spatial lag: Y (pixels)
G: X
0.03
0.02
0.01
0.00
0
5
10
15
Spatial lag: X (pixels)

Treatment: TCA , Domain: IBD , File: Image 245.czi

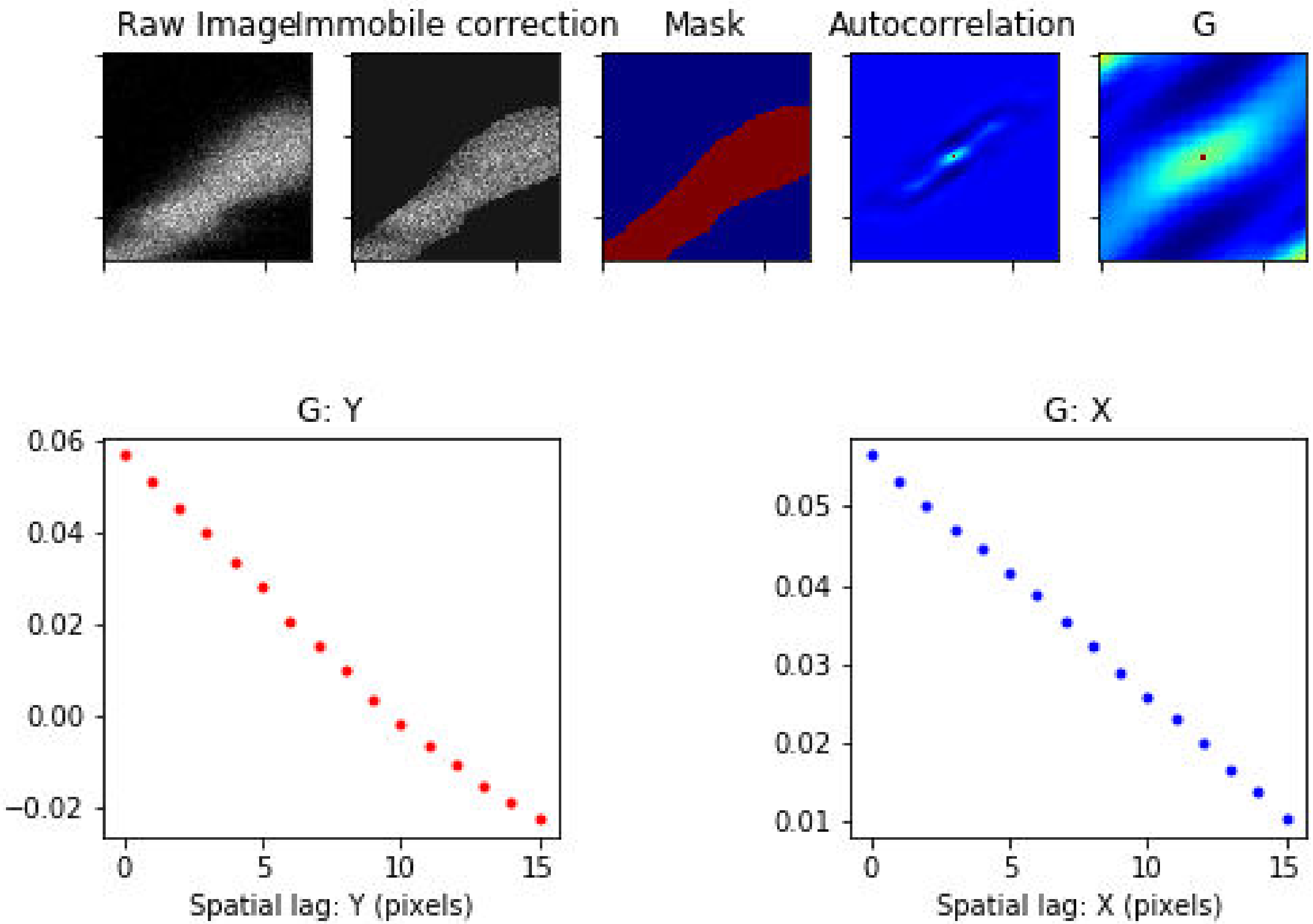

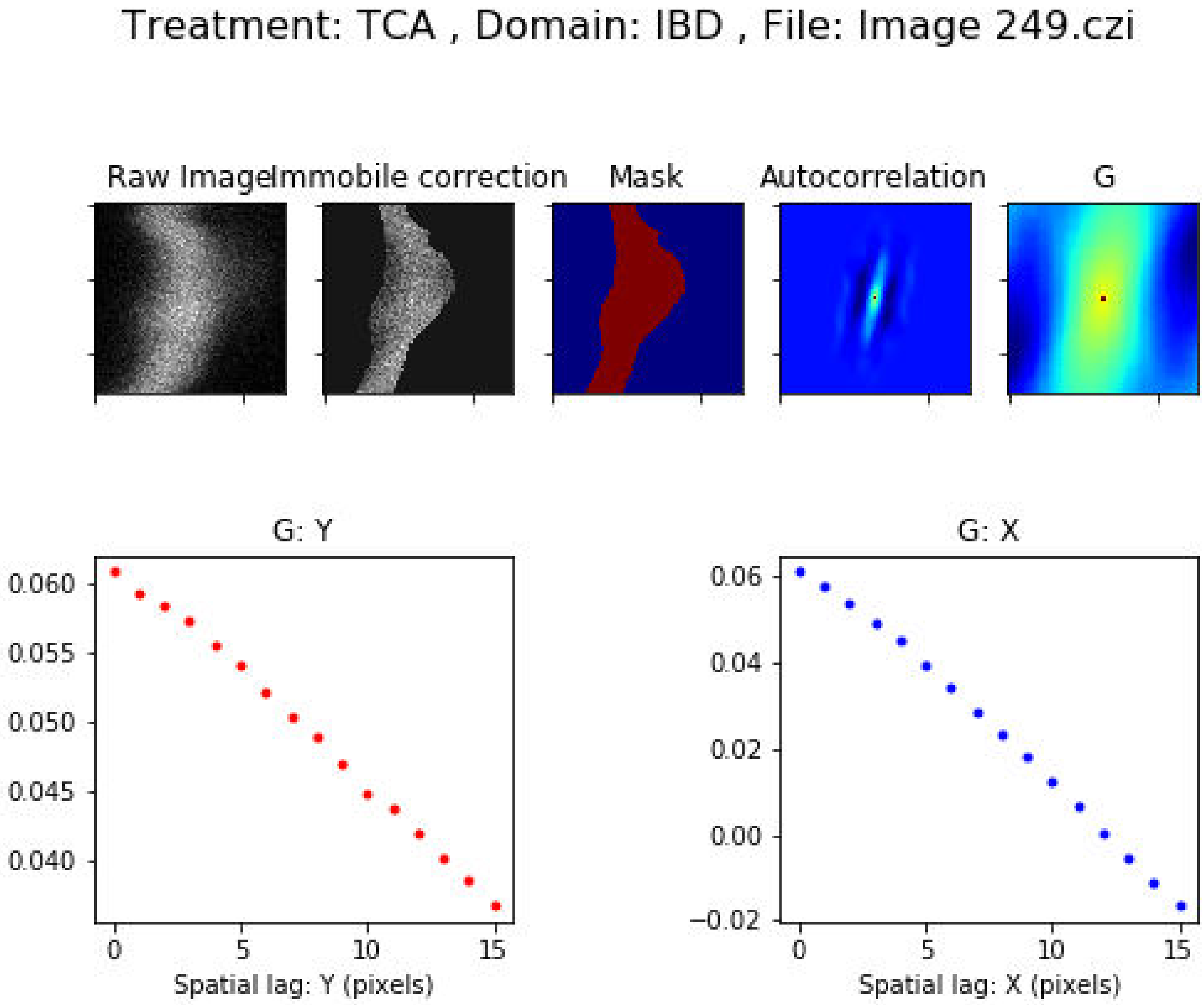

Treatment: TCA , Domain: IBD , File: Image 249.czi
Raw Image
Immobile correction
Mask
Autocorrelation
G
G: Y
0.060
0.055
0.050
0.045
0.040
0 5 10 15
Spatial lag: Y (pixels)
G: X
0.06
0.04
0.02
0.00
−0.02
0 5 10 15
Spatial lag: X (pixels)

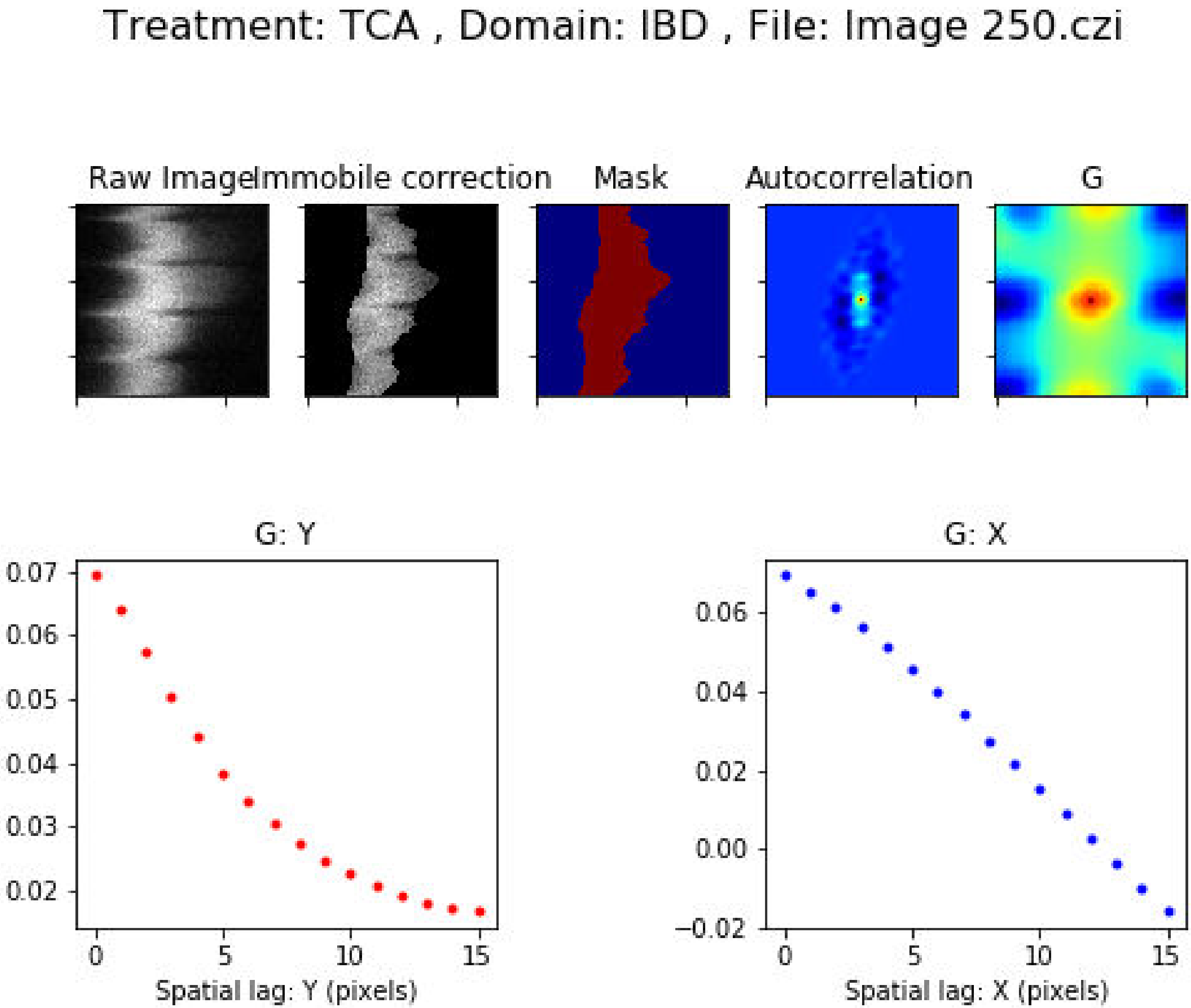
Treatment: TCA , Domain: IBD , File: Image 250.czi
Raw Image
Immobile correction
Mask
Autocorrelation
G
G: Y
0.07
0.06
0.05
0.04
0.03
0.02
0
5
10
15
Spatial lag: Y (pixels)
G: X
0.06
0.04
0.02
0.00
−0.02
0
5
10
15
Spatial lag: X (pixels)

### Treatment: TCA , Domain: IBD , File: Image 253.czi

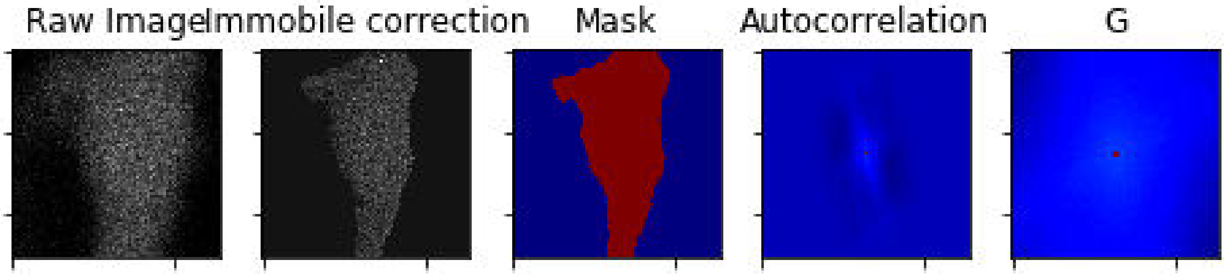


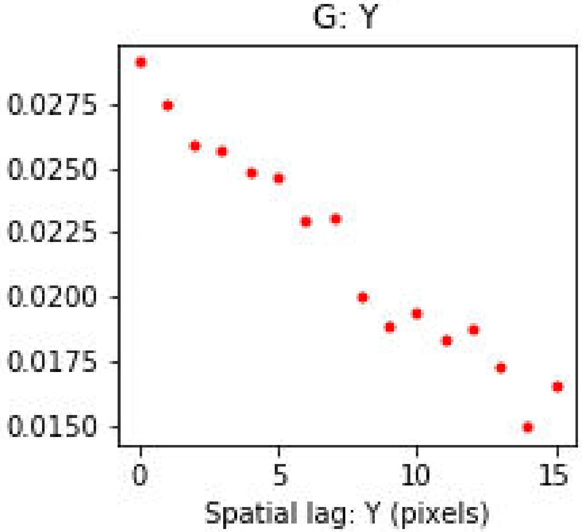


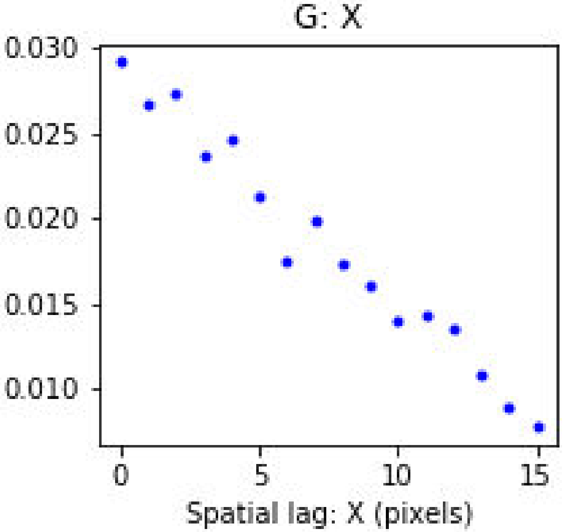

Treatment: TCA , Domain: IBD , File: Image 254.czi

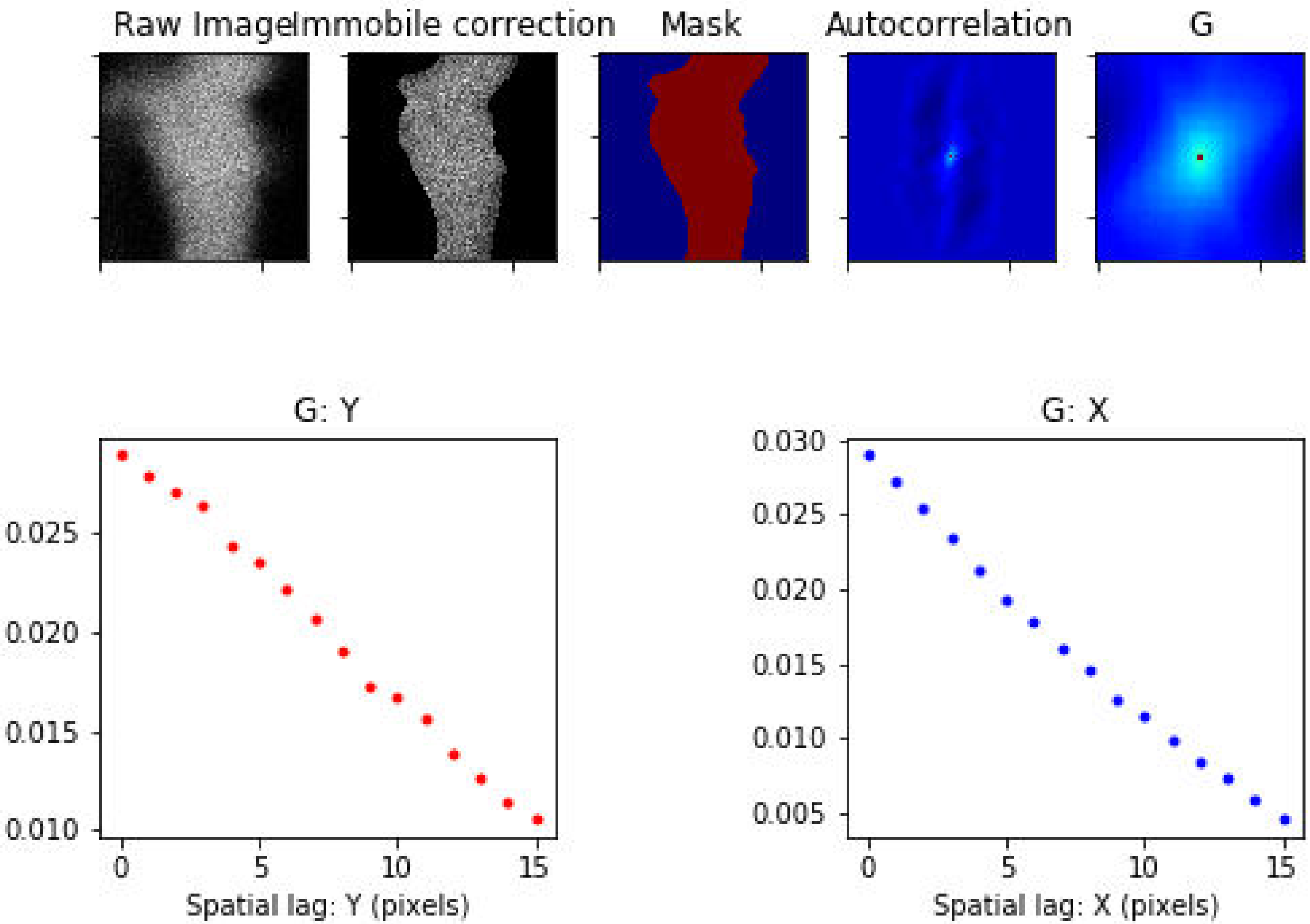

Treatment: TCA , Domain: MZ , File: Image 249.czi

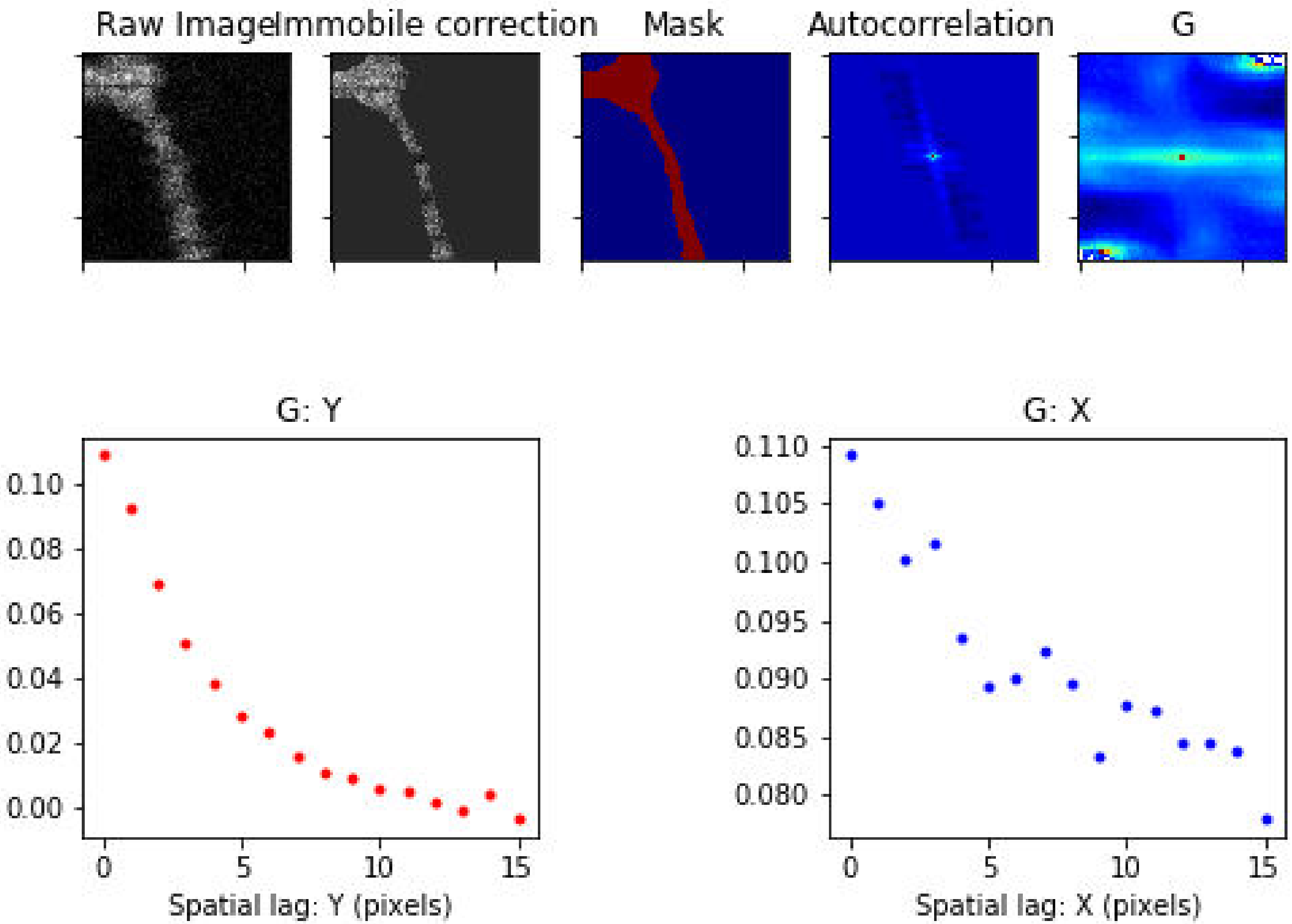

### Treatment: TCA , Domain: MZ , File: Image 255.czi

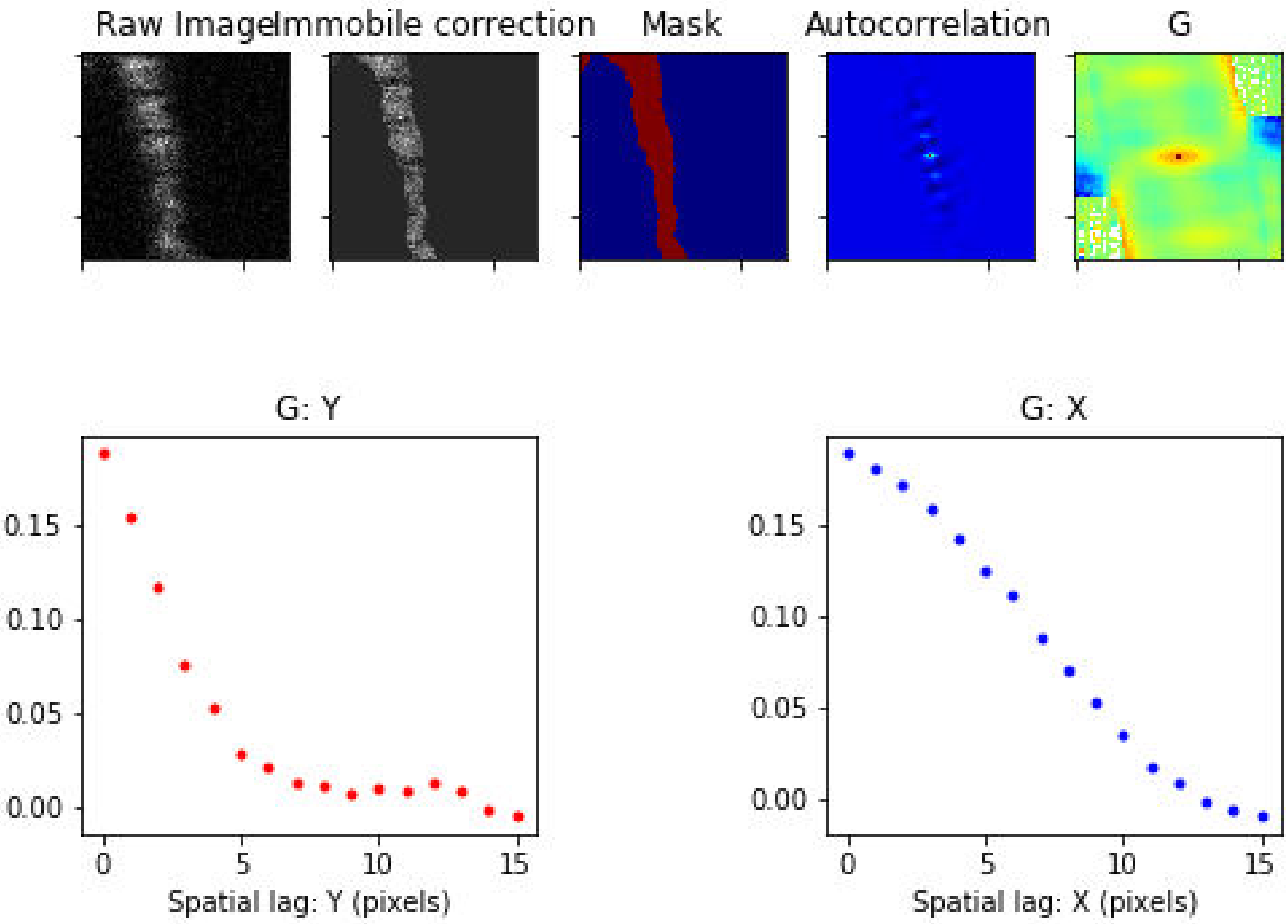

Treatment: TCA , Domain: MZ , File: Image 276.czi

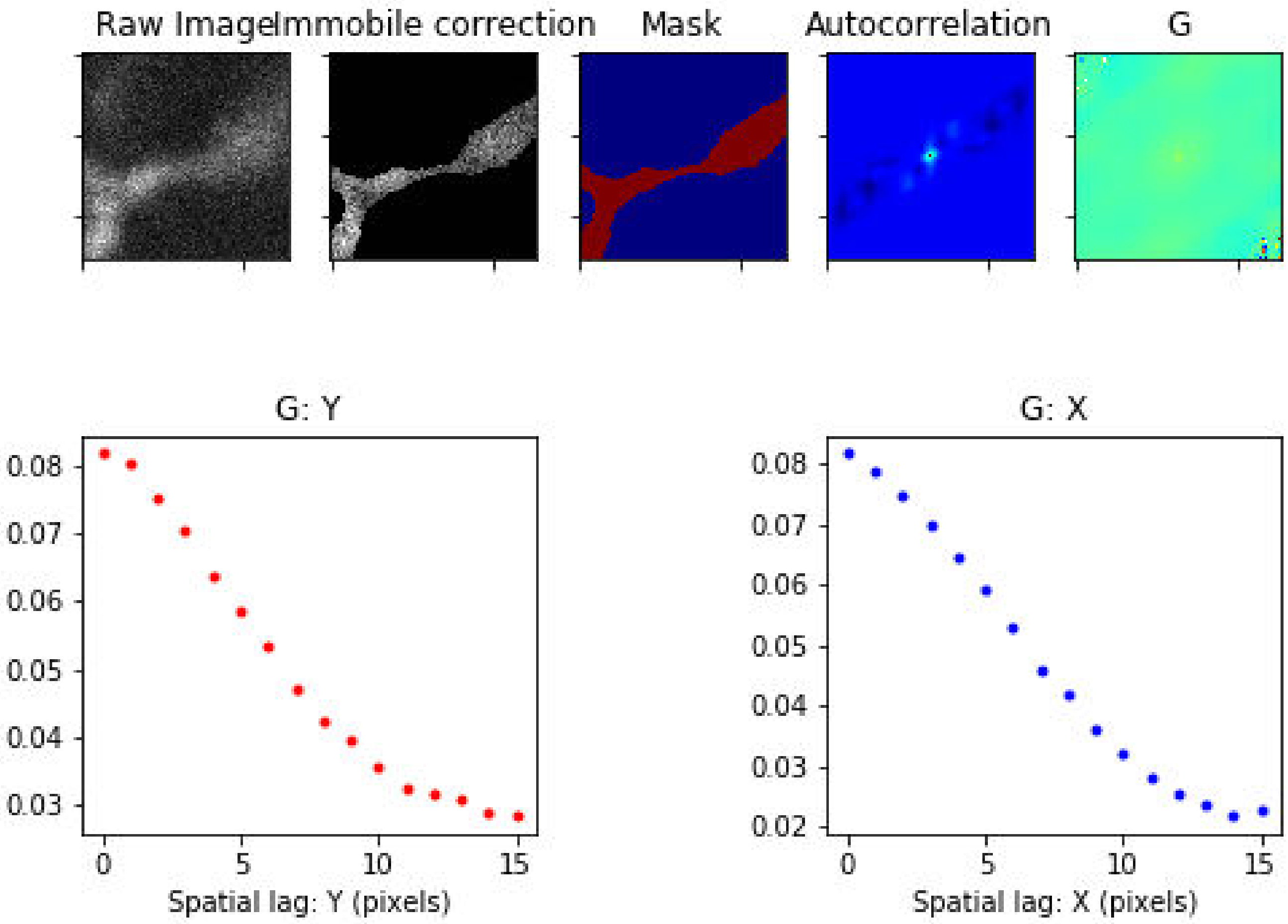

Treatment: TCA , Domain: MZ , File: Image 277.czi

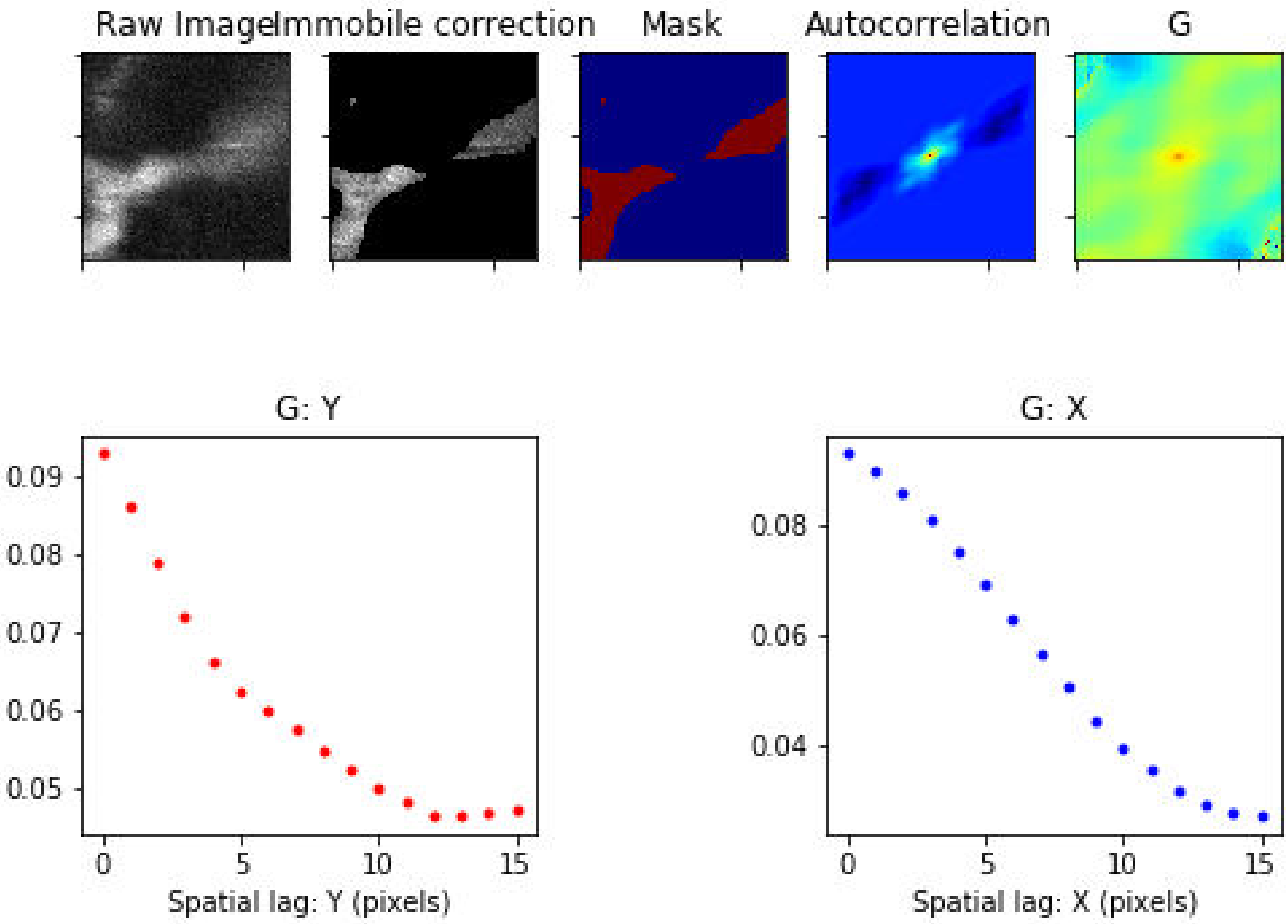

Treatment: TCA , Domain: MZ , File: Image 278.czi

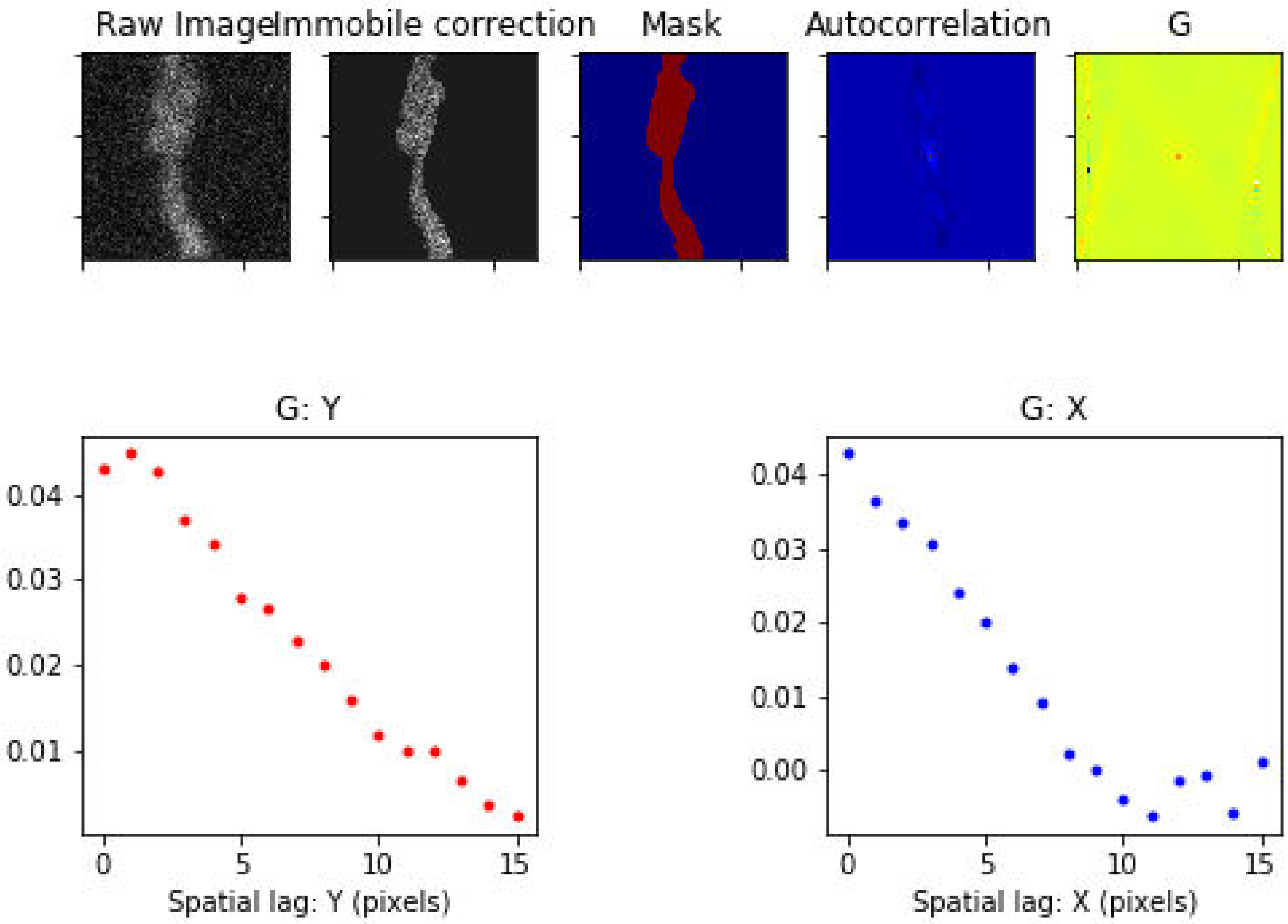

Treatment: TCA , Domain: MZ , File: Image 279.czi

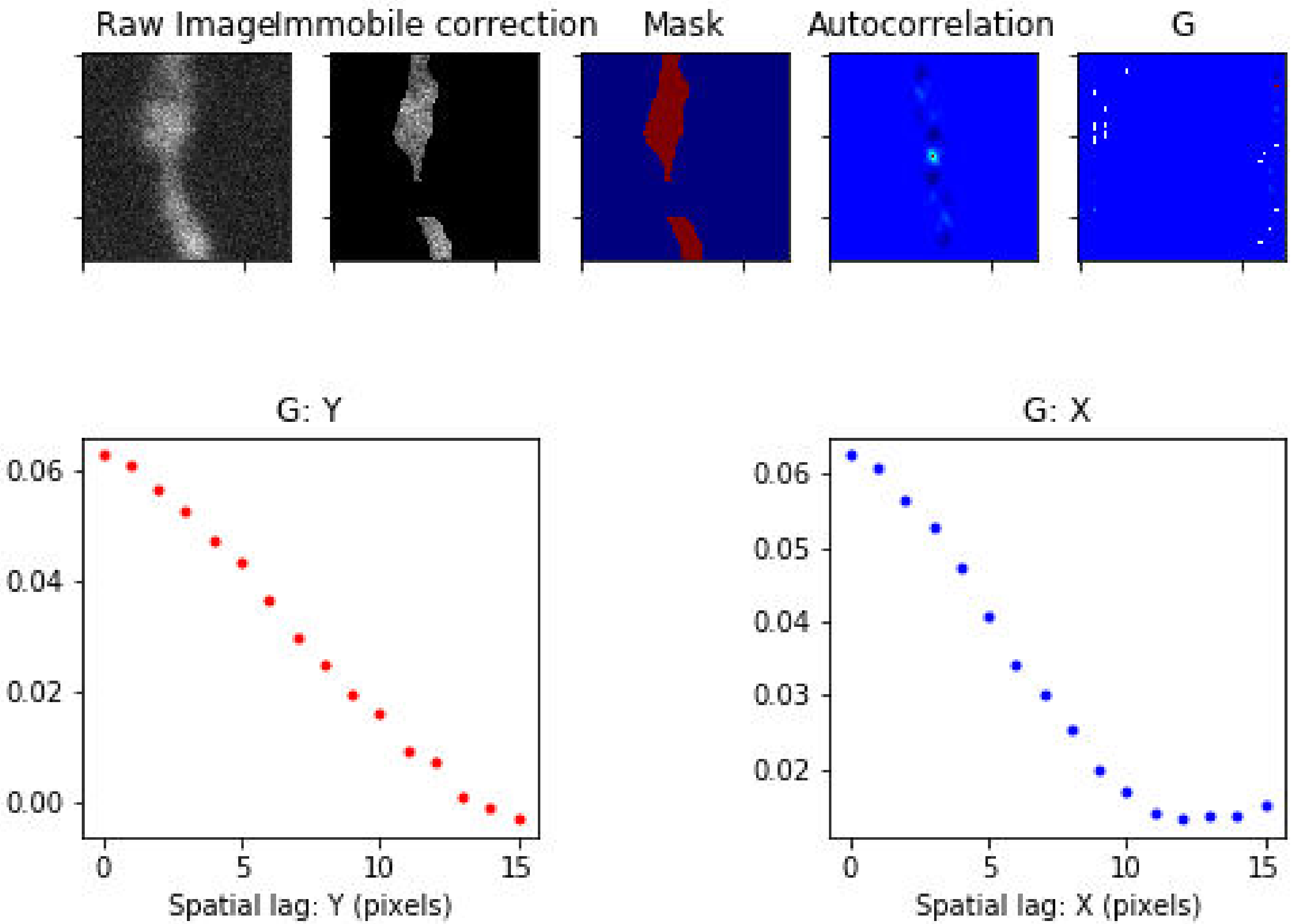

### Treatment: TCA , Domain: MZ , File: Image 280.czi

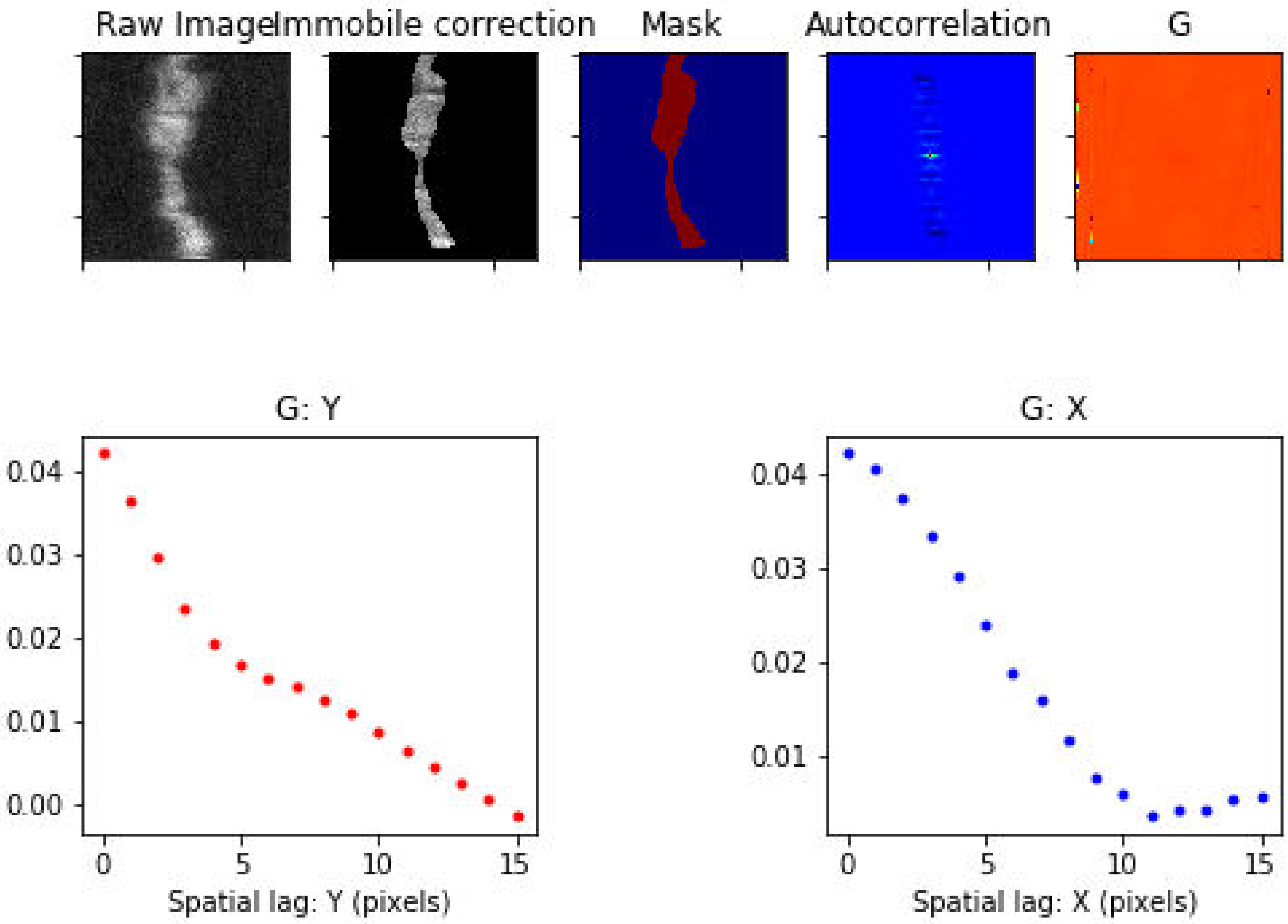

Treatment: TCA , Domain: MZ , File: Image 281.czi

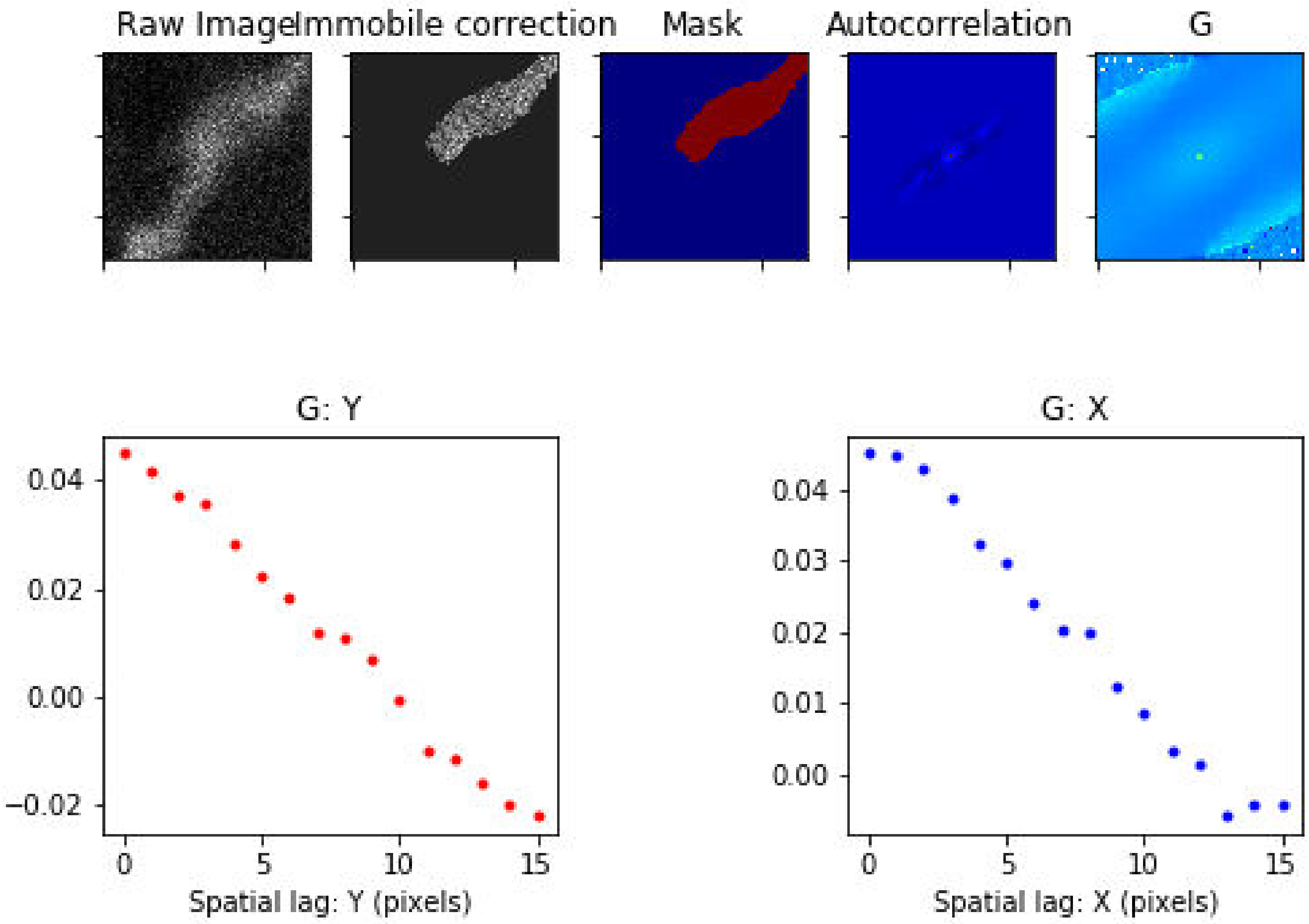

Treatment: TCA , Domain: MZ , File: Image 282.czi

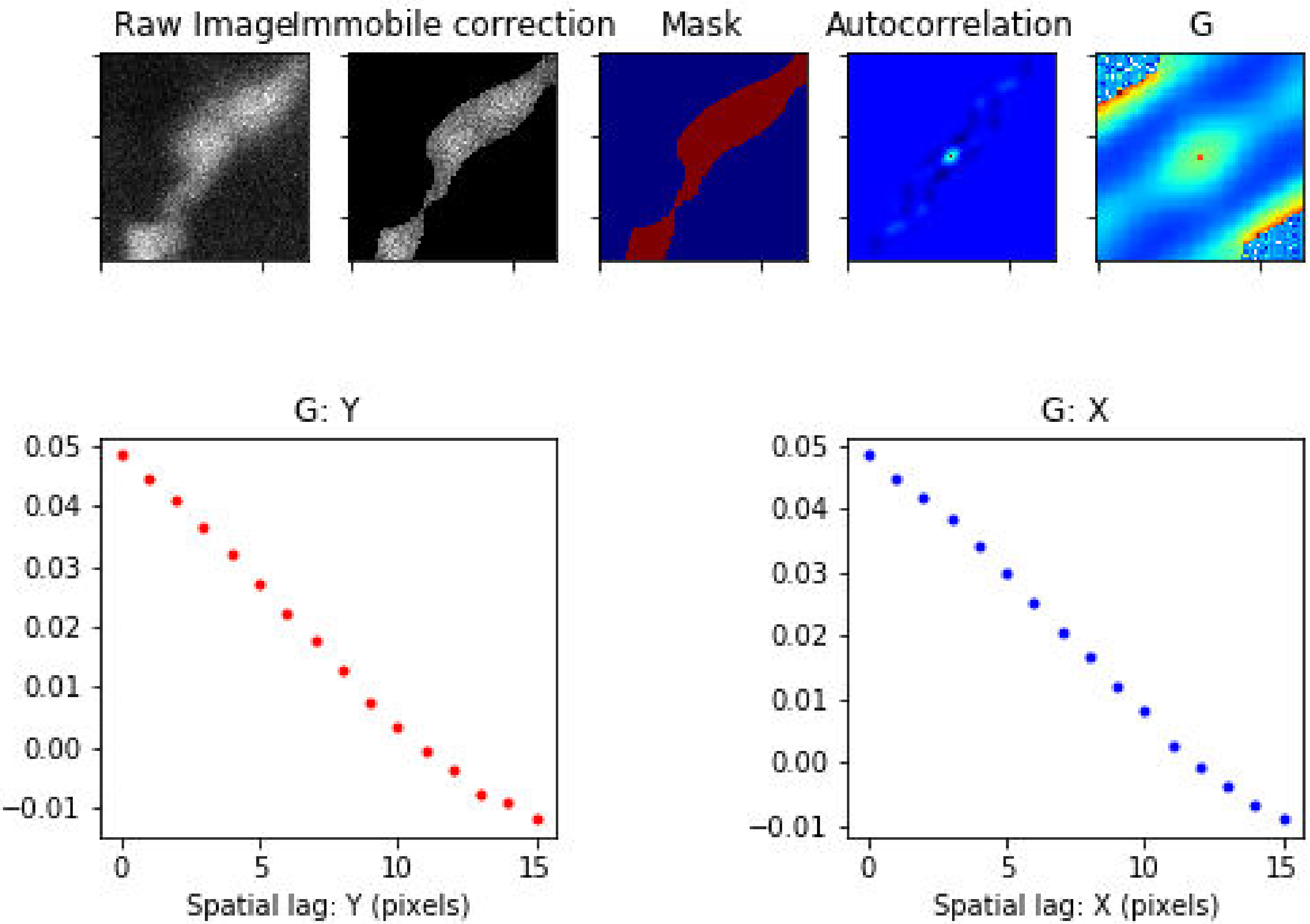

Treatment: TCA , Domain: MZ , File: Image 283.czi

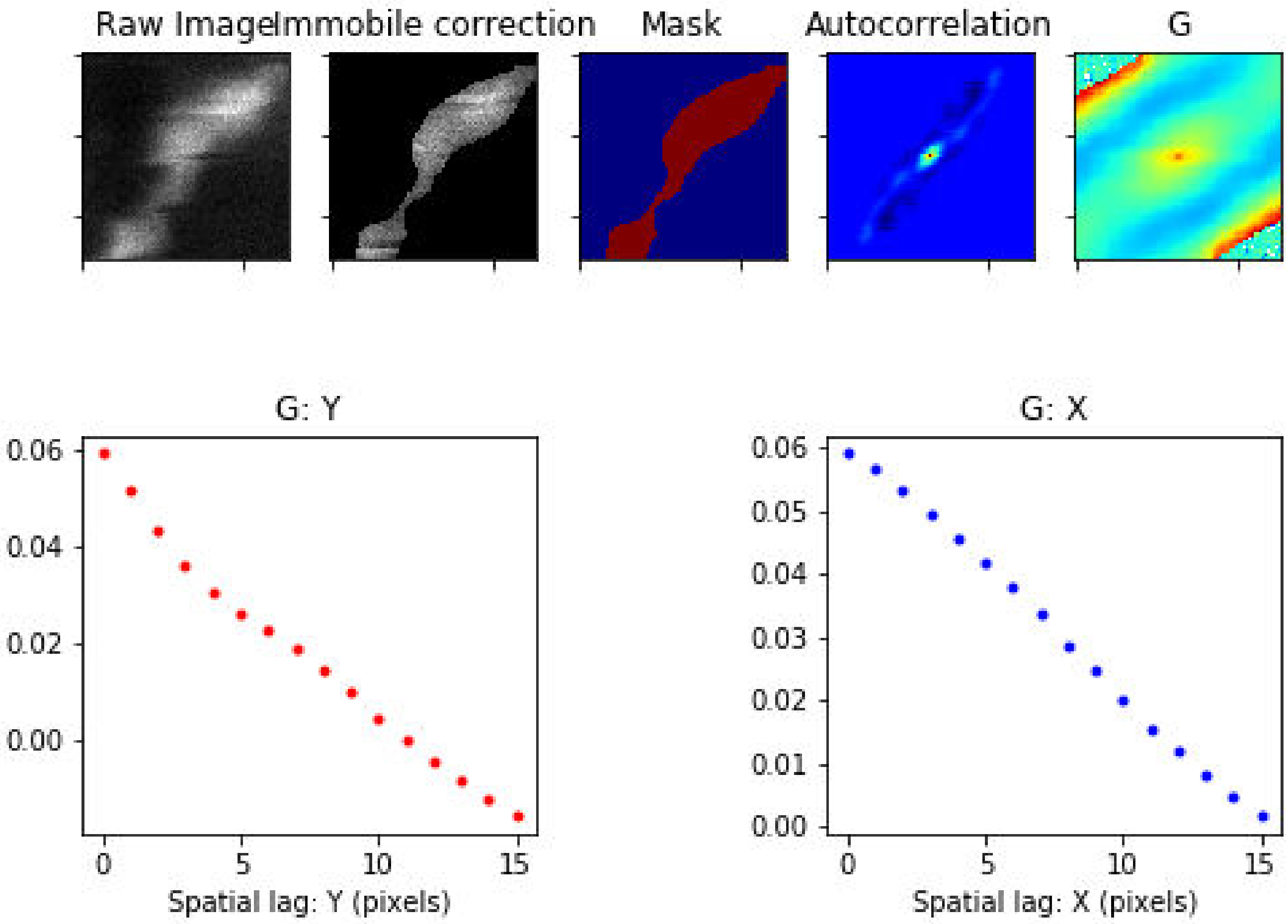

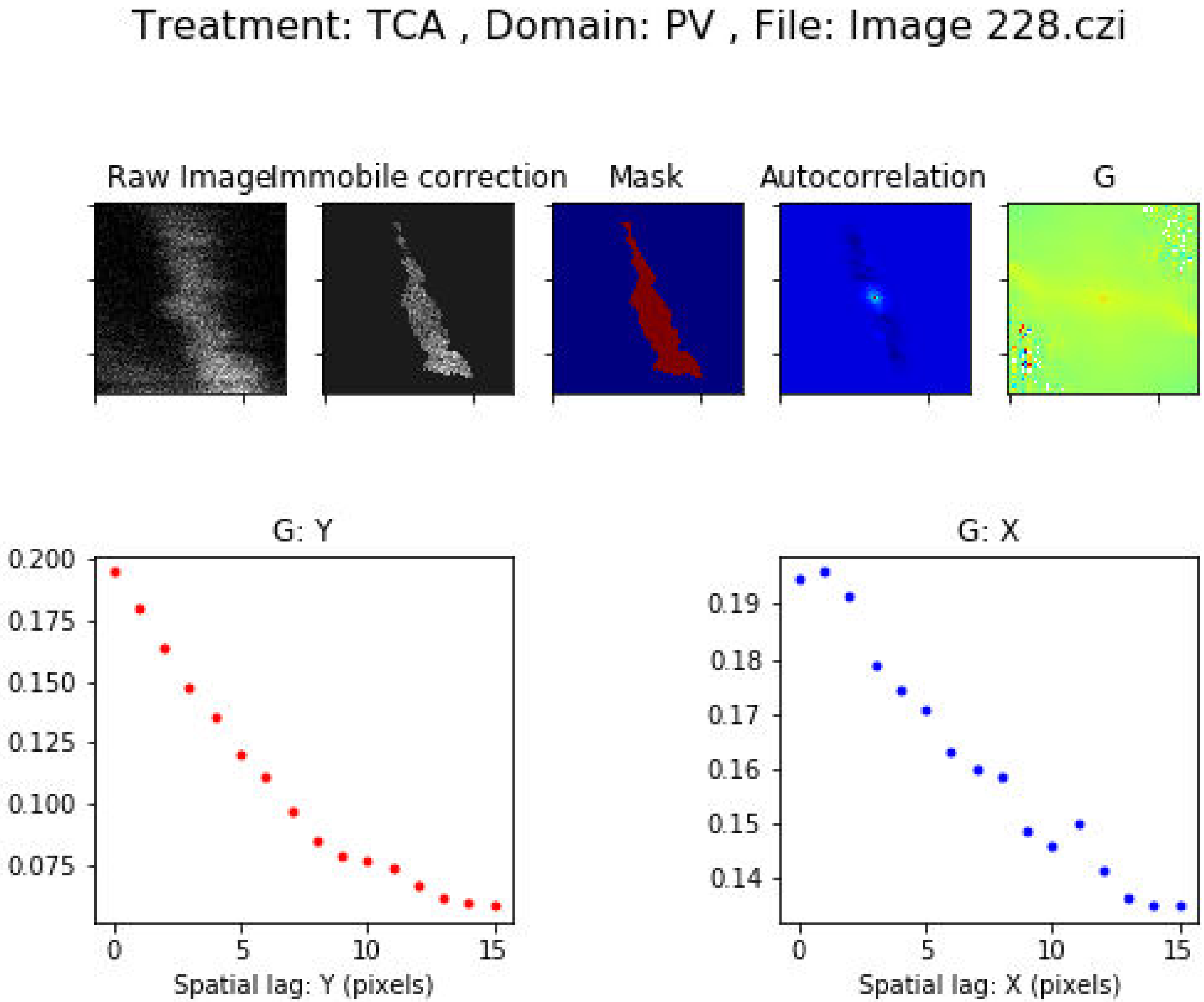
Treatment: TCA , Domain: PV , File: Image 228.czi
Raw Image
Immobile correction
Mask
Autocorrelation
G
G: Y
0.200
0.175
0.150
0.125
0.100
0.075
0
5
10
15
Spatial lag: Y (pixels)
G: X
0.19
0.18
0.17
0.16
0.15
0.14
0
5
10
15
Spatial lag: X (pixels)

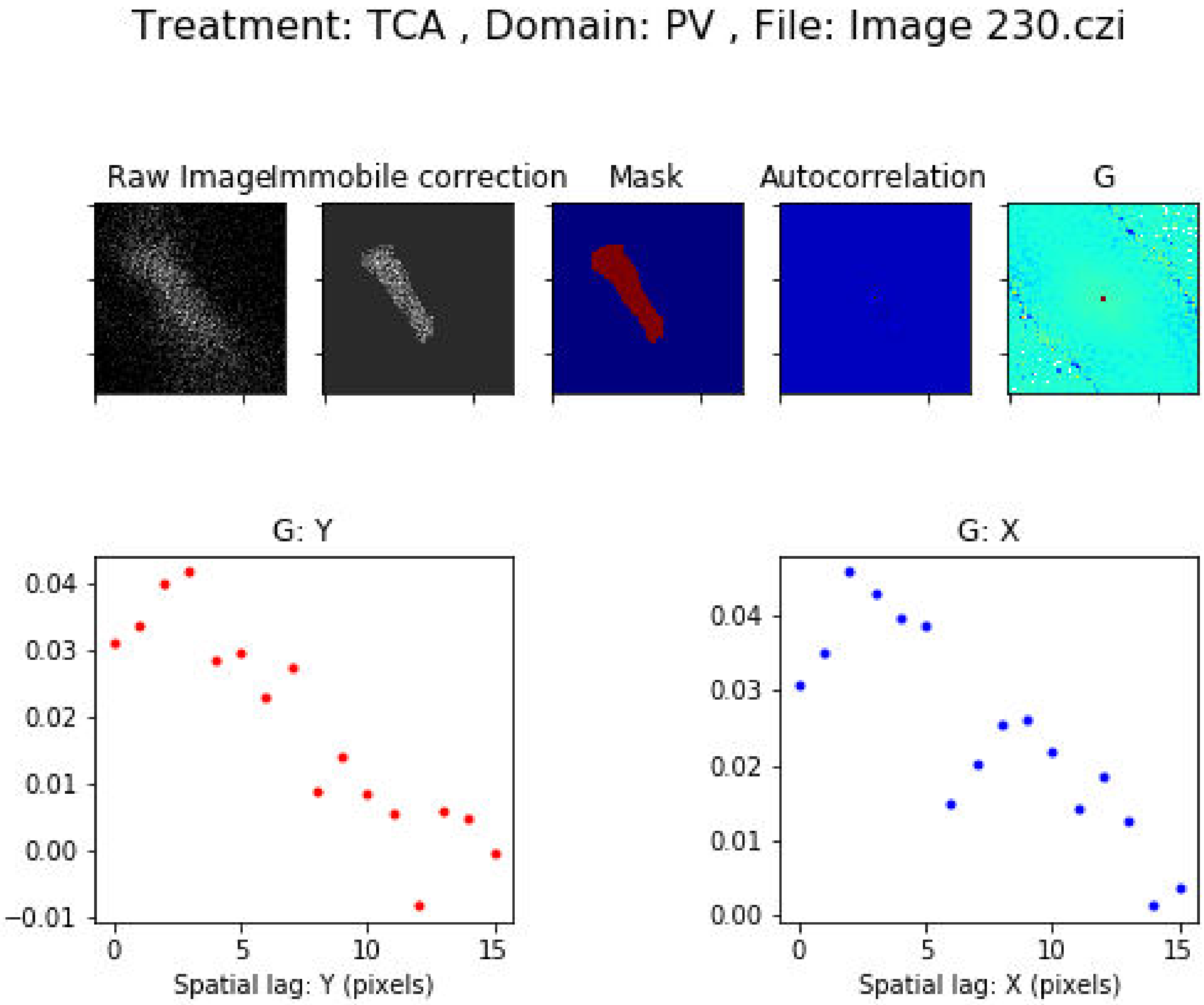
Treatment: TCA , Domain: PV , File: Image 230.czi
Raw Image
Immobile correction
Mask
Autocorrelation
G
G: Y
0.04
0.03
0.02
0.01
0.00
−0.01
0
5
10
15
Spatial lag: Y (pixels)
G: X
0.04
0.03
0.02
0.01
0.00
0
5
10
15
Spatial lag: X (pixels)

Treatment: TCA , Domain: PV , File: Image 263.czi

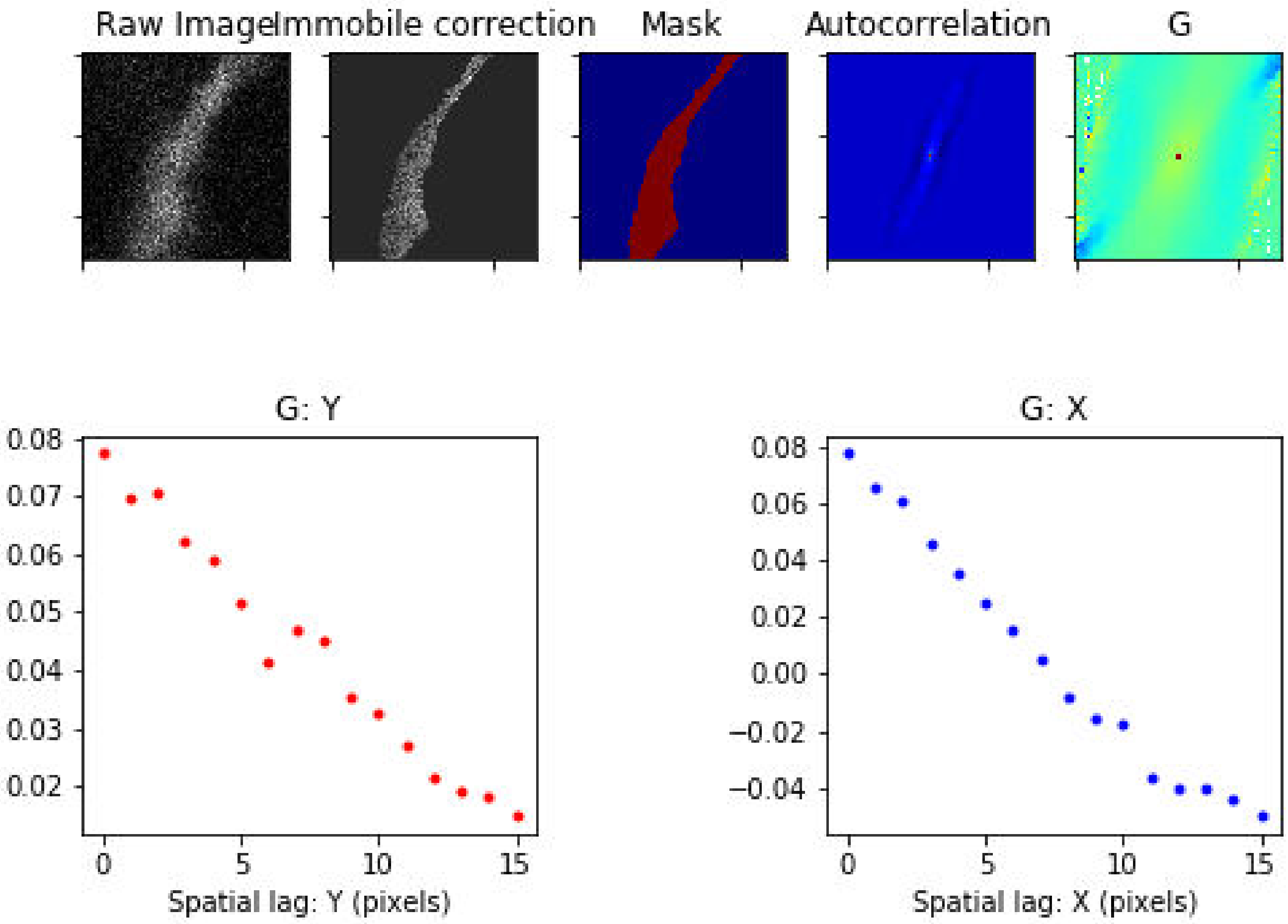

Treatment: TCA , Domain: PV , File: Image 264.czi

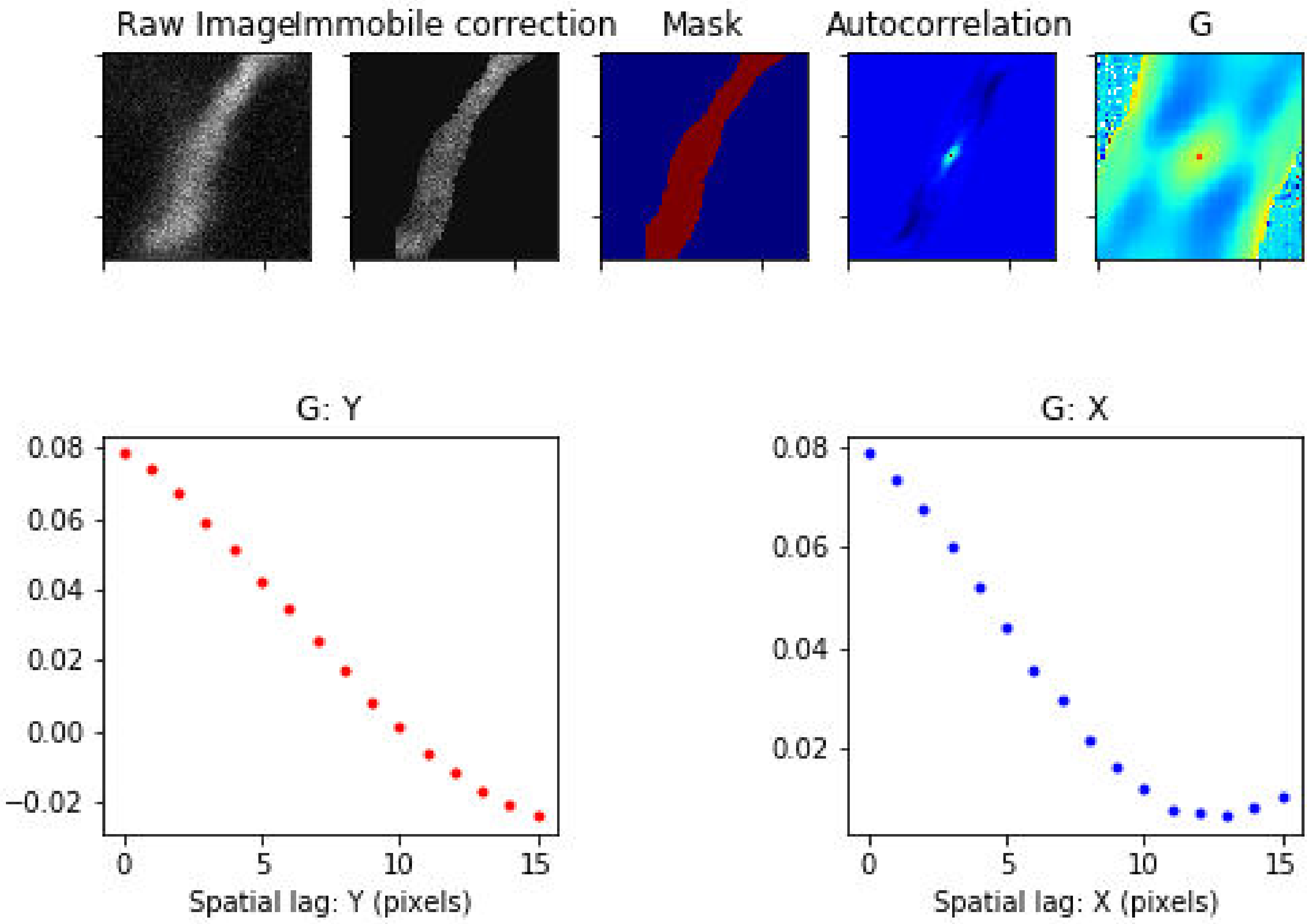

Treatment: TCA , Domain: PV , File: Image 265.czi

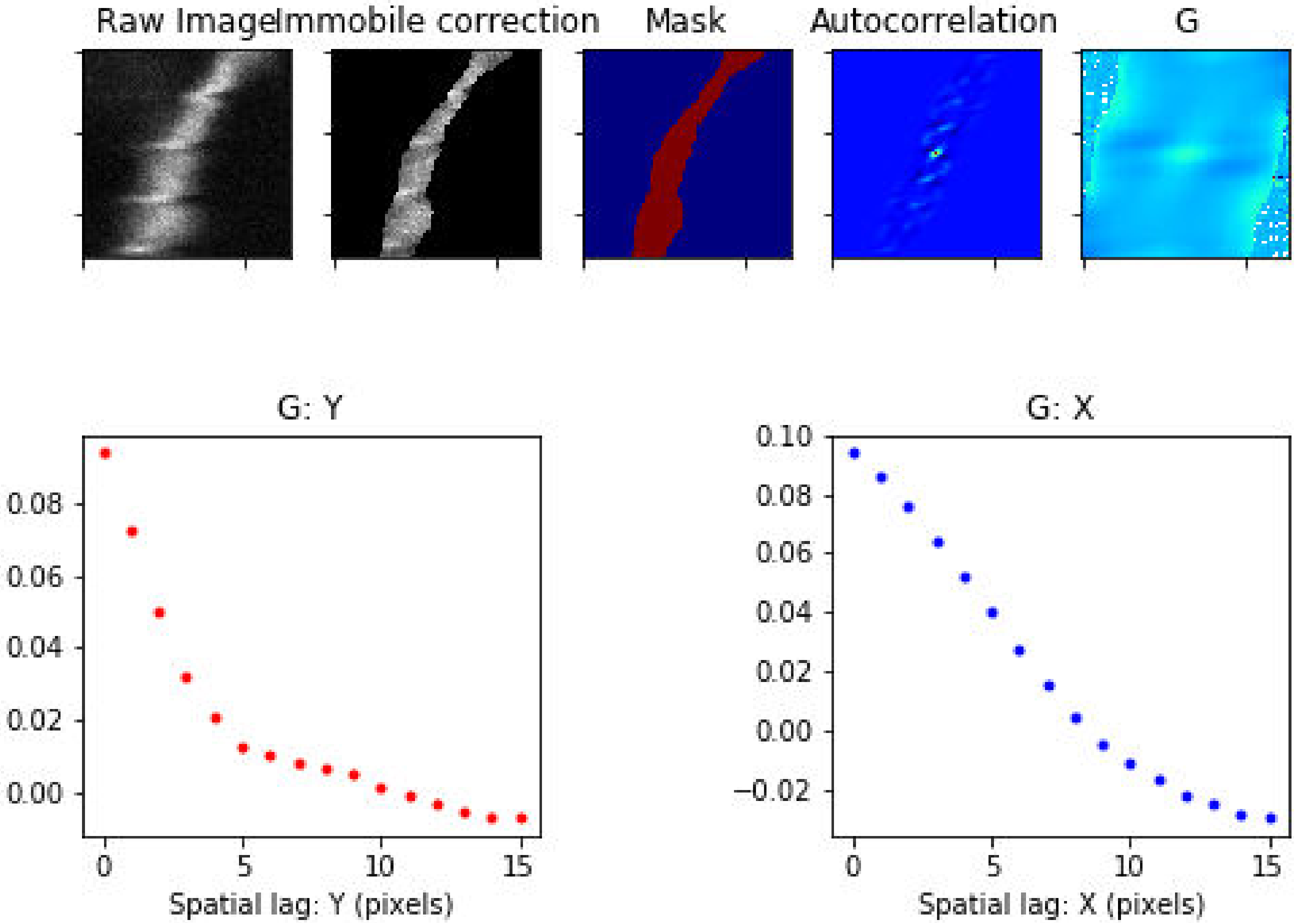

Treatment: TCA , Domain: PV , File: Image 266.czi

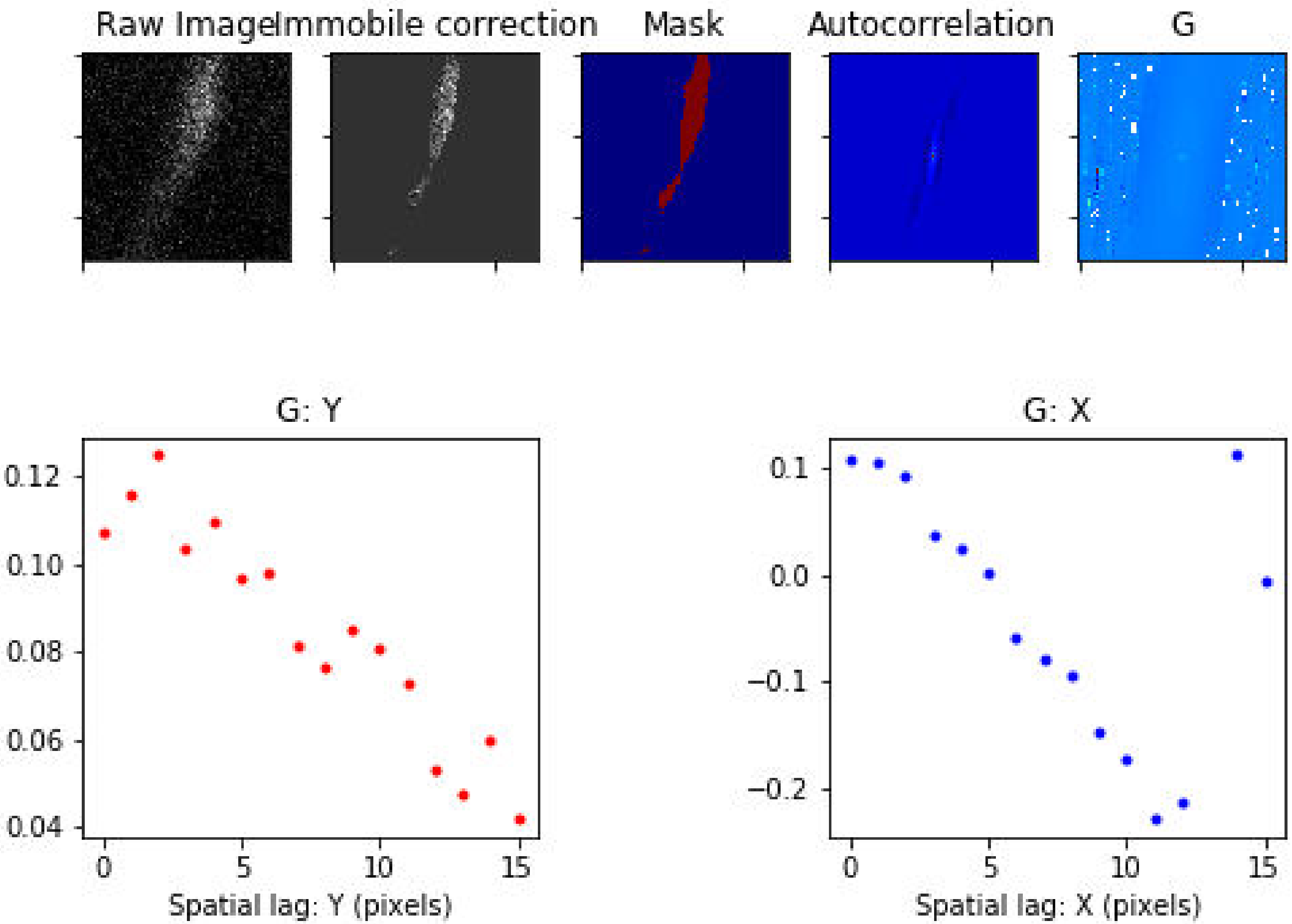

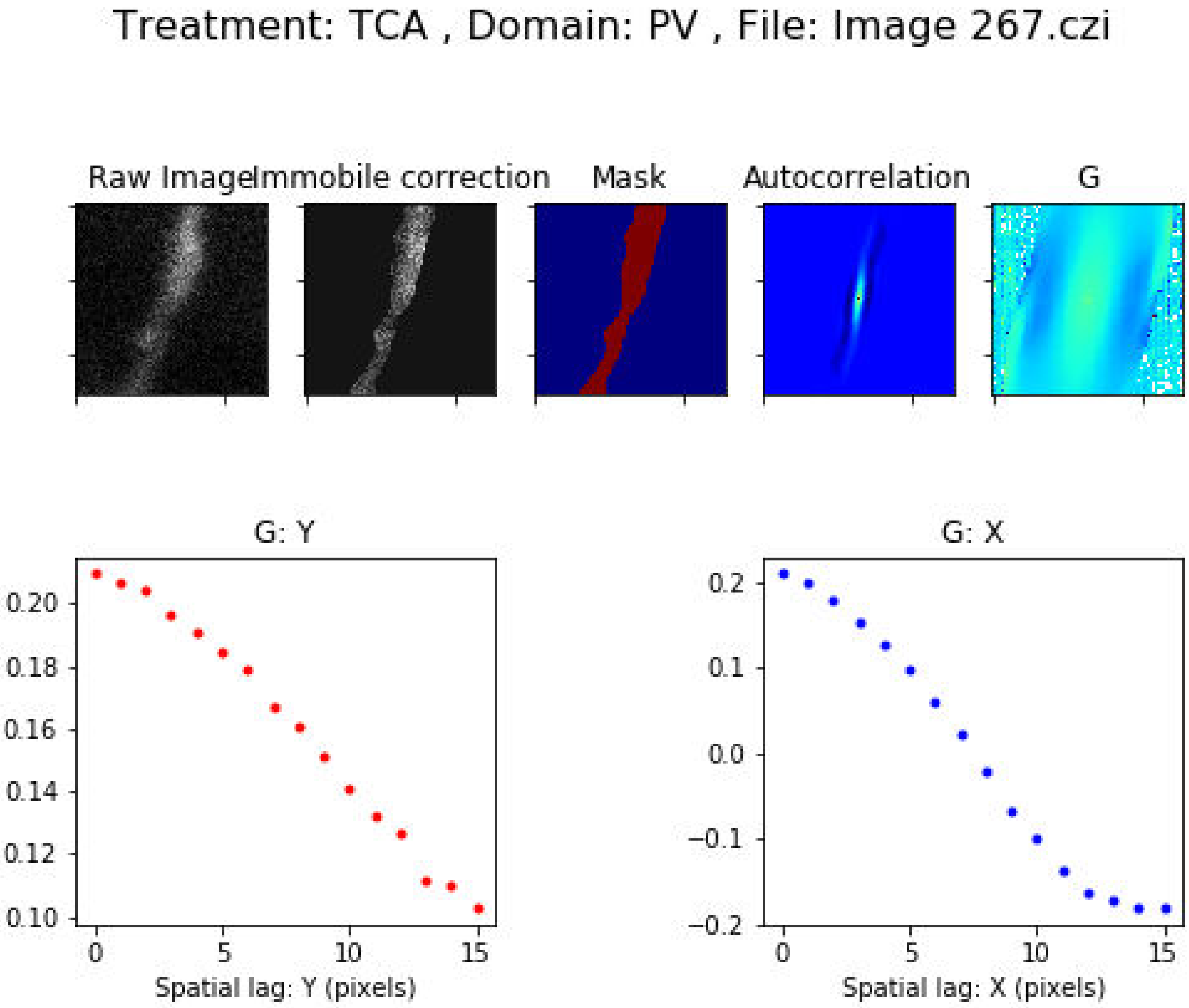
Treatment: TCA , Domain: PV , File: Image 267.czi
Raw Image
Immobile correction
Mask
Autocorrelation
G
G: Y
0.20
0.18
0.16
0.14
0.12
0.10
0
5
10
15
Spatial lag: Y (pixels)
G: X
0.2
0.1
0.0
−0.1
−0.2
0
5
10
15
Spatial lag: X (pixels)

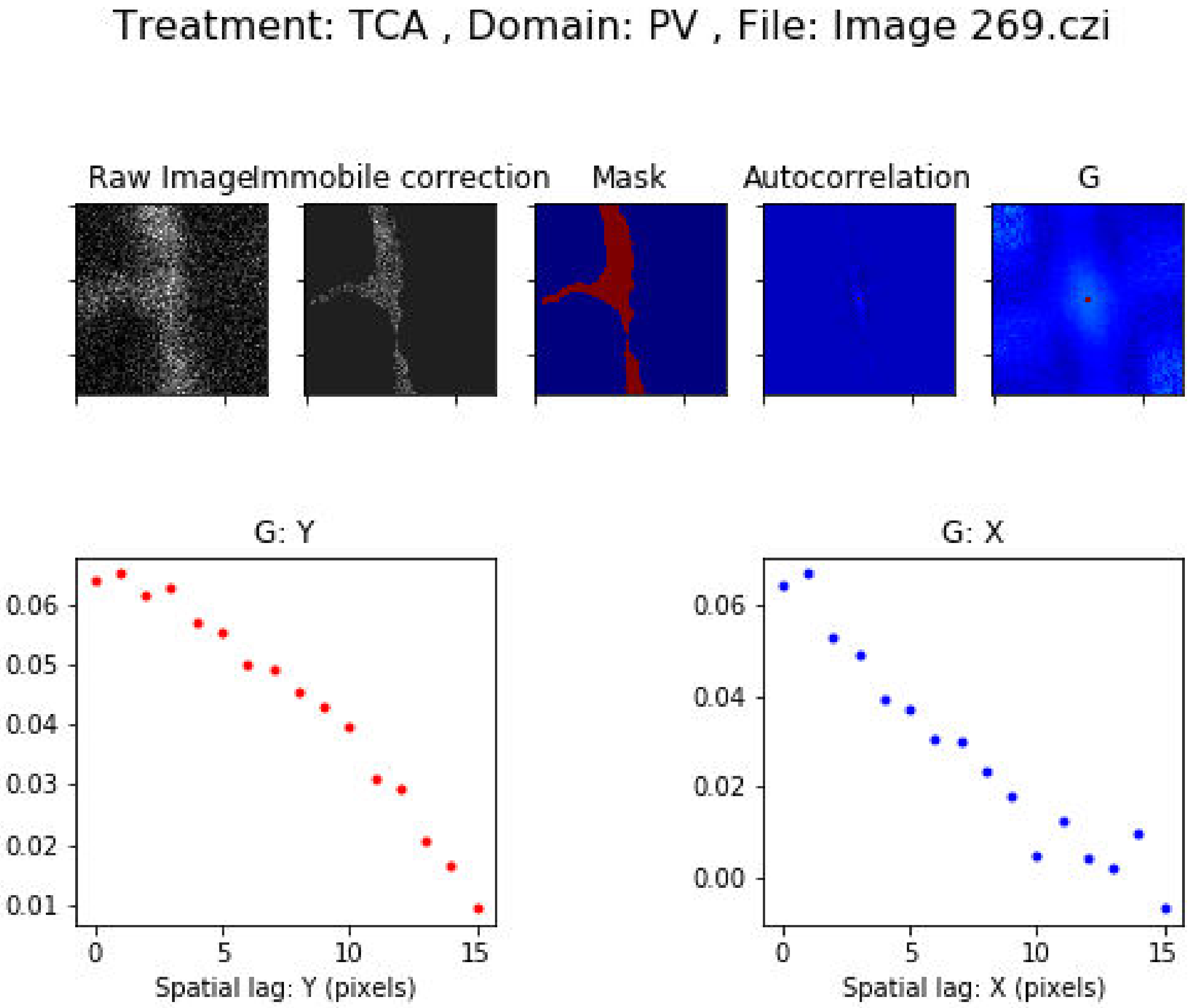

Treatment: TCA , Domain: PV , File: Image 269.czi
Raw Image
Immobile correction
Mask
Autocorrelation
G
G: Y
0.06
0.05
0.04
0.03
0.02
0.01
0
5
10
15
Spatial lag: Y (pixels)
G: X
0.06
0.04
0.02
0.00
0
5
10
15
Spatial lag: X (pixels)

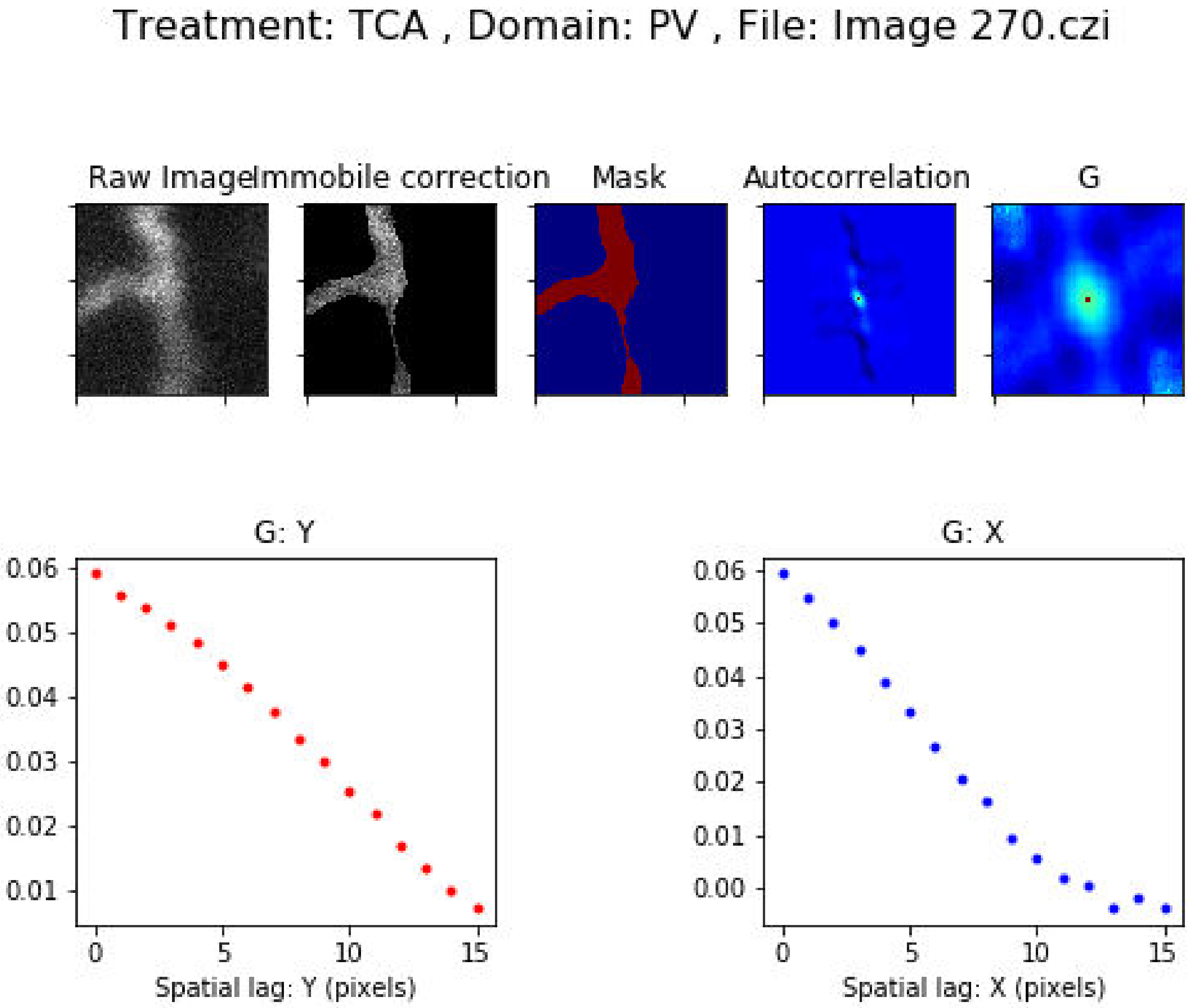
Treatment: TCA , Domain: PV , File: Image 270.czi
Raw Image
Immobile correction
Mask
Autocorrelation
G
G: Y
0.06
0.05
0.04
0.03
0.02
0.01
0
5
10
15
Spatial lag: Y (pixels)
G: X
0.06
0.05
0.04
0.03
0.02
0.01
0.00
0
5
10
15
Spatial lag: X (pixels)

Treatment: TCA , Domain: PV , File: Image 271.czi

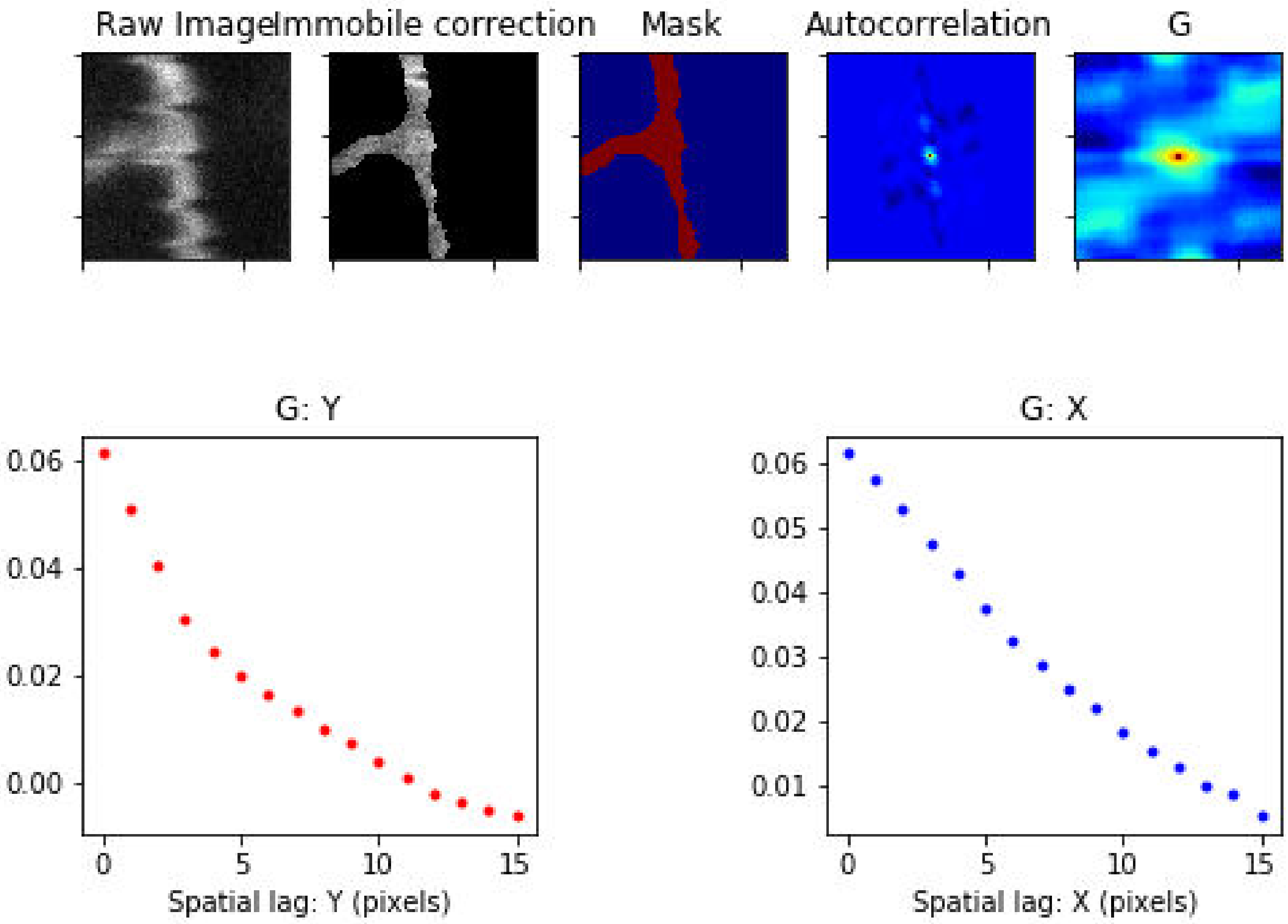

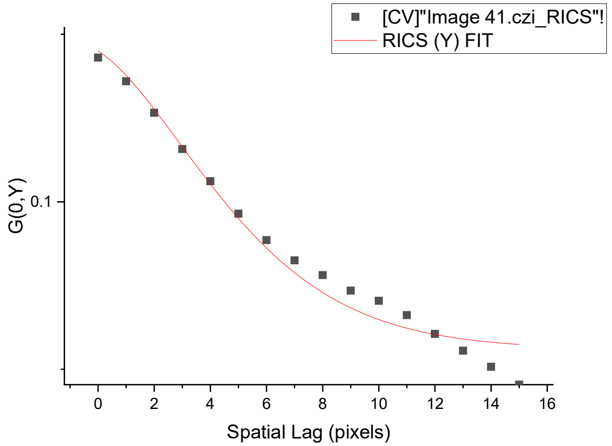
[CV]"Image 41.czi_RICS"!
RICS (Y) FIT
G(0,Y)
Spatial Lag (pixels)

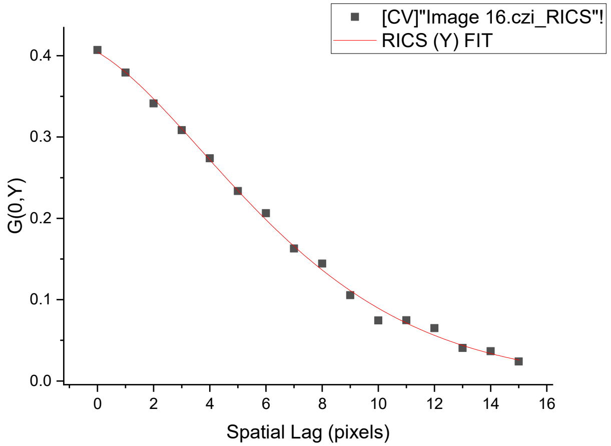
[CV]"Image 16.czi_RICS"!
RICS (Y) FIT
G(0,Y)
Spatial Lag (pixels)

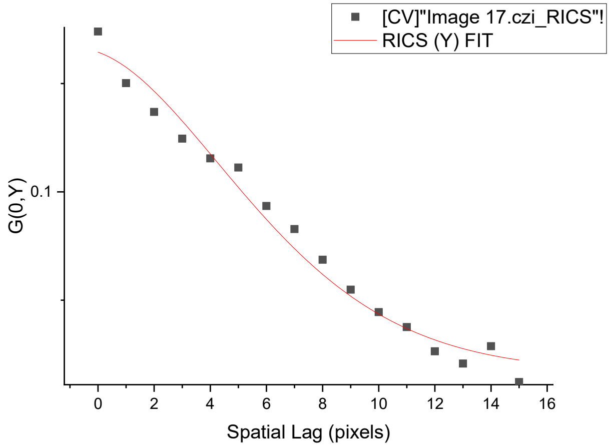
[CV]"Image 17.czi_RICS"!
RICS (Y) FIT
G(0,Y)
Spatial Lag (pixels)

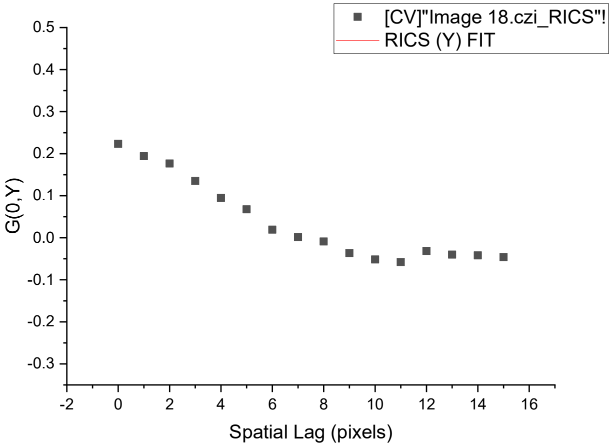
[CV]"Image 18.czi_RICS"!
RICS (Y) FIT
G(0,Y)
Spatial Lag (pixels)

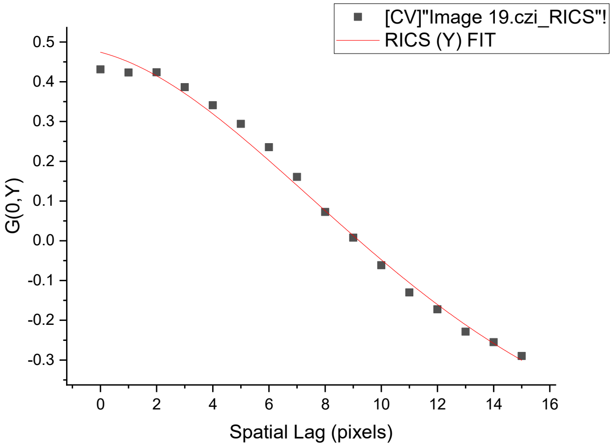
[CV]"Image 19.czi_RICS"!
RICS (Y) FIT
G(0,Y)
Spatial Lag (pixels)

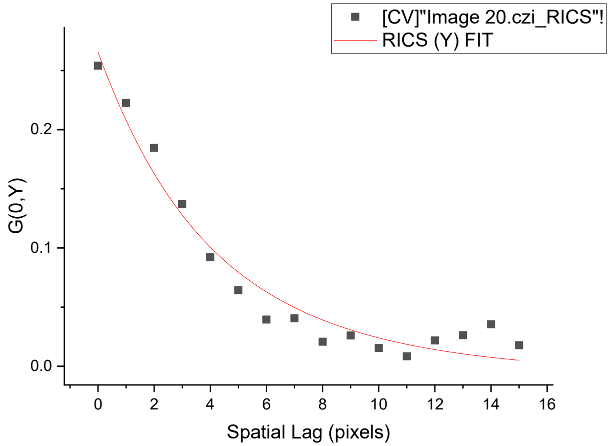
[CV]"Image 20.czi_RICS"!
RICS (Y) FIT
G(0,Y)
Spatial Lag (pixels)

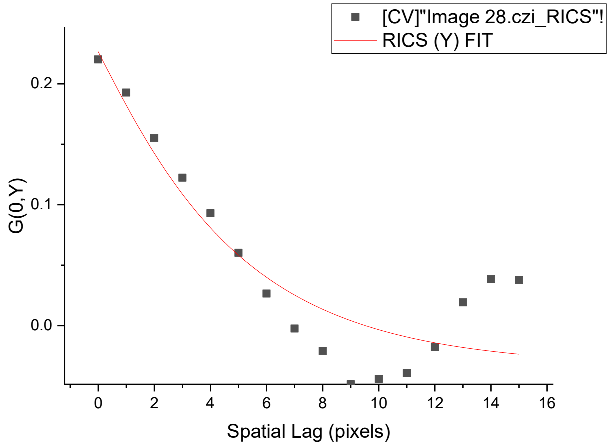
[CV]"Image 28.czi_RICS"!
RICS (Y) FIT
G(0,Y)
Spatial Lag (pixels)

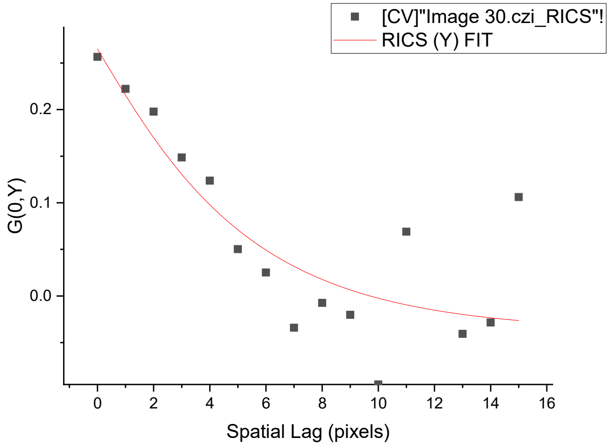
[CV]"Image 30.czi_RICS"!
RICS (Y) FIT
G(0,Y)
Spatial Lag (pixels)

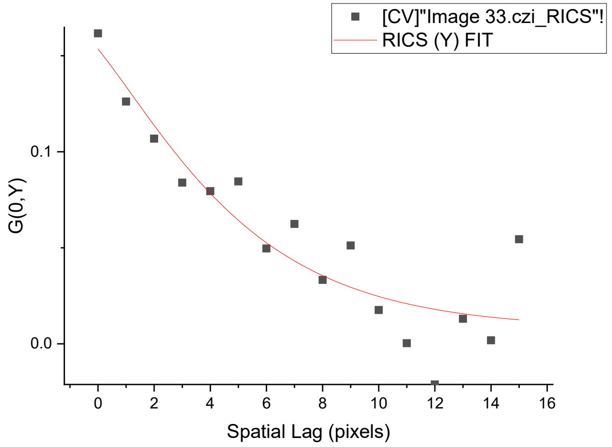
[CV]"Image 33.czi_RICS"!
RICS (Y) FIT
G(0,Y)
Spatial Lag (pixels)

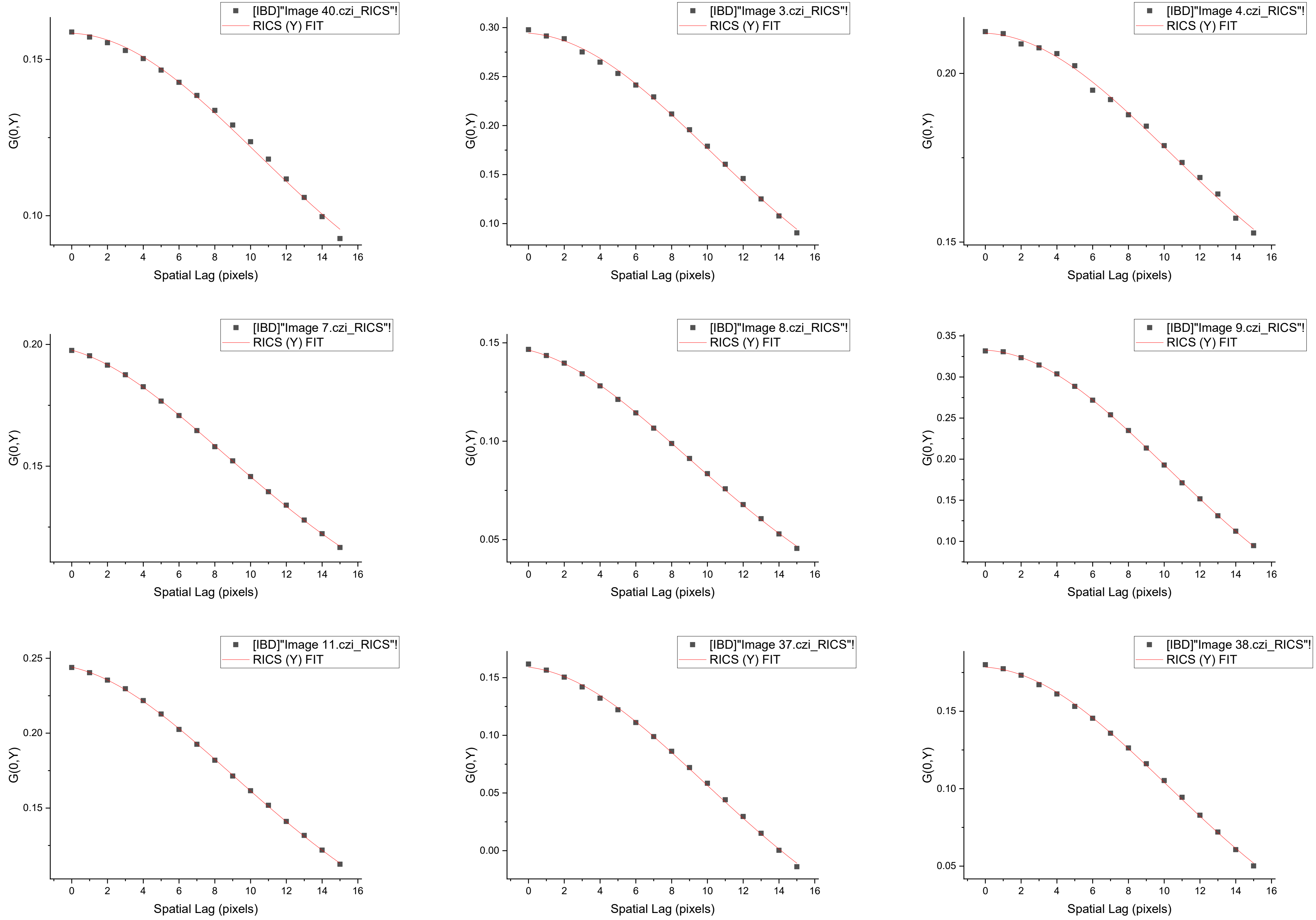
[IBD]"Image 40.czi_RICS"!
RICS (Y) FIT
G(0,Y)
Spatial Lag (pixels)
[IBD]"Image 3.czi_RICS"!
RICS (Y) FIT
G(0,Y)
Spatial Lag (pixels)
[IBD]"Image 4.czi_RICS"!
RICS (Y) FIT
G(0,Y)
Spatial Lag (pixels)
[IBD]"Image 7.czi_RICS"!
RICS (Y) FIT
G(0,Y)
Spatial Lag (pixels)
[IBD]"Image 8.czi_RICS"!
RICS (Y) FIT
G(0,Y)
Spatial Lag (pixels)
[IBD]"Image 9.czi_RICS"!
RICS (Y) FIT
G(0,Y)
Spatial Lag (pixels)
[IBD]"Image 11.czi_RICS"!
RICS (Y) FIT
G(0,Y)
Spatial Lag (pixels)
[IBD]"Image 37.czi_RICS"!
RICS (Y) FIT
G(0,Y)
Spatial Lag (pixels)
[IBD]"Image 38.czi_RICS"!
RICS (Y) FIT
G(0,Y)
Spatial Lag (pixels)

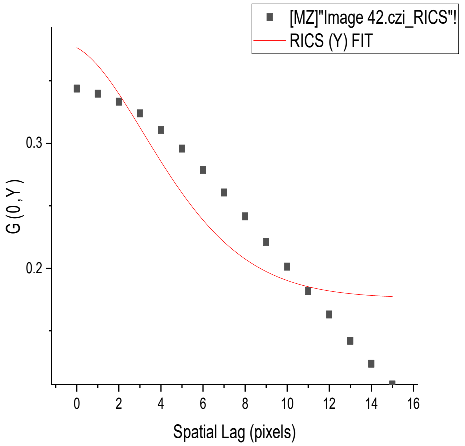
[MZ]"Image 42.czi_RICS"!
RICS (Y) FIT
G (0, Y)
0.3
0.2
0 2 4 6 8 10 12 14 16
Spatial Lag (pixels)

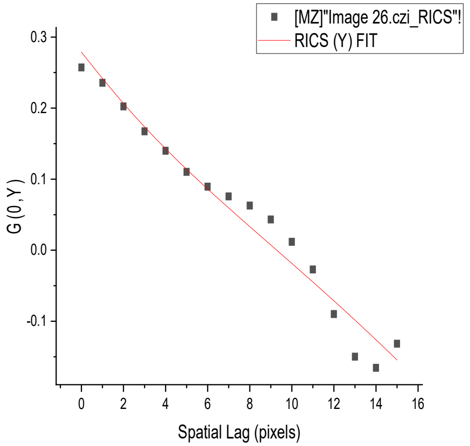
[MZ]"Image 26.czi_RICS"!
RICS (Y) FIT
G (0, Y)
0.3
0.2
0.1
0.0
-0.1
0 2 4 6 8 10 12 14 16
Spatial Lag (pixels)

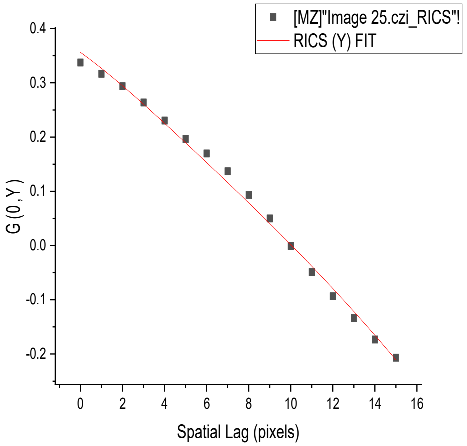
[MZ]"Image 25.czi_RICS"!
RICS (Y) FIT
G (0, Y)
0.4
0.3
0.2
0.1
0.0
-0.1
-0.2
0 2 4 6 8 10 12 14 16
Spatial Lag (pixels)

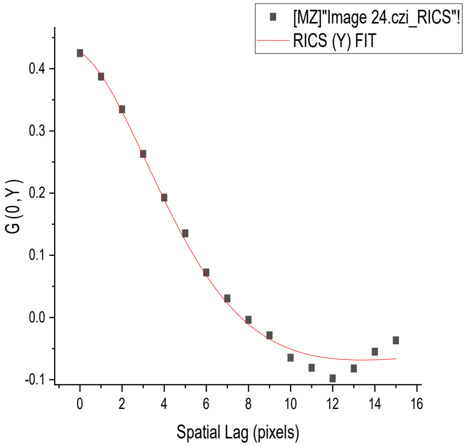
[MZ]"Image 24.czi_RICS"!
RICS (Y) FIT
G (0, Y)
0.4
0.3
0.2
0.1
0.0
-0.1
0 2 4 6 8 10 12 14 16
Spatial Lag (pixels)

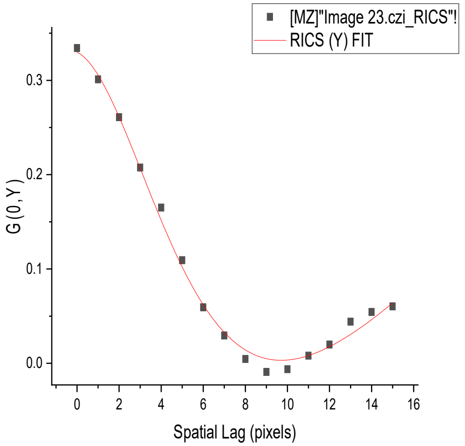
[MZ]"Image 23.czi_RICS"!
RICS (Y) FIT
G (0, Y)
0.3
0.2
0.1
0.0
0 2 4 6 8 10 12 14 16
Spatial Lag (pixels)

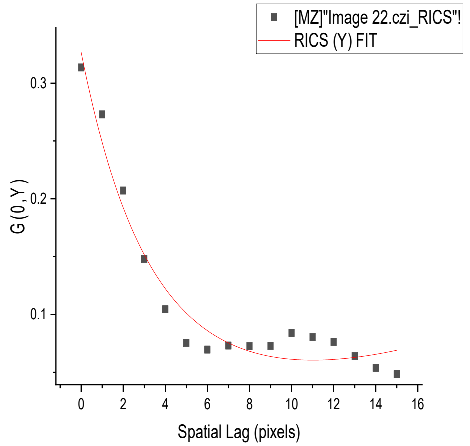
[MZ]"Image 22.czi_RICS"!
RICS (Y) FIT
G (0, Y)
0.3
0.2
0.1
0 2 4 6 8 10 12 14 16
Spatial Lag (pixels)

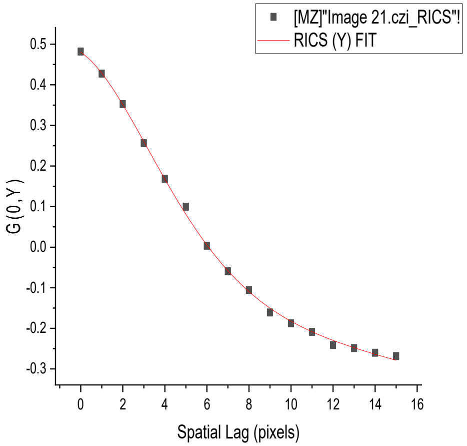
[MZ]"Image 21.czi_RICS"!
RICS (Y) FIT
G (0, Y)
0.5
0.4
0.3
0.2
0.1
0.0
-0.1
-0.2
-0.3
0 2 4 6 8 10 12 14 16
Spatial Lag (pixels)

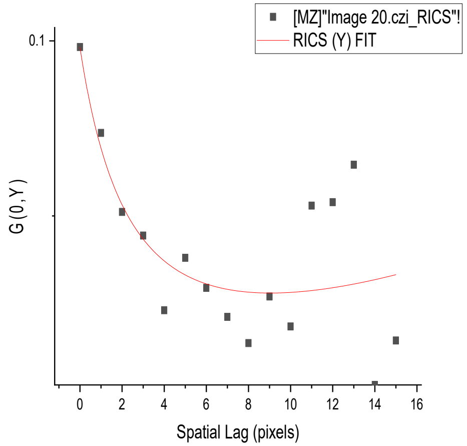
[MZ]"Image 20.czi_RICS"!
RICS (Y) FIT
G (0, Y)
0.1
0 2 4 6 8 10 12 14 16
Spatial Lag (pixels)

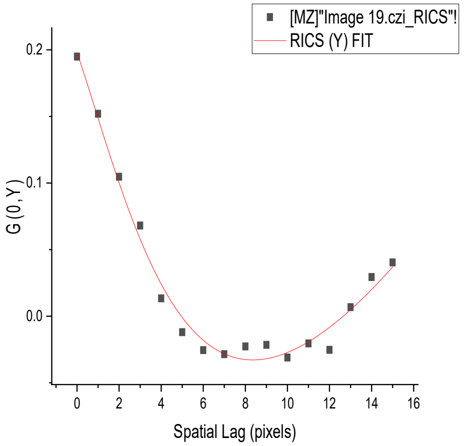
[MZ]"Image 19.czi_RICS"!
RICS (Y) FIT
G (0, Y)
0.2
0.1
0.0
0 2 4 6 8 10 12 14 16
Spatial Lag (pixels)

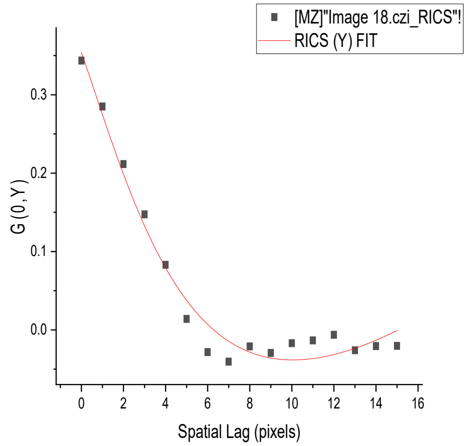
[MZ]"Image 18.czi_RICS"!
RICS (Y) FIT
G (0, Y)
0.3
0.2
0.1
0.0
0 2 4 6 8 10 12 14 16
Spatial Lag (pixels)

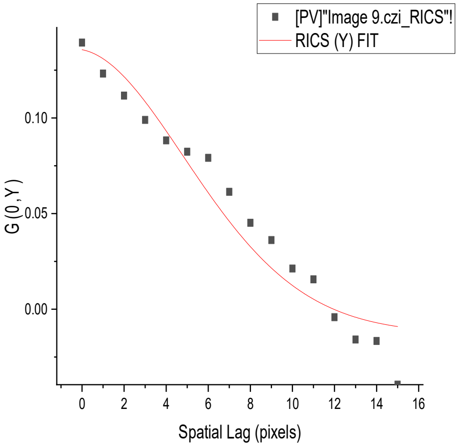
[PV]"Image 9.czi_RICS"!
RICS (Y) FIT
G (0, Y)
Spatial Lag (pixels)

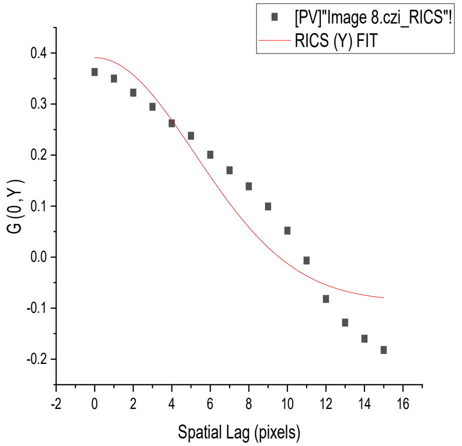
[PV]"Image 8.czi_RICS"!
RICS (Y) FIT
G (0, Y)
Spatial Lag (pixels)

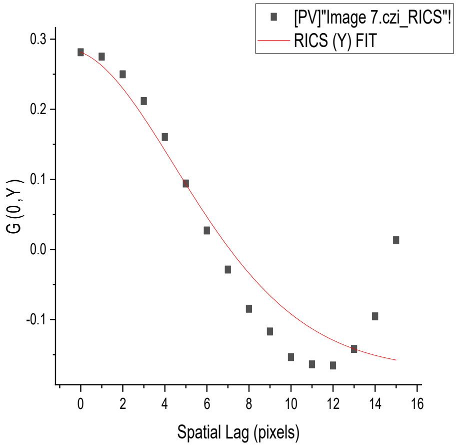
[PV]"Image 7.czi_RICS"!
RICS (Y) FIT
G (0, Y)
Spatial Lag (pixels)

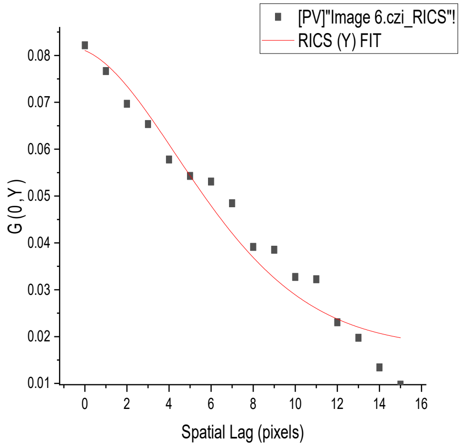
[PV]"Image 6.czi_RICS"!
RICS (Y) FIT
G (0, Y)
Spatial Lag (pixels)

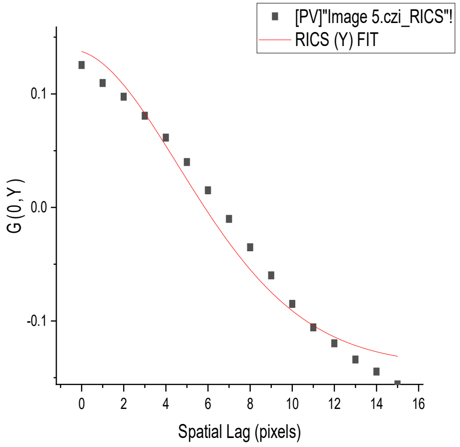
[PV]"Image 5.czi_RICS"!
RICS (Y) FIT
G (0, Y)
Spatial Lag (pixels)

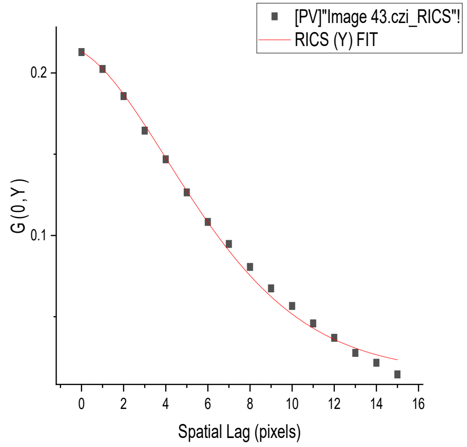
[PV]"Image 43.czi_RICS"!
RICS (Y) FIT
G (0, Y)
Spatial Lag (pixels)

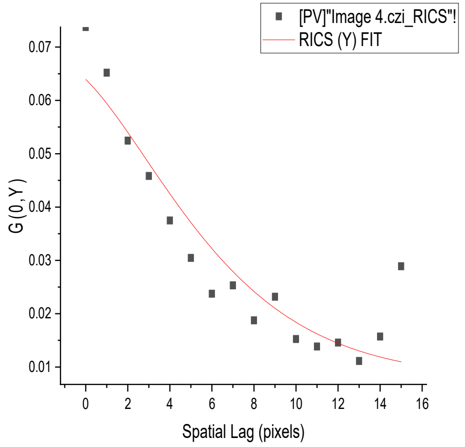
[PV]"Image 4.czi_RICS"!
RICS (Y) FIT
G (0, Y)
Spatial Lag (pixels)

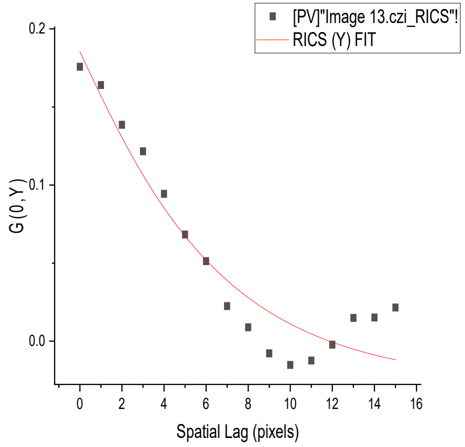
[PV]"Image 13.czi_RICS"!
RICS (Y) FIT
G (0, Y)
Spatial Lag (pixels)

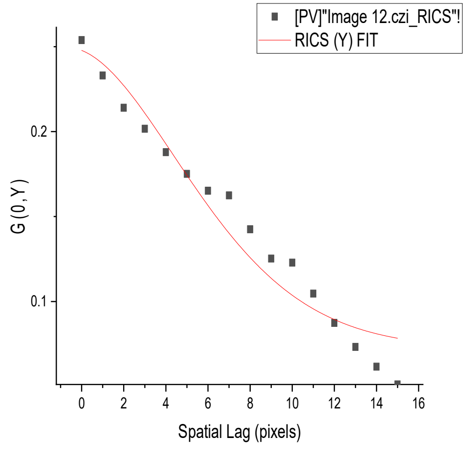
[PV]"Image 12.czi_RICS"!
RICS (Y) FIT
G (0, Y)
Spatial Lag (pixels)

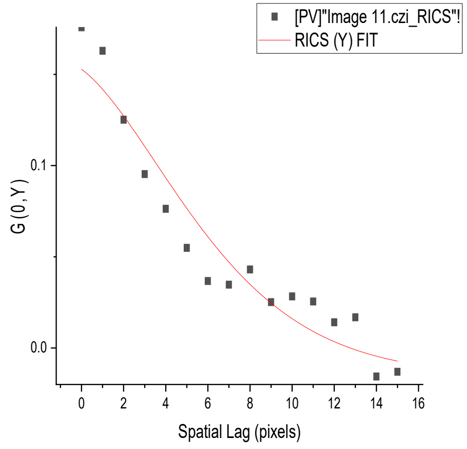
[PV]"Image 11.czi_RICS"!
RICS (Y) FIT
G (0, Y)
Spatial Lag (pixels)

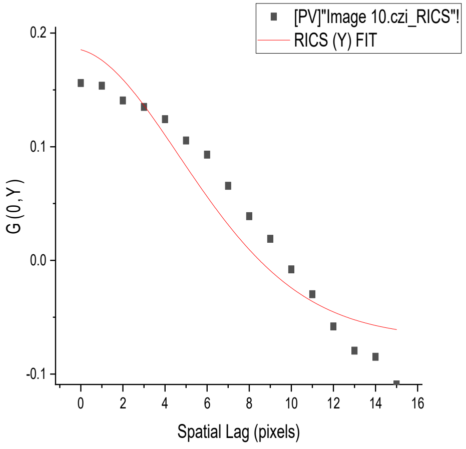
[PV]"Image 10.czi_RICS"!
RICS (Y) FIT
G (0, Y)
Spatial Lag (pixels)

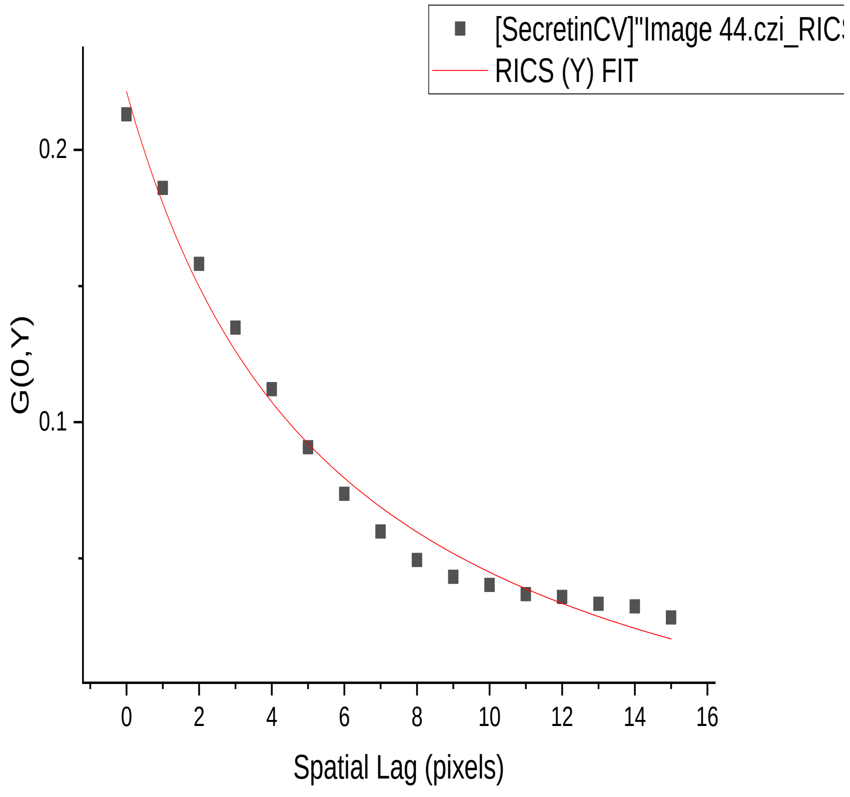
[SecretinCV]"Image 44.czi_RIC
RICS (Y) FIT
G(0,Y)
Spatial Lag (pixels)

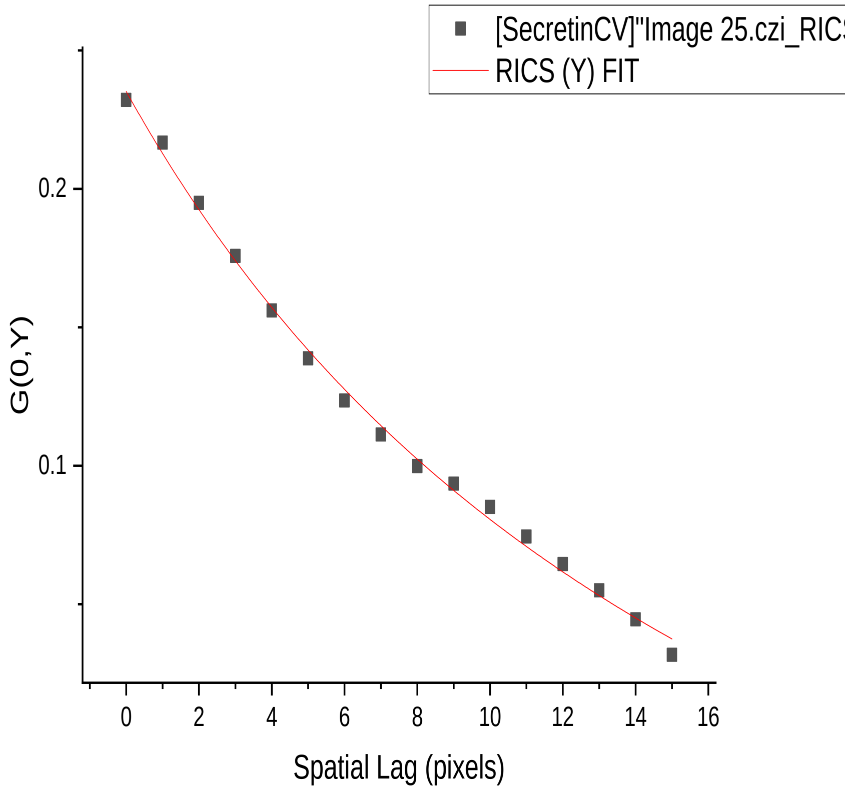
[SecretinCV]"Image 25.czi_RIC
RICS (Y) FIT
G(0,Y)
Spatial Lag (pixels)

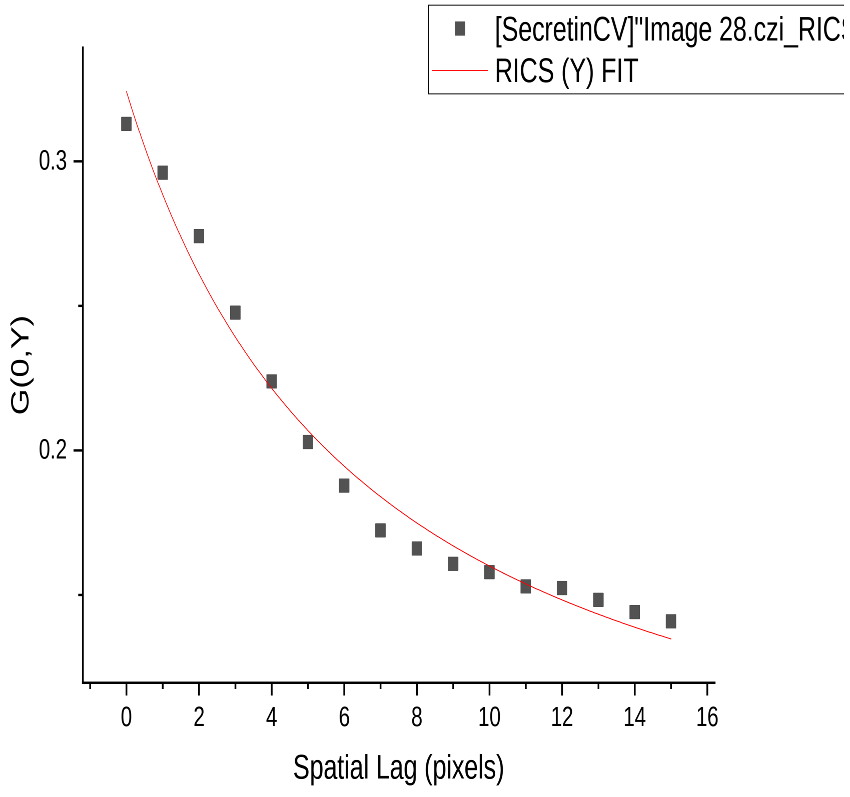
[SecretinCV]"Image 28.czi_RIC
RICS (Y) FIT
G(0,Y)
Spatial Lag (pixels)

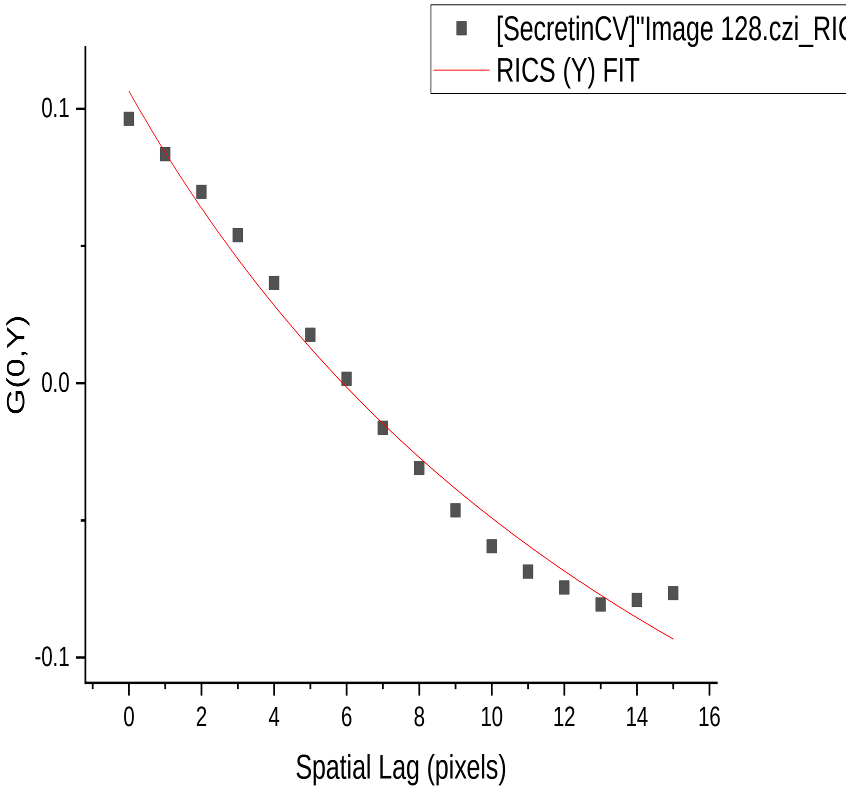
[SecretinCV]"Image 128.czi_RI
RICS (Y) FIT
G(0,Y)
Spatial Lag (pixels)

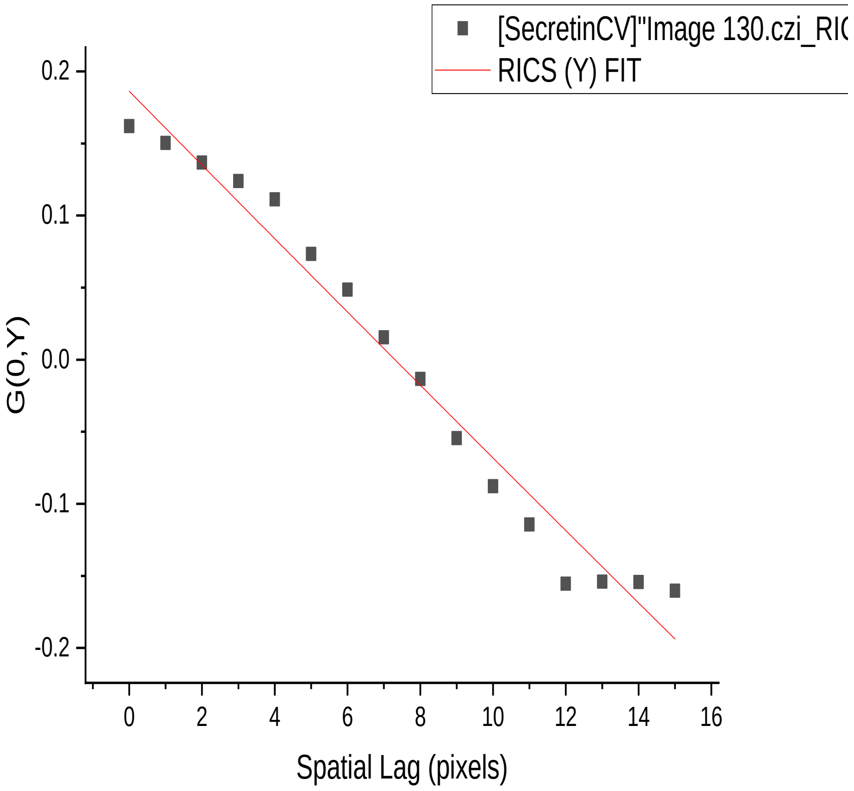
[SecretinCV]"Image 130.czi_RI
RICS (Y) FIT
G(0,Y)
Spatial Lag (pixels)

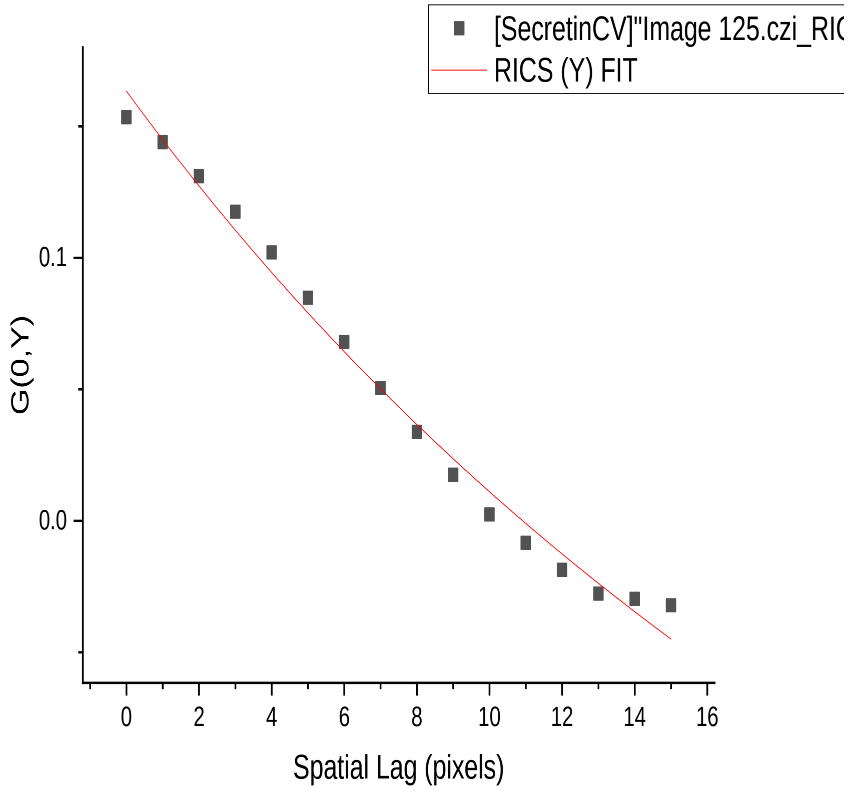
[SecretinCV]"Image 125.czi_RI
RICS (Y) FIT
G(0,Y)
Spatial Lag (pixels)

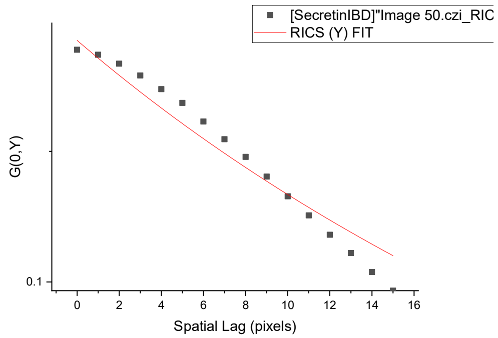
[SecretinIBD]"Image 50.czi_RIC
RICS (Y) FIT
G(0,Y)
Spatial Lag (pixels)

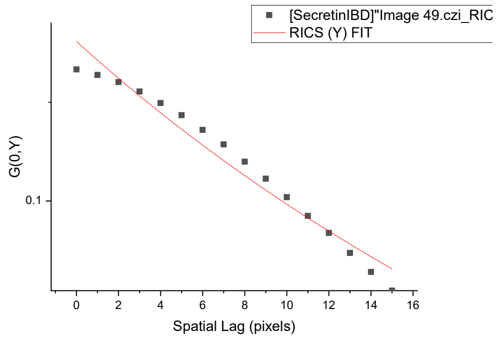
[SecretinIBD]"Image 49.czi_RIC
RICS (Y) FIT
G(0,Y)
Spatial Lag (pixels)

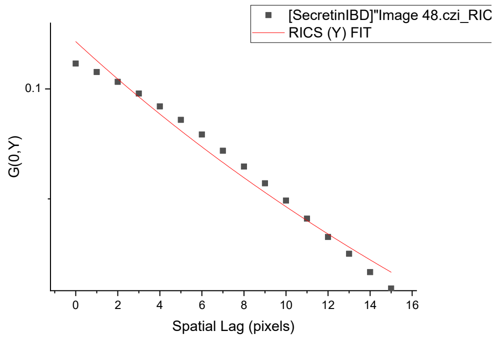
[SecretinIBD]"Image 48.czi_RIC
RICS (Y) FIT
G(0,Y)
Spatial Lag (pixels)

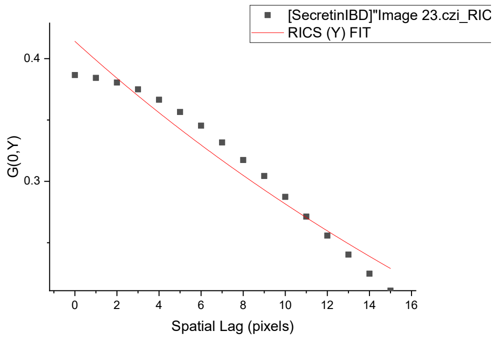
[SecretinIBD]"Image 23.czi_RIC
RICS (Y) FIT
G(0,Y)
Spatial Lag (pixels)

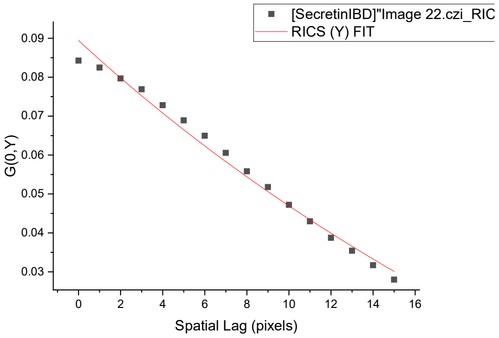
[SecretinIBD]"Image 22.czi_RIC
RICS (Y) FIT
G(0,Y)
Spatial Lag (pixels)

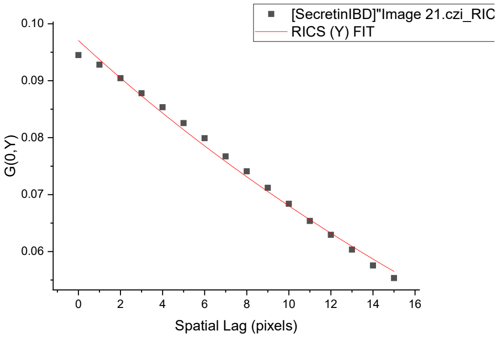
[SecretinIBD]"Image 21.czi_RIC
RICS (Y) FIT
G(0,Y)
Spatial Lag (pixels)

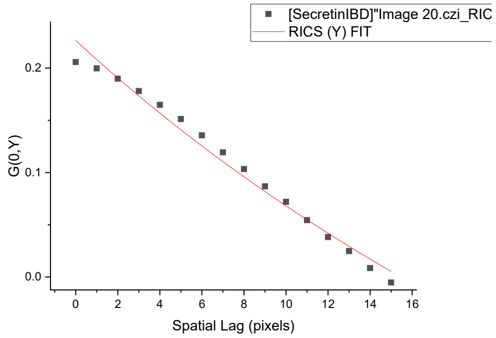
[SecretinIBD]"Image 20.czi_RIC
RICS (Y) FIT
G(0,Y)
Spatial Lag (pixels)

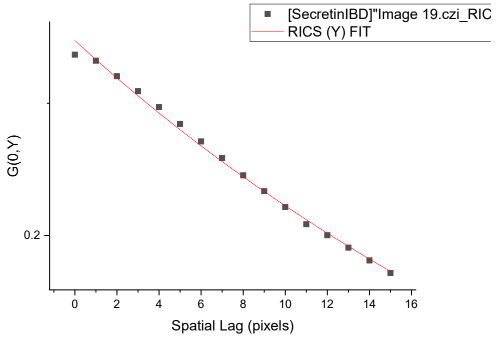
[SecretinIBD]"Image 19.czi_RIC
RICS (Y) FIT
G(0,Y)
Spatial Lag (pixels)

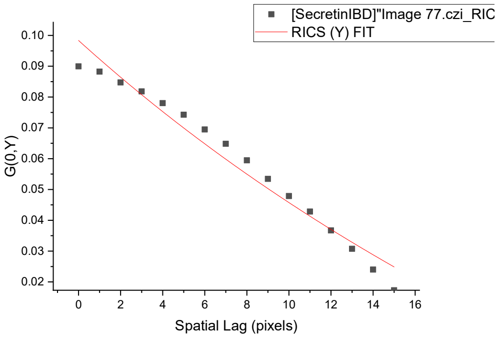
[SecretinIBD]"Image 77.czi_RIC
RICS (Y) FIT
G(0,Y)
Spatial Lag (pixels)

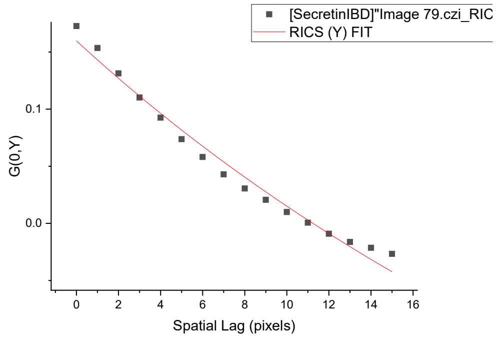
[SecretinIBD]"Image 79.czi_RIC
RICS (Y) FIT
G(0,Y)
Spatial Lag (pixels)

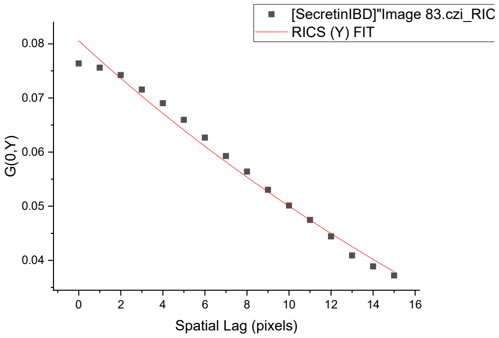
[SecretinIBD]"Image 83.czi_RIC
RICS (Y) FIT
G(0,Y)
Spatial Lag (pixels)

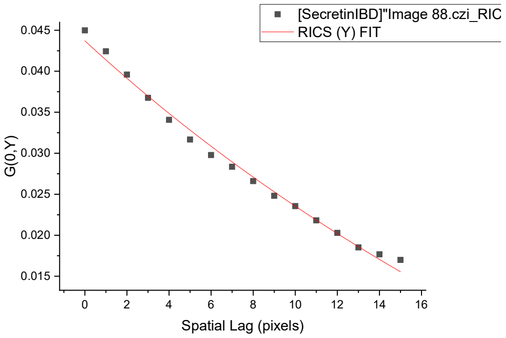
[SecretinIBD]"Image 88.czi_RIC
RICS (Y) FIT
G(0,Y)
Spatial Lag (pixels)

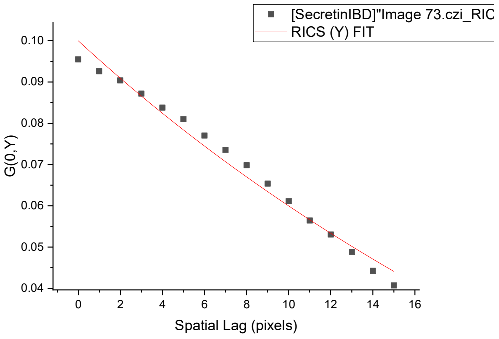
[SecretinIBD]"Image 73.czi_RIC
RICS (Y) FIT
G(0,Y)
Spatial Lag (pixels)

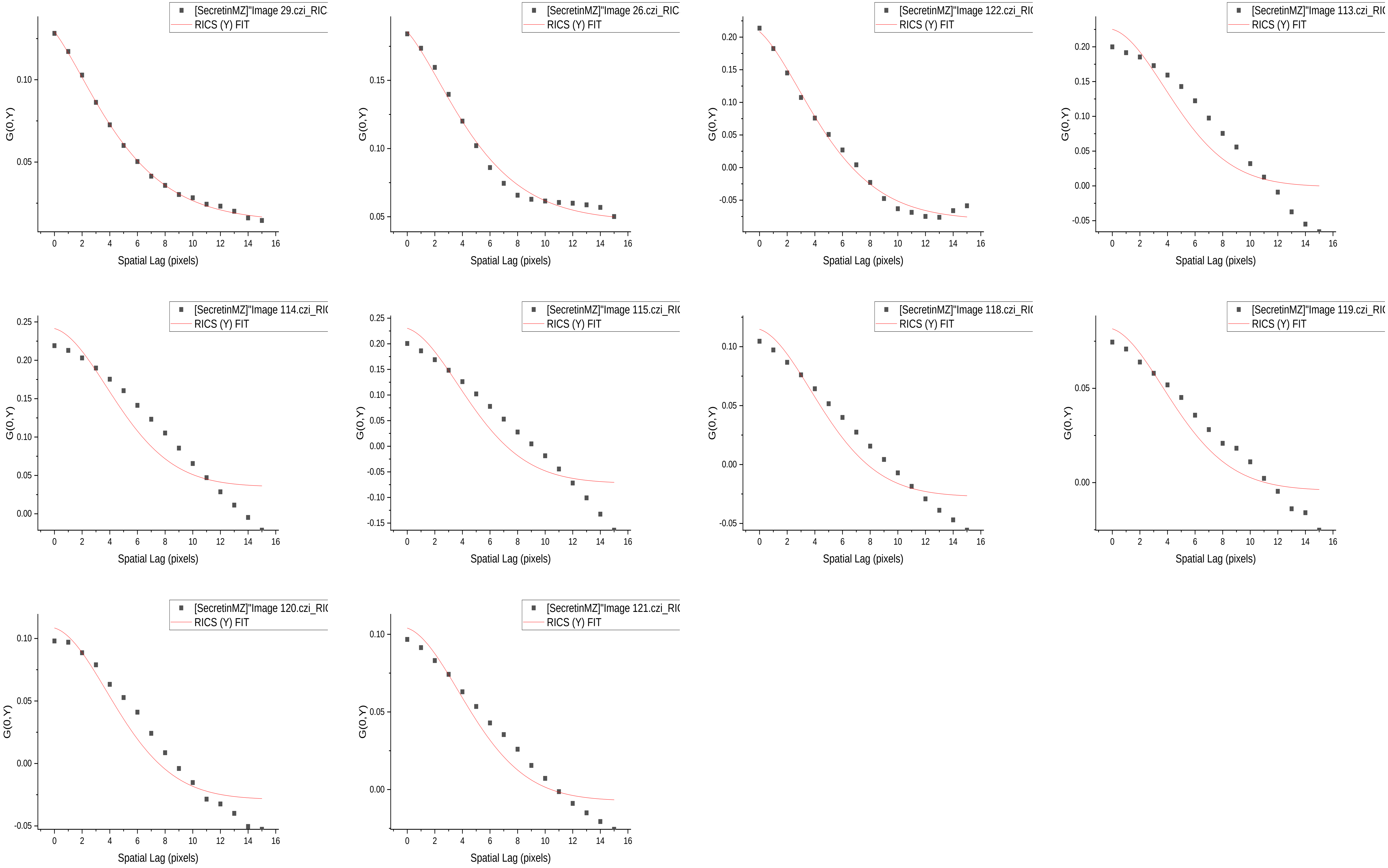
[SecretinMZ]"Image 29.czi_RIC
RICS (Y) FIT
G(0,Y)
Spatial Lag (pixels)
[SecretinMZ]"Image 26.czi_RIC
RICS (Y) FIT
G(0,Y)
Spatial Lag (pixels)
[SecretinMZ]"Image 122.czi_RI
RICS (Y) FIT
G(0,Y)
Spatial Lag (pixels)
[SecretinMZ]"Image 113.czi_RI
RICS (Y) FIT
G(0,Y)
Spatial Lag (pixels)
[SecretinMZ]"Image 114.czi_RI
RICS (Y) FIT
G(0,Y)
Spatial Lag (pixels)
[SecretinMZ]"Image 115.czi_RI
RICS (Y) FIT
G(0,Y)
Spatial Lag (pixels)
[SecretinMZ]"Image 118.czi_RI
RICS (Y) FIT
G(0,Y)
Spatial Lag (pixels)
[SecretinMZ]"Image 119.czi_RI
RICS (Y) FIT
G(0,Y)
Spatial Lag (pixels)
[SecretinMZ]"Image 120.czi_RI
RICS (Y) FIT
G(0,Y)
Spatial Lag (pixels)
[SecretinMZ]"Image 121.czi_RI
RICS (Y) FIT
G(0,Y)
Spatial Lag (pixels)

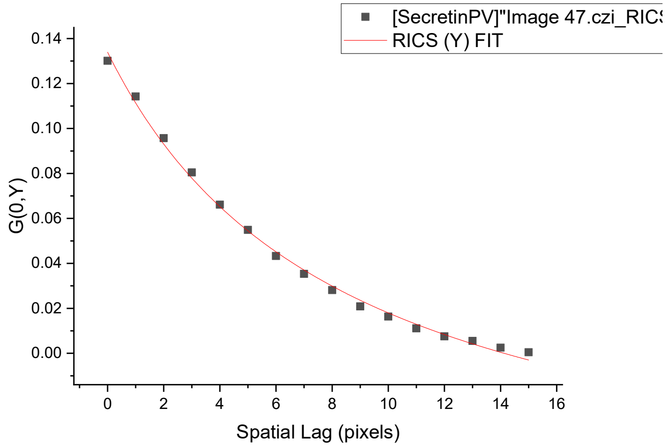
[SecretinPV]"Image 47.czi_RICS
RICS (Y) FIT
G(0,Y)
Spatial Lag (pixels)

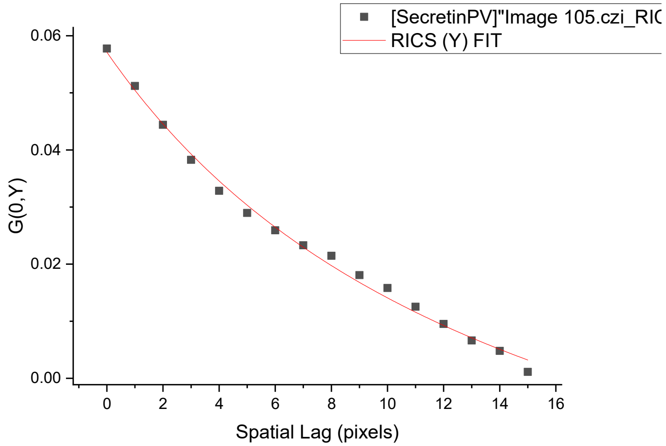
[SecretinPV]"Image 105.czi_RIC
RICS (Y) FIT
G(0,Y)
Spatial Lag (pixels)

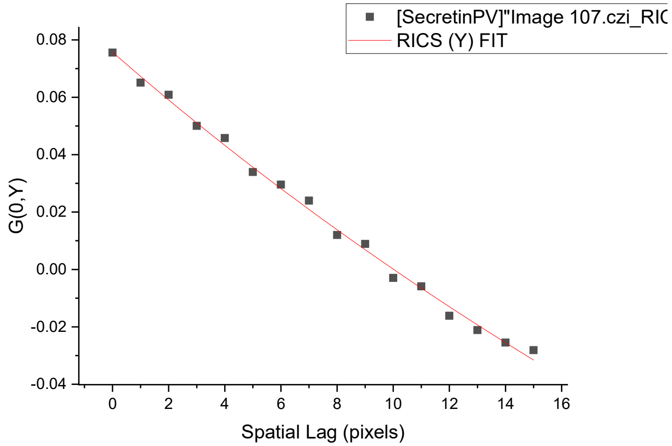
[SecretinPV]"Image 107.czi_RIC
RICS (Y) FIT
G(0,Y)
Spatial Lag (pixels)

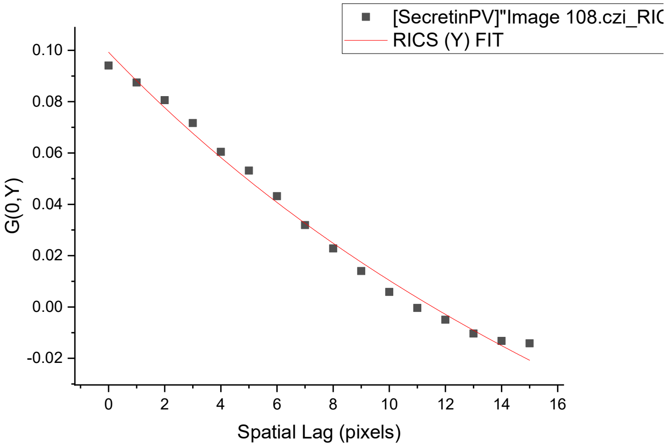
[SecretinPV]"Image 108.czi_RIC
RICS (Y) FIT
G(0,Y)
Spatial Lag (pixels)

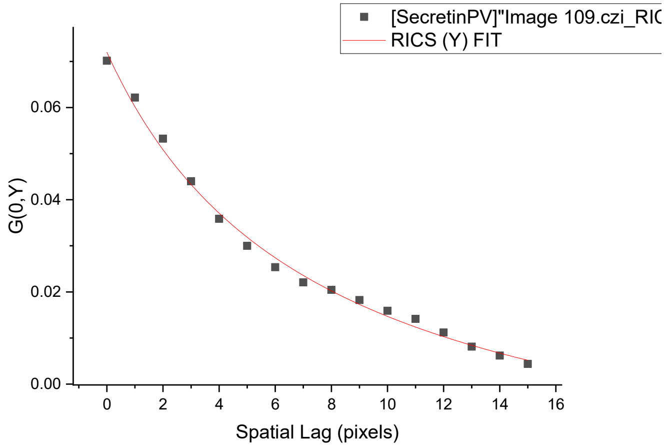
[SecretinPV]"Image 109.czi_RIC
RICS (Y) FIT
G(0,Y)
Spatial Lag (pixels)

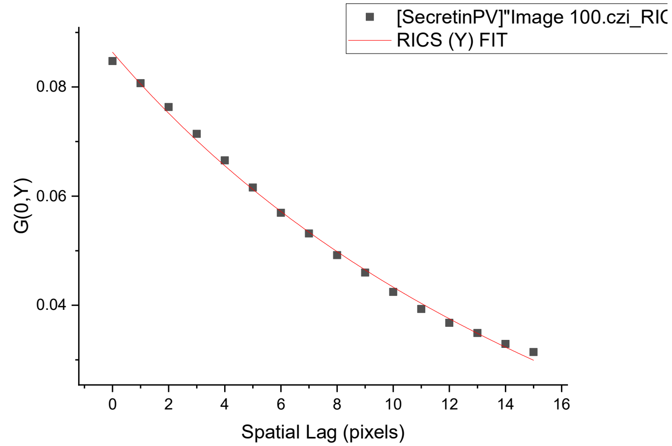
[SecretinPV]"Image 100.czi_RIC
RICS (Y) FIT
G(0,Y)
Spatial Lag (pixels)

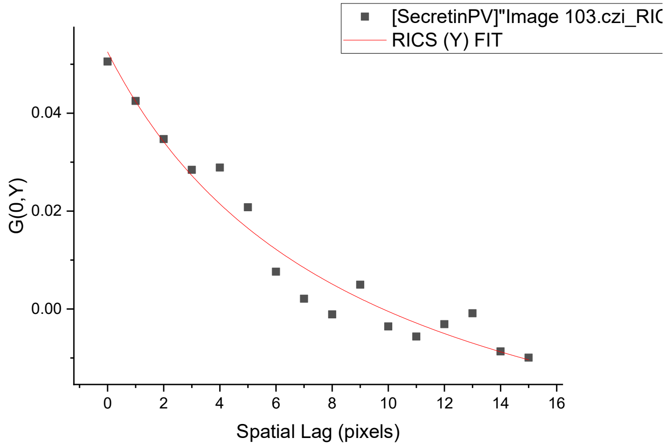
[SecretinPV]"Image 103.czi_RIC
RICS (Y) FIT
G(0,Y)
Spatial Lag (pixels)

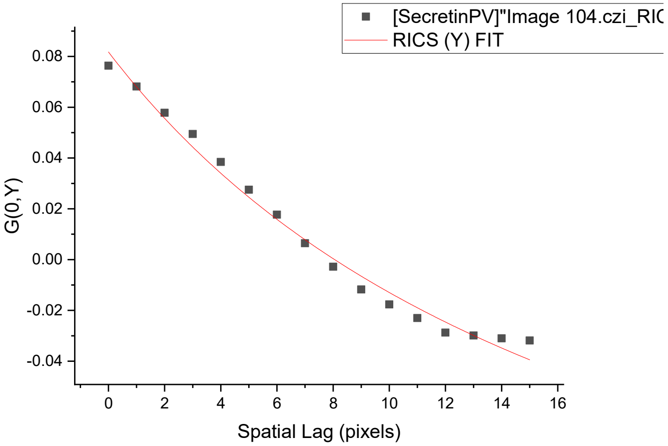
[SecretinPV]"Image 104.czi_RIC
RICS (Y) FIT
G(0,Y)
Spatial Lag (pixels)

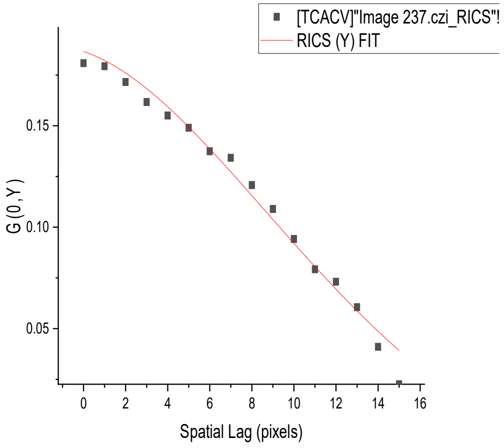
[TCACV]"Image 237.czi_RICS"!
RICS (Y) FIT
G (0, Y)
Spatial Lag (pixels)

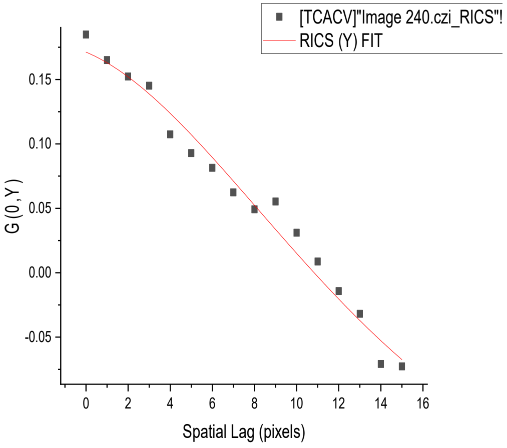
[TCACV]"Image 240.czi_RICS"!
RICS (Y) FIT
G (0, Y)
Spatial Lag (pixels)

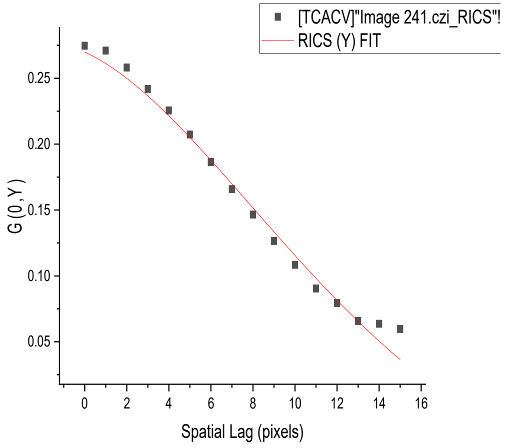
[TCACV]"Image 241.czi_RICS"!
RICS (Y) FIT
G (0, Y)
Spatial Lag (pixels)

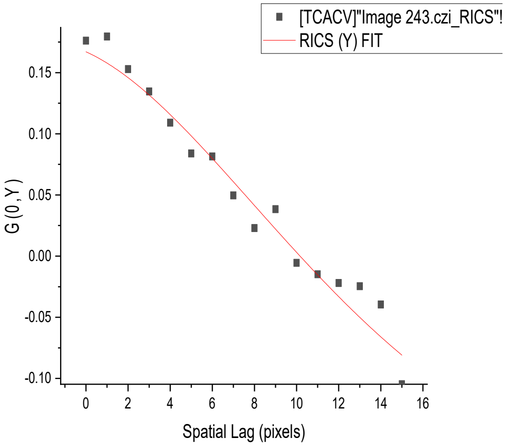
[TCACV]"Image 243.czi_RICS"!
RICS (Y) FIT
G (0, Y)
Spatial Lag (pixels)

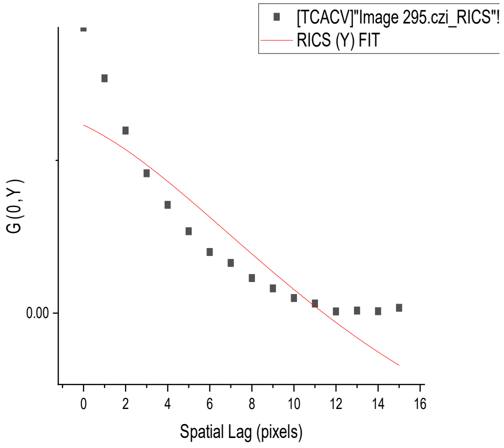
[TCACV]"Image 295.czi_RICS"!
RICS (Y) FIT
G (0, Y)
Spatial Lag (pixels)

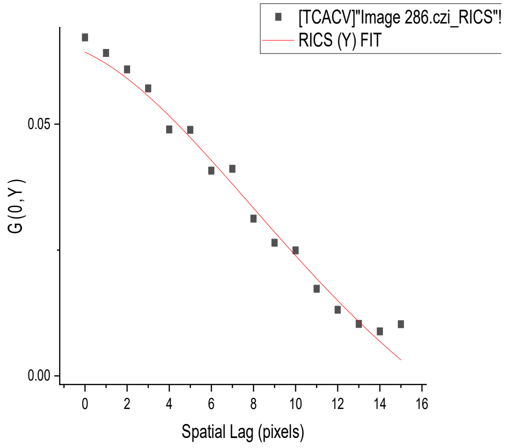
[TCACV]"Image 286.czi_RICS"!
RICS (Y) FIT
G (0, Y)
Spatial Lag (pixels)

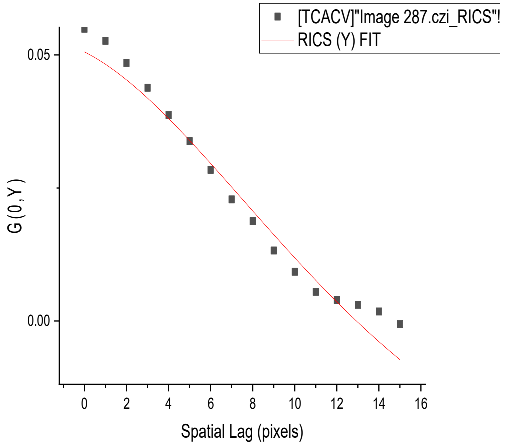
[TCACV]"Image 287.czi_RICS"!
RICS (Y) FIT
G (0, Y)
Spatial Lag (pixels)

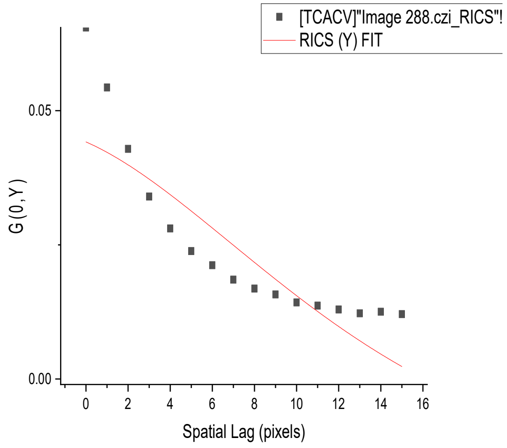
[TCACV]"Image 288.czi_RICS"!
RICS (Y) FIT
G (0, Y)
Spatial Lag (pixels)

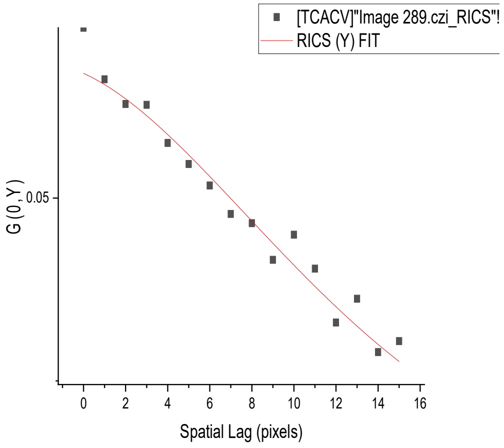
[TCACV]"Image 289.czi_RICS"!
RICS (Y) FIT
G (0, Y)
Spatial Lag (pixels)

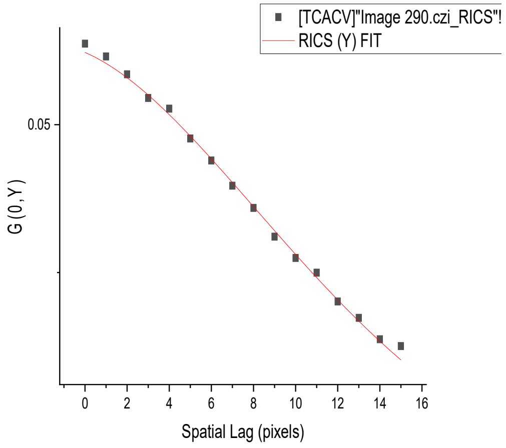
[TCACV]"Image 290.czi_RICS"!
RICS (Y) FIT
G (0, Y)
Spatial Lag (pixels)

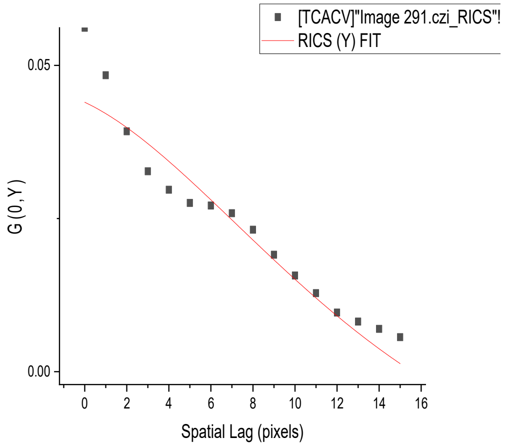
[TCACV]"Image 291.czi_RICS"!
RICS (Y) FIT
G (0, Y)
Spatial Lag (pixels)

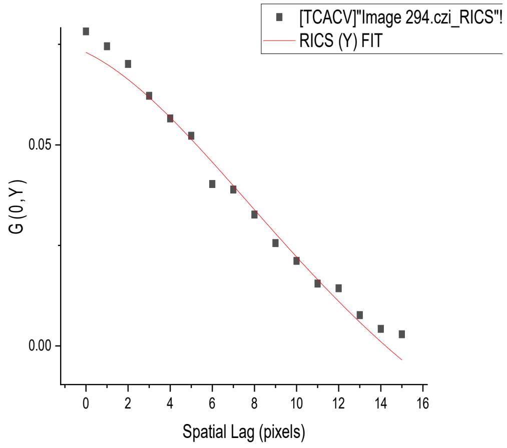
[TCACV]"Image 294.czi_RICS"!
RICS (Y) FIT
G (0, Y)
Spatial Lag (pixels)

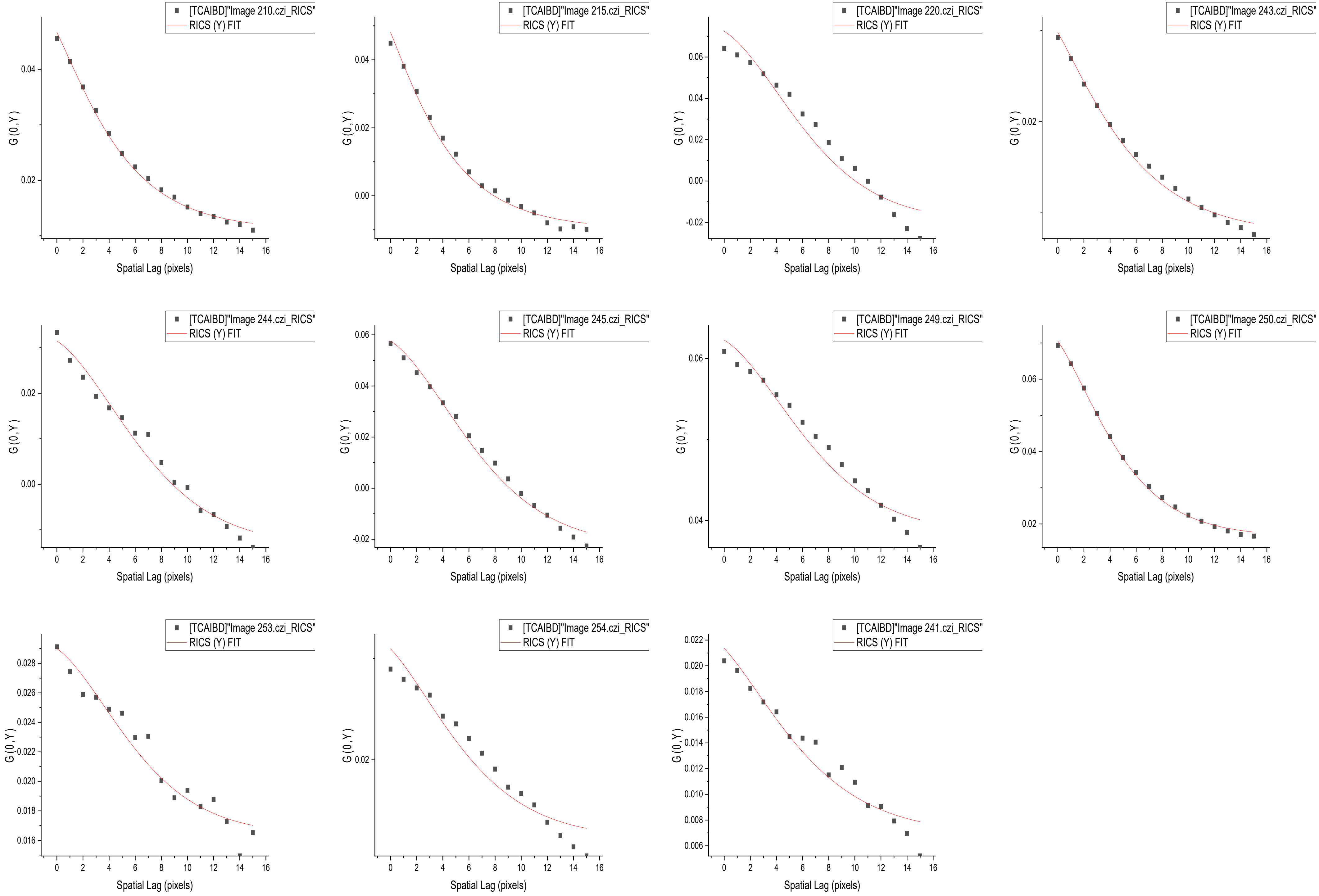
[TCAIBD]"Image 210.czi_RICS"
RICS (Y) FIT
[TCAIBD]"Image 215.czi_RICS"
RICS (Y) FIT
[TCAIBD]"Image 220.czi_RICS"
RICS (Y) FIT
[TCAIBD]"Image 243.czi_RICS"
RICS (Y) FIT
[TCAIBD]"Image 244.czi_RICS"
RICS (Y) FIT
[TCAIBD]"Image 245.czi_RICS"
RICS (Y) FIT
[TCAIBD]"Image 249.czi_RICS"
RICS (Y) FIT
[TCAIBD]"Image 250.czi_RICS"
RICS (Y) FIT
[TCAIBD]"Image 253.czi_RICS"
RICS (Y) FIT
[TCAIBD]"Image 254.czi_RICS"
RICS (Y) FIT
[TCAIBD]"Image 241.czi_RICS"
RICS (Y) FIT
G(0,Y)
Spatial Lag (pixels)

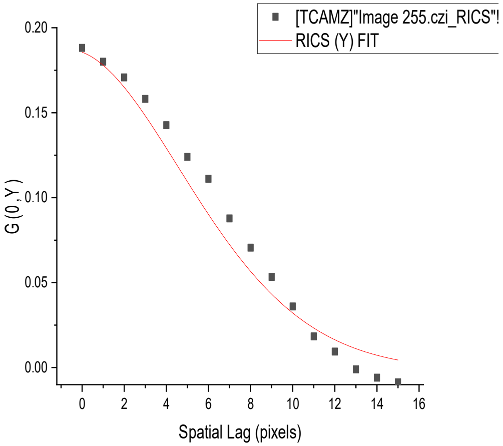
[TCAMZ]"Image 255.czi_RICS"!
RICS (Y) FIT
G (0, Y)
0.20
0.15
0.10
0.05
0.00
0 2 4 6 8 10 12 14 16
Spatial Lag (pixels)

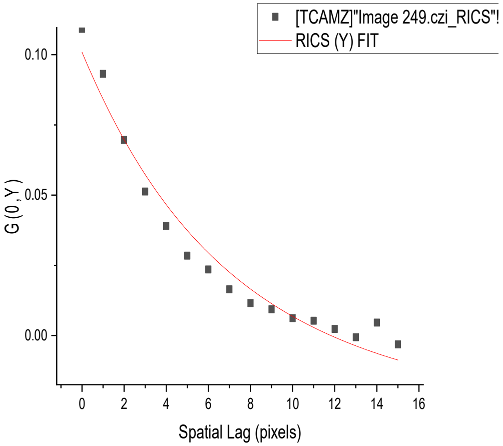
[TCAMZ]"Image 249.czi_RICS"!
RICS (Y) FIT
G (0, Y)
0.10
0.05
0.00
0 2 4 6 8 10 12 14 16
Spatial Lag (pixels)

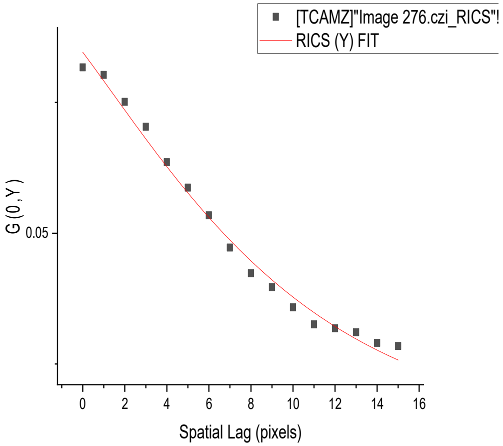
[TCAMZ]"Image 276.czi_RICS"!
RICS (Y) FIT
G (0, Y)
0.05
0 2 4 6 8 10 12 14 16
Spatial Lag (pixels)

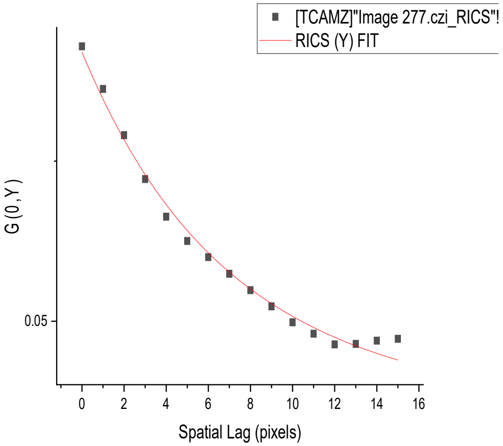
[TCAMZ]"Image 277.czi_RICS"!
RICS (Y) FIT
G (0, Y)
0.05
0 2 4 6 8 10 12 14 16
Spatial Lag (pixels)

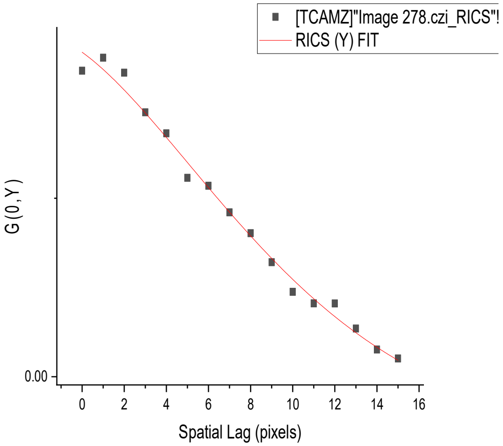
[TCAMZ]"Image 278.czi_RICS"!
RICS (Y) FIT
G (0, Y)
0.00
0 2 4 6 8 10 12 14 16
Spatial Lag (pixels)

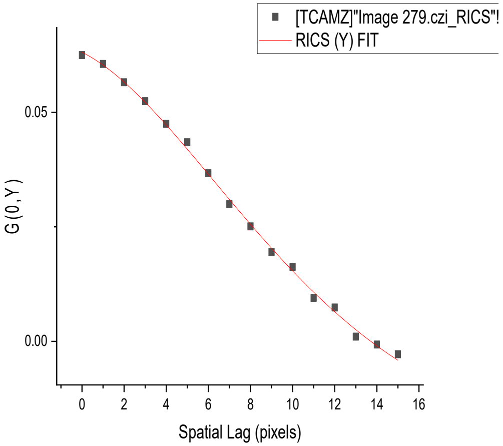
[TCAMZ]"Image 279.czi_RICS"!
RICS (Y) FIT
G (0, Y)
0.05
0.00
0 2 4 6 8 10 12 14 16
Spatial Lag (pixels)

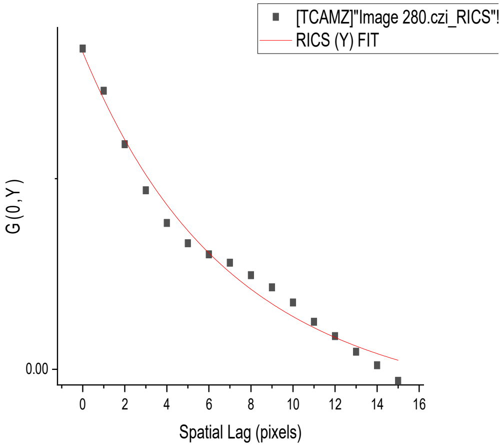
[TCAMZ]"Image 280.czi_RICS"!
RICS (Y) FIT
G (0, Y)
0.00
0 2 4 6 8 10 12 14 16
Spatial Lag (pixels)

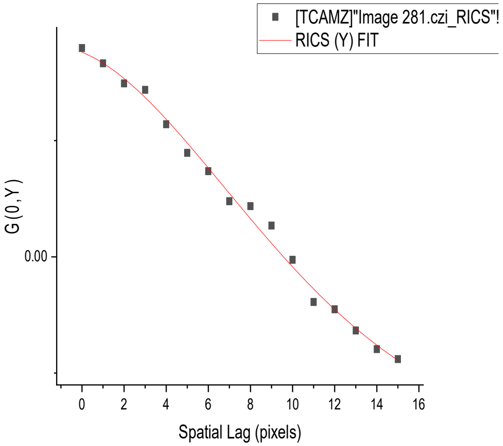
[TCAMZ]"Image 281.czi_RICS"!
RICS (Y) FIT
G (0, Y)
0.00
0 2 4 6 8 10 12 14 16
Spatial Lag (pixels)

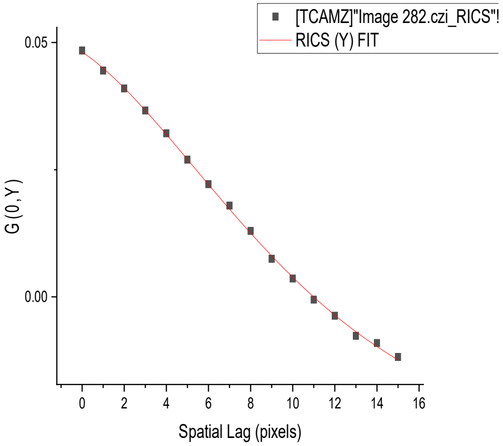
[TCAMZ]"Image 282.czi_RICS"!
RICS (Y) FIT
G (0, Y)
0.05
0.00
0 2 4 6 8 10 12 14 16
Spatial Lag (pixels)

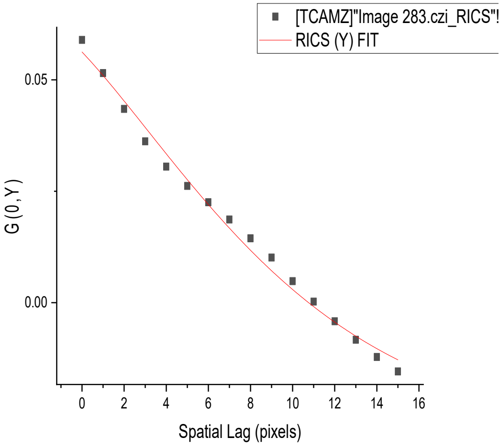
[TCAMZ]"Image 283.czi_RICS"!
RICS (Y) FIT
G (0, Y)
0.05
0.00
0 2 4 6 8 10 12 14 16
Spatial Lag (pixels)

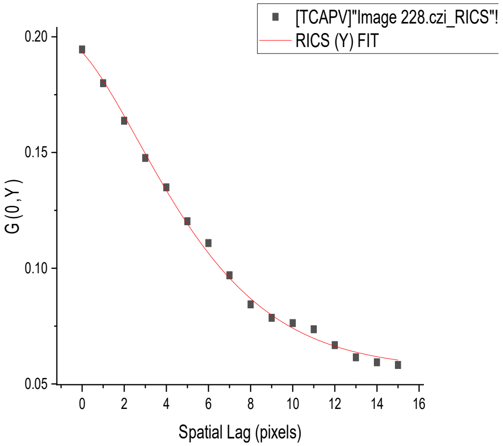
[TCAPV]"Image 228.czi_RICS"!
RICS (Y) FIT
G (0, Y)
Spatial Lag (pixels)

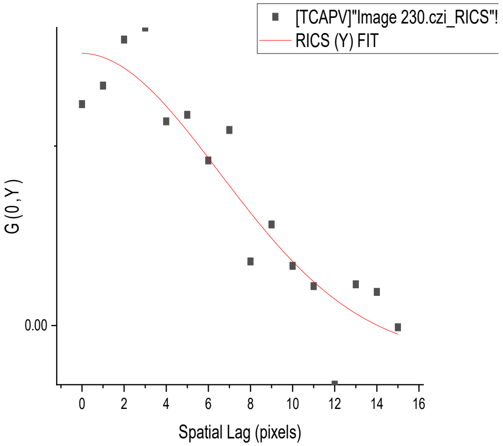
[TCAPV]"Image 230.czi_RICS"!
RICS (Y) FIT
G (0, Y)
Spatial Lag (pixels)

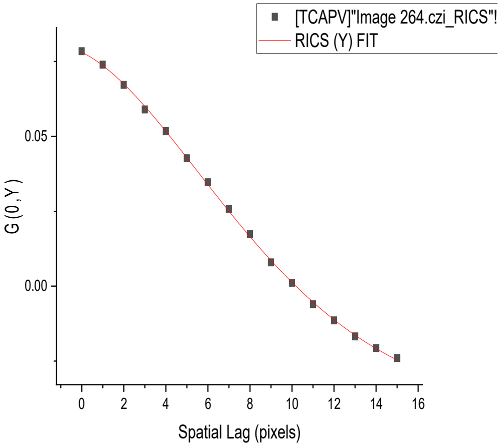
[TCAPV]"Image 264.czi_RICS"!
RICS (Y) FIT
G (0, Y)
Spatial Lag (pixels)

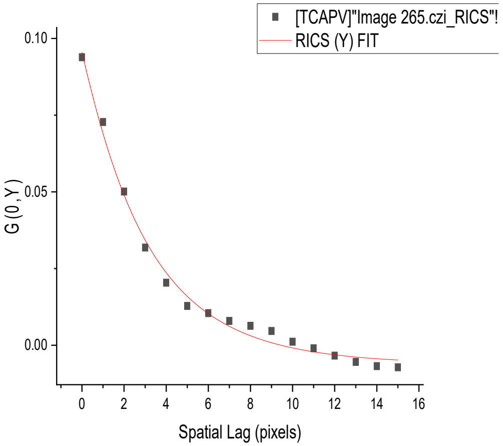
[TCAPV]"Image 265.czi_RICS"!
RICS (Y) FIT
G (0, Y)
Spatial Lag (pixels)

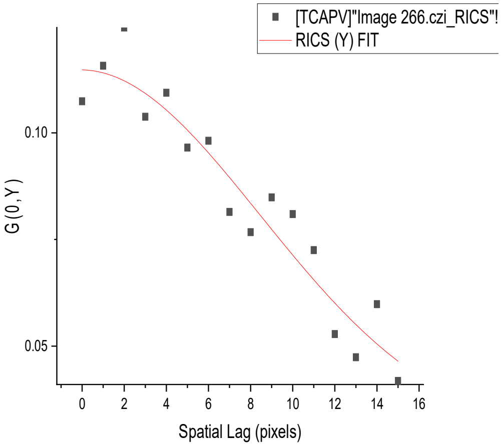
[TCAPV]"Image 266.czi_RICS"!
RICS (Y) FIT
G (0, Y)
Spatial Lag (pixels)

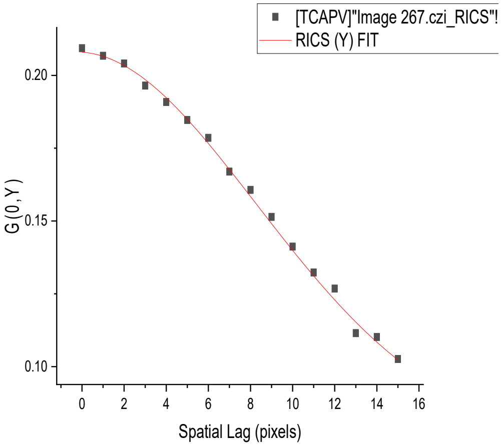
[TCAPV]"Image 267.czi_RICS"!
RICS (Y) FIT
G (0, Y)
Spatial Lag (pixels)

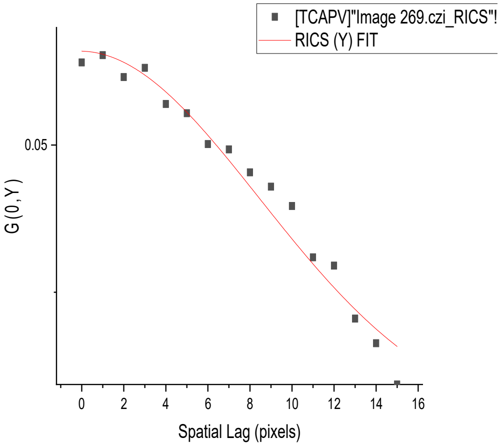
[TCAPV]"Image 269.czi_RICS"!
RICS (Y) FIT
G (0, Y)
Spatial Lag (pixels)

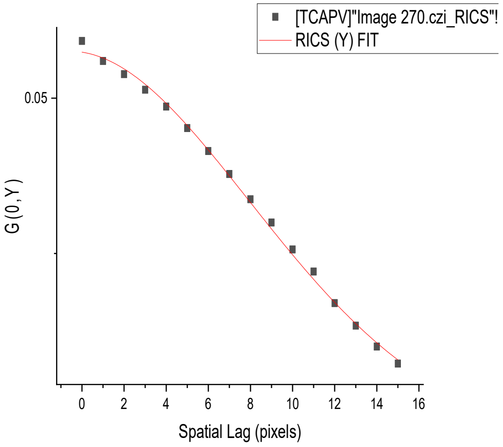
[TCAPV]"Image 270.czi_RICS"!
RICS (Y) FIT
G (0, Y)
Spatial Lag (pixels)

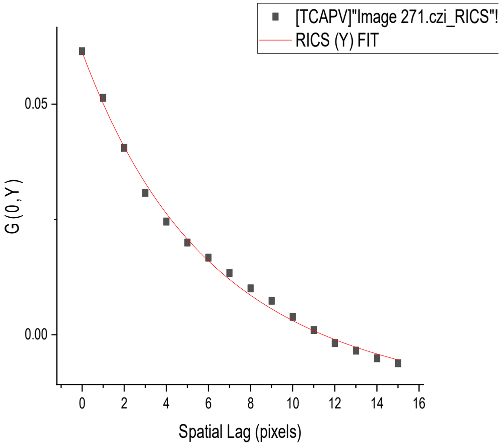
[TCAPV]"Image 271.czi_RICS"!
RICS (Y) FIT
G (0, Y)
Spatial Lag (pixels)

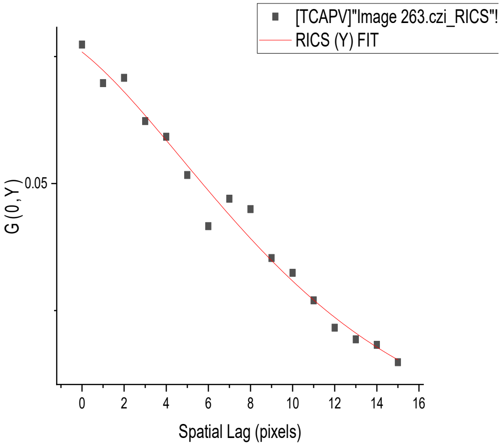
[TCAPV]"Image 263.czi_RICS"!
RICS (Y) FIT
G (0, Y)
Spatial Lag (pixels)

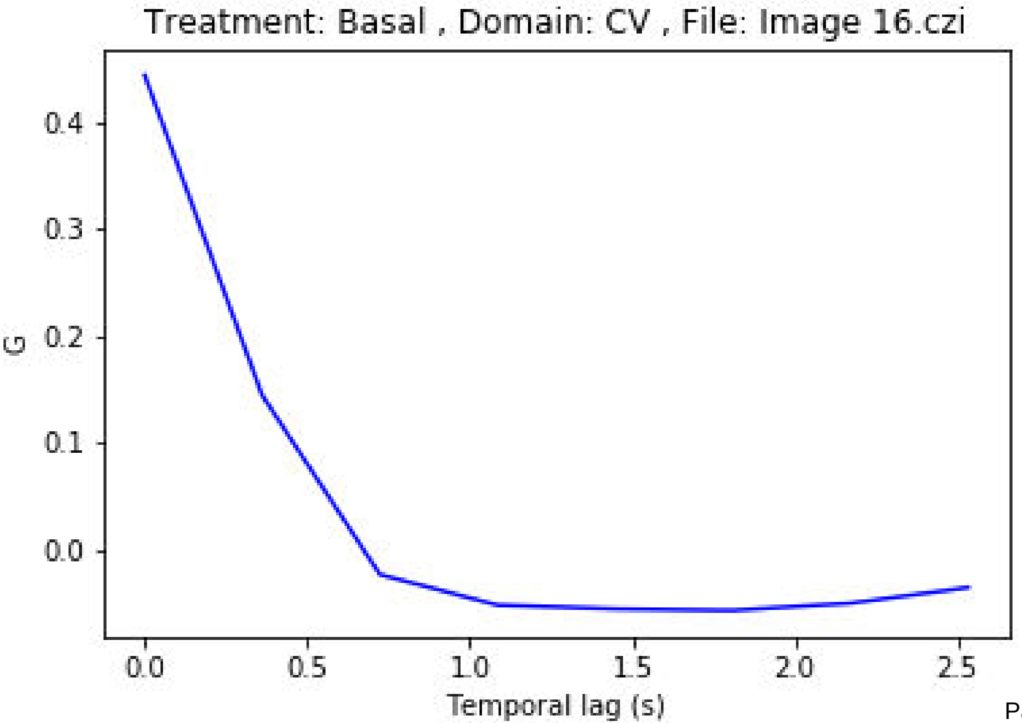
Treatment: Basal , Domain: CV , File: Image 16.czi
G
0.4
0.3
0.2
0.1
0.0
0.0
0.5
1.0
1.5
2.0
2.5
Temporal lag (s)

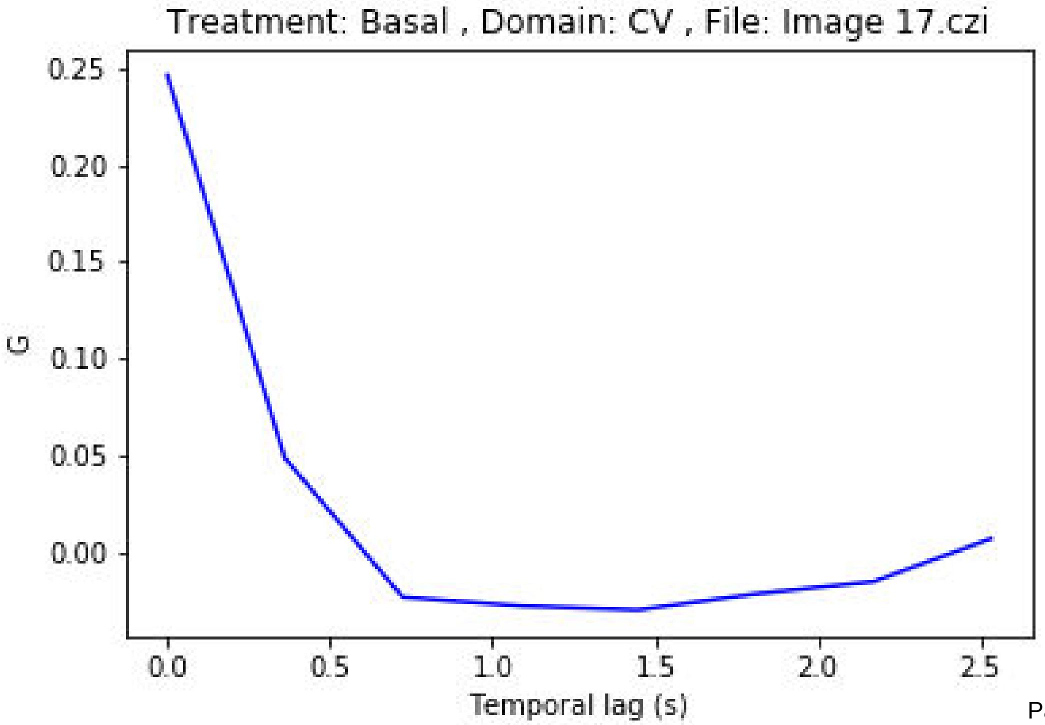
Treatment: Basal , Domain: CV , File: Image 17.czi
G
0.25
0.20
0.15
0.10
0.05
0.00
0.0
0.5
1.0
1.5
2.0
2.5
Temporal lag (s)

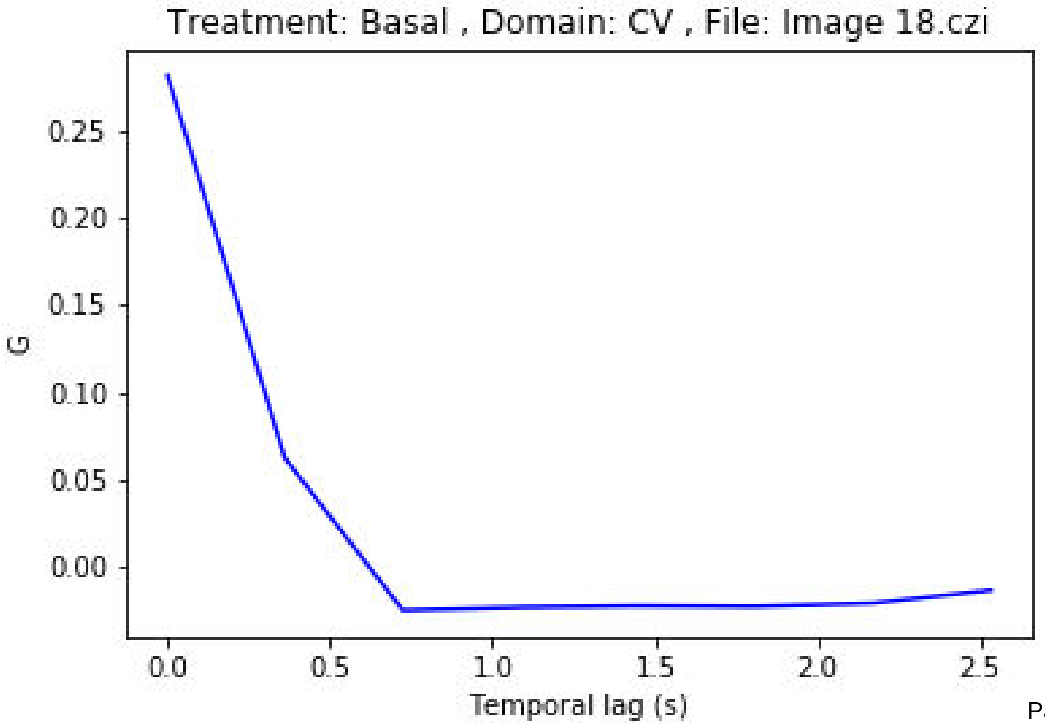
Treatment: Basal , Domain: CV , File: Image 18.czi
G
0.25
0.20
0.15
0.10
0.05
0.00
0.0
0.5
1.0
1.5
2.0
2.5
Temporal lag (s)

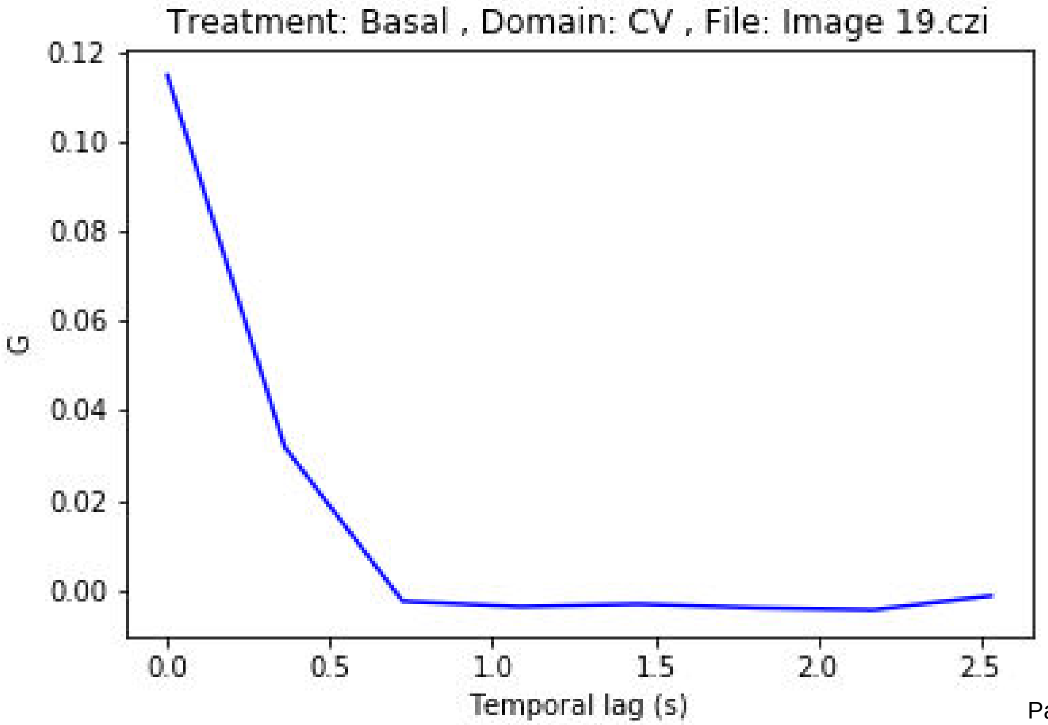
Treatment: Basal , Domain: CV , File: Image 19.czi
0.12
0.10
0.08
0.06
0.04
0.02
0.00
G
0.0
0.5
1.0
1.5
2.0
2.5
Temporal lag (s)

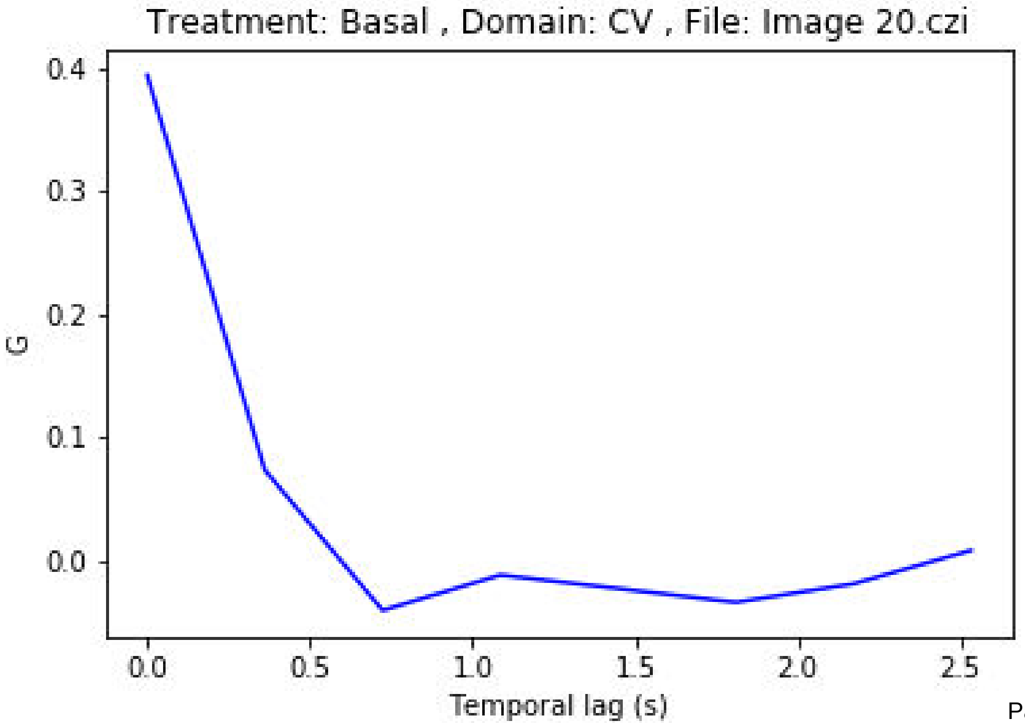
Treatment: Basal , Domain: CV , File: Image 20.czi
G
0.4
0.3
0.2
0.1
0.0
0.0
0.5
1.0
1.5
2.0
2.5
Temporal lag (s)

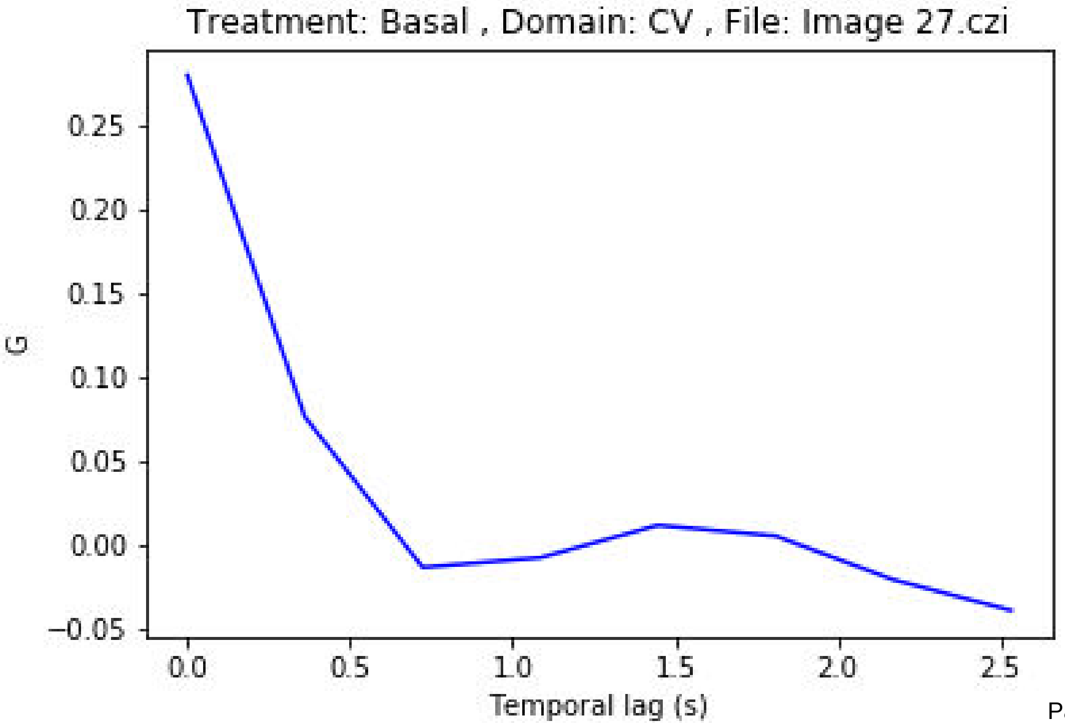
Treatment: Basal , Domain: CV , File: Image 27.czi
G
0.25
0.20
0.15
0.10
0.05
0.00
−0.05
0.0
0.5
1.0
1.5
2.0
2.5
Temporal lag (s)

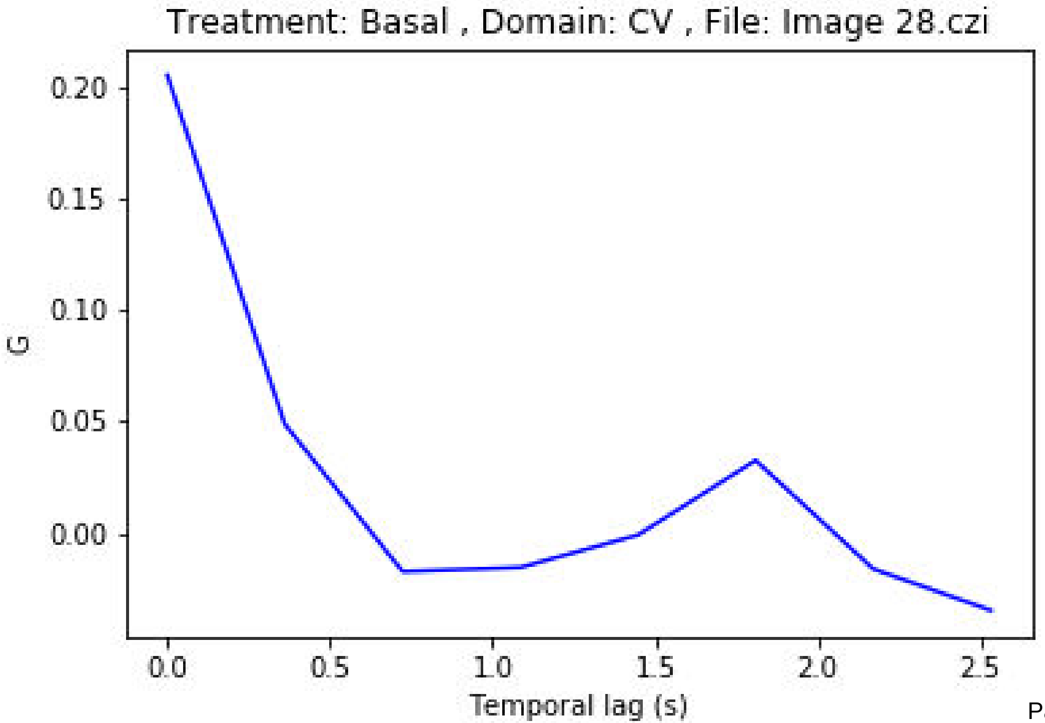
Treatment: Basal , Domain: CV , File: Image 28.czi
0.20
0.15
0.10
0.05
0.00
G
0.0
0.5
1.0
1.5
2.0
2.5
Temporal lag (s)

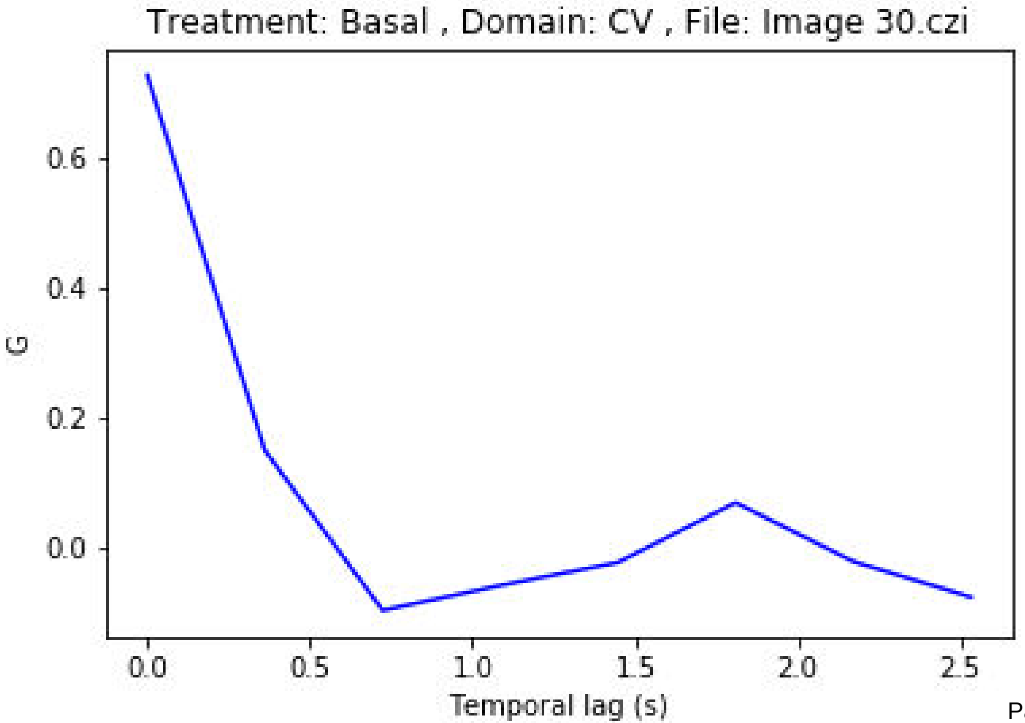
Treatment: Basal , Domain: CV , File: Image 30.czi
0.6
0.4
0.2
0.0
G
0.0
0.5
1.0
1.5
2.0
2.5
Temporal lag (s)

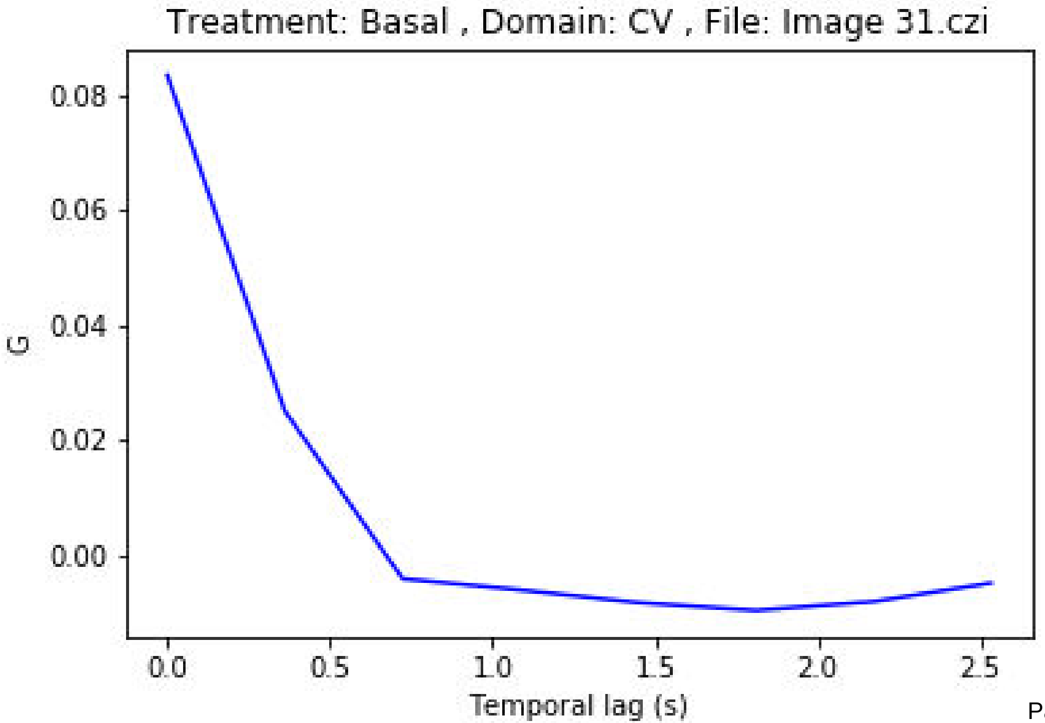
Treatment: Basal , Domain: CV , File: Image 31.czi
0.08
0.06
0.04
0.02
0.00
G
0.0
0.5
1.0
1.5
2.0
2.5
Temporal lag (s)

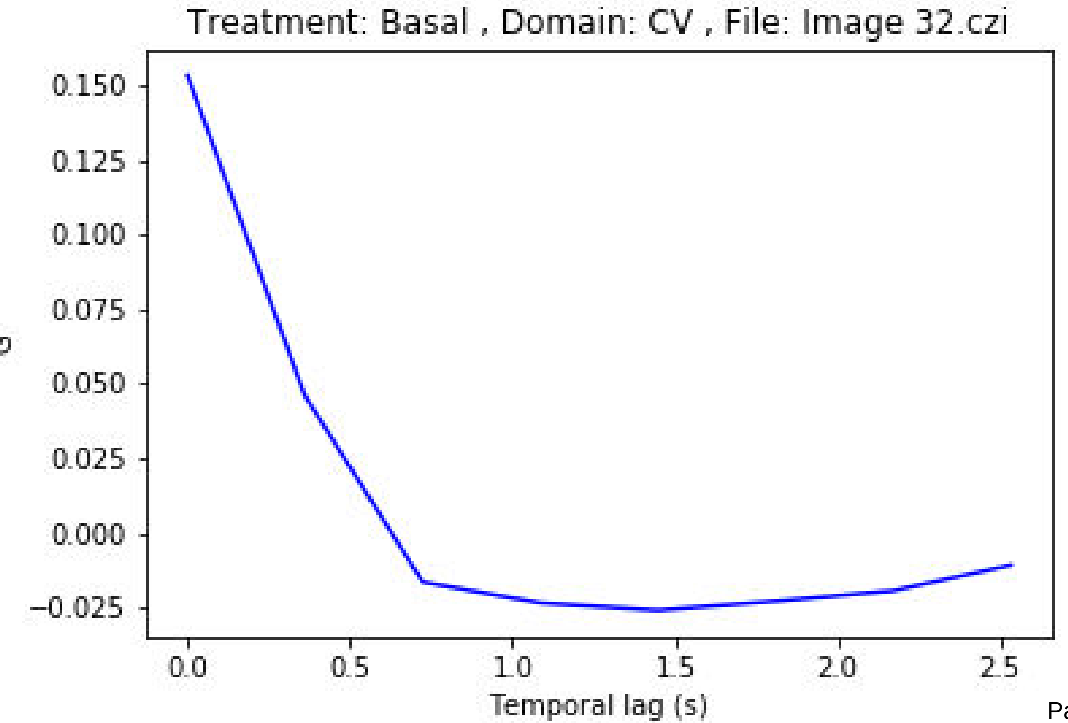
Treatment: Basal , Domain: CV , File: Image 32.czi
0.150
0.125
0.100
0.075
0.050
0.025
0.000
−0.025
0.0
0.5
1.0
1.5
2.0
2.5
Temporal lag (s)

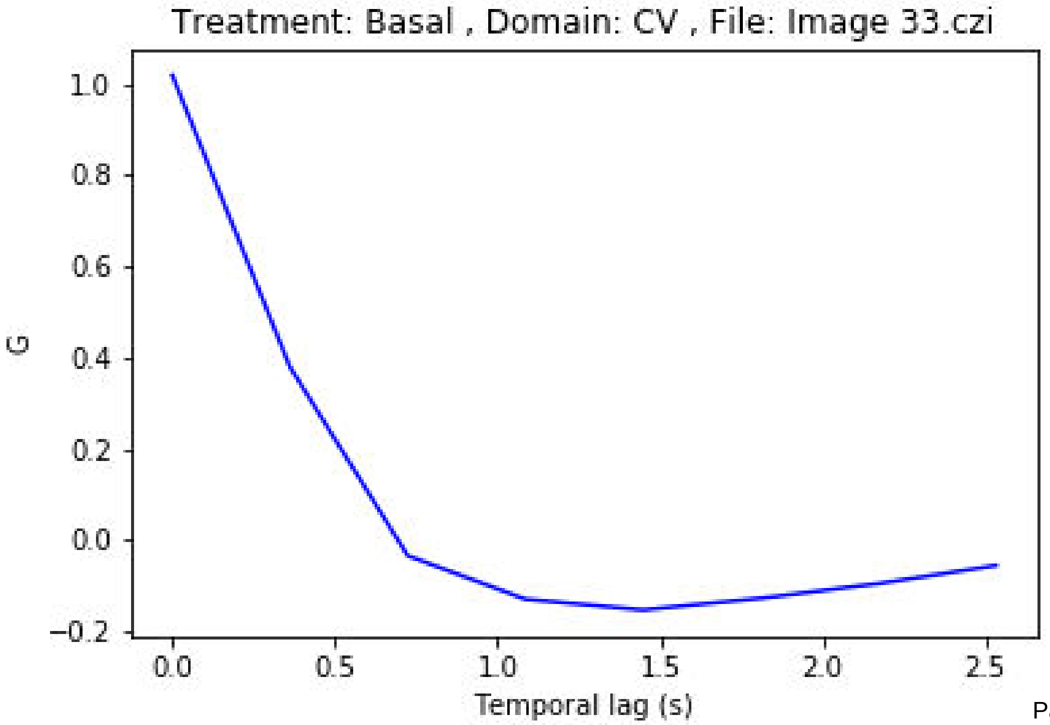
Treatment: Basal , Domain: CV , File: Image 33.czi
1.0
0.8
0.6
0.4
0.2
0.0
−0.2
G
0.0
0.5
1.0
1.5
2.0
2.5
Temporal lag (s)

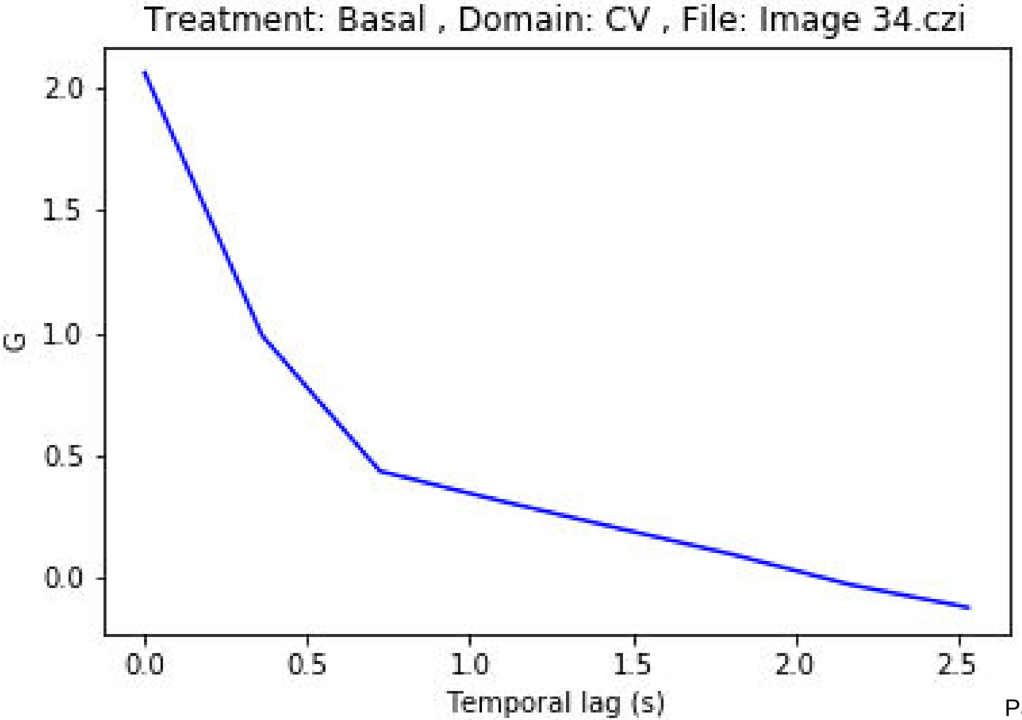
Treatment: Basal , Domain: CV , File: Image 34.czi
G
2.0
1.5
1.0
0.5
0.0
0.0
0.5
1.0
1.5
2.0
2.5
Temporal lag (s)

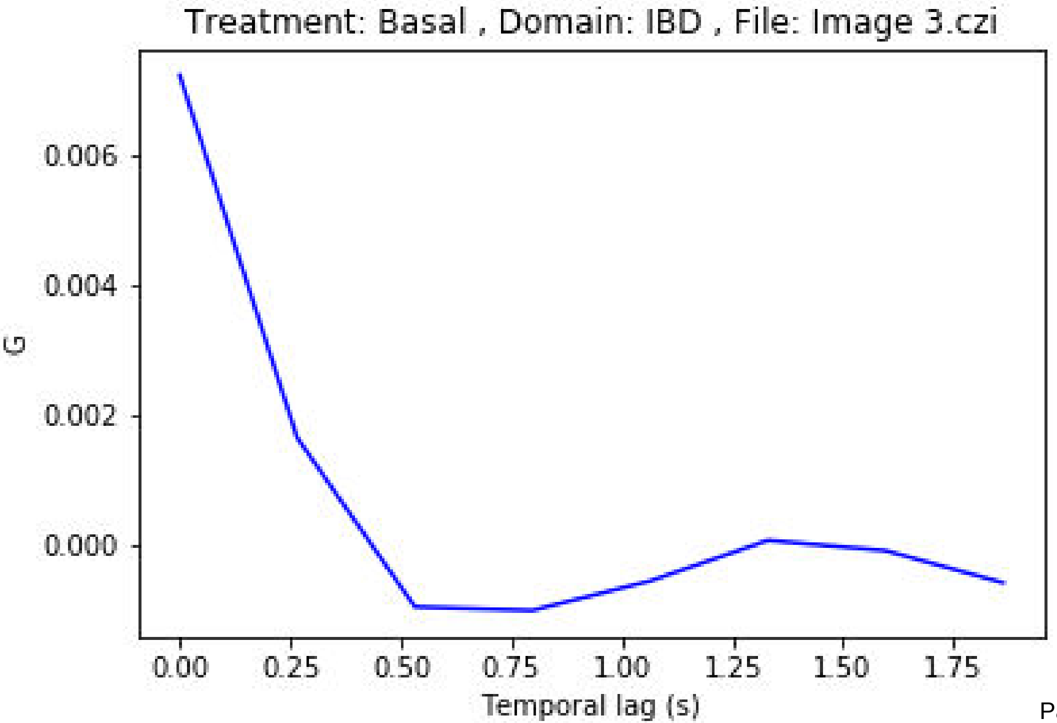

Treatment: Basal , Domain: IBD , File: Image 3.czi
0.006
0.004
0.002
0.000
G
0.00
0.25
0.50
0.75
1.00
1.25
1.50
1.75
Temporal lag (s)

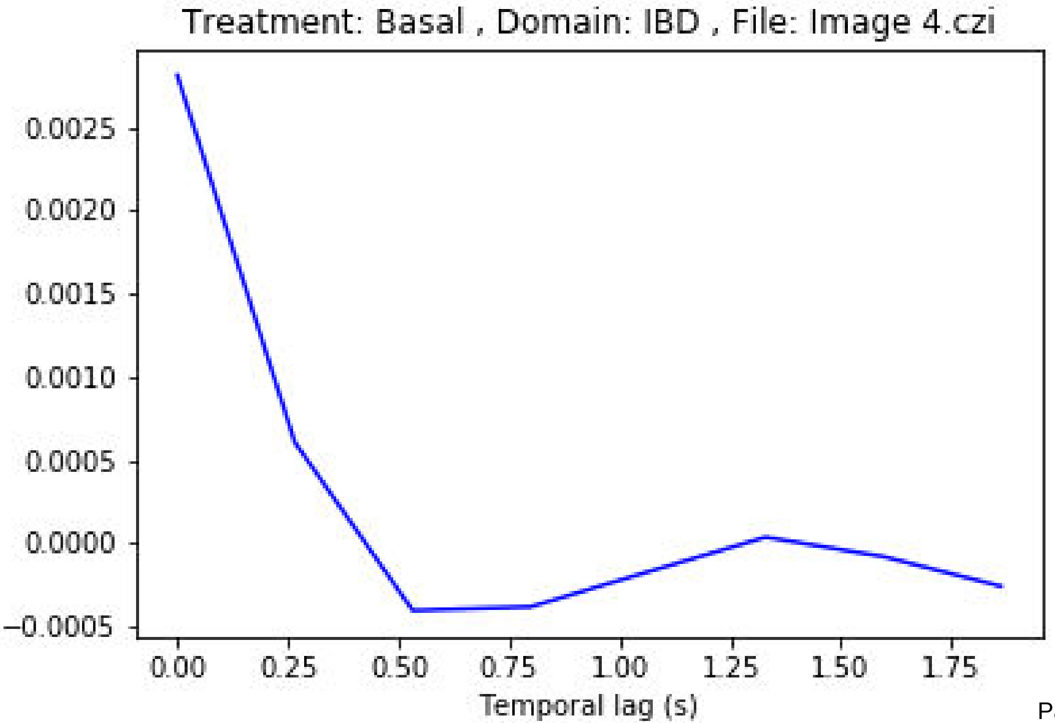
Treatment: Basal , Domain: IBD , File: Image 4.czi
0.0025
0.0020
0.0015
0.0010
0.0005
0.0000
−0.0005
0.00
0.25
0.50
0.75
1.00
1.25
1.50
1.75
Temporal lag (s)

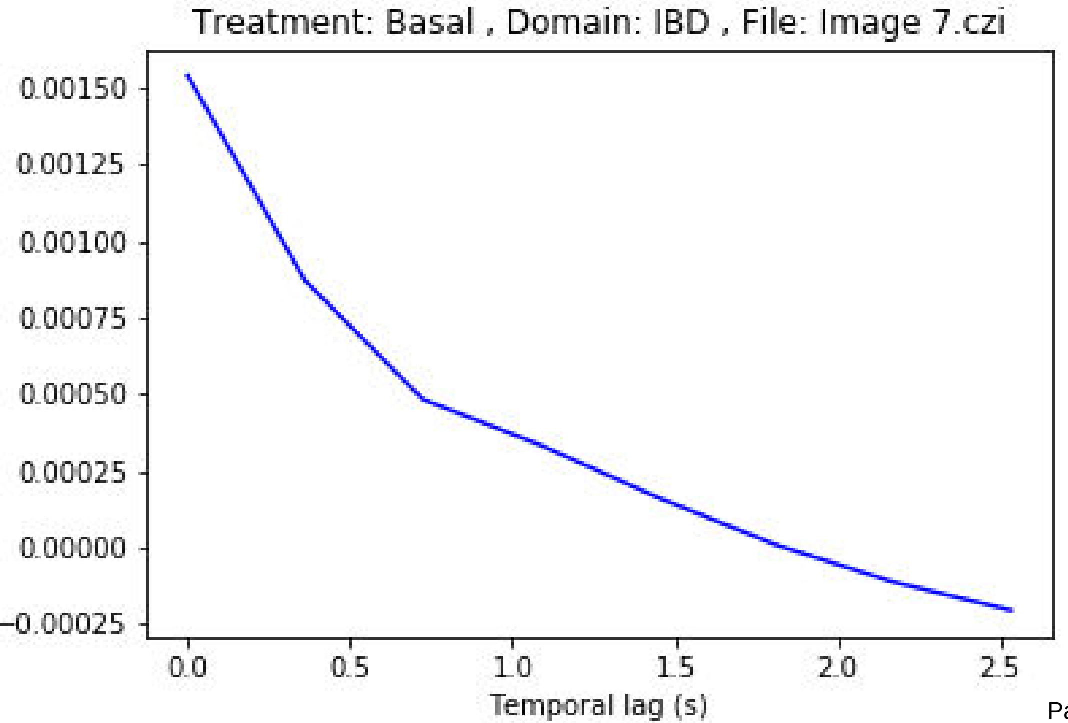

Treatment: Basal , Domain: IBD , File: Image 7.czi
0.00150
0.00125
0.00100
0.00075
0.00050
0.00025
0.00000
−0.00025
0.0
0.5
1.0
1.5
2.0
2.5
Temporal lag (s)

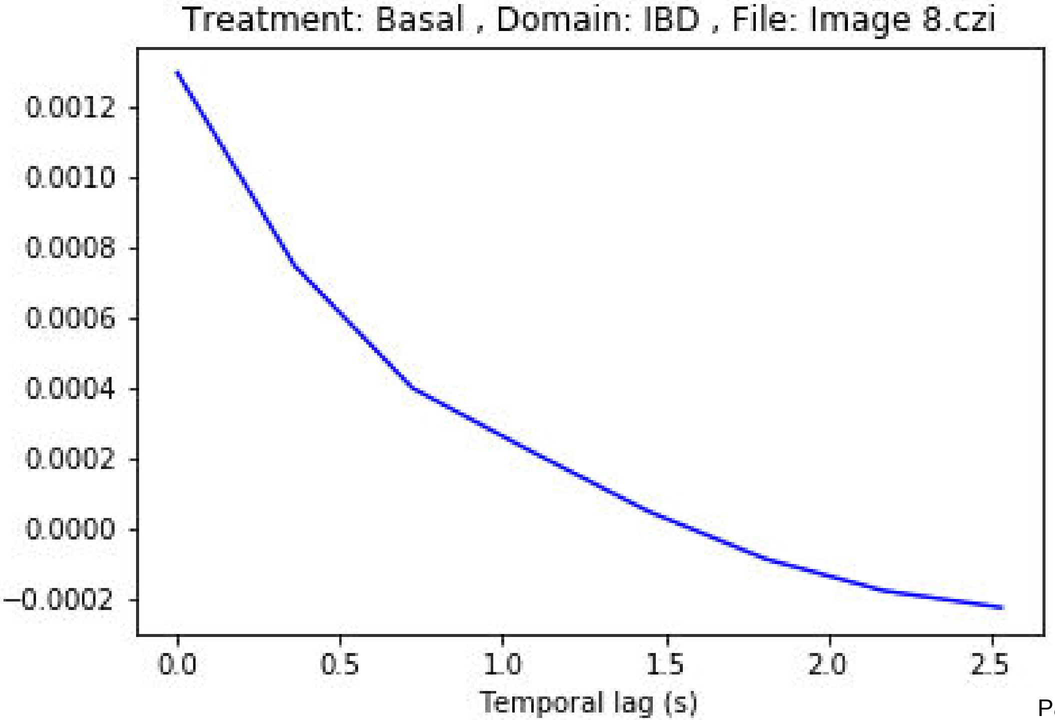

Treatment: Basal , Domain: IBD , File: Image 8.czi
0.0012
0.0010
0.0008
0.0006
0.0004
0.0002
0.0000
−0.0002
0.0
0.5
1.0
1.5
2.0
2.5
Temporal lag (s)

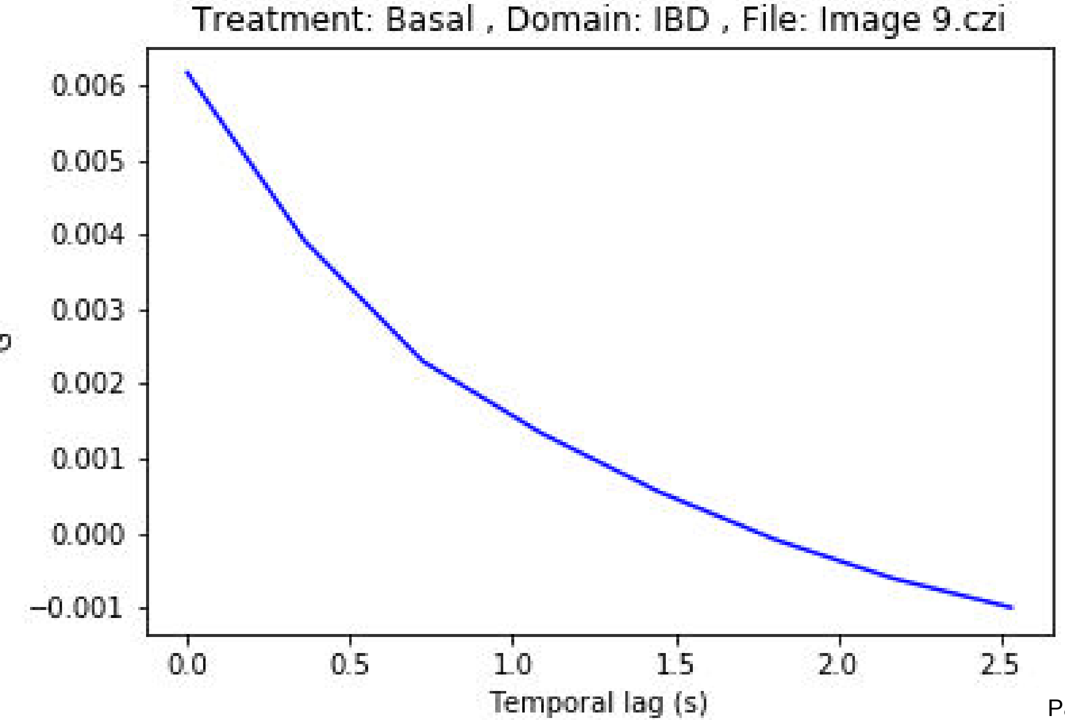
Treatment: Basal , Domain: IBD , File: Image 9.czi
0.006
0.005
0.004
0.003
0.002
0.001
0.000
−0.001
0.0
0.5
1.0
1.5
2.0
2.5
Temporal lag (s)

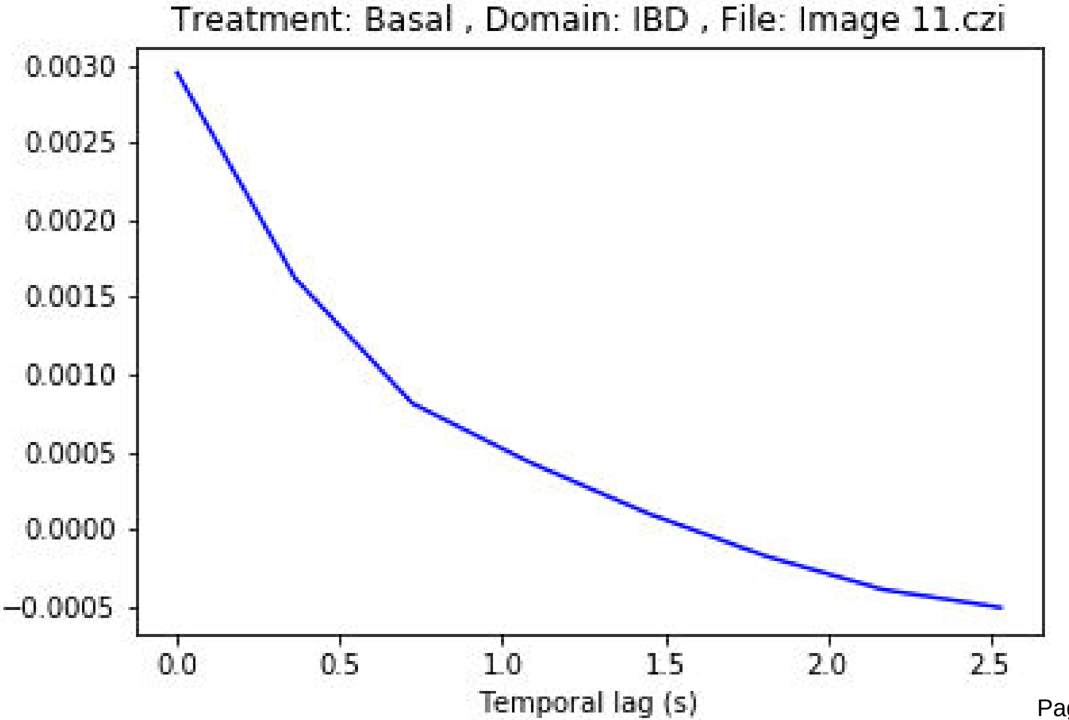
Treatment: Basal , Domain: IBD , File: Image 11.czi
0.0030
0.0025
0.0020
0.0015
0.0010
0.0005
0.0000
−0.0005
0.0
0.5
1.0
1.5
2.0
2.5
Temporal lag (s)

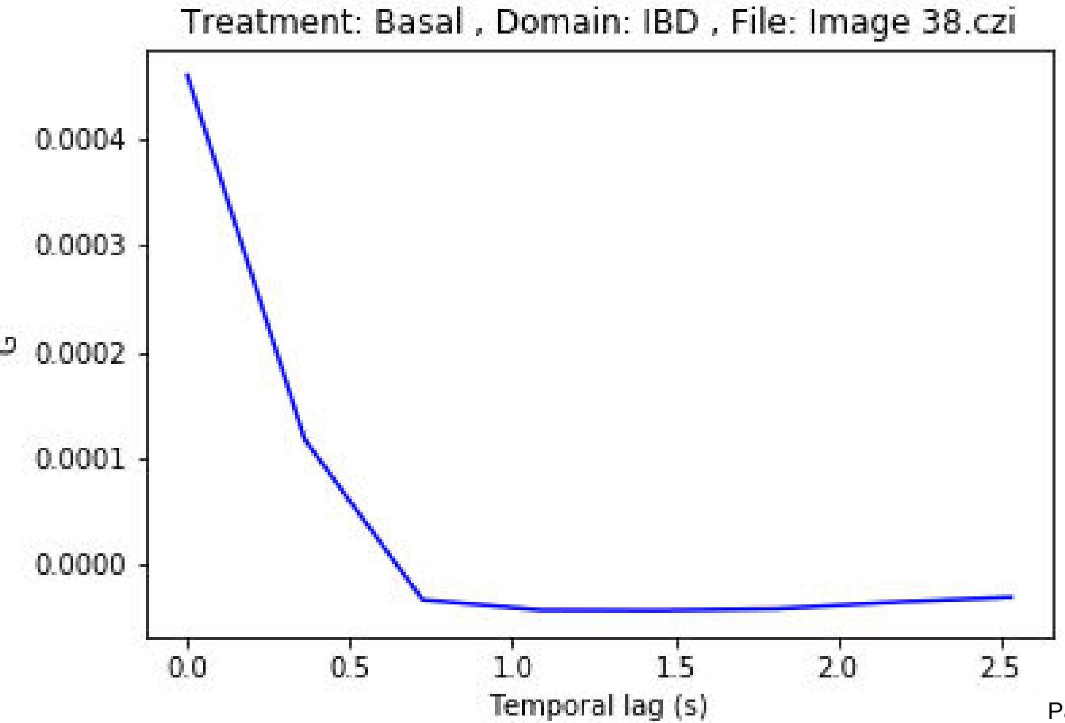
Treatment: Basal , Domain: IBD , File: Image 38.czi
0.0004
0.0003
0.0002
0.0001
0.0000
0.0
0.5
1.0
1.5
2.0
2.5
Temporal lag (s)

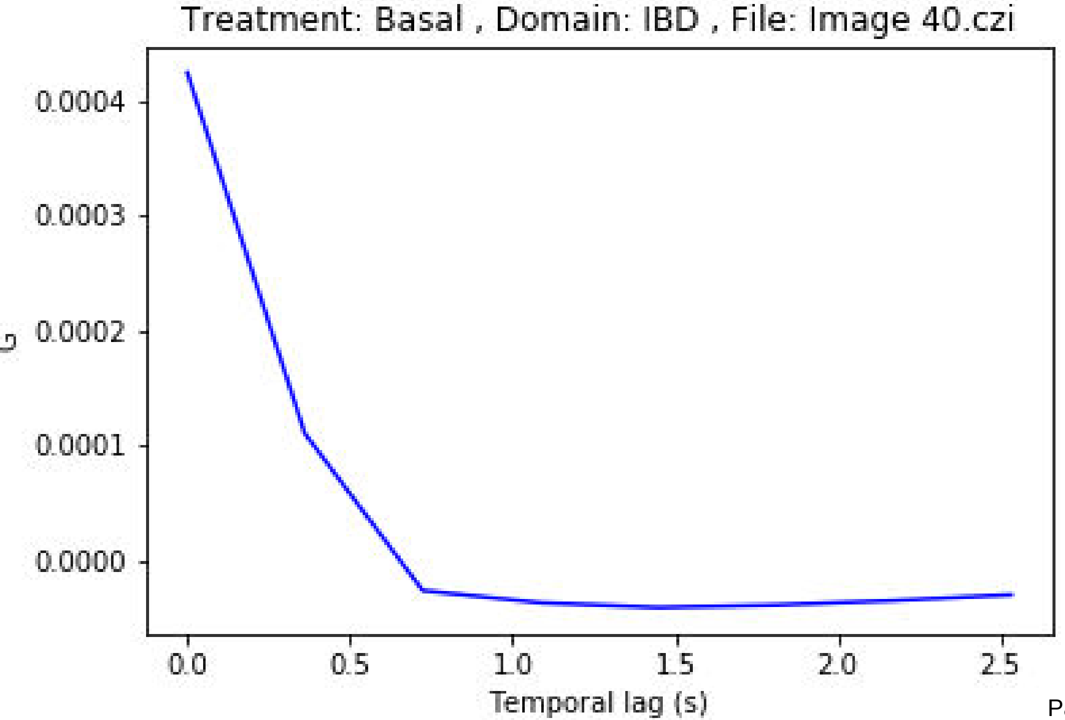

Treatment: Basal , Domain: IBD , File: Image 40.czi
G
0.0004
0.0003
0.0002
0.0001
0.0000
0.0
0.5
1.0
1.5
2.0
2.5
Temporal lag (s)

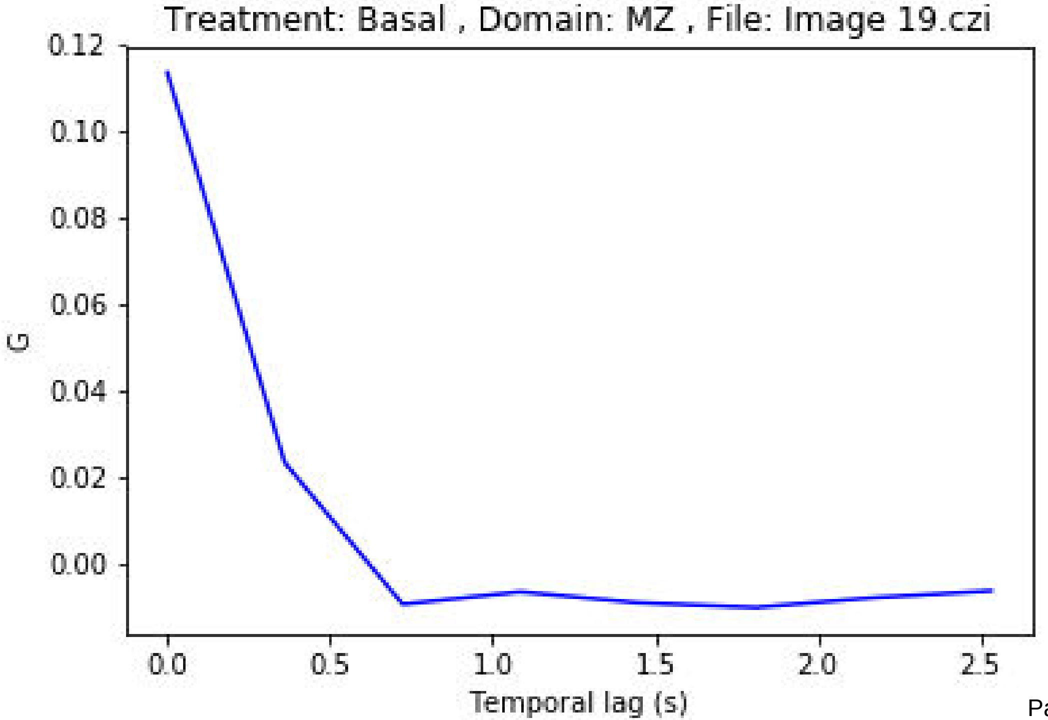
Treatment: Basal , Domain: MZ , File: Image 19.czi
0.12
0.10
0.08
0.06
0.04
0.02
0.00
G
0.0
0.5
1.0
1.5
2.0
2.5
Temporal lag (s)

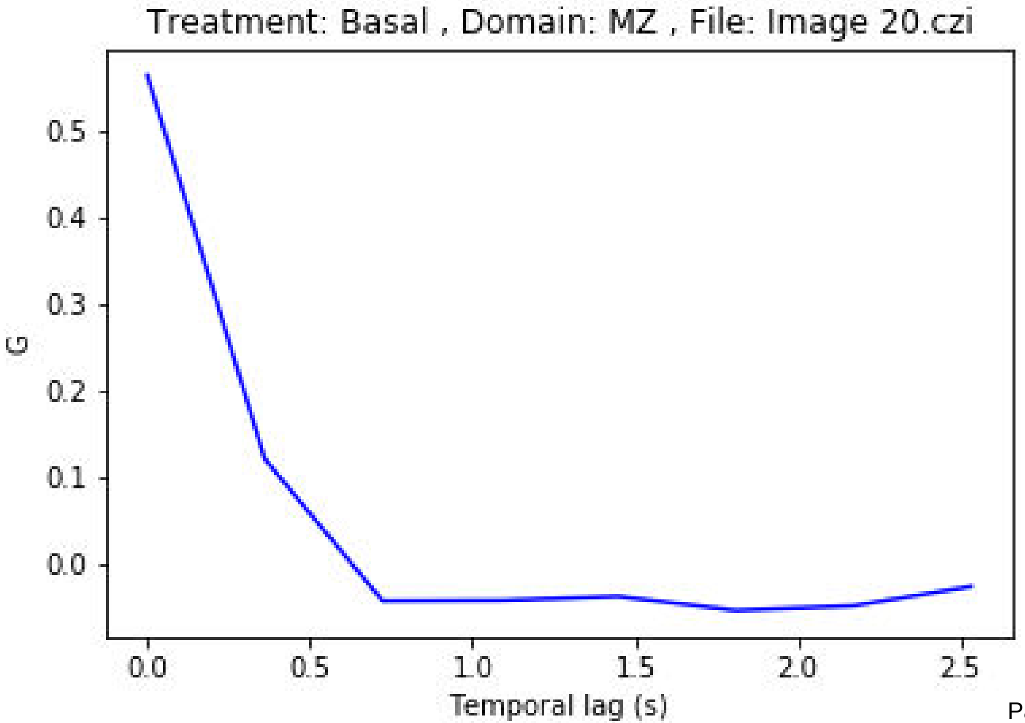

Treatment: Basal , Domain: MZ , File: Image 20.czi
G
0.5
0.4
0.3
0.2
0.1
0.0
0.0
0.5
1.0
1.5
2.0
2.5
Temporal lag (s)

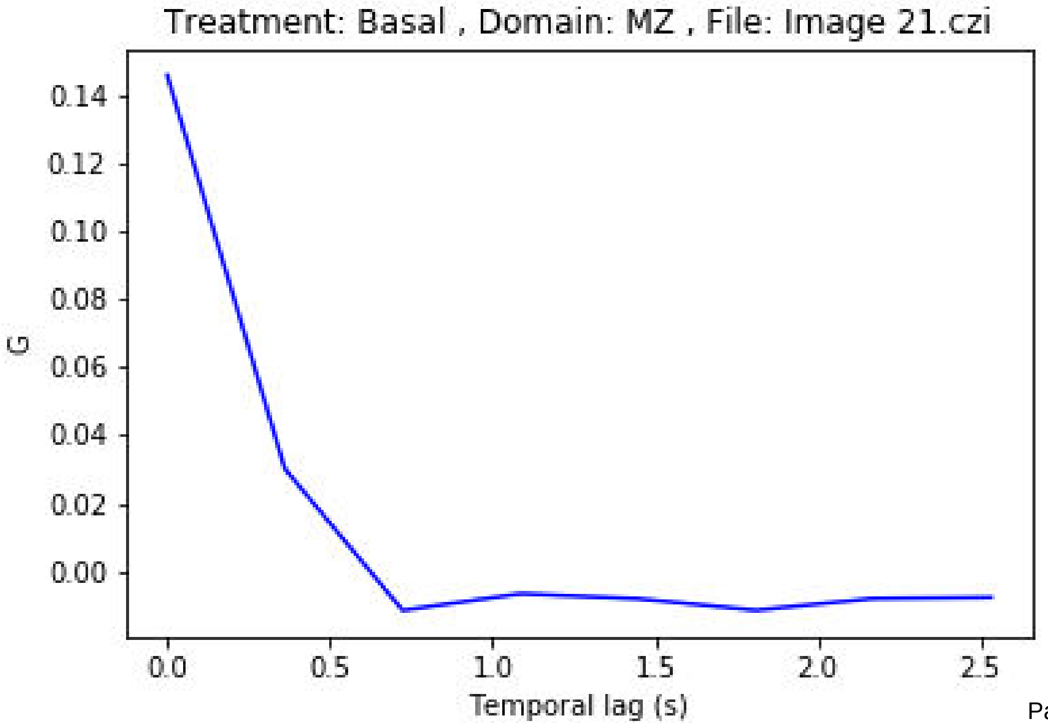
Treatment: Basal , Domain: MZ , File: Image 21.czi
0.14
0.12
0.10
0.08
0.06
0.04
0.02
0.00
G
0.0
0.5
1.0
1.5
2.0
2.5
Temporal lag (s)

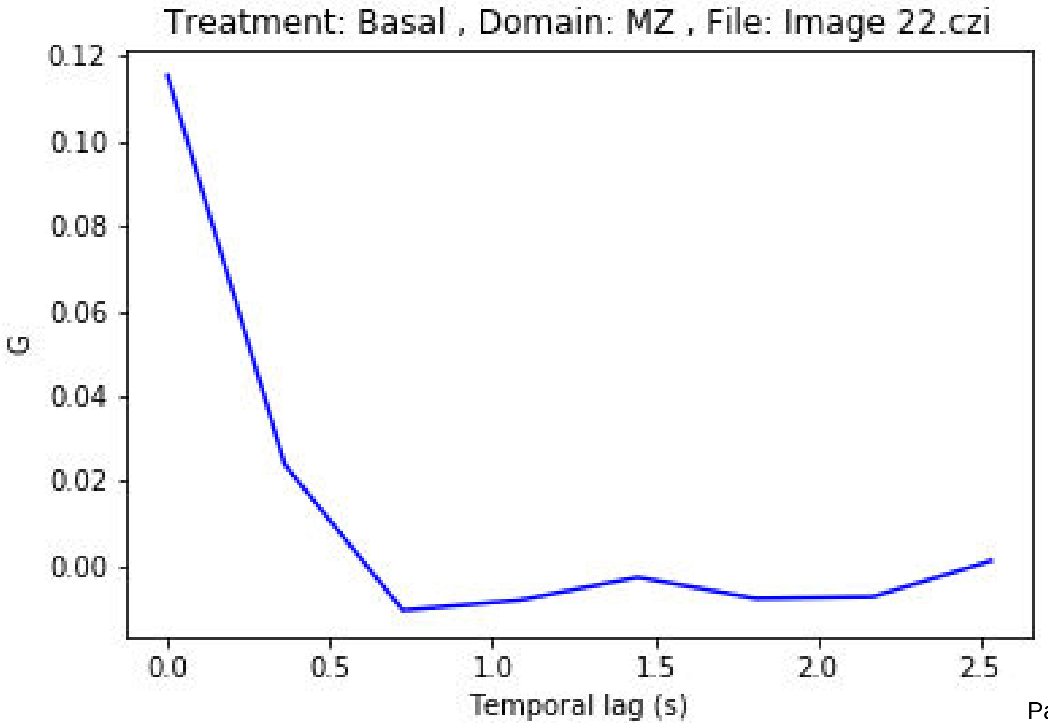
Treatment: Basal , Domain: MZ , File: Image 22.czi
0.12
0.10
0.08
0.06
0.04
0.02
0.00
G
0.0
0.5
1.0
1.5
2.0
2.5
Temporal lag (s)

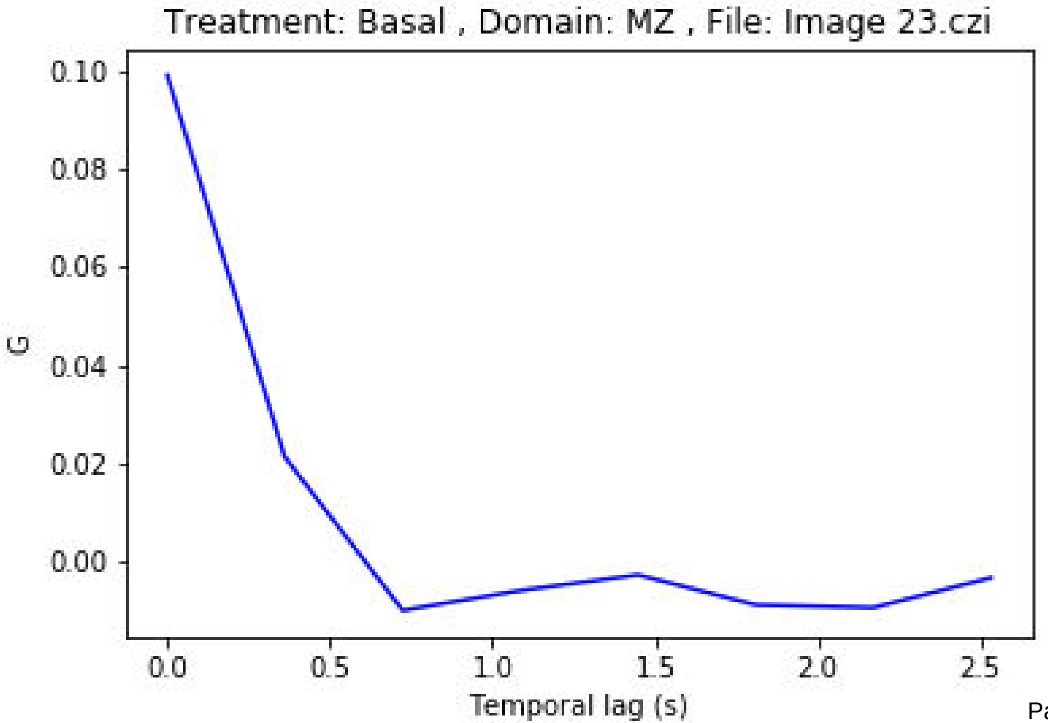
Treatment: Basal , Domain: MZ , File: Image 23.czi
0.10
0.08
0.06
0.04
0.02
0.00
G
0.0
0.5
1.0
1.5
2.0
2.5
Temporal lag (s)

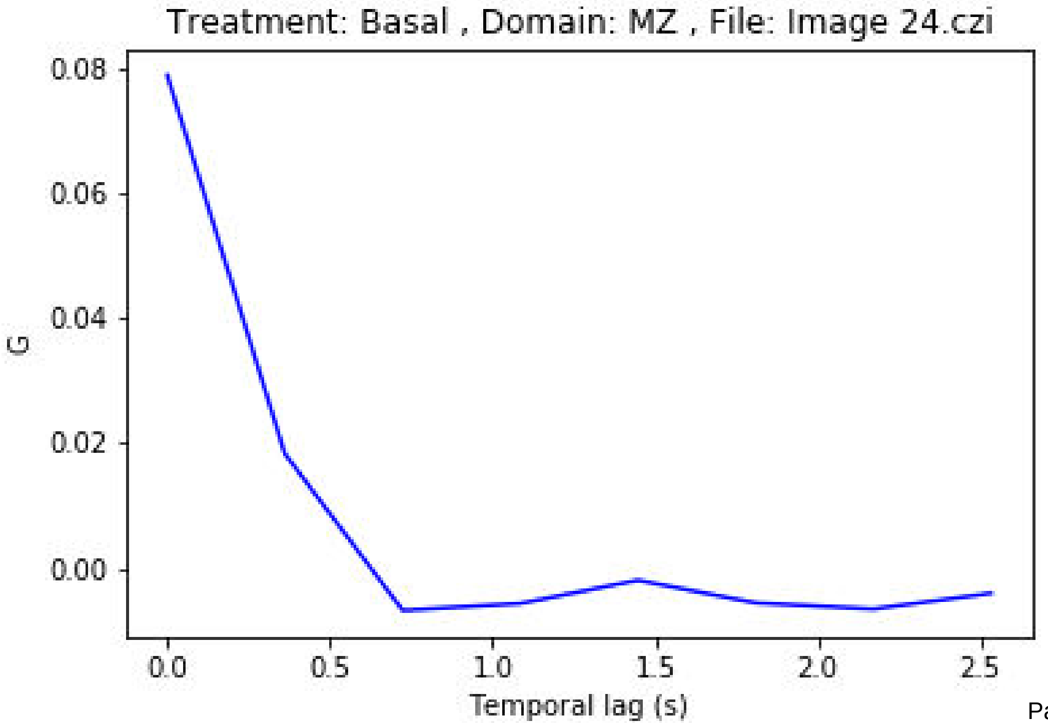
Treatment: Basal , Domain: MZ , File: Image 24.czi
0.08
0.06
0.04
0.02
0.00
G
0.0
0.5
1.0
1.5
2.0
2.5
Temporal lag (s)

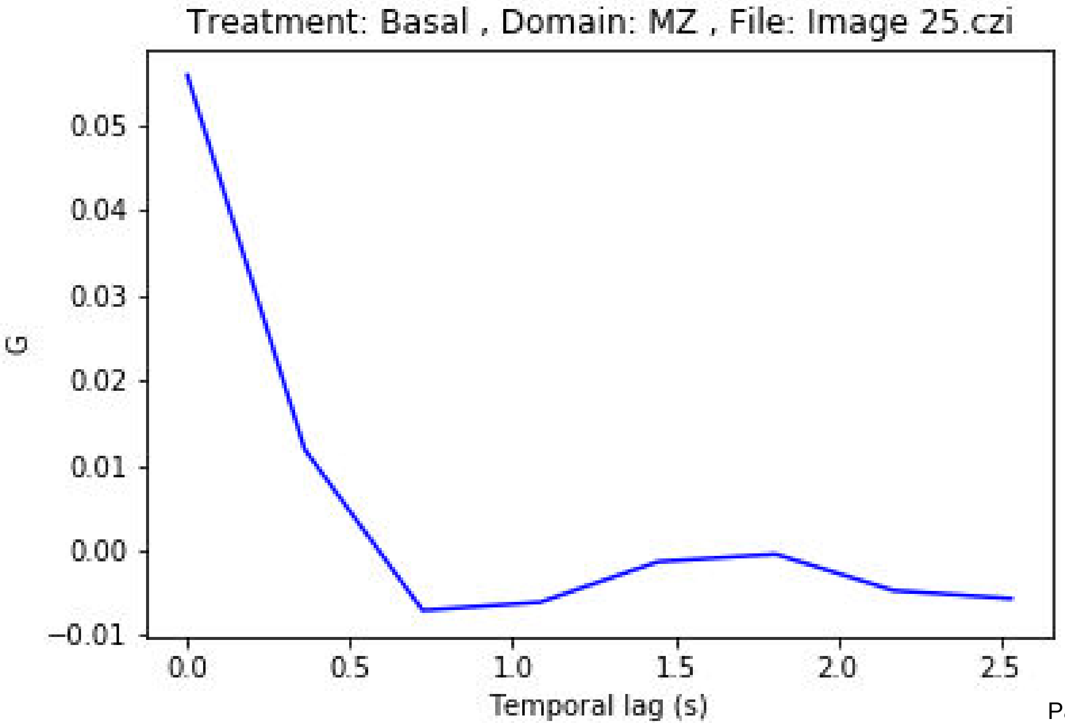
Treatment: Basal , Domain: MZ , File: Image 25.czi
0.05
0.04
0.03
0.02
0.01
0.00
−0.01
G
0.0
0.5
1.0
1.5
2.0
2.5
Temporal lag (s)

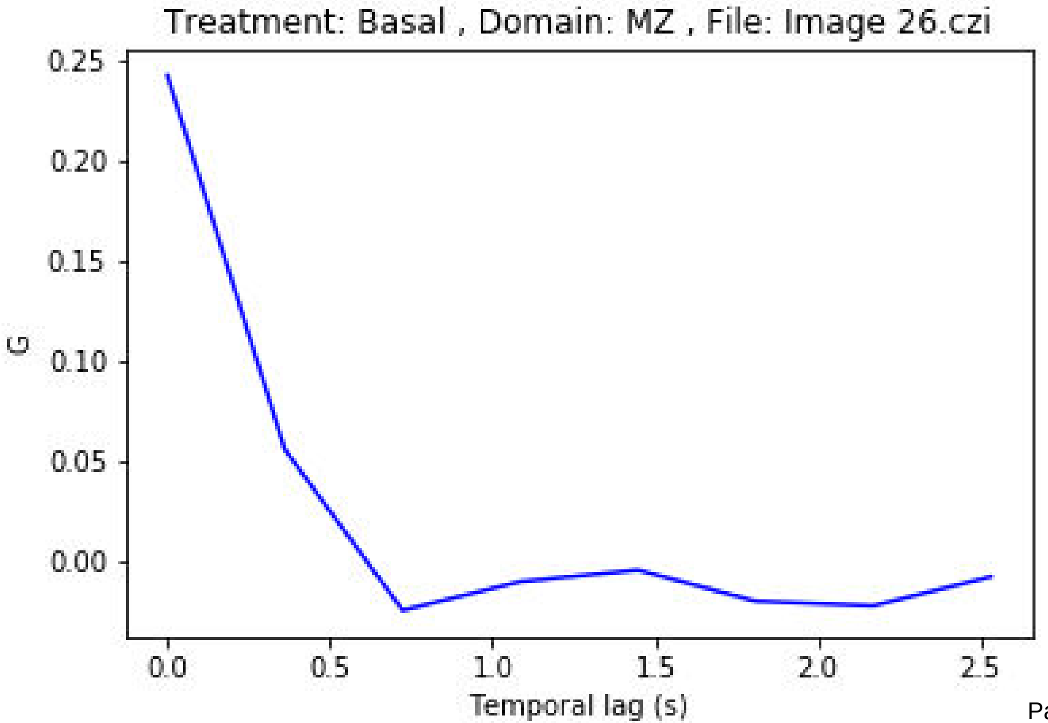

Treatment: Basal , Domain: MZ , File: Image 26.czi
G
0.25
0.20
0.15
0.10
0.05
0.00
0.0
0.5
1.0
1.5
2.0
2.5
Temporal lag (s)

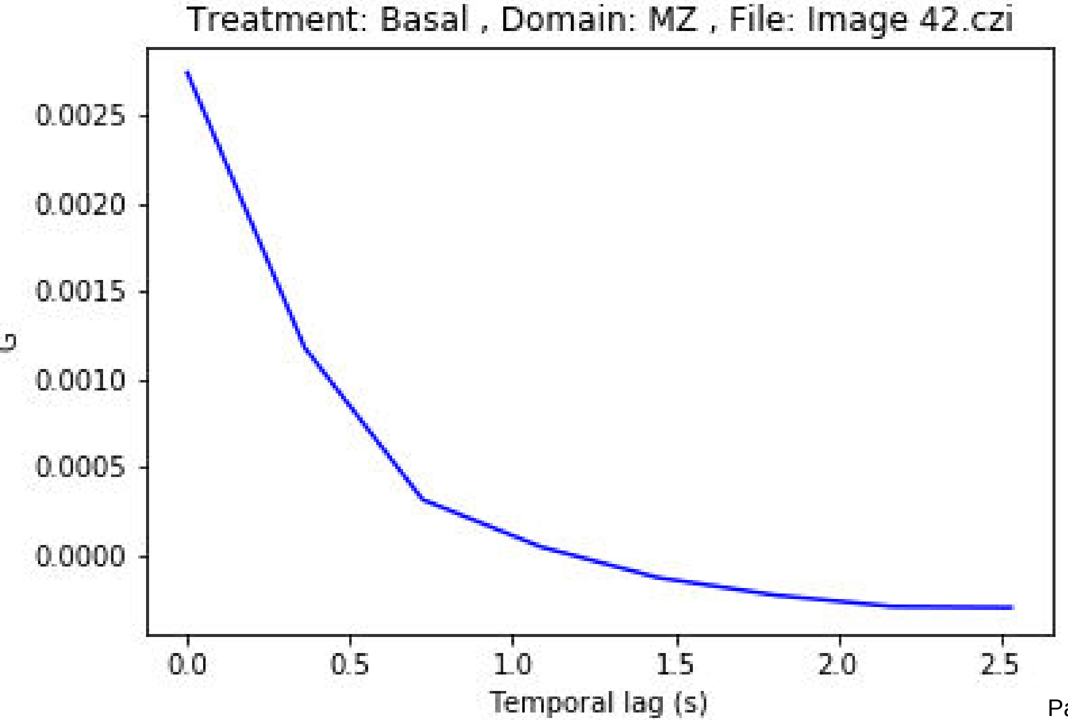

Treatment: Basal , Domain: MZ , File: Image 42.czi
0.0025
0.0020
0.0015
0.0010
0.0005
0.0000
G
0.0
0.5
1.0
1.5
2.0
2.5
Temporal lag (s)

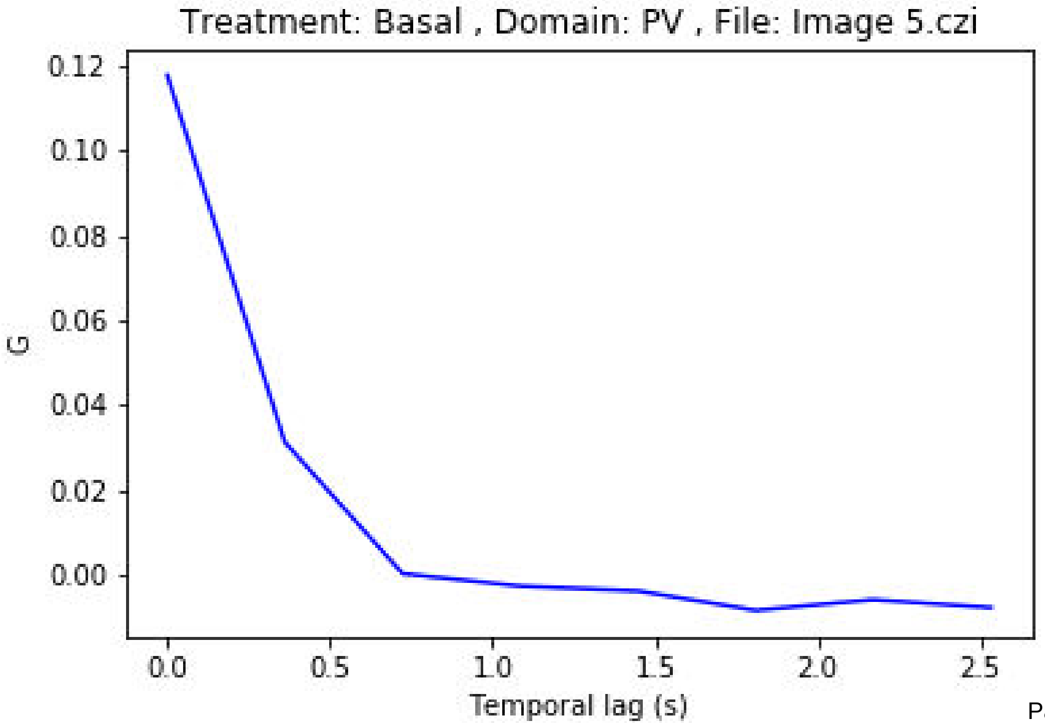
Treatment: Basal , Domain: PV , File: Image 5.czi
0.12
0.10
0.08
0.06
0.04
0.02
0.00
G
0.0
0.5
1.0
1.5
2.0
2.5
Temporal lag (s)

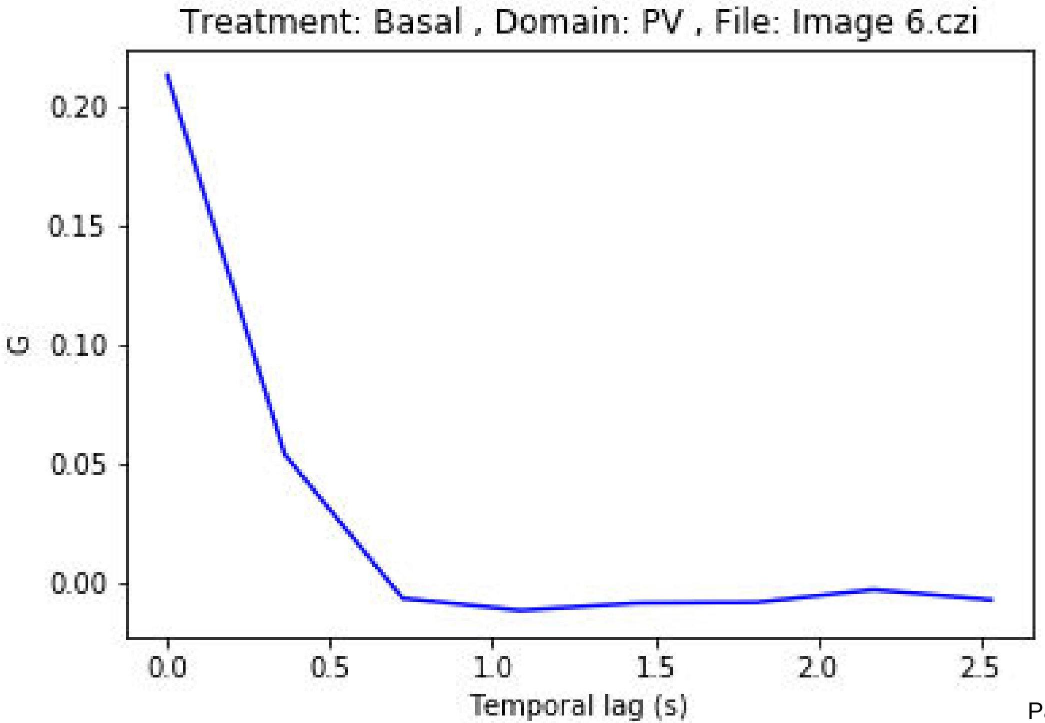

Treatment: Basal , Domain: PV , File: Image 6.czi
G
0.20
0.15
0.10
0.05
0.00
0.0
0.5
1.0
1.5
2.0
2.5
Temporal lag (s)

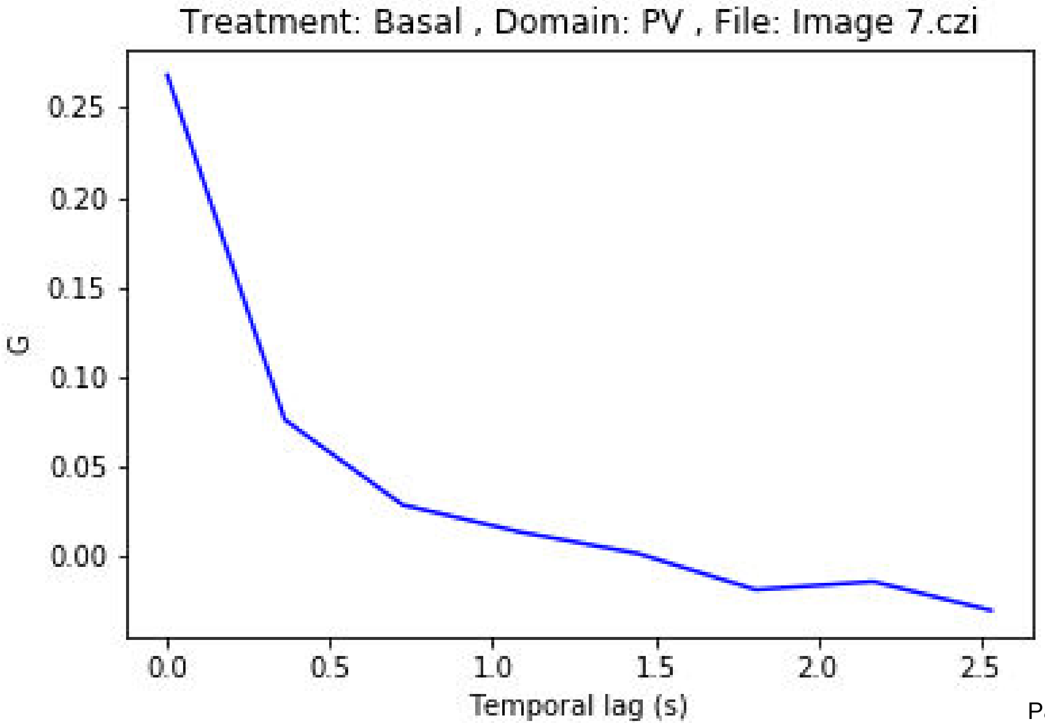
Treatment: Basal , Domain: PV , File: Image 7.czi
G
0.25
0.20
0.15
0.10
0.05
0.00
0.0
0.5
1.0
1.5
2.0
2.5
Temporal lag (s)

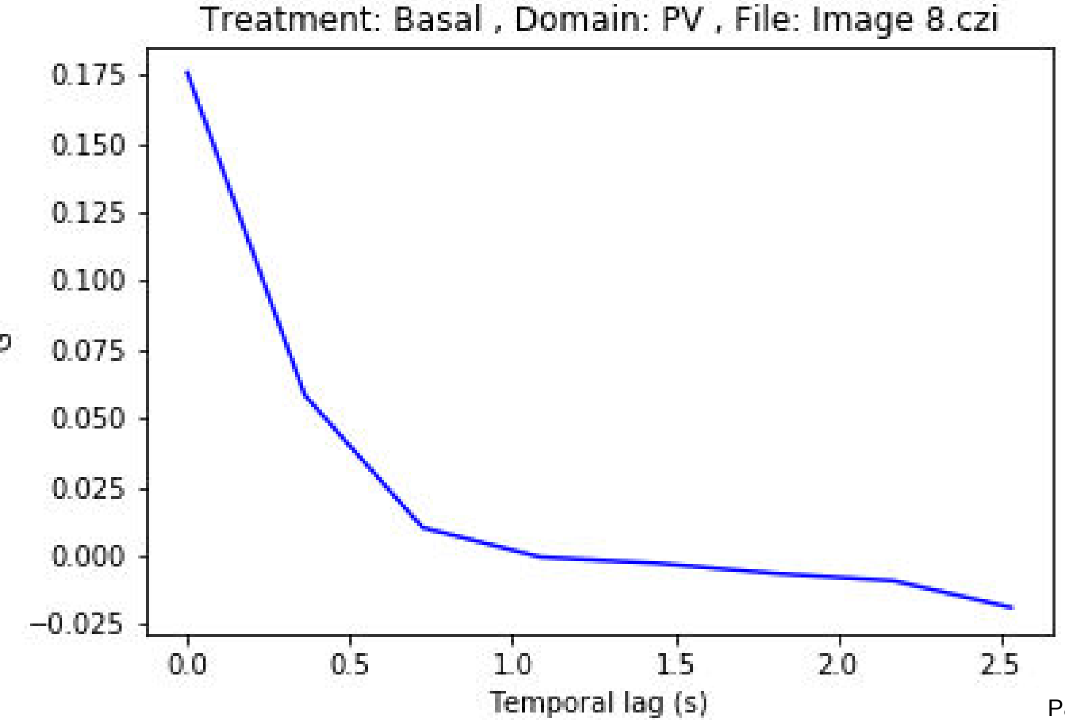
Treatment: Basal , Domain: PV , File: Image 8.czi
0.175
0.150
0.125
0.100
0.075
0.050
0.025
0.000
−0.025
0.0
0.5
1.0
1.5
2.0
2.5
Temporal lag (s)

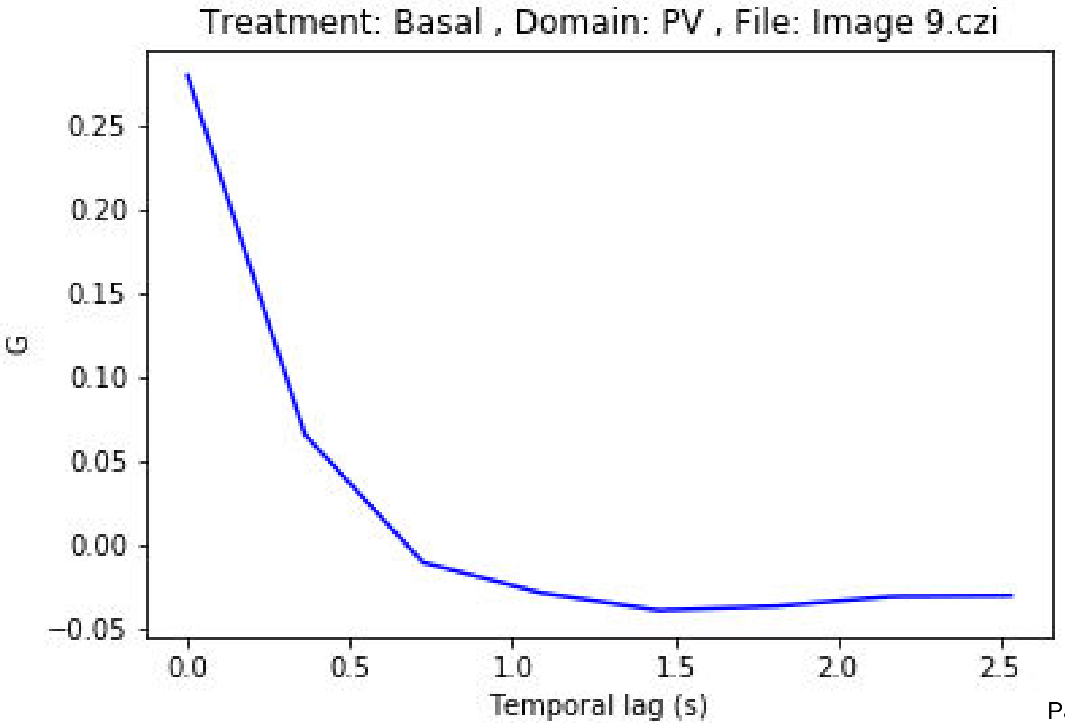

Treatment: Basal , Domain: PV , File: Image 9.czi
G
0.25
0.20
0.15
0.10
0.05
0.00
−0.05
0.0
0.5
1.0
1.5
2.0
2.5
Temporal lag (s)

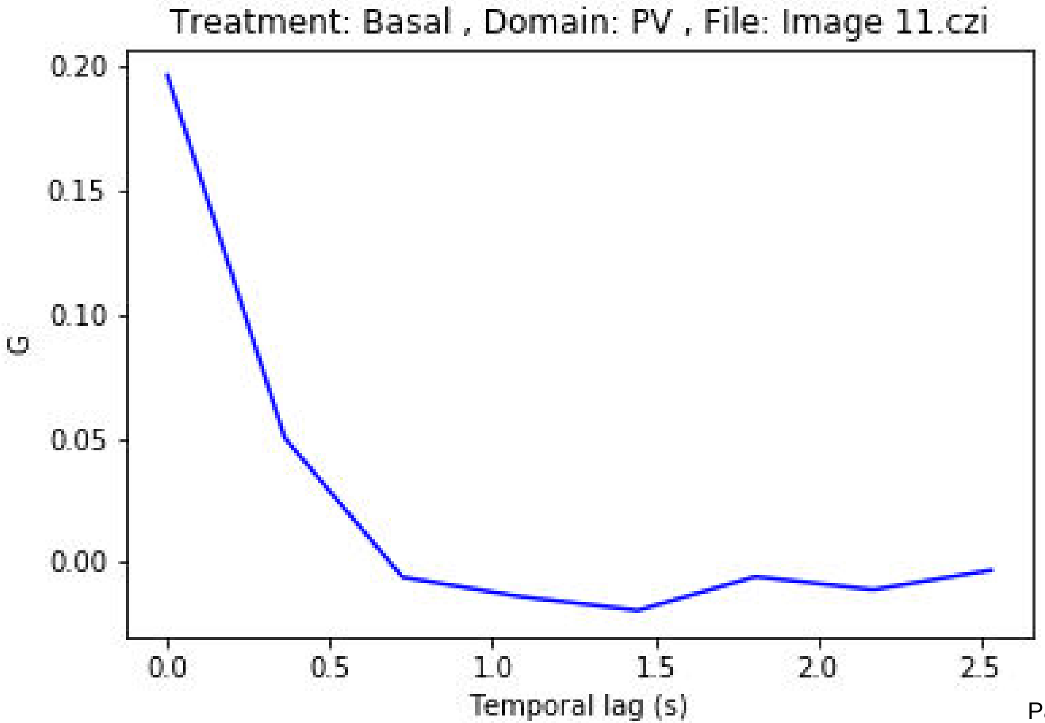
Treatment: Basal , Domain: PV , File: Image 11.czi
0.20
0.15
0.10
0.05
0.00
G
0.0
0.5
1.0
1.5
2.0
2.5
Temporal lag (s)

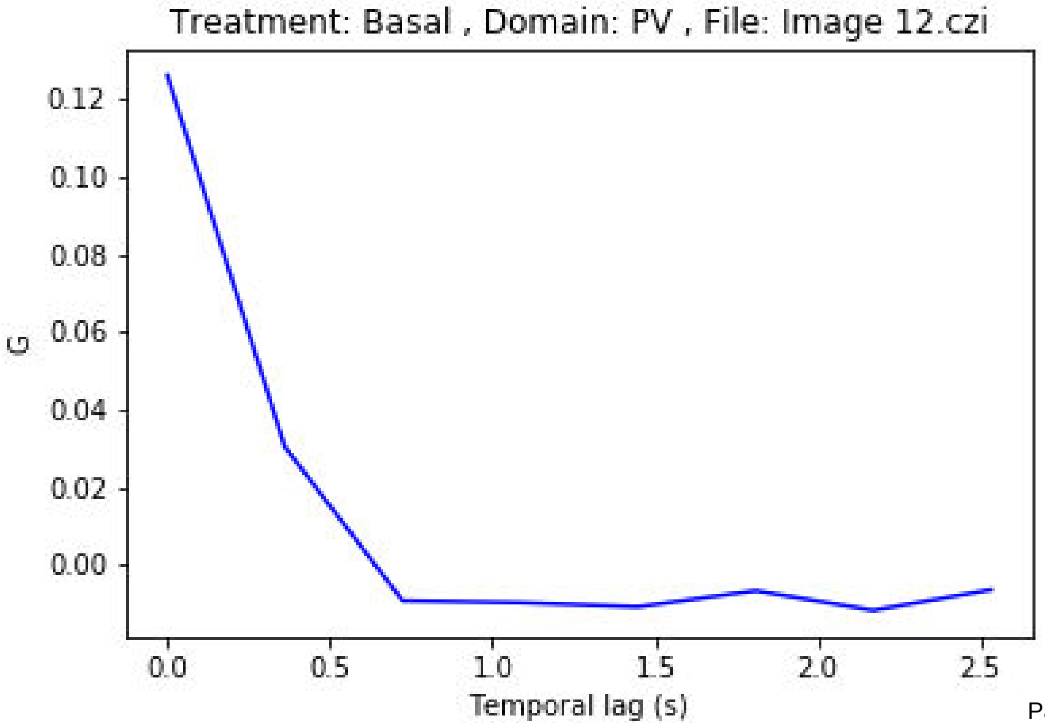
Treatment: Basal , Domain: PV , File: Image 12.czi
G
0.12
0.10
0.08
0.06
0.04
0.02
0.00
0.0
0.5
1.0
1.5
2.0
2.5
Temporal lag (s)

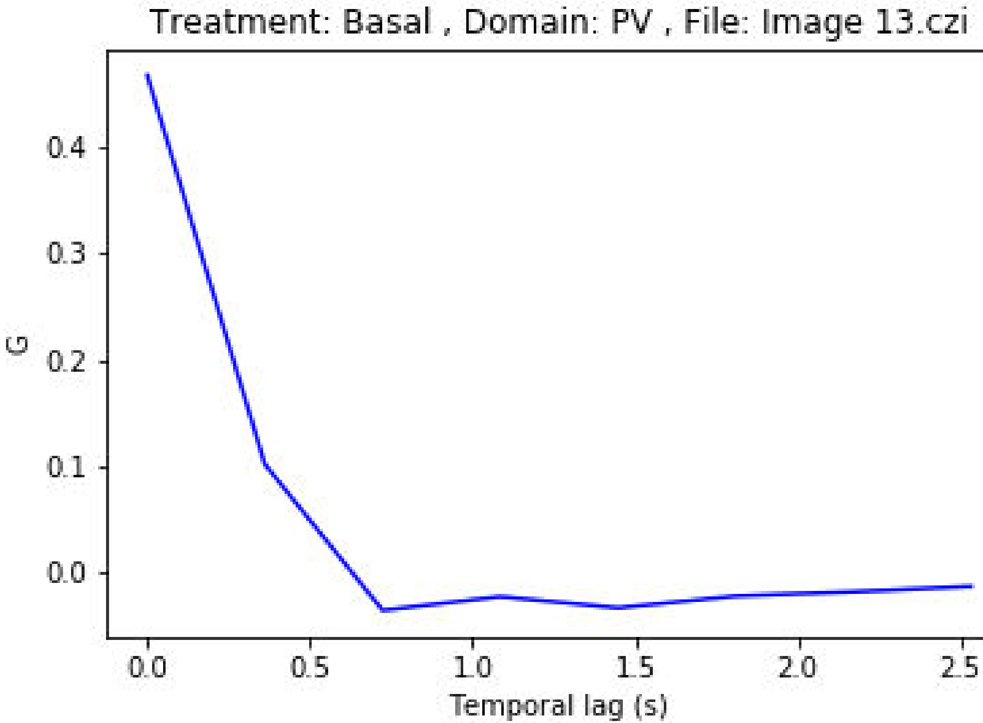
Treatment: Basal , Domain: PV , File: Image 13.czi
G
0.4
0.3
0.2
0.1
0.0
0.0
0.5
1.0
1.5
2.0
2.5
Temporal lag (s)

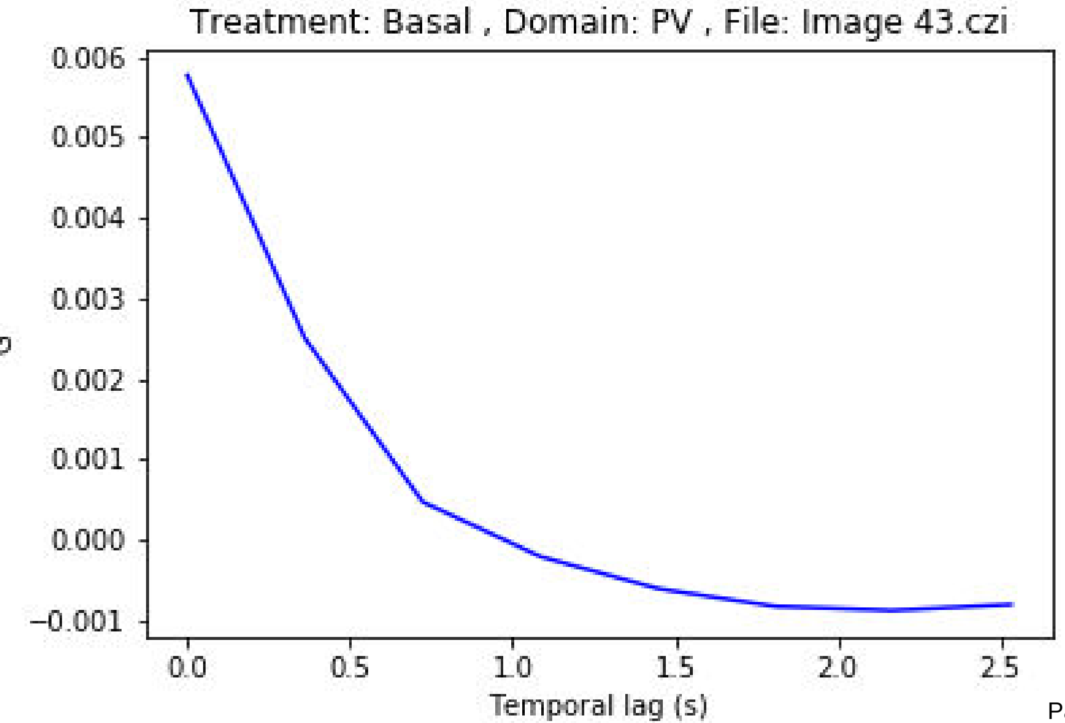
Treatment: Basal , Domain: PV , File: Image 43.czi
0.006
0.005
0.004
0.003
0.002
0.001
0.000
−0.001
0.0
0.5
1.0
1.5
2.0
2.5
Temporal lag (s)

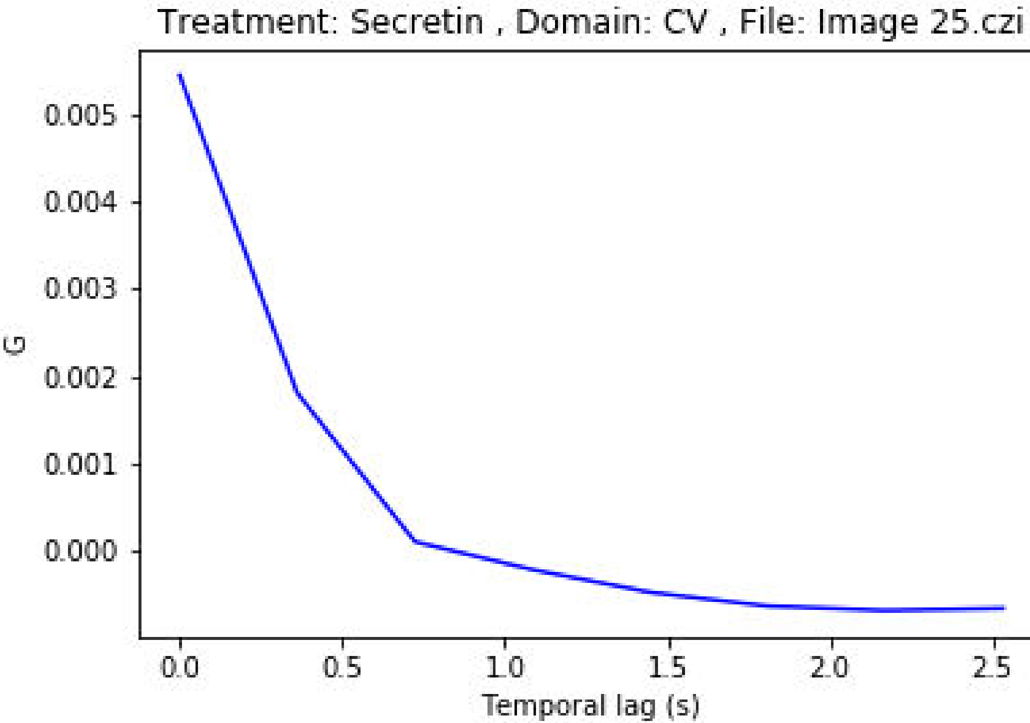
Treatment: Secretin , Domain: CV , File: Image 25.czi
0.005
0.004
0.003
0.002
0.001
0.000
G
0.0
0.5
1.0
1.5
2.0
2.5
Temporal lag (s)

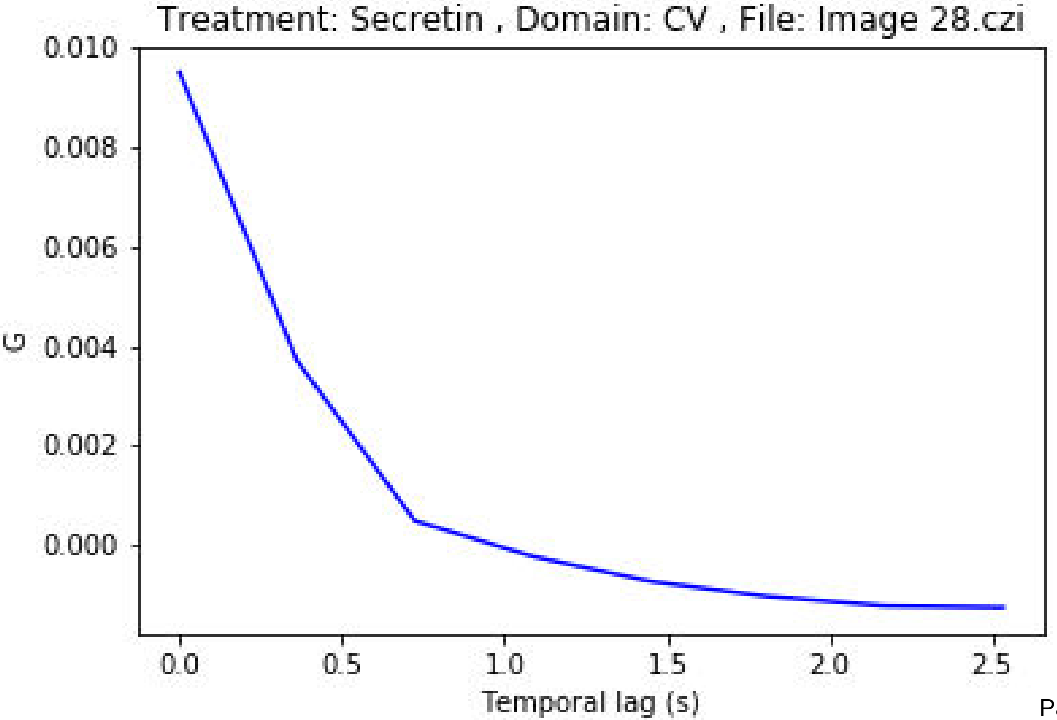
Treatment: Secretin , Domain: CV , File: Image 28.czi
0.010
0.008
0.006
0.004
0.002
0.000
G
0.0
0.5
1.0
1.5
2.0
2.5
Temporal lag (s)

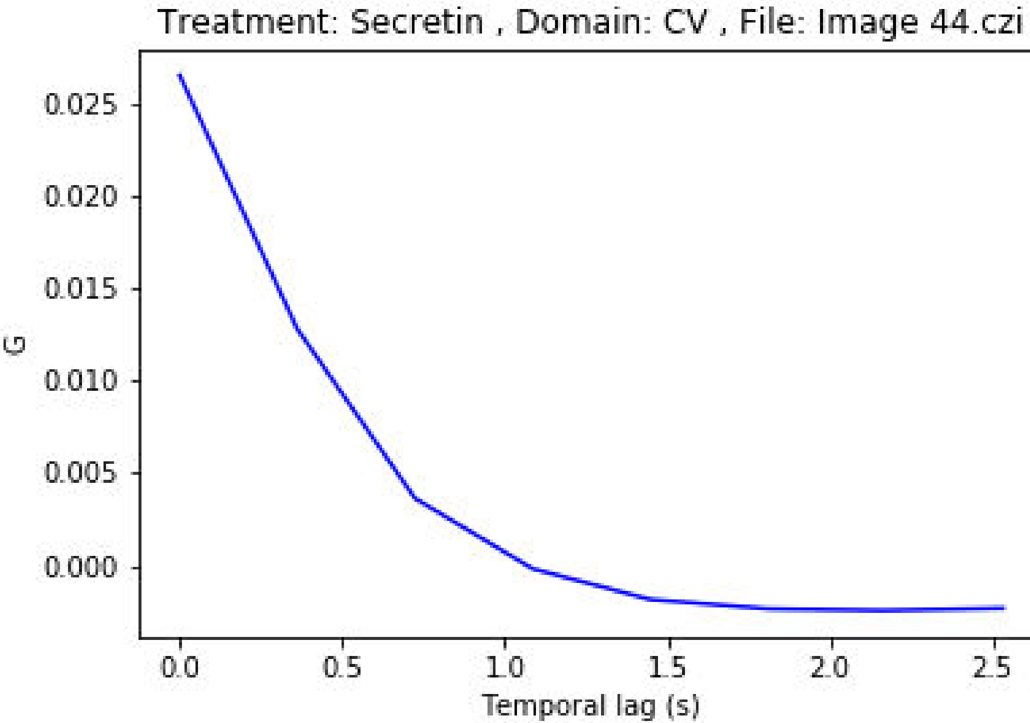
Treatment: Secretin , Domain: CV , File: Image 44.czi
0.025
0.020
0.015
0.010
0.005
0.000
G
0.0
0.5
1.0
1.5
2.0
2.5
Temporal lag (s)

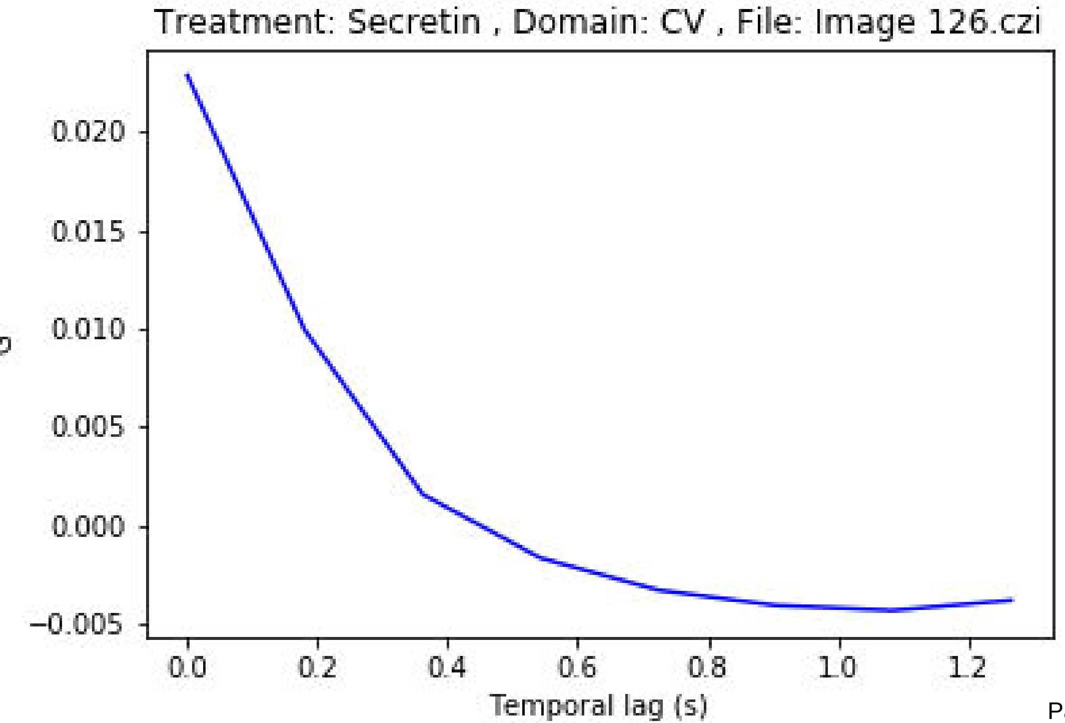

Treatment: Secretin , Domain: CV , File: Image 126.czi
0.020
0.015
0.010
0.005
0.000
−0.005
0.0
0.2
0.4
0.6
0.8
1.0
1.2
Temporal lag (s)

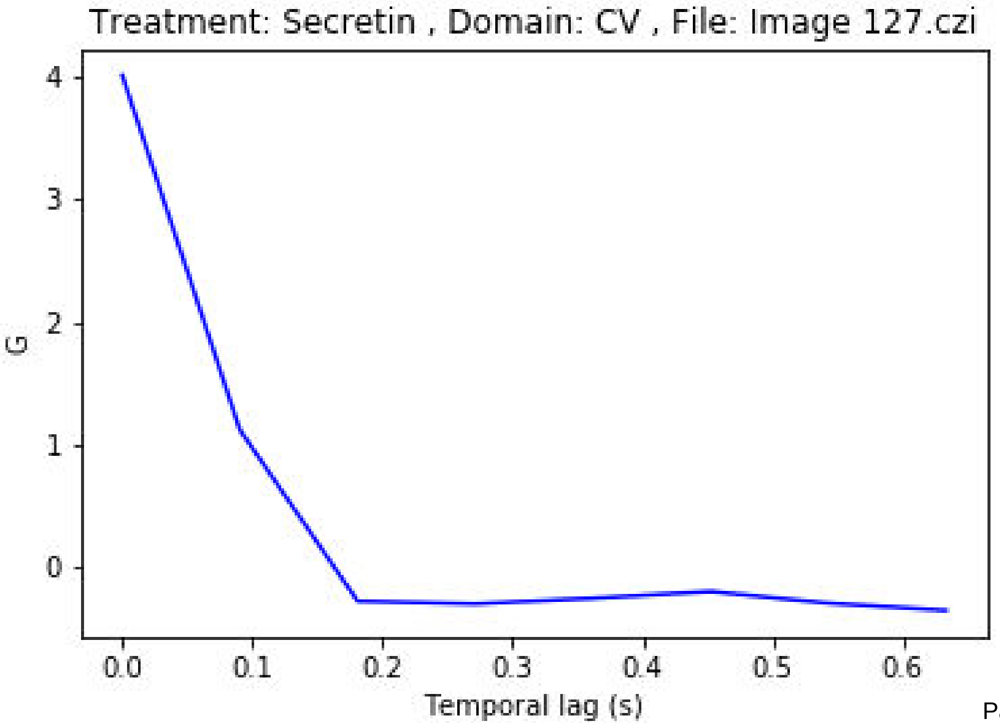
Treatment: Secretin , Domain: CV , File: Image 127.czi
G
4
3
2
1
0
0.0
0.1
0.2
0.3
0.4
0.5
0.6
Temporal lag (s)

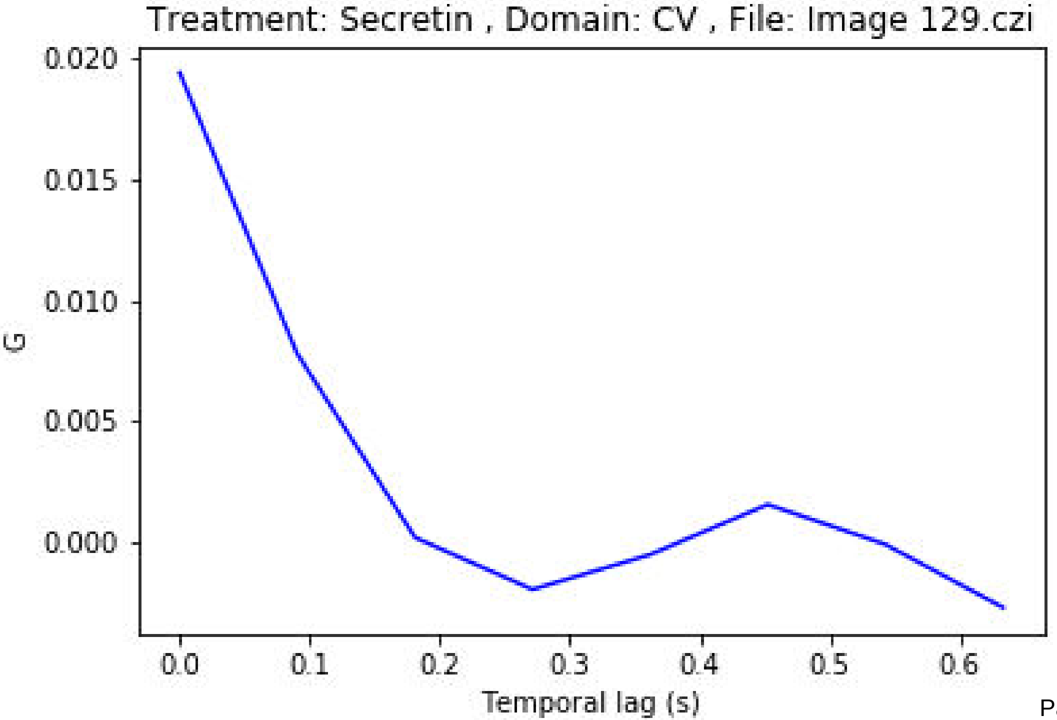
Treatment: Secretin , Domain: CV , File: Image 129.czi
0.020
0.015
0.010
0.005
0.000
G
0.0
0.1
0.2
0.3
0.4
0.5
0.6
Temporal lag (s)

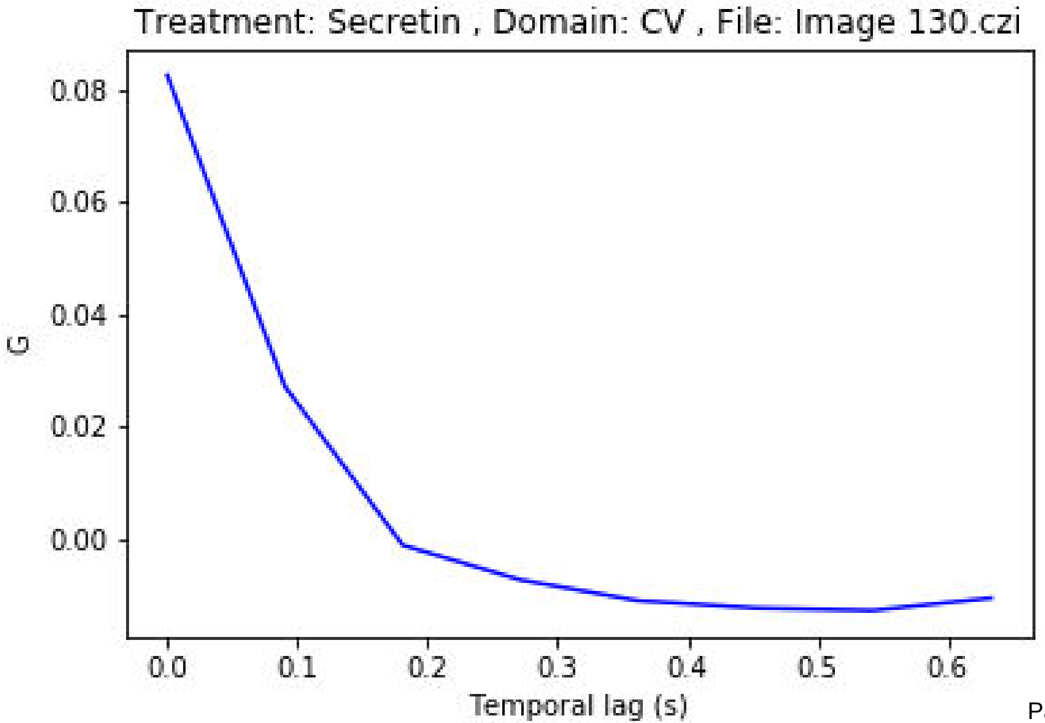

Treatment: Secretin , Domain: CV , File: Image 130.czi
0.08
0.06
0.04
0.02
0.00
G
0.0
0.1
0.2
0.3
0.4
0.5
0.6
Temporal lag (s)

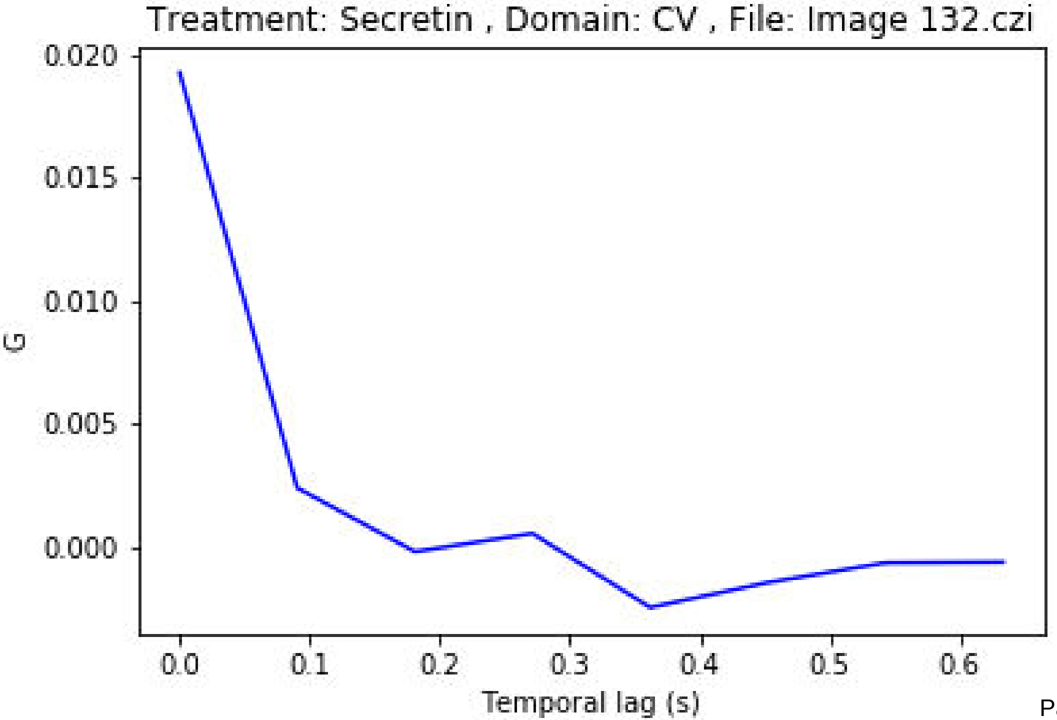
Treatment: Secretin , Domain: CV , File: Image 132.czi
0.020
0.015
0.010
0.005
0.000
G
0.0
0.1
0.2
0.3
0.4
0.5
0.6
Temporal lag (s)

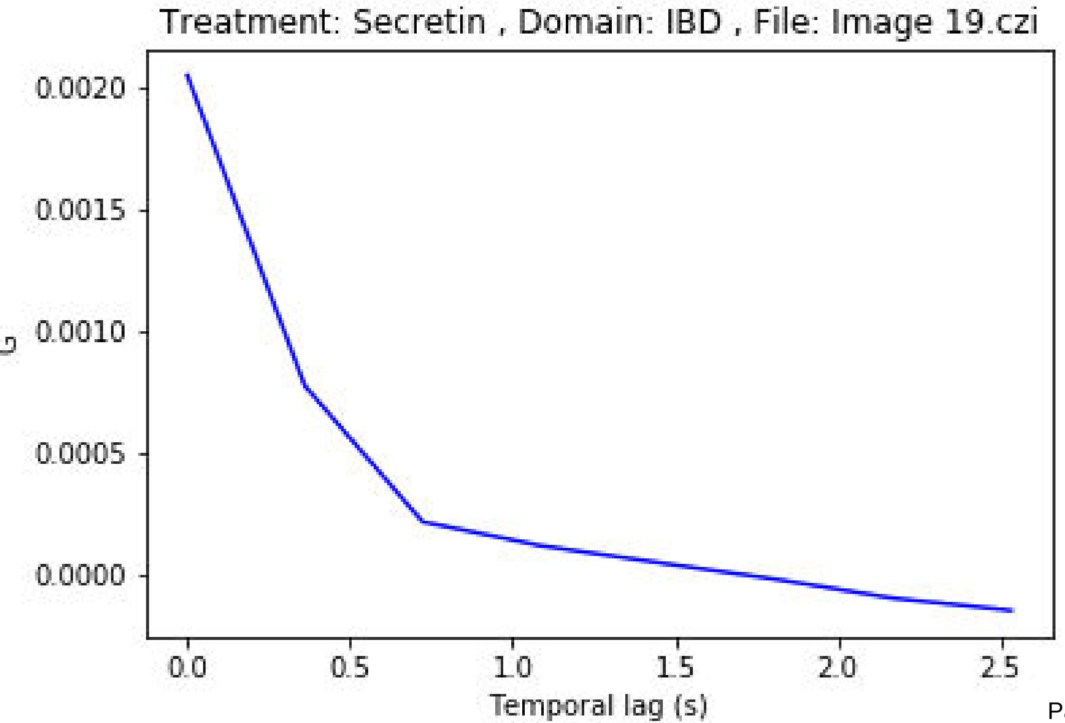

Treatment: Secretin , Domain: IBD , File: Image 19.czi
G
0.0020
0.0015
0.0010
0.0005
0.0000
0.0
0.5
1.0
1.5
2.0
2.5
Temporal lag (s)

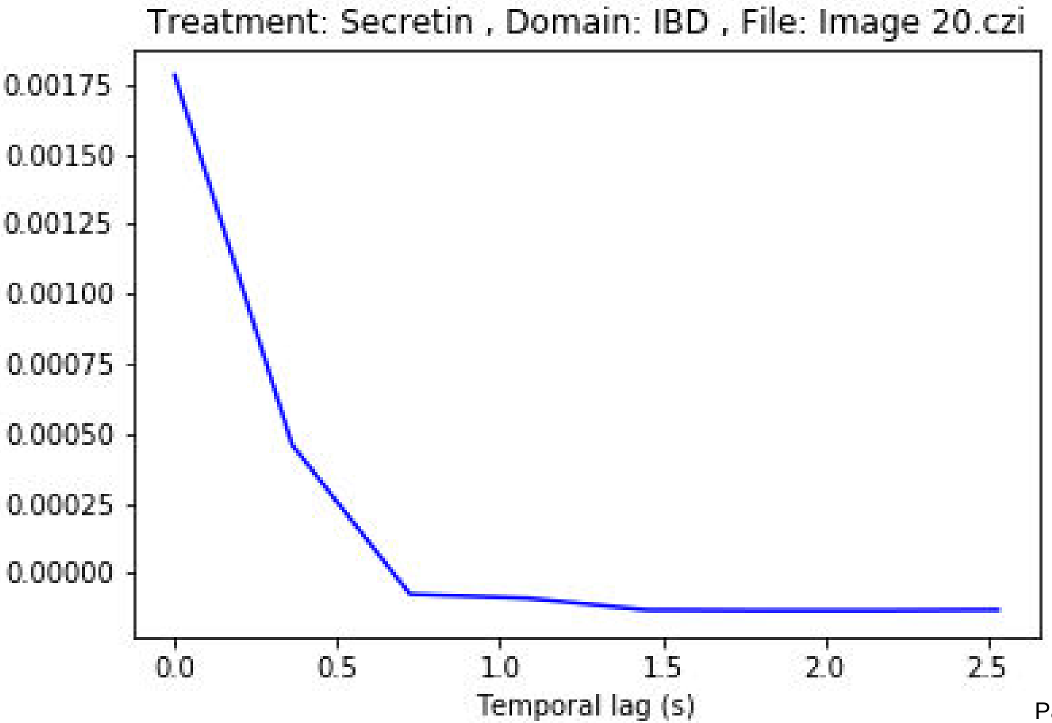
Treatment: Secretin , Domain: IBD , File: Image 20.czi
0.00175
0.00150
0.00125
0.00100
0.00075
0.00050
0.00025
0.00000
0.0
0.5
1.0
1.5
2.0
2.5
Temporal lag (s)

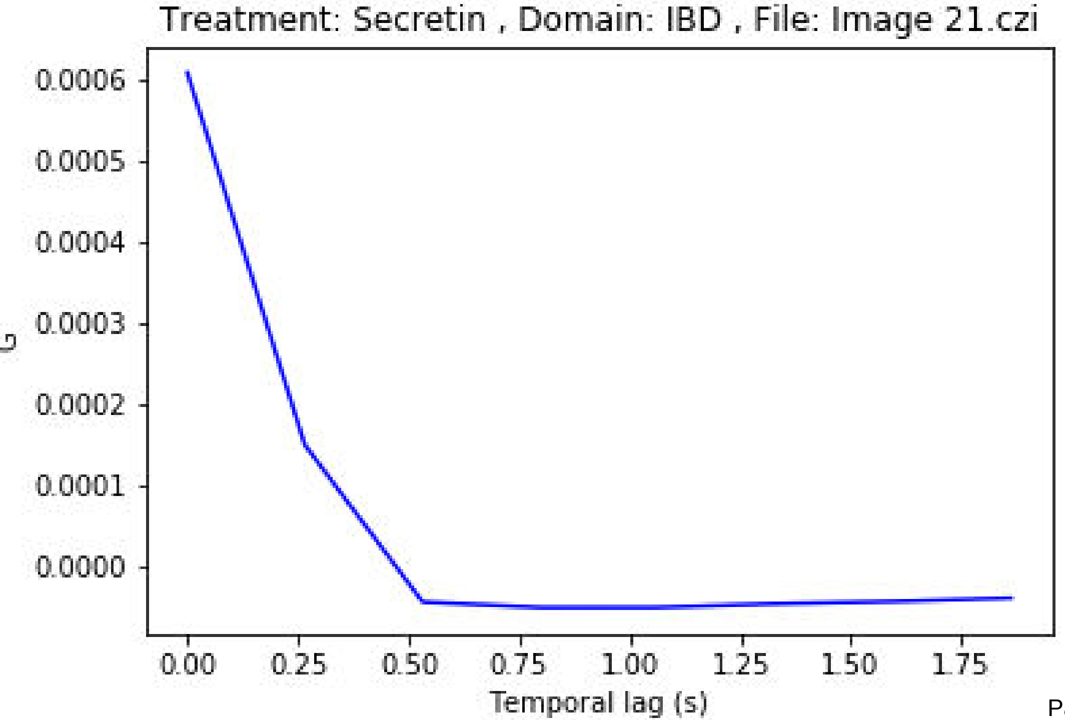
Treatment: Secretin , Domain: IBD , File: Image 21.czi
0.0006
0.0005
0.0004
0.0003
0.0002
0.0001
0.0000
G
0.00
0.25
0.50
0.75
1.00
1.25
1.50
1.75
Temporal lag (s)

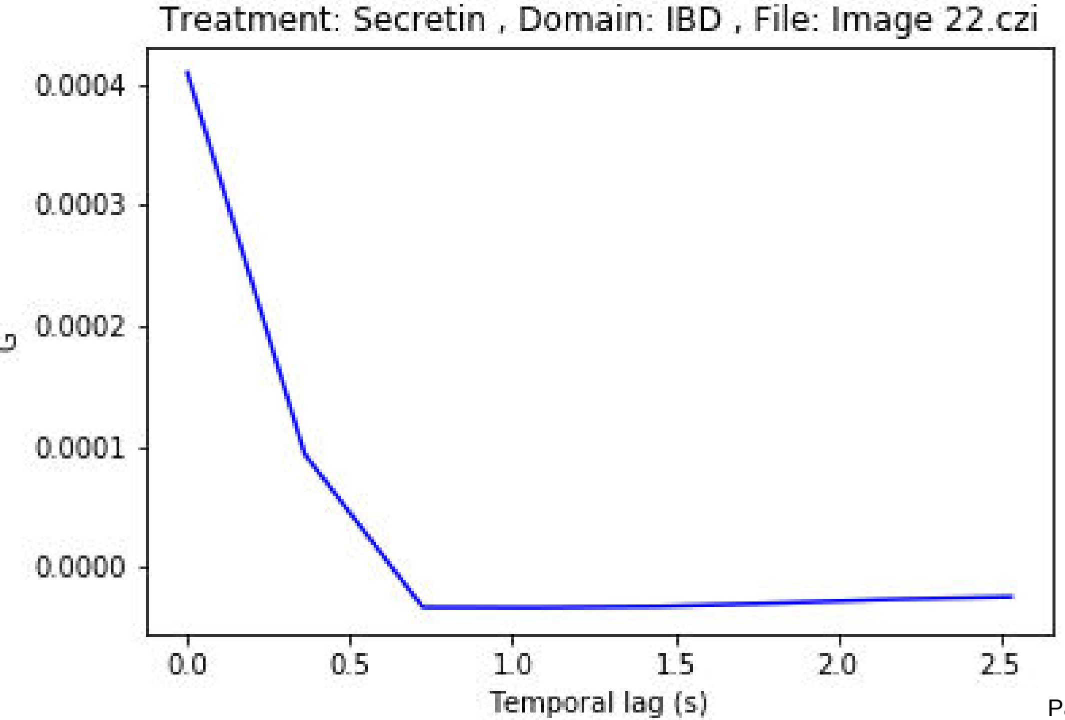

Treatment: Secretin , Domain: IBD , File: Image 22.czi
G
0.0004
0.0003
0.0002
0.0001
0.0000
0.0
0.5
1.0
1.5
2.0
2.5
Temporal lag (s)

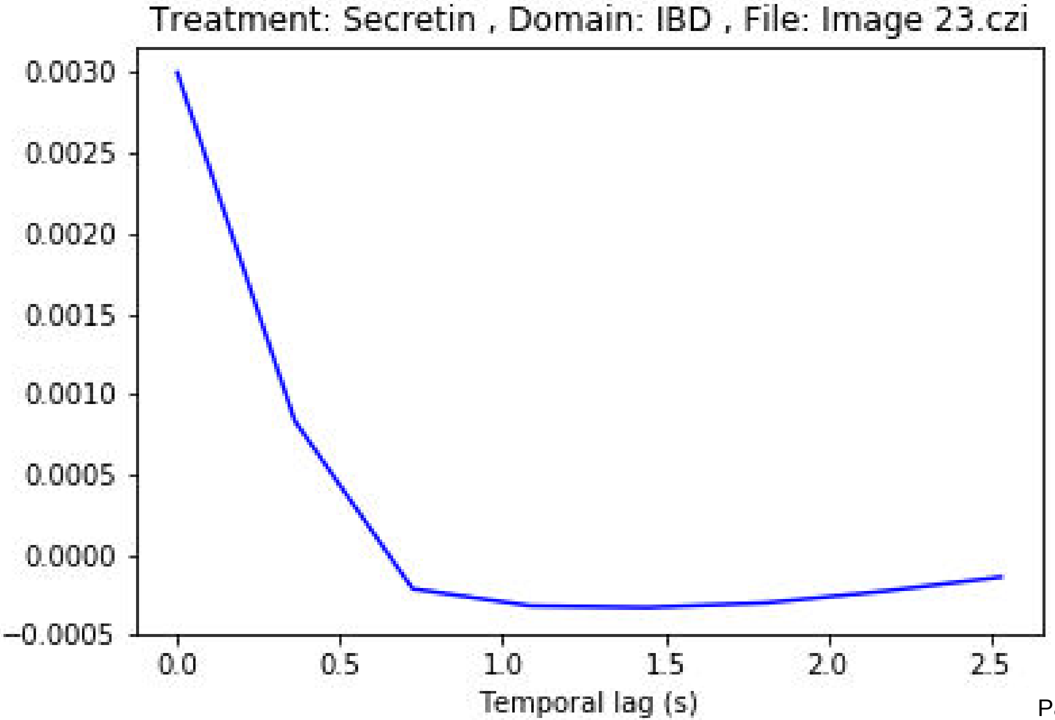
Treatment: Secretin , Domain: IBD , File: Image 23.czi
0.0030
0.0025
0.0020
0.0015
0.0010
0.0005
0.0000
−0.0005
0.0
0.5
1.0
1.5
2.0
2.5
Temporal lag (s)

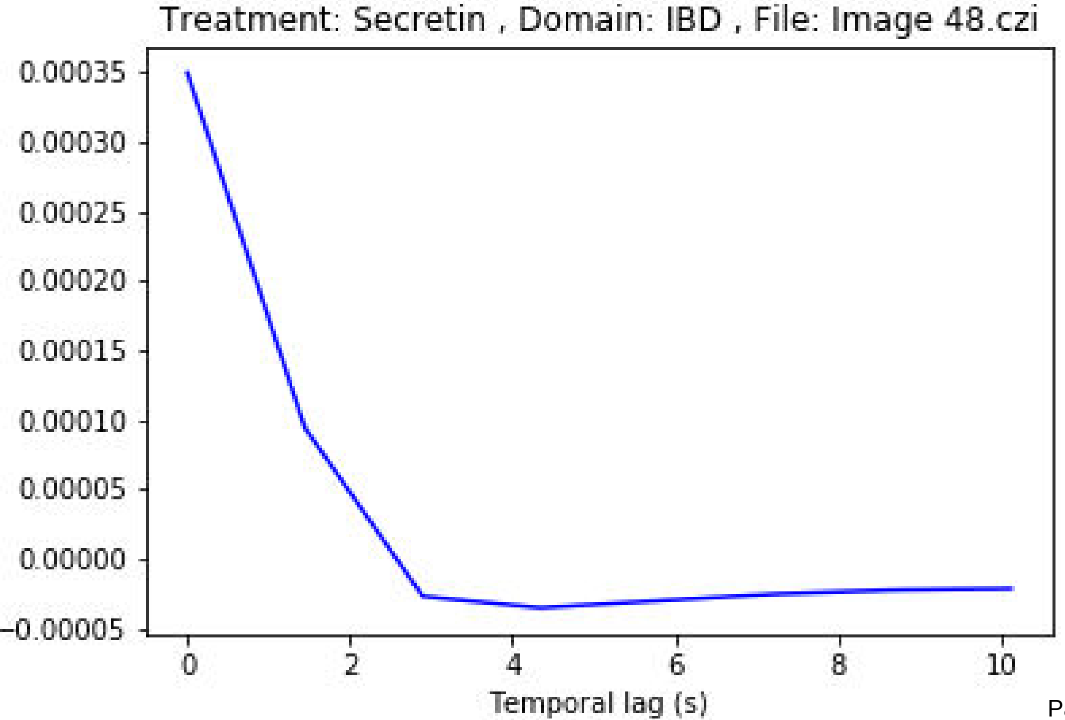
Treatment: Secretin , Domain: IBD , File: Image 48.czi
0.00035
0.00030
0.00025
0.00020
0.00015
0.00010
0.00005
0.00000
−0.00005
0
2
4
6
8
10
Temporal lag (s)

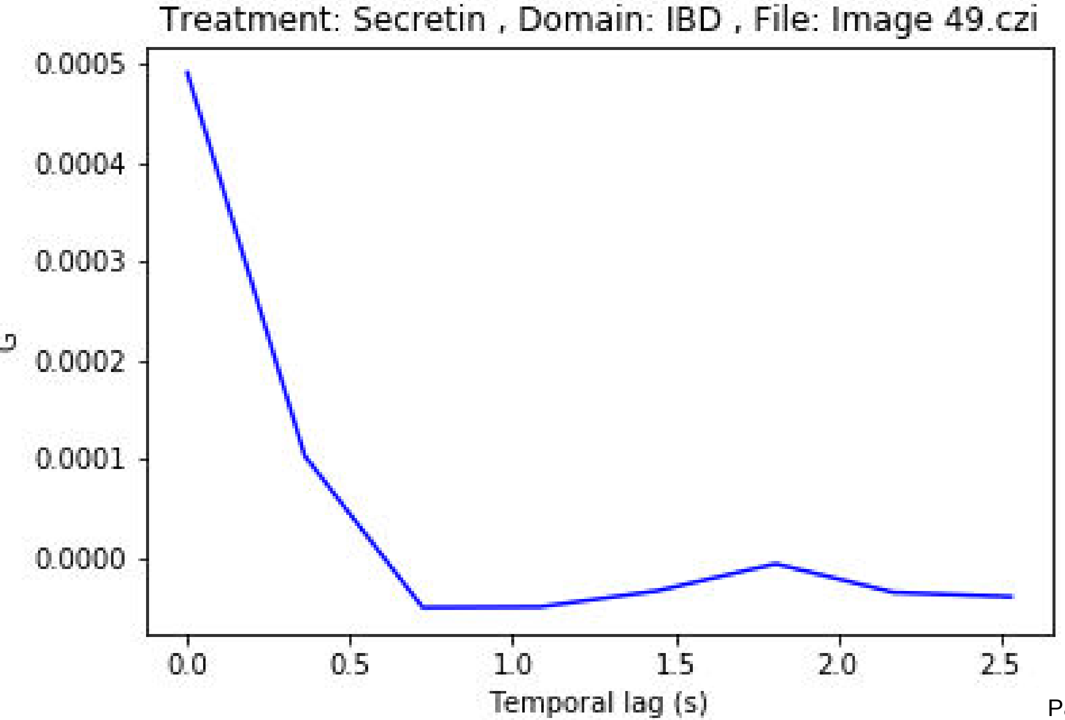
Treatment: Secretin , Domain: IBD , File: Image 49.czi
0.0005
0.0004
0.0003
0.0002
0.0001
0.0000
G
0.0
0.5
1.0
1.5
2.0
2.5
Temporal lag (s)

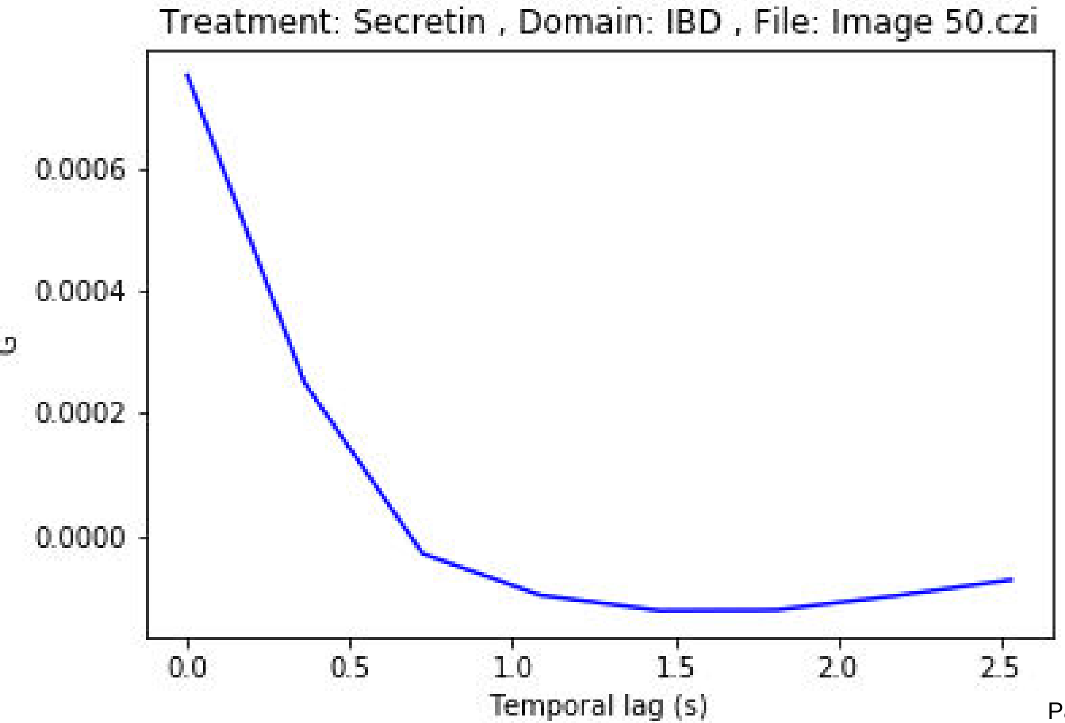
Treatment: Secretin , Domain: IBD , File: Image 50.czi
0.0006
0.0004
0.0002
0.0000
G
0.0
0.5
1.0
1.5
2.0
2.5
Temporal lag (s)

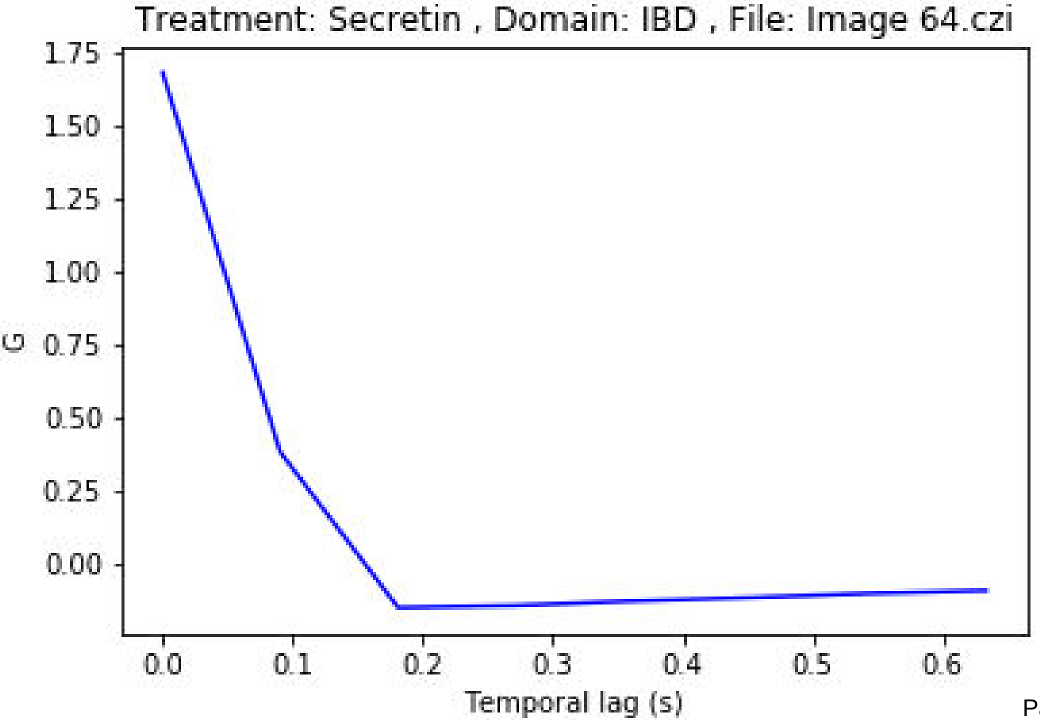
Treatment: Secretin , Domain: IBD , File: Image 64.czi
1.75
1.50
1.25
1.00
0.75
0.50
0.25
0.00
G
0.0
0.1
0.2
0.3
0.4
0.5
0.6
Temporal lag (s)

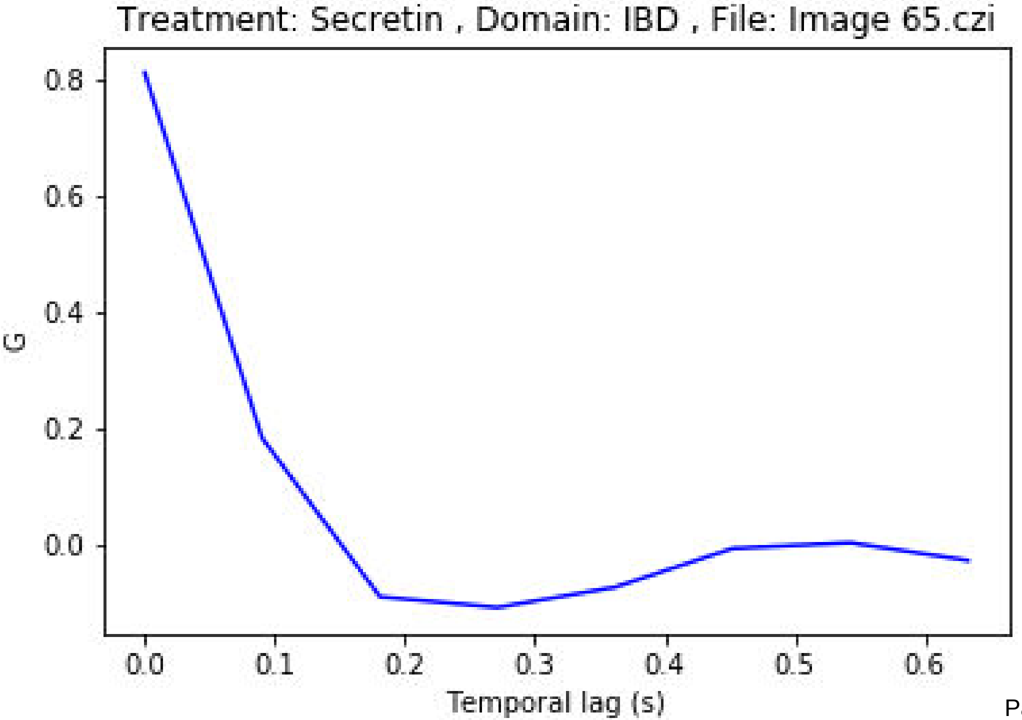

Treatment: Secretin , Domain: IBD , File: Image 65.czi
0.8
0.6
0.4
0.2
0.0
G
0.0
0.1
0.2
0.3
0.4
0.5
0.6
Temporal lag (s)

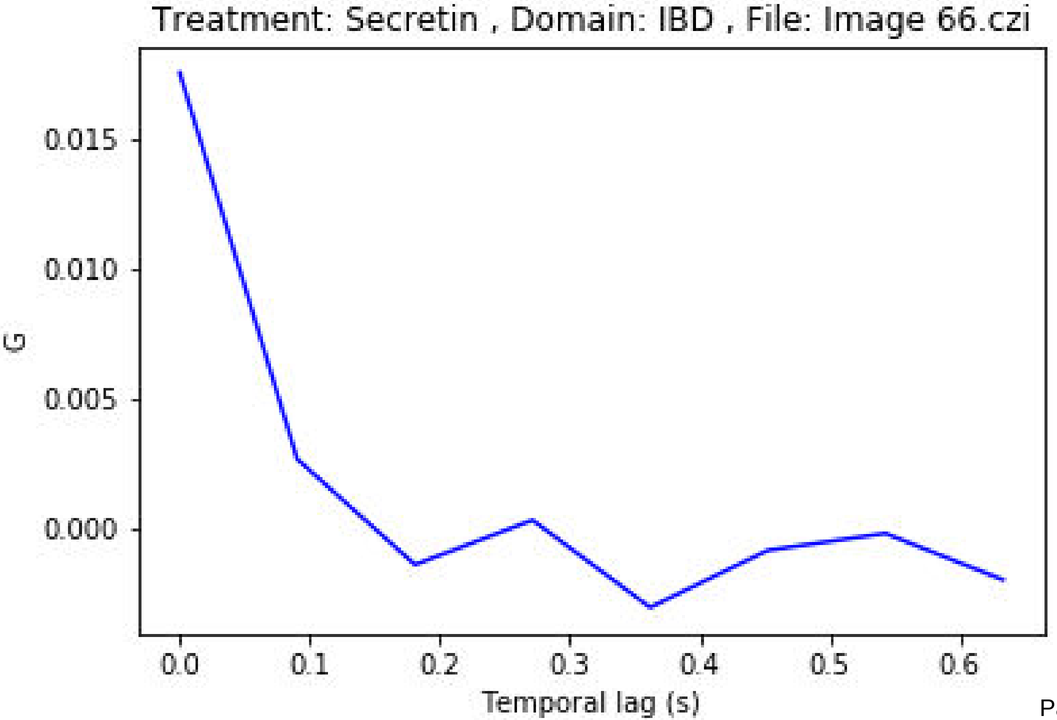

Treatment: Secretin , Domain: IBD , File: Image 66.czi
0.015
0.010
0.005
0.000
G
0.0
0.1
0.2
0.3
0.4
0.5
0.6
Temporal lag (s)

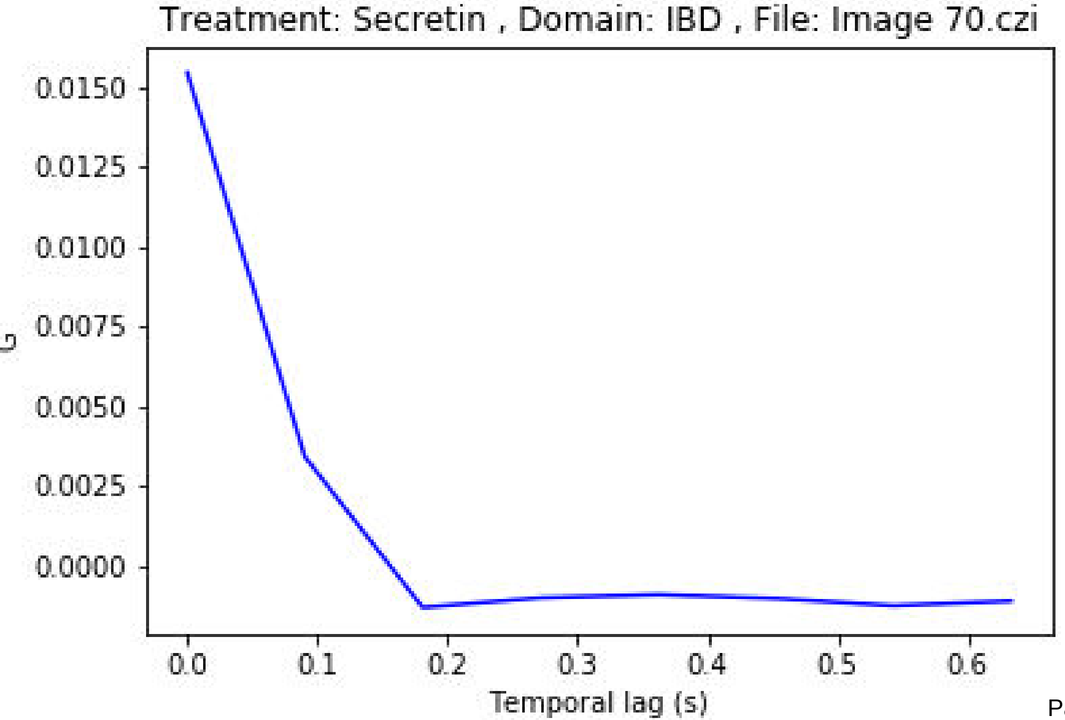
Treatment: Secretin , Domain: IBD , File: Image 70.czi
G
0.0150
0.0125
0.0100
0.0075
0.0050
0.0025
0.0000
0.0
0.1
0.2
0.3
0.4
0.5
0.6
Temporal lag (s)

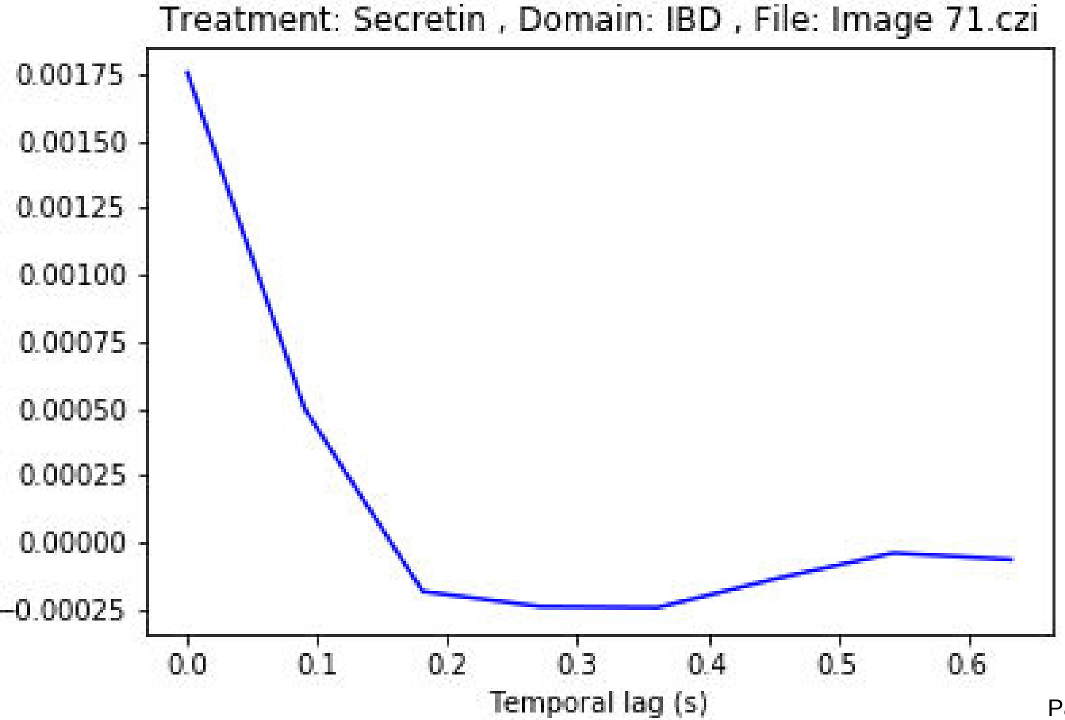
Treatment: Secretin , Domain: IBD , File: Image 71.czi
0.00175
0.00150
0.00125
0.00100
0.00075
0.00050
0.00025
0.00000
−0.00025
0.0
0.1
0.2
0.3
0.4
0.5
0.6
Temporal lag (s)

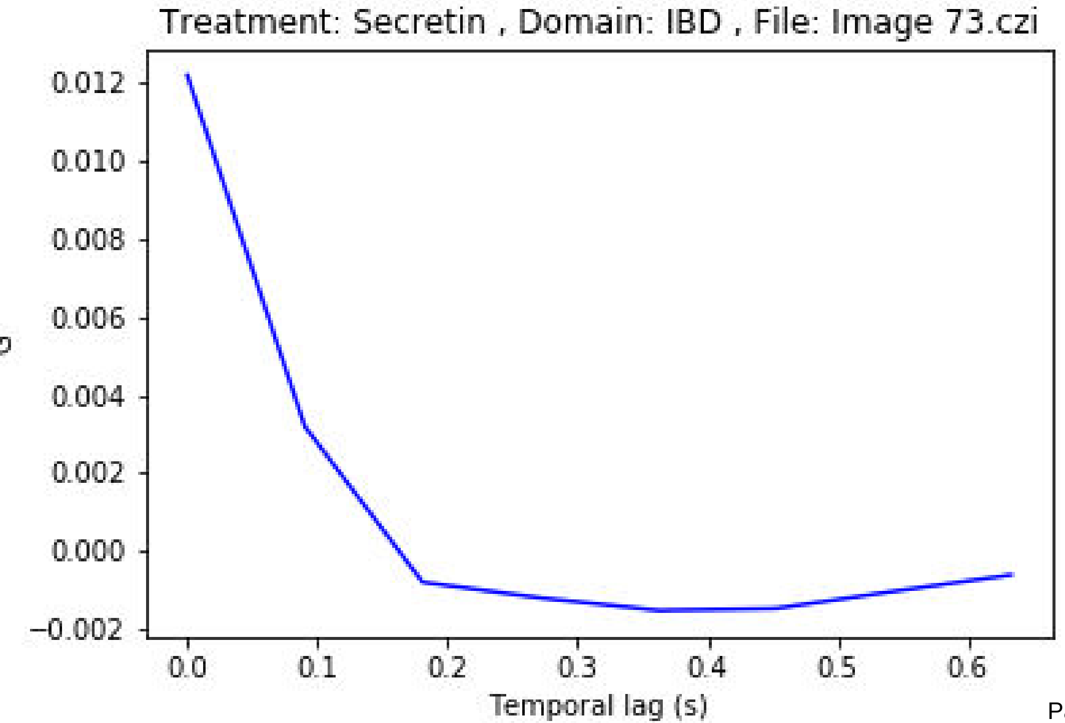

Treatment: Secretin , Domain: IBD , File: Image 73.czi
0.012
0.010
0.008
0.006
0.004
0.002
0.000
−0.002
0.0
0.1
0.2
0.3
0.4
0.5
0.6
Temporal lag (s)

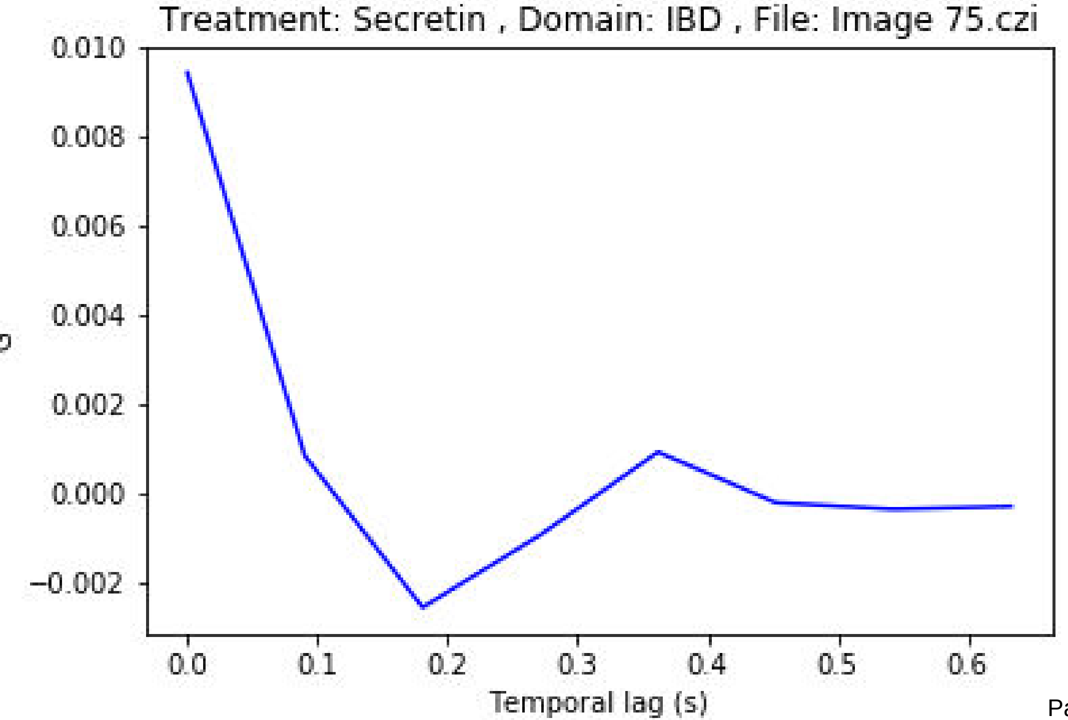
Treatment: Secretin , Domain: IBD , File: Image 75.czi
0.010
0.008
0.006
0.004
0.002
0.000
−0.002
0.0
0.1
0.2
0.3
0.4
0.5
0.6
Temporal lag (s)

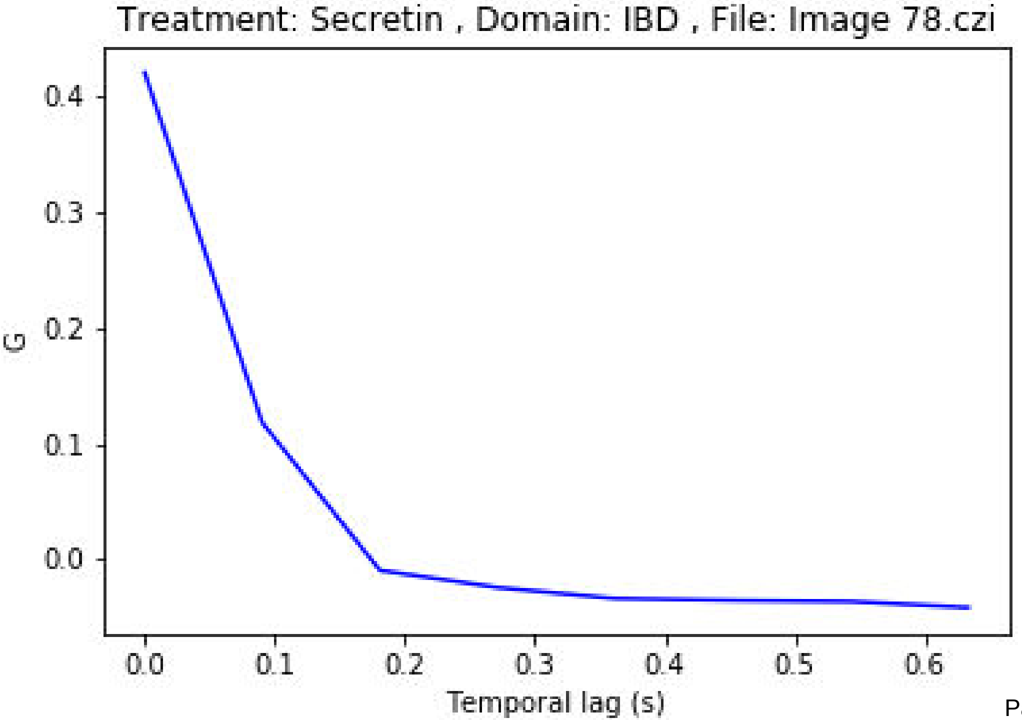
Treatment: Secretin , Domain: IBD , File: Image 78.czi
G
0.4
0.3
0.2
0.1
0.0
0.0
0.1
0.2
0.3
0.4
0.5
0.6
Temporal lag (s)

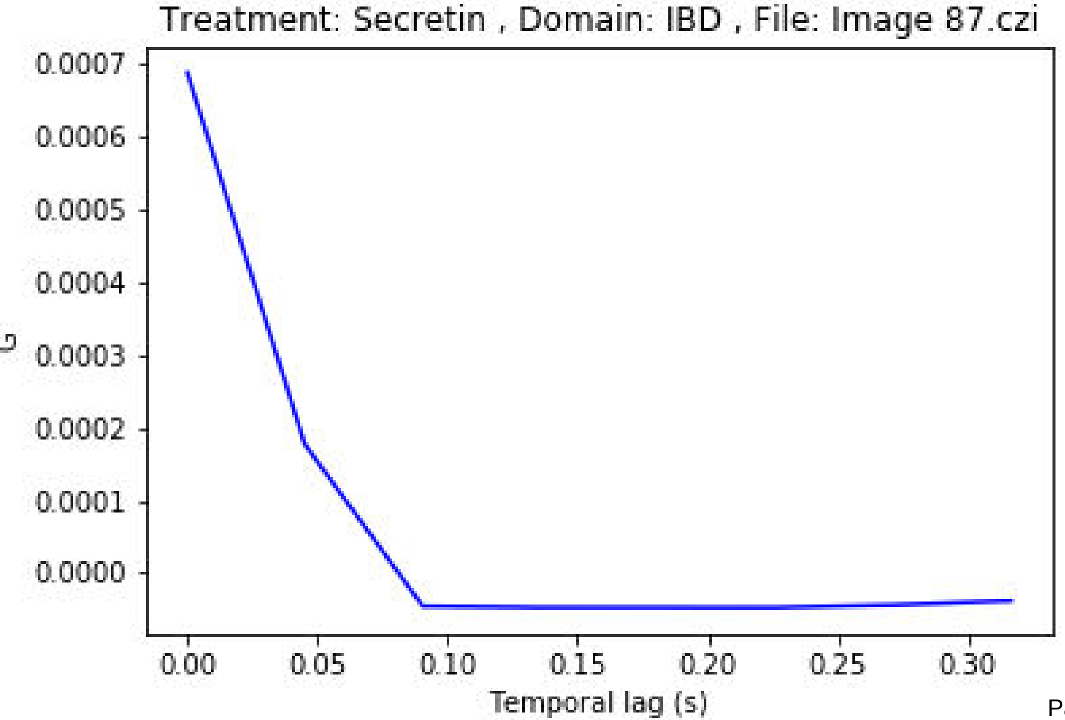
Treatment: Secretin , Domain: IBD , File: Image 87.czi
G
0.0007
0.0006
0.0005
0.0004
0.0003
0.0002
0.0001
0.0000
0.00
0.05
0.10
0.15
0.20
0.25
0.30
Temporal lag (s)

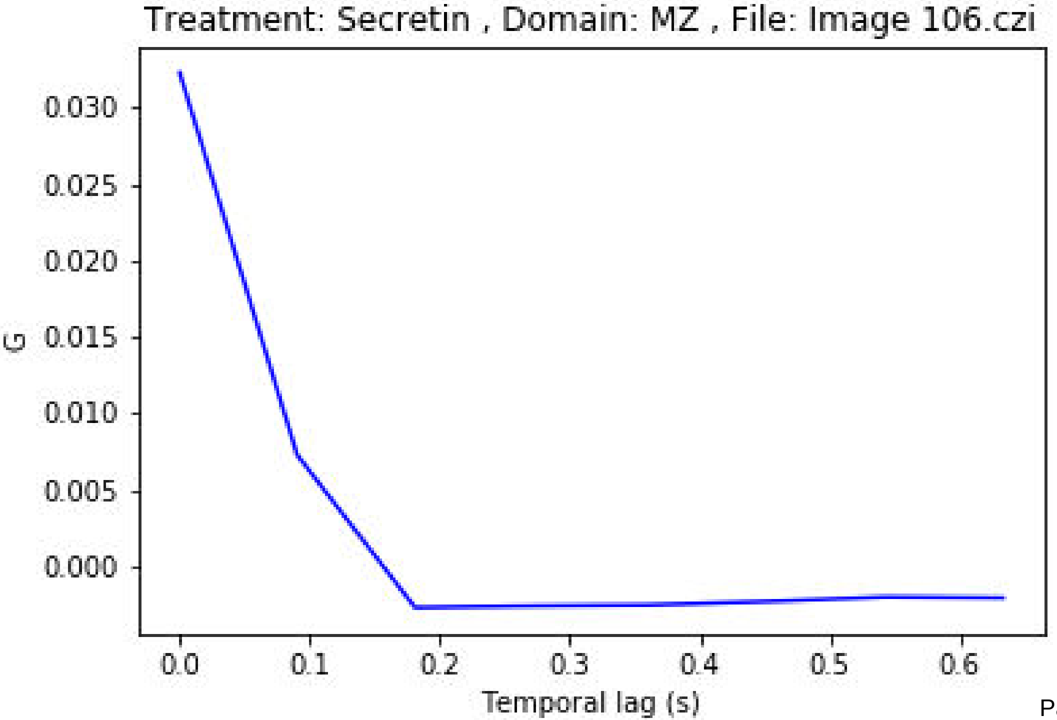

Treatment: Secretin , Domain: MZ , File: Image 106.czi
0.030
0.025
0.020
0.015
0.010
0.005
0.000
G
0.0
0.1
0.2
0.3
0.4
0.5
0.6
Temporal lag (s)

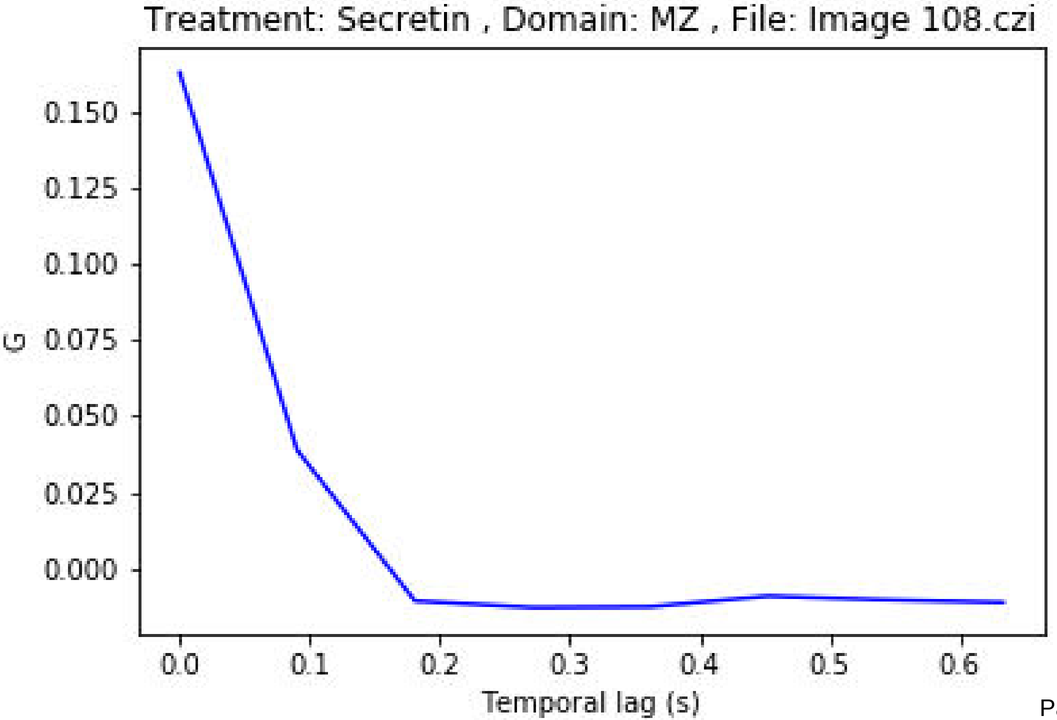
Treatment: Secretin , Domain: MZ , File: Image 108.czi
0.150
0.125
0.100
0.075
0.050
0.025
0.000
G
0.0
0.1
0.2
0.3
0.4
0.5
0.6
Temporal lag (s)

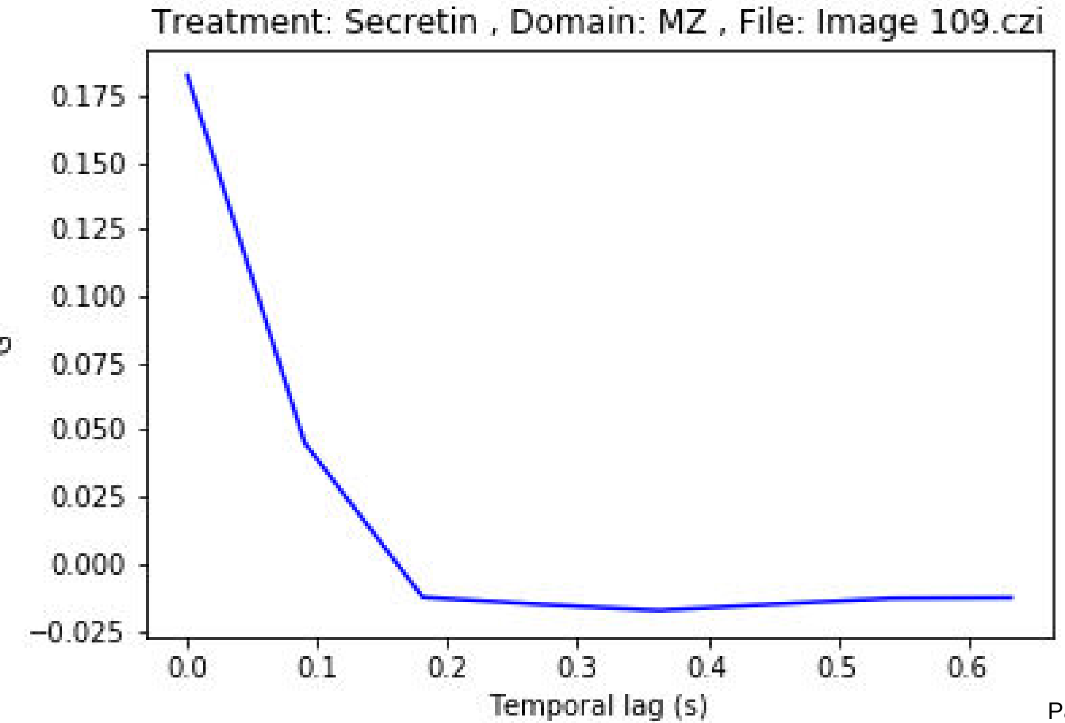
Treatment: Secretin , Domain: MZ , File: Image 109.czi
0.175
0.150
0.125
0.100
0.075
0.050
0.025
0.000
−0.025
0.0
0.1
0.2
0.3
0.4
0.5
0.6
Temporal lag (s)

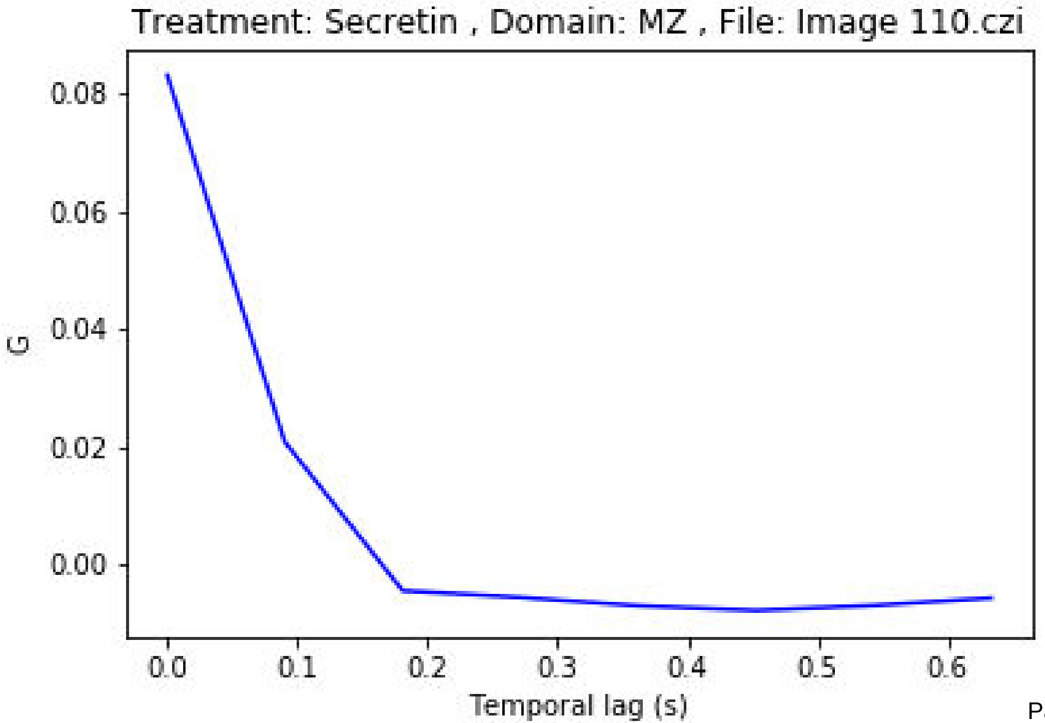
Treatment: Secretin , Domain: MZ , File: Image 110.czi
0.08
0.06
0.04
0.02
0.00
G
0.0
0.1
0.2
0.3
0.4
0.5
0.6
Temporal lag (s)

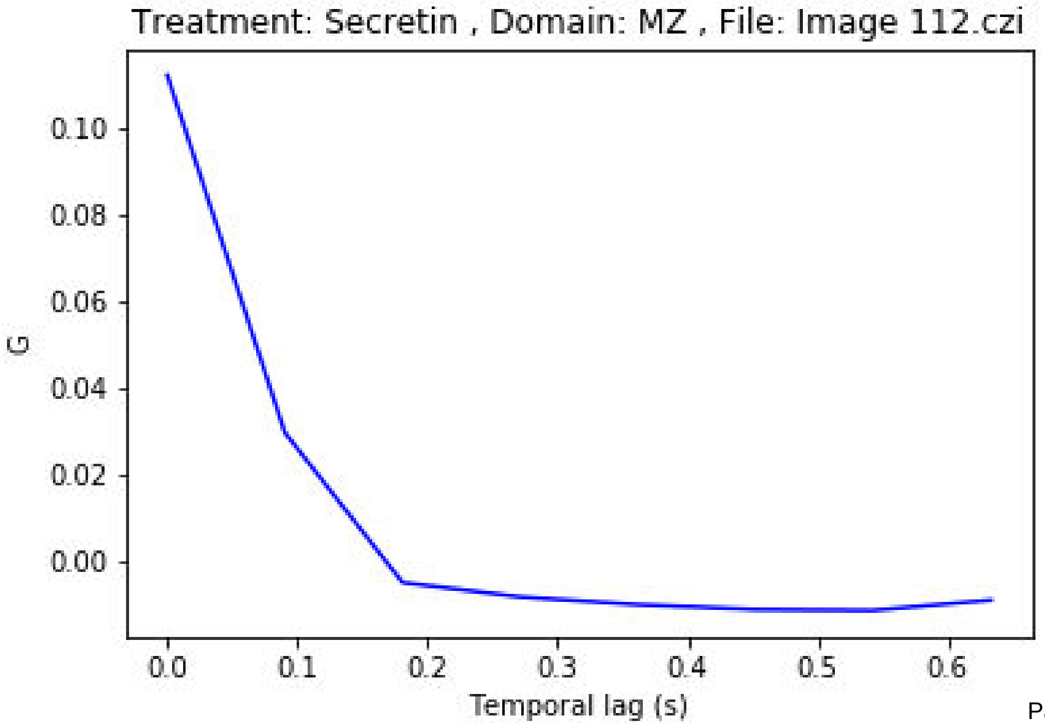
Treatment: Secretin , Domain: MZ , File: Image 112.czi
0.10
0.08
0.06
0.04
0.02
0.00
G
0.0
0.1
0.2
0.3
0.4
0.5
0.6
Temporal lag (s)

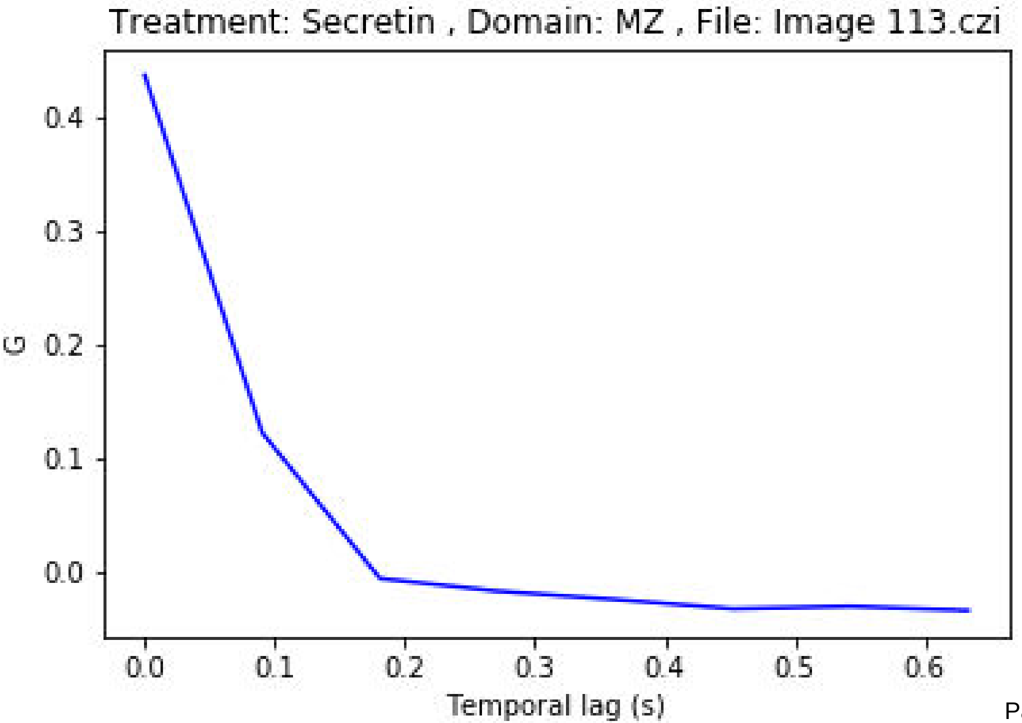
Treatment: Secretin , Domain: MZ , File: Image 113.czi
G
0.4
0.3
0.2
0.1
0.0
0.0
0.1
0.2
0.3
0.4
0.5
0.6
Temporal lag (s)

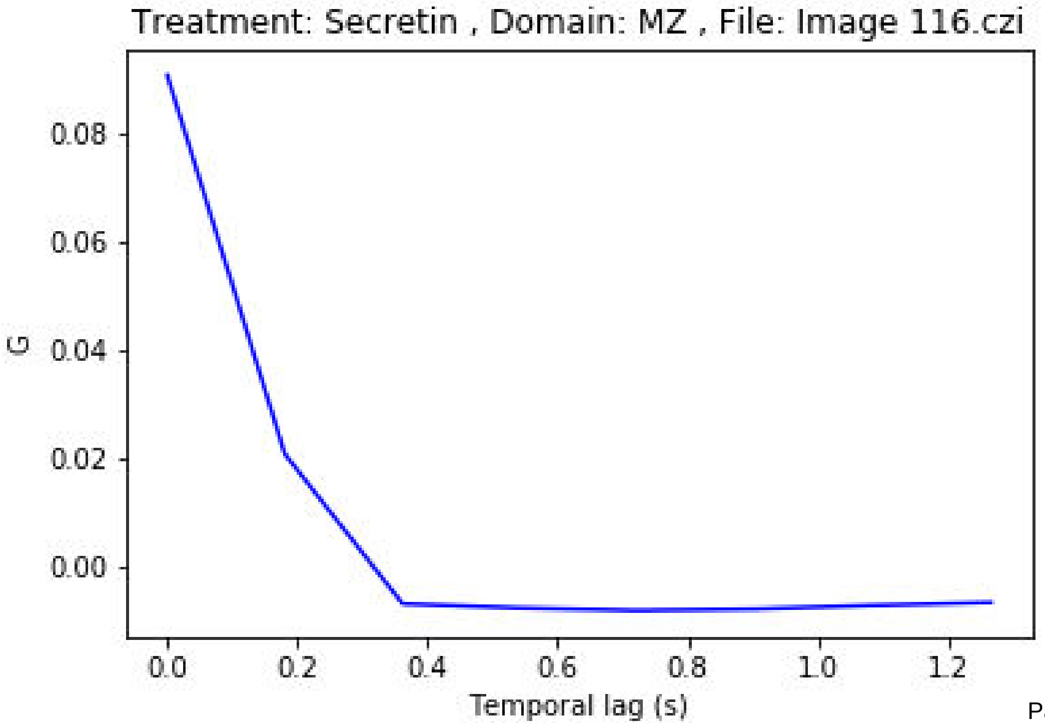
Treatment: Secretin , Domain: MZ , File: Image 116.czi
G
0.08
0.06
0.04
0.02
0.00
0.0
0.2
0.4
0.6
0.8
1.0
1.2
Temporal lag (s)

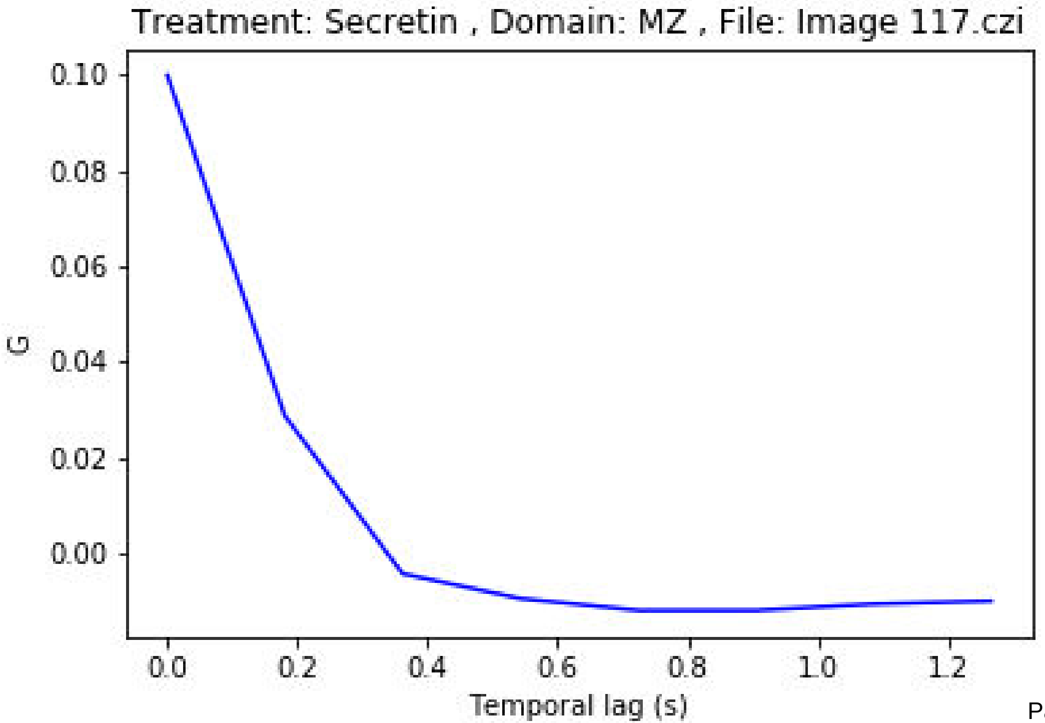
Treatment: Secretin , Domain: MZ , File: Image 117.czi
G
0.10
0.08
0.06
0.04
0.02
0.00
0.0
0.2
0.4
0.6
0.8
1.0
1.2
Temporal lag (s)

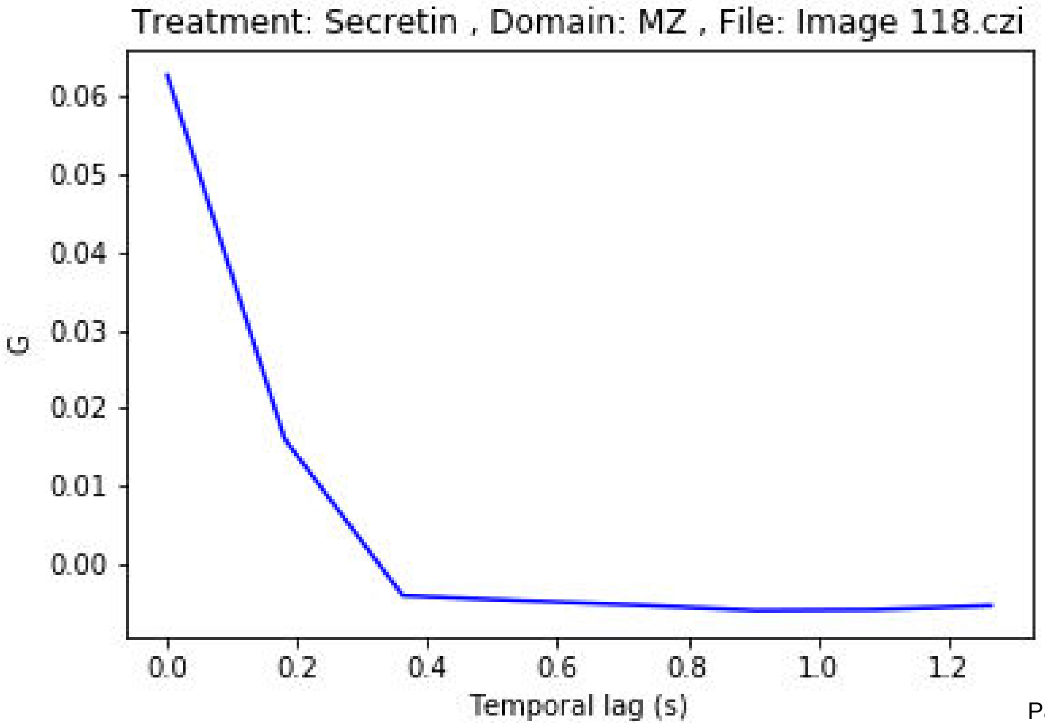
Treatment: Secretin , Domain: MZ , File: Image 118.czi
G
0.06
0.05
0.04
0.03
0.02
0.01
0.00
0.0
0.2
0.4
0.6
0.8
1.0
1.2
Temporal lag (s)

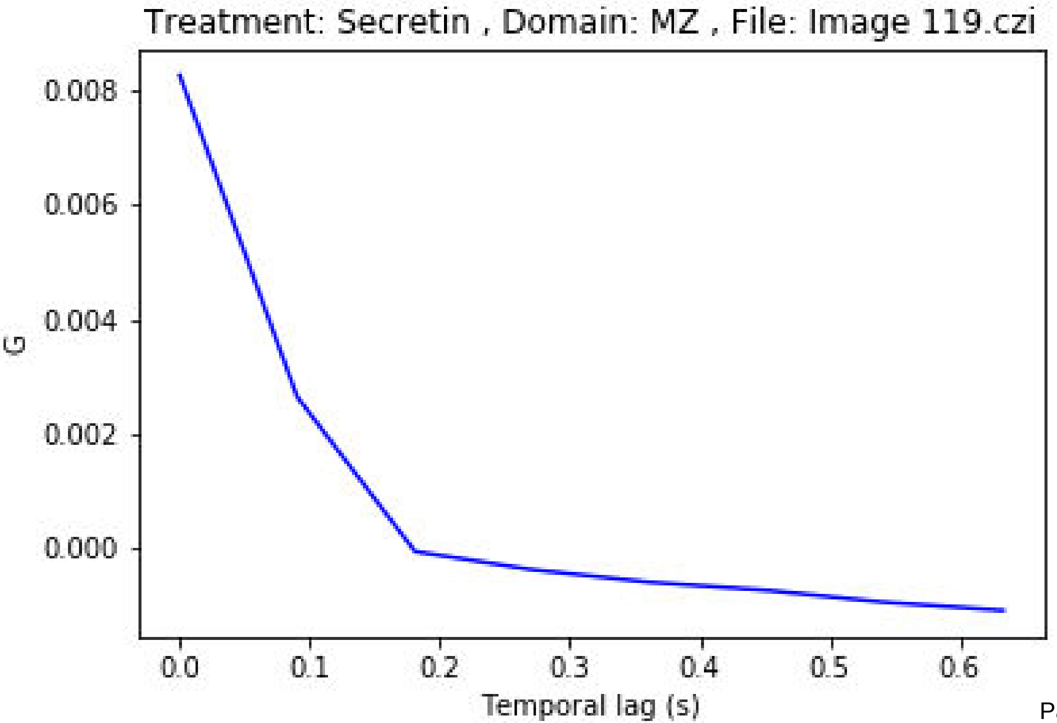

Treatment: Secretin , Domain: MZ , File: Image 119.czi
0.008
0.006
0.004
0.002
0.000
G
0.0
0.1
0.2
0.3
0.4
0.5
0.6
Temporal lag (s)

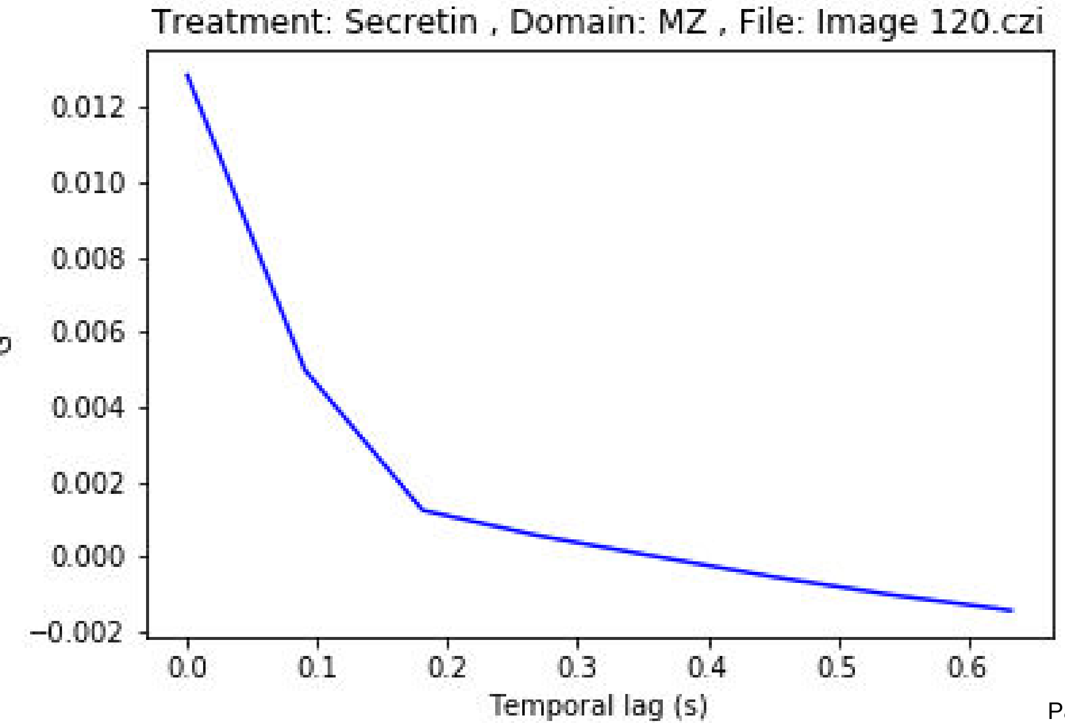
Treatment: Secretin , Domain: MZ , File: Image 120.czi
0.012
0.010
0.008
0.006
0.004
0.002
0.000
−0.002
0.0
0.1
0.2
0.3
0.4
0.5
0.6
Temporal lag (s)

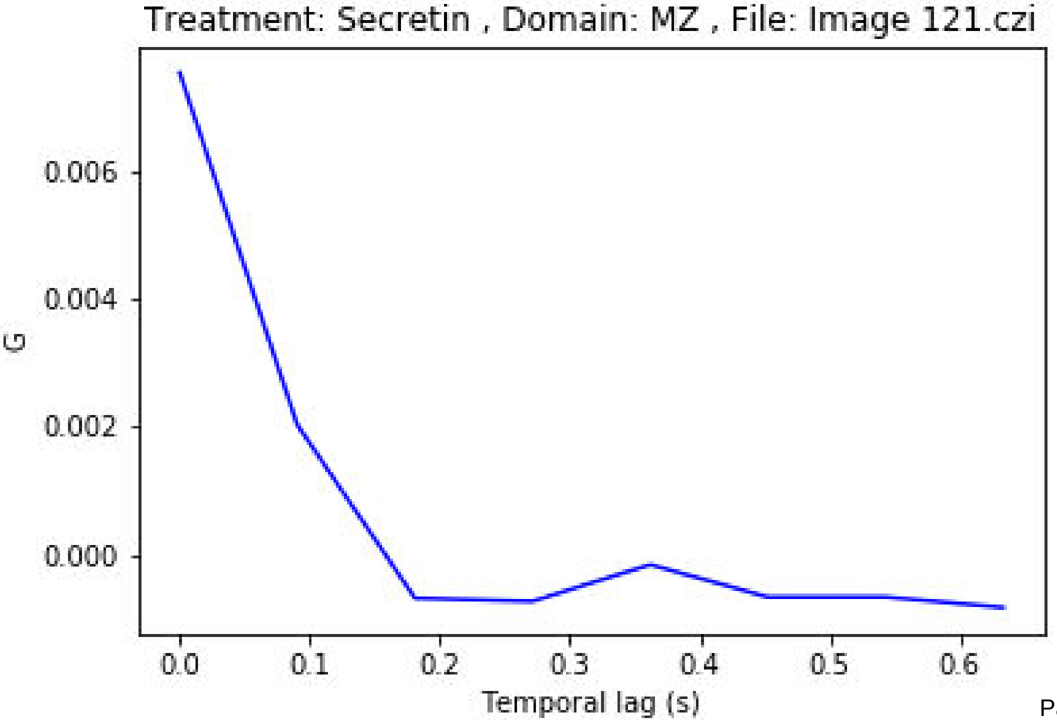
Treatment: Secretin , Domain: MZ , File: Image 121.czi
0.006
0.004
0.002
0.000
G
0.0
0.1
0.2
0.3
0.4
0.5
0.6
Temporal lag (s)

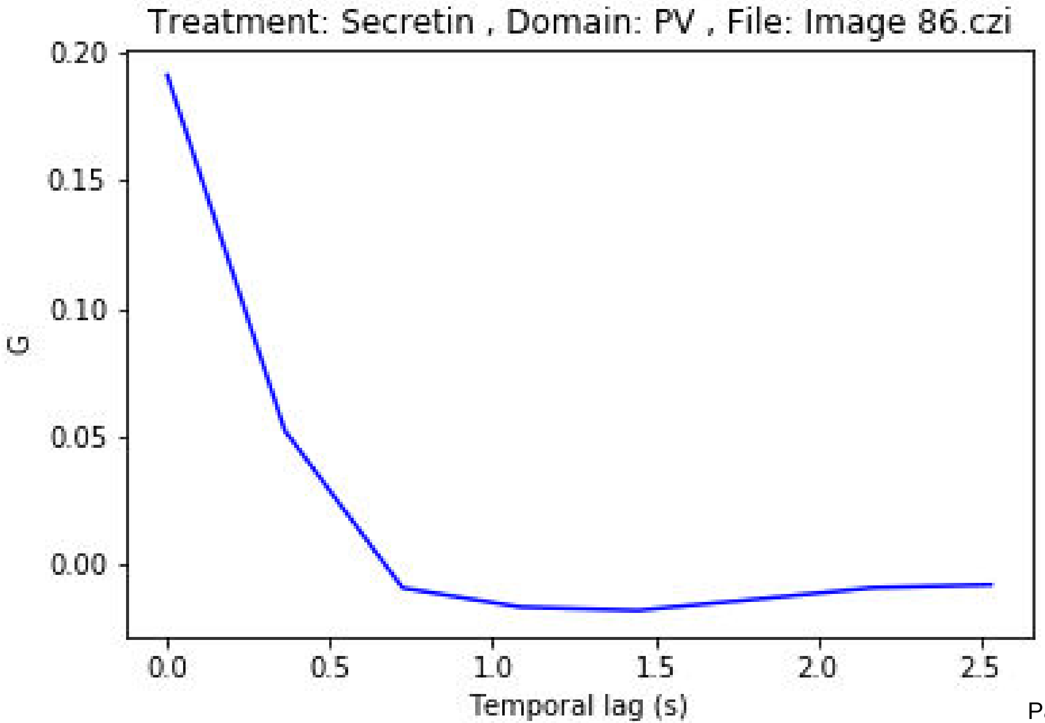
Treatment: Secretin , Domain: PV , File: Image 86.czi
0.20
0.15
0.10
0.05
0.00
G
0.0
0.5
1.0
1.5
2.0
2.5
Temporal lag (s)

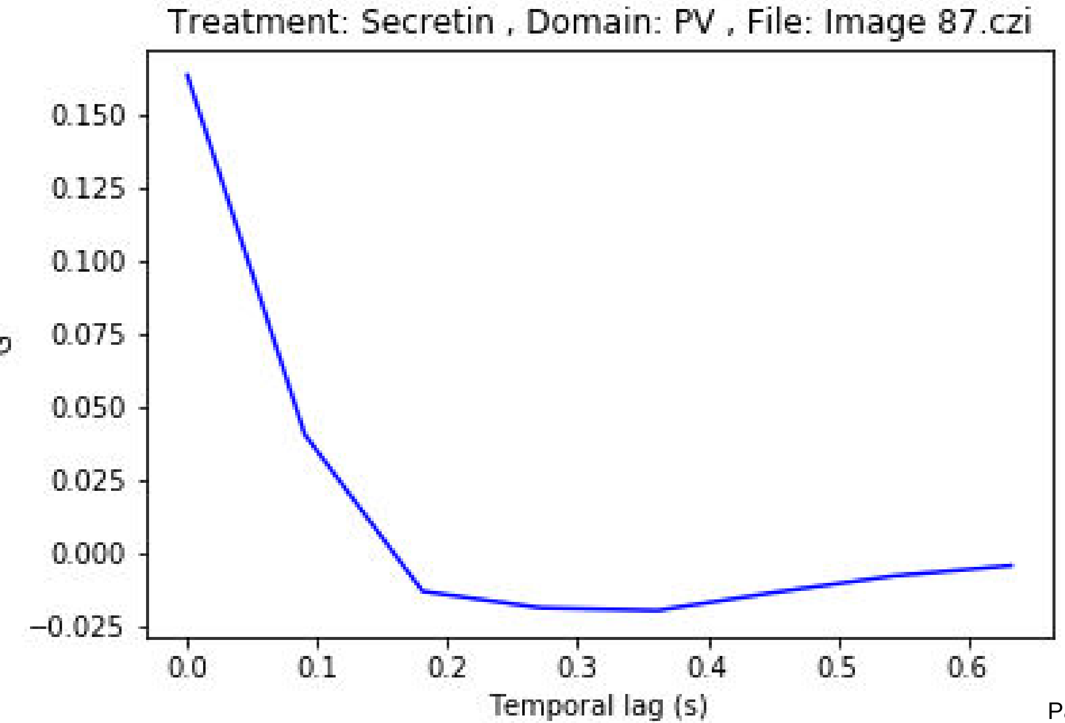

Treatment: Secretin , Domain: PV , File: Image 87.czi
0.150
0.125
0.100
0.075
0.050
0.025
0.000
−0.025
0.0
0.1
0.2
0.3
0.4
0.5
0.6
Temporal lag (s)

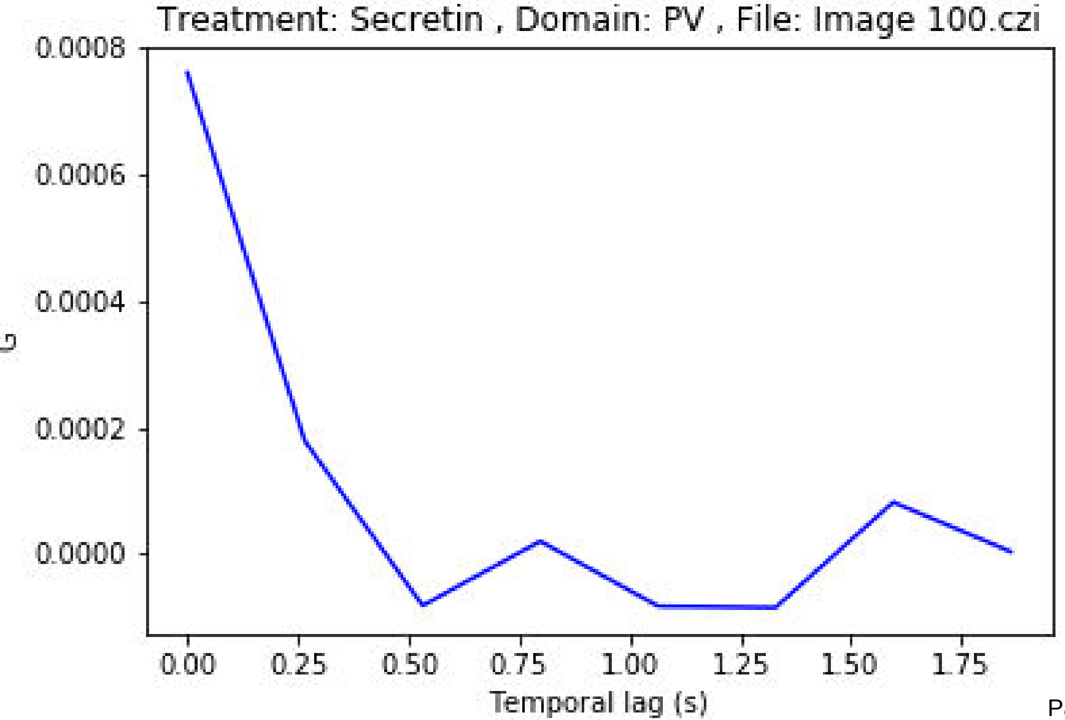
Treatment: Secretin , Domain: PV , File: Image 100.czi
0.0008
0.0006
0.0004
0.0002
0.0000
G
0.00
0.25
0.50
0.75
1.00
1.25
1.50
1.75
Temporal lag (s)

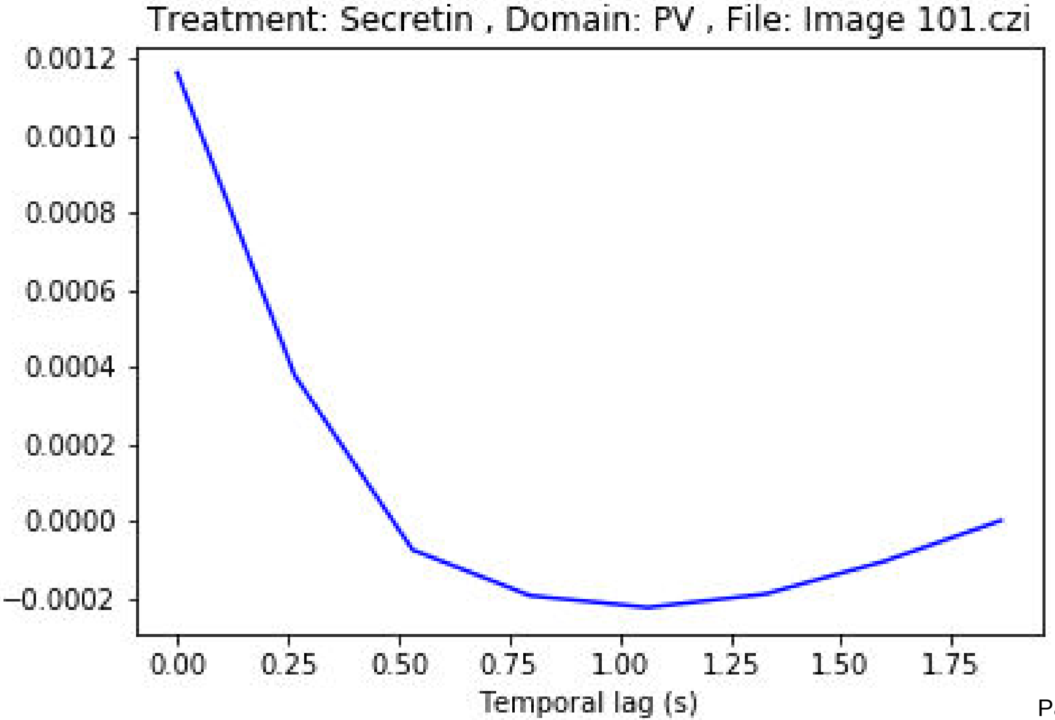

Treatment: Secretin , Domain: PV , File: Image 101.czi
0.0012
0.0010
0.0008
0.0006
0.0004
0.0002
0.0000
−0.0002
0.00
0.25
0.50
0.75
1.00
1.25
1.50
1.75
Temporal lag (s)

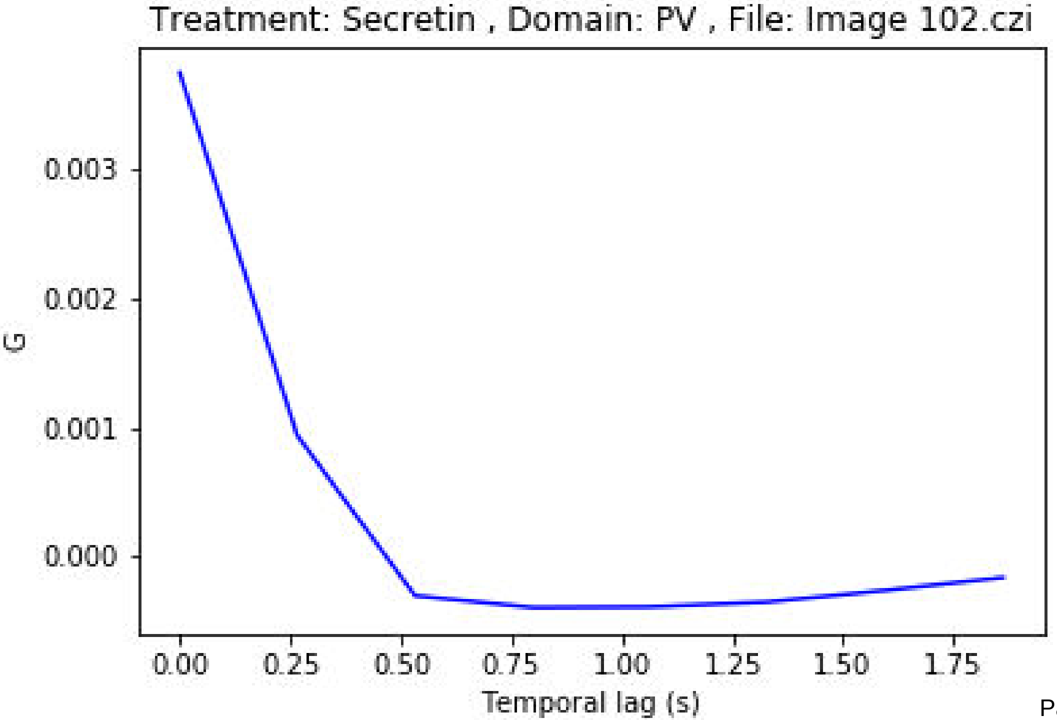
Treatment: Secretin , Domain: PV , File: Image 102.czi
0.003
0.002
0.001
0.000
G
0.00
0.25
0.50
0.75
1.00
1.25
1.50
1.75
Temporal lag (s)

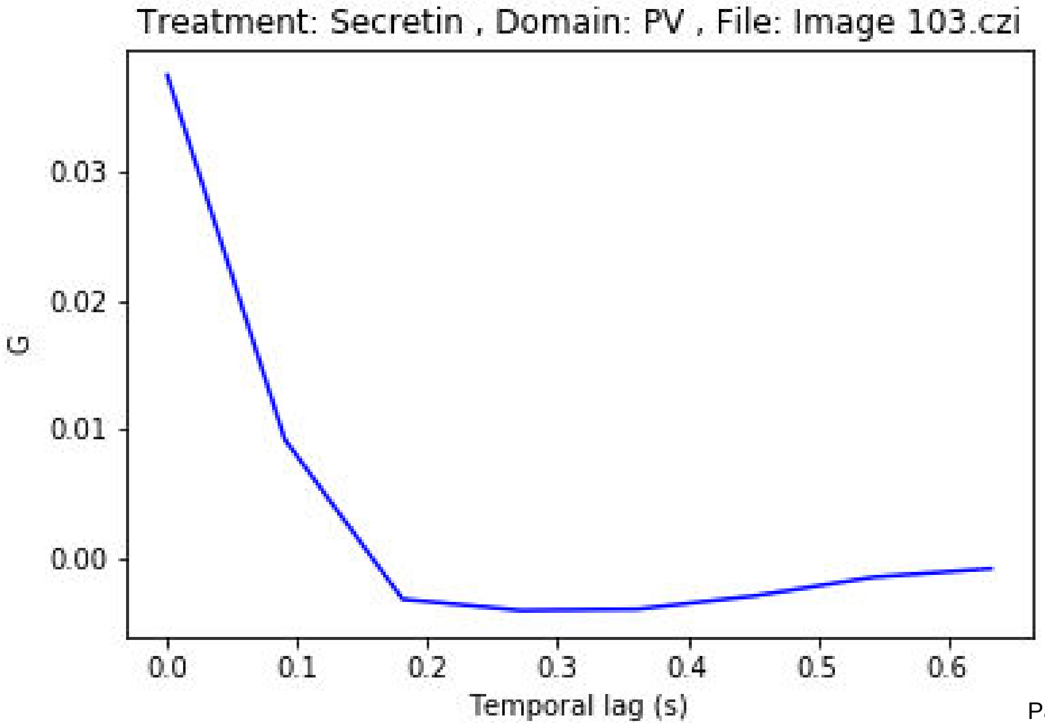

Treatment: Secretin , Domain: PV , File: Image 103.czi
0.03
0.02
0.01
0.00
G
0.0
0.1
0.2
0.3
0.4
0.5
0.6
Temporal lag (s)

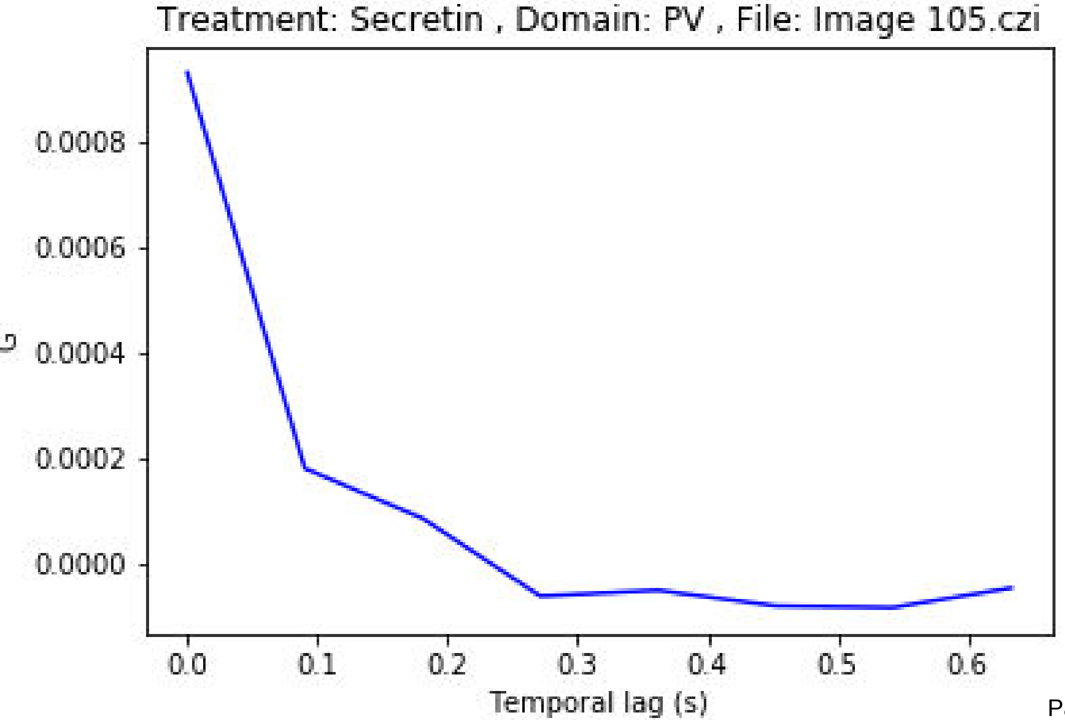
Treatment: Secretin , Domain: PV , File: Image 105.czi
G
0.0008
0.0006
0.0004
0.0002
0.0000
0.0
0.1
0.2
0.3
0.4
0.5
0.6
Temporal lag (s)

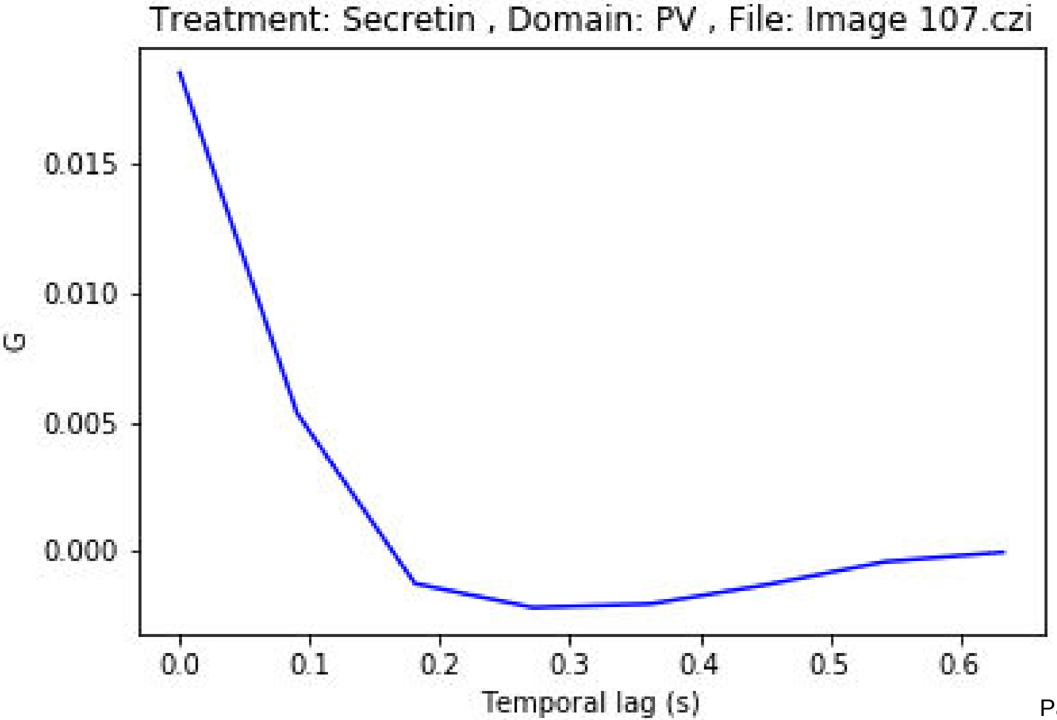

Treatment: Secretin , Domain: PV , File: Image 107.czi
0.015
0.010
0.005
0.000
G
0.0
0.1
0.2
0.3
0.4
0.5
0.6
Temporal lag (s)

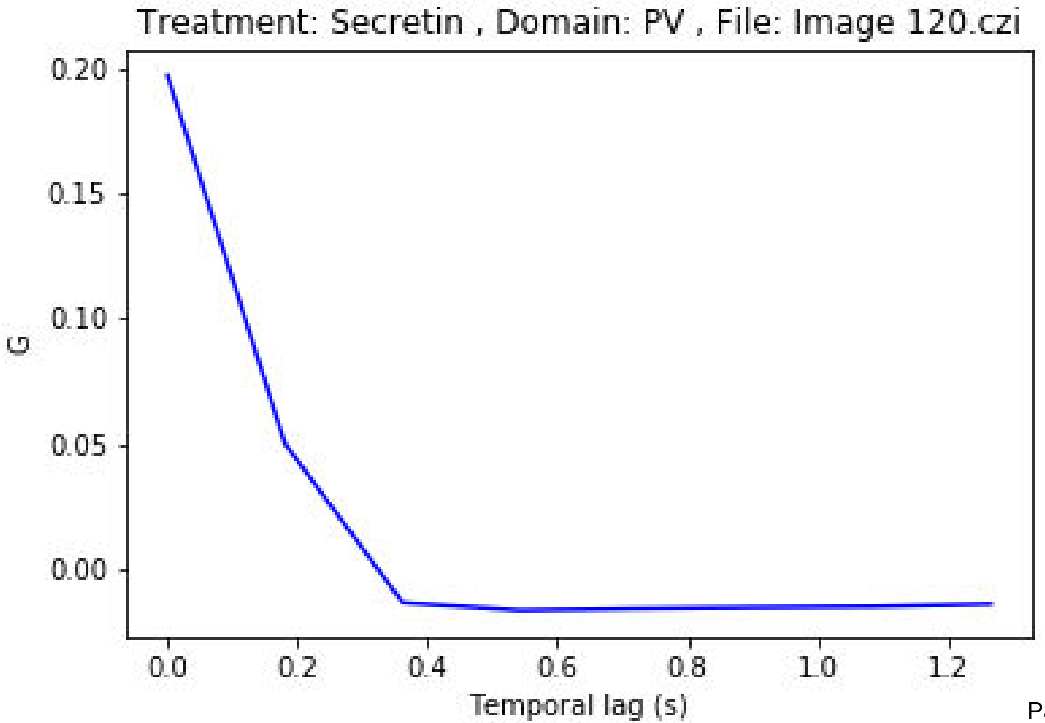
Treatment: Secretin , Domain: PV , File: Image 120.czi
0.20
0.15
0.10
0.05
0.00
G
0.0
0.2
0.4
0.6
0.8
1.0
1.2
Temporal lag (s)

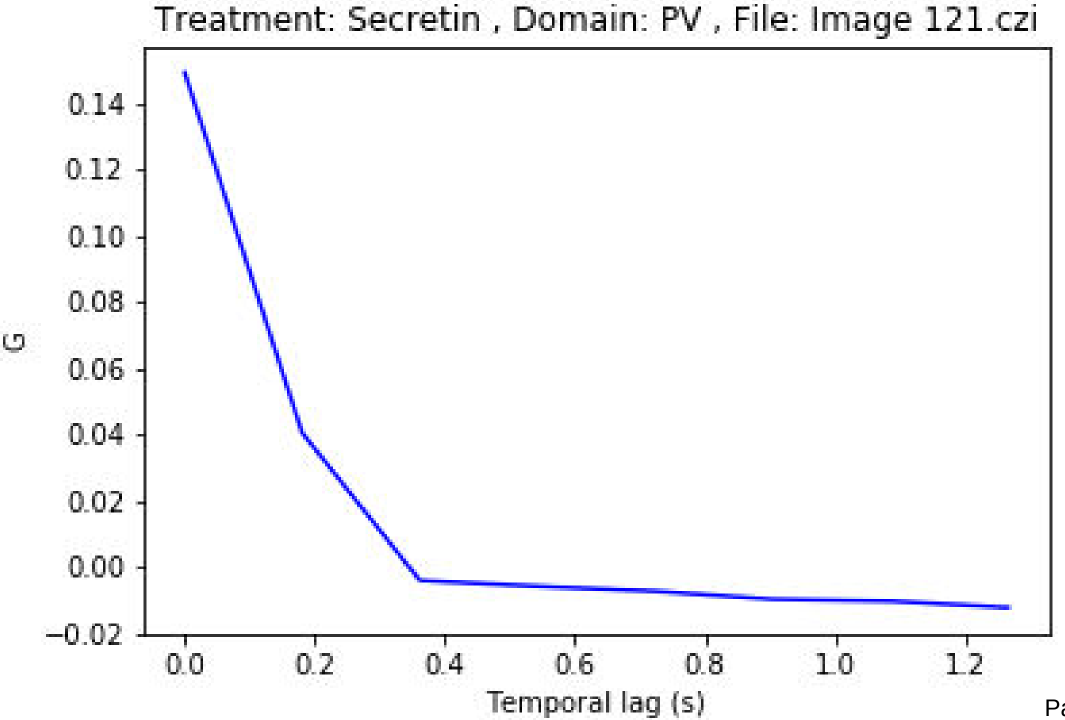
Treatment: Secretin , Domain: PV , File: Image 121.czi
0.14
0.12
0.10
0.08
0.06
0.04
0.02
0.00
−0.02
G
0.0
0.2
0.4
0.6
0.8
1.0
1.2
Temporal lag (s)

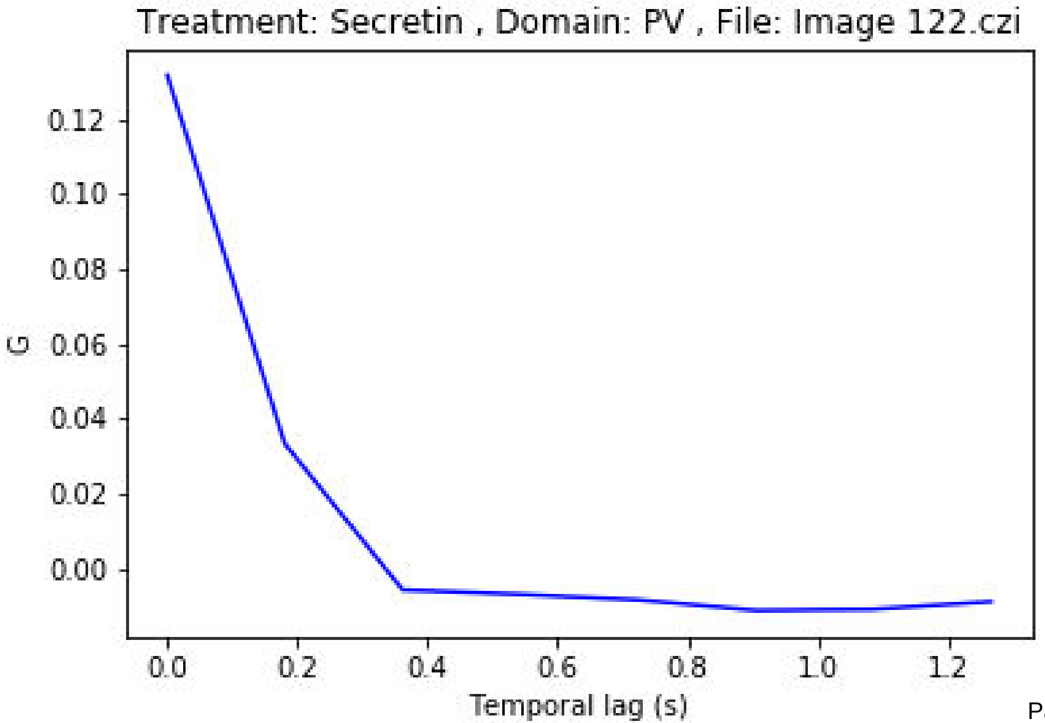
Treatment: Secretin , Domain: PV , File: Image 122.czi
G
0.12
0.10
0.08
0.06
0.04
0.02
0.00
0.0
0.2
0.4
0.6
0.8
1.0
1.2
Temporal lag (s)

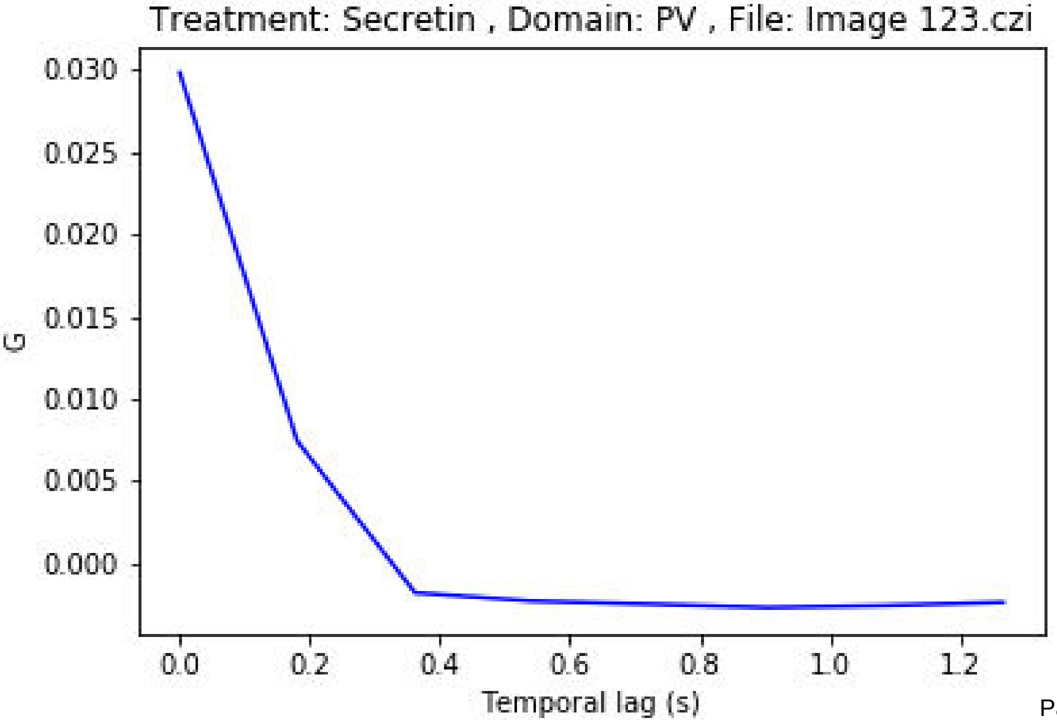
Treatment: Secretin , Domain: PV , File: Image 123.czi
0.030
0.025
0.020
0.015
0.010
0.005
0.000
G
0.0
0.2
0.4
0.6
0.8
1.0
1.2
Temporal lag (s)

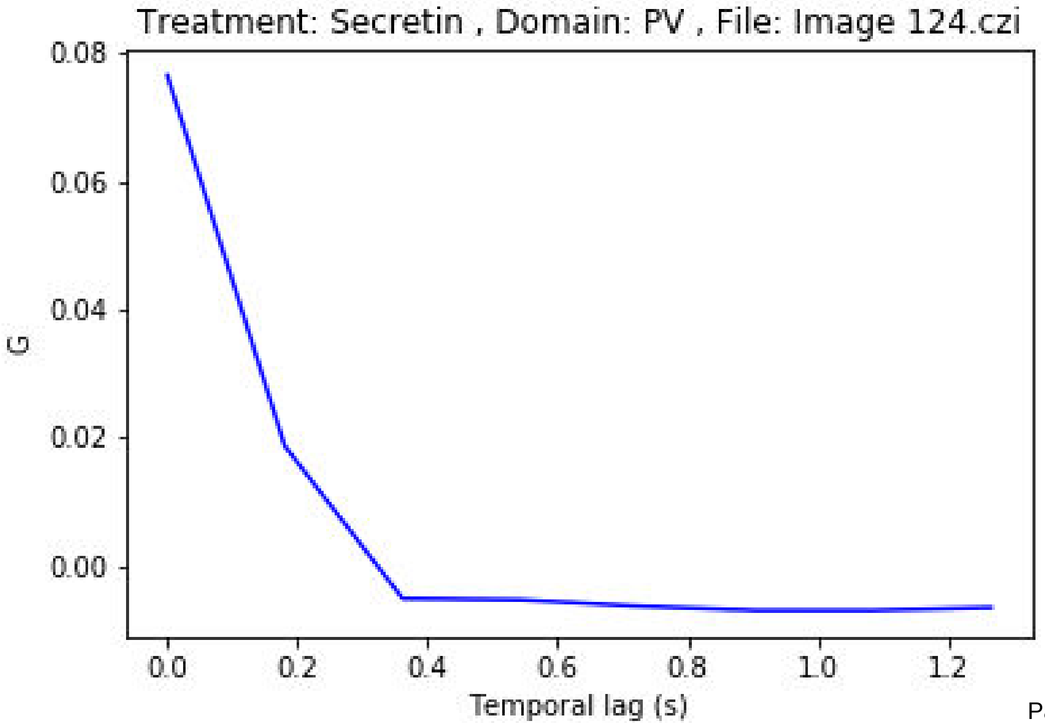
Treatment: Secretin , Domain: PV , File: Image 124.czi
0.08
0.06
0.04
0.02
0.00
G
0.0
0.2
0.4
0.6
0.8
1.0
1.2
Temporal lag (s)

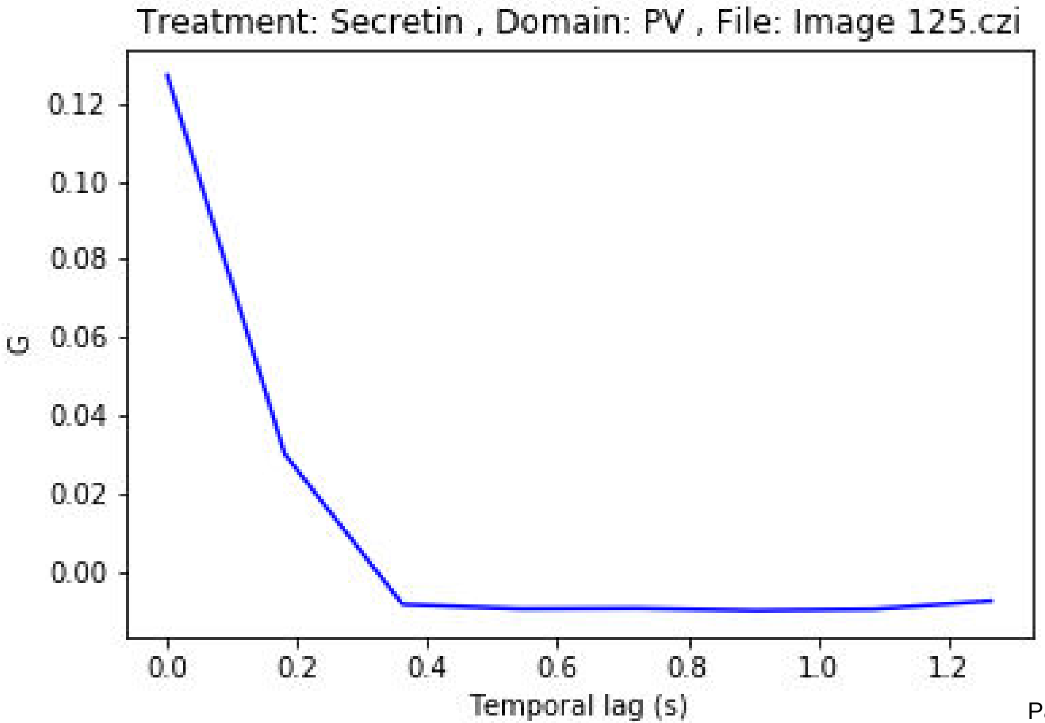
Treatment: Secretin , Domain: PV , File: Image 125.czi
G
0.12
0.10
0.08
0.06
0.04
0.02
0.00
0.0
0.2
0.4
0.6
0.8
1.0
1.2
Temporal lag (s)

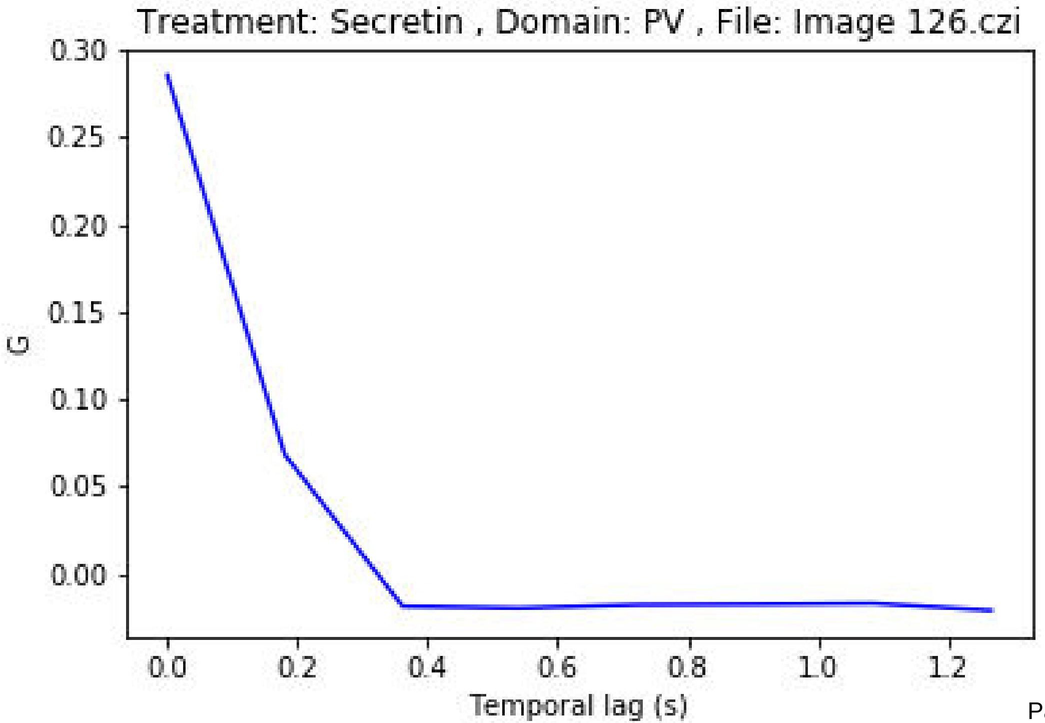
Treatment: Secretin , Domain: PV , File: Image 126.czi
0.30
0.25
0.20
0.15
0.10
0.05
0.00
G
0.0
0.2
0.4
0.6
0.8
1.0
1.2
Temporal lag (s)

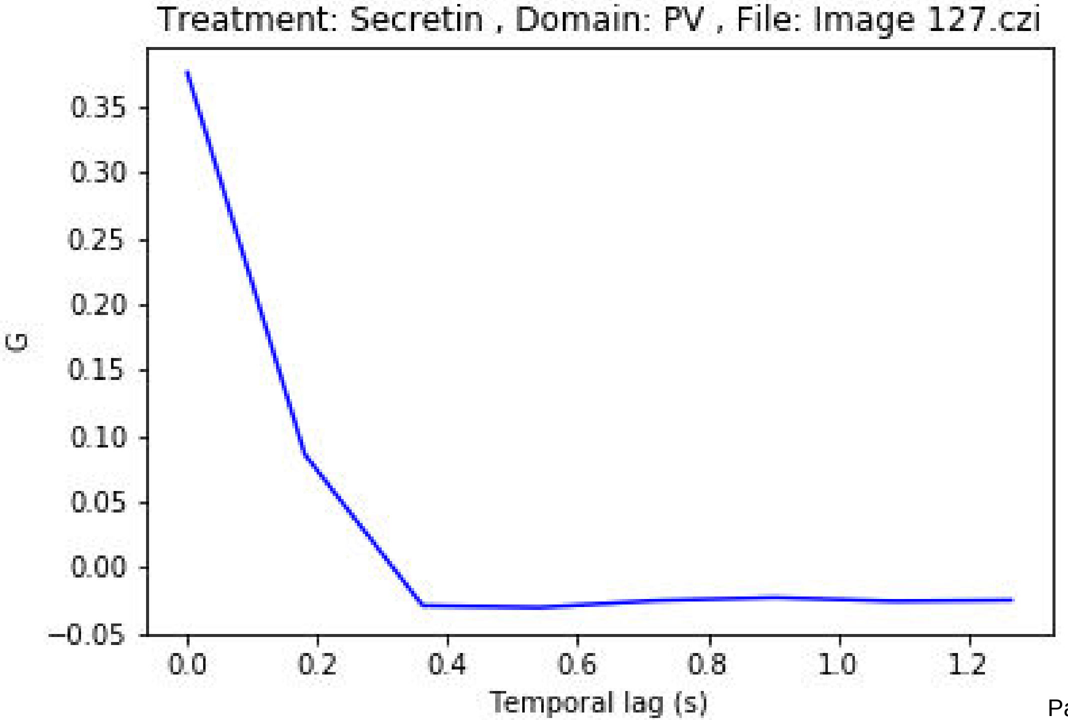

Treatment: Secretin , Domain: PV , File: Image 127.czi
0.35
0.30
0.25
0.20
0.15
0.10
0.05
0.00
−0.05
G
0.0
0.2
0.4
0.6
0.8
1.0
1.2
Temporal lag (s)

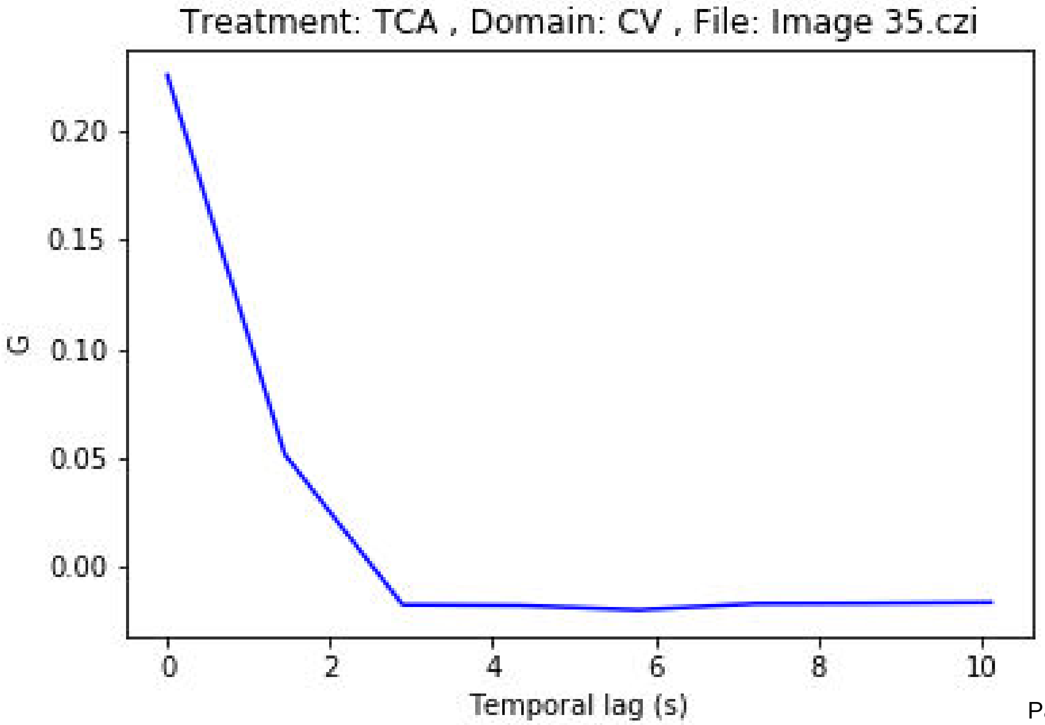
Treatment: TCA , Domain: CV , File: Image 35.czi
G
0.20
0.15
0.10
0.05
0.00
0
2
4
6
8
10
Temporal lag (s)

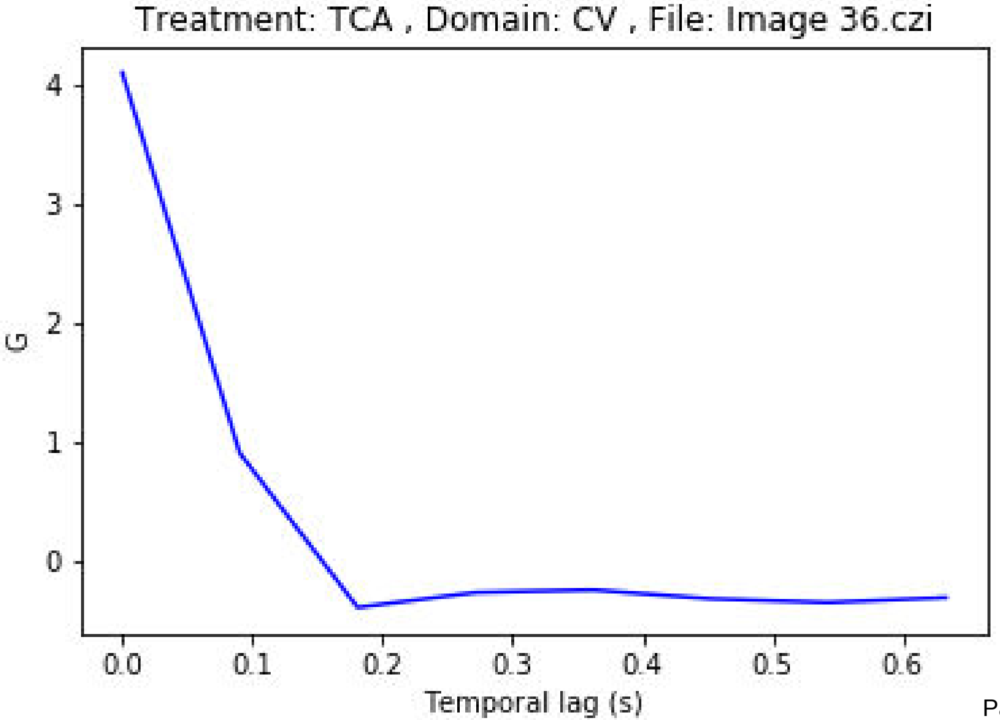
Treatment: TCA , Domain: CV , File: Image 36.czi
G
4
3
2
1
0
0.0
0.1
0.2
0.3
0.4
0.5
0.6
Temporal lag (s)

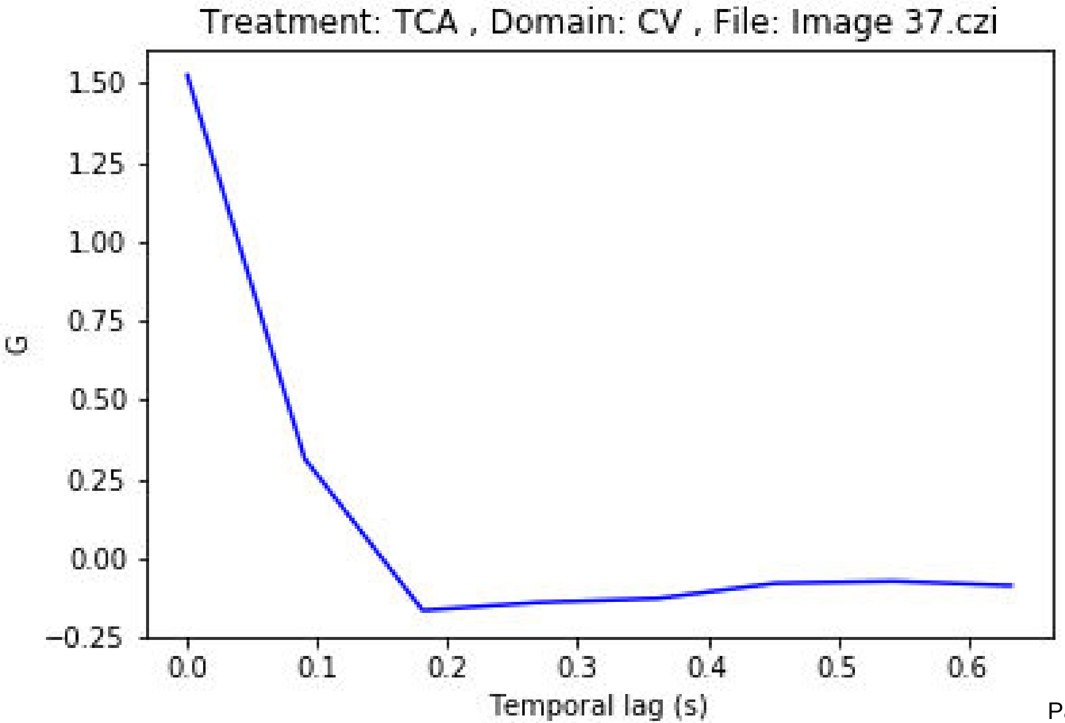
Treatment: TCA , Domain: CV , File: Image 37.czi
G
1.50
1.25
1.00
0.75
0.50
0.25
0.00
−0.25
0.0
0.1
0.2
0.3
0.4
0.5
0.6
Temporal lag (s)

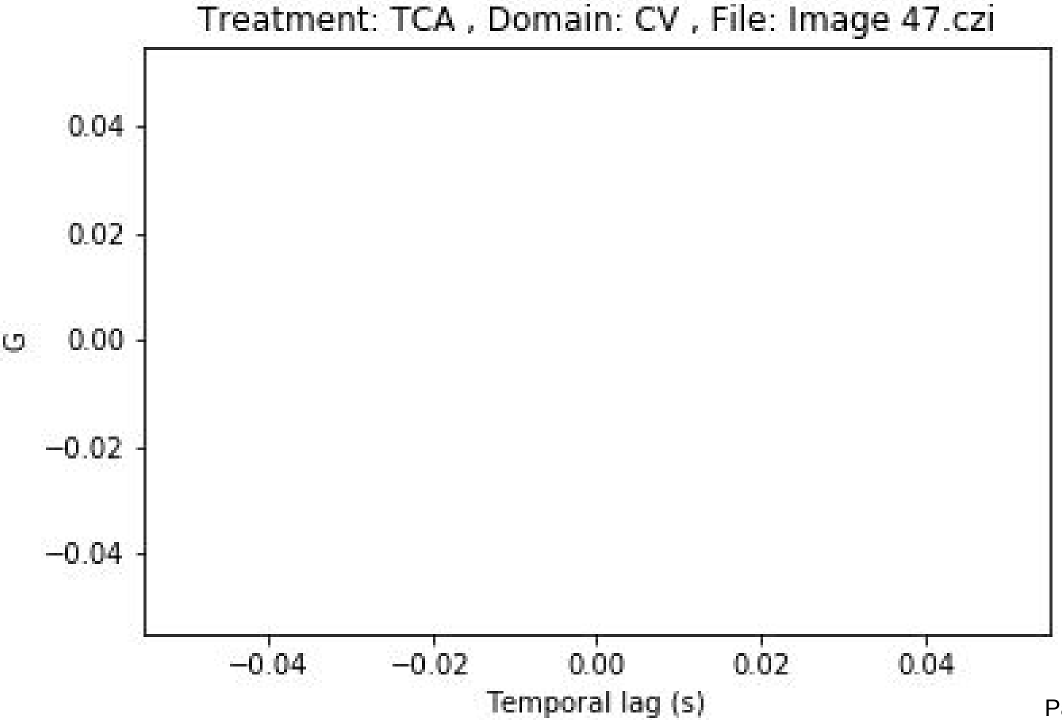
Treatment: TCA , Domain: CV , File: Image 47.czi
G
0.04
0.02
0.00
−0.02
−0.04
−0.04
−0.02
0.00
0.02
0.04
Temporal lag (s)

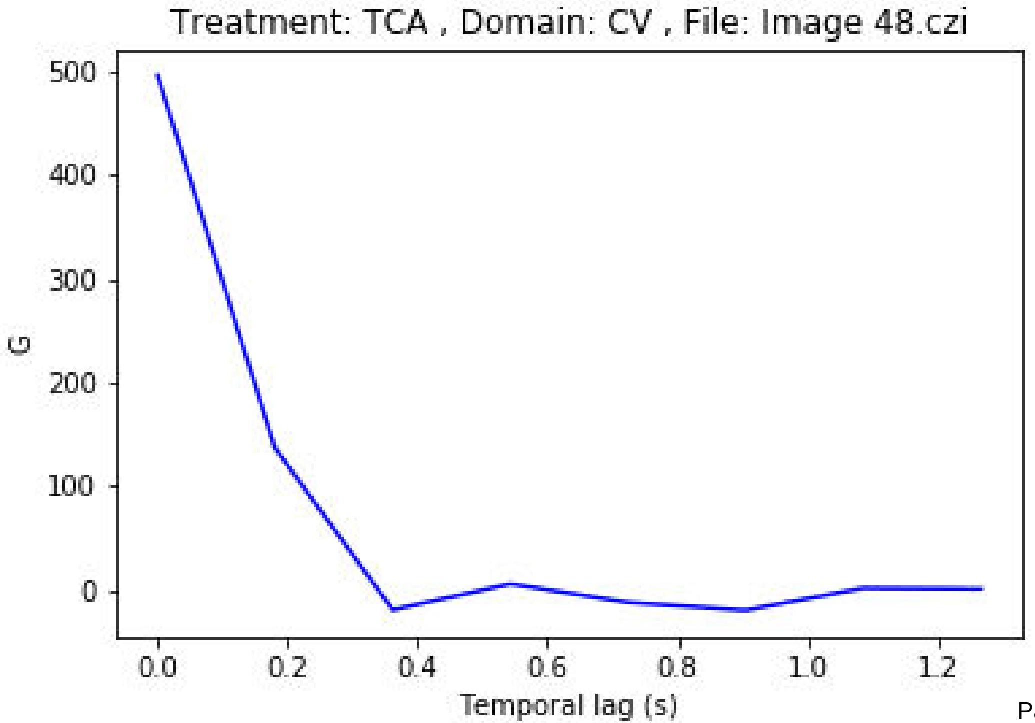
Treatment: TCA , Domain: CV , File: Image 48.czi
G
500
400
300
200
100
0
0.0
0.2
0.4
0.6
0.8
1.0
1.2
Temporal lag (s)

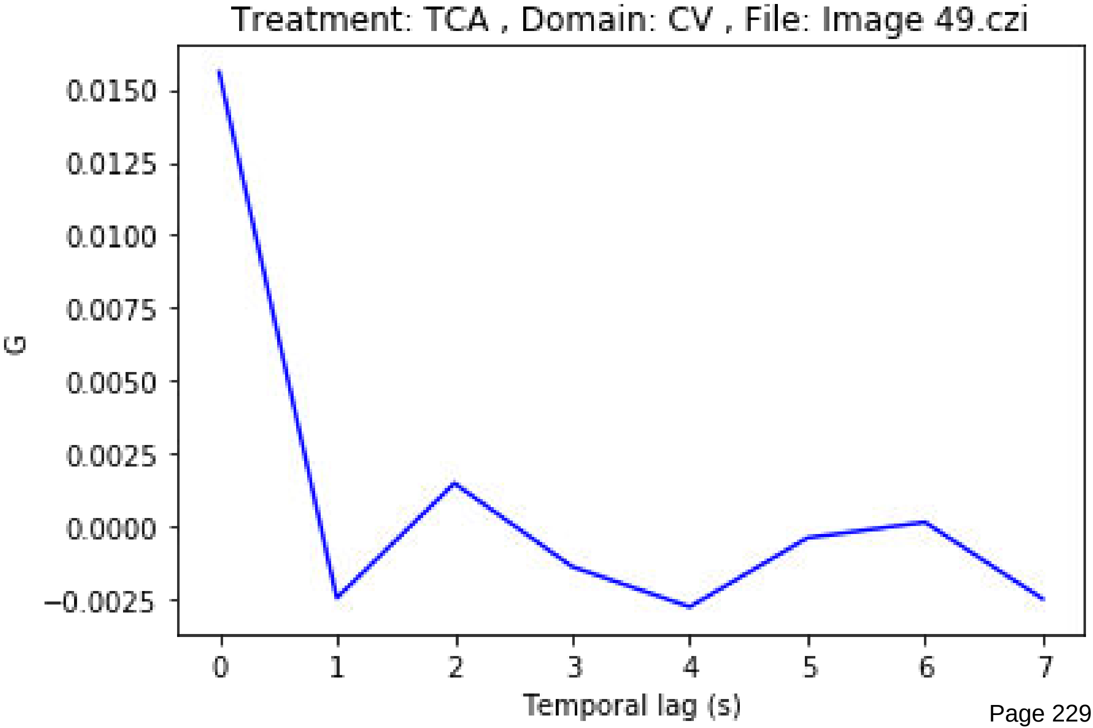
Treatment: TCA , Domain: CV , File: Image 49.czi
0.0150
0.0125
0.0100
0.0075
0.0050
0.0025
0.0000
−0.0025
G
0
1
2
3
4
5
6
7
Temporal lag (s)

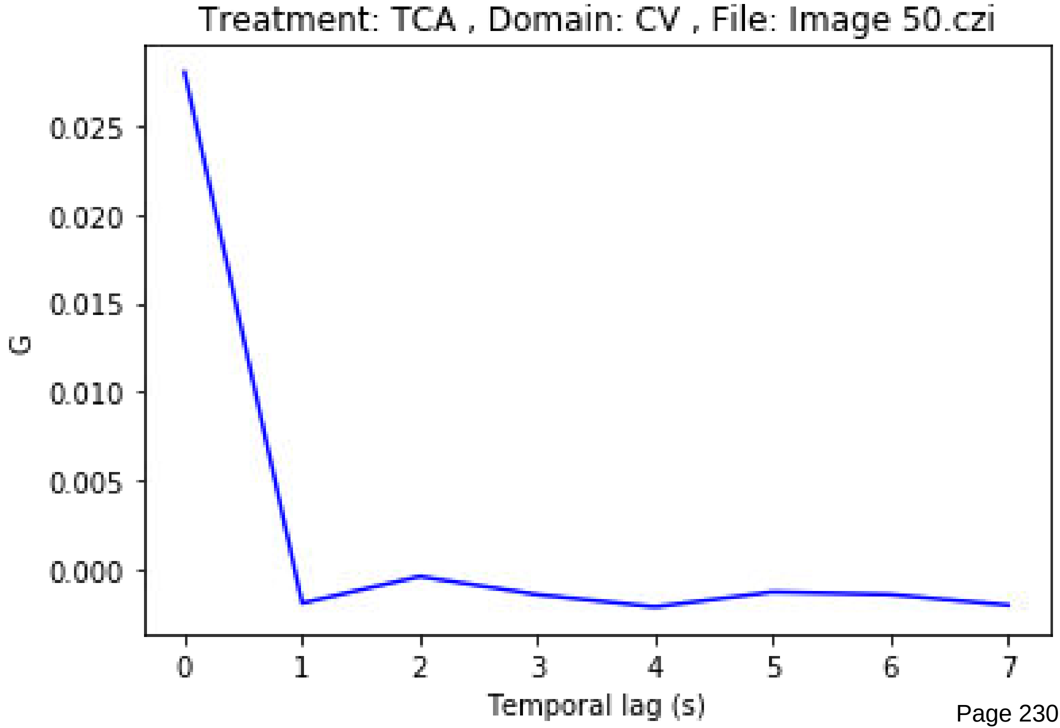
Treatment: TCA , Domain: CV , File: Image 50.czi
0.025
0.020
0.015
0.010
0.005
0.000
G
0
1
2
3
4
5
6
7
Temporal lag (s)

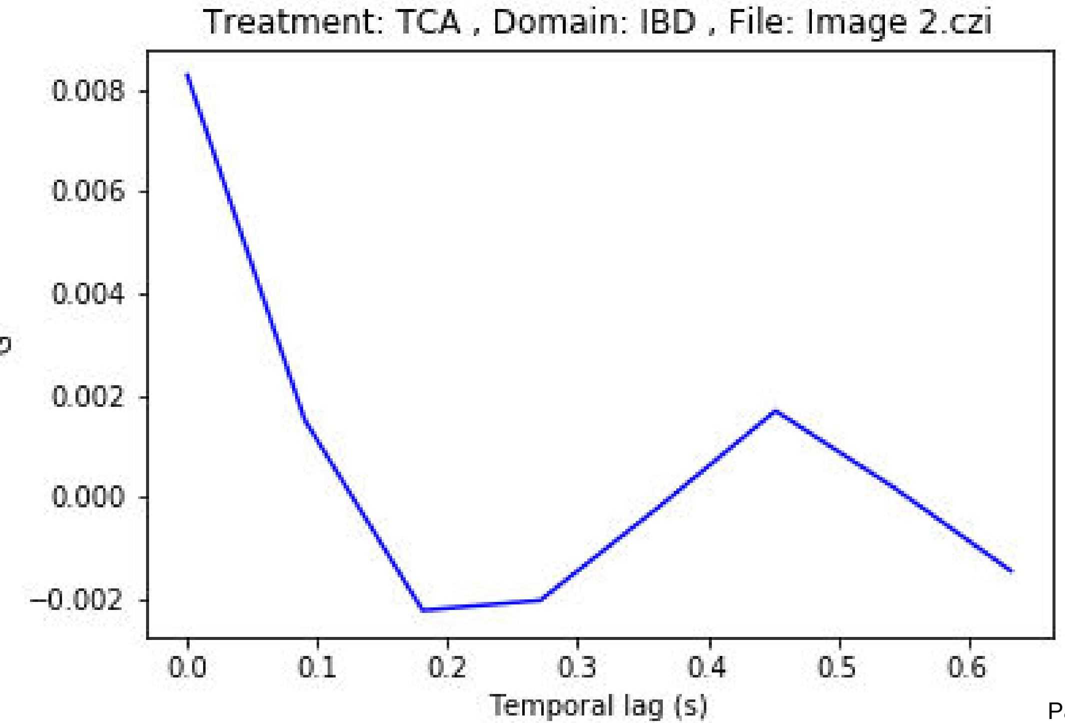
Treatment: TCA , Domain: IBD , File: Image 2.czi
0.008
0.006
0.004
0.002
0.000
−0.002
0.0
0.1
0.2
0.3
0.4
0.5
0.6
Temporal lag (s)

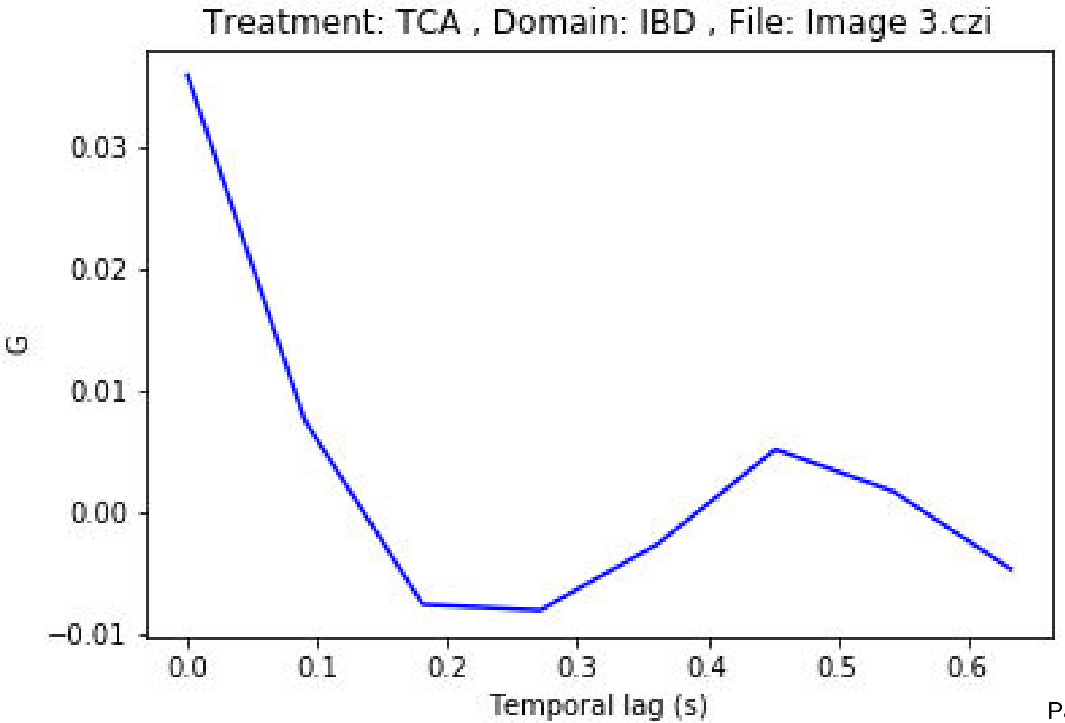
Treatment: TCA , Domain: IBD , File: Image 3.czi
G
0.03
0.02
0.01
0.00
−0.01
0.0
0.1
0.2
0.3
0.4
0.5
0.6
Temporal lag (s)

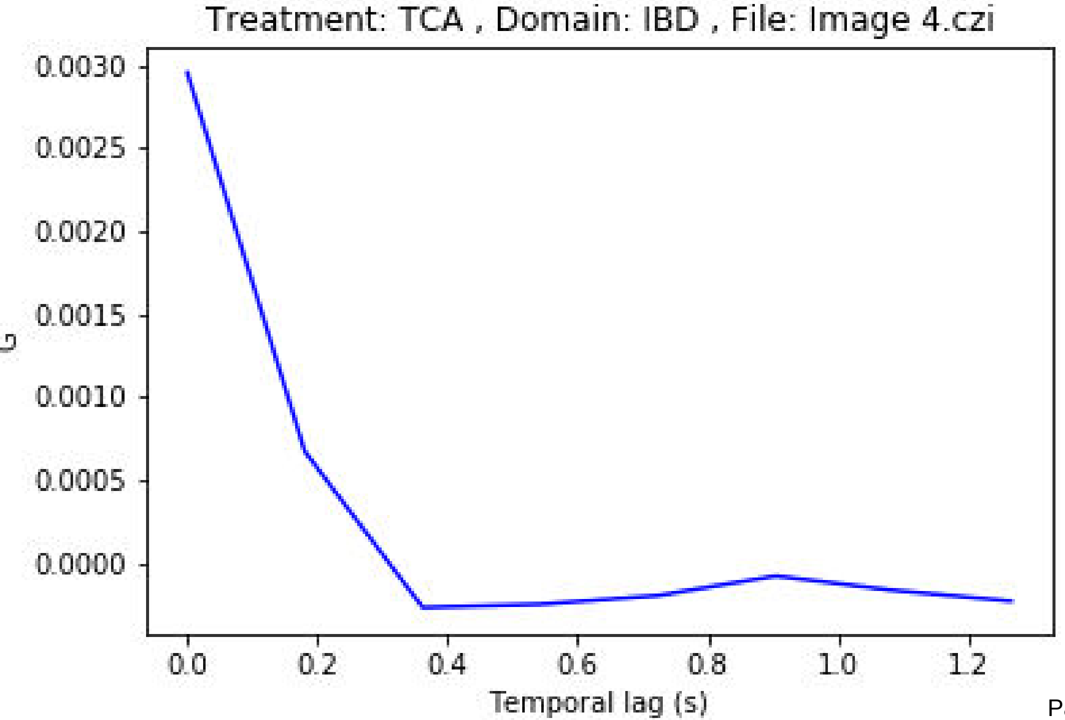
Treatment: TCA , Domain: IBD , File: Image 4.czi
0.0030
0.0025
0.0020
0.0015
0.0010
0.0005
0.0000
G
0.0
0.2
0.4
0.6
0.8
1.0
1.2
Temporal lag (s)

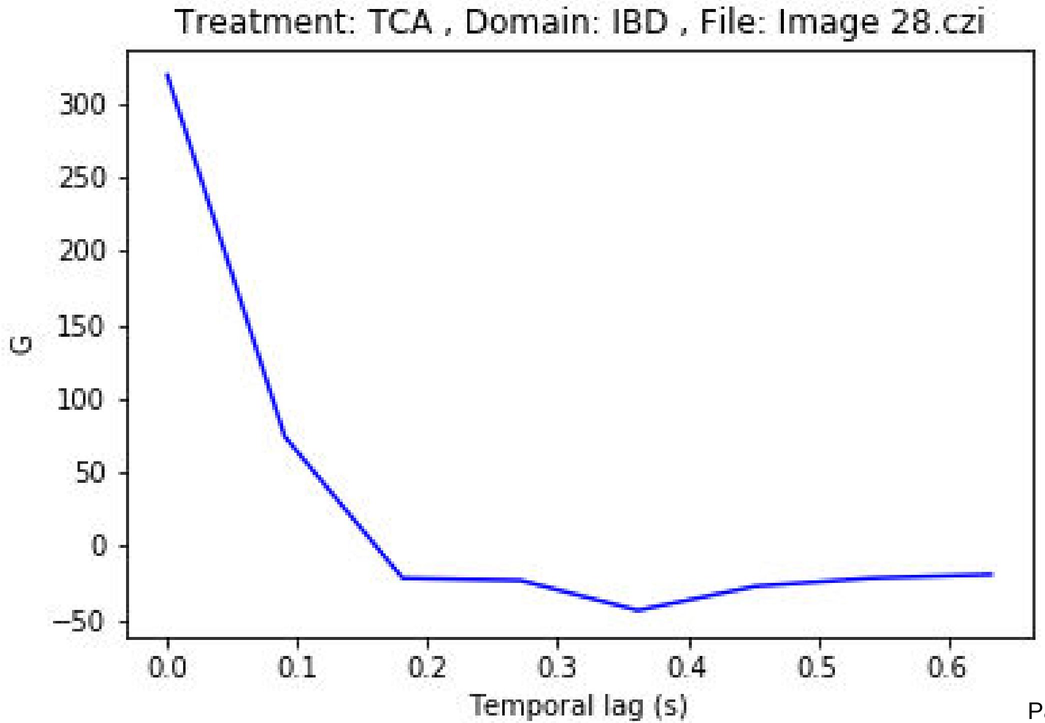

Treatment: TCA , Domain: IBD , File: Image 28.czi
G
300
250
200
150
100
50
0
−50
0.0
0.1
0.2
0.3
0.4
0.5
0.6
Temporal lag (s)

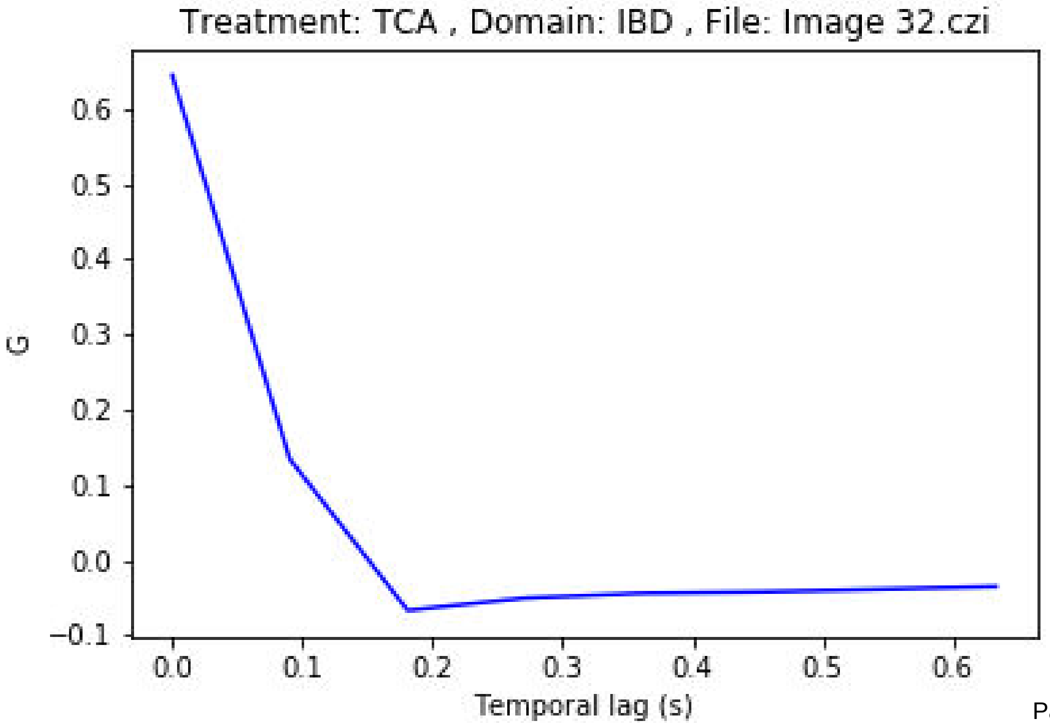
Treatment: TCA , Domain: IBD , File: Image 32.czi
G
0.6
0.5
0.4
0.3
0.2
0.1
0.0
−0.1
0.0
0.1
0.2
0.3
0.4
0.5
0.6
Temporal lag (s)

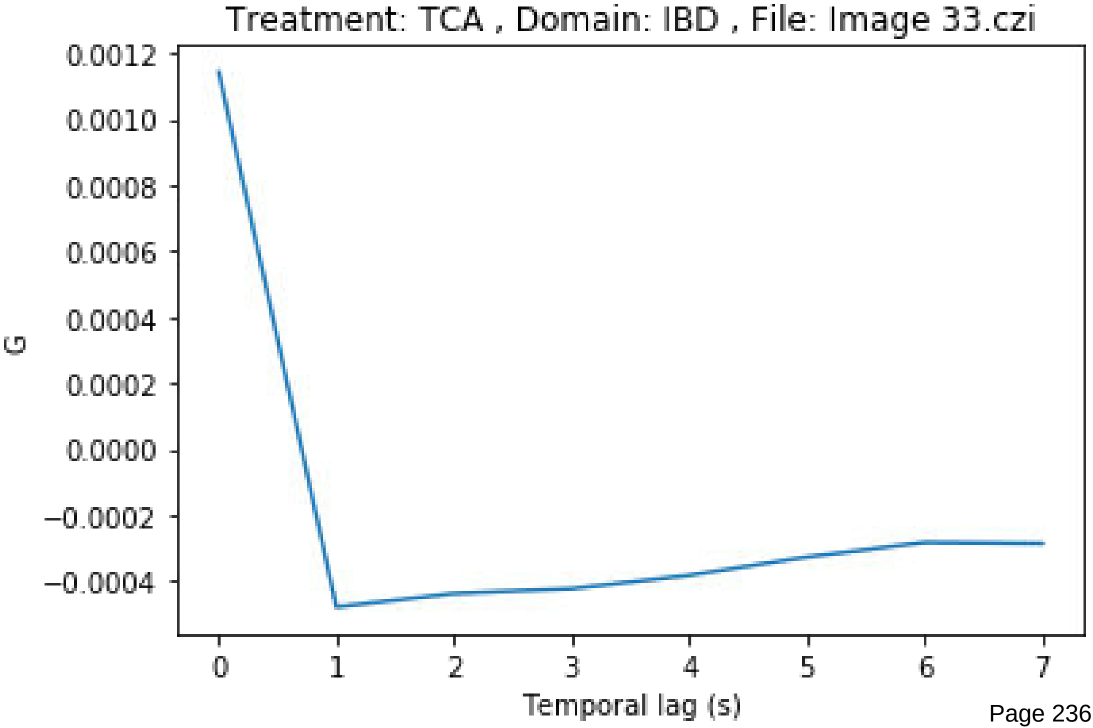
Treatment: TCA , Domain: IBD , File: Image 33.czi
0.0012
0.0010
0.0008
0.0006
0.0004
0.0002
0.0000
−0.0002
−0.0004
G
0
1
2
3
4
5
6
7
Temporal lag (s)

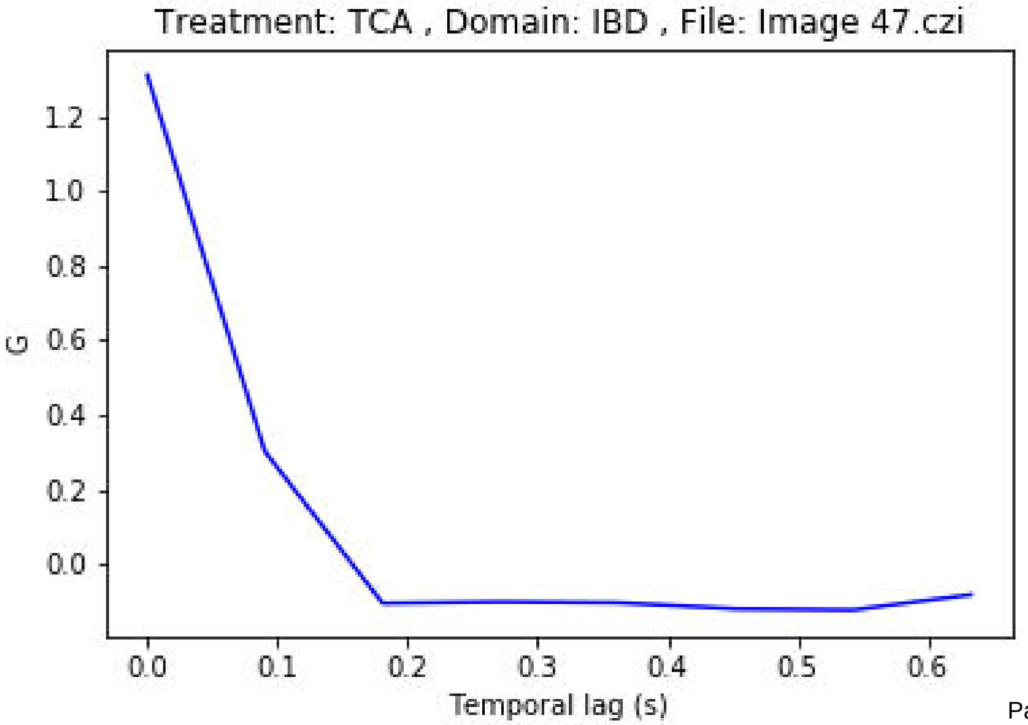

Treatment: TCA , Domain: IBD , File: Image 47.czi
G
1.2
1.0
0.8
0.6
0.4
0.2
0.0
0.0
0.1
0.2
0.3
0.4
0.5
0.6
Temporal lag (s)

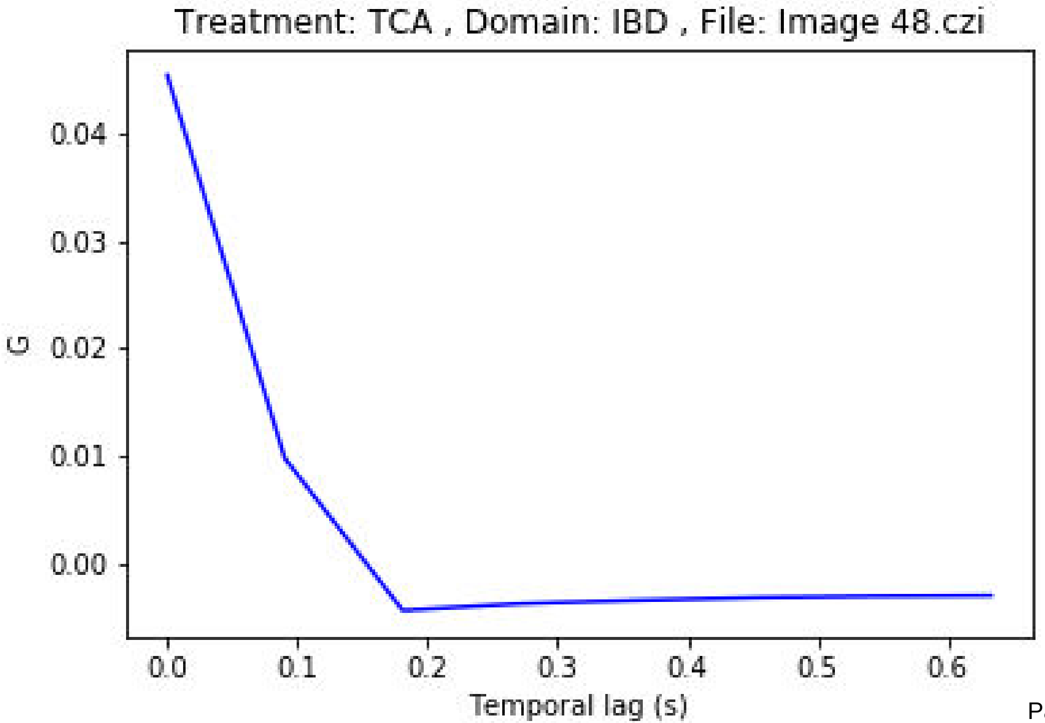

Treatment: TCA , Domain: IBD , File: Image 48.czi
0.04
0.03
0.02
0.01
0.00
G
0.0
0.1
0.2
0.3
0.4
0.5
0.6
Temporal lag (s)

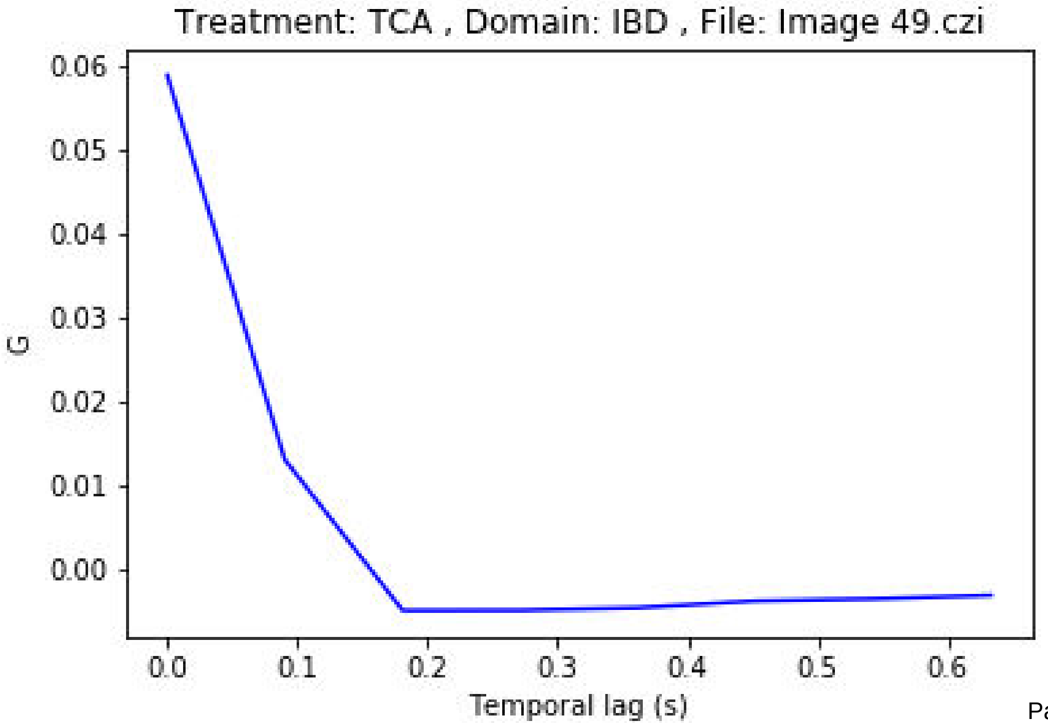
Treatment: TCA , Domain: IBD , File: Image 49.czi
0.06
0.05
0.04
0.03
0.02
0.01
0.00
G
0.0
0.1
0.2
0.3
0.4
0.5
0.6
Temporal lag (s)

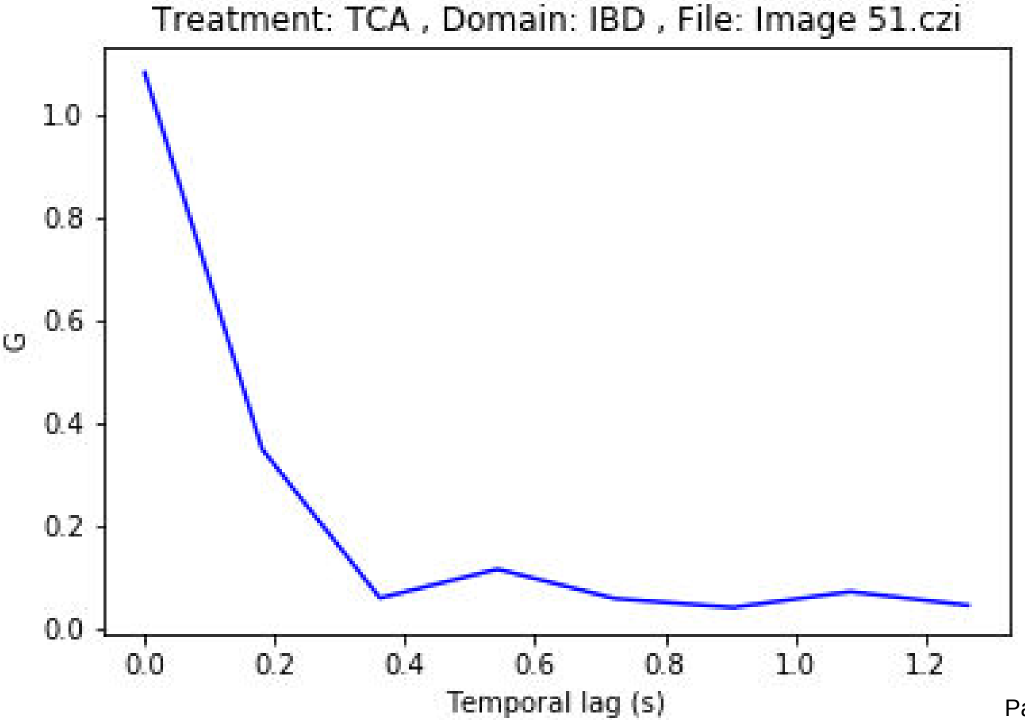

Treatment: TCA , Domain: IBD , File: Image 51.czi
G
1.0
0.8
0.6
0.4
0.2
0.0
0.0
0.2
0.4
0.6
0.8
1.0
1.2
Temporal lag (s)

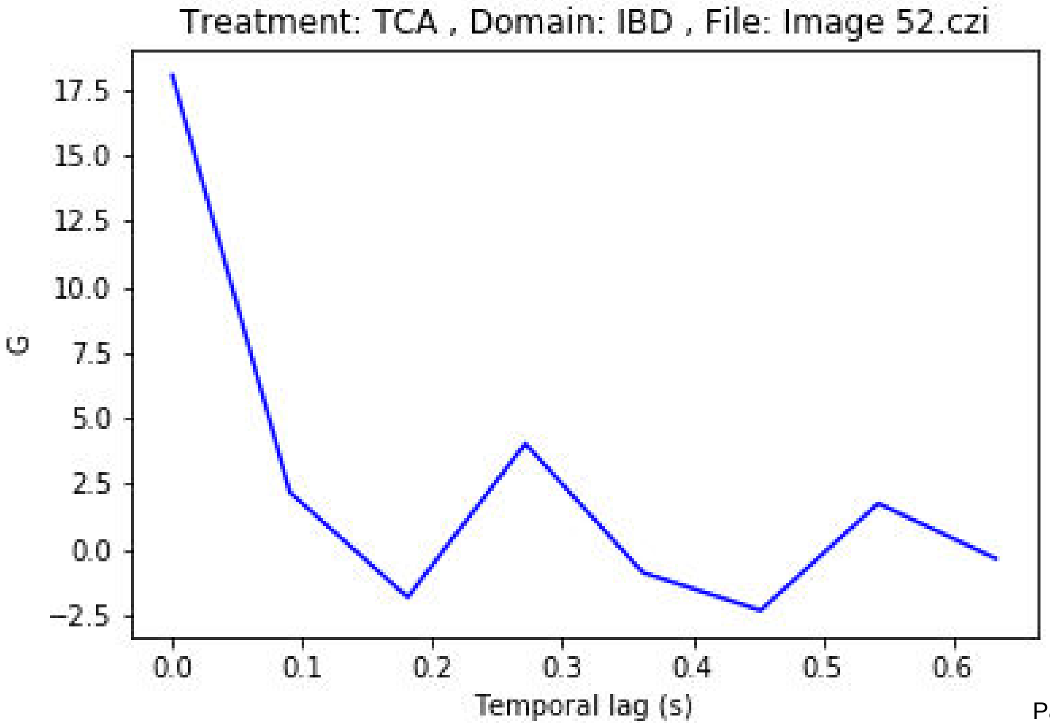

Treatment: TCA , Domain: IBD , File: Image 52.czi
G
17.5
15.0
12.5
10.0
7.5
5.0
2.5
0.0
−2.5
0.0
0.1
0.2
0.3
0.4
0.5
0.6
Temporal lag (s)

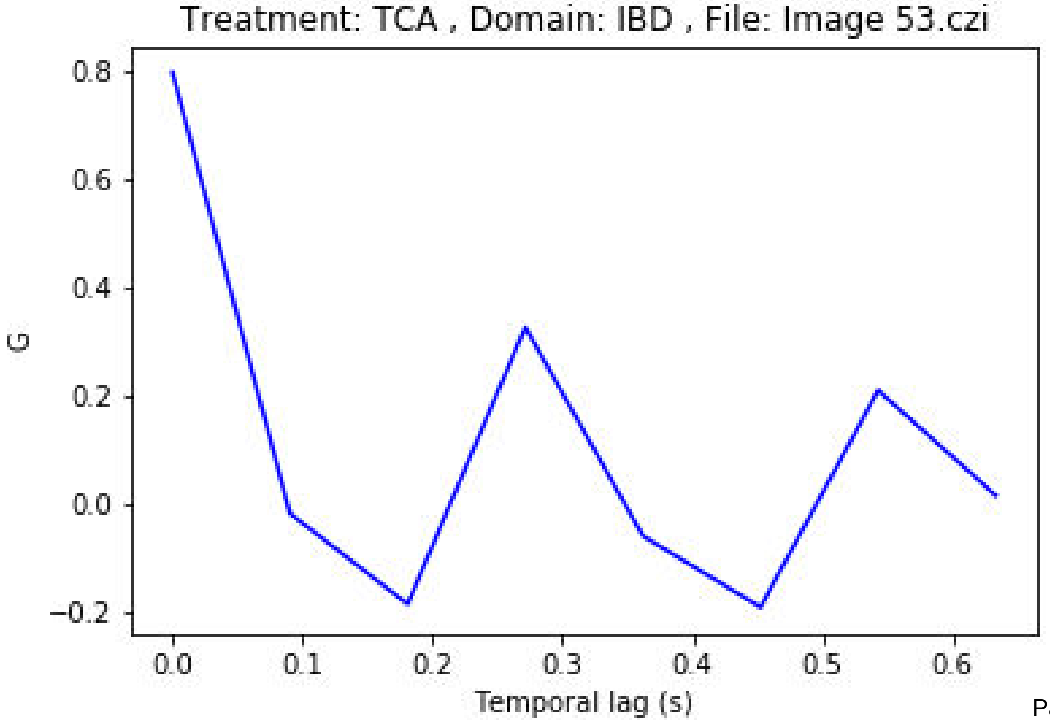

Treatment: TCA , Domain: IBD , File: Image 53.czi
0.8
0.6
0.4
0.2
0.0
−0.2
G
0.0
0.1
0.2
0.3
0.4
0.5
0.6
Temporal lag (s)

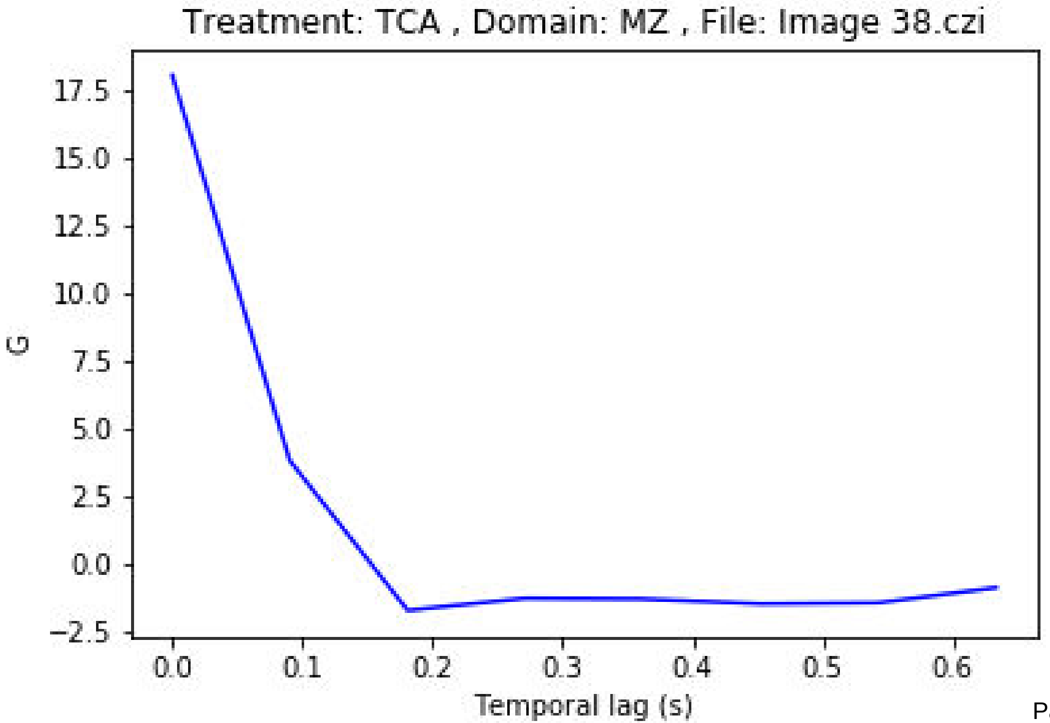

Treatment: TCA , Domain: MZ , File: Image 38.czi
G
Temporal lag (s)
17.5
15.0
12.5
10.0
7.5
5.0
2.5
0.0
−2.5
0.0
0.1
0.2
0.3
0.4
0.5
0.6

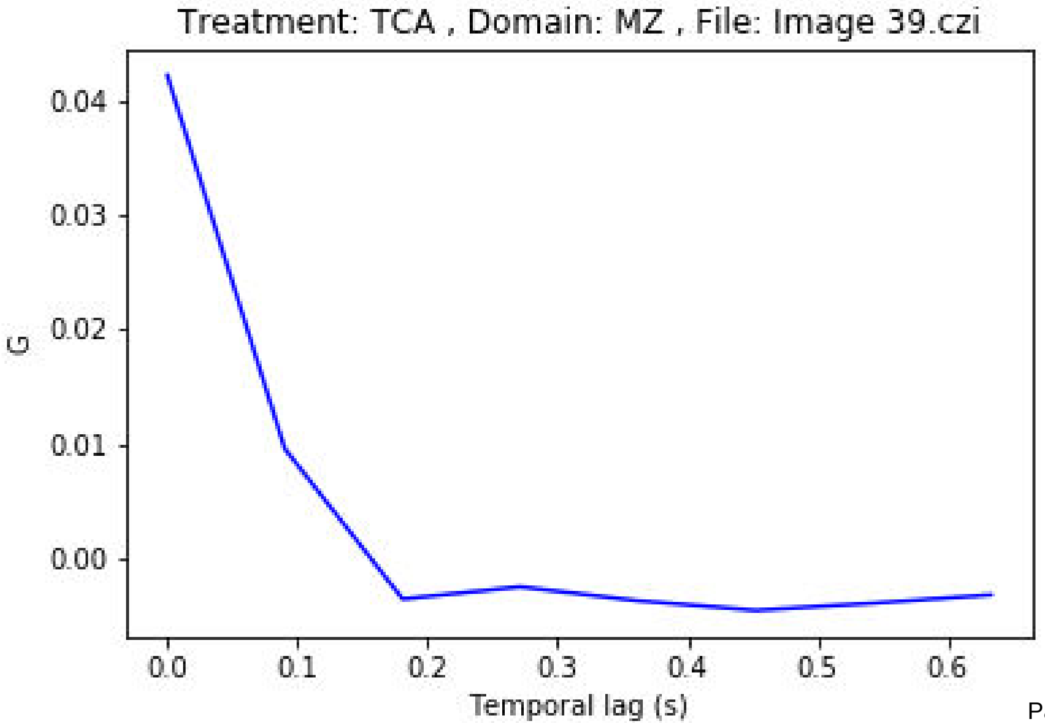

Treatment: TCA , Domain: MZ , File: Image 39.czi
G
0.04
0.03
0.02
0.01
0.00
0.0
0.1
0.2
0.3
0.4
0.5
0.6
Temporal lag (s)

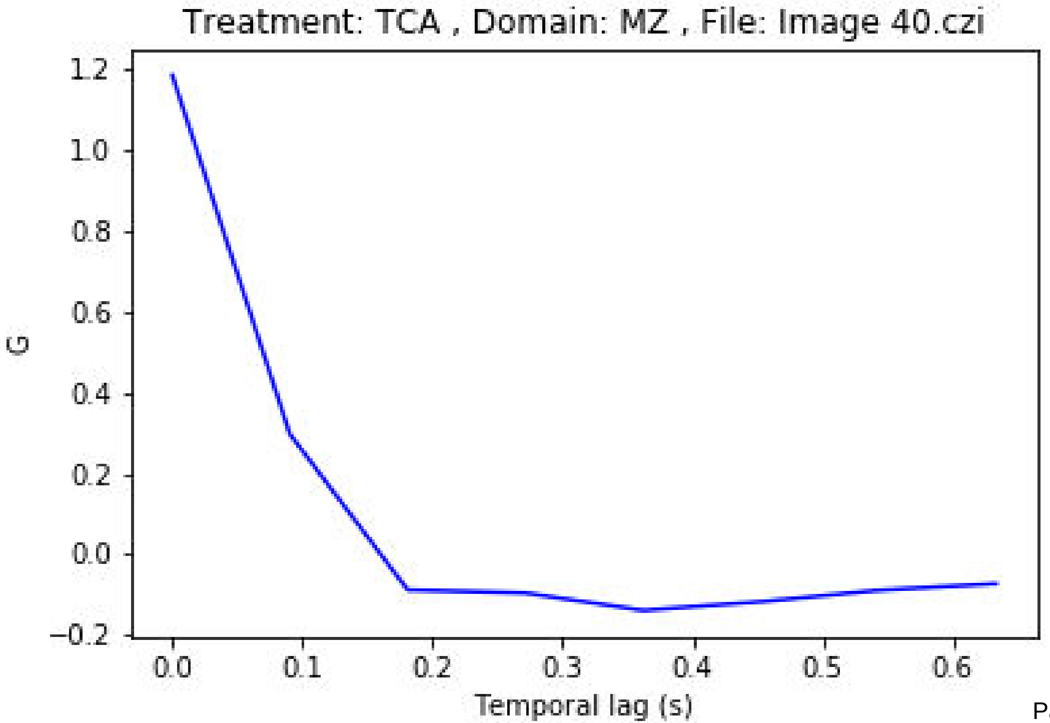

Treatment: TCA , Domain: MZ , File: Image 40.czi
G
1.2
1.0
0.8
0.6
0.4
0.2
0.0
−0.2
0.0
0.1
0.2
0.3
0.4
0.5
0.6
Temporal lag (s)

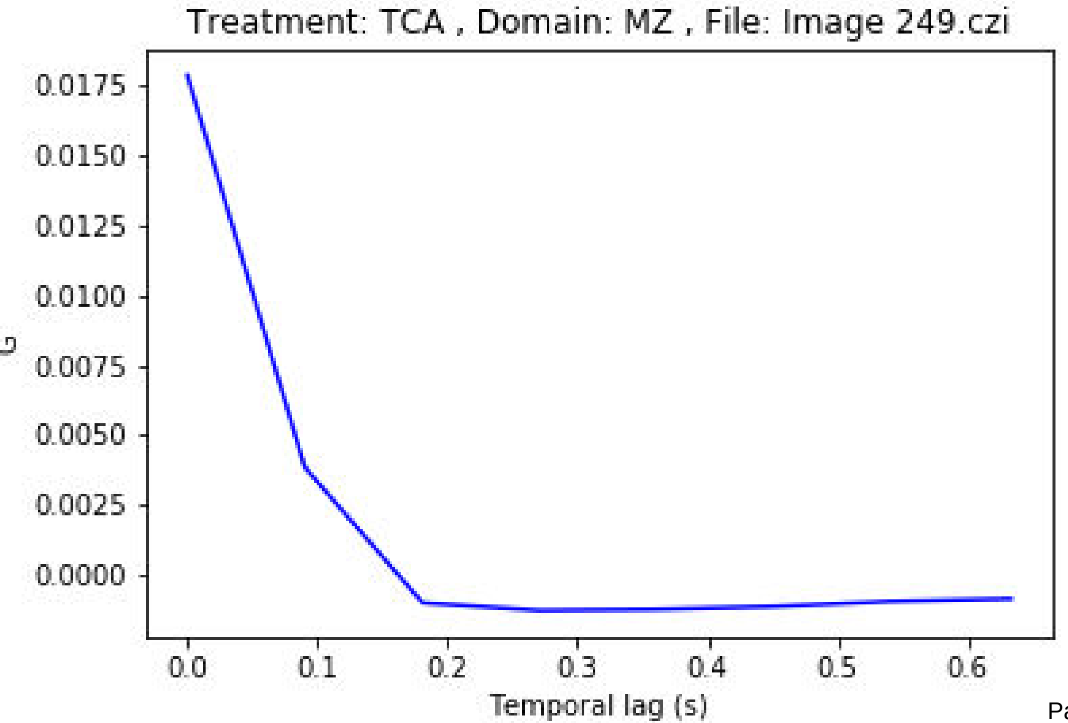

Treatment: TCA , Domain: MZ , File: Image 249.czi
G
0.0175
0.0150
0.0125
0.0100
0.0075
0.0050
0.0025
0.0000
0.0
0.1
0.2
0.3
0.4
0.5
0.6
Temporal lag (s)

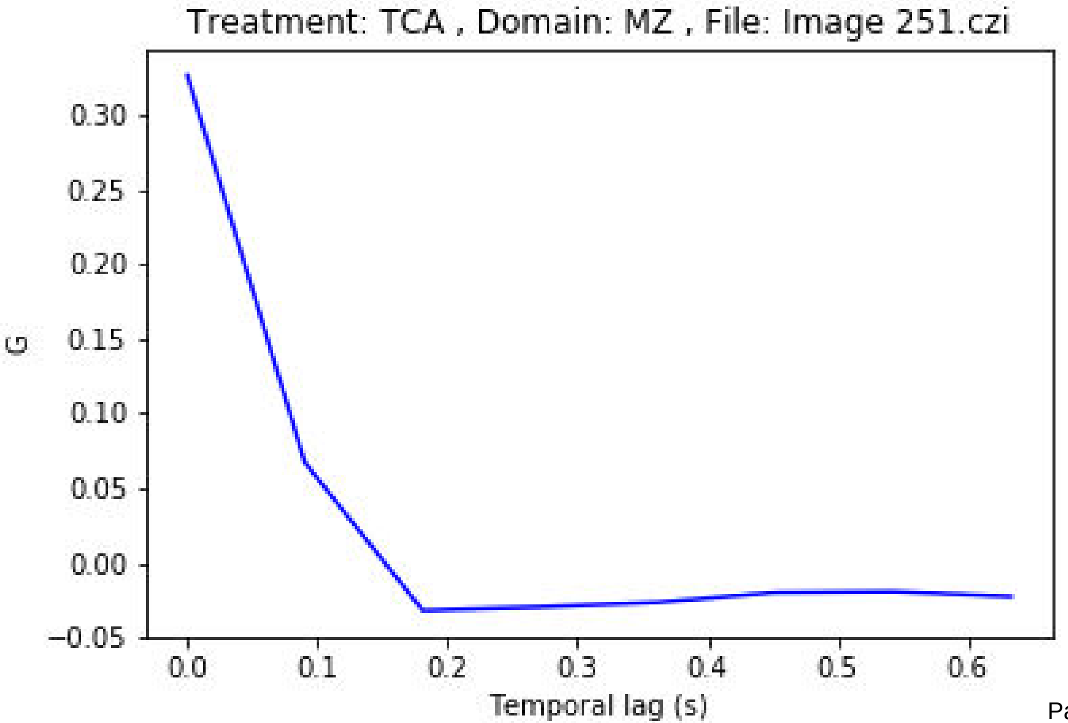
Treatment: TCA , Domain: MZ , File: Image 251.czi
G
0.30
0.25
0.20
0.15
0.10
0.05
0.00
−0.05
0.0
0.1
0.2
0.3
0.4
0.5
0.6
Temporal lag (s)

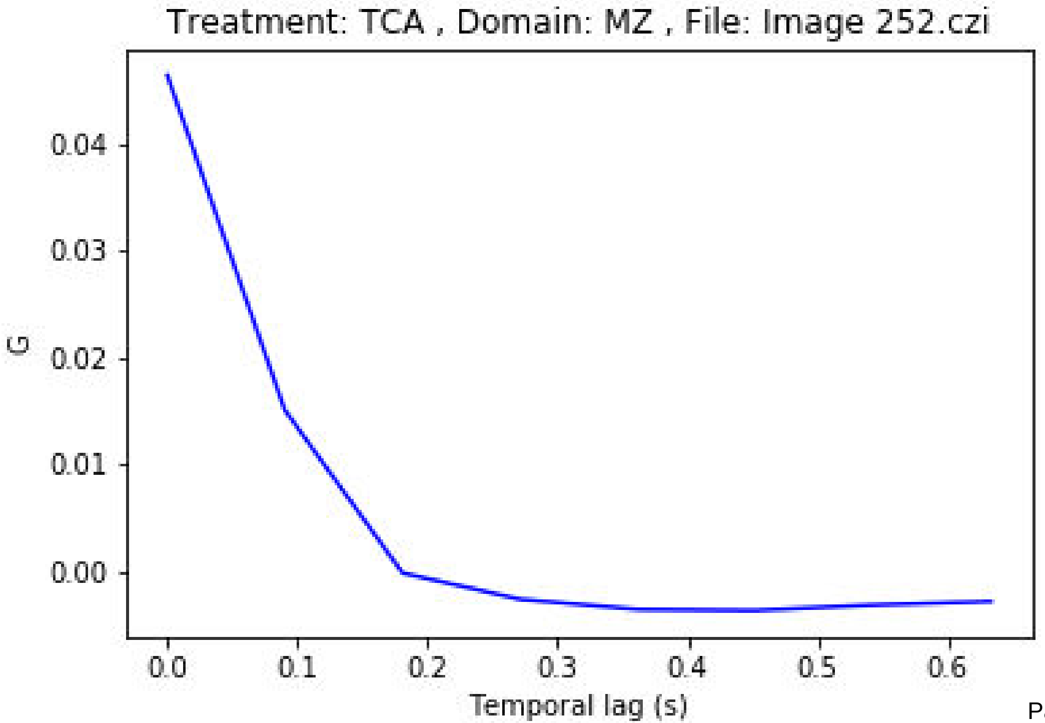

Treatment: TCA , Domain: MZ , File: Image 252.czi
G
0.04
0.03
0.02
0.01
0.00
0.0
0.1
0.2
0.3
0.4
0.5
0.6
Temporal lag (s)

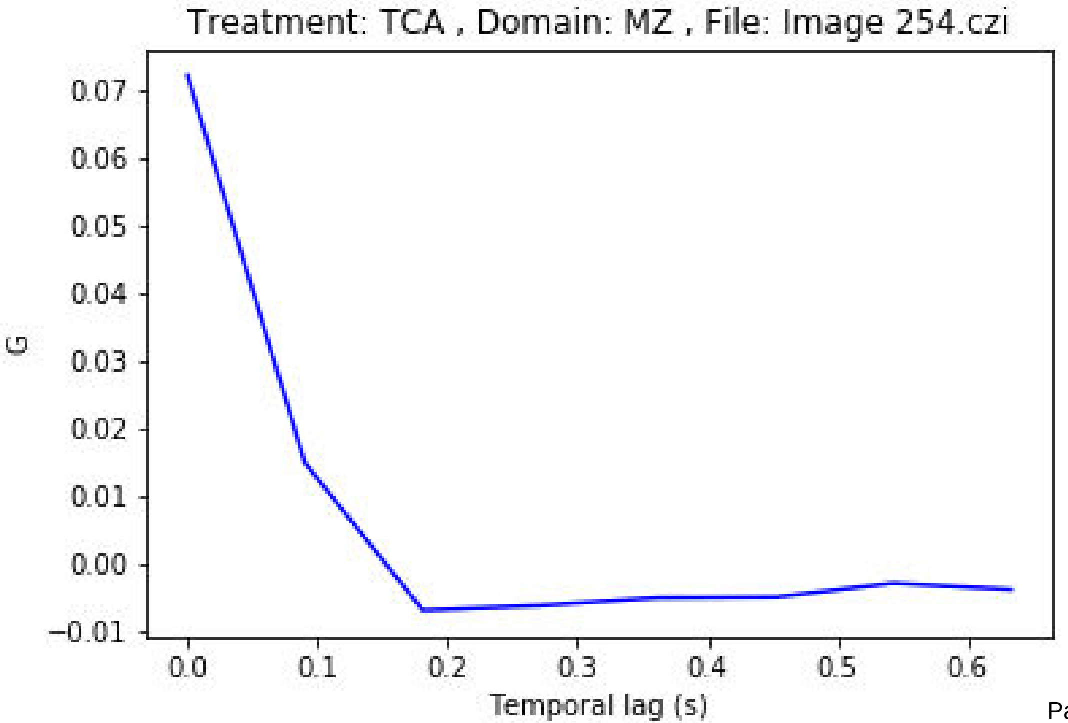

Treatment: TCA , Domain: MZ , File: Image 254.czi
G
0.07
0.06
0.05
0.04
0.03
0.02
0.01
0.00
−0.01
0.0
0.1
0.2
0.3
0.4
0.5
0.6
Temporal lag (s)

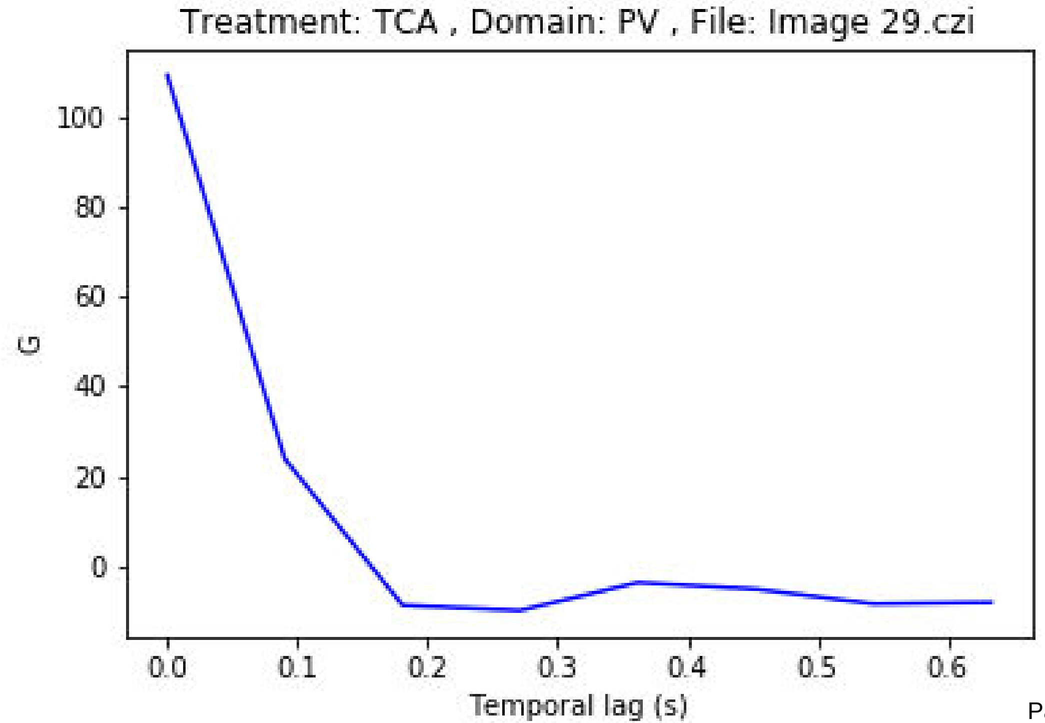

Treatment: TCA , Domain: PV , File: Image 29.czi
100
80
60
40
20
0
G
0.0
0.1
0.2
0.3
0.4
0.5
0.6
Temporal lag (s)

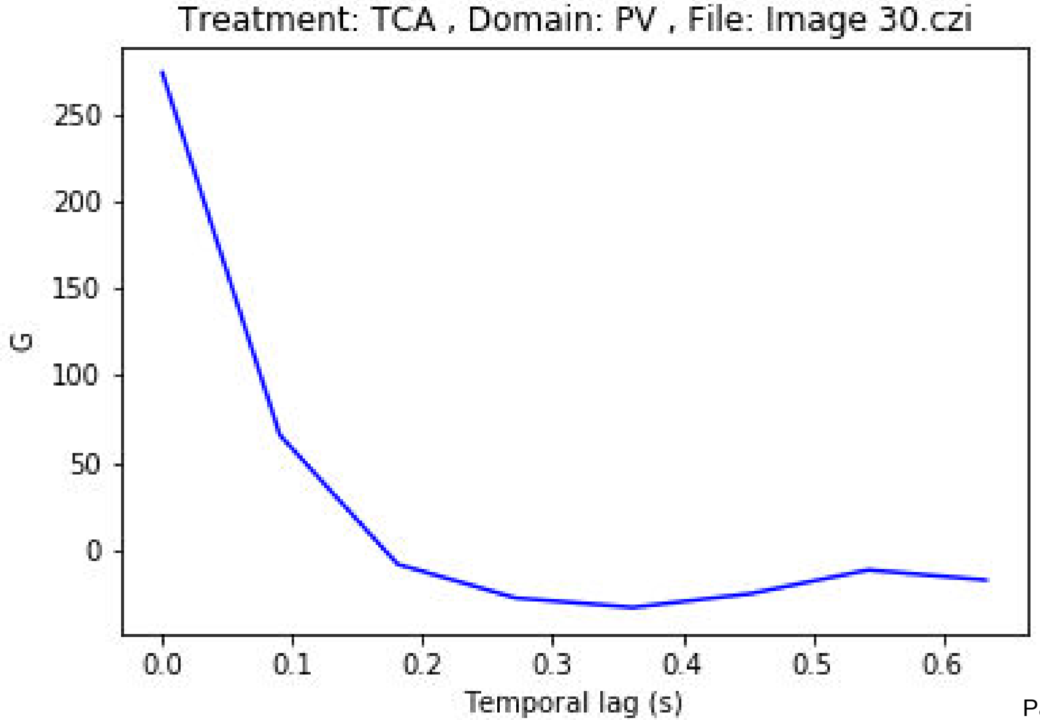
Treatment: TCA , Domain: PV , File: Image 30.czi
250
200
150
100
50
0
G
0.0
0.1
0.2
0.3
0.4
0.5
0.6
Temporal lag (s)

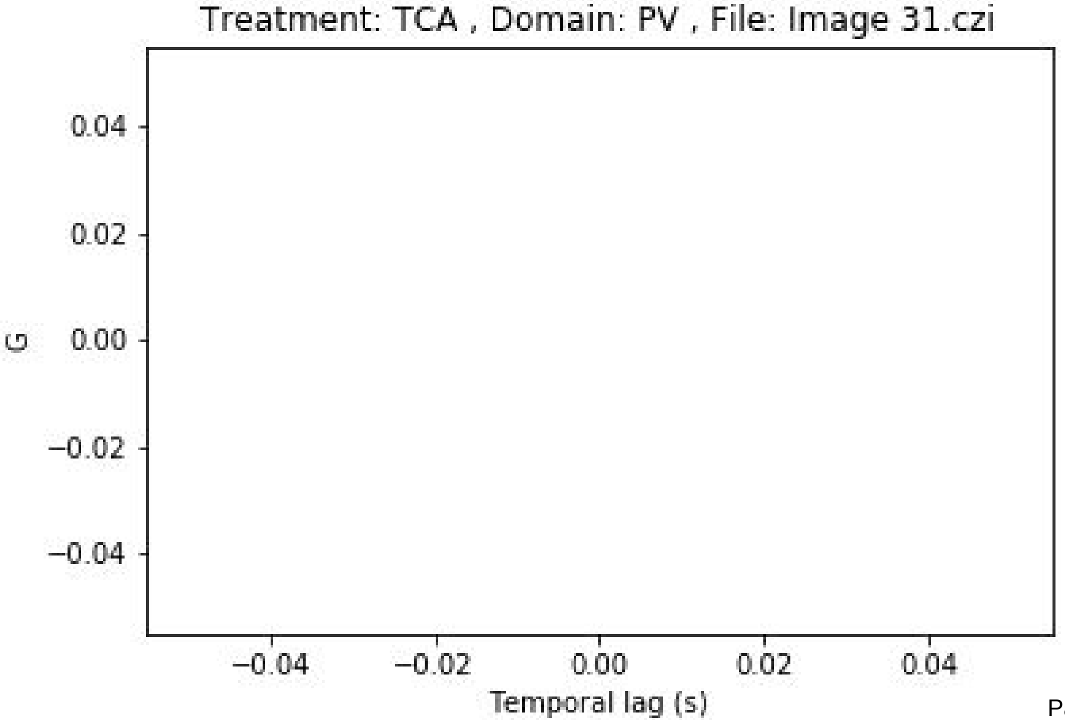
Treatment: TCA , Domain: PV , File: Image 31.czi
0.04
0.02
0.00
−0.02
−0.04
G
−0.04
−0.02
0.00
0.02
0.04
Temporal lag (s)

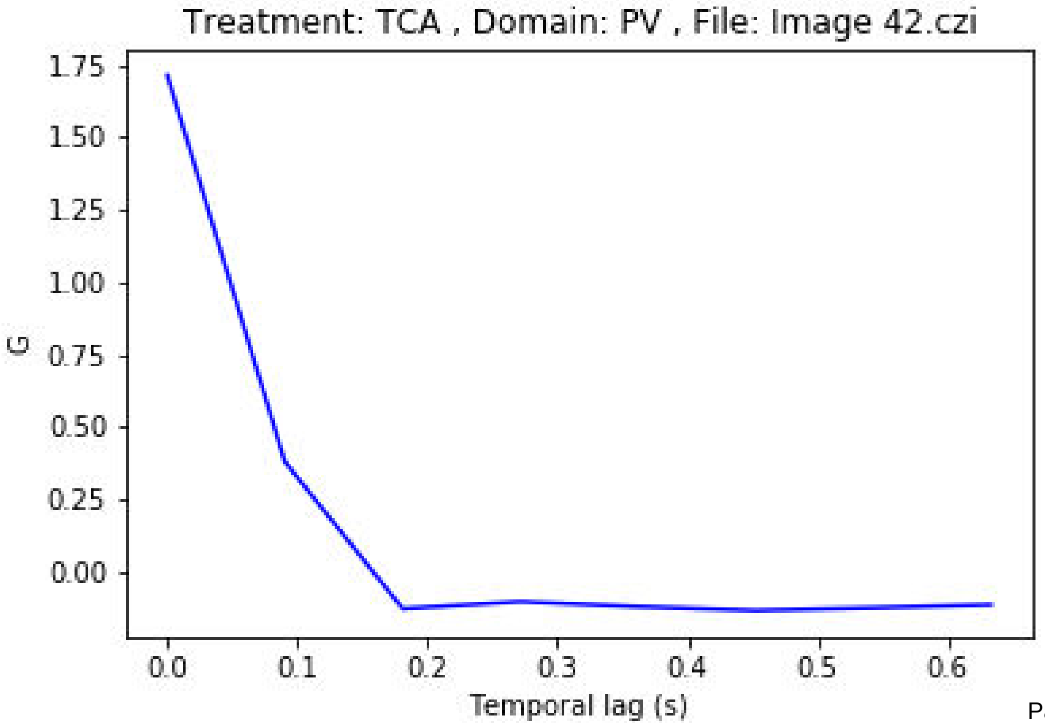
Treatment: TCA , Domain: PV , File: Image 42.czi
1.75
1.50
1.25
1.00
0.75
0.50
0.25
0.00
G
0.0
0.1
0.2
0.3
0.4
0.5
0.6
Temporal lag (s)

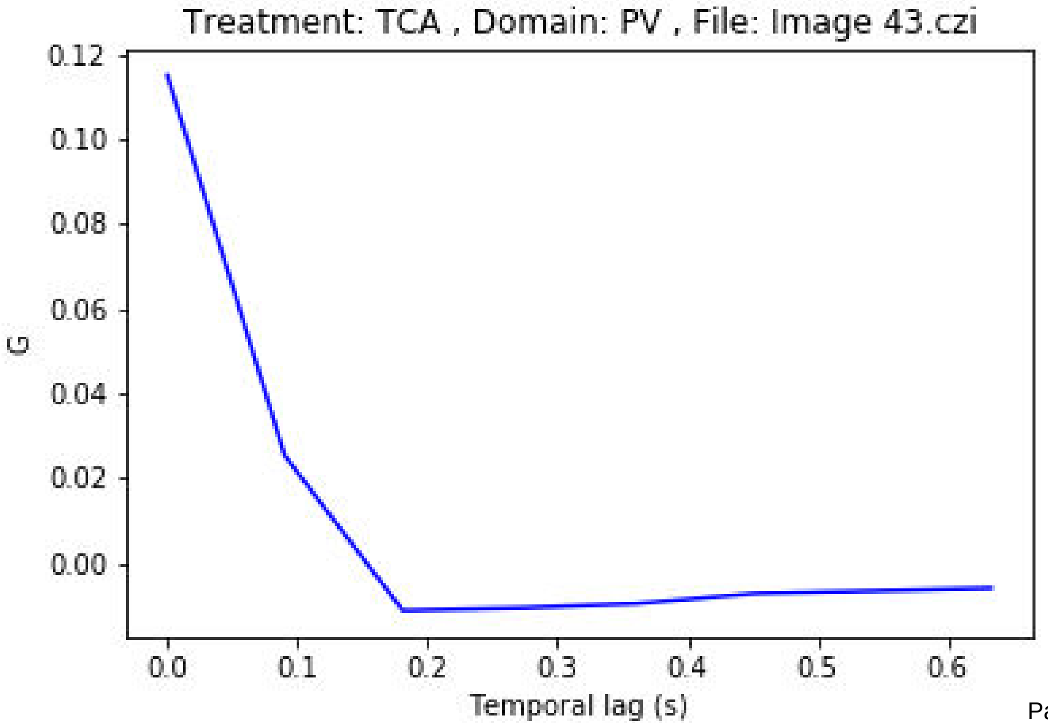
Treatment: TCA , Domain: PV , File: Image 43.czi
0.12
0.10
0.08
0.06
0.04
0.02
0.00
G
0.0
0.1
0.2
0.3
0.4
0.5
0.6
Temporal lag (s)

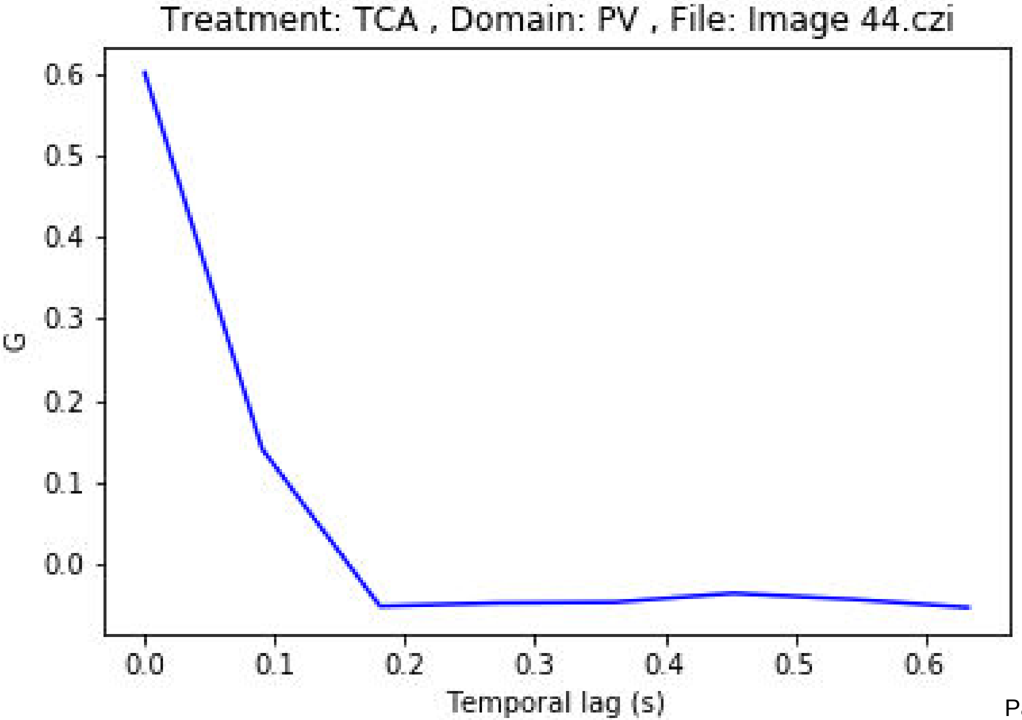
Treatment: TCA , Domain: PV , File: Image 44.czi
G
0.6
0.5
0.4
0.3
0.2
0.1
0.0
0.0
0.1
0.2
0.3
0.4
0.5
0.6
Temporal lag (s)

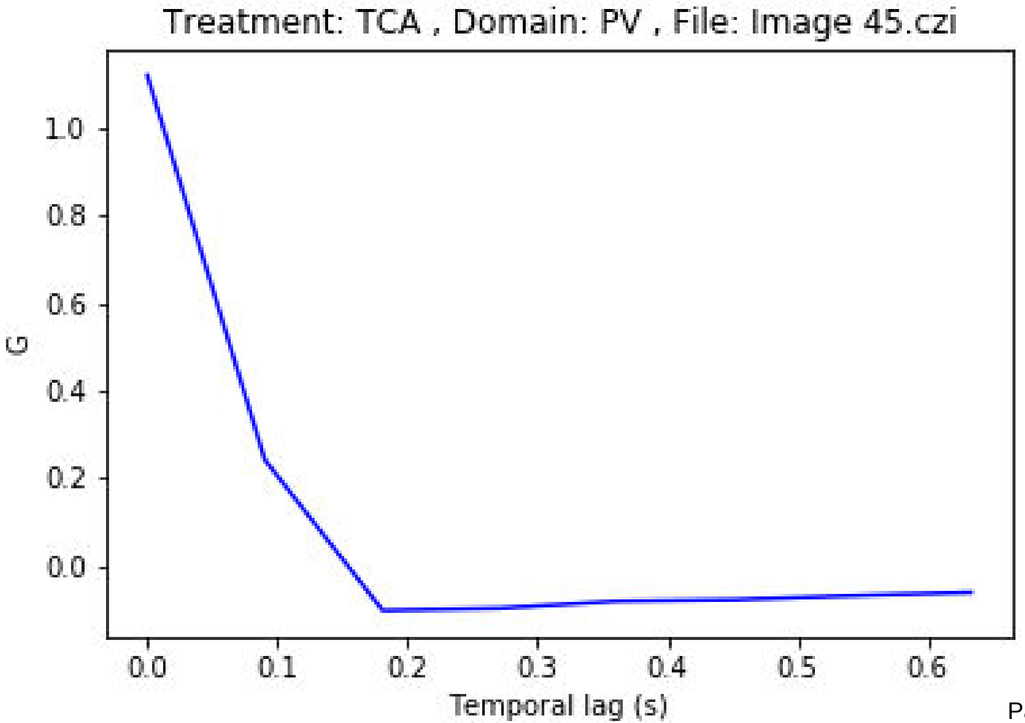
Treatment: TCA , Domain: PV , File: Image 45.czi
G
1.0
0.8
0.6
0.4
0.2
0.0
0.0
0.1
0.2
0.3
0.4
0.5
0.6
Temporal lag (s)

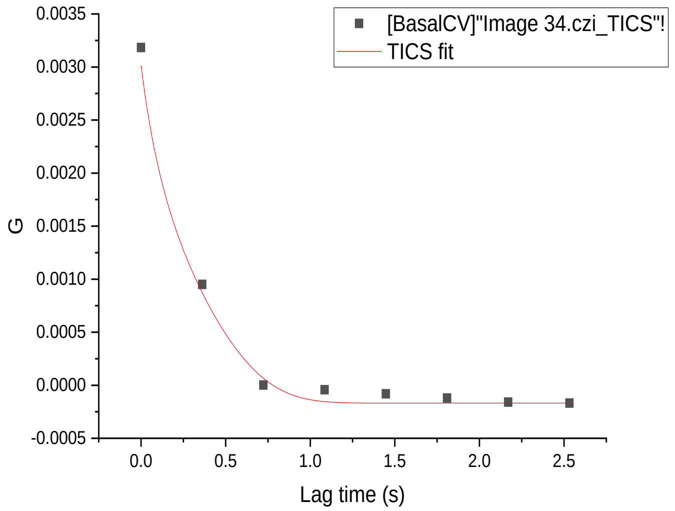
[BasalCV]"Image 34.czi_TICS"!
TICS fit
G
Lag time (s)
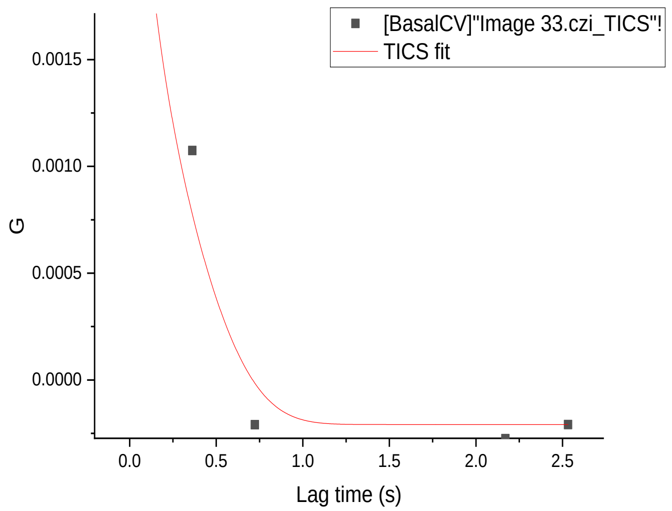
[BasalCV]"Image 33.czi_TICS"!
TICS fit
G
Lag time (s)
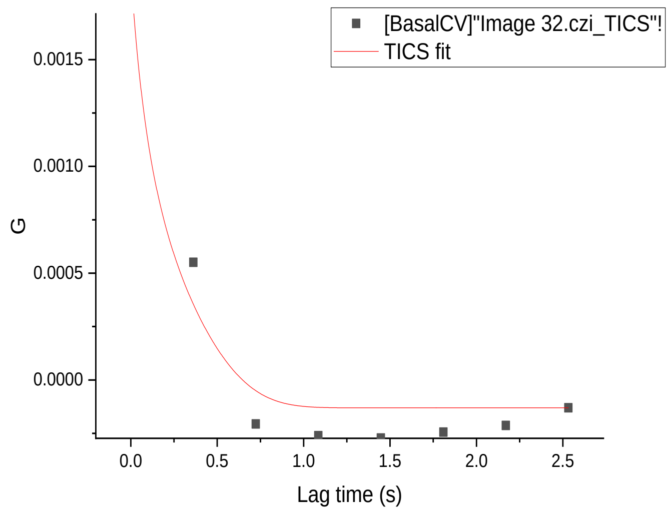
[BasalCV]"Image 32.czi_TICS"!
TICS fit
G
Lag time (s)
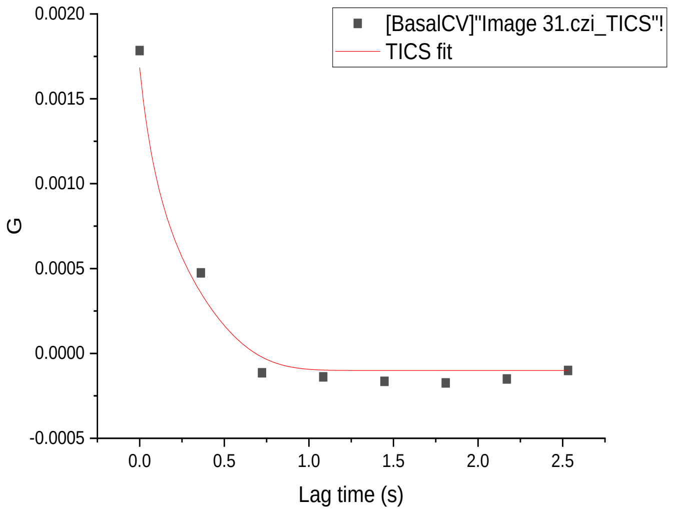
[BasalCV]"Image 31.czi_TICS"!
TICS fit
G
Lag time (s)
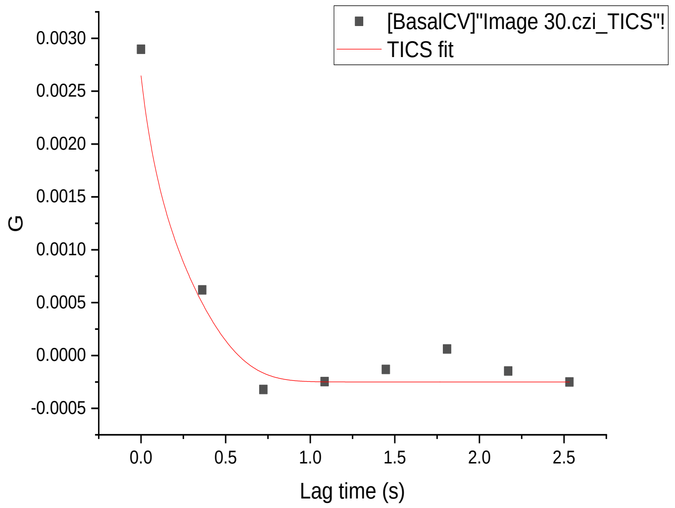
[BasalCV]"Image 30.czi_TICS"!
TICS fit
G
Lag time (s)
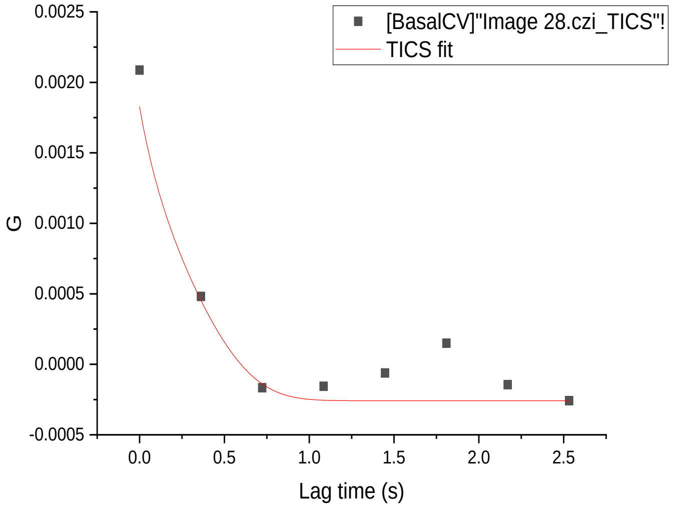
[BasalCV]"Image 28.czi_TICS"!
TICS fit
G
Lag time (s)
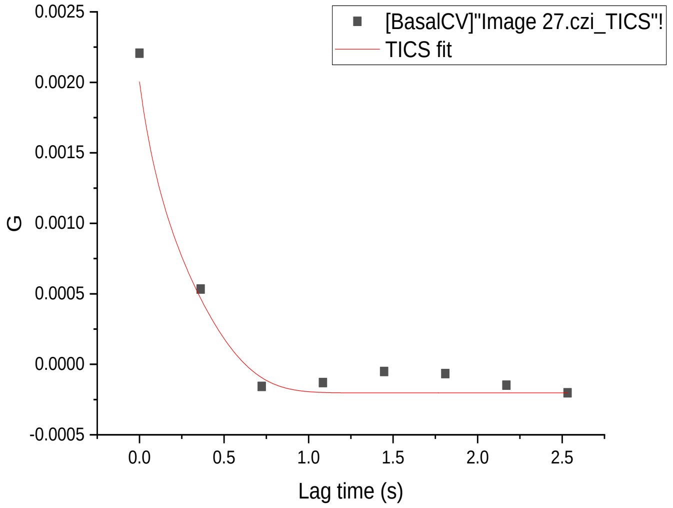
[BasalCV]"Image 27.czi_TICS"!
TICS fit
G
Lag time (s)
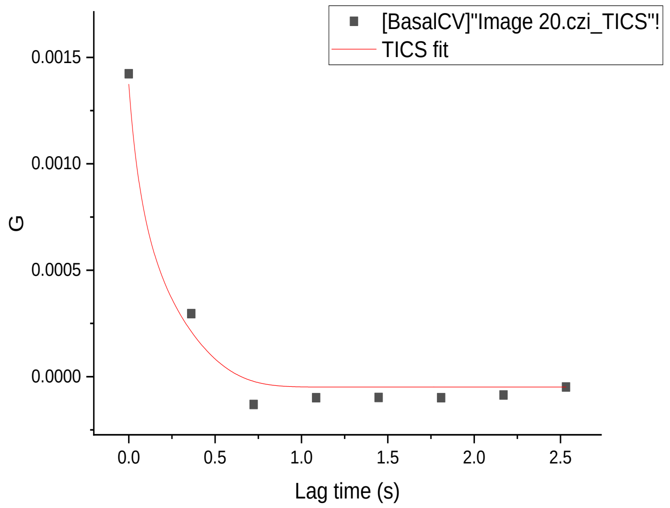
[BasalCV]"Image 20.czi_TICS"!
TICS fit
G
Lag time (s)
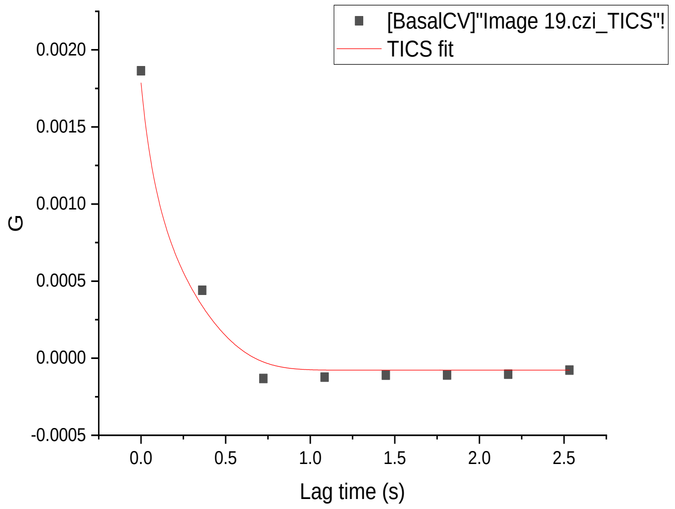
[BasalCV]"Image 19.czi_TICS"!
TICS fit
G
Lag time (s)
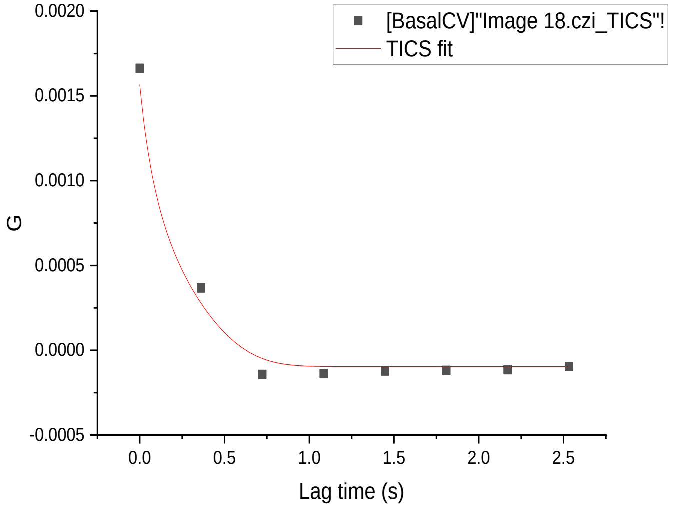
[BasalCV]"Image 18.czi_TICS"!
TICS fit
G
Lag time (s)
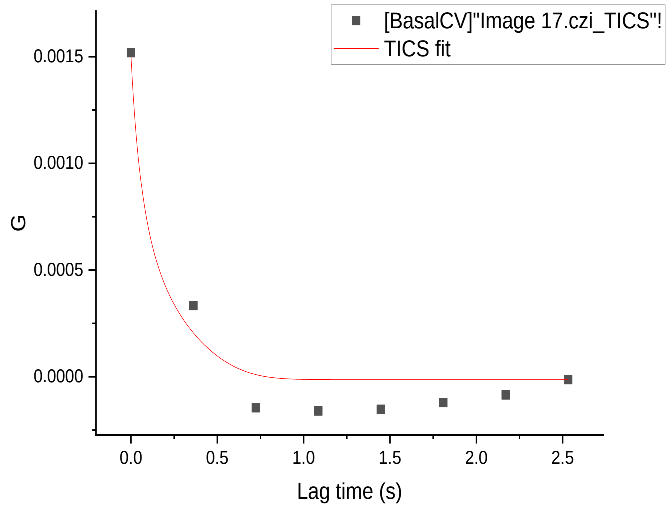
[BasalCV]"Image 17.czi_TICS"!
TICS fit
G
Lag time (s)
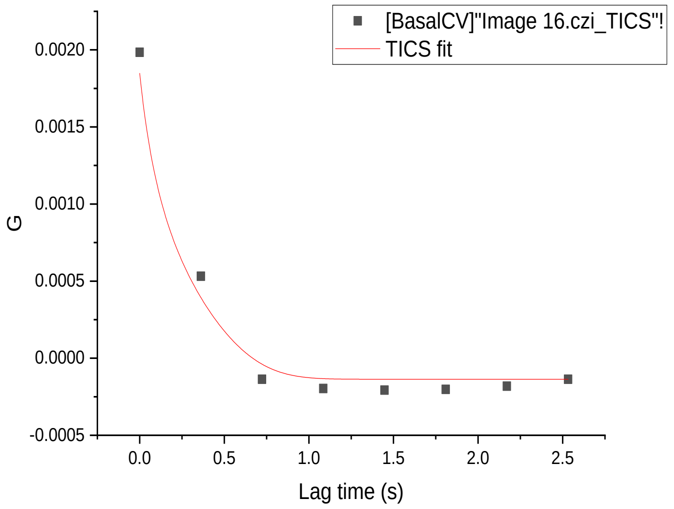
[BasalCV]"Image 16.czi_TICS"!
TICS fit
G
Lag time (s)

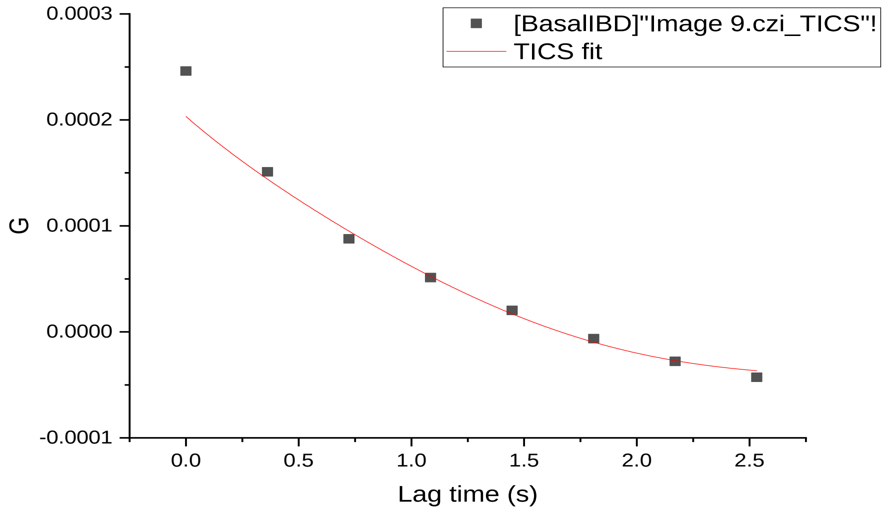
[BasalIBD]"Image 9.czi_TICS"!
TICS fit
G
0.0003
0.0002
0.0001
0.0000
-0.0001
0.0 0.5 1.0 1.5 2.0 2.5
Lag time (s)

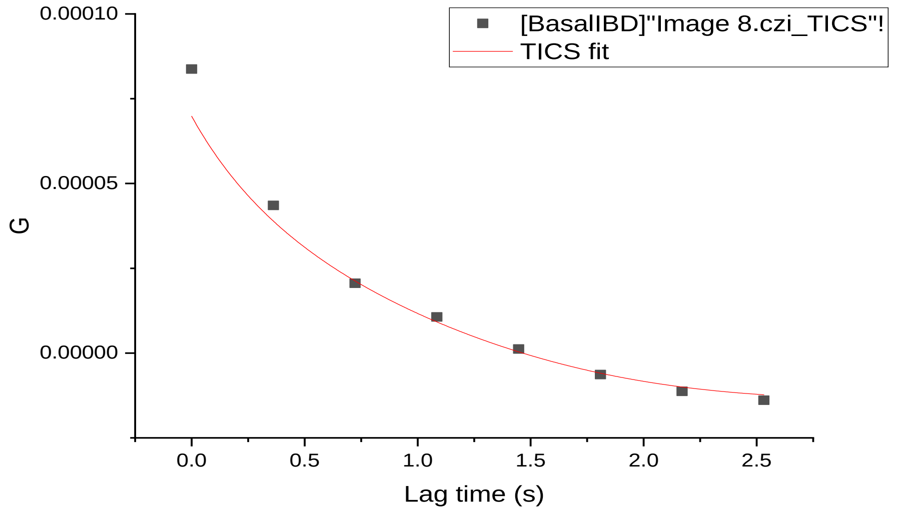
[BasalIBD]"Image 8.czi_TICS"!
TICS fit
G
0.00010
0.00005
0.00000
0.0 0.5 1.0 1.5 2.0 2.5
Lag time (s)

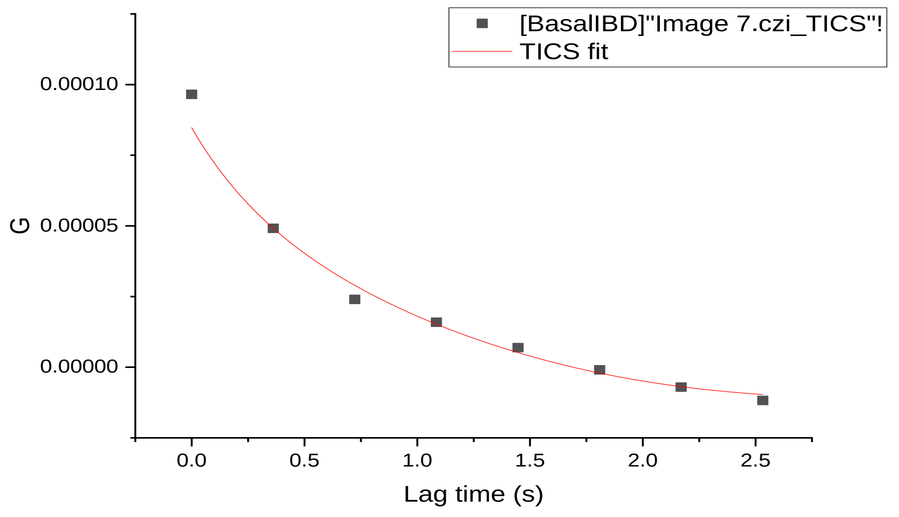
[BasalIBD]"Image 7.czi_TICS"!
TICS fit
G
0.00010
0.00005
0.00000
0.0 0.5 1.0 1.5 2.0 2.5
Lag time (s)

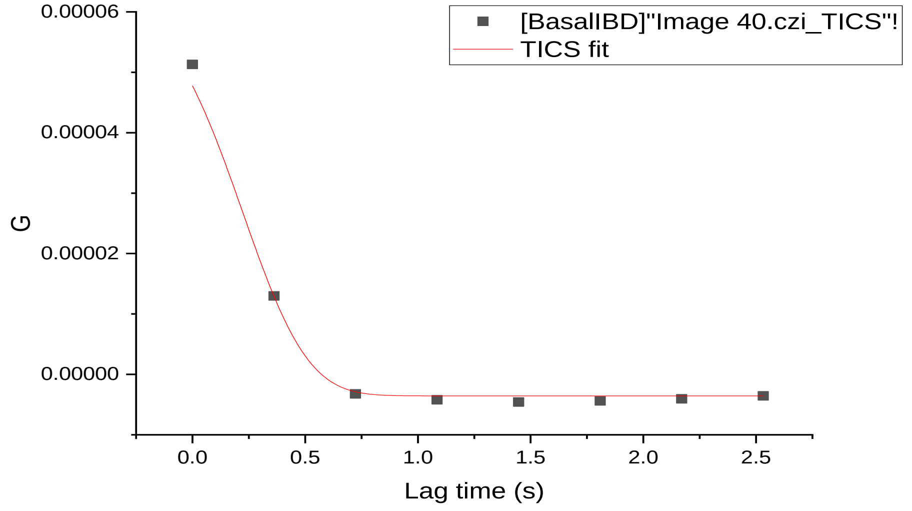
[BasalIBD]"Image 40.czi_TICS"!
TICS fit
G
0.00006
0.00004
0.00002
0.00000
0.0 0.5 1.0 1.5 2.0 2.5
Lag time (s)

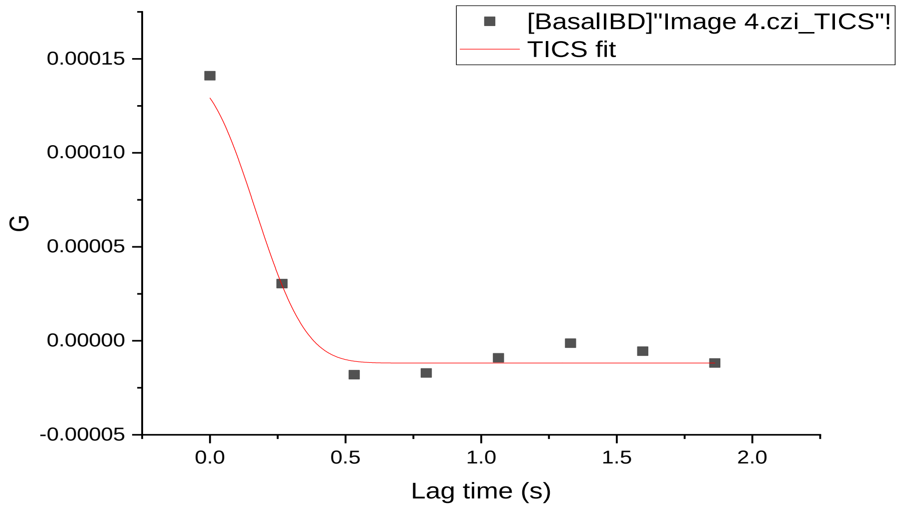
[BasalIBD]"Image 4.czi_TICS"!
TICS fit
G
0.00015
0.00010
0.00005
0.00000
-0.00005
0.0 0.5 1.0 1.5 2.0
Lag time (s)

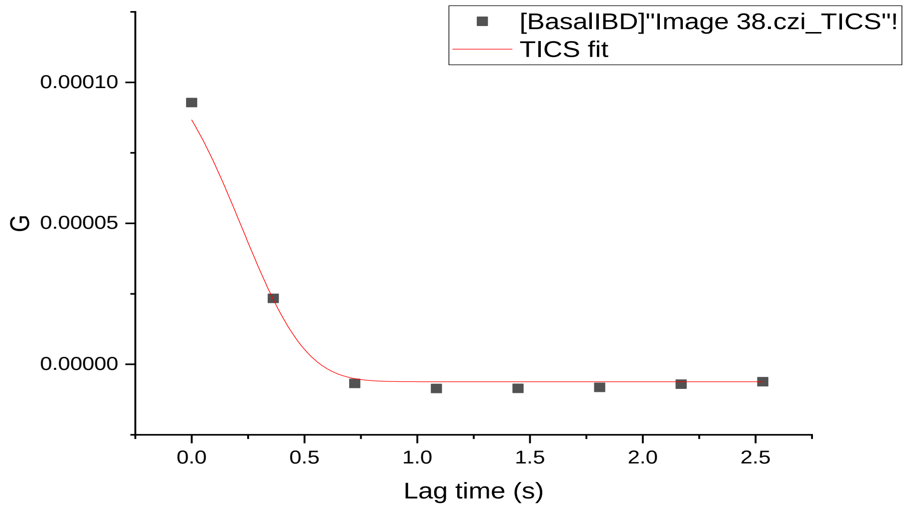
[BasalIBD]"Image 38.czi_TICS"!
TICS fit
G
0.00010
0.00005
0.00000
0.0 0.5 1.0 1.5 2.0 2.5
Lag time (s)

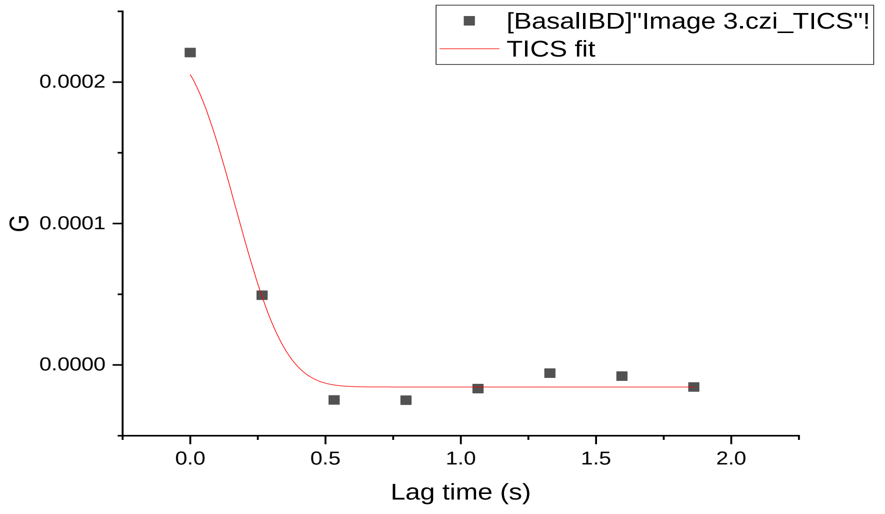
[BasalIBD]"Image 3.czi_TICS"!
TICS fit
G
0.0002
0.0001
0.0000
0.0 0.5 1.0 1.5 2.0
Lag time (s)

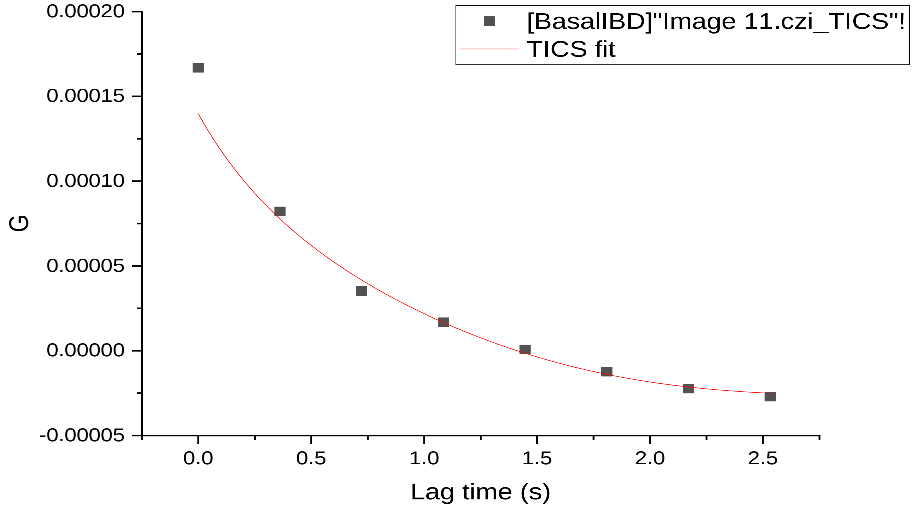
[BasalIBD]"Image 11.czi_TICS"!
TICS fit
G
0.00020
0.00015
0.00010
0.00005
0.00000
-0.00005
0.0 0.5 1.0 1.5 2.0 2.5
Lag time (s)

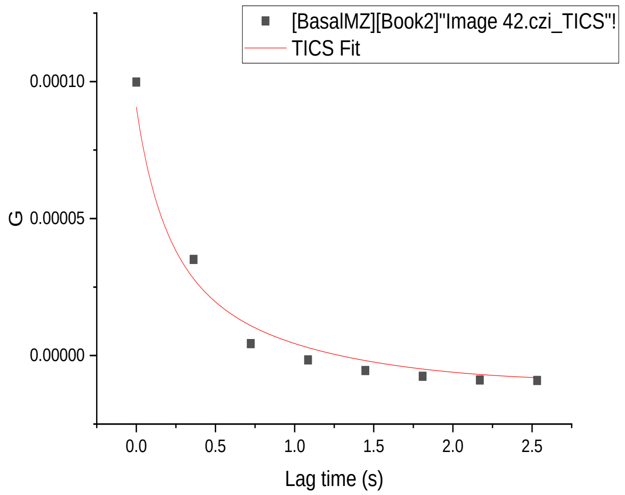
[BasalMZ][Book2]"Image 42.czi_TICS"!
TICS Fit
G
Lag time (s)

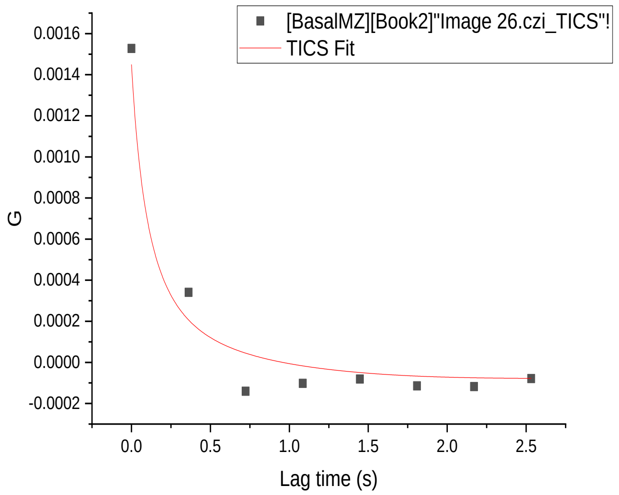
[BasalMZ][Book2]"Image 26.czi_TICS"!
TICS Fit
G
Lag time (s)

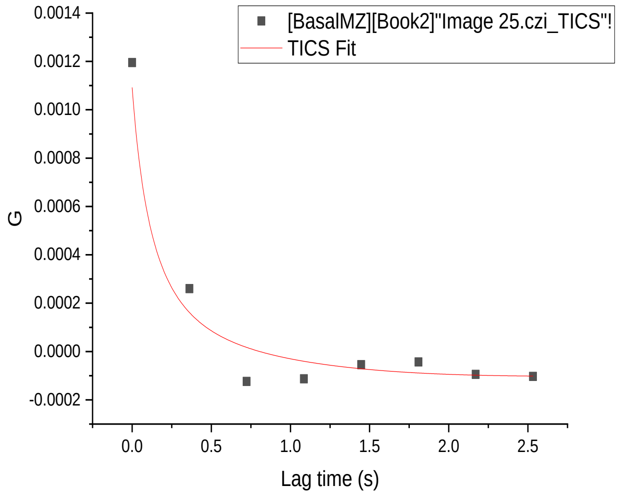
[BasalMZ][Book2]"Image 25.czi_TICS"!
TICS Fit
G
Lag time (s)

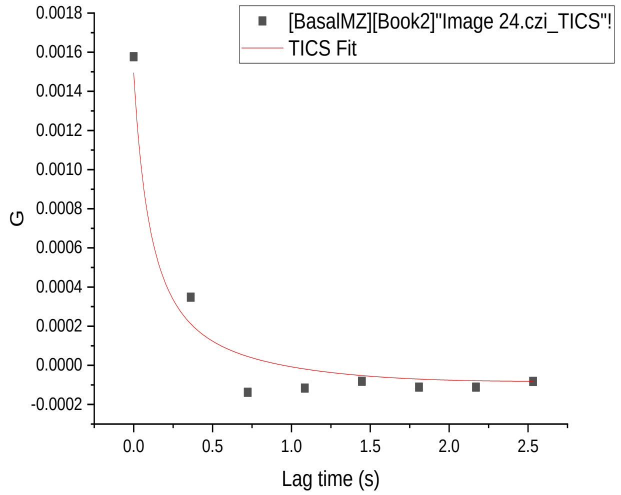
[BasalMZ][Book2]"Image 24.czi_TICS"!
TICS Fit
G
Lag time (s)

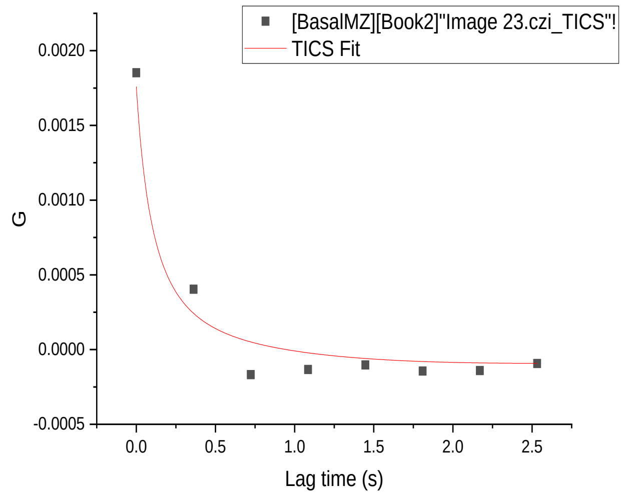
[BasalMZ][Book2]"Image 23.czi_TICS"!
TICS Fit
G
Lag time (s)

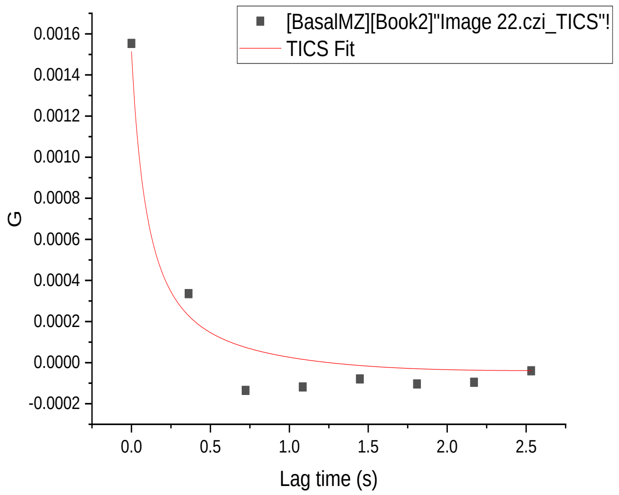
[BasalMZ][Book2]"Image 22.czi_TICS"!
TICS Fit
G
Lag time (s)

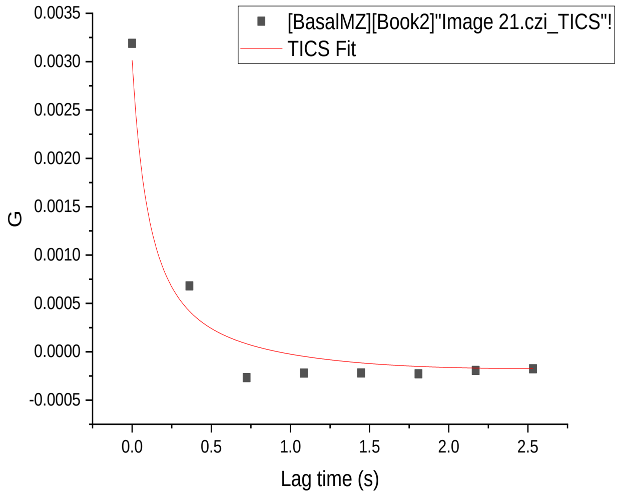
[BasalMZ][Book2]"Image 21.czi_TICS"!
TICS Fit
G
Lag time (s)

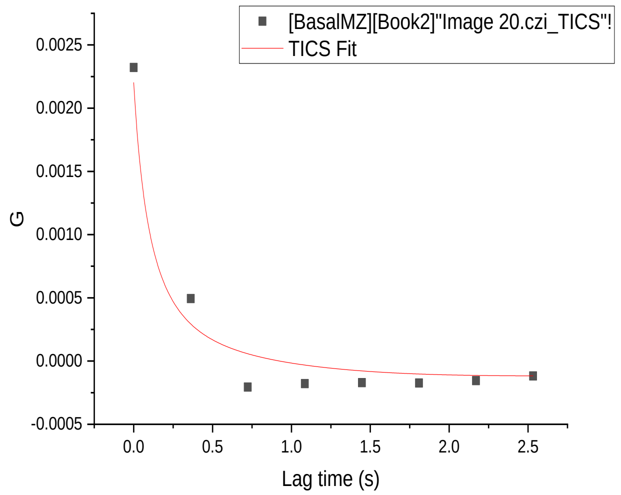
[BasalMZ][Book2]"Image 20.czi_TICS"!
TICS Fit
G
Lag time (s)

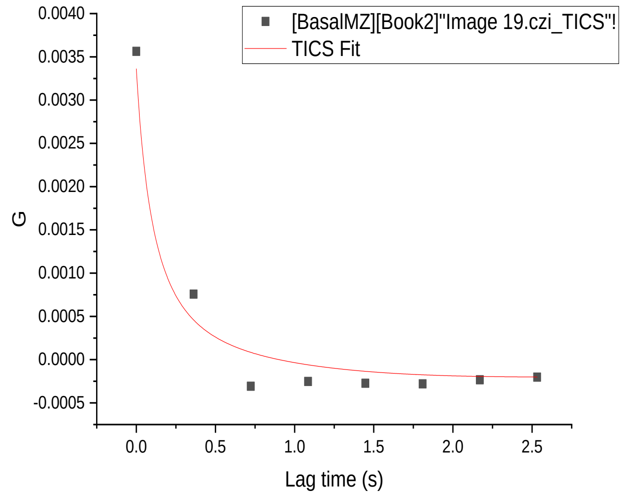
[BasalMZ][Book2]"Image 19.czi_TICS"!
TICS Fit
G
Lag time (s)

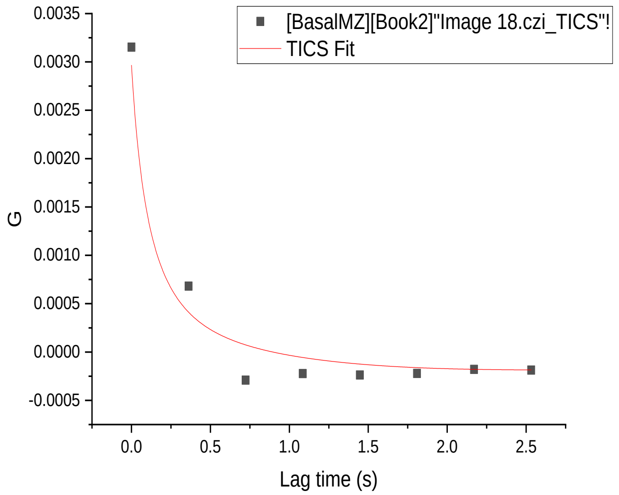
[BasalMZ][Book2]"Image 18.czi_TICS"!
TICS Fit
G
Lag time (s)

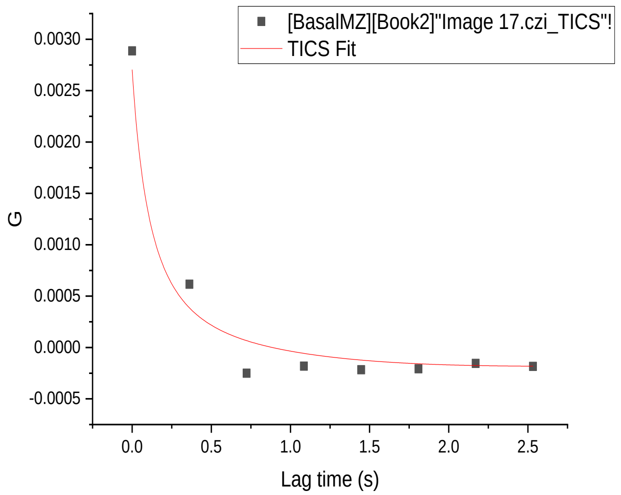
[BasalMZ][Book2]"Image 17.czi_TICS"!
TICS Fit
G
Lag time (s)

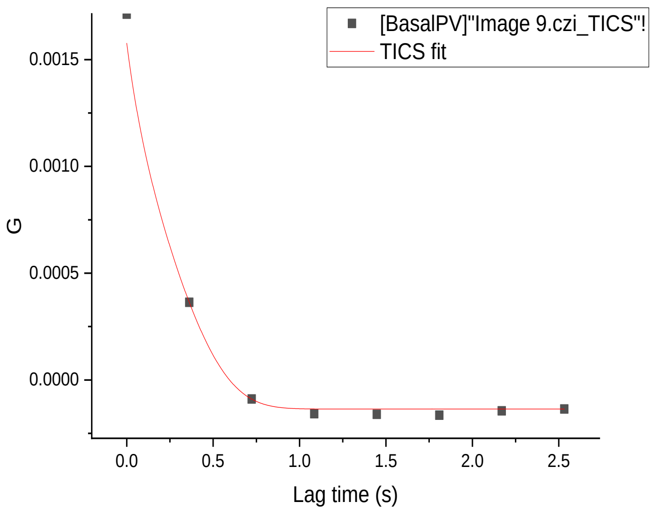
[BasalPV]"Image 9.czi_TICS"!
TICS fit
G
0.0015
0.0010
0.0005
0.0000
0.0 0.5 1.0 1.5 2.0 2.5
Lag time (s)

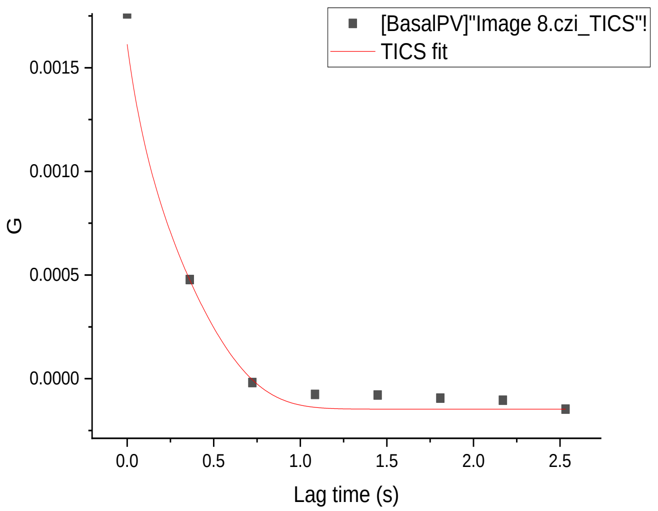
[BasalPV]"Image 8.czi_TICS"!
TICS fit
G
0.0015
0.0010
0.0005
0.0000
0.0 0.5 1.0 1.5 2.0 2.5
Lag time (s)

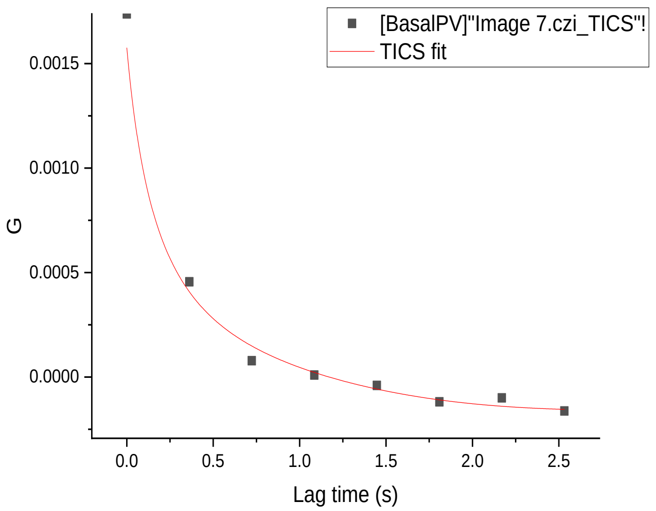
[BasalPV]"Image 7.czi_TICS"!
TICS fit
G
0.0015
0.0010
0.0005
0.0000
0.0 0.5 1.0 1.5 2.0 2.5
Lag time (s)

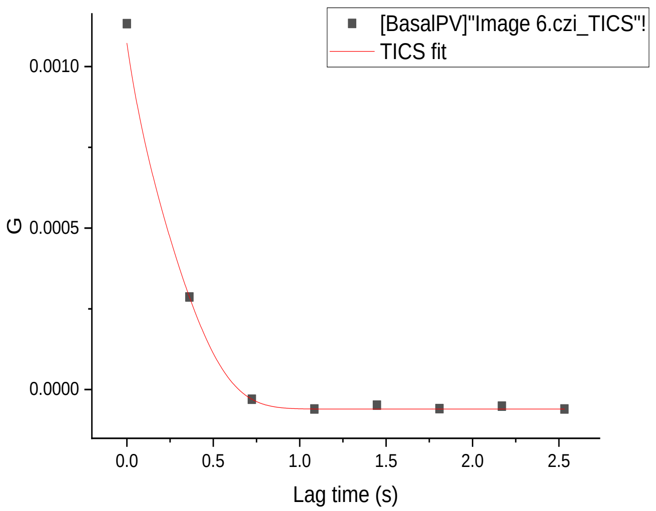
[BasalPV]"Image 6.czi_TICS"!
TICS fit
G
0.0010
0.0005
0.0000
0.0 0.5 1.0 1.5 2.0 2.5
Lag time (s)

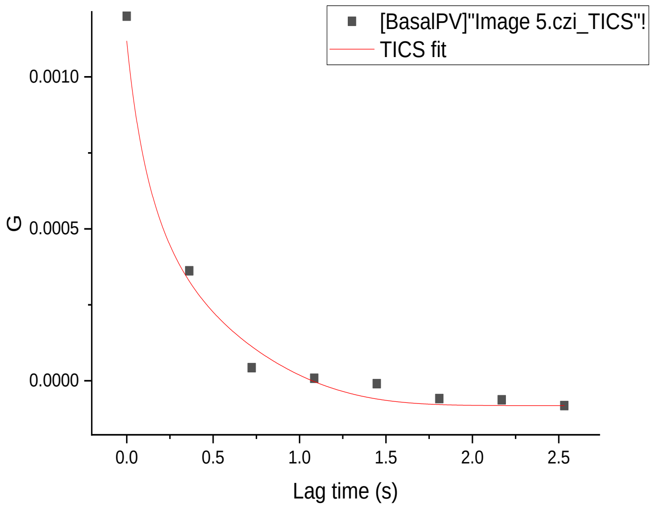
[BasalPV]"Image 5.czi_TICS"!
TICS fit
G
0.0010
0.0005
0.0000
0.0 0.5 1.0 1.5 2.0 2.5
Lag time (s)

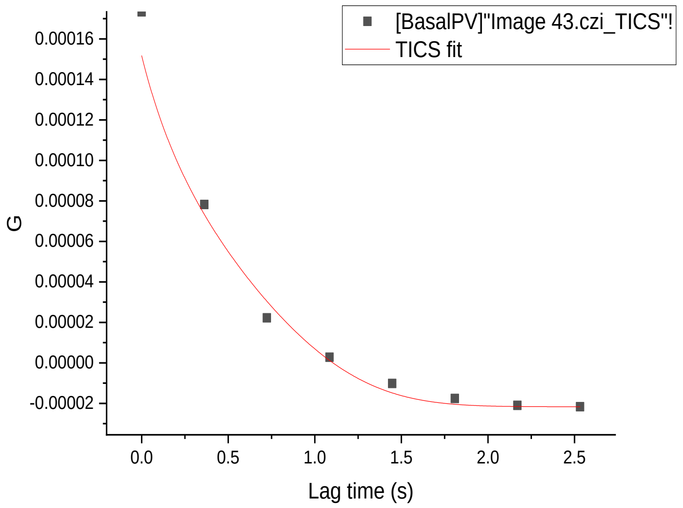
[BasalPV]"Image 43.czi_TICS"!
TICS fit
G
0.00016
0.00014
0.00012
0.00010
0.00008
0.00006
0.00004
0.00002
0.00000
-0.00002
0.0 0.5 1.0 1.5 2.0 2.5
Lag time (s)

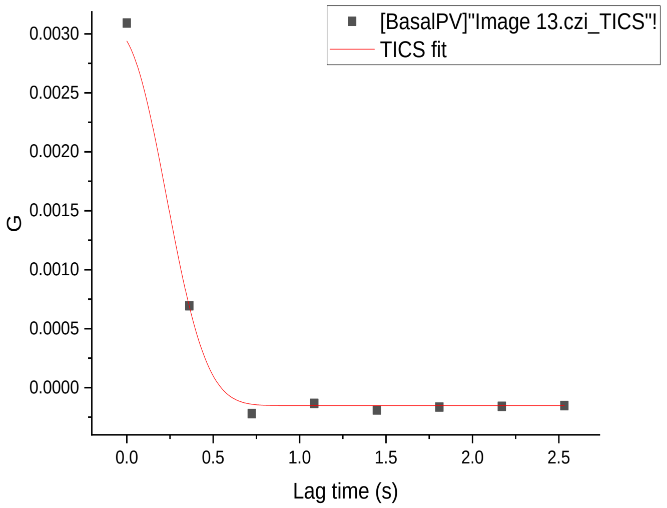
[BasalPV]"Image 13.czi_TICS"!
TICS fit
G
0.0030
0.0025
0.0020
0.0015
0.0010
0.0005
0.0000
0.0 0.5 1.0 1.5 2.0 2.5
Lag time (s)

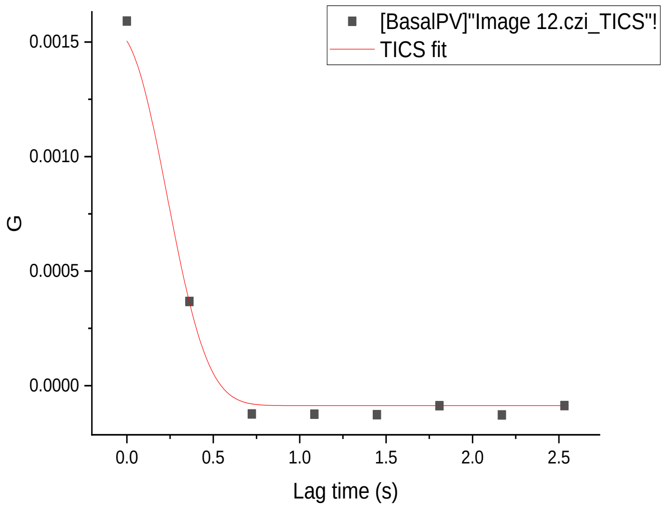
[BasalPV]"Image 12.czi_TICS"!
TICS fit
G
0.0015
0.0010
0.0005
0.0000
0.0 0.5 1.0 1.5 2.0 2.5
Lag time (s)

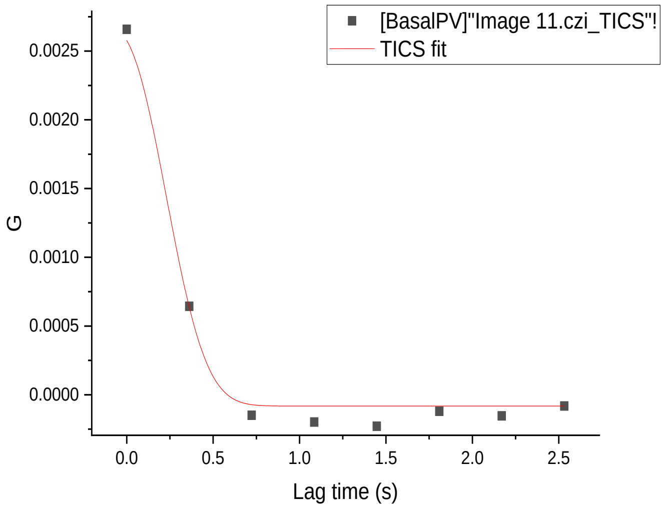
[BasalPV]"Image 11.czi_TICS"!
TICS fit
G
0.0025
0.0020
0.0015
0.0010
0.0005
0.0000
0.0 0.5 1.0 1.5 2.0 2.5
Lag time (s)

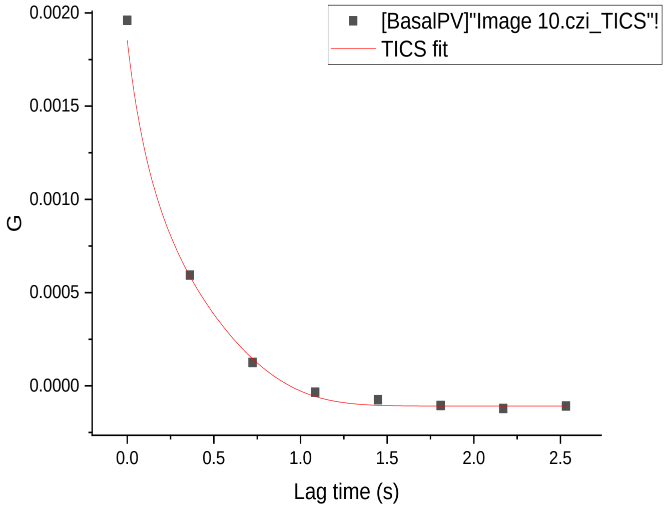
[BasalPV]"Image 10.czi_TICS"!
TICS fit
G
0.0020
0.0015
0.0010
0.0005
0.0000
0.0 0.5 1.0 1.5 2.0 2.5
Lag time (s)

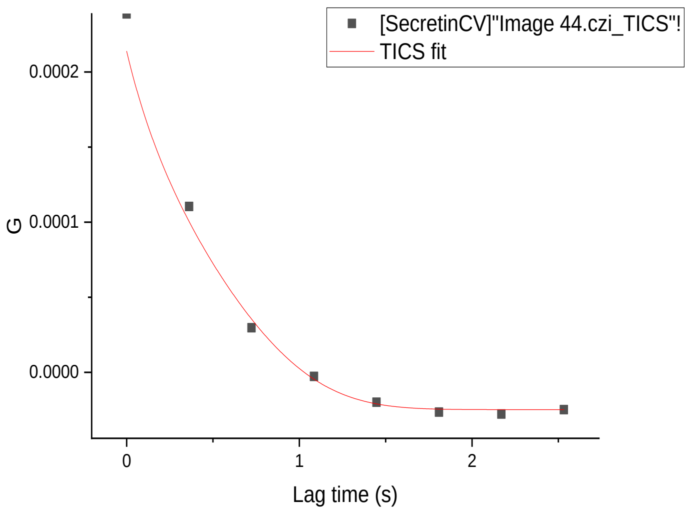
[SecretinCV]"Image 44.czi_TICS"!
TICS fit
G
0.0002
0.0001
0.0000
0
1
2
Lag time (s)

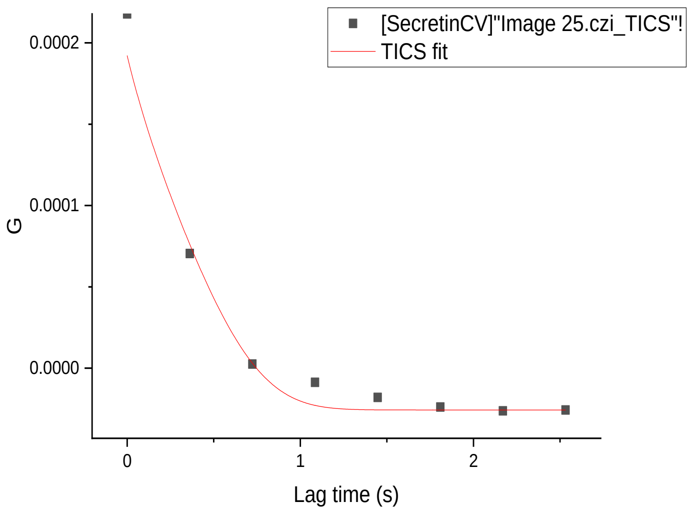
[SecretinCV]"Image 25.czi_TICS"!
TICS fit
G
0.0002
0.0001
0.0000
0
1
2
Lag time (s)

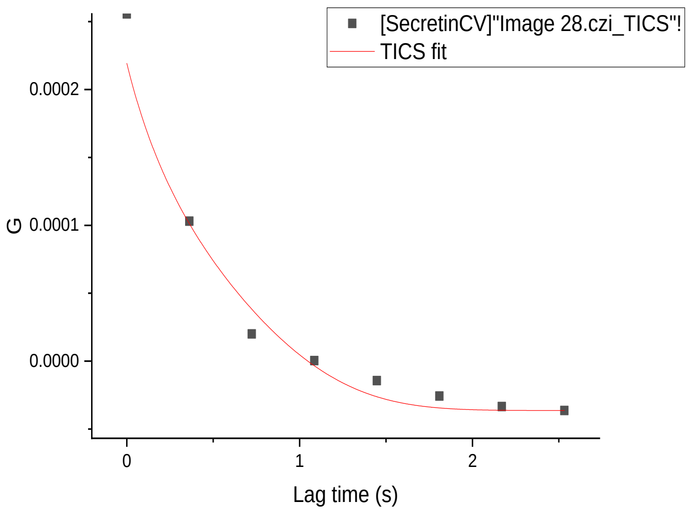
[SecretinCV]"Image 28.czi_TICS"!
TICS fit
G
0.0002
0.0001
0.0000
0
1
2
Lag time (s)

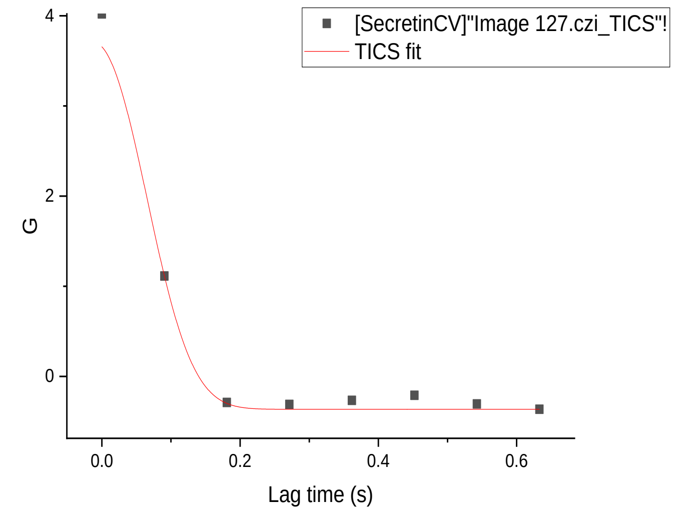
[SecretinCV]"Image 127.czi_TICS"!
TICS fit
G
4
2
0
0.0
0.2
0.4
0.6
Lag time (s)

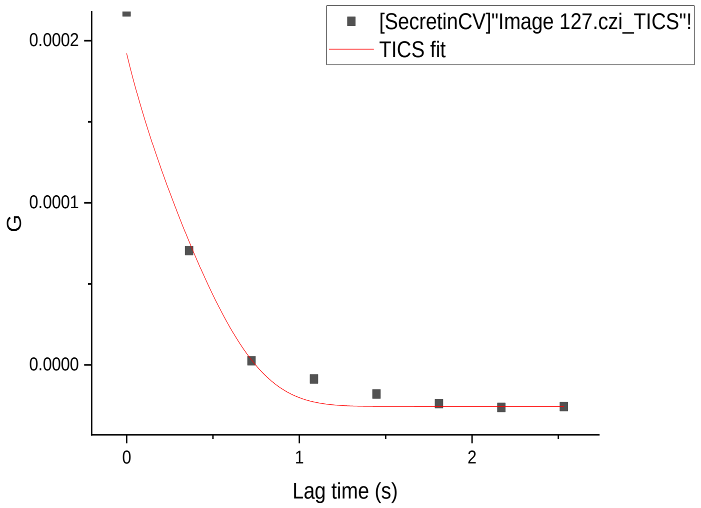
[SecretinCV]"Image 127.czi_TICS"!
TICS fit
G
0.0002
0.0001
0.0000
0
1
2
Lag time (s)

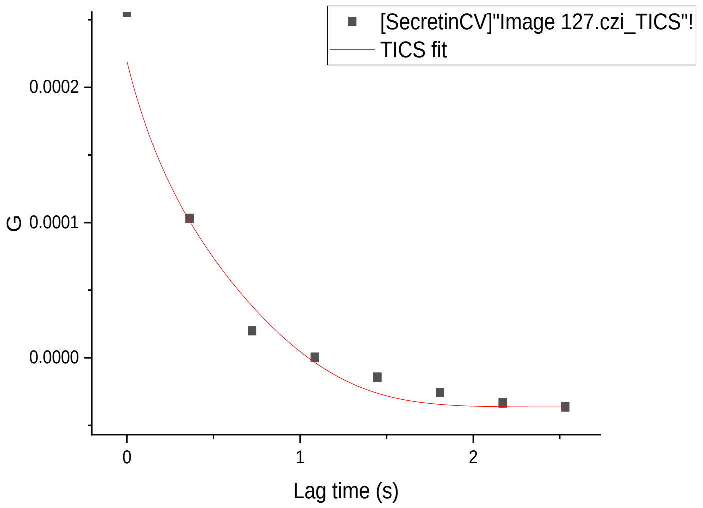
[SecretinCV]"Image 127.czi_TICS"!
TICS fit
G
0.0002
0.0001
0.0000
0
1
2
Lag time (s)

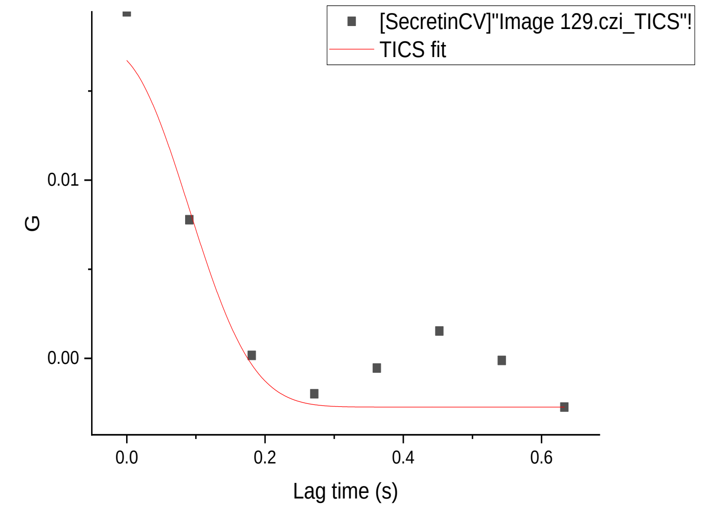
[SecretinCV]"Image 129.czi_TICS"!
TICS fit
G
0.01
0.00
0.0
0.2
0.4
0.6
Lag time (s)

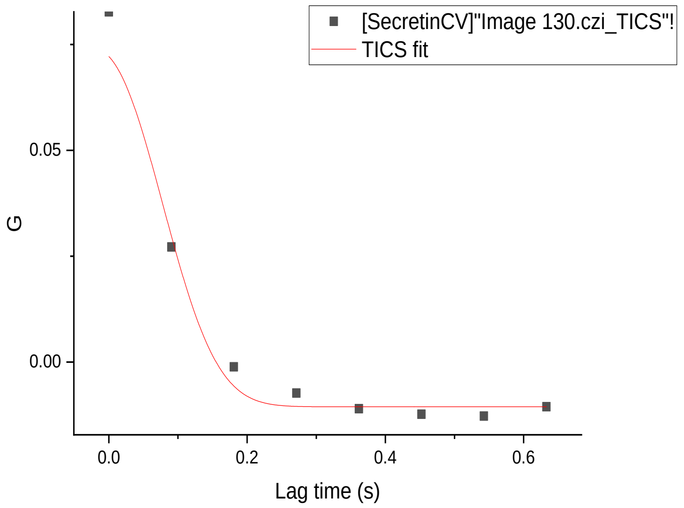
[SecretinCV]"Image 130.czi_TICS"!
TICS fit
G
0.05
0.00
0.0
0.2
0.4
0.6
Lag time (s)

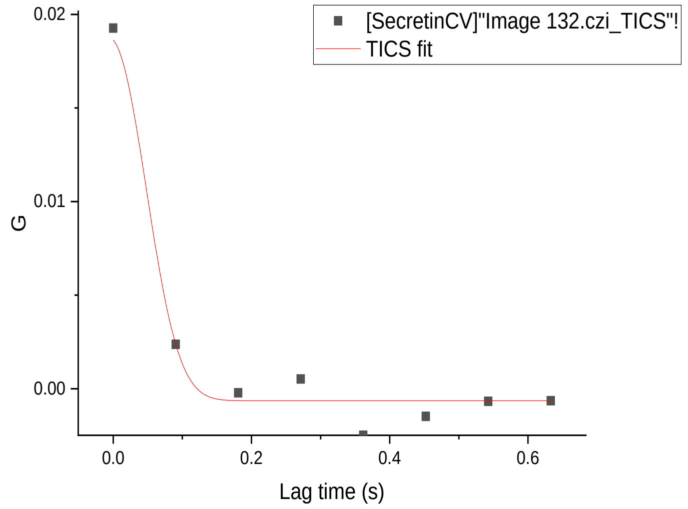
[SecretinCV]"Image 132.czi_TICS"!
TICS fit
G
0.02
0.01
0.00
0.0
0.2
0.4
0.6
Lag time (s)

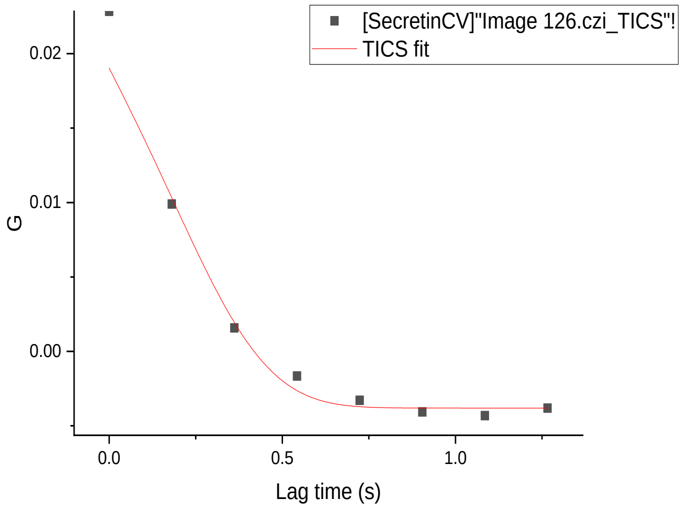
[SecretinCV]"Image 126.czi_TICS"!
TICS fit
G
0.02
0.01
0.00
0.0
0.5
1.0
Lag time (s)

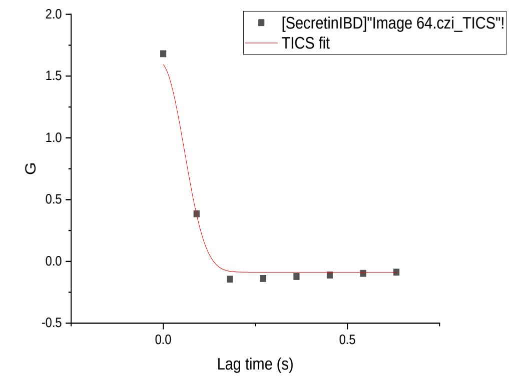
[SecretinIBD]"Image 64.czi_TICS"!
TICS fit
G
Lag time (s)

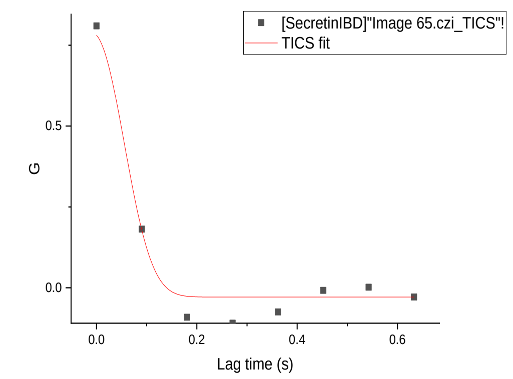
[SecretinIBD]"Image 65.czi_TICS"!
TICS fit
G
Lag time (s)

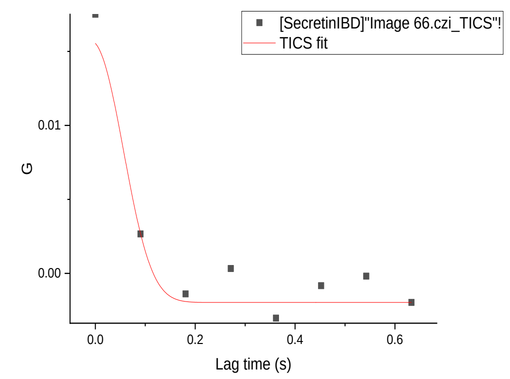
[SecretinIBD]"Image 66.czi_TICS"!
TICS fit
G
Lag time (s)

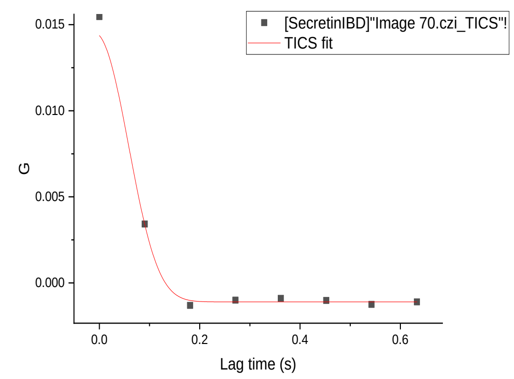
[SecretinIBD]"Image 70.czi_TICS"!
TICS fit
G
Lag time (s)

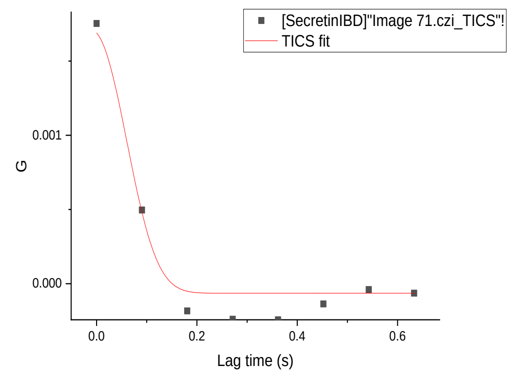
[SecretinIBD]"Image 71.czi_TICS"!
TICS fit
G
Lag time (s)

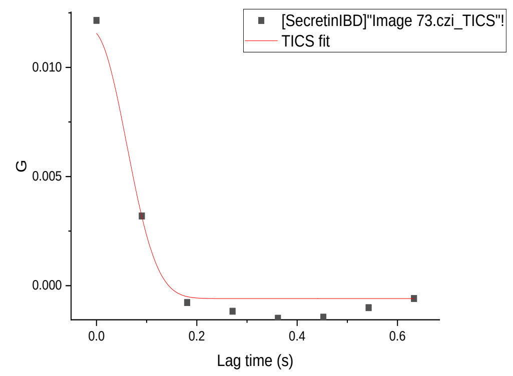
[SecretinIBD]"Image 73.czi_TICS"!
TICS fit
G
Lag time (s)

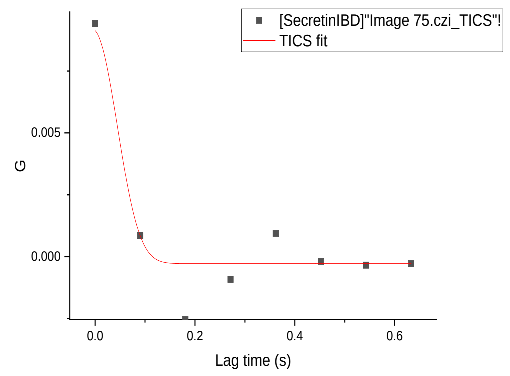
[SecretinIBD]"Image 75.czi_TICS"!
TICS fit
G
Lag time (s)

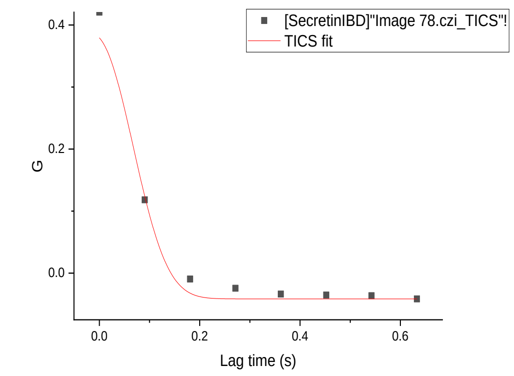
[SecretinIBD]"Image 78.czi_TICS"!
TICS fit
G
Lag time (s)

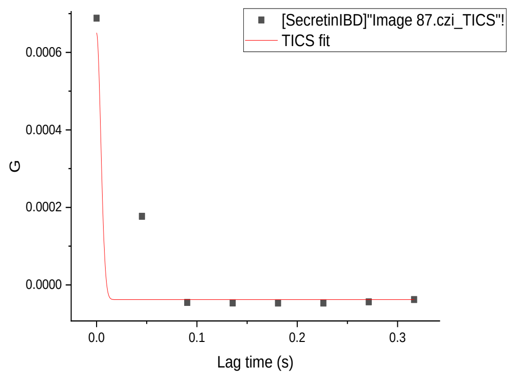
[SecretinIBD]"Image 87.czi_TICS"!
TICS fit
G
Lag time (s)

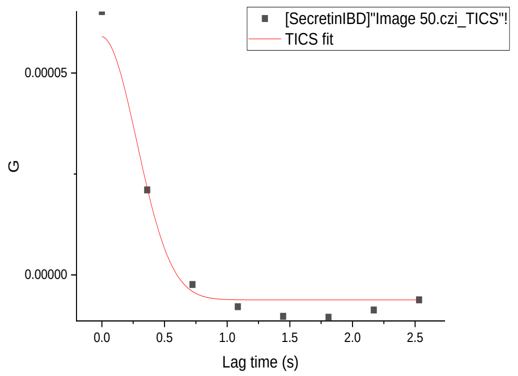
[SecretinIBD]"Image 50.czi_TICS"!
TICS fit
G
Lag time (s)

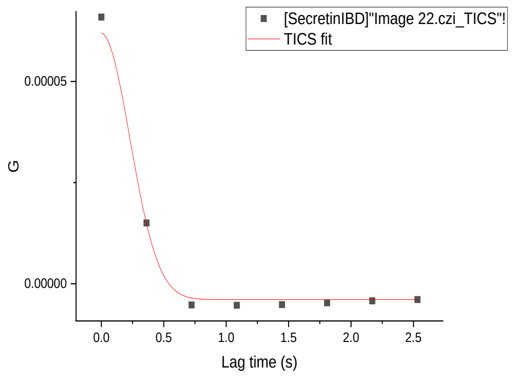
[SecretinIBD]"Image 22.czi_TICS"!
TICS fit
G
Lag time (s)

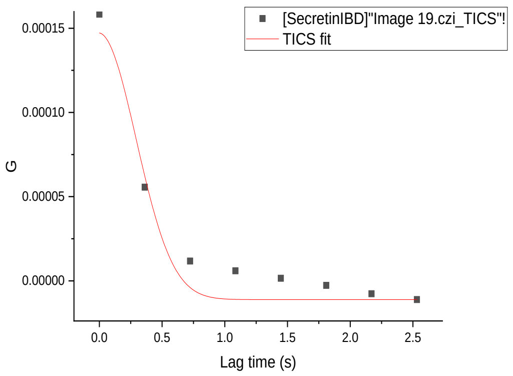
[SecretinIBD]"Image 19.czi_TICS"!
TICS fit
G
Lag time (s)

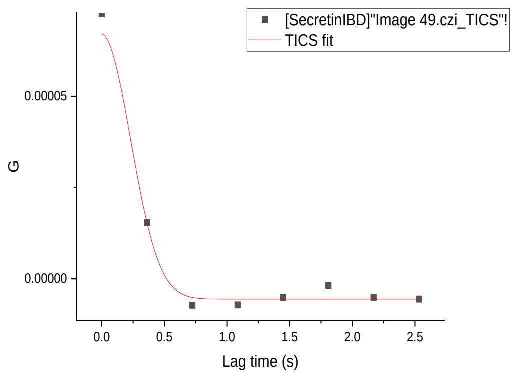
[SecretinIBD]"Image 49.czi_TICS"!
TICS fit
G
Lag time (s)

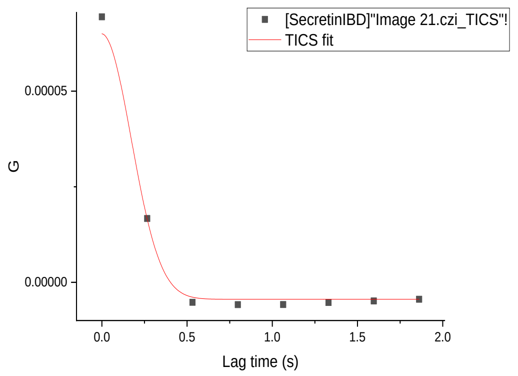
[SecretinIBD]"Image 21.czi_TICS"!
TICS fit
G
Lag time (s)

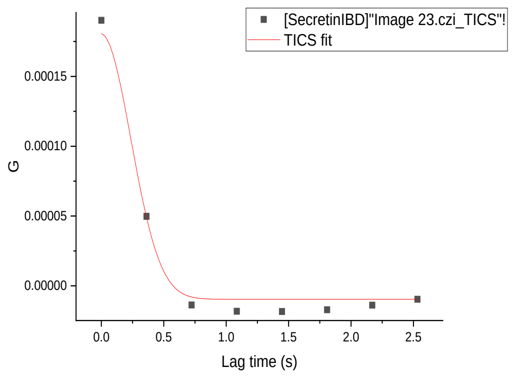
[SecretinIBD]"Image 23.czi_TICS"!
TICS fit
G
Lag time (s)

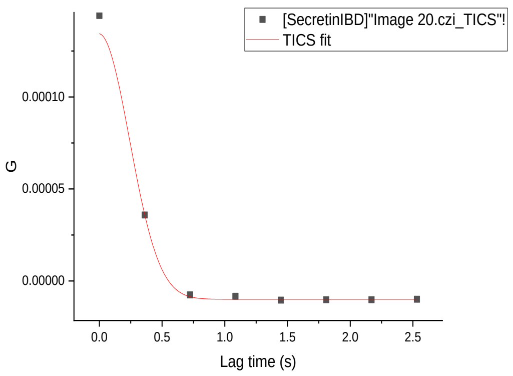
[SecretinIBD]"Image 20.czi_TICS"!
TICS fit
G
Lag time (s)

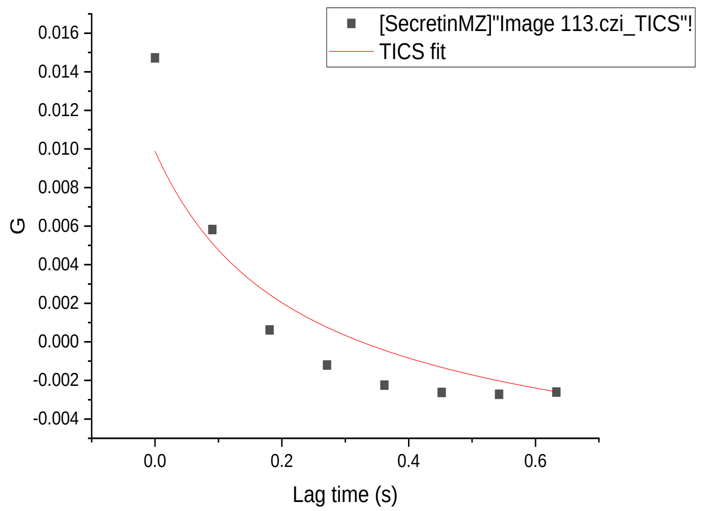
[SecretinMZ]"Image 113.czi_TICS"!
TICS fit
G
Lag time (s)
0.016
0.014
0.012
0.010
0.008
0.006
0.004
0.002
0.000
-0.002
-0.004
0.0
0.2
0.4
0.6

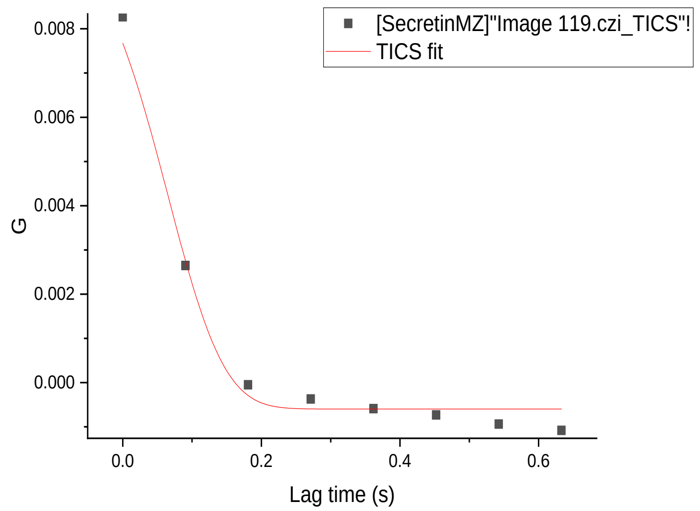
[SecretinMZ]"Image 119.czi_TICS"!
TICS fit
G
Lag time (s)
0.008
0.006
0.004
0.002
0.000
0.0
0.2
0.4
0.6

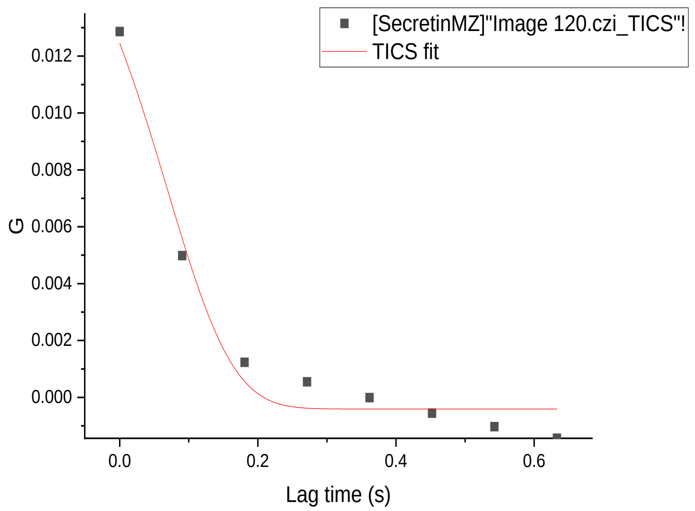
[SecretinMZ]"Image 120.czi_TICS"!
TICS fit
G
Lag time (s)
0.012
0.010
0.008
0.006
0.004
0.002
0.000
0.0
0.2
0.4
0.6

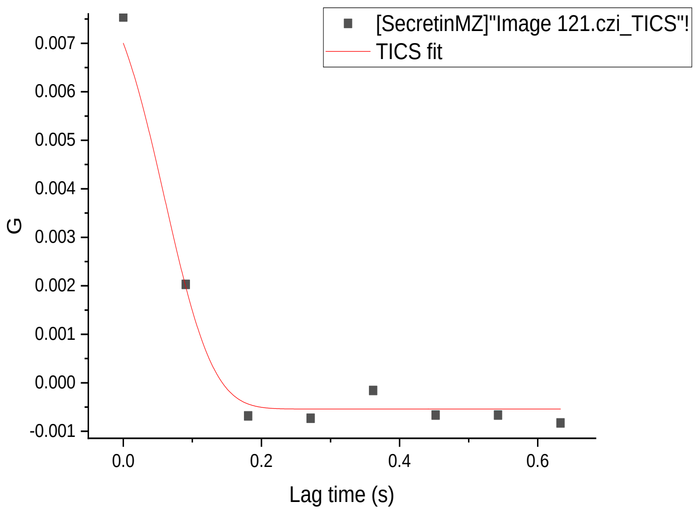
[SecretinMZ]"Image 121.czi_TICS"!
TICS fit
G
Lag time (s)
0.007
0.006
0.005
0.004
0.003
0.002
0.001
0.000
-0.001
0.0
0.2
0.4
0.6

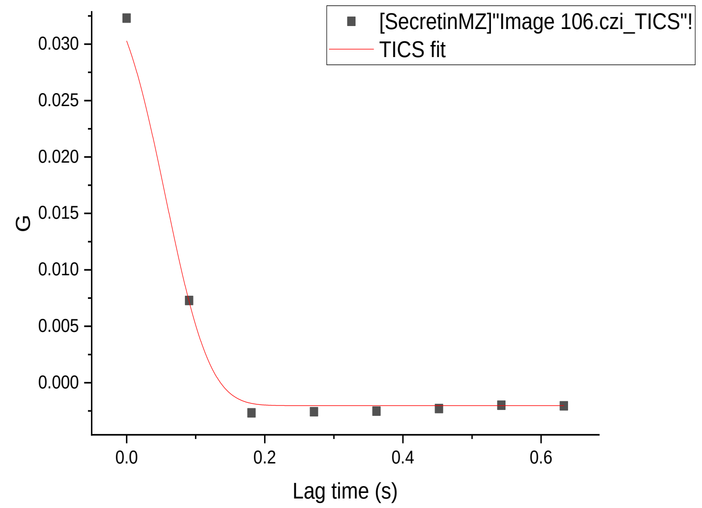
[SecretinMZ]"Image 106.czi_TICS"!
TICS fit
G
Lag time (s)
0.030
0.025
0.020
0.015
0.010
0.005
0.000
0.0
0.2
0.4
0.6

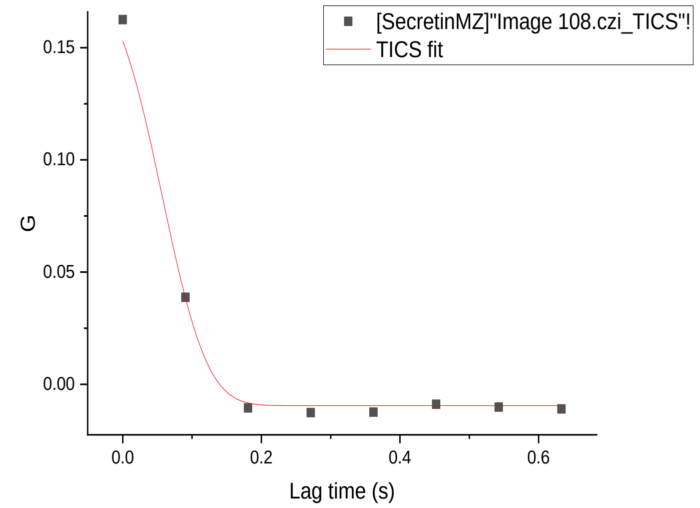
[SecretinMZ]"Image 108.czi_TICS"!
TICS fit
G
Lag time (s)
0.15
0.10
0.05
0.00
0.0
0.2
0.4
0.6

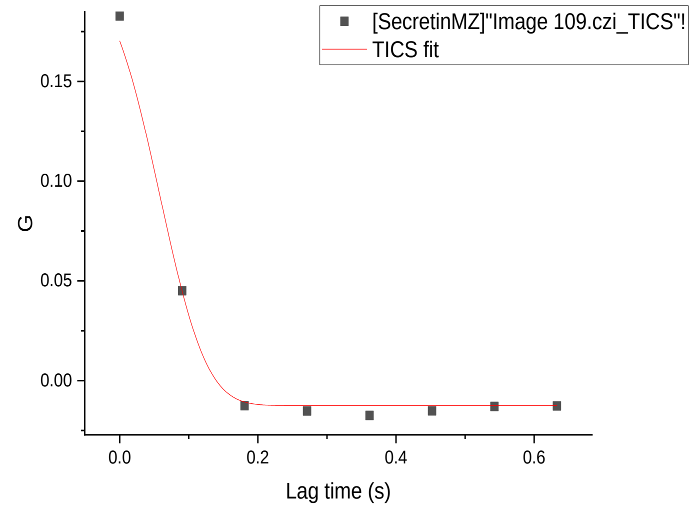
[SecretinMZ]"Image 109.czi_TICS"!
TICS fit
G
Lag time (s)
0.15
0.10
0.05
0.00
0.0
0.2
0.4
0.6

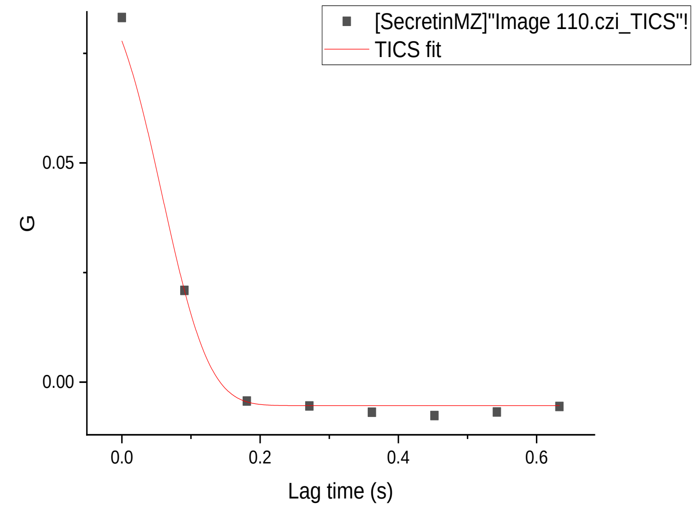
[SecretinMZ]"Image 110.czi_TICS"!
TICS fit
G
Lag time (s)
0.05
0.00
0.0
0.2
0.4
0.6

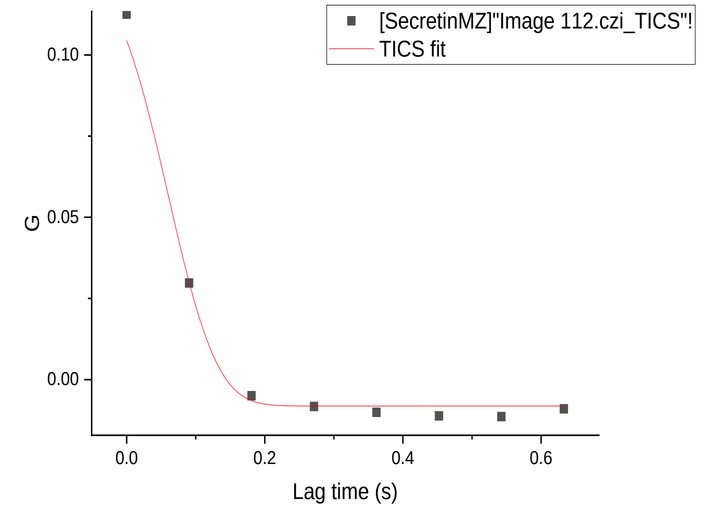
[SecretinMZ]"Image 112.czi_TICS"!
TICS fit
G
Lag time (s)
0.10
0.05
0.00
0.0
0.2
0.4
0.6

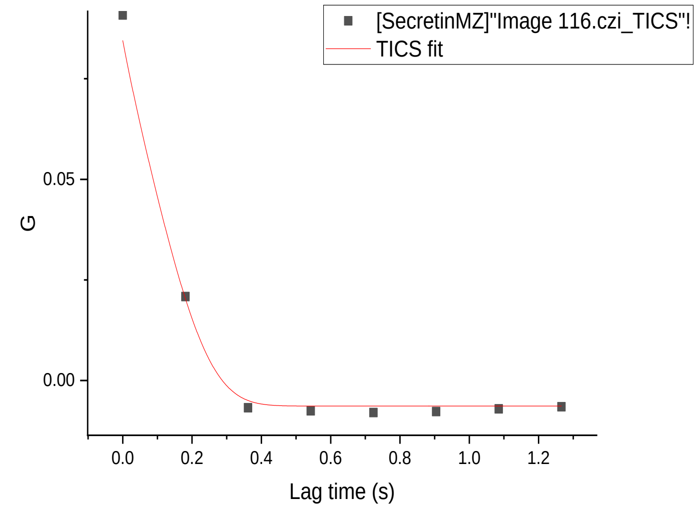
[SecretinMZ]"Image 116.czi_TICS"!
TICS fit
G
Lag time (s)
0.05
0.00
0.0
0.2
0.4
0.6
0.8
1.0
1.2

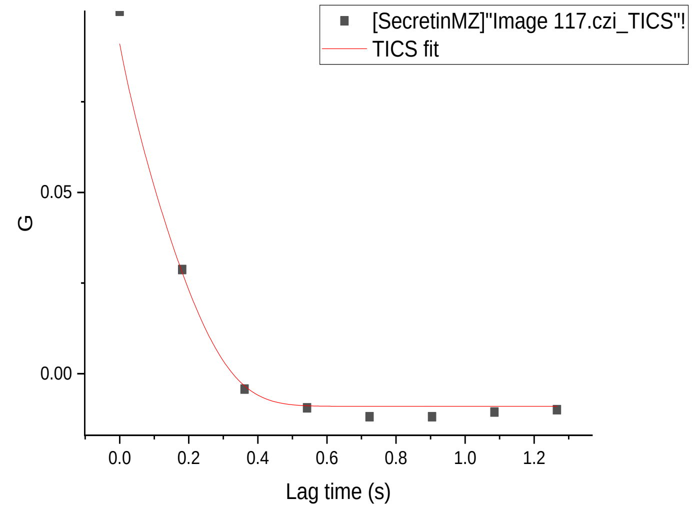
[SecretinMZ]"Image 117.czi_TICS"!
TICS fit
G
Lag time (s)
0.05
0.00
0.0
0.2
0.4
0.6
0.8
1.0
1.2

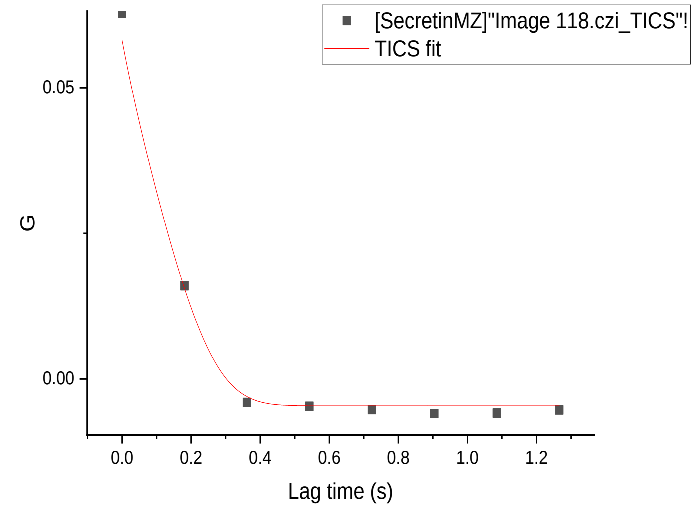
[SecretinMZ]"Image 118.czi_TICS"!
TICS fit
G
Lag time (s)
0.05
0.00
0.0
0.2
0.4
0.6
0.8
1.0
1.2

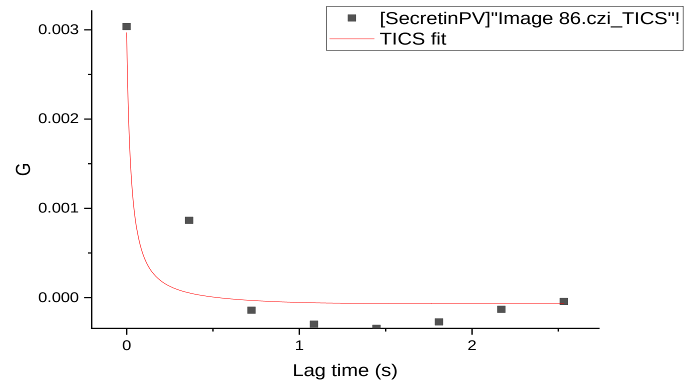
[SecretinPV]"Image 86.czi_TICS"!
TICS fit
G
Lag time (s)

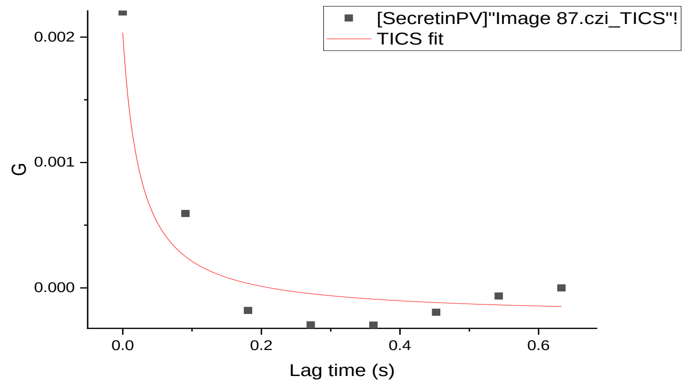
[SecretinPV]"Image 87.czi_TICS"!
TICS fit
G
Lag time (s)

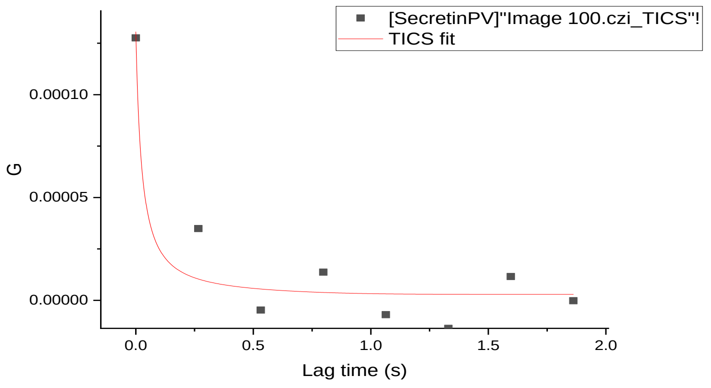
[SecretinPV]"Image 100.czi_TICS"!
TICS fit
G
Lag time (s)

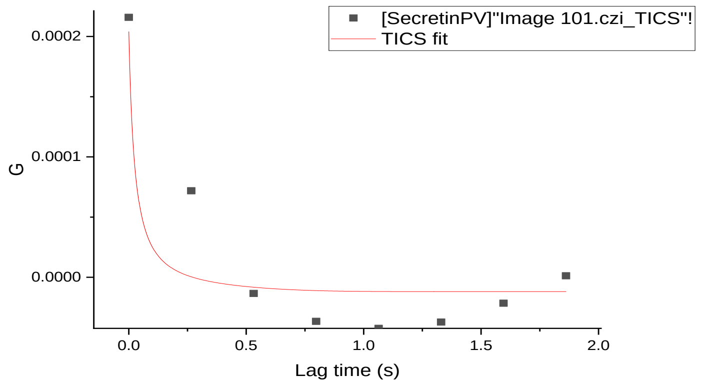
[SecretinPV]"Image 101.czi_TICS"!
TICS fit
G
Lag time (s)

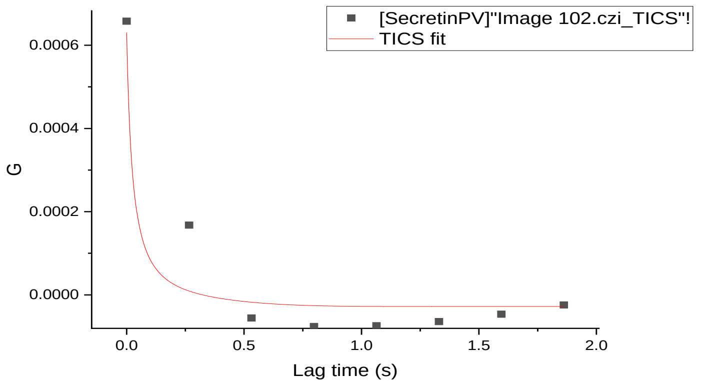
[SecretinPV]"Image 102.czi_TICS"!
TICS fit
G
Lag time (s)

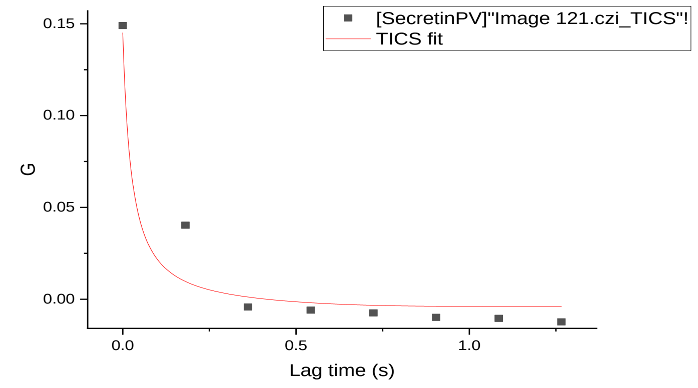
[SecretinPV]"Image 121.czi_TICS"!
TICS fit
G
Lag time (s)

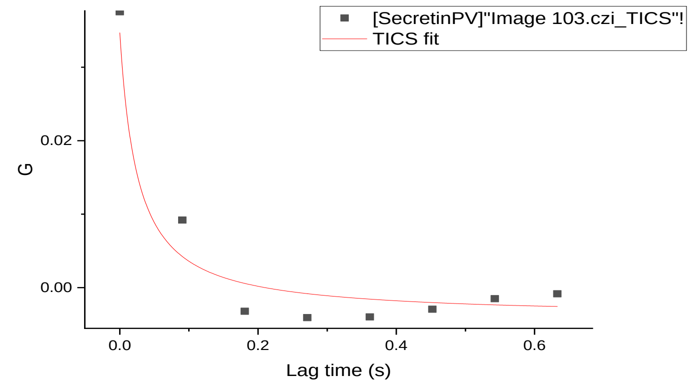
[SecretinPV]"Image 103.czi_TICS"!
TICS fit
G
Lag time (s)

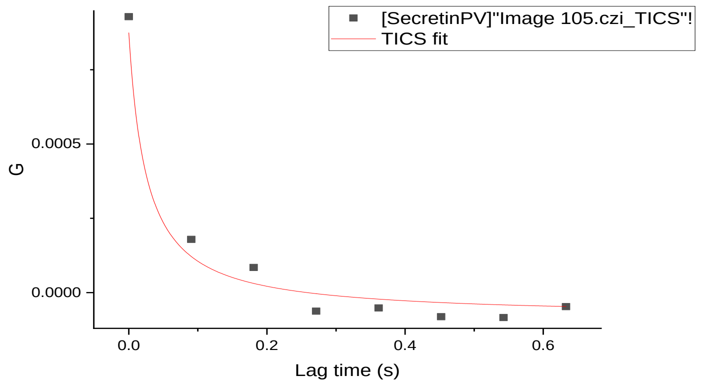
[SecretinPV]"Image 105.czi_TICS"!
TICS fit
G
Lag time (s)

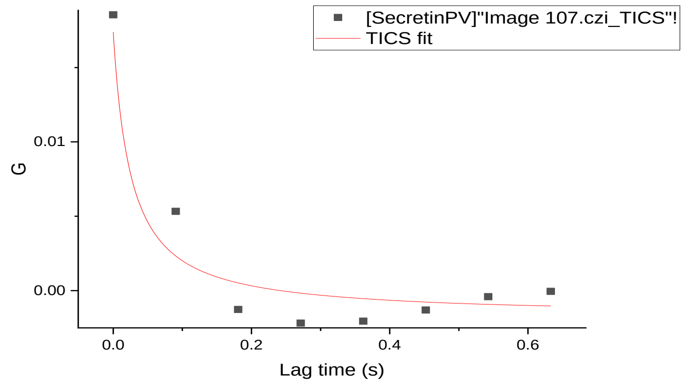
[SecretinPV]"Image 107.czi_TICS"!
TICS fit
G
Lag time (s)

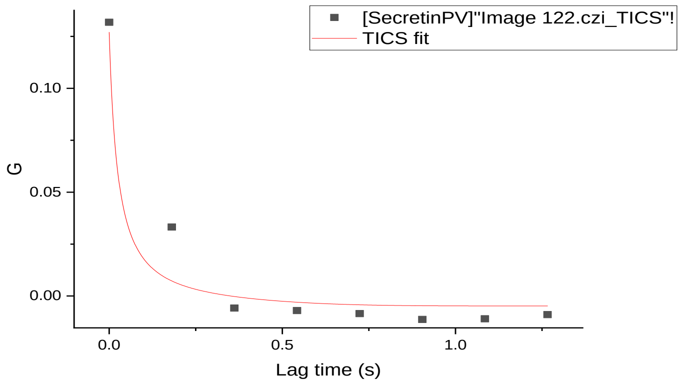
[SecretinPV]"Image 122.czi_TICS"!
TICS fit
G
Lag time (s)

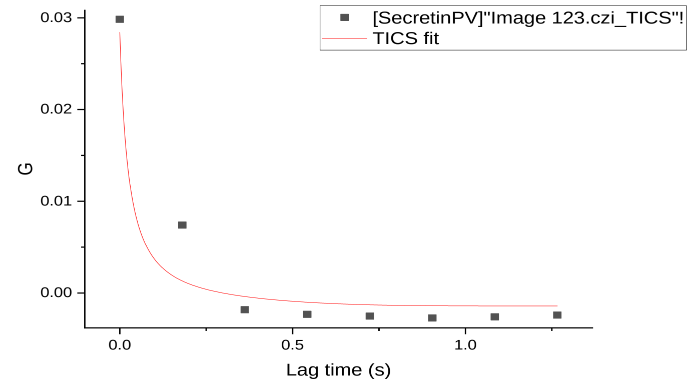
[SecretinPV]"Image 123.czi_TICS"!
TICS fit
G
Lag time (s)

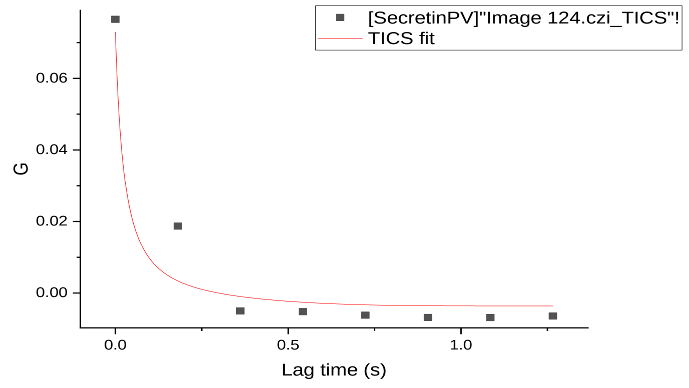
[SecretinPV]"Image 124.czi_TICS"!
TICS fit
G
Lag time (s)

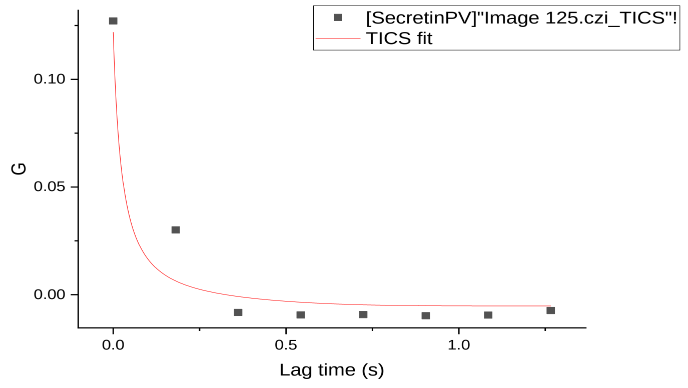
[SecretinPV]"Image 125.czi_TICS"!
TICS fit
G
Lag time (s)

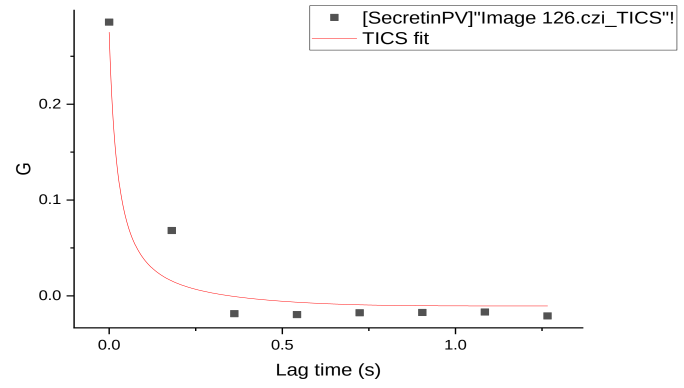
[SecretinPV]"Image 126.czi_TICS"!
TICS fit
G
Lag time (s)

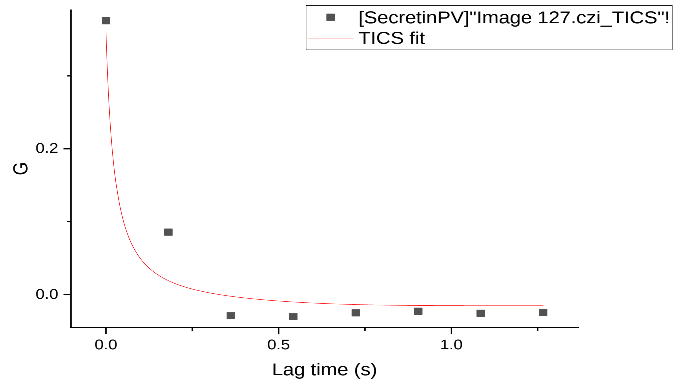
[SecretinPV]"Image 127.czi_TICS"!
TICS fit
G
Lag time (s)

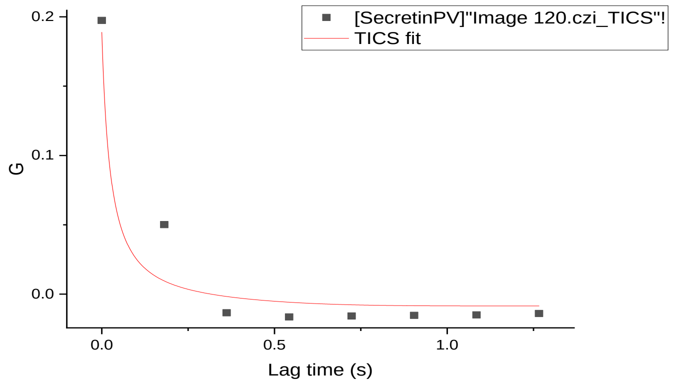
[SecretinPV]"Image 120.czi_TICS"!
TICS fit
G
Lag time (s)

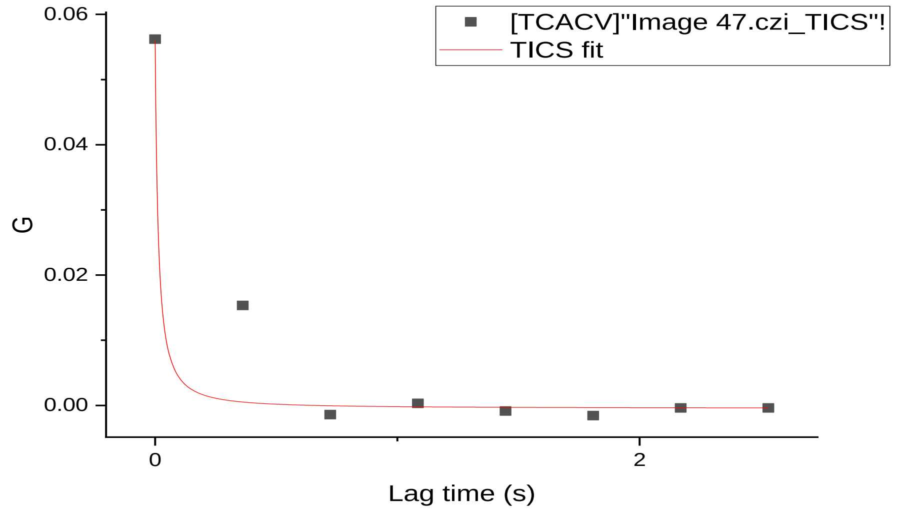
[TCACV]"Image 47.czi_TICS"!
TICS fit
G
Lag time (s)
0.06
0.04
0.02
0.00
0
2

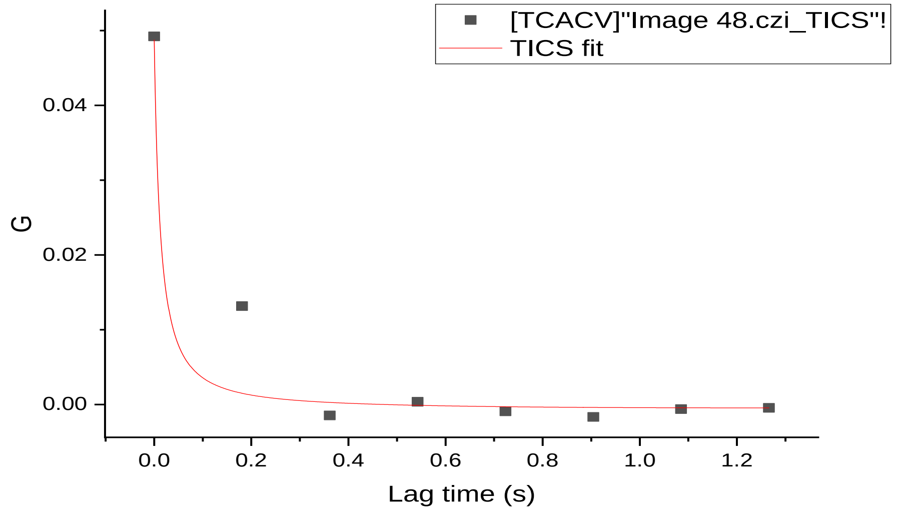
[TCACV]"Image 48.czi_TICS"!
TICS fit
G
Lag time (s)
0.04
0.02
0.00
0.0
0.2
0.4
0.6
0.8
1.0
1.2

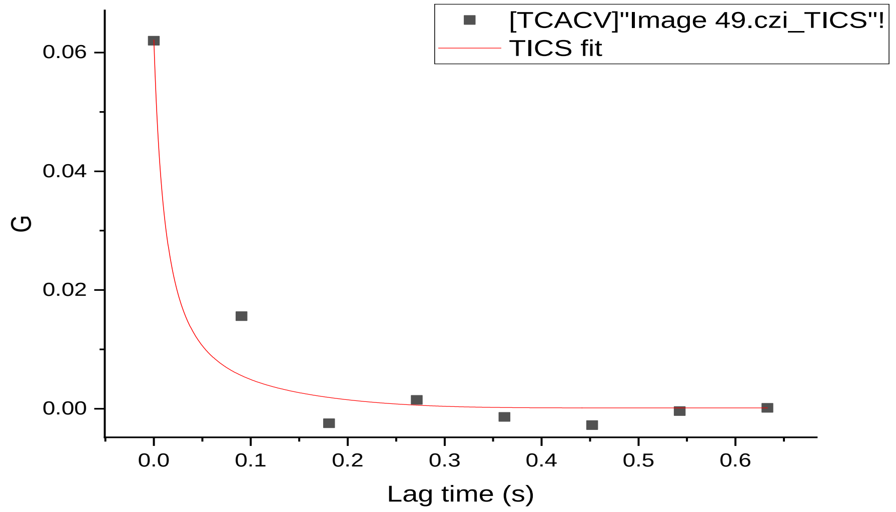
[TCACV]"Image 49.czi_TICS"!
TICS fit
G
Lag time (s)
0.06
0.04
0.02
0.00
0.0
0.1
0.2
0.3
0.4
0.5
0.6

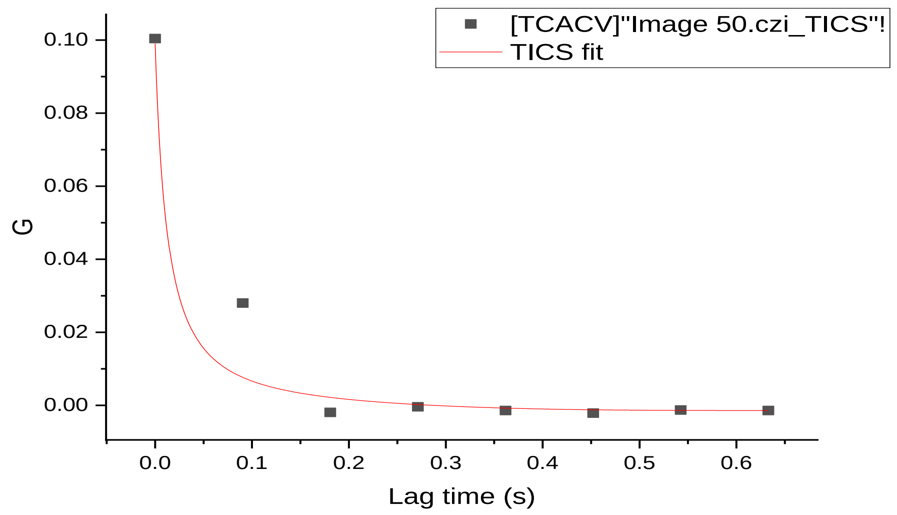
[TCACV]"Image 50.czi_TICS"!
TICS fit
G
Lag time (s)
0.10
0.08
0.06
0.04
0.02
0.00
0.0
0.1
0.2
0.3
0.4
0.5
0.6

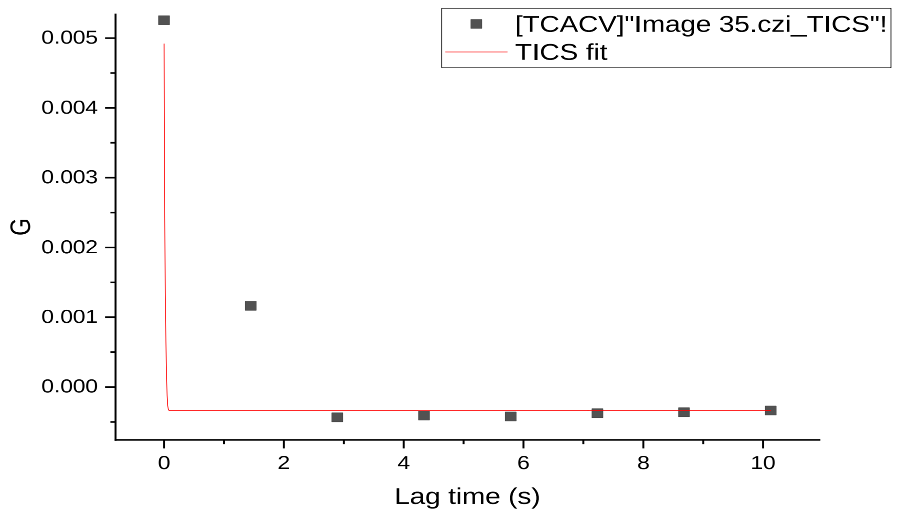
[TCACV]"Image 35.czi_TICS"!
TICS fit
G
Lag time (s)
0.005
0.004
0.003
0.002
0.001
0.000
0
2
4
6
8
10

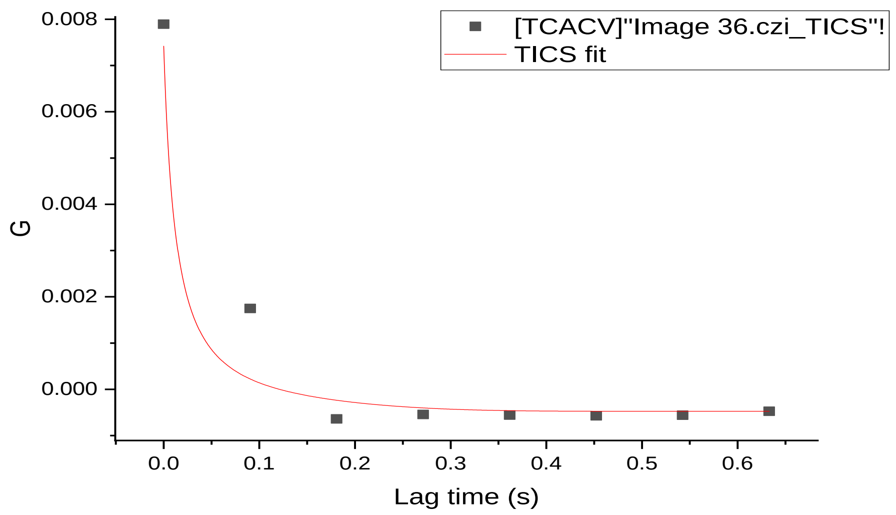
[TCACV]"Image 36.czi_TICS"!
TICS fit
G
Lag time (s)
0.008
0.006
0.004
0.002
0.000
0.0
0.1
0.2
0.3
0.4
0.5
0.6

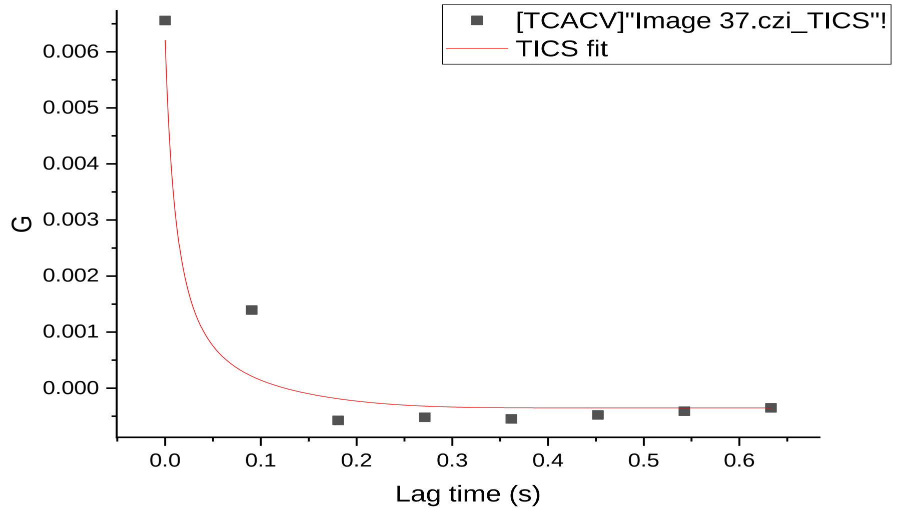
[TCACV]"Image 37.czi_TICS"!
TICS fit
G
Lag time (s)
0.006
0.005
0.004
0.003
0.002
0.001
0.000
0.0
0.1
0.2
0.3
0.4
0.5
0.6

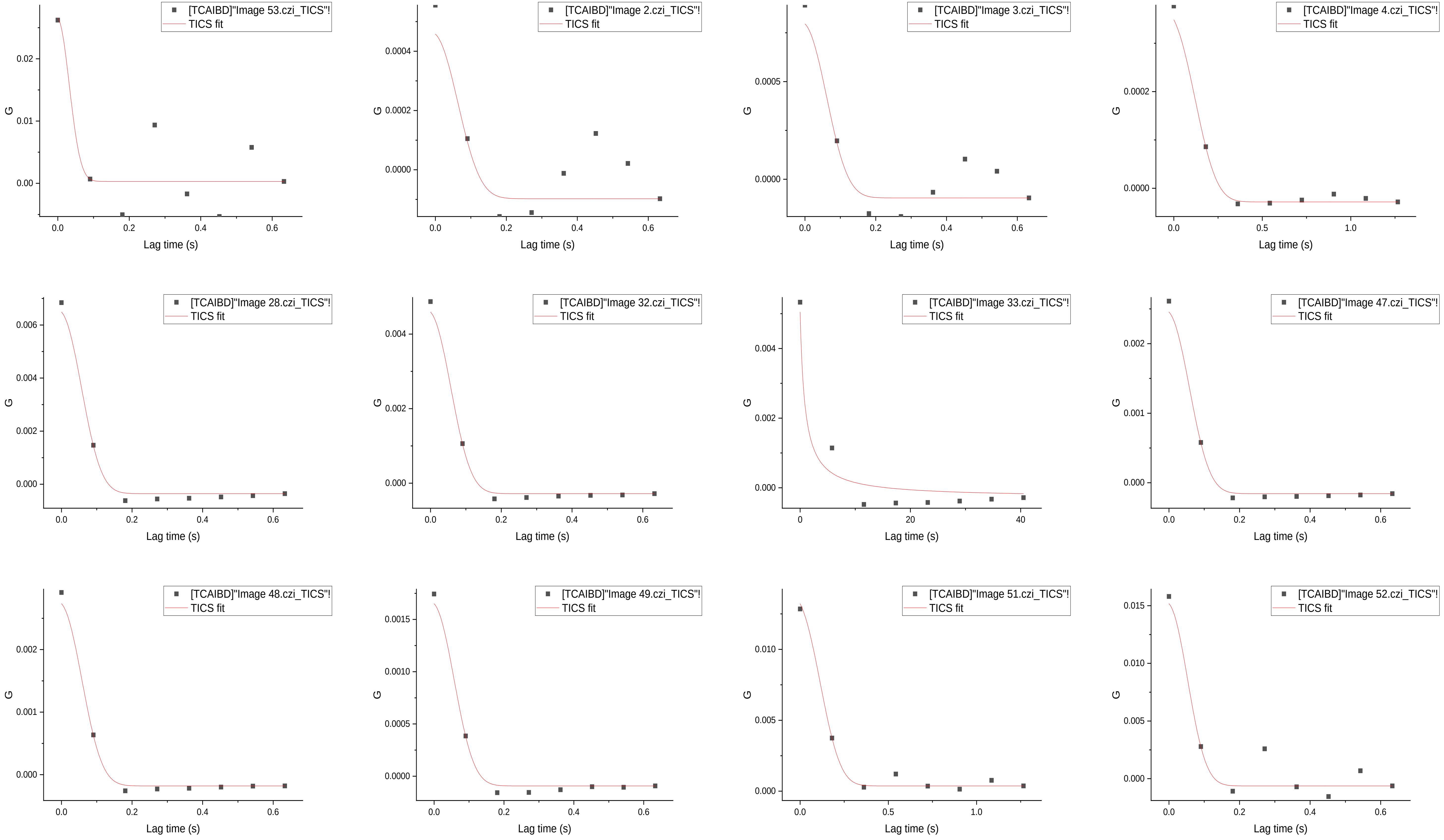
[TCAIBD]"Image 53.czi_TICS"!
TICS fit
[TCAIBD]"Image 2.czi_TICS"!
TICS fit
[TCAIBD]"Image 3.czi_TICS"!
TICS fit
[TCAIBD]"Image 4.czi_TICS"!
TICS fit
[TCAIBD]"Image 28.czi_TICS"!
TICS fit
[TCAIBD]"Image 32.czi_TICS"!
TICS fit
[TCAIBD]"Image 33.czi_TICS"!
TICS fit
[TCAIBD]"Image 47.czi_TICS"!
TICS fit
[TCAIBD]"Image 48.czi_TICS"!
TICS fit
[TCAIBD]"Image 49.czi_TICS"!
TICS fit
[TCAIBD]"Image 51.czi_TICS"!
TICS fit
[TCAIBD]"Image 52.czi_TICS"!
TICS fit
G
Lag time (s)

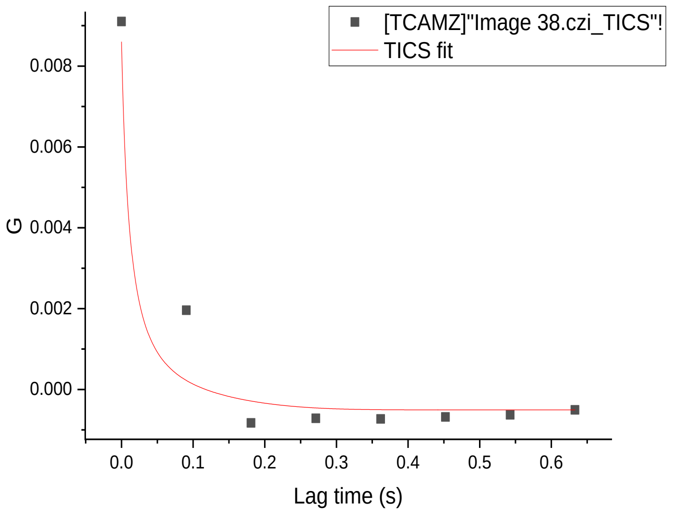
[TCAMZ]"Image 38.czi_TICS"!
TICS fit
G
0.008
0.006
0.004
0.002
0.000
0.0 0.1 0.2 0.3 0.4 0.5 0.6
Lag time (s)

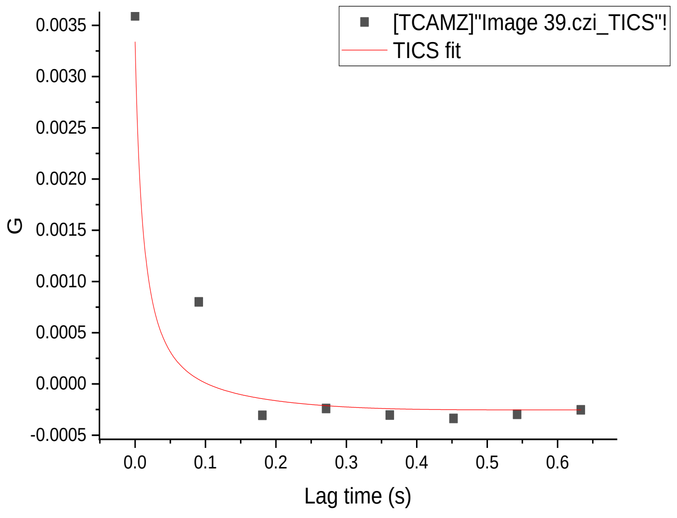
[TCAMZ]"Image 39.czi_TICS"!
TICS fit
G
0.0035
0.0030
0.0025
0.0020
0.0015
0.0010
0.0005
0.0000
-0.0005
0.0 0.1 0.2 0.3 0.4 0.5 0.6
Lag time (s)

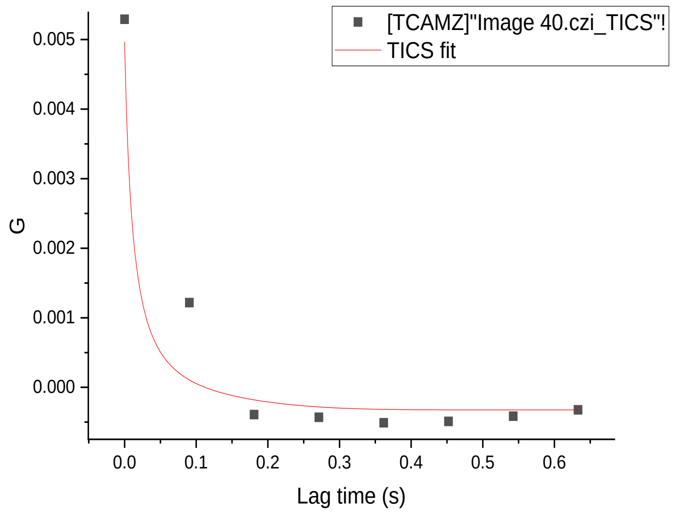
[TCAMZ]"Image 40.czi_TICS"!
TICS fit
G
0.005
0.004
0.003
0.002
0.001
0.000
0.0 0.1 0.2 0.3 0.4 0.5 0.6
Lag time (s)

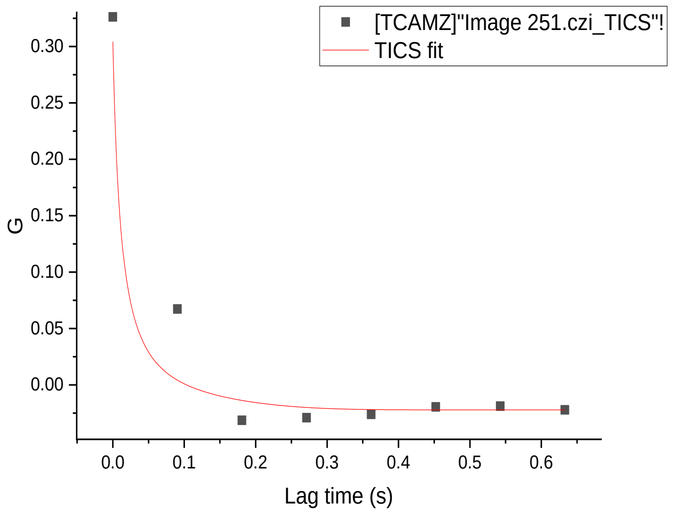
[TCAMZ]"Image 251.czi_TICS"!
TICS fit
G
0.30
0.25
0.20
0.15
0.10
0.05
0.00
0.0 0.1 0.2 0.3 0.4 0.5 0.6
Lag time (s)

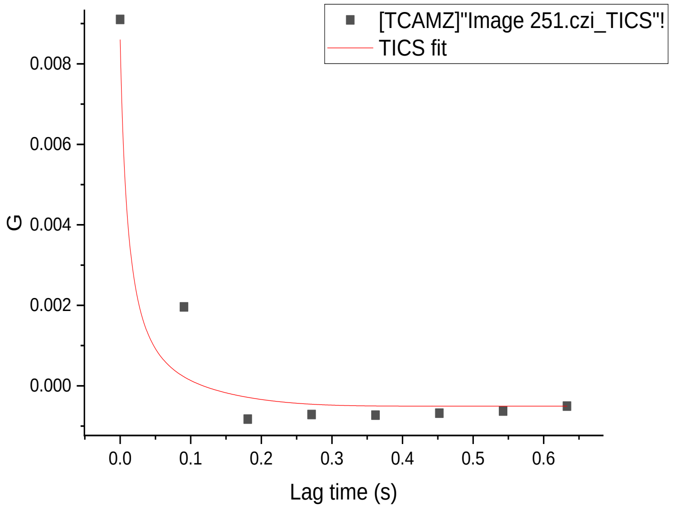
[TCAMZ]"Image 251.czi_TICS"!
TICS fit
G
0.008
0.006
0.004
0.002
0.000
0.0 0.1 0.2 0.3 0.4 0.5 0.6
Lag time (s)

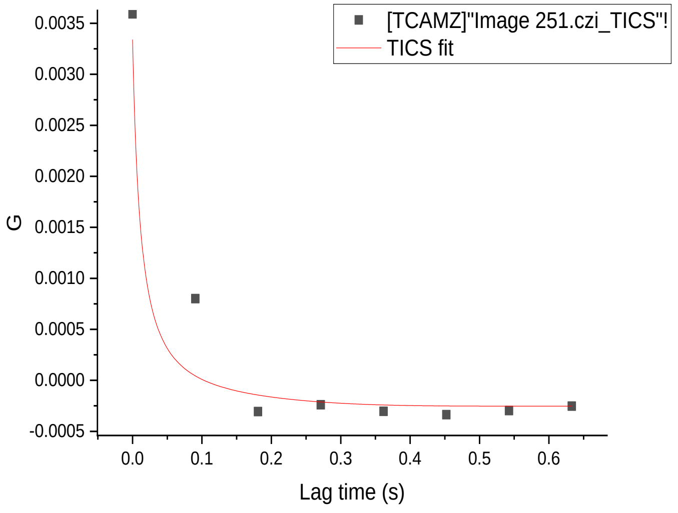
[TCAMZ]"Image 251.czi_TICS"!
TICS fit
G
0.0035
0.0030
0.0025
0.0020
0.0015
0.0010
0.0005
0.0000
-0.0005
0.0 0.1 0.2 0.3 0.4 0.5 0.6
Lag time (s)

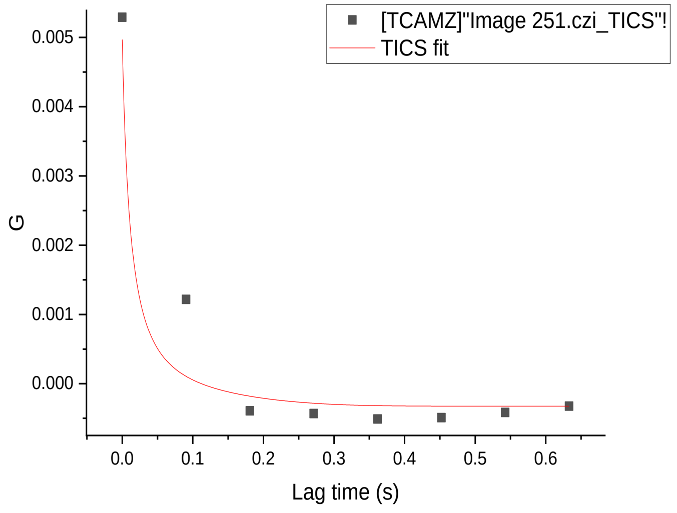
[TCAMZ]"Image 251.czi_TICS"!
TICS fit
G
0.005
0.004
0.003
0.002
0.001
0.000
0.0 0.1 0.2 0.3 0.4 0.5 0.6
Lag time (s)

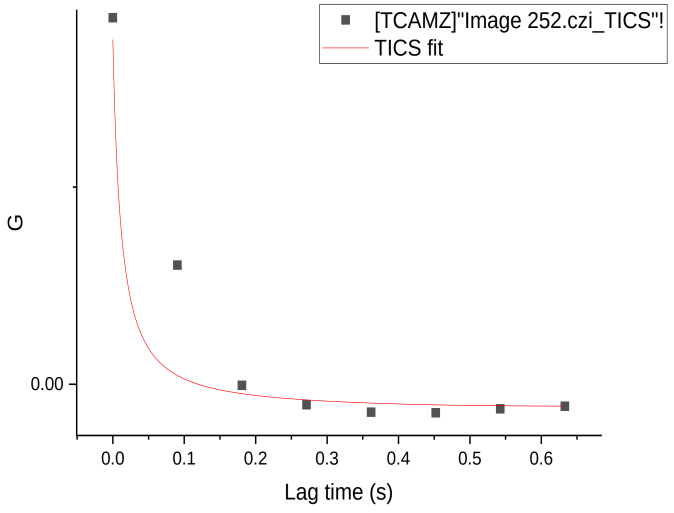
[TCAMZ]"Image 252.czi_TICS"!
TICS fit
G
0.00
0.0 0.1 0.2 0.3 0.4 0.5 0.6
Lag time (s)

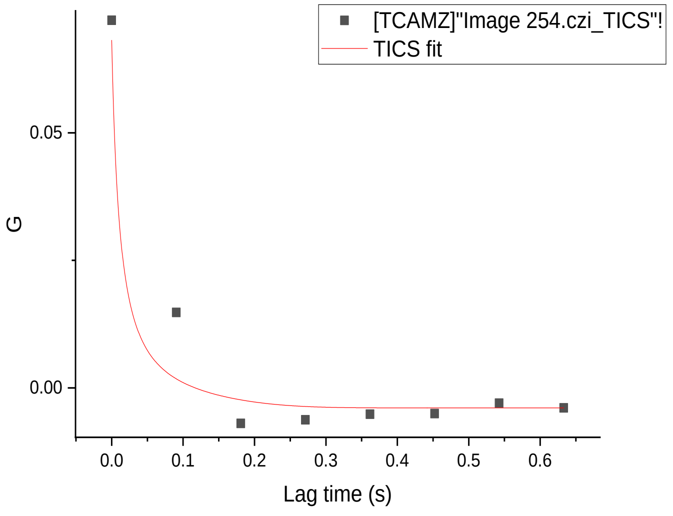
[TCAMZ]"Image 254.czi_TICS"!
TICS fit
G
0.05
0.00
0.0 0.1 0.2 0.3 0.4 0.5 0.6
Lag time (s)

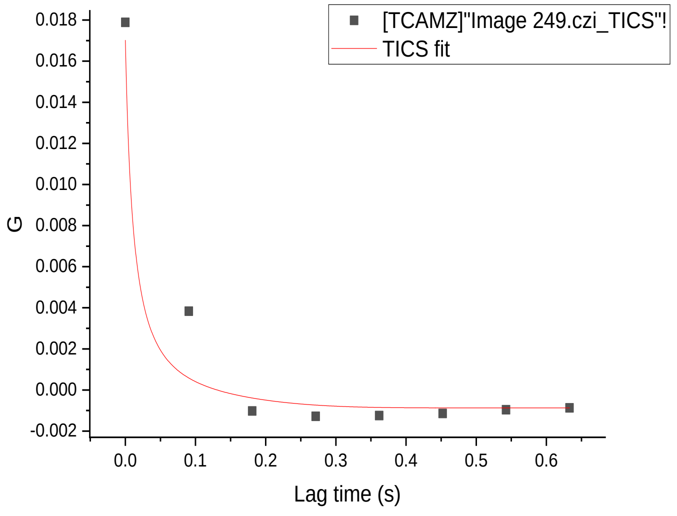
[TCAMZ]"Image 249.czi_TICS"!
TICS fit
G
0.018
0.016
0.014
0.012
0.010
0.008
0.006
0.004
0.002
0.000
-0.002
0.0 0.1 0.2 0.3 0.4 0.5 0.6
Lag time (s)

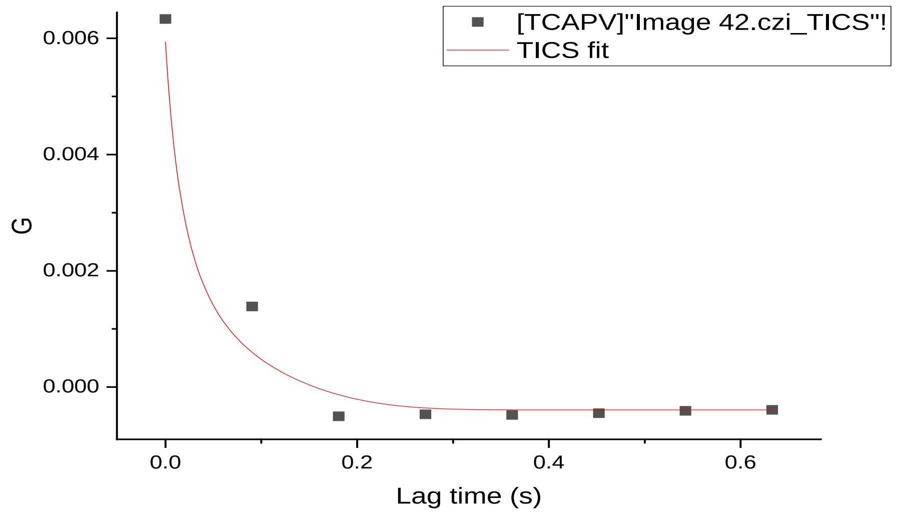
[TCAPV]"Image 42.czi_TICS"!
TICS fit
G
0.006
0.004
0.002
0.000
0.0
0.2
0.4
0.6
Lag time (s)

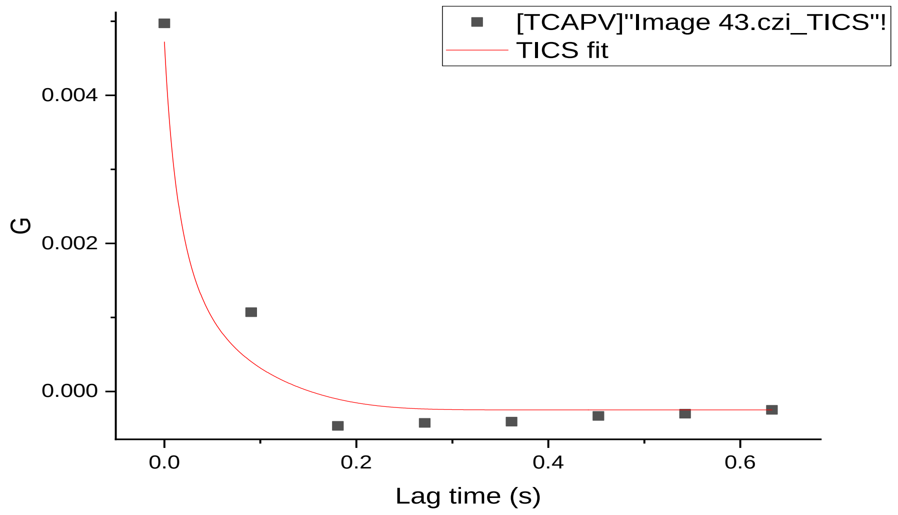
[TCAPV]"Image 43.czi_TICS"!
TICS fit
G
0.004
0.002
0.000
0.0
0.2
0.4
0.6
Lag time (s)

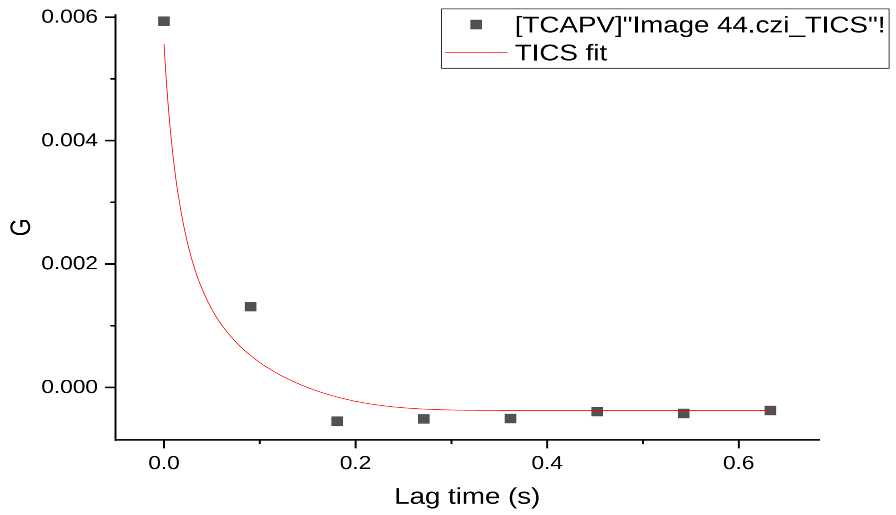
[TCAPV]"Image 44.czi_TICS"!
TICS fit
G
0.006
0.004
0.002
0.000
0.0
0.2
0.4
0.6
Lag time (s)

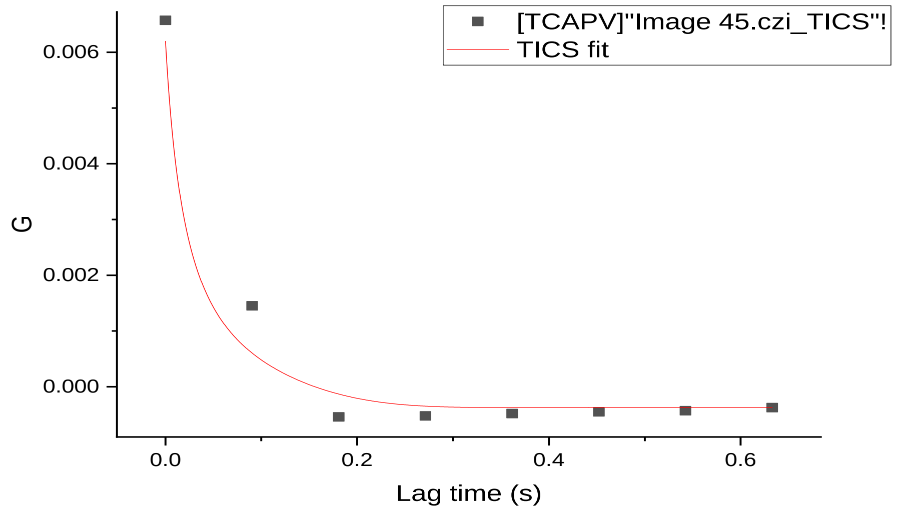
[TCAPV]"Image 45.czi_TICS"!
TICS fit
G
0.006
0.004
0.002
0.000
0.0
0.2
0.4
0.6
Lag time (s)

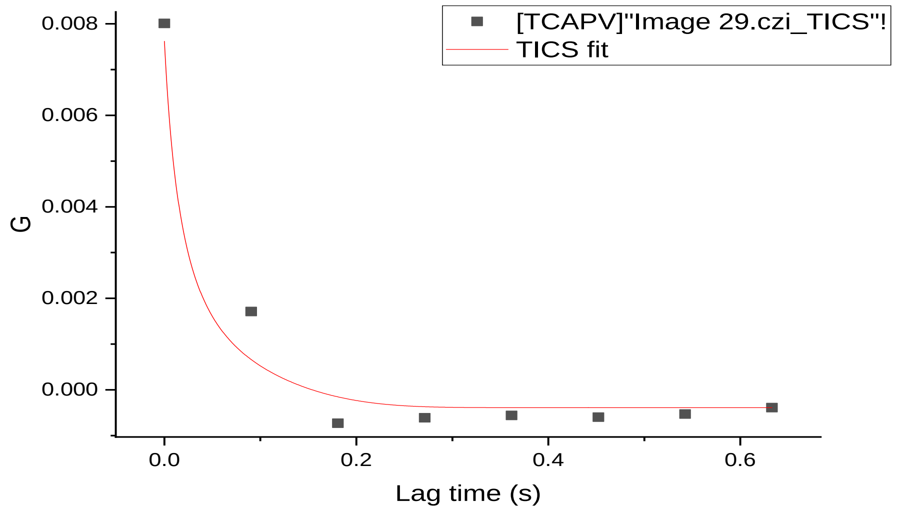
[TCAPV]"Image 29.czi_TICS"!
TICS fit
G
0.008
0.006
0.004
0.002
0.000
0.0
0.2
0.4
0.6
Lag time (s)

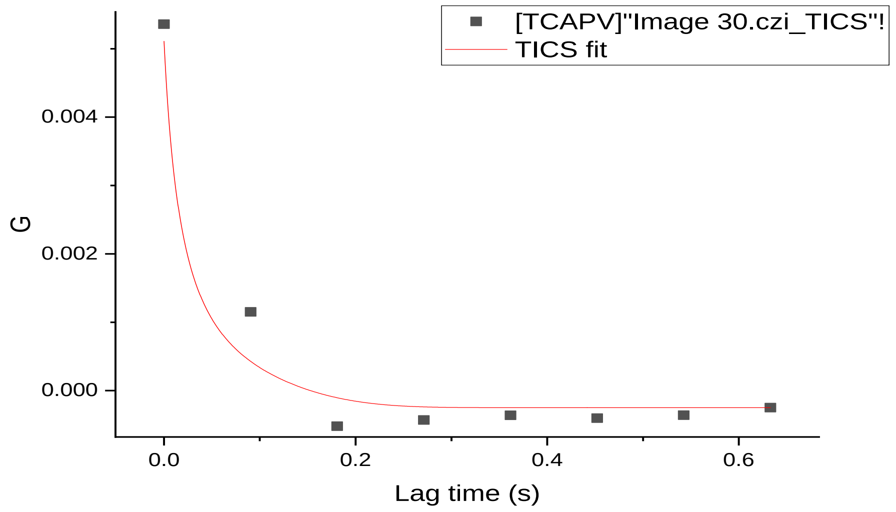
[TCAPV]"Image 30.czi_TICS"!
TICS fit
G
0.004
0.002
0.000
0.0
0.2
0.4
0.6
Lag time (s)

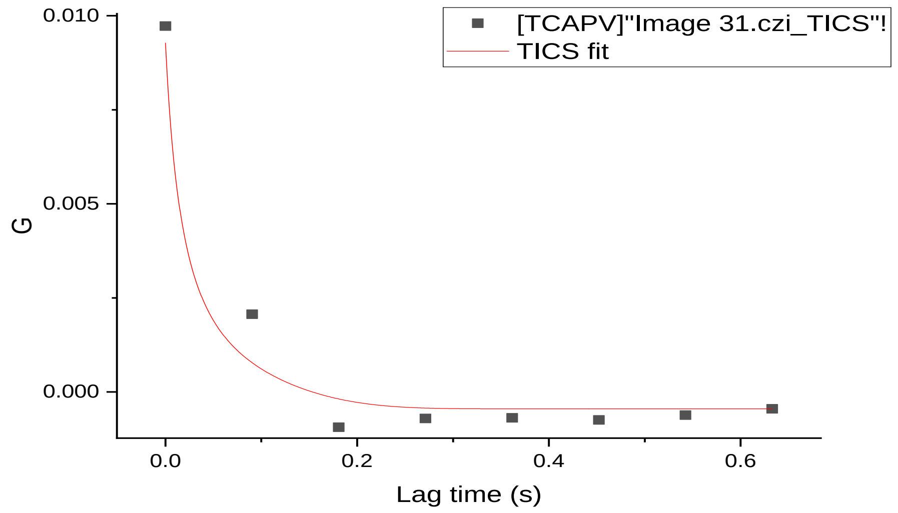
[TCAPV]"Image 31.czi_TICS"!
TICS fit
G
0.010
0.005
0.000
0.0
0.2
0.4
0.6
Lag time (s)
